# A Spatiotemporal Embryonic Atlas of Bombyx mori Integrating Single-Nucleus RNA Sequencing and Spatial Transcriptomics

**DOI:** 10.64898/2026.11.20.768035

**Authors:** Yabin Dou, Tengjiao Li, Yi Zhang, Zigang Shen, Changrui Liu, Maoshaung Ran, Sheng Li, Jie Chen, Keqin Chen, Qunxin Huang, Yanchao Shi, Shuaibin Wu, Letian Li, Yunlin Tang, Lei Chen, Shuo Li, Lunfu Shen, Yue Qin, Guo Chen, Chunxia Wang, Qing Lv, Xianzhi Meng, Junhong Wei, Mengxian Long, Jialing Bao, Chunfeng Li, Zeyang Zhou, Jie Chen, Xiaodong Li, Benhua Xiong, Tian Li, Guoqing Pan

**Affiliations:** State Key Laboratory of Resource Insects, Key Laboratory of Sericultural Biology and Genetic Breeding, Ministry of Agriculture and Rural Affairs, Southwest University, Chongqing 400715, China; Chongqing Key Laboratory of Microsporidia Infection and Control, Southwest University, Chongqing 400715, China; Chongqing Sericulture Science and Technology Research Institute, Chongqing Sericulture Technology Extension Station, Chongqing 400700, China; Guangxi Key Laboratory of Sericulture Ecology and Applied Intelligent Technology, Guangxi Collaborative Innovation Center of Modern Sericulture and Silk, Hechi University, Hechi 546300, China; Department of Microbiology, College of Basic Medical Sciences, Army Medical University, Chongqing 400700, China

## Abstract

The silkworm, *Bombyx mori*, is an important lepidopteran model and economically valuable insect. However, the cellular composition and dynamic spatiotemporal features of its embryonic development remain to be systematically characterized. Here, we integrated single-nucleus RNA sequencing (snRNA-seq) with spatial transcriptomics to construct an embryonic atlas of *B. mori* at 1, 3, 5, 7, and 9 days post-oviposition (dpo). We systematically identified the major cell types present during embryonic development and mapped their spatial distributions. We then focused on cellular heterogeneity and differentiation within the embryonic midgut lineage, identifying several candidate midgut-associated marker genes and characterizing their spatial expression patterns. These findings expand the molecular marker resources available for multiple cell types in *B. mori* embryos. This study provides valuable data and a framework for improving our understanding of cellular heterogeneity and organogenesis during embryonic development in *B. mori* and other lepidopteran insects.

## Introduction

Embryogenesis is a tightly orchestrated process through which a single fertilized egg undergoes proliferation and differentiation to form a complex multicellular organism with specialized tissues and organs^1–3^. Deciphering the transcriptional programs governing cell fate decisions and lineage trajectories is fundamental to developmental biology^4,5^. In recent years, single-cell RNA sequencing (scRNA-seq) and spatial transcriptomics has revolutionized this field, facilitating the construction of comprehensive developmental atlases for model organisms including *Drosophila* (fruit fly), zebrafish, and mice^6–8^. These resources offer unprecedented insights into the dynamic changes in gene expression and cellular heterogeneity during development.

However, our understanding of insect embryonic development remains heavily biased toward the dipteran model organism *Drosophila*. As the most diverse group within the animal kingdom, insects exhibit striking interordinal differences in developmental timing, cleavage modes, and organogenesis^9–11^. *B. mori* belongs to Lepidoptera-the second-largest insect order, encompassing critical agricultural pests and pollinators^12,13^. In contrast to the rapid embryogenesis of *Drosophila* (≈24 hours), *B. mori* undergoes an extended developmental period (≈9-10 days) with distinct morphological landmarks, rendering it an ideal model for investigating the prolonged dynamics of fate specification and overlapping organogenesis events^14–16^. Despite the significant economic importance of the *B. mori* and its widespread use as a model organism in insect physiology and developmental biology, existing Spatio-temporal Omics studies have primarily focused on individual tissues. Ma et al. used scRNA-seq to characterize silk gland cell heterogeneity and gene expression profiles in *B. mori* embryos and larvae, and combined these data with spatial transcriptomics to resolve silk gland architecture at high resolution. Feng et al. performed snRNA-seq on the fat body, hemocytes, midgut, and other tissues infected with *B. mori* nucleopolyhedrovirus (BmNPV), generating cell-type atlases and providing a resource for investigating gene expression responses to viral infection across cell types. Liu et al. constructed spatiotemporal atlases of wing discs at multiple developmental stages to elucidate the effects of hormones on wing developent.^17–22^. Consequently, a comprehensive high-resolution, time-resolved transcriptional atlas spanning the entire process of embryonic morphogenesis remains lacking, Which impede the systematic dissection of developmental dynamics and organogenesis mechanisms at the single-cell and spatial scales.

Current understanding of silkworm embryogenesis primarily relies on traditional histology and bulk transcriptome analyses, which mask cellular heterogeneity and asynchronous developmental events^23–25^. Critical questions regarding the sequential specification of cell lineages remain unresolved. For instance, comprehensively elucidating the development of the embryonic midgut, the maturation markers of diverse cell types during embryogenesis, and historical gene localization patterns would be challenging without a high-resolution atlas. Elucidating these cells and markers is not only pivotal for developmental biology but also provides foundational insights for modulating insect growth and immunity.

This study generated a comprehensive single-nucleus transcriptomic atlas of *B. mori* embryonic development, spanning the period from gastrulation to the pre-hatching stage. We analyzed single-nucleus gene-expression profiles from more than 80,000 cells collected at five key developmental stages—1, 3, 5, 7, and 9 dpo—and performed spatial transcriptomic sequencing on embryonic samples from the corresponding stages. We established an annotation framework for embryonic cell types and systematically identified the major cell populations present at each developmental time point. By integrating single-nucleus transcriptomic analyses with spatial validation of gene expression, we characterized in detail midgut cell types associated with digestion, nutrient absorption, and immune defense. This study provides an important data resource and conceptual foundation for investigating *B. mori* embryogenesis and the cell types involved in embryonic immunity.

## Result

### Panoramic Spatiotemporal Atlas of *B. mori* Embryo

The embryonic development of *B. mori* typically spans 9 days from oviposition to larval hatching. To systematically dissect the developmental trajectory of *B. mori* embryos, we performed snRNA-seq and Stereo-seq to construct comprehensive, multi-dimensional transcriptomic profiles at five critical embryonic time points: 1 dpo, 3 dpo, 5 dpo, 7 dpo, and 9 dpo (Fig 1A). Following rigorous quality control filtering, 81,772 high-quality single-nucleus transcriptomes were retained, exhibiting a median of 1,450 genes detected per nucleus. Through dimensionality reduction and uniform manifold approximation and projection (UMAP) analysis, embryonic cells from the five time points were computationally clustered into 42 transcriptionally distinct clusters (Fig 1B, 1D, Tab 1). Spatial transcriptomic sequencing data were analyzed using a spatial clustering strategy with a 20 × 20 pixel bin size (bin 20). Silkworm eggs corresponding to different developmental days were cropped from the capture area of the array. Dimensionality reduction and clustering of the dataset identified 26 clusters and their corresponding spots across samples, providing an initial characterization of spatial organization within the eggs (Fig 1C). The temporal dynamics of cell-type proportions showed that the number of cell clusters increased progressively throughout development (Fig 1E, Supplementary Data 1), reflecting the increasing cellular heterogeneity and complexity within the embryos.

**Fig. 1.**
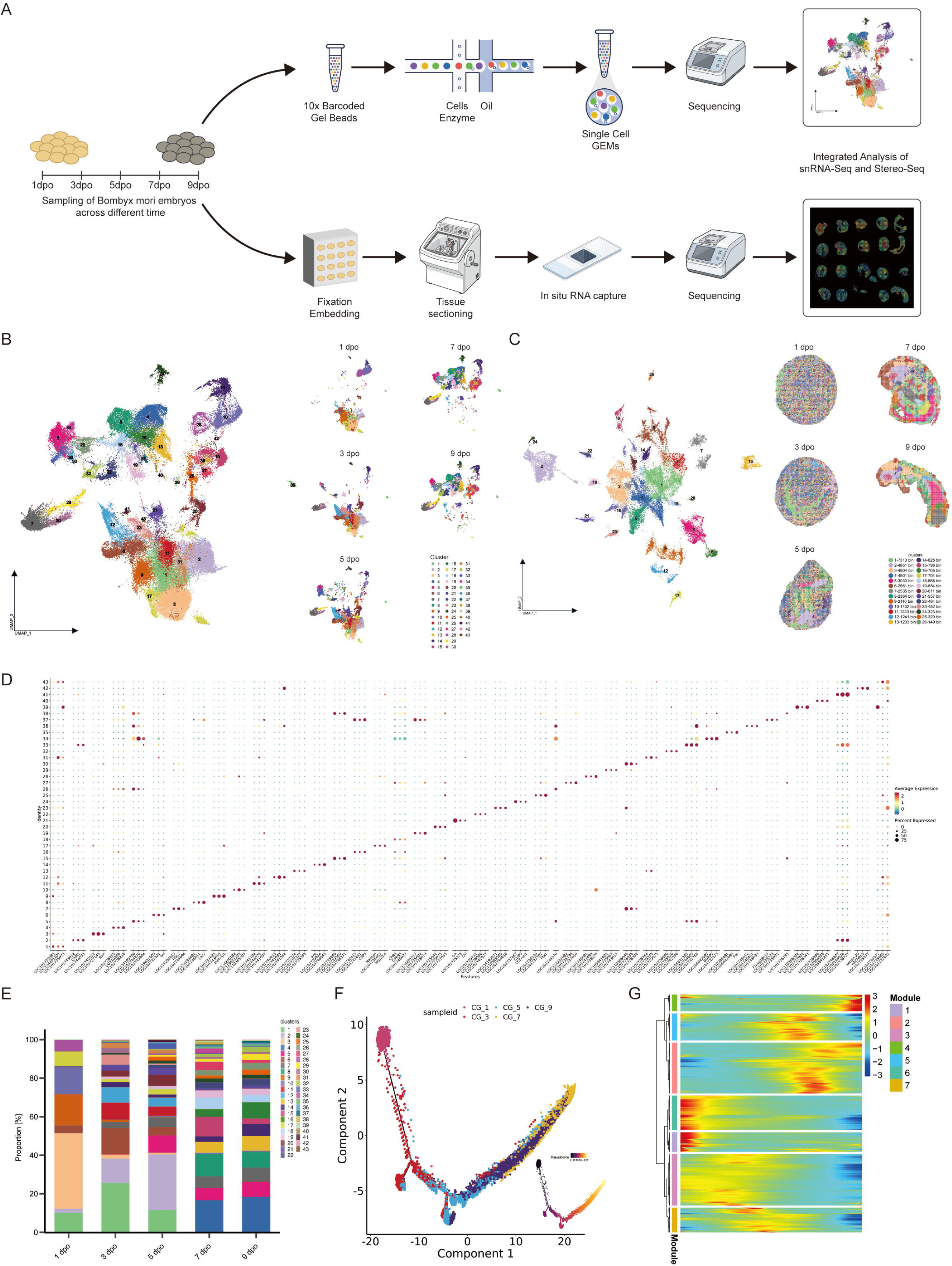
Global single-nucleus transcriptomic and spatial transcriptomic profiles of silkworm embryos. **A**, Schematic workflow of the study. **B**, Global UMAP plot of 81,831 cells from silkworm eggs collected at five developmental time points. The insets show the cellular distributions at 1, 3, 5, 7, and 9 days post-oviposition **C**, UMAP visualization of Stereo-seq data following dimensionality reduction and clustering. Individual spots correspond to bins, differentiated by color according to cluster identity. The adjacent plot depicts the spatial localization of these clusters within the tissue sections. **D**, Dot plot showing the three representative top-ranked genes in each cell cluster. **E**, Proportional representation of each cell cluster in the samples collected at each developmental stage. **F**, Pseudotemporal trajectories of samples from the five developmental time points. Each color represents one developmental time point, and each dot represents a cell. The inset on the right displays a gradient from dark to light, indicating the progression of differentiation from early to late stages. **G**, Heatmap showing gene expression patterns along the pseudotemporal trajectory of the global atlas. The horizontal axis represents developmental progression from early to late stages, from left to right. The color gradient from blue to red indicates increasing gene expression levels. The gene expression profiles were divided into seven modules.

Pseudotime analysis of the panoramic single-nucleus atlas revealed that cells at 1 dpo correspond to the embryonic establishment stage, forming relatively independent clusters along the developmental trajectory. At 3 and 5 dpo, silkworm embryos are in the stages of somite differentiation and organogenesis, while cells from these two days exhibit rapid shifts in cellular heterogeneity across the developmental trajectory. In contrast, cells at 7 and 9 dpo are largely fully developed, with their developmental states tending toward stability and relatively attenuated pseudotemporal trajectory dynamics (Fig 1F, 1G). Based on the pseudotime heatmap (Fig 1G) and functional enrichment analysis (GO and KEGG) of the corresponding modules, we summarized developmental features from 1 to 9 dpo (Supplementary Data 2-4). In the early stages of pseudotime, Module 1 and Module 6 exhibited high expression of RNA processing, ribosome biogenesis, and protein synthesis during the early stage, suggesting that the cells may be in a preparatory state conducive to proliferation. Module 3 then showed a transient increase during the early-to-intermediate phase, consistent with cell-cycle progression, DNA damage repair, and chromatin remodeling. In parallel, Module 7 gradually increased, indicating the activation of programs associated with cell-fate determination, tissue development, transcriptional regulation, and cell adhesion,possibly indicating that cells begin to further differentiate and develop. At the intermediate stage, Module 5 reached a distinct expression peak, suggesting the establishment of neuronal processes, synaptic structures, neurotransmitter signaling, and neuroendocrine signaling. Subsequently, Module 2 increased progressively during the intermediate-to-late phase, reflecting the gradual maturation of membrane trafficking, ion homeostasis, and metabolic adaptation. At the end of pseudotime, Module 4 increased sharply, with enrichment for mitochondrial protein import, respiratory-chain complex assembly, oxidative phosphorylation, and energy metabolism. This pattern suggests that cells ultimately transition toward enhanced mitochondrial function and metabolic maturation. Overall, these results suggest a continuous transition in cell state, from high biosynthetic activity and preparation for proliferation to genome remodeling and cell-fate determination, followed by the establishment of neural and tissue functions, membrane-system adaptation, and mitochondrial energy maturation.

We found that dimensionality reduction and clustering of all samples failed to adequately capture the underlying cellular heterogeneity of *B. mori* embryos across a range of resolutions. The temporal heterogeneity inherent in this complex dataset poses significant challenges for marker gene identification and manual cell type annotation following global dimensionality reduction. In light of this, we performed clustering on individual time-point samples using the global dataset framework and leveraged spatial transcriptomics mapping to refine cell annotations. This methodology significantly aided in elucidating the identity of embryonic cell types.

### Spatiotemporal Atlas and Characterization of *B. mori* Embryos across Developmental Time Points

Given that embryogenesis is a complex and highly orchestrated process, and the developmental timeline of *B. mori* embryos is much longer than that of *Drosophila*, we independently performed dimensionality reduction clustering and cell type annotation on single-nucleus datasets from each of the five distinct time points to achieve more precise annotation of *B. mori* embryonic cell types. Cell types at the five developmental time points were defined based on the following criteria: (1) marker genes from *B. mori*, *Drosophila*, and other Lepidoptera species, (2) the top 200 marker genes of each cell cluster marker (Supplementary Data 5), and (3) Gene Set Variation Analysis (GSVA), Gene Ontology (GO) and Kyoto Encyclopedia of Genes and Genomes (KEGG) enrichment analyses of these marker genes (Supplementary Data 6).

### Spatiotemporal Atlas of *B. mori* eggs at 1 dpo

A total of 15,586 cells were profiled at 1 dpo. Following dimensionality reduction and clustering analysis, 12 cell clusters were identified (Fig 2A). Cluster identities were assigned based on specific gene expression signatures (Fig 2E, Supplementary Data 5) and manual annotation, identifying 5 ectodermal clusters (including 2 neuroectodermal clusters), 2 mesodermal clusters, 2 endodermal clusters, metabolic clusters and unknown cluster (Tab 2).

**Fig. 2.**
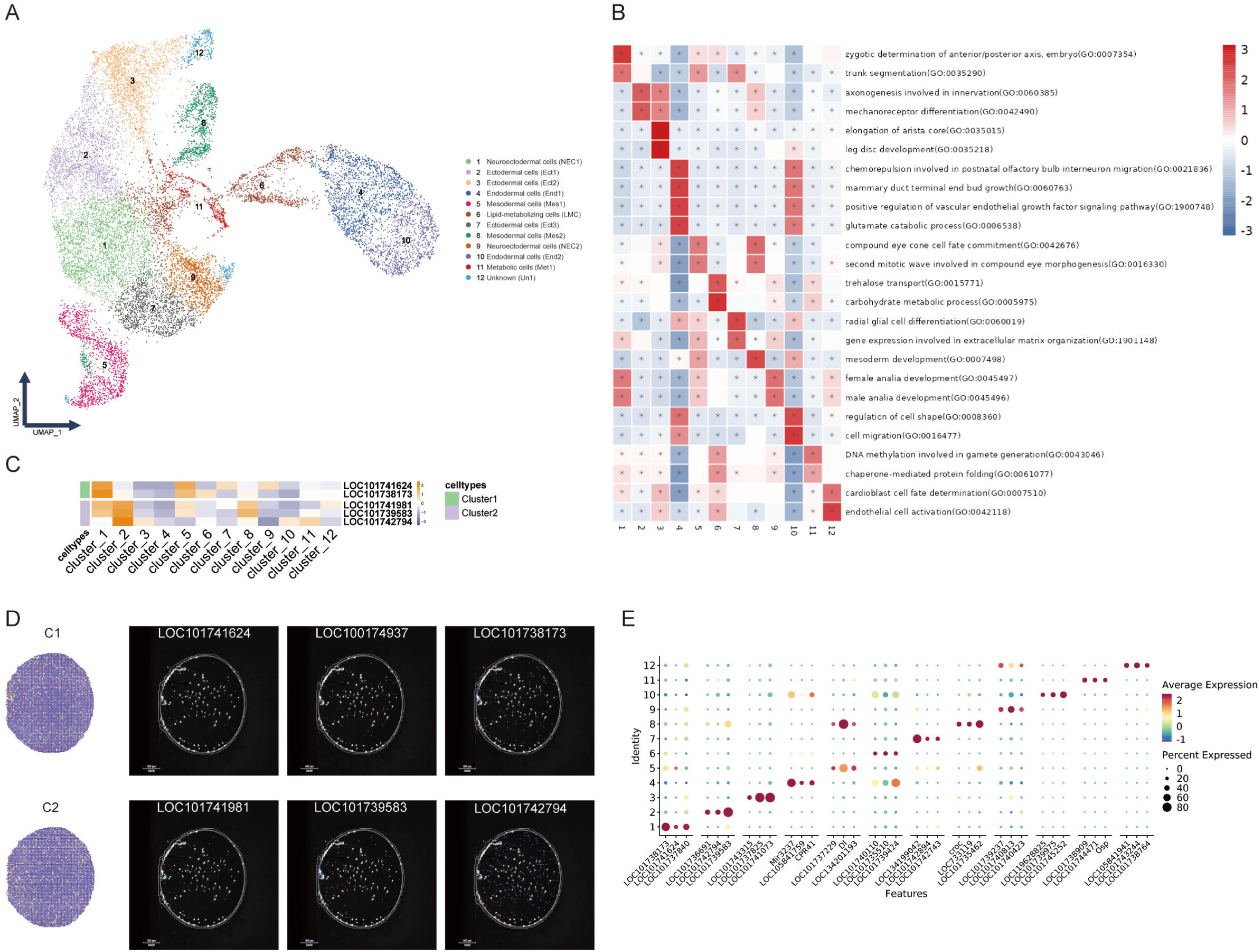
Cellular atlas of *Bombyx mori* embryos at 1 dpo. **A**, UMAP visualization of dimensionality reduction and cell-type annotations for the 1 dpo silkworm egg samples. **B**, GSVA heatmap of the cell clusters at 1 dpo. Based on the pathway activity scores calculated for individual cells using GSVA, differential testing was performed by comparing each cell group with all other cell groups. For each group, the two pathways with the highest *t*-statistics among those satisfying *p* < 0.05 were selected for heatmap visualization. **C**, Heatmap of selected marker genes at 1 dpo. These genes were used for the spatial localization shown in Fig D. **D**, Spatial transcriptomic mapping of selected cell clusters at 1 dpo and the spatial localization of representative genes on the spatial transcriptomic array. For each cell cluster, the first panel shows the RCTD mapping result. The color gradient from blue to red represents the inferred weight or relative abundance of each cell cluster in individual spatial spots, estimated by RCTD using the single-cell reference data. **E**, Dot plot showing representative marker genes on the UMAP plot at 1 dpo.

The marker genes of Cluster 1 predominantly exhibited expression patterns associated with neurogenesis and neuropeptide signal transduction. Among these, *neuroendocrine convertase 1* (LOC101741624) encodes neuroendocrine convertase 1, an enzyme involved in the tissue-specific proteolytic processing and maturation of prohormones and neuropeptide precursors in humans^26^. *CNGR-A26* (LOC10174937) encodes a neuropeptide receptor. Previous studies have demonstrated that neuropeptides and their G protein-coupled receptors are broadly involved in neuronal regulation, growth and development, and various physiological processes in insects^27,28^. In addition, *protein krueppel* (LOC101738173) is closely associated with the differentiation of neural precursor cells, tissue formation, and cell fate determination^29^. Based on the expression characteristics of these neurogenesis-and neuropeptide signaling-related genes, Cluster 1 was provisionally annotated as Neuroectodermal Cells (Ned1).

Cluster 2 was annotated as Ectodermal Cells (Ect1). This cluster exhibited high expression of *protein gooseberry* (LOC101741981) and *Zic family member odd paired* (LOC101739583). Studies in *Drosophila* have shown that *gooseberry* and *odd paired* are involved in the formation of neural and muscle tissues, respectively, during embryonic ectoderm development, and play important roles in germ-layer differentiation and tissue patterning^30^. In addition, Cluster 2 showed high expression of *Actn* (LOC101742794). Alpha-actinin is a key structural component of the sarcomeric Z-disc that crosslinks actin filaments between adjacent sarcomeres, thereby maintaining sarcomere integrity and facilitating the efficient transmission of contractile force^31^. Previous studies have demonstrated that this gene is expressed in both neural and muscle tissues of the *Drosophila* head. Collectively, the characteristic gene expression profile of Cluster 2 primarily reflects the differentiation of ectoderm-derived tissues, particularly neural-and muscle-related tissues. Therefore, Cluster 2 was provisionally annotated as Ect1.

GSVA revealed that (Fig 2B), at 1 dpo, processes associated with anterior–posterior axis specification and trunk segmentation reflected embryonic regionalization and the initial establishment of segmental structures. Enrichment of axonogenesis, glutamate metabolism, and sensory receptor differentiation further suggested that the nervous and sensory systems were beginning to form. In addition, cell migration, regulation of cell morphology, and extracellular matrix organization were prominent, indicating that embryonic cells were undergoing early tissue remodeling through migration, adhesion, and cytoskeletal reorganization. It is important to note that terms such as *mammary duct terminal end bud growth* and *postnatal olfactory bulb interneuron migration* should not be interpreted literally as referring to mammalian organs. Rather, in cross-species GO annotation, these terms generally represent conserved developmental, migratory, or signaling-regulatory genes that are annotated under those categories.

In the spatial transcriptomics data, mapping the single-nucleus cluster annotations onto the spatial transcriptomic sections showed that, on embryonic day 1, the corresponding signals were generally dispersed, with no clearly delineated regions of enrichment (Fig 2C, 2D). This pattern may be associated with the presence of free yolk granules in the early embryo. In light of the physiological characteristics of *B. mori* embryos at 1 dpo, 24 h post-oviposition corresponds to a developmental stage preceding germ-band formation, which may account for the relatively dispersed distribution of gene-localization signals.

### Spatiotemporal Atlas of *B. mori* eggs at 3 dpo

At 3 dpo, a total of 17,099 cells were collected. Dimensionality reduction and clustering analysis identified 22 cell clusters (Fig 3A), and based on the gene expression profiles of each cluster (Fig 3E, Supplementary Data 5), we identified multiple cell types, including neurons, hemocytes, intestinal cells, and somitic differentiation cells (Tab 3). Compared with 1 dpo, clusters exhibiting intestinal, appendage, and epithelial characteristics were evident by 3 dpo, suggesting that embryonic organogenesis and tissue differentiation had become more pronounced at this stage.

**Fig. 3.**
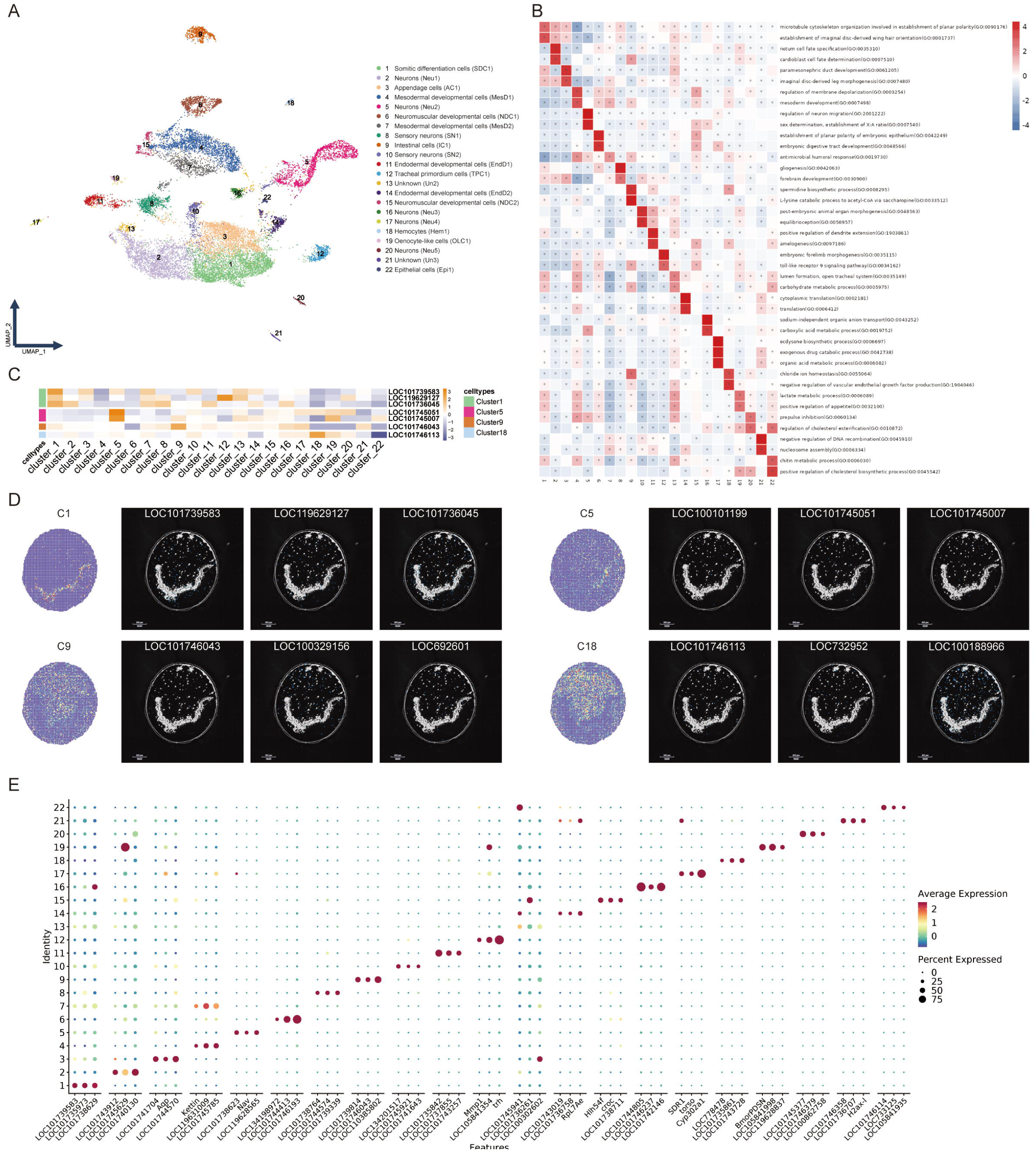
Cellular atlas of *Bombyx mori* embryos at 3 dpo. **A**, UMAP visualization of dimensionality reduction and cell-type annotations for the 3 dpo silkworm egg samples. **B**, GSVA heatmap of the cell clusters at 3 dpo. Based on the pathway activity scores calculated for individual cells using GSVA, differential testing was performed by comparing each cell group with all other cell groups. For each group, the two pathways with the highest *t*-statistics among those satisfying *p* < 0.05 were selected for heatmap visualization. **C**, Heatmap of selected marker genes at 3 dpo. These genes were used for the spatial localization shown in Fig D. **D**, Spatial transcriptomic mapping of selected cell clusters at 3 dpo and the spatial localization of representative genes on the spatial transcriptomic array. For each cell cluster, the first panel shows the RCTD mapping result. The color gradient from blue to red represents the inferred weight or relative abundance of each cell cluster in individual spatial spots, estimated by RCTD using the single-cell reference data. **E**, Dot plot showing representative marker genes on the UMAP plot at 3 dpo.

Cluster 1 exhibited transcriptional features associated with appendage development and segmental patterning. *opa* (LOC101739583) encodes a member of the Zic family of transcription factors. In *Drosophila*, *opa* is classified as a pair-rule gene, is highly expressed during early embryonic development, and contributes to the establishment of embryonic segmentation patterns^32^. *Knirps-related protein* (LOC119629127) encodes a short-range transcriptional repressor associated with the nuclear receptor superfamily. Studies in *Drosophila* have implicated this gene in developmental processes including abdominal cell migration, tracheal branch morphogenesis, and posterior segment formation^33,34^. In addition, Cluster 1 showed high expression of *four-jointed box kinase* (LOC101736045), which plays important roles in leg and wing growth and differentiation, as well as normal eye development in *Drosophila*^35,36^. Based on the expression of genes associated with segmentation, appendage growth, and morphogenesis, Cluster 1 was provisionally annotated as Somitic Differentiation Cells (SDC1).

Cluster 5 was annotated as Neurons (Neu1). This cluster exhibited high expression of *para* (LOC100101199), which encodes a voltage-gated sodium channel that plays important roles in maintaining neuronal excitability and regulating nervous system development in *Drosophila*^37^. In addition, *RhoGAP100F* (LOC101745051) is involved in axonal growth, neuronal morphogenesis, and neural network formation^38,39^, whereas *Septin4* (LOC101745007) is predominantly expressed in the wings, head, central nervous system, and neurons of *Drosophila*^40^. Collectively, these genes indicate that Cluster 5 possesses pronounced molecular features associated with neuronal differentiation and neural function.

Cluster 9 exhibited prominent features associated with nutrient transport, digestion, and metabolism and was therefore annotated as Intestinal Cells (IC1). This cluster expressed *putative inorganic phosphate cotransporter* (LOC101746043), which encodes a member of the inorganic phosphate cotransporter family. Previous studies have shown that related transporters participate in phosphate absorption in the mammalian small intestine^41,42^. *St4* (LOC100329156) encodes a sugar transporter that is highly expressed or enriched in the intestinal tissues of aphids and may be involved in the transport of carbohydrates such as glucose and fructose^43^. Overall, these genes are primarily involved in nutrient absorption, transmembrane transport, digestion, and metabolism, which are consistent with the major physiological functions of the midgut and support the annotation of Cluster 9 as IC1.

Cluster 18 exhibited characteristic features associated with hemocytes and innate immunity. This cluster expressed *integrin alpha-PS3-like* (LOC101746113), which encodes an integrin α-subunit-related transmembrane receptor capable of mediating cell adhesion and signal transduction. Studies in *Drosophila* have shown that integrin-related molecules participate in hemocyte adhesion to and phagocytosis of pathogens, embryonic mesodermal cell migration, and epithelial tissue formation. In the *B. mori*, this gene has also been reported as a hemocyte-associated marker^44,45^. In addition, *CLIP3* (LOC732952) encodes a Clip domain-containing serine protease that participates in the regulation of innate immune signaling pathways in insects and may mediate pathogen recognition and activation of immune responses through proteolytic cascades^46^. Based on the co-expression of genes associated with cell adhesion, phagocytosis, and innate immunity, Cluster 18 was provisionally annotated as Hemocytes (Hem1).

By 3 dpo, embryonic development was characterized primarily by the establishment of cell polarity and the formation of organ primordia (Fig 3B). At this stage, developmental processes had shifted from body-axis specification and segmentation toward the oriented arrangement of cells, cell-fate specification, and morphogenesis of organ primordia. Microtubule cytoskeleton-and planar cell polarity-associated processes may contribute to maintaining the spatial organization of epithelial cells and tissue boundaries. Enrichment of terms related to neuronal migration, gliogenesis, and forebrain development suggests further differentiation of the nervous system. Meanwhile, terms associated with antibacterial humoral responses and Toll-like receptors began to emerge, suggesting that gene modules involved in embryonic innate defense may be established in parallel with organogenesis.

By day 3 of embryonic development, the germ band was clearly established. Spatial mapping of the single-nucleus transcriptomic data showed that C1 SDC1 were primarily localized to arc-shaped regions associated with the germ band (Fig 3C, 3D). The spatial signals of representative candidate genes partially overlapped with these regions, supporting an association between C1 and germ-band and segment formation. C5 Neu2 were mainly distributed in localized regions adjacent to the germ band. Their candidate genes showed relatively sparse, localized signals, suggesting that these cells may occupy regions associated with early neural differentiation, although no distinct neural tissue-like spatial organization was yet apparent. C9 IC1 were primarily projected to the interior of the embryo, where signals from their candidate genes were detectable but broadly dispersed. C18 Hem1 showed a comparatively widespread and scattered distribution, consistent with the dispersed occurrence of hemocytes.

### Spatiotemporal Atlas of *B. mori* eggs at 5 dpo

5 dpo in the *B. mori* corresponds to the period of embryonic inversion and represents a critical stage of organogenesis and tissue differentiation. Dimensionality reduction and clustering analysis of the 16,824 collected cells identified 22 cell clusters (Fig 4A). Based on the characteristic gene expression patterns of each cluster and functional studies in related species, several clusters were provisionally annotated (Fig 4E, Supplementary Data 5). At 5 dpo, embryos contain multiple cell types, including muscle cells, epithelial cells, hemocytes, intestinal cells, and sensory neurons (Tab 4).

**Fig. 4.**
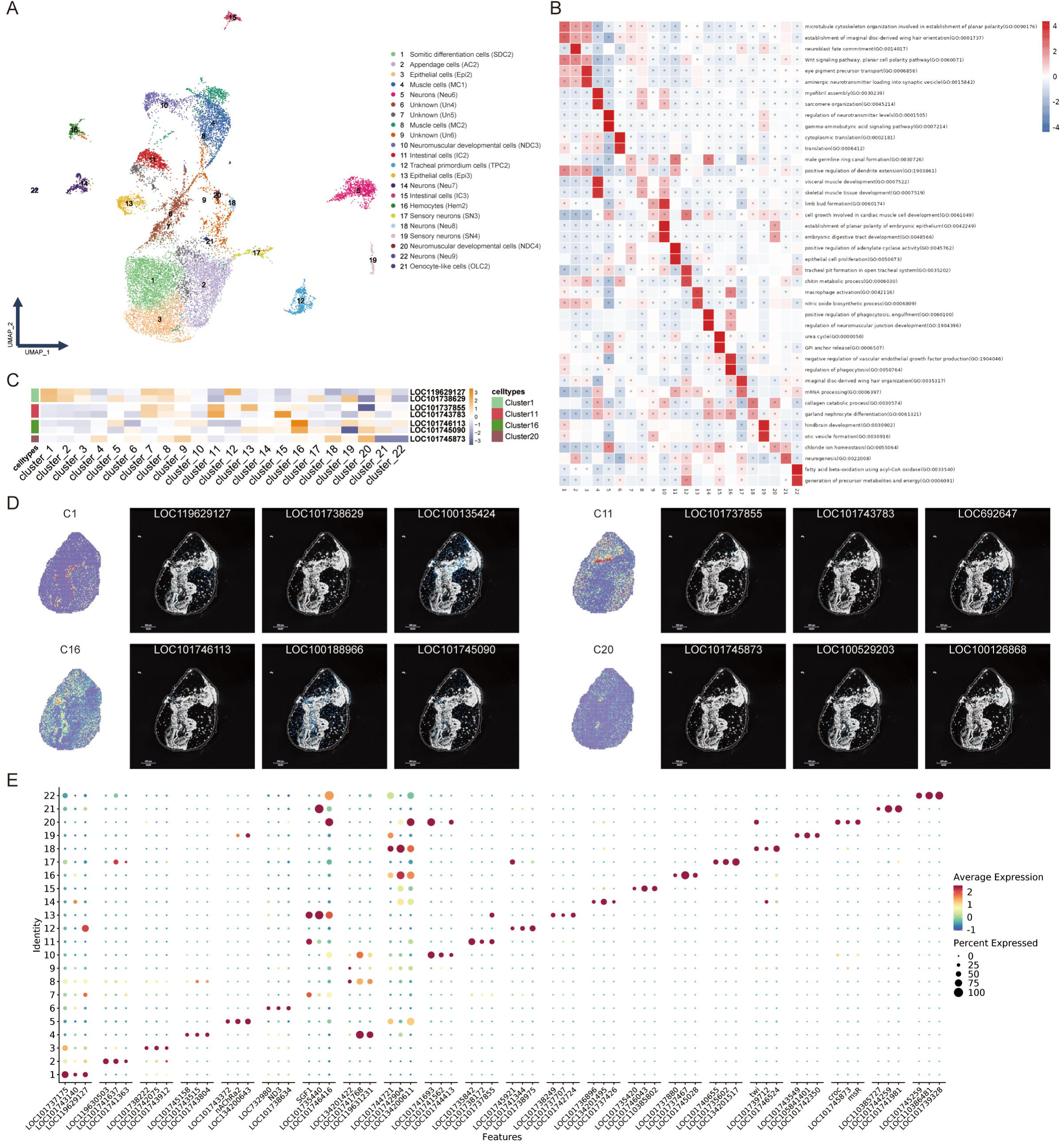
Cellular atlas of *Bombyx mori* embryos at 5 dpo. **A**, UMAP visualization of dimensionality reduction and cell-type annotations for the 5 dpo silkworm egg samples. **B**, GSVA heatmap of the cell clusters at 5 dpo. Based on the pathway activity scores calculated for individual cells using GSVA, differential testing was performed by comparing each cell group with all other cell groups. For each group, the two pathways with the highest *t*-statistics among those satisfying *p* < 0.05 were selected for heatmap visualization. **C**, Heatmap of selected marker genes at 5 dpo. These genes were used for the spatial localization shown in Fig D. **D**, Spatial transcriptomic mapping of selected cell clusters at 5 dpo and the spatial localization of representative genes on the spatial transcriptomic array. For each cell cluster, the first panel shows the RCTD mapping result. The color gradient from blue to red represents the inferred weight or relative abundance of each cell cluster in individual spatial spots, estimated by RCTD using the single-cell reference data. **E**, Dot plot showing representative marker genes on the UMAP plot at 5 dpo.

Cluster 1 exhibited prominent features associated with segmentation and posterior regional differentiation. *mid* (LOC101738629) belongs to the T-box family of transcription factors and is primarily involved in the regulation of cell migration. It also plays important roles in segment boundary formation and abdominal appendage development in *Drosophila*^47^. In addition, *abdominal A* (LOC100135424) has been reported in *Drosophila* as an important regulatory factor determining the identity of posterior embryonic segments and is involved in the differentiation of posterior appendages, including the legs and abdomen^48^. Based on the reported functions of these genes in segmental patterning, regional specification, and appendage development, Cluster 1 was provisionally annotated as Somitic Differentiation Cells (SDC2).

Cluster 4 exhibited prominent expression of genes associated with myogenesis. *Myogenic-determination protein nautilus* (LOC101743515) is an important myogenic determination factor in *Drosophila* and plays a central regulatory role in insect muscle development^49^. This cluster also expressed *glutamate receptor ionotropic, kainate 3* (LOC101743804). In mammals, this gene encodes a ligand-gated ion channel involved in rapid excitatory synaptic transmission, suggesting that the corresponding cells may possess features related to excitatory signal responsiveness^50^. Given the prominent expression of *nautilus* and its established role in myogenesis, Cluster 4 exhibited strong myogenic differentiation characteristics and was therefore provisionally annotated as Muscle Cells (MC1).

Clusters 11 and 15 both exhibited expression profiles associated with intestinal tissues and were therefore annotated as Intestinal Cells (IC2 and IC3). Cluster 11 expressed *transcription factor GATA-6* (LOC101737855), *LKRSDH* (LOC101743783), and *GlcNAcase3* (LOC692647). GATA family transcription factors are closely associated with endodermal differentiation and digestive organ development^51^. *LKRSDH* is involved in amino acid metabolism, whereas *GlcNAcase3* participates in the hydrolysis of carbohydrate-related substrates^52–54^. The expression of these genes suggests that Cluster 11 may possess active digestive and metabolic functions. Cluster 15 showed high expression of *putative inorganic phosphate cotransporter* (LOC101746043) and *facilitated trehalose transporter Tret1-like* (LOC101743252), which are associated with the transmembrane transport of nutrients such as inorganic phosphate and trehalose, respectively^55,56^. These features are consistent with the roles of intestinal epithelial cells in nutrient absorption and material transport. Cluster 16 was annotated as Hemocytes (Hem2). This cluster expressed *integrin alpha-PS3-like* (LOC101746113), which encodes an integrin-related receptor involved in cell adhesion, migration, and immune signal transduction and plays important roles in hemocyte function and innate immunity in insects^57^. The cluster also expressed *Scr-c* (LOC100188966). This gene belongs to the C-type lectin-like receptor family within the scavenger receptor superfamily and can recognize and internalize various ligands^58^. It is involved in pathogen recognition, phagocytosis, innate immune defense, and the maintenance of cellular metabolic homeostasis. The expression pattern of these genes supports the interpretation that Cluster 16 possesses molecular features characteristic of hemocytes and immune effector cells.

Cluster 20 co-expressed multiple genes associated with neural and muscle development and was therefore annotated as Neuromuscular Developmental Cells (NDC4). *Sodium/potassium-transporting ATPase subunit beta-1* (LOC101745873) is predominantly expressed in the embryonic trachea, central nervous system, and neurons of the peripheral nervous system in *Drosophila*^59^. *MsR* (LOC692474) is highly expressed in body-wall and visceral muscles and may be involved in the regulation of muscle contraction mediated by neuropeptides^60^. In addition, *twi* (LOC100529203) has been shown to participate in mesoderm formation, muscle differentiation, and tissue remodeling in the *B. mori*, and plays an important regulatory role in the differentiation of body-wall muscles and silk-gland attachment muscles^61^. *Run* (LOC100126868), a member of the Runx family of transcription factors in insects, is involved in embryonic segmentation and the specification of muscle and neural cell fates^62^. Based on the co-expression of genes associated with neuronal development, muscle differentiation, and embryonic tissue patterning, Cluster 20 was provisionally identified as NDC4.

At 5 dpo, embryonic development had begun to shift from morphogenesis toward tissue functionalization (Fig 4B). This transition was characterized by changes in pathways associated with Wnt/planar cell polarity signaling, neuroblast fate determination, neurotransmitter regulation, myofibril assembly, sarcomere organization, and visceral and skeletal muscle development, suggesting a shift from the formation of organ primordia toward the structural and functional differentiation of tissues. Enrichment of terms related to GABA signaling, amine neurotransmitter loading, and regulation of neurotransmitter levels suggests that the nervous system was gradually acquiring the machinery required for signal transmission. Meanwhile, myofibril-and sarcomere-associated pathways indicate the onset of contractile tissue assembly. In addition, the emergence of terms related to chitin metabolism, epithelial cell proliferation, phagocytosis, nitric oxide synthesis, and immune cell-like functions suggests concurrent remodeling of the epidermal structure and innate defense modules.

At 5 dpo, spatial transcriptomic analysis revealed relatively well-defined embryonic morphology (Fig 4C, 4D). C1 SDC2 exhibited a segmental distribution along both sides of the embryo, while C11 IC2 had clustered in the ventral region. C20 NDC4 associated with neuromuscular development were primarily localized to discrete regions within the embryo, whereas the C16 Hem2 showed a more dispersed distribution. Candidate genes associated with each cell cluster displayed detectable spatial signals in the sections, with varying degrees of localization. Overall, these findings provide preliminary support for the correspondence between the identified cell clusters and specific embryonic regions.

### Spatiotemporal Atlas of *B. mori* eggs at 7 dpo

At 7 dpo, a total of 16,221 cells were captured. Dimensionality reduction and clustering analysis identified 24 cell clusters (Fig 5A). Based on the characteristic gene expression patterns of each cluster and relevant studies (Fig 5E, Supplementary Data 5), several cell populations were provisionally annotated, including hemocytes, neurons, intestinal cells, and muscle cells, and fat body-and silk gland-related cell clusters were also identified (Tab 5). Cluster 2 was annotated as Intestinal Cells (IC4). This cluster specifically expressed *Apn4* (LOC100127099) and *peptidoglycan recognition protein* (LOC101739748). Aminopeptidase N is an exopeptidase found primarily in the larval midgut of lepidopteran insects, where it participates in the digestion of proteins and peptides and is closely associated with the activity of *Bacillus thuringiensis* toxins^63^. Peptidoglycan recognition proteins (PGRPs) are important pattern-recognition receptors in insects. They recognize peptidoglycan in microbial cell walls and mediate innate immune responses^64^. PGRP-L1, in particular, is considered an important regulator of gut immunity and microbiota homeostasis in the *B. mori* and may regulate antimicrobial peptide production and the clearance of specific microorganisms through the IMD signaling pathway^65^. The expression of genes associated with digestion and intestinal immunity supports the annotation of Cluster 2 as IC4.

**Fig. 5.**
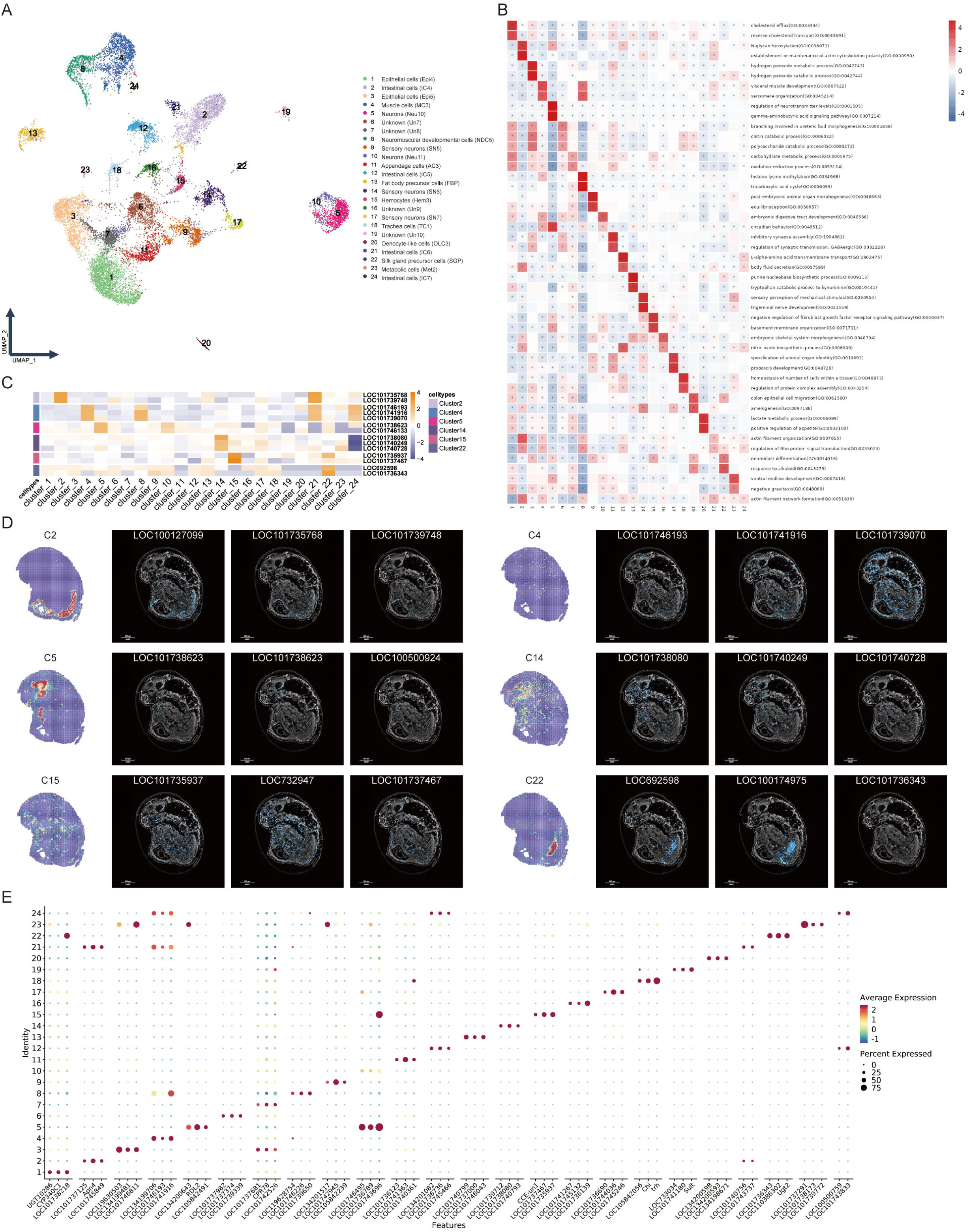
Cellular atlas of *Bombyx mori* embryos at 7 dpo. **A**, UMAP visualization of dimensionality reduction and cell-type annotations for the 7 dpo silkworm egg samples. **B**, GSVA heatmap of the cell clusters at 7 dpo. Based on the pathway activity scores calculated for individual cells using GSVA, differential testing was performed by comparing each cell group with all other cell groups. For each group, the two pathways with the highest *t*-statistics among those satisfying *p* < 0.05 were selected for heatmap visualization. **C**, Heatmap of selected marker genes at 7 dpo. These genes were used for the spatial localization shown in Fig D. **D**, Spatial transcriptomic mapping of selected cell clusters at 7 dpo and the spatial localization of representative genes on the spatial transcriptomic array. For each cell cluster, the first panel shows the RCTD mapping result. The color gradient from blue to red represents the inferred weight or relative abundance of each cell cluster in individual spatial spots, estimated by RCTD using the single-cell reference data. **E**, Dot plot showing representative marker genes on the UMAP plot at 7 dpo.

Cluster 4 exhibited prominent features associated with myogenesis and muscle function. *TBX1* (LOC101746193) encodes a member of the T-box family of transcription factors. Studies indicate that *Tbx1* is the ortholog of *organ-1* in *Drosophila* and is closely associated with muscle formation^66^. *PKCδ*(LOC101741916), whose phosphorylation is important for sarcomere assembly and normal muscle function^67^. *Ryanodine receptor* (LOC101739070) encodes a calcium-release channel required for normal muscle function in *Drosophila* and may participate in excitation-contraction coupling^68^. Taken together, the expression of genes associated with muscle formation, sarcomere assembly, and contraction supports the annotation of Cluster 4 as Muscle Cells (MC3).

Cluster 5 was annotated as Neurons (Neu10). This cluster expressed *transcriptional repressor scratch 2* (LOC101738623) and *protein CEPU-1* (LOC101746133). Both genes are associated with nervous system development and the regulation of neural functions, suggesting that this cell population possesses molecular features related to neuronal differentiation and neural regulation^69,70^. Cluster 15 was annotated as Hemocytes (Hem3). *Hemolectin* (LOC101735937) is specifically expressed in embryonic and larval hemocytes of *Drosophila* and may participate in hemostasis and coagulation^71,72^. *Scavenger receptor type C* (LOC100188966) encodes a C-type scavenger receptor. Previous studies have shown that this gene is expressed in both hemocytes and the Malpighian tubules of fifth-instar silkworms, with relatively high expression levels maintained in hemocytes^73^. These genes are involved in coagulation, pathogen recognition, and immune defense, and their expression profiles indicate that Cluster 15 possesses typical molecular characteristics of hemocytes. Cluster 22 exhibited distinct features of silk gland cells. This cluster specifically expressed *seroin 1* (LOC692598) and *Ugt2* (LOC100174975). *Seroin 1* is primarily expressed in the *B. mori* silk gland and exhibits antimicrobial activity^74^. *Ugt2* encodes a UDP-glycosyltransferase, whose glycosylation activity may contribute to the detoxification and excretion of endogenous metabolites and harmful compounds, the regulation of hormone levels, and growth and development^75^. Previous studies have shown that several members of the UDP-glycosyltransferase family display tissue-specific expression in the midgut and silk glands of fifth-instar silkworms^76^. Given the silk-gland-specific expression pattern of *seroin 1* and the potential functions of *Ugt2*, Cluster 22 was provisionally annotated as Silk Gland Precursor Cells (SGP).

At 7 dpo, lipid and carbohydrate metabolism had increased, accompanied by further maturation of tissue architecture and neural function (Fig 5B). Enrichment of terms related to sarcomere organization and visceral muscle development indicated the continued formation of contractile tissues. Meanwhile, terms associated with GABA signaling, inhibitory synapse assembly, neurotransmitter regulation, and balance/mechanosensory functions suggested that the nervous system had progressed beyond the early stages of neuronal production and migration toward the establishment of neural connections and the acquisition of sensory function.

The enrichment of chitin, polysaccharide, and carbohydrate catabolism, together with the tricarboxylic acid cycle and redox processes, indicated the simultaneous demands of structural-material turnover and energy metabolism. In addition, cholesterol transport, N-glycan modification, and changes in cytoskeletal polarity may have been associated with the remodeling of membrane structures, cell surfaces, and tissue morphology.

Spatial mapping of cell clusters in 7dpo *B. mori* embryos revealed differences in the distributions of several cell populations (Fig 5C, 5D). C2 IC4 were primarily localized to a discrete band-like region in the lower part of the embryo. C5 Neu10 and C14 SN6 were both prominent in the anterior region, showing partial spatial overlap. C15 exhibited a relatively dispersed distribution, broadly consistent with the spatial pattern of Hem3, whereas C22 SGP showed localized enrichment. Together, these findings suggest that cell populations in day 7 embryos exhibit regionally distinct spatial distributions.

### Spatiotemporal Atlas of *B. mori* eggs at 9 dpo

At 9 dpo, the embryo enters the preparatory stage preceding hatching of the first-instar larva, during which the major organs and tissues gradually mature. Single-nucleus transcriptome sequencing captured a total of 16,042 cells. Dimensionality reduction and clustering analysis identified 32 cell clusters (Fig 6). Based on the characteristic gene expression patterns of each cluster and existing functional studies, several cell populations were provisionally annotated (Fig 5E, Supplementary Data 5). Before the silkworm eggs hatch, the number and types of cells are already quite diverse, including various neurons, epithelial cells, intestinal cells, hemocytes, silk gland cells, fat body cells, and others (Tab 6).

**Fig. 6.**
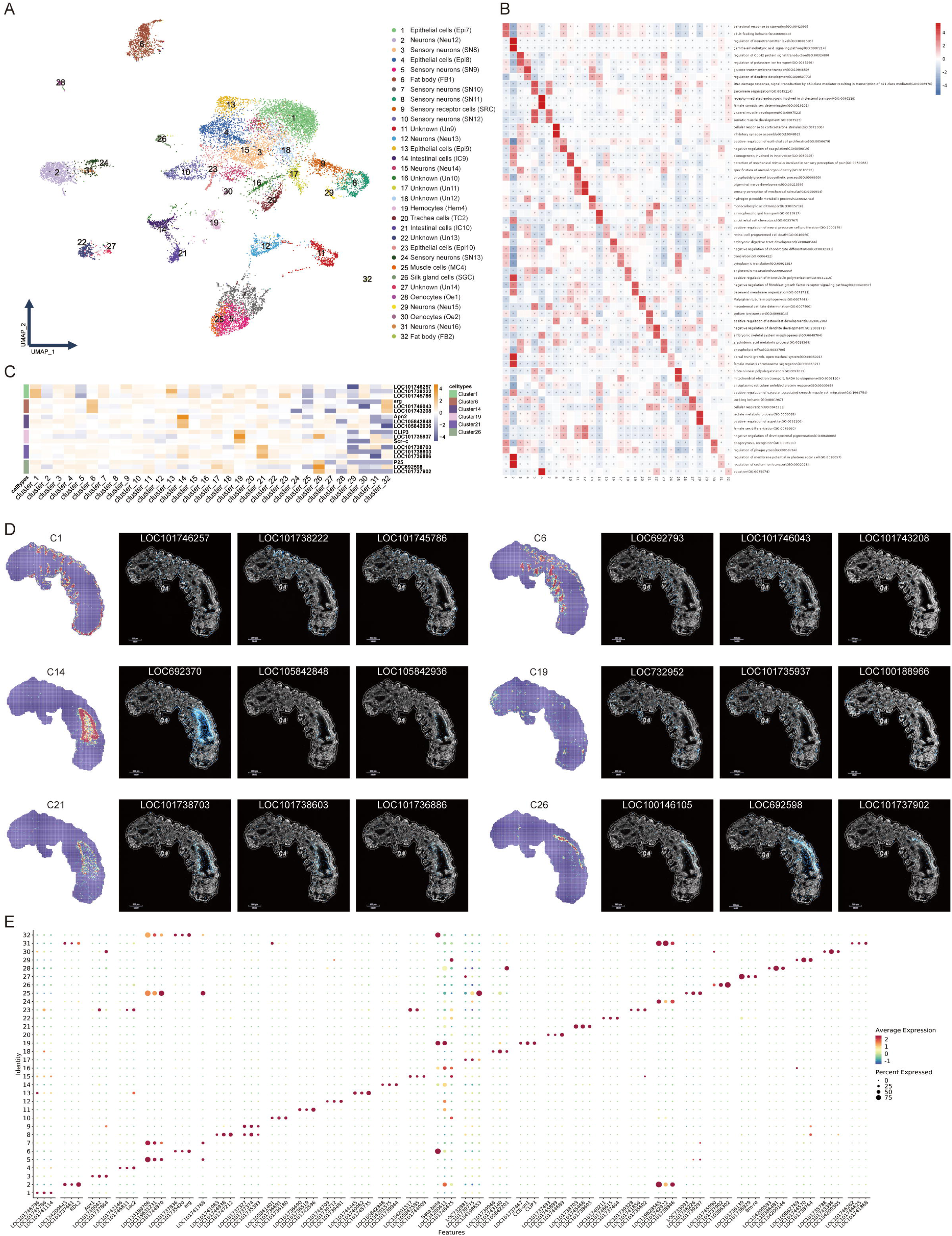
Cellular atlas of *Bombyx mori* embryos at 9 dpo. **A**, UMAP visualization of dimensionality reduction and cell-type annotations for the 9 dpo silkworm egg samples. **B**, GSVA heatmap of the cell clusters at 9 dpo. Based on the pathway activity scores calculated for individual cells using GSVA, differential testing was performed by comparing each cell group with all other cell groups. For each group, the two pathways with the highest *t*-statistics among those satisfying *p* < 0.05 were selected for heatmap visualization. **C**, Heatmap of selected marker genes at 9 dpo. These genes were used for the spatial localization shown in Fig D. **D**, Spatial transcriptomic mapping of selected cell clusters at 9 dpo and the spatial localization of representative genes on the spatial transcriptomic array. For each cell cluster, the first panel shows the RCTD mapping result. The color gradient from blue to red represents the inferred weight or relative abundance of each cell cluster in individual spatial spots, estimated by RCTD using the single-cell reference data. **E**, Dot plot showing representative marker genes on the UMAP plot at 9 dpo.

Cluster 1 was annotated as Epithelial Cells (Epi 7). This cluster expressed *Cda4* (LOC101746257), which encodes a chitin deacetylase and is an important member of the insect chitin-modifying enzyme family. Cda4 catalyzes the deacetylation of chitin to produce chitosan and thereby contributes to the regulation of chitin fiber structure and function. Previous studies have shown that members of the chitin deacetylase family play important roles in insect cuticle formation, molting, and the maintenance of body-wall structural integrity^77^. Cluster 1 also expressed *MCT9* (LOC101738222). In the *B. mori*, MCT9 is primarily associated with uric acid metabolism and the physiological functions of epithelial tissues^78^. Given the roles of these genes in chitin metabolism, epithelial structural maintenance, and material transport, the expression profile of Cluster 1 is consistent with the molecular characteristics of epithelial cells.

Cluster 6 was annotated as Fat Body (FB1). This cluster expressed *alanine-glyoxylate* (LOC101737836), which encodes alanine-glyoxylate aminotransferase. This enzyme participates in amino acid metabolism and catalyzes transamination reactions involving alanine and glyoxylate, thereby contributing to glycine and pyruvate metabolism^79^. *Arg* (LOC692793) encodes arginase, which catalyzes the hydrolysis of arginine to produce ornithine and urea and is an important enzyme in arginine metabolism and urea cycle-related processes. Previous studies have shown that *arg* is expressed in both fat body and muscle tissues^80^. In addition, this cluster highly expressed LOC101743208 and multiple genes associated with lipoproteins and lipases, suggesting active lipid storage, lipid mobilization, and systemic metabolic regulation. Based on its features related to amino acid metabolism, nitrogen metabolism, and lipid metabolism, Cluster 6 was annotated as FB1.

Cluster 19 exhibited prominent characteristics associated with hemocytes and innate immunity and was therefore annotated as Hemocytes (Hem4). This cluster expressed *CLIP3* (LOC732952), which encodes a serine protease containing a Clip domain. CLIP family proteins participate in insect development and innate immune processes through proteolytic cascades, including melanization, pathogen recognition, and antimicrobial peptide production^81^. Cluster 19 also expressed *hemolectin* (LOC101735937) and *scavenger receptor type C* (LOC100188966). *Hemolectin* is a hemocyte-associated marker gene in insects and participates in cell aggregation, hemostasis, and coagulation, whereas the C-type scavenger receptor is involved in the recognition of pathogen-associated molecules, phagocytosis, and the maintenance of immune homeostasis. The co-expression of these genes collectively supports the hemocyte identity of Cluster 19.

Cluster 26 was annotated as Silk Gland Cells (SGC). This cluster specifically expressed *dimm* (LOC101736973), which is specifically expressed in the *B. mori* silk gland and may cooperate with *fib-H* to regulate the synthesis of the fibroin heavy chain^82,83^. *P25* (LOC100146105) is also closely associated with the formation of fibroin complexes^84^. In addition, Cluster 26 highly expressed silk-gland-associated marker genes, including *seroin 1* (LOC692598) and *Ugt2* (LOC100174975). *Seroin 1* is primarily expressed in the silk gland and exhibits antimicrobial activity^74^, whereas *Ugt2* participates in glycosylation and may be involved in the metabolism of endogenous compounds, detoxification, and developmental regulation^75^. Based on the expression of genes associated with fibroin synthesis, silk-gland-specific expression, and metabolic regulation, Cluster 26 was identified as SGC.

The embryonic development at 9 dpo is defined by established cellular homeostasis, ongoing metabolic adaptation, and progressive functional maturation of organ systems. GSVA highlighted processes related to neural regulation, transport, cell proliferation, and protein quality control (Fig 6B). Enrichment of terms associated with GABA signaling, neurotransmitter regulation, dendrite and axon development, and mechanosensation suggested continued maturation of the nervous system. Potassium, sodium, and glucose transport, in turn, reflected the importance of ion homeostasis and nutrient utilization. Terms related to sarcomere organization and visceral and somatic muscle development indicated that movement-associated tissues were still maturing.

Meanwhile, translation, cytoplasmic translation, mitochondrial electron transport, and cellular respiration suggested that protein synthesis and energy production remained active. The p53-associated DNA damage response, linear polyubiquitination, and the endoplasmic reticulum unfolded protein response pointed to ongoing cellular quality control and homeostatic regulation during this period of rapid development.

At 9 dpo, embryos had completed morphological development and entered the pre-hatching stage, correspondingly, the yolk and other intra-ovular contents were markedly depleted or nearly exhausted. To enhance histological visibility during sample preparation for spatial transcriptomics, the chorion was manually removed from selected 9-dpo embryos. As a result, these specimens exhibited altered gross morphology compared with intact-egg samples described previously (Fig 6C, 6D). The results showed that C1 Epi7 were broadly distributed along the outer boundary of the embryo, whereas C6 FB1 were localized to multiple regions in the mid-body. C19 Hem exhibited a relatively dispersed spatial pattern. Both C14 IC9 and C21 IC10 clusters were concentrated in the posterior region, where the gut lumen was clearly visible. C26 SGC mapped to a band-like region above the gut, consistent with the expected anatomical location of the silk glands. Overall, cell-cluster mapping in day 9 embryos revealed clear regional differences. Some candidate-gene signals corresponded to the regions assigned to their respective clusters, broadly supporting the cell identities inferred from the single-nucleus data.

It is worth noting that among the markers identified in our single-nucleus atlas, certain clusters harbor genes with undefined functions or ambiguous annotations. The application of joint snRNA-seq and spatial transcriptomic deconvolution enabled the assignment of some markers to distinct lineages. However, residual ambiguity persists for others, necessitating their provisional designation as unknown cell states. Defining the precise identity and functionality of these clusters will require subsequent validation through targeted functional experiments, refined 3D spatiotemporal modeling, and phylogenetic comparisons of homologous sequences.

### Identification and Functional Annotation of Midgut Lineage-Specific Cell Populations in the *B. mori* Embryo

In the preceding analyses, some cell populations remained difficult to classify. To assess the ability of our dataset to resolve heterogeneity within a specific lineage, we re-clustered 8,601 putative midgut-associated cells identified across developmental stages. This analysis yielded 22 clusters, which we manually annotated based on characteristic gene-expression patterns and previously reported midgut cell markers (Fig 7A, Tab 7, Supplementary Data 7). At early embryonic stages, we identified endodermal cells and the midgut primordium. The endodermal cell clusters (Clusters 1, 4, and 7) comprised 2,511 cells and collectively expressed multiple regulators associated with early embryonic development and cell-fate determination. These included Hox-A2, which is associated with embryonic brain development^85^; the transcription factor Nanoso, which may contribute to embryonic abdominal formation^86^, and the transcription factor GATA-6, which is highly expressed in the *B. mori* midgut^51^. Cluster 16 was annotated as the midgut primordium and was detected primarily in 3 dpo embryos. This cluster showed high expression of *abd-A*, *abd-B*, and *sox21b*, genes associated in *Drosophila* with the development of the midgut^87^, body segments^88^, or intestine^89^. It also expressed several Notch pathway-associated genes^90^, including *serrate* and the E(spl) family members *E(spl)malpha* and *E(spl)mbeta* ^91^. Together, these expression features are consistent with molecular programs involved in intestinal regionalization and cell-fate determination. We also identified two transitional midgut cell clusters with distinct expression profiles, Cluster 8 and Cluster 2. Cluster 8 was detected primarily in 3 and 5 dpo embryos and expressed SOX-14 (LOC101735842), a regulator associated in *Drosophila* with 20-hydroxyecdysone (20E)-dependent embryonic and tissue development^92,93^, *scabrous* (LOC101738052), a protein that can bind to and modify Notch receptor activity^94^, and dendritic arbor reduction protein 1 (LOC105842151), which is associated with the regulation of intestinal stem-cell proliferation^95^. This expression profile suggests that Cluster 8 may represent an earlier stage of differentiation. Cluster 2 was detected primarily in 7 and 9 dpo embryos and showed high expression of digestion-related genes, including *maltase A1*, *myrosinase 1*, and *glucosidase*, as well as the development-associated genes Hox-B4 (LOC101745466)^96^ and Zn finger homeodomain 2 (LOC101747010)^97^. These features suggest that Cluster 2 may be undergoing differentiation toward functional midgut cells. Accordingly, we provisionally annotated Cluster 8 and Cluster 2 as early-and late-stage transitional midgut cells, respectively. Apn4 (LOC100127099) is a previously reported marker of *B. mori* midgut cells and has been shown to be specifically expressed in larval midgut tissue^98^. Clusters 3, 6, 9, 20, and 21 all showed high *Apn4* expression. Further analysis of functional gene-expression profiles indicated that Clusters 3, 6, 20, and 21 exhibited transcriptional features of midgut epithelial cells, including expression of genes associated with digestion, absorption, and transport, such as ABC transporters, the trehalose transporter Tret1, and trypsin-4. By contrast, Cluster 9 displayed features associated with endocrine cells and specifically expressed several neuropeptide receptors, including NGR-A13 and NGR-A5. Previous single-nucleus transcriptomic studies of the larval *B. mori* midgut have reported that midgut endocrine cells express multiple neuropeptide receptors^99,100^. Cluster 9 also expressed juvenile hormone esterase (LOC101735768), which is associated with hormone metabolism and endocrine regulation^101^, and the transcription factor senseless-2 (LOC101737547), which is expressed in the embryonic midgut of *Drosophila*^102,103^. Cluster 11 expressed *notch-like protein* (LOC100174829), a previously reported marker of *B. mori* midgut stem cells^99,100^, as well as midgut epithelial cell-associated genes, including *myrosinase 1*^104^, *maltase A1*, and the chitinase *chitinase 7*. This expression profile suggests that Cluster 11 may already exhibit some features of differentiation; we therefore provisionally annotated it as a midgut stem cell-like population.

**Fig. 7.**
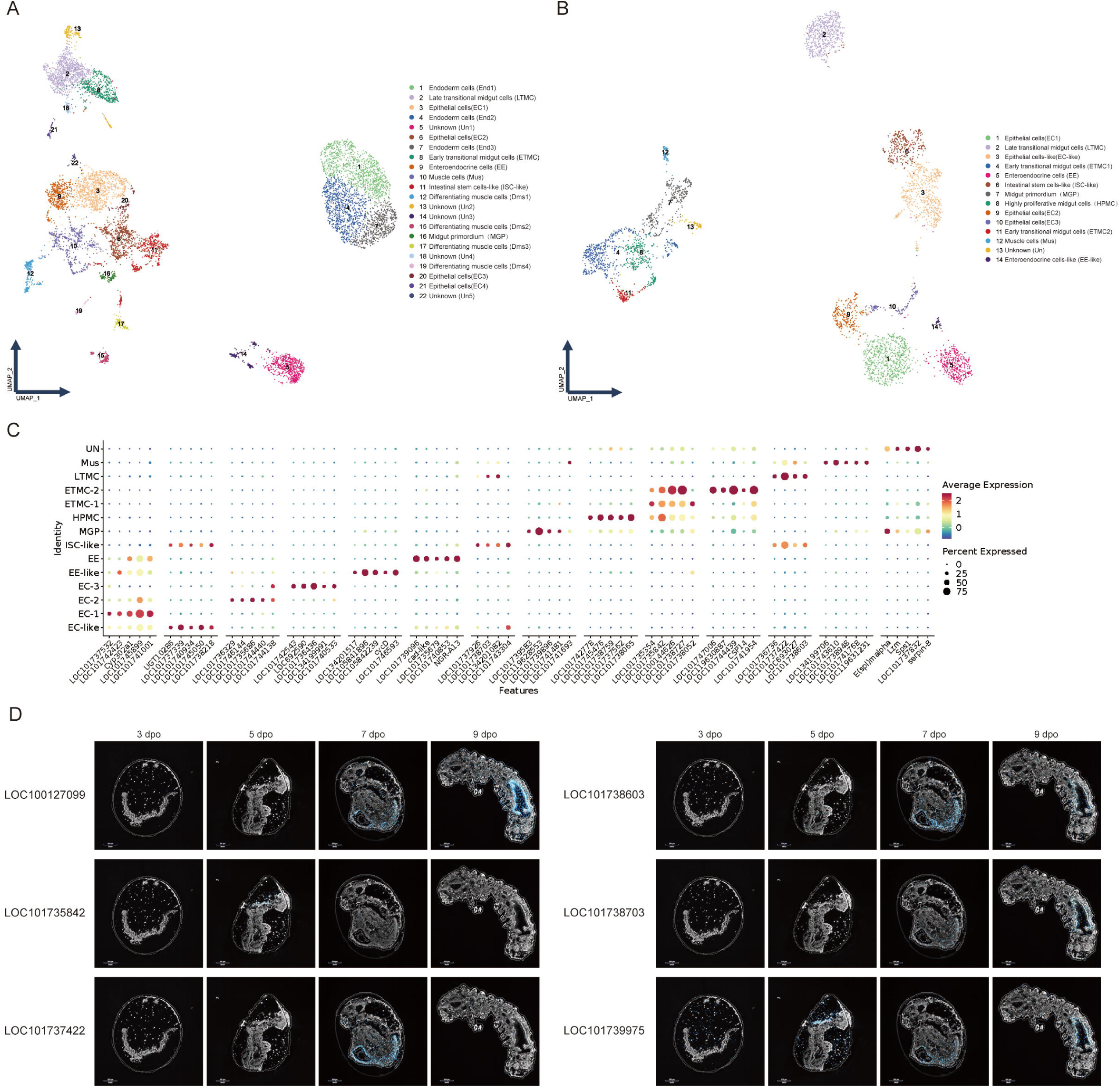
Global atlas of midgut-lineage cells. **A**, UMAP visualization of midgut-associated cells across all developmental stages. Based on the cell-type annotations obtained at the five developmental time points, putative midgut cell populations were selected for dimensionality reduction, clustering, and cell-type annotation. **B**, UMAP visualization of subclusters within the global midgut-cell atlas. The midgut cell clusters identified in **A** were subjected to further subclustering and annotation. **C**, Dot plot of marker genes for the midgut cell clusters. The five top-ranked marker genes for the midgut cell populations identified in **B** are shown. **D**, Spatiotemporal localization of midgut marker genes in spatial transcriptomic data. The spatial localization patterns of selected midgut marker genes in spatial transcriptomic samples collected at 3, 5, 7, and 9 dpo are shown.

Given that the transcriptional profiles of some clusters remained insufficiently resolved, we performed a second round of clustering on the 4,099 cells initially identified as midgut cells to further characterize transcriptional heterogeneity within the midgut. This analysis identified 14 midgut cell subtypes (Fig 7B, 7C, Tab 8, Supplementary Data 8). Based on Apn4 expression, we identified six late-stage midgut cell clusters (Clusters 1, 3, 5, 9, 10, and 14). The midgut epithelial cell clusters (Clusters 1, 9, and 10) were further classified into EC1–EC3 and collectively exhibited transcriptional features primarily associated with digestion and absorption. EC1 expressed several cuticular proteins, including Cuticular protein RR-2 motif 102, Cuticular protein RR-2 motif 100, and Cuticular protein RR-2 motif 88. EC2 and EC3, in contrast, specifically expressed multiple aminopeptidases. Their expression profiles differed: EC2 showed more prominent features associated with amino acid transport, whereas EC3 was more strongly associated with digestive functions. The endocrine cell cluster (Cluster 5; EE) displayed an expression profile consistent with that described above. Two additional clusters (Clusters 3 and 14) both highly expressed the midgut stem-cell marker notch-like protein, while exhibiting transcriptional features of epithelial and endocrine cells, respectively. This suggests that they may represent transitional states of differentiation; accordingly, we provisionally annotated them as EC-like and EE-like, respectively. Subclustering also identified a cell cluster (Cluster 8) found predominantly in 3 and 5 dpo embryos and characterized by a strong proliferation-associated expression profile. This cluster highly expressed the cell-cycle-related genes CDK1, Cyclin B, and Cyclin B3^105,106^, as well as the mitosis-associated genes Kinesin-like protein 3A^107^, Kinesin-like protein KIF18A^108^, and Protein regulator of cytokinesis 1^109^. These features suggest that Cluster 8 may represent an actively proliferating cell population involved in early midgut formation. Overall, the transcriptional profiles of the midgut primordium, transitional midgut cells, and midgut stem cell-like cells were broadly consistent with those described above.

We further characterized the spatial expression patterns of midgut-associated genes in *B. mori* embryos using spatial transcriptomic analysis (Fig 7D). The previously reported midgut marker Apn4 (LOC100127099) was predominantly enriched in the midgut region at late embryonic stages (7–9 dpo), supporting its use as a candidate marker for midgut cells at these stages. By integrating the cell-cluster annotations from the single-nucleus transcriptomic analysis with spatial expression profiles, we identified additional candidate markers that may aid in identifying midgut-associated cell populations. Sodium/potassium/calcium exchanger 4 (LOC101739975) showed a midgut-specific spatial expression pattern from 5 to 9 dpo. The transcription factor SOX-14 (LOC101735842) was predominantly enriched in the midgut region in the upper-left portion of the 5 dpo embryo section. Adhesion G-protein coupled receptor G2 (LOC101737422) was mainly enriched in the abdominal midgut region at 7 dpo. The spatial expression patterns of myogenesis-regulating glycosidase (LOC101738603) and putative inorganic phosphate cotransporter (LOC101738703) resembled that of Apn4, with both genes predominantly enriched in the midgut region at 7–9 dpo. These findings provide spatial evidence for an association between these genes and midgut-associated cell populations and suggest that they may help identify midgut cell populations at different developmental stages.

## Discussion

This study integrated single-nucleus and spatial transcriptomic data to construct a cellular atlas of *B. mori* embryos spanning developmental stages from 1 dpo to the pre-hatching stage. By systematically identifying the major cell populations and their transcriptional profiles at different developmental stages, we characterized the progressive changes in embryonic cellular composition, from early regionalization to tissue differentiation and pre-hatching maturation. Spatial transcriptomic analysis further provided preliminary information on the tissue localization of selected cell populations and genes. Overall, this atlas provides a reference resource for investigating *B. mori* embryogenesis at cellular resolution and establishes a foundation for future studies of tissues development and the establishment of the immune system.

Integration of cell-type annotations and functional analyses across developmental stages showed that *B. mori* embryos undergo a progressive transition from early germ-layer-associated states to organ primordium formation, tissue differentiation, and pre-hatching maturation. This overall pattern is consistent with previous descriptions of *B. mori* embryonic morphogenesis and organ development^15,25,110^. The single-nucleus transcriptomic data further extend these observations by providing cell-level information on population composition and molecular characteristics across developmental stages.

At 1 dpo, we identified cell populations associated with the ectoderm, endoderm, mesoderm, and neuroectoderm. GSVA revealed prominent activity of processes related to anterior–posterior axis specification, trunk segmentation, cell migration, and extracellular matrix organization, suggesting that the embryo may be undergoing regionalization and tissue remodeling at this stage. By 3 dpo, cellular composition had become more diverse, with populations associated with segment differentiation, appendages, neurons, sensory neurons, the gut, and tracheal primordia. Concurrent enrichment of processes related to cell polarity, the microtubule cytoskeleton, neuronal migration, and gliogenesis reflected the progression of embryonic organization and nervous system differentiation.

At 5–7 dpo, cell populations associated with muscle, epithelium, the nervous system, and neuromuscular development emerged progressively. By 7 dpo, tracheal cells, silk gland precursor cells, and fat body precursor cells were also detected. These changes are consistent with tissue differentiation and organogenesis during mid-to-late embryogenesis. Enrichment of pathways related to somite organization, myofibril assembly, planar cell polarity, and epithelial cell proliferation suggests that differentiation of tissues such as muscle and the body wall may proceed alongside embryonic morphogenesis. Expression signatures associated with neuronal connectivity and neurotransmitter regulation further suggest that the nervous system is progressively establishing the molecular basis required for signal transmission. It should be noted, however, that pathway enrichment reflects the expression characteristics of associated genes and does not, by itself, demonstrate that the corresponding tissues have acquired mature function. These findings therefore require further validation through histological or functional studies.

At 9 dpo, we detected cell populations corresponding to sensory neurons, fat body, and silk glands. Enrichment of pathways related to ion and glucose transport, protein translation, mitochondrial energy metabolism, and cellular quality control suggests that pre-hatching embryos maintain active metabolic and homeostatic processes. Overall, the observed changes in cell populations are consistent with embryonic progression from regionalization and organ primordium formation to tissue maturation, and provide cellular and molecular insights into these morphological processes. Compared with single-cell studies of other insect embryos, existing Drosophila embryonic atlases have been used to characterize embryonic cell composition, germ-layer differentiation, axial patterning, and stage-specific expression changes^6,111,112^. The early regionalization, nervous system differentiation, and organ primordium formation observed in this study correspond to several fundamental processes in insect embryogenesis, suggesting that some developmental features may be conserved. However, *B. mori* belongs to the order Lepidoptera and differs from the dipteran *Drosophila* in its phylogenetic background, embryonic development, and tissue biology^113^. Thus, the *B. mori* embryonic atlas is not merely a supplement to existing Drosophila atlases; it also provides a reference for comparing developmental processes and cell-type characteristics across insect groups. Future comparative studies could examine which cell types and molecular programs are conserved across species and which are associated with developmental features specific to *B. mori* or Lepidoptera. Spatial transcriptomic analysis provided preliminary tissue-localization information for cell populations and candidate genes identified by single-nucleus sequencing. Whereas single-nucleus transcriptomics primarily resolves transcriptional differences among cell populations, spatial data help interpret expression signals in the context of embryonic tissue architecture.

A cellular and spatial reference atlas of normal *B. mori* embryos can provide a baseline for future analyses of developmental abnormalities and infection-associated changes. In future studies, datasets from infected embryos at matched developmental stages could be integrated with the present atlas to investigate infection-associated changes in cellular composition, cell-type-specific transcriptional responses, and spatial organization. Developmental stage should be matched as closely as possible, because infection-induced developmental delay may affect cellular composition and gene expression. It is therefore important to distinguish infection-specific effects from changes attributable to differences in developmental stage.

This study has several limitations. First, cell-type annotation was based primarily on known marker genes and related functional information. The functions of some differentially expressed genes remain unknown or incompletely annotated, limiting the assignment of cell identity and the interpretation of biological function for certain populations. Accordingly, some populations can currently be classified only as unknown or provisionally annotated cell types. Future studies could combine more comprehensive spatiotemporal expression data, homologous gene comparisons, and functional experiments to systematically validate highly expressed candidate genes with uncharacterized functions and progressively refine cell-type annotations. Second, the spatial transcriptomic analysis was based primarily on sections from the central region of the embryo, resulting in limited spatial coverage. Differences in section position and orientation among eggs may also affect comparisons of spatial distribution patterns. Future studies could increase the number of serial sections, perform inter-section spatial registration and three-dimensional reconstruction, and validate the localization of candidate cell populations and genes using tissue clearing and in situ hybridization. These approaches would facilitate the construction of a more comprehensive three-dimensional spatiotemporal cellular atlas of the *B. mori* embryo.

In summary, this study establishes a cellular atlas spanning multiple key developmental stages of the *B. mori* embryo. It characterizes the transcriptional profiles and stage-dependent compositional changes of major cell populations, as well as selected spatial distribution patterns, and provides candidate cell-type markers and cell-population-associated expression clues for genes with uncharacterized functions. This atlas offers a reference for investigating cell types and developmental processes in *B. mori* embryos, and provides a foundation for future comparisons of embryogenesis across insect species and analyses of infection-associated changes in embryonic cellular composition and transcriptional states.

## Methods

### Experimental animal preparation and sample collection

The “305” strain of *B. mori* (domestic silkworm) was provided by the State Key Laboratory of Resource Insects, Southwest University. Silkworms were reared under standard laboratory conditions following a natural photoperiod. Larvae were fed fresh mulberry leaves until mating and oviposition. At 1, 3, 5, 7, and 9 days post-oviposition (dpo: 1 dpo corresponds to 24 h after oviposition), 1,000 silkworm eggs were randomly selected at each time point for single-nucleus suspension preparation. All procedures for nucleus suspension preparation were performed on ice. Frozen tissue samples were minced in 1 mL of nuclear lysis buffer (NST), which contained 0.1% NP-40, 10 mM Tris-HCl (pH 7.4), 146 mM NaCl, 1 mM CaCl₂, 21 mM MgCl₂, and 1 U/μL RNase inhibitor (Thermo Scientific,Cat.No.EO0382). The lysis duration was 7 minutes. Complete cell lysis was confirmed by trypan blue staining and microscopic examination. Subsequently, 1 mL of ST Wash buffer (10 mM Tris-HCl, 146 mM NaCl, 1 mM CaCl₂, 21 mM MgCl₂, 0.01% BSA (MACS, Cat.No.1000076), and 40 U/mL RNase inhibitor was added. The lysate was filtered through a 40-µm cell strainer (Falcon, cat.no. 352340), and the filtrate was transferred to a 15 mL centrifuge tube. The strainer was rinsed with additional ST Wash buffer, and the rinse solution was combined with the nuclear filtrate. The mixture was centrifuged at 500 × g for 5 minutes at 4°C using a horizontal rotor. The nuclear pellet was resuspended in 5 mL of PBS containing 1% BSA, followed by another centrifugation step. Finally, nuclei were resuspended in 100 μL of PBS with 1% BSA. Nuclear viability and concentration were assessed via trypan blue staining and microscopic counting.

### Single-nucleus RNA sequencing library construction

The nuclear suspension was diluted to a concentration of 700-1,200 nuclei/μL using PBS with 1% BSA. Single-nucleus RNA sequencing libraries were prepared according to the manufacturer’s protocol for the 10× Genomics Chromium Next GEM Single Cell 3ʹ Reagent Kit v3.1 (Cat.no. PN-1000268). The constructed libraries were sequenced on the Illumina Nova 6000 PE150 platform for high-throughput sequencing.

### snRNA-seq data processing

The FASTQ files were processed and aligned to the NCBI *B. mori* reference genome (GCF_030269925.1) using Cell Ranger software (version 9.0.0) from 10x Genomics, with unique molecular identifier (UMI) counts summarized for each barcode. The UMI count matrix was then analyzed using Seurat^114^ (version 4.0.0) R package. To remove low-quality cells and likely multiplet captures, a set of criteria were conducted: Cells were filtered by (1) gene numbers (gene numbers < 200), (2) UMI (UMI < 0.7), (4) percentage of mitochondrial RNA UMIs (proportion of UMIs mapped to mitochondrial genes > 5%). Subsequently, the DoubletFinder^115^ package (version 2.0.3) was used to identify potential doublets. To obtain the normalized gene expression data, library size normalization was processed using the NormalizeData function. Top 2000 highly variable genes (HVGs) were calculated using the Seurat function FindVariableGenes (mean.function=FastExpMean, dispersion.function= FastLogVMR). Principal-component analysis (PCA) was performed to reduce the dimensionality with RunPCA function. Then, the RunHarmony function in harmony^116^ (version 1.0) R package was performed to remove the batch effects. Graph-based clustering was performed to cluster cells according to their gene expression profile with the FindClusters function. Cells were visualized using a 2-dimensional Uniform Manifold www.oebiotech.com Approximation and Projection (UMAP) algorithm with the RunUMAP function. The FindAllMarkers function (test.use = presto) was used to identify marker genes of each cluster. GO enrichment and KEGG pathway enrichment analysis of DEGs were respectively performed using R(version 4.0.3) based on the hypergeometric distribution. The sequencing and bioinformatics analysis were provided by OE Biotech Co., Ltd. (Shanghai, China).

### Pseudotime Analysis

The developmental pseudotime was determined with the Monocle2^117^ package(version 2.9.0). The raw count was first converted from Seurat object into CellDataSet object with the importCDS function in Monocle. The differentialGeneTest function of the Monocle2 package was used to select ordering genes (qval < 0.01) which were likely to be informative in the ordering of cells along the pseudotime trajectory. The dimensional reduction clustering analysis was performed with the reduceDimension function, followed by trajectory inference with the orderCells function using default parameters. Gene expression was plotted with the plot_genes_in_pseudotime function to track changes over pseudo-time.

### Gene Set Variation Analysis (GSVA)

To perform the Gene Set Variation Analysis, the GSEABase package (version 1.44.0) was used to load the gene set file which was downloaded and processed from KEGG database (https://www.kegg.jp/) and Gene Ontology database (https://geneontology.org/). To assign pathway activity estimates to individual cells, we applied GSVA^118^ using standard settings, as implemented in the GSVA package (version 1.30.0). The differences in pathway activities scored per cell were calculated with LIMMA package (version 3.38.3).

### Spatial transcriptomic sequencing library construction

*B. mori* egg samples from different developmental days were orderly arranged in embedding molds, and the tissues were embedded in OCT Medium (SAKURA, cat. no. 4583), snap-frozen on dry ice, and stored at-80°C. OCT-embedded samples were sent to Shanghai OE Biotech Co., Ltd. (Shanghai, China) for spatial transcriptomic library preparation, sequencing and data analysis. Briefly, frozen sections of 10 μm thickness were prepared using a Leica CM1950 Microtome Cryostat (Leica Microsystems, Germany) at-20 °C, and then mounted on Stereo-seq permeabilization chips (STOmics, cat.no.201SP11118) or transcriptomics chips (STOmics,cat.no.201ST13114 for 1×1 cm chips). The following spatial transcriptomics procedures, including methanol fixation, H&E staining, image scanning, tissue permeabilization, reverse transcription, tissue removal, cDNA recovery and purification, were performed according to the manufacturer’s manual and previous publications^119,120^. DNA library construction was performed using Stereo-seq 16 Barcode library kit (STOmics, cat.no.101KL160) and final libraries were sequenced on the BGI DNBSEQ-T7 sequencing platform using PE75 mode.

### Spatial transcriptomic sequencing data processing

The Stereo-seq analysis workflow (SAW, version 8.2.1, https://github.com/STOmics/SAW) was used to process Stereo-seq spatial RNA-seq output and brightfield microscope images in order to detect tissue, align reads to *B. mori* GCF_030269925.1 reference genome, generate feature-CID (coordinate identifier) matrices, perform clustering and gene expression analysis, and place spots in spatial context on the slide image. In this step, a bin size of 50 (50 × 50 spots) was used as the analytical unit for the annotation of Stereo-seq slides. The molecular identifier (MID) count matrix was then processed using Seurat^114^ (version 4.3.0) R package. Sctransform^121^ was used to normalize data and identify top 3000 highly variable genes (HVGs). Principal component analysis (PCA) was conducted to reduce dimensionality on the log transformed gene-barcode matrices of top variable genes. Then, the RunHarmony function in harmony^116^ (version 1.0) R package was performed to remove the batch effects. Graph-based clustering was performed to cluster bins according to their gene expression profile with the FindClusters function. Bins were visualized using a 2-dimensional Uniform Manifold Approximation and Projection (UMAP) algorithm with the RunUMAP function. The FindAllMarkers function (test.use = bimod) was used to identify marker genes of each cluster. GO enrichment and KEGG pathway enrichment analysis of DEGs were respectively performed using R (version 4.0.3) based on the hypergeometric distribution. The sequencing and bioinformatics analysis were provided by OE Biotech Co., Ltd. (Shanghai, China).

### RCTD Analysis

According to our single-nucleus RNA sequencing dataset, RCTD^122^ (version 1.1.0) was applied to infer the cell-type composition of each bin. Specifically, the default parameters were used in the creat.RCTD function, except for minimum number of cell > 1 for each cell types and minimum of unique molecular identifier (UMI) counts > 1 per pixel, and doublet_mode parameter was set to FALSE in the run.RCTD function.

## Data availability

The raw data of RNA sequencing have been deposited in the Genome Sequence Archive in National Genomics Data Center ^123,124^(https://ngdc.cncb.ac.cn/gsa) under the number CRA053965 (single-cell transcriptome) and CRA053967 (spatial transcriptome).

## Acknowledgements

The authors gratefully appreciate the individuals and organizations that provided the software used in this study, as well as Shanghai OE Biotech Co., Ltd for single-cell transcriptome and spatial transcriptome sequencing.

## Author contributions

Writing-original draft preparation: YBD, TJL, YZ, JC^b^; writing-review and editing: GQP, TL, XDL, JC^b^, ZGS; Conceptualization: GQP, TL, XDL, BHX, LFS, CFL, ZYZ; Resources: BHX, CFL, LFS, SL, YQ, ZGS; methodology: YZ, YBD, TJL, KQC, GC, CXW; Data Analysis: YBD, YZ, TJL, ZGS, CRL, MSR, SL, JC^a^, KQC, QXH, YCS, SBW, LTL, YLT, QL, XZM; Data curation: GQP, TL, XDL, JC^b^, SL, MXL, JLB, JHW, LC.

---

**Table 1.** Cluster Composition of the Comprehensive Single-nucleus Atlas.

| clusters | cell_number | Freq (%) |
|---|---|---|
| 1 | 7965 | 9.74 |
| 2 | 7360 | 9.00 |
| 3 | 6657 | 8.14 |
| 4 | 5589 | 6.83 |
| 5 | 3751 | 4.59 |
| 6 | 3686 | 4.51 |
| 7 | 3604 | 4.41 |
| 8 | 3159 | 3.86 |
| 9 | 2730 | 3.34 |
| 10 | 2426 | 2.97 |
| 11 | 2372 | 2.90 |
| 12 | 2262 | 2.77 |
| 13 | 2178 | 2.66 |
| 14 | 2154 | 2.63 |
| 15 | 2148 | 2.63 |
| 16 | 2002 | 2.45 |
| 17 | 1915 | 2.34 |
| 18 | 1628 | 1.99 |
| 19 | 1602 | 1.96 |
| 20 | 1538 | 1.88 |
| 21 | 1523 | 1.86 |
| 22 | 1322 | 1.62 |
| 23 | 1082 | 1.32 |
| 24 | 1028 | 1.26 |
| 25 | 1012 | 1.24 |
| 26 | 996 | 1.22 |
| 27 | 948 | 1.16 |
| 28 | 909 | 1.11 |
| 29 | 908 | 1.11 |
| 30 | 856 | 1.05 |
| 31 | 815 | 1.00 |
| 32 | 799 | 0.98 |
| 33 | 623 | 0.76 |
| 34 | 425 | 0.52 |
| 35 | 401 | 0.49 |
| 36 | 304 | 0.37 |
| 37 | 215 | 0.26 |
| 38 | 205 | 0.25 |
| 39 | 193 | 0.24 |
| 40 | 174 | 0.21 |
| 41 | 164 | 0.20 |
| 42 | 101 | 0.12 |
| 43 | 43 | 0.05 |

**Table 2.** Identification Results and Statistics of Cell Clusters at 1dpo.

| Sample_Cluster | Name | Abbreviation | cell_number | freq(%) |
|---|---|---|---|---|
| 1_1 | Neuroectodermal Cells | Ned1 | 2545 | 16.33 |
| 1_2 | Ectodermal Cells | Ect1 | 1912 | 12.27 |
| 1_3 | Ectodermal Cells | Ect2 | 1887 | 12.11 |
| 1_4 | Endodermal Cells | End1 | 1541 | 9.89 |
| 1_5 | Mesodermal Cells | Mes1 | 1342 | 8.61 |
| 1_6 | Lipid-metabolizing Cells | LMC | 1271 | 8.15 |
| 1_7 | Ectodermal Cells | Ect3 | 1247 | 8.00 |
| 1_8 | Mesodermal Cells | Mes2 | 997 | 6.40 |
| 1_9 | Neuroectodermal Cells | Ned2 | 990 | 6.35 |
| 1_10 | Endodermal Cells | End2 | 970 | 6.22 |
| 1_11 | Metabolic Cells | Met1 | 445 | 2.86 |
| 1_12 | Unknown | Un1 | 439 | 2.82 |

**Table 3.** Identification Results and Statistics of Cell Clusters at 3dpo.

| Sample_Cluster | Name | Abbreviation | cell_number | freq(%) |
|---|---|---|---|---|
| 3_1 | Somitic Differentiation Cells | SDC1 | 3385 | 19.80 |
| 3_2 | Neurons | Neu1 | 2337 | 13.67 |
| 3_3 | Appendage Cells | AC1 | 2056 | 12.02 |
| 3_4 | Mesodermal Developmental Cells | MDC1 | 2029 | 11.87 |
| 3_5 | Neurons | Neu2 | 1673 | 9.78 |
| 3_6 | Neuromuscular Developmental Cells | NDC1 | 717 | 4.19 |
| 3_7 | Mesodermal Developmental Cells | MDC2 | 696 | 4.07 |
| 3_8 | Sensory Neurons | SN1 | 654 | 3.82 |
| 3_9 | Intestinal Cells | IC1 | 521 | 3.05 |
| 3_10 | Sensory Neurons | SN2 | 451 | 2.64 |
| 3_11 | Endodermal Developmental Cells | EDC1 | 445 | 2.60 |
| 3_12 | Tracheal Primordium Cells | TPC1 | 441 | 2.58 |
| 3_13 | Unknown | Un2 | 324 | 1.89 |
| 3_14 | Endodermal Developmental Cells | EDC2 | 298 | 1.74 |
| 3_15 | Neuromuscular Developmental Cells | NDC2 | 232 | 1.36 |
| 3_16 | Neurons | Neu3 | 218 | 1.27 |
| 3_17 | Neurons | Neu4 | 137 | 0.80 |
| 3_18 | Hemocytes | Hem1 | 130 | 0.76 |
| 3_19 | Oenocyte-like Cells | OLC1 | 103 | 0.60 |
| 3_20 | Neurons | Neu5 | 100 | 0.58 |
| 3_21 | Unknown | Un3 | 89 | 0.52 |
| 3_22 | Epithelial Cells | Epi1 | 63 | 0.37 |

**Table 4.** Identification Results and Statistics of Cell Clusters at 5dpo.

| Sample_Cluster | Name | Abbreviation | cell_number | freq(%) |
|---|---|---|---|---|
| 5_1 | Somitic Differentiation Cells | SDC2 | 3045 | 18.10 |
| 5_2 | Appendage Cells | AC2 | 2745 | 16.32 |
| 5_3 | Epithelial Cells | Epi2 | 1790 | 10.64 |
| 5_4 | Muscle Cells | MC1 | 1438 | 8.55 |
| 5_5 | Neurons | Neu6 | 994 | 5.91 |
| 5_6 | Unknown | Un4 | 862 | 5.12 |
| 5_7 | Unknown | Un5 | 689 | 4.10 |
| 5_8 | Muscle Cells | MC2 | 680 | 4.04 |
| 5_9 | Unknown | Un6 | 667 | 3.96 |
| 5_10 | Neuromuscular Developmental Cells | NDC3 | 653 | 3.88 |
| 5_11 | Intestinal Cells | IC2 | 649 | 3.86 |
| 5_12 | Tracheal Primordium Cells | TPC2 | 564 | 3.35 |
| 5_13 | Epithelial Cells | Epi3 | 471 | 2.80 |
| 5_14 | Neurons | Neu7 | 317 | 1.88 |
| 5_15 | Intestinal Cells | IC3 | 250 | 1.49 |
| 5_16 | Hemocytes | Hem2 | 249 | 1.48 |
| 5_17 | Sensory Neurons | SN3 | 238 | 1.41 |
| 5_18 | Neurons | Neu8 | 158 | 0.94 |
| 5_19 | Sensory Neurons | SN4 | 152 | 0.90 |
| 5_20 | Neuromuscular Developmental Cells | NDC4 | 99 | 0.59 |
| 5_21 | Neurons | Neu9 | 57 | 0.34 |
| 5_22 | Oenocyte-like Cells | OLC2 | 57 | 0.34 |

**Table 5.** Identification Results and Statistics of Cell Clusters at 7dpo.

| Sample_Cluster | Name | Abbreviation | cell_number | freq(%) |
|---|---|---|---|---|
| 7_1 | Epithelial Cells | Epi4 | 2587 | 15.95 |
| 7_2 | Intestinal Cells | IC4 | 1622 | 10.00 |
| 7_3 | Epithelial Cells | Epi5 | 1564 | 9.64 |
| 7_4 | Muscle Cells | MC3 | 1466 | 9.04 |
| 7_5 | Neurons | Neu10 | 1084 | 6.68 |
| 7_6 | Unknown | Un7 | 997 | 6.15 |
| 7_7 | Unknown | Un8 | 763 | 4.70 |
| 7_8 | Neuromuscular Developmental Cells | NDC5 | 736 | 4.54 |
| 7_9 | Sensory Neurons | SN5 | 695 | 4.28 |
| 7_10 | Neurons | Neu11 | 693 | 4.27 |
| 7_11 | Appendage Cells | AC3 | 689 | 4.25 |
| 7_12 | Intestinal Cells | IC5 | 642 | 3.96 |
| 7_13 | Fat Body Precursor Cells | FBP | 497 | 3.06 |
| 7_14 | Sensory Neurons | SN6 | 470 | 2.90 |
| 7_15 | Hemocytes | Hem3 | 302 | 1.86 |
| 7_16 | Unknown | Un9 | 277 | 1.71 |
| 7_17 | Sensory Neurons | SN7 | 257 | 1.58 |
| 7_18 | Trachea Cells | TC1 | 247 | 1.52 |
| 7_19 | Unknown | Un10 | 149 | 0.92 |
| 7_20 | Oenocyte-like Cells | OLC3 | 131 | 0.81 |
| 7_21 | Intestinal Cells | IC6 | 128 | 0.79 |
| 7_22 | Silk Gland Precursor Cells | SGP | 113 | 0.70 |
| 7_23 | Metabolic Cells | Met2 | 67 | 0.41 |
| 7_24 | Intestinal Cells | IC7 | 45 | 0.28 |

**Table 6.** Identification Results and Statistics of Cell Clusters at 9dpo.

| Sample_Cluster | Name | Abbreviation | cell_number | freq(%) |
|---|---|---|---|---|
| 9_1 | Epithelial Cells | Epi7 | 2515 | 15.68 |
| 9_2 | Neurons | Neu12 | 1128 | 7.03 |
| 9_3 | Sensory Neurons | SN8 | 1108 | 6.91 |
| 9_4 | Epithelial Cells | Epi8 | 1023 | 6.38 |
| 9_5 | Sensory Neurons | SN9 | 995 | 6.20 |
| 9_6 | Fat Body | FB1 | 939 | 5.85 |
| 9_7 | Sensory Neurons | SN10 | 889 | 5.54 |
| 9_8 | Sensory Neurons | SN11 | 591 | 3.68 |
| 9_9 | Sensory Receptor Cells | SRC | 522 | 3.25 |
| 9_10 | Sensory Neurons | SN12 | 492 | 3.07 |
| 9_11 | Unknown | Un11 | 482 | 3.00 |
| 9_12 | Neurons | Neu13 | 457 | 2.85 |
| 9_13 | Epithelial Cells | Epi9 | 444 | 2.77 |
| 9_14 | Intestinal Cells | IC9 | 433 | 2.70 |
| 9_15 | Neurons | Neu14 | 420 | 2.62 |
| 9_16 | Unknown | Un12 | 407 | 2.54 |
| 9_17 | Unknown | Un13 | 396 | 2.47 |
| 9_18 | Unknown | Un14 | 356 | 2.22 |
| 9_19 | Hemocytes | Hem4 | 327 | 2.04 |
| 9_20 | Trachea Cells | TC2 | 307 | 1.91 |
| 9_21 | Intestinal Cells | IC10 | 305 | 1.90 |
| 9_22 | Unknown | Un15 | 302 | 1.88 |
| 9_23 | Epithelial Cells | Epi10 | 292 | 1.82 |
| 9_24 | Sensory Neurons | SN13 | 197 | 1.23 |
| 9_25 | Muscle Cells | MC4 | 186 | 1.16 |
| 9_26 | Silk Gland Cells | SGC | 121 | 0.75 |
| 9_27 | Intestinal Cells | Un16 | 100 | 0.62 |
| 9_28 | Oenocytes | Oe1 | 93 | 0.58 |
| 9_29 | Neurons | Neu15 | 79 | 0.49 |
| 9_30 | Oenocytes | Oe2 | 58 | 0.36 |
| 9_31 | Neurons | Neu16 | 46 | 0.29 |
| 9_32 | Fat Body | FB2 | 32 | 0.20 |

**Table 7.**
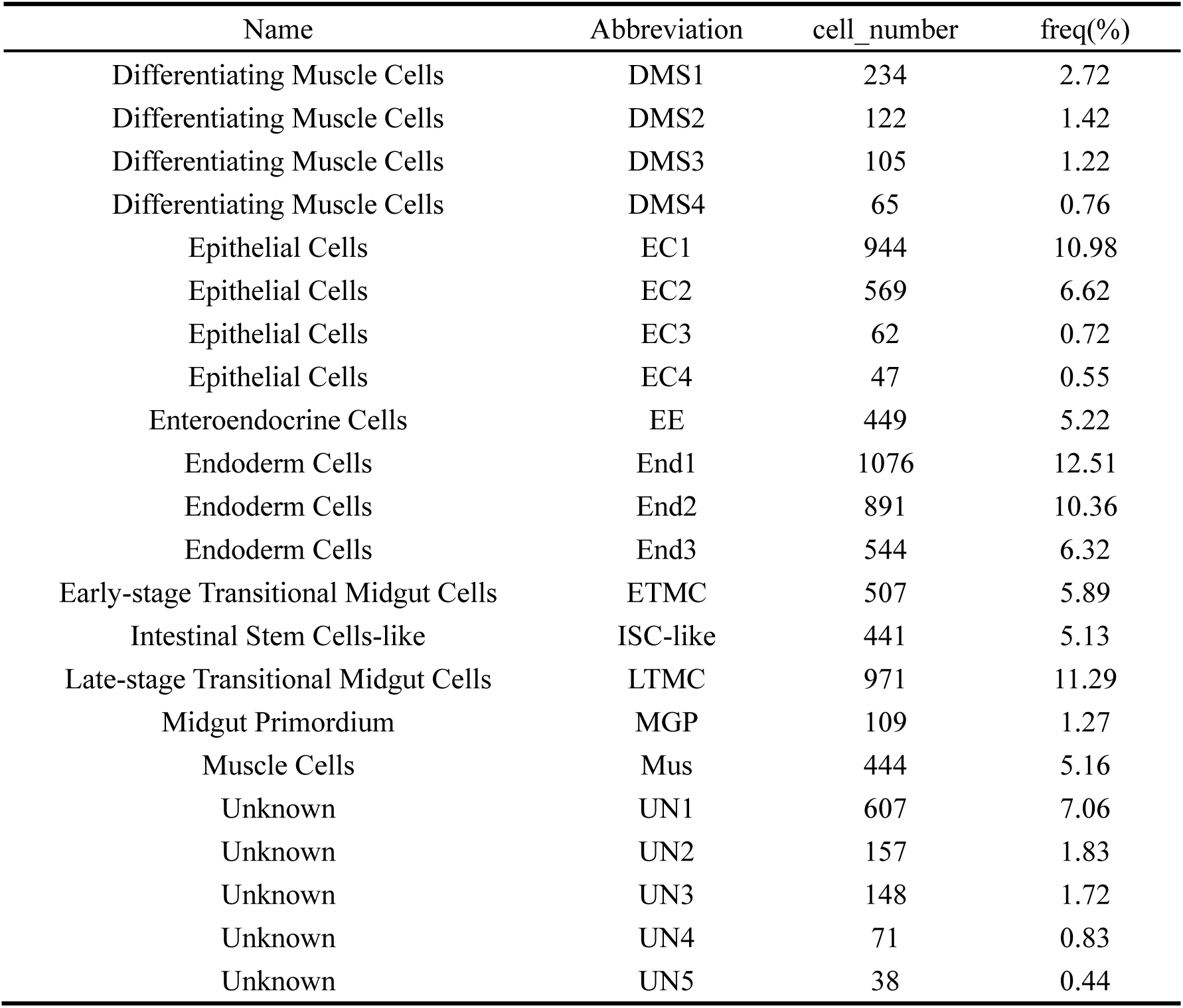
Identification and Statistics of Putative Midgut Cell Clusters in the Whole-Dataset Panorama.

**Table 8.** Fine-Grained Subclustering and Statistics of Curated Midgut Cell.

| Name | Abbreviation | cell_number | freq(%) |
|---|---|---|---|
| Epithelial Cells | EC-1 | 657 | 16.03 |
| Epithelial Cells | EC-2 | 230 | 5.61 |
| Epithelial Cells | EC-3 | 117 | 2.85 |
| Epithelial Cells-like | EC-like | 540 | 13.17 |
| Enteroendocrine Cells | EE | 380 | 9.27 |
| Enteroendocrine Cells-like | EE-like | 28 | 0.68 |
| Early-stage Transitional Midgut Cells | ETMC-1 | 464 | 11.32 |
| Early-stage Transitional Midgut Cells | ETMC-2 | 109 | 2.66 |
| Highly Proliferative Midgut Cells | HPMC | 274 | 6.68 |
| Intestinal Stem Cells-like | ISC-like | 289 | 7.05 |
| Late-stage Transitional Midgut Cells | LTMC | 602 | 14.69 |
| Midgut Primordium | MGP | 281 | 6.86 |
| Muscle Cells | Mus | 70 | 1.71 |
| Unknown | UN | 58 | 1.41 |

