## Supplemental Data 1-8 for "A Spatiotemporal Embryonic Atlas of Bombyx mori Integrating Single-Nucleus RNA Sequencing and Spatial Transcriptomics": Cluster1 GO top100.pdf

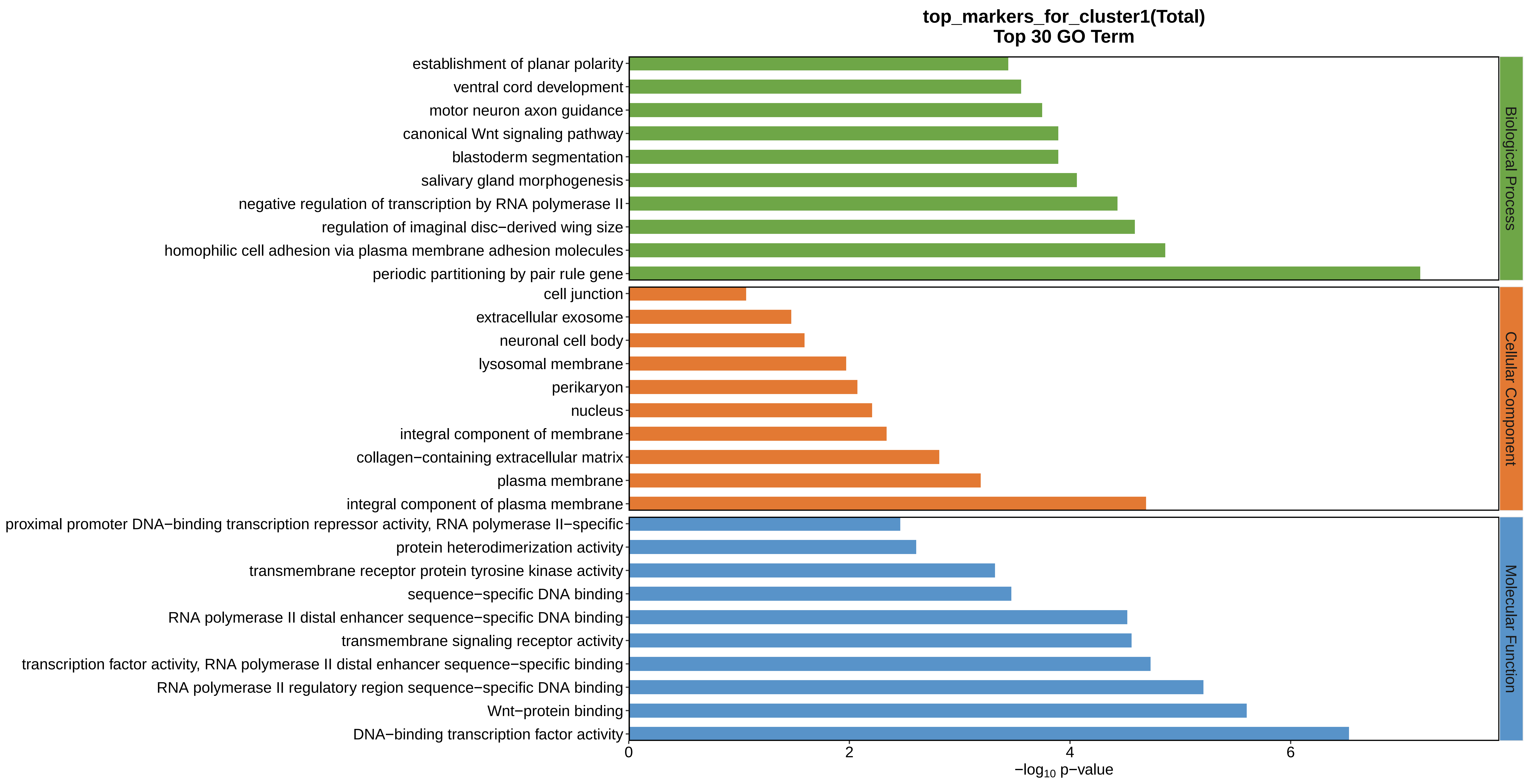
top_markers_for_cluster1(Total)
Top 30 GO Term
establishment of planar polarity
ventral cord development
motor neuron axon guidance
canonical Wnt signaling pathway
blastoderm segmentation
salivary gland morphogenesis
negative regulation of transcription by RNA polymerase II
regulation of imaginal disc−derived wing size
homophilic cell adhesion via plasma membrane adhesion molecules
periodic partitioning by pair rule gene
cell junction
extracellular exosome
neuronal cell body
lysosomal membrane
perikaryon
nucleus
integral component of membrane
collagen−containing extracellular matrix
plasma membrane
integral component of plasma membrane
proximal promoter DNA−binding transcription repressor activity, RNA polymerase II−specific
protein heterodimerization activity
transmembrane receptor protein tyrosine kinase activity
sequence−specific DNA binding
RNA polymerase II distal enhancer sequence−specific DNA binding
transmembrane signaling receptor activity
transcription factor activity, RNA polymerase II distal enhancer sequence−specific binding
RNA polymerase II regulatory region sequence−specific DNA binding
Wnt−protein binding
DNA−binding transcription factor activity
Biological Process
Cellular Component
Molecular Function
0
2
4
6
−log10 p−value
