## Supplemental Data 1-8 for "A Spatiotemporal Embryonic Atlas of Bombyx mori Integrating Single-Nucleus RNA Sequencing and Spatial Transcriptomics": Cluster2 GO top100.pdf

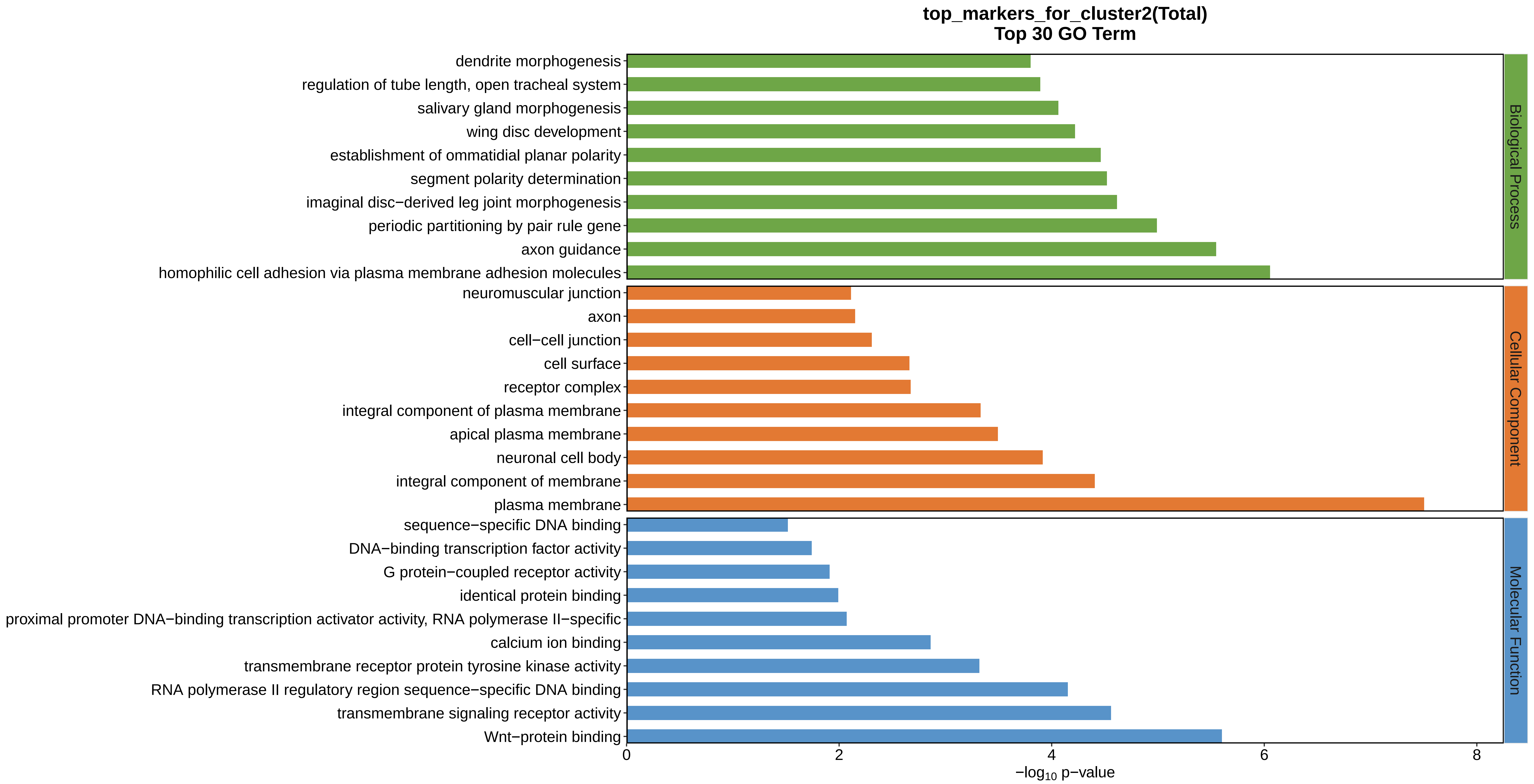
top_markers_for_cluster2(Total)
Top 30 GO Term
dendrite morphogenesis
regulation of tube length, open tracheal system
salivary gland morphogenesis
wing disc development
establishment of ommatidial planar polarity
segment polarity determination
imaginal disc−derived leg joint morphogenesis
periodic partitioning by pair rule gene
axon guidance
homophilic cell adhesion via plasma membrane adhesion molecules
neuromuscular junction
axon
cell−cell junction
cell surface
receptor complex
integral component of plasma membrane
apical plasma membrane
neuronal cell body
integral component of membrane
plasma membrane
sequence−specific DNA binding
DNA−binding transcription factor activity
G protein−coupled receptor activity
identical protein binding
proximal promoter DNA−binding transcription activator activity, RNA polymerase II−specific
calcium ion binding
transmembrane receptor protein tyrosine kinase activity
RNA polymerase II regulatory region sequence−specific DNA binding
transmembrane signaling receptor activity
Wnt−protein binding
Biological Process
Cellular Component
Molecular Function
0
2
4
6
8
−log10 p−value
