## Supplemental Data 1-8 for "A Spatiotemporal Embryonic Atlas of Bombyx mori Integrating Single-Nucleus RNA Sequencing and Spatial Transcriptomics": Cluster3 GO top100.pdf

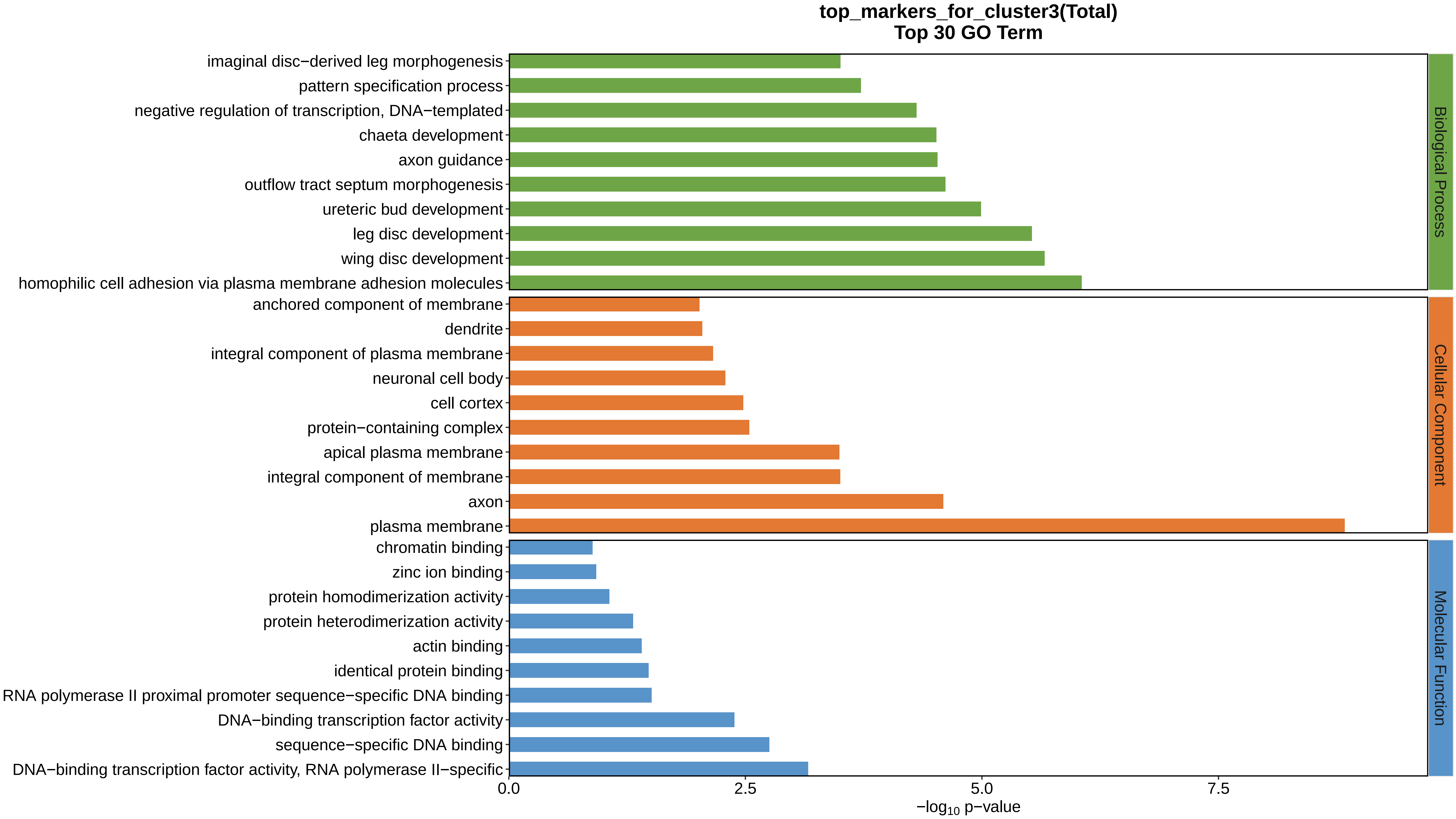

top_markers_for_cluster3(Total)
Top 30 GO Term
imaginal disc−derived leg morphogenesis
pattern specification process
negative regulation of transcription, DNA−templated
chaeta development
axon guidance
outflow tract septum morphogenesis
ureteric bud development
leg disc development
wing disc development
homophilic cell adhesion via plasma membrane adhesion molecules
anchored component of membrane
dendrite
integral component of plasma membrane
neuronal cell body
cell cortex
protein−containing complex
apical plasma membrane
integral component of membrane
axon
plasma membrane
chromatin binding
zinc ion binding
protein homodimerization activity
protein heterodimerization activity
actin binding
identical protein binding
RNA polymerase II proximal promoter sequence−specific DNA binding
DNA−binding transcription factor activity
sequence−specific DNA binding
DNA−binding transcription factor activity, RNA polymerase II−specific
Biological Process
Cellular Component
Molecular Function
0.0
2.5
5.0
7.5
−log10 p−value
