## Supplemental Data 1-8 for "A Spatiotemporal Embryonic Atlas of Bombyx mori Integrating Single-Nucleus RNA Sequencing and Spatial Transcriptomics": Cluster4 GO top100.pdf

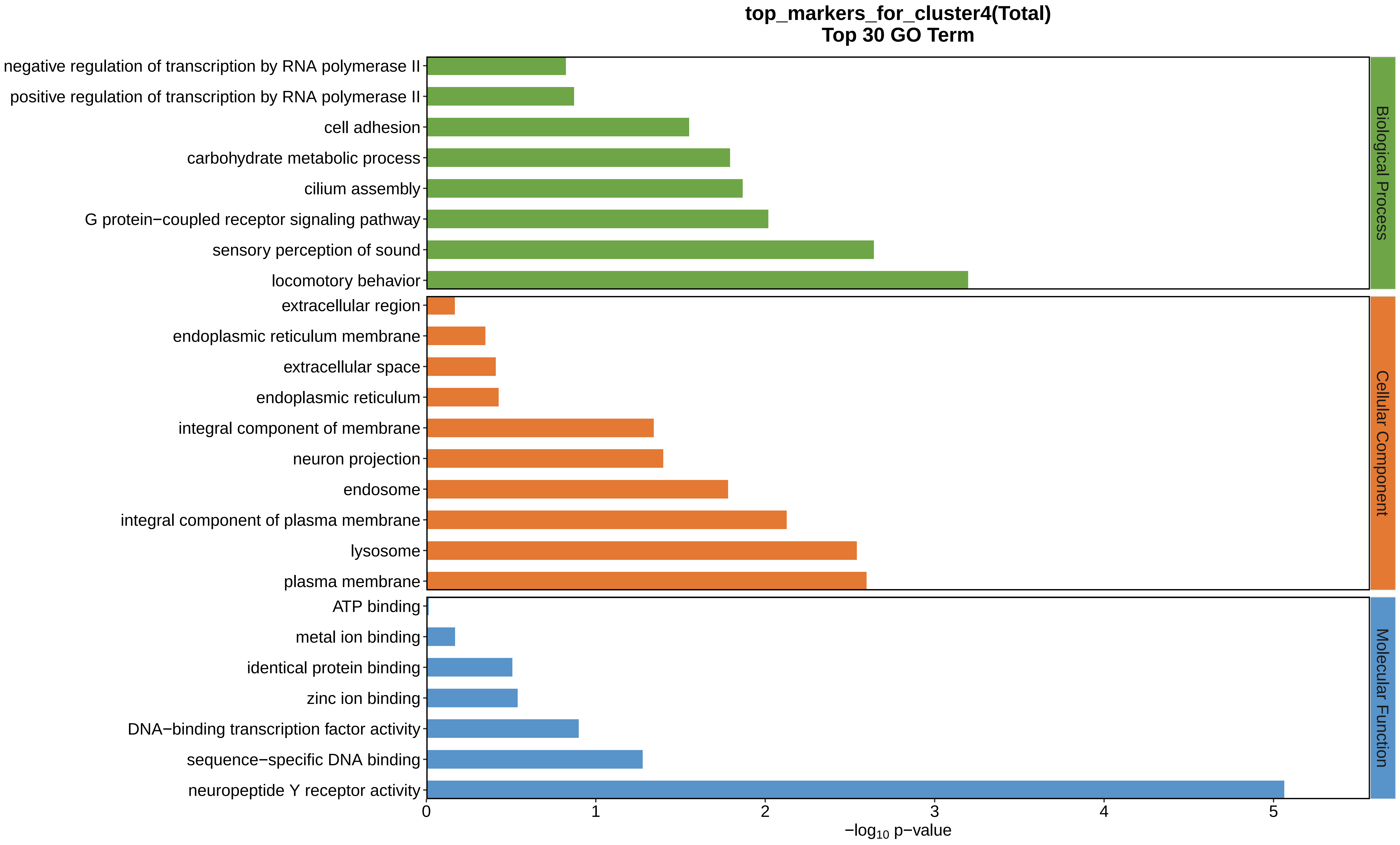
top_markers_for_cluster4(Total)
Top 30 GO Term
negative regulation of transcription by RNA polymerase II
positive regulation of transcription by RNA polymerase II
cell adhesion
carbohydrate metabolic process
cilium assembly
G protein−coupled receptor signaling pathway
sensory perception of sound
locomotory behavior
extracellular region
endoplasmic reticulum membrane
extracellular space
endoplasmic reticulum
integral component of membrane
neuron projection
endosome
integral component of plasma membrane
lysosome
plasma membrane
ATP binding
metal ion binding
identical protein binding
zinc ion binding
DNA−binding transcription factor activity
sequence−specific DNA binding
neuropeptide Y receptor activity
Biological Process
Cellular Component
Molecular Function
0
1
2
3
4
5
−log10 p−value
