## Supplemental Data 1-8 for "A Spatiotemporal Embryonic Atlas of Bombyx mori Integrating Single-Nucleus RNA Sequencing and Spatial Transcriptomics": Cluster5 GO top100.pdf

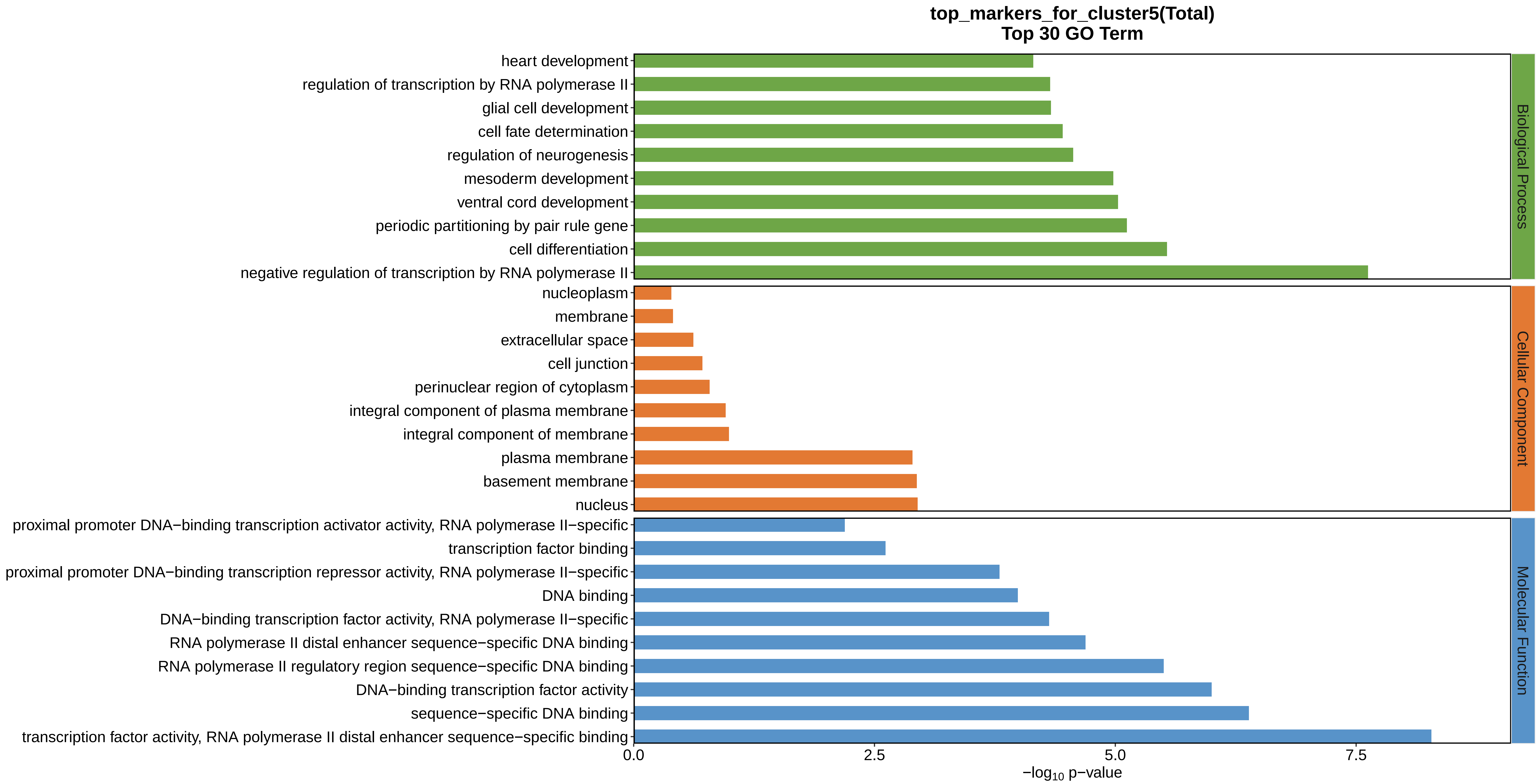
top_markers_for_cluster5(Total)
Top 30 GO Term
heart development
regulation of transcription by RNA polymerase II
glial cell development
cell fate determination
regulation of neurogenesis
mesoderm development
ventral cord development
periodic partitioning by pair rule gene
cell differentiation
negative regulation of transcription by RNA polymerase II
nucleoplasm
membrane
extracellular space
cell junction
perinuclear region of cytoplasm
integral component of plasma membrane
integral component of membrane
plasma membrane
basement membrane
nucleus
proximal promoter DNA−binding transcription activator activity, RNA polymerase II−specific
transcription factor binding
proximal promoter DNA−binding transcription repressor activity, RNA polymerase II−specific
DNA binding
DNA−binding transcription factor activity, RNA polymerase II−specific
RNA polymerase II distal enhancer sequence−specific DNA binding
RNA polymerase II regulatory region sequence−specific DNA binding
DNA−binding transcription factor activity
sequence−specific DNA binding
transcription factor activity, RNA polymerase II distal enhancer sequence−specific binding
Biological Process
Cellular Component
Molecular Function
0.0
2.5
5.0
7.5
−log10 p−value
