## Supplemental Data 1-8 for "A Spatiotemporal Embryonic Atlas of Bombyx mori Integrating Single-Nucleus RNA Sequencing and Spatial Transcriptomics": Cluster6 GO top100.pdf

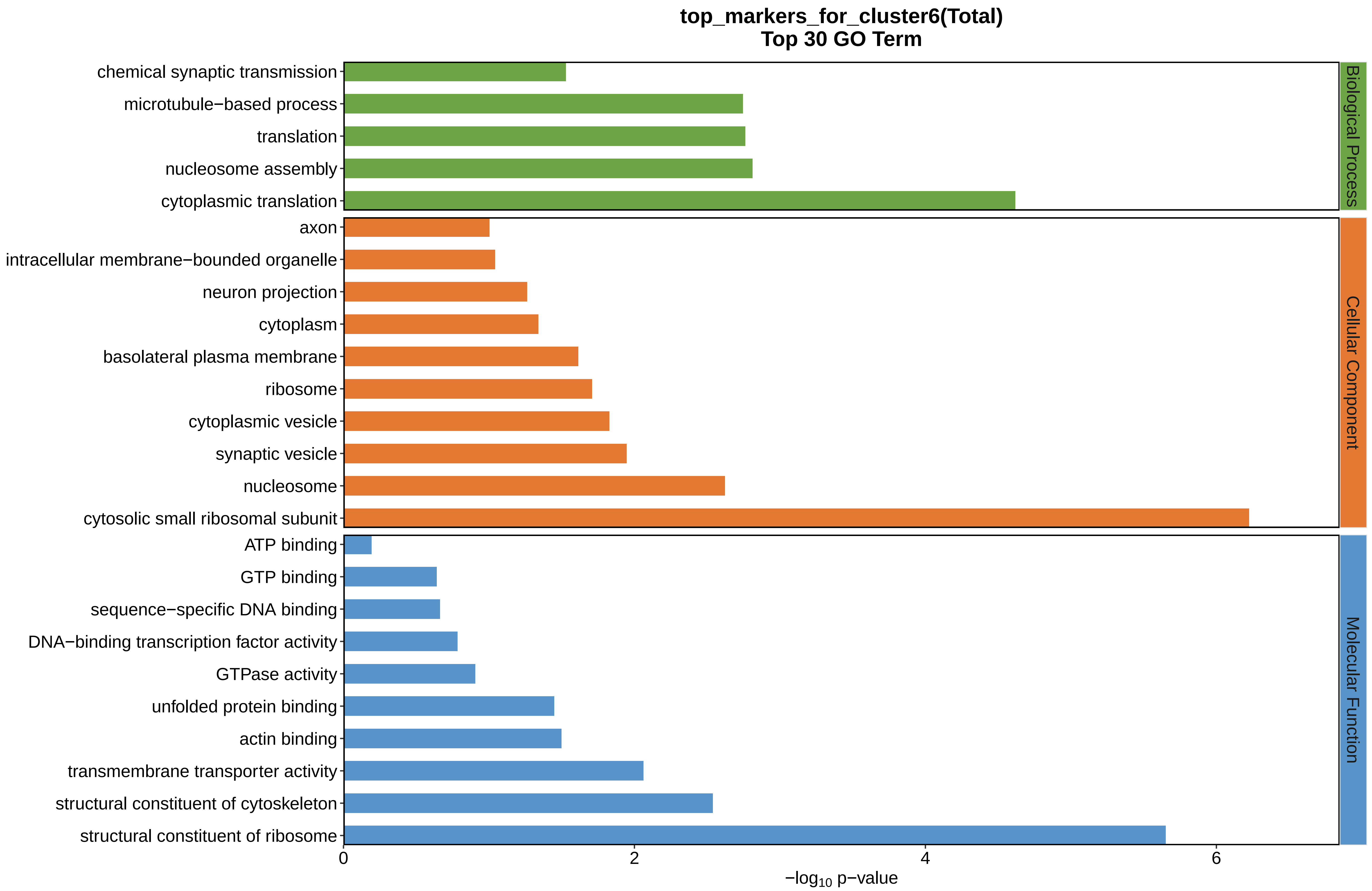
top_markers_for_cluster6(Total)
Top 30 GO Term
chemical synaptic transmission
microtubule−based process
translation
nucleosome assembly
cytoplasmic translation
axon
intracellular membrane−bounded organelle
neuron projection
cytoplasm
basolateral plasma membrane
ribosome
cytoplasmic vesicle
synaptic vesicle
nucleosome
cytosolic small ribosomal subunit
ATP binding
GTP binding
sequence−specific DNA binding
DNA−binding transcription factor activity
GTPase activity
unfolded protein binding
actin binding
transmembrane transporter activity
structural constituent of cytoskeleton
structural constituent of ribosome
Biological Process
Cellular Component
Molecular Function
0
2
4
6
$-\log_{10}$ p−value
