## Supplemental Data 1-8 for "A Spatiotemporal Embryonic Atlas of Bombyx mori Integrating Single-Nucleus RNA Sequencing and Spatial Transcriptomics": Cluster7 GO top100.pdf

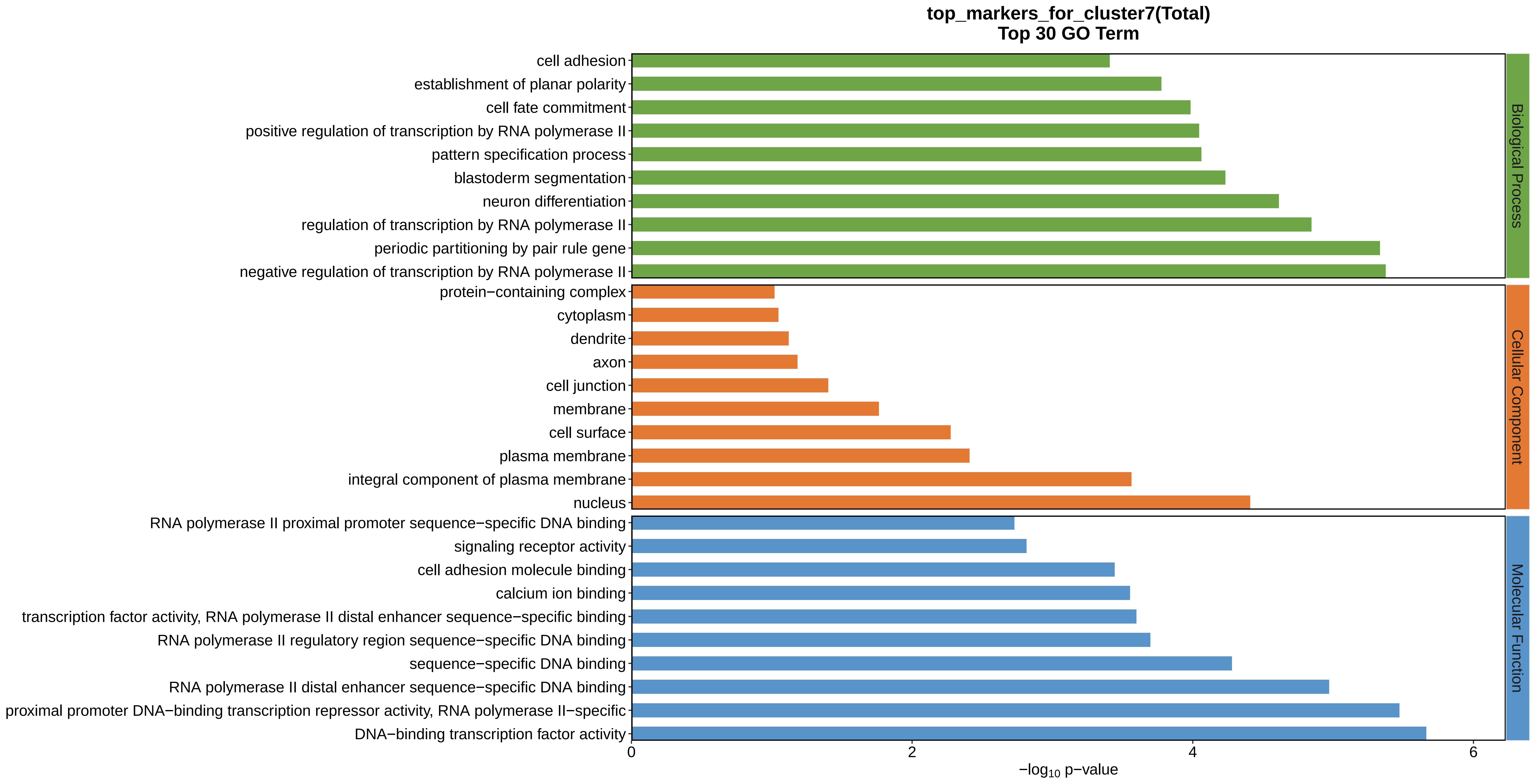
top_markers_for_cluster7(Total)
Top 30 GO Term
cell adhesion
establishment of planar polarity
cell fate commitment
positive regulation of transcription by RNA polymerase II
pattern specification process
blastoderm segmentation
neuron differentiation
regulation of transcription by RNA polymerase II
periodic partitioning by pair rule gene
negative regulation of transcription by RNA polymerase II
protein−containing complex
cytoplasm
dendrite
axon
cell junction
membrane
cell surface
plasma membrane
integral component of plasma membrane
nucleus
RNA polymerase II proximal promoter sequence−specific DNA binding
signaling receptor activity
cell adhesion molecule binding
calcium ion binding
transcription factor activity, RNA polymerase II distal enhancer sequence−specific binding
RNA polymerase II regulatory region sequence−specific DNA binding
sequence−specific DNA binding
RNA polymerase II distal enhancer sequence−specific DNA binding
proximal promoter DNA−binding transcription repressor activity, RNA polymerase II−specific
DNA−binding transcription factor activity
Biological Process
Cellular Component
Molecular Function
0
2
4
6
−log10 p−value
