## Supplemental Data 1-8 for "A Spatiotemporal Embryonic Atlas of Bombyx mori Integrating Single-Nucleus RNA Sequencing and Spatial Transcriptomics": Cluster8 GO top100.pdf

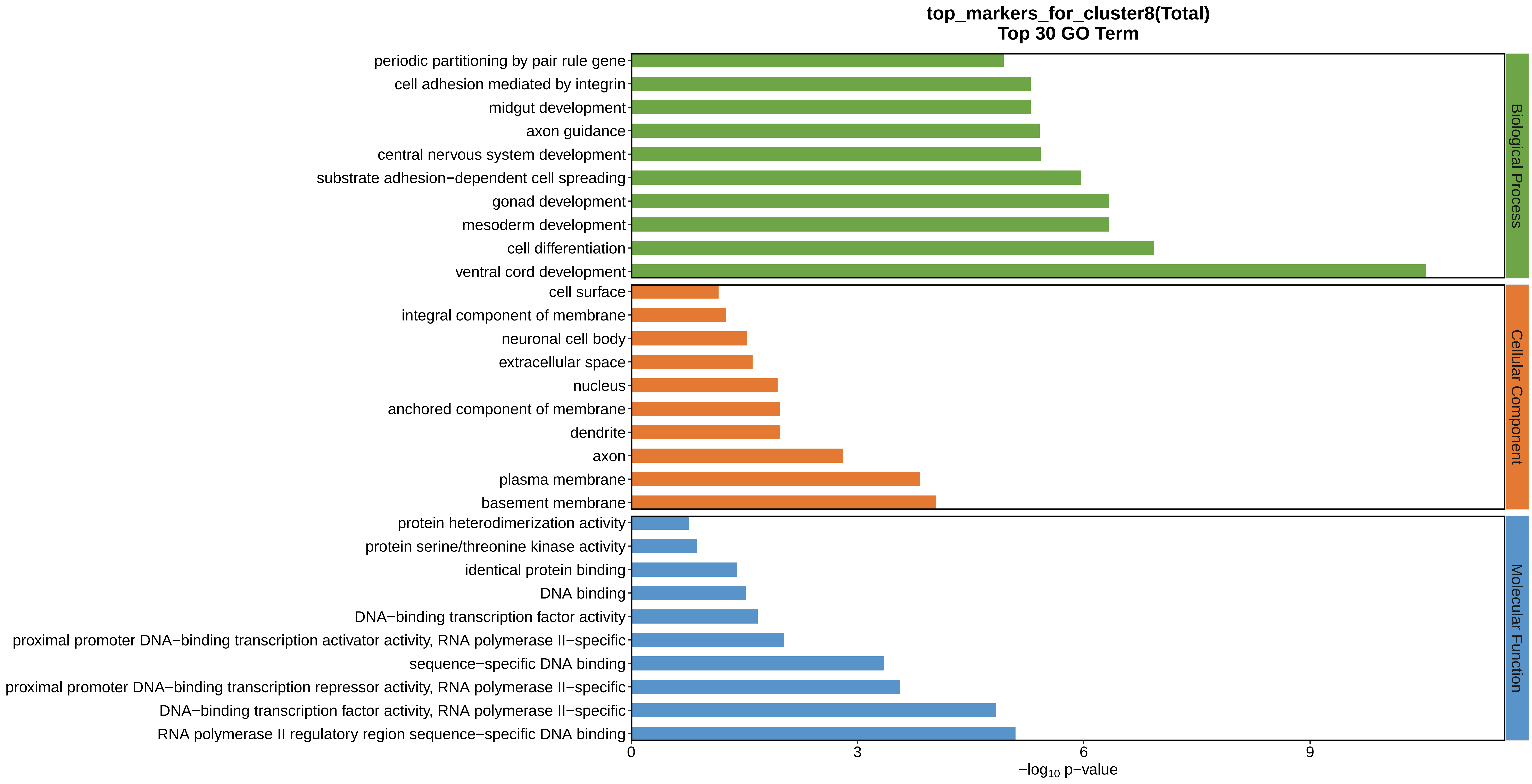

top_markers_for_cluster8(Total)
Top 30 GO Term
periodic partitioning by pair rule gene
cell adhesion mediated by integrin
midgut development
axon guidance
central nervous system development
substrate adhesion−dependent cell spreading
gonad development
mesoderm development
cell differentiation
ventral cord development
cell surface
integral component of membrane
neuronal cell body
extracellular space
nucleus
anchored component of membrane
dendrite
axon
plasma membrane
basement membrane
protein heterodimerization activity
protein serine/threonine kinase activity
identical protein binding
DNA binding
DNA−binding transcription factor activity
proximal promoter DNA−binding transcription activator activity, RNA polymerase II−specific
sequence−specific DNA binding
proximal promoter DNA−binding transcription repressor activity, RNA polymerase II−specific
DNA−binding transcription factor activity, RNA polymerase II−specific
RNA polymerase II regulatory region sequence−specific DNA binding
Biological Process
Cellular Component
Molecular Function
0
3
6
9
−log10 p−value
