## Supplemental Data 1-8 for "A Spatiotemporal Embryonic Atlas of Bombyx mori Integrating Single-Nucleus RNA Sequencing and Spatial Transcriptomics": Cluster9 GO top100.pdf

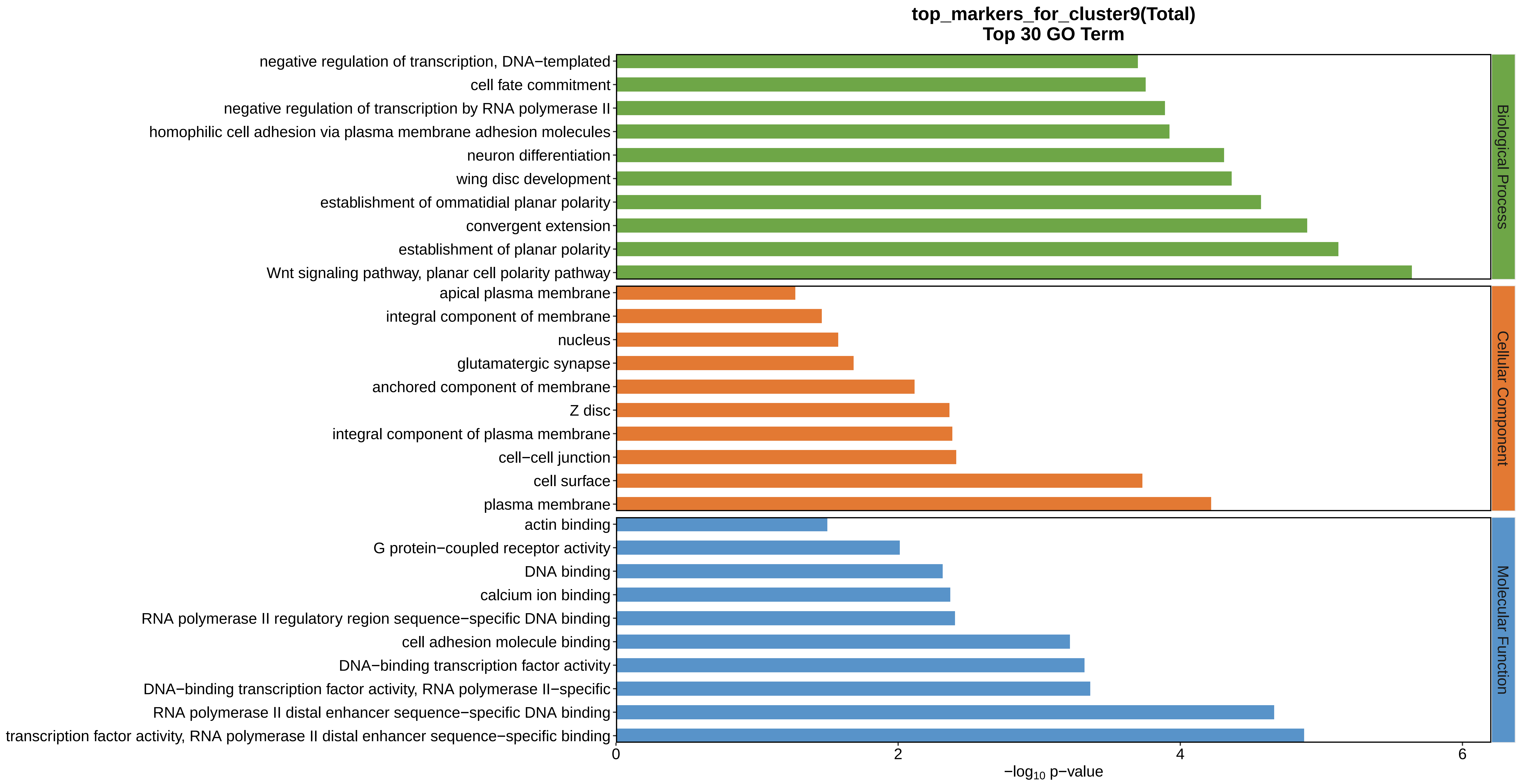
top_markers_for_cluster9(Total)
Top 30 GO Term
negative regulation of transcription, DNA−templated
cell fate commitment
negative regulation of transcription by RNA polymerase II
homophilic cell adhesion via plasma membrane adhesion molecules
neuron differentiation
wing disc development
establishment of ommatidial planar polarity
convergent extension
establishment of planar polarity
Wnt signaling pathway, planar cell polarity pathway
apical plasma membrane
integral component of membrane
nucleus
glutamatergic synapse
anchored component of membrane
Z disc
integral component of plasma membrane
cell−cell junction
cell surface
plasma membrane
actin binding
G protein−coupled receptor activity
DNA binding
calcium ion binding
RNA polymerase II regulatory region sequence−specific DNA binding
cell adhesion molecule binding
DNA−binding transcription factor activity
DNA−binding transcription factor activity, RNA polymerase II−specific
RNA polymerase II distal enhancer sequence−specific DNA binding
transcription factor activity, RNA polymerase II distal enhancer sequence−specific binding
Biological Process
Cellular Component
Molecular Function
0
2
4
6
−log10 p−value
