## Supplemental Data 1-8 for "A Spatiotemporal Embryonic Atlas of Bombyx mori Integrating Single-Nucleus RNA Sequencing and Spatial Transcriptomics": Cluster10 GO top100.pdf

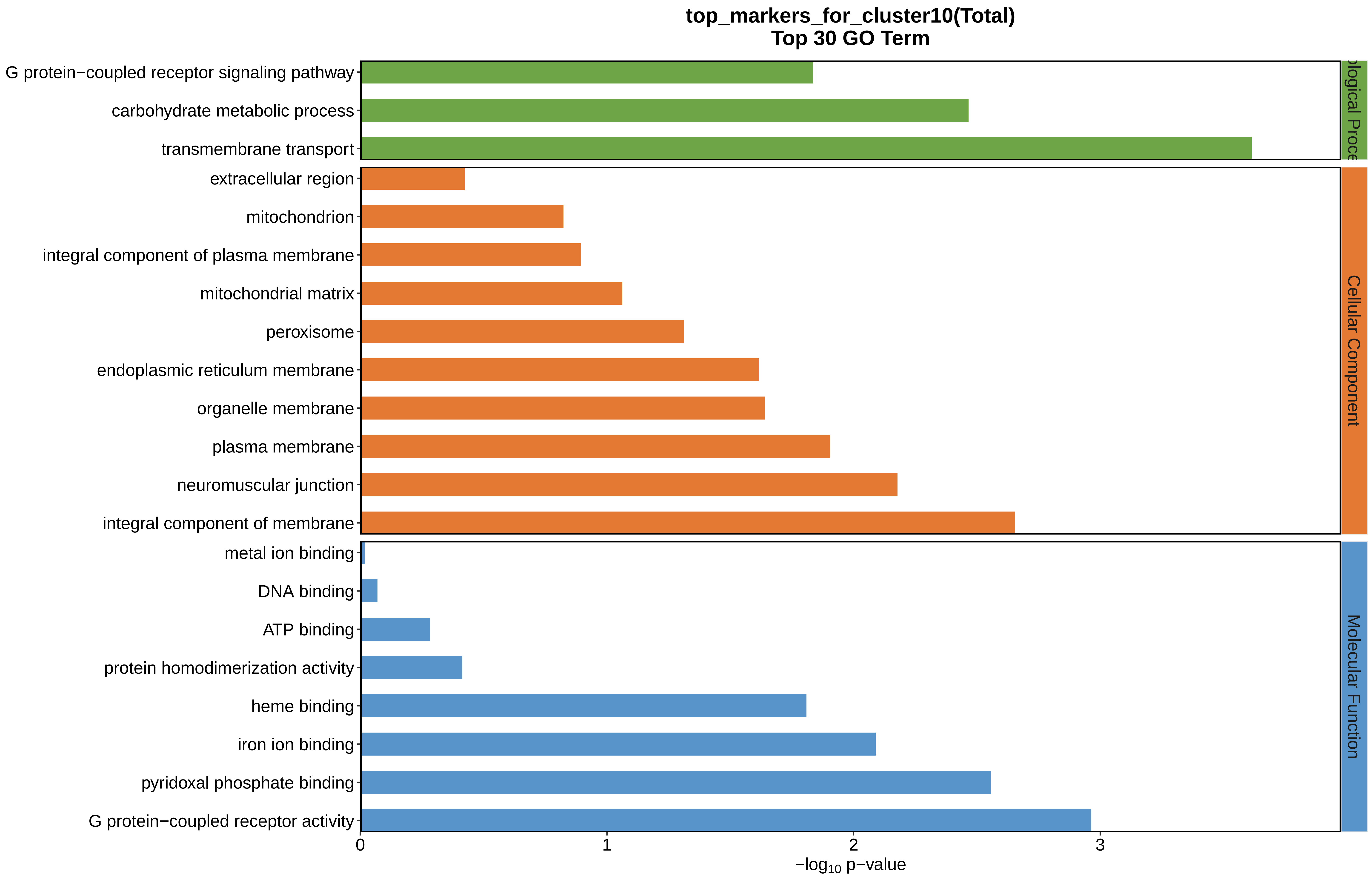

top_markers_for_cluster10(Total)
Top 30 GO Term
G protein−coupled receptor signaling pathway
carbohydrate metabolic process
transmembrane transport
extracellular region
mitochondrion
integral component of plasma membrane
mitochondrial matrix
peroxisome
endoplasmic reticulum membrane
organelle membrane
plasma membrane
neuromuscular junction
integral component of membrane
metal ion binding
DNA binding
ATP binding
protein homodimerization activity
heme binding
iron ion binding
pyridoxal phosphate binding
G protein−coupled receptor activity
ological Proce
Cellular Component
Molecular Function
0
1
2
3
$-\log_{10}$ p−value
