## Supplemental Data 1-8 for "A Spatiotemporal Embryonic Atlas of Bombyx mori Integrating Single-Nucleus RNA Sequencing and Spatial Transcriptomics": Cluster11 GO top100.pdf

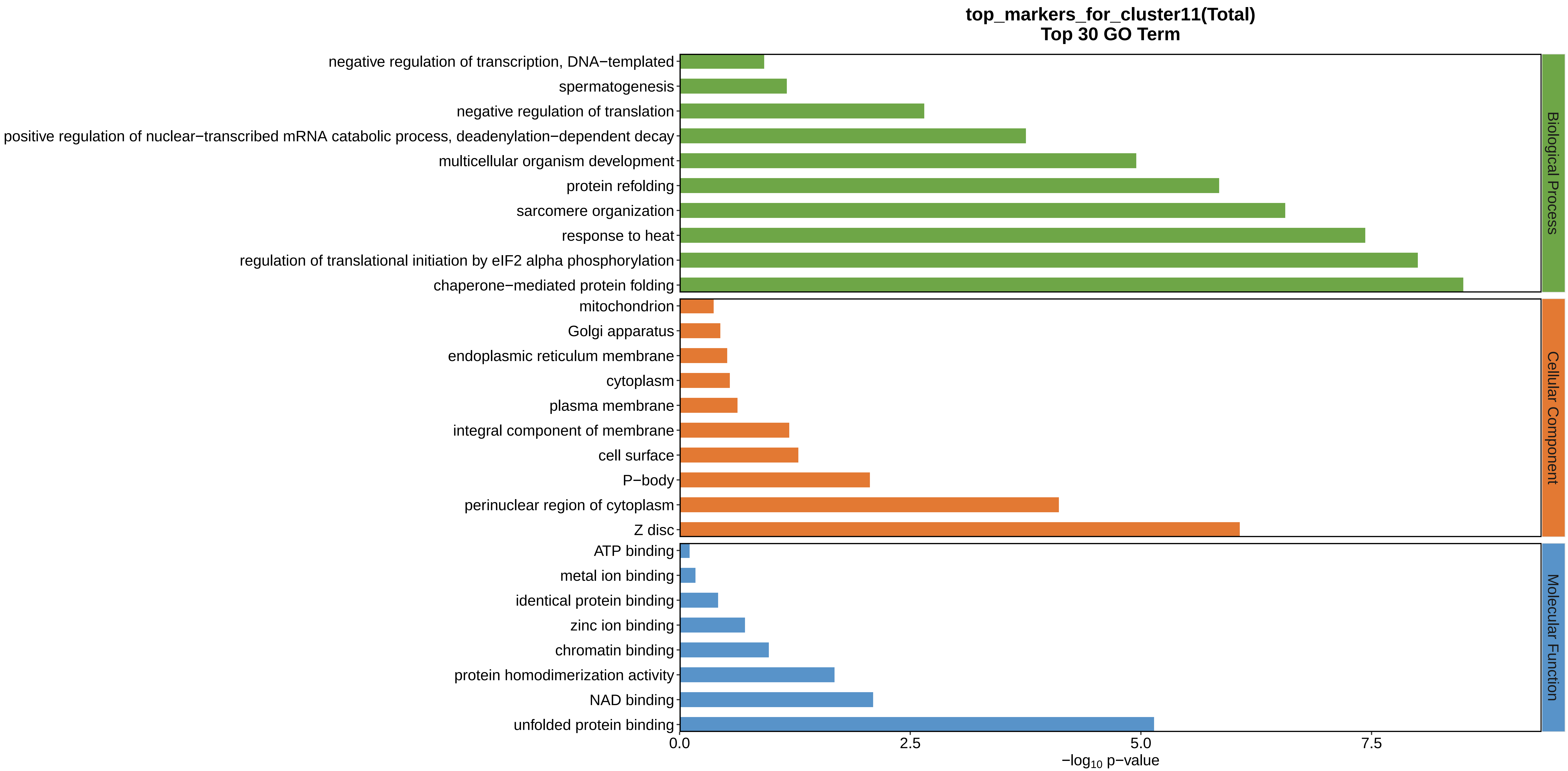
top_markers_for_cluster11(Total)
Top 30 GO Term
negative regulation of transcription, DNA−templated
spermatogenesis
negative regulation of translation
positive regulation of nuclear−transcribed mRNA catabolic process, deadenylation−dependent decay
multicellular organism development
protein refolding
sarcomere organization
response to heat
regulation of translational initiation by eIF2 alpha phosphorylation
chaperone−mediated protein folding
mitochondrion
Golgi apparatus
endoplasmic reticulum membrane
cytoplasm
plasma membrane
integral component of membrane
cell surface
P−body
perinuclear region of cytoplasm
Z disc
ATP binding
metal ion binding
identical protein binding
zinc ion binding
chromatin binding
protein homodimerization activity
NAD binding
unfolded protein binding
0.0
2.5
5.0
7.5
−log10 p−value
Biological Process
Cellular Component
Molecular Function
