## Supplemental Data 1-8 for "A Spatiotemporal Embryonic Atlas of Bombyx mori Integrating Single-Nucleus RNA Sequencing and Spatial Transcriptomics": Cluster12 GO top100.pdf

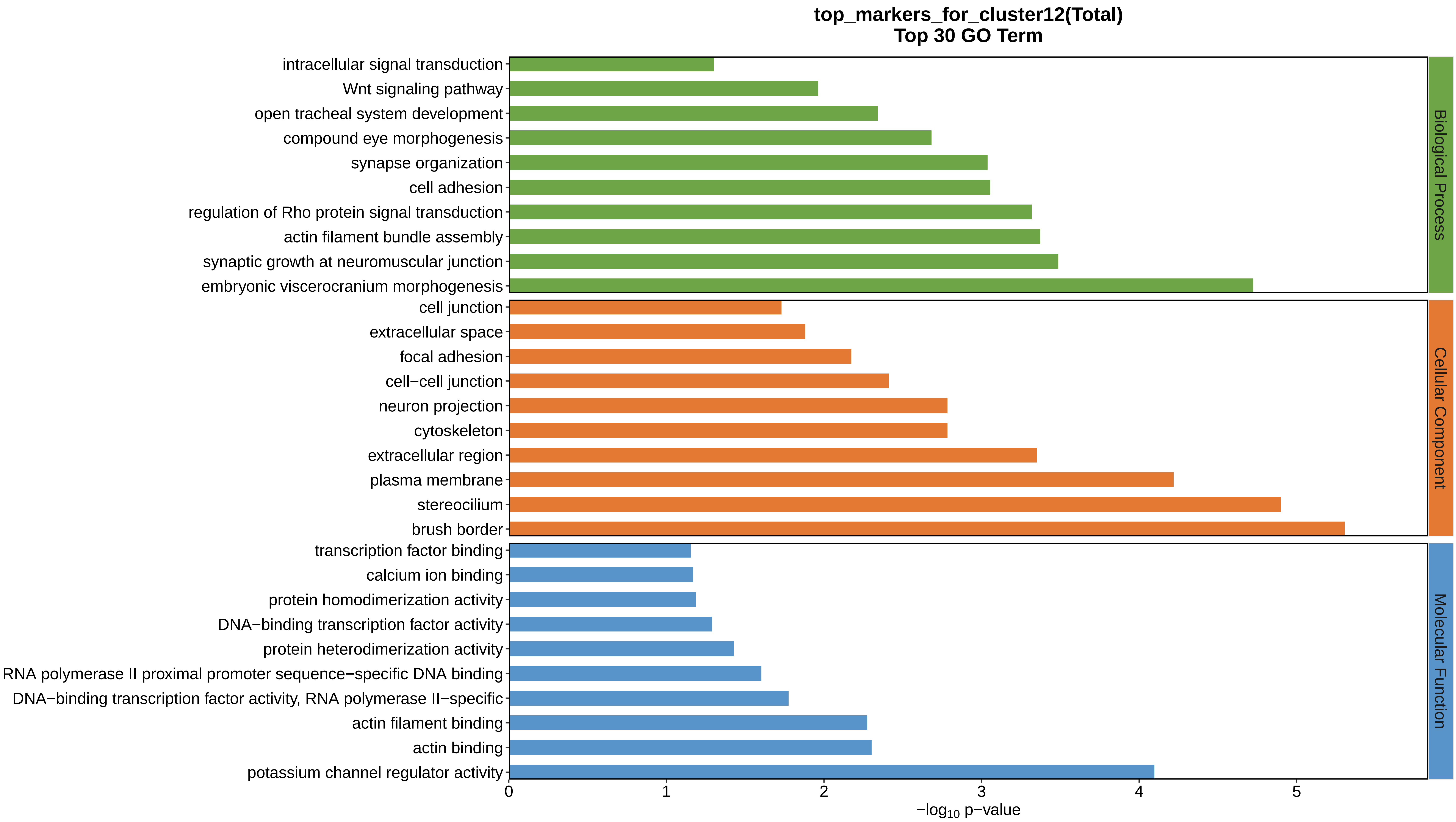
top_markers_for_cluster12(Total)
Top 30 GO Term
intracellular signal transduction
Wnt signaling pathway
open tracheal system development
compound eye morphogenesis
synapse organization
cell adhesion
regulation of Rho protein signal transduction
actin filament bundle assembly
synaptic growth at neuromuscular junction
embryonic viscerocranium morphogenesis
cell junction
extracellular space
focal adhesion
cell−cell junction
neuron projection
cytoskeleton
extracellular region
plasma membrane
stereocilium
brush border
transcription factor binding
calcium ion binding
protein homodimerization activity
DNA−binding transcription factor activity
protein heterodimerization activity
RNA polymerase II proximal promoter sequence−specific DNA binding
DNA−binding transcription factor activity, RNA polymerase II−specific
actin filament binding
actin binding
potassium channel regulator activity
Biological Process
Cellular Component
Molecular Function
0
1
2
3
4
5
−log10 p−value
