## Supplemental Data 1-8 for "A Spatiotemporal Embryonic Atlas of Bombyx mori Integrating Single-Nucleus RNA Sequencing and Spatial Transcriptomics": Cluster1 GO top100.pdf

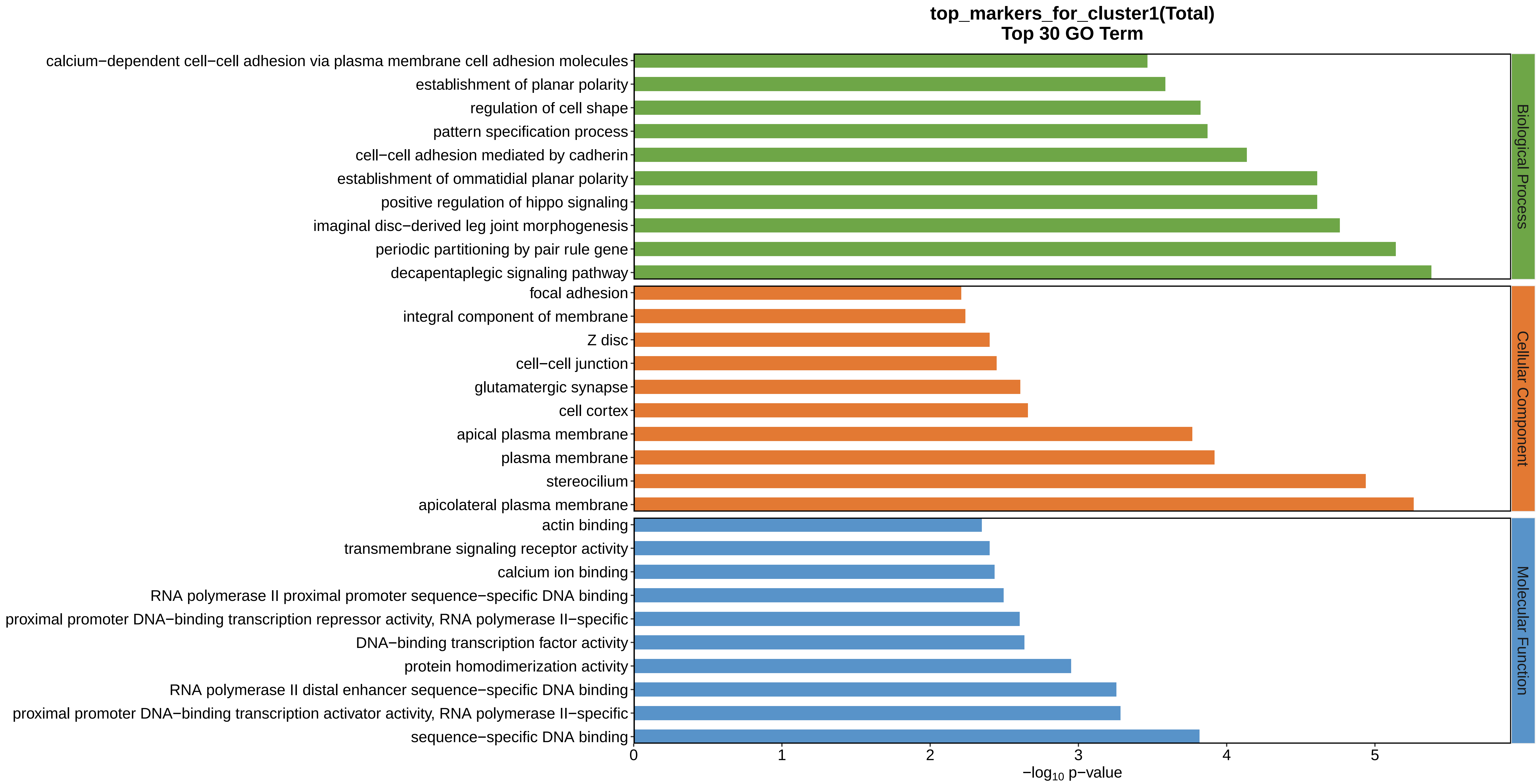
top_markers_for_cluster1(Total)
Top 30 GO Term
calcium−dependent cell−cell adhesion via plasma membrane cell adhesion molecules
establishment of planar polarity
regulation of cell shape
pattern specification process
cell−cell adhesion mediated by cadherin
establishment of ommatidial planar polarity
positive regulation of hippo signaling
imaginal disc−derived leg joint morphogenesis
periodic partitioning by pair rule gene
decapentaplegic signaling pathway
focal adhesion
integral component of membrane
Z disc
cell−cell junction
glutamatergic synapse
cell cortex
apical plasma membrane
plasma membrane
stereocilium
apicolateral plasma membrane
actin binding
transmembrane signaling receptor activity
calcium ion binding
RNA polymerase II proximal promoter sequence−specific DNA binding
proximal promoter DNA−binding transcription repressor activity, RNA polymerase II−specific
DNA−binding transcription factor activity
protein homodimerization activity
RNA polymerase II distal enhancer sequence−specific DNA binding
proximal promoter DNA−binding transcription activator activity, RNA polymerase II−specific
sequence−specific DNA binding
0
1
2
3
4
5
$-\log_{10}$ p−value
Biological Process
Cellular Component
Molecular Function
