## Supplemental Data 1-8 for "A Spatiotemporal Embryonic Atlas of Bombyx mori Integrating Single-Nucleus RNA Sequencing and Spatial Transcriptomics": Cluster2 GO top100.pdf

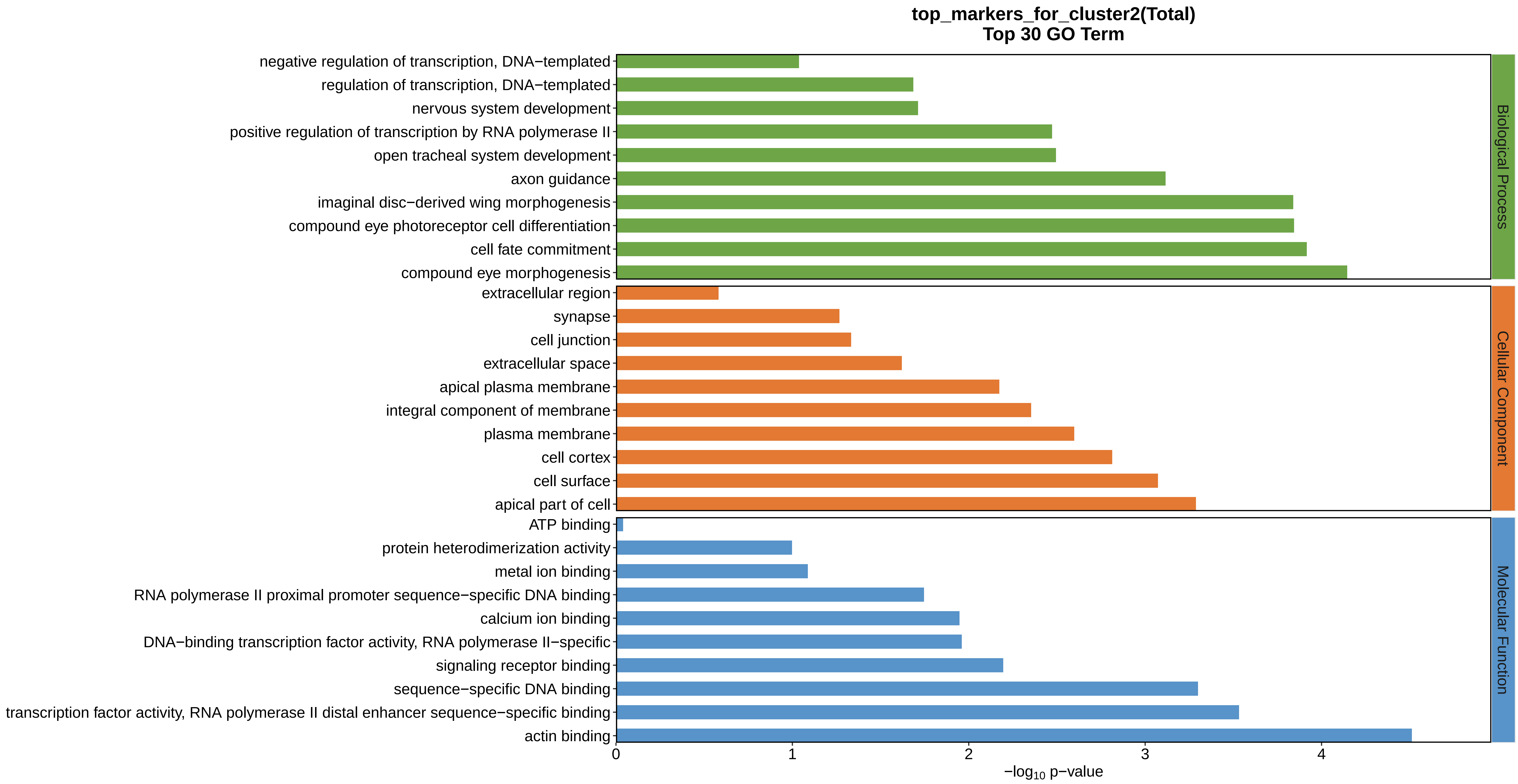
top_markers_for_cluster2(Total)
Top 30 GO Term
negative regulation of transcription, DNA−templated
regulation of transcription, DNA−templated
nervous system development
positive regulation of transcription by RNA polymerase II
open tracheal system development
axon guidance
imaginal disc−derived wing morphogenesis
compound eye photoreceptor cell differentiation
cell fate commitment
compound eye morphogenesis
extracellular region
synapse
cell junction
extracellular space
apical plasma membrane
integral component of membrane
plasma membrane
cell cortex
cell surface
apical part of cell
ATP binding
protein heterodimerization activity
metal ion binding
RNA polymerase II proximal promoter sequence−specific DNA binding
calcium ion binding
DNA−binding transcription factor activity, RNA polymerase II−specific
signaling receptor binding
sequence−specific DNA binding
transcription factor activity, RNA polymerase II distal enhancer sequence−specific binding
actin binding
Biological Process
Cellular Component
Molecular Function
0
1
2
3
4
$-\log_{10}$ p−value
