## Supplemental Data 1-8 for "A Spatiotemporal Embryonic Atlas of Bombyx mori Integrating Single-Nucleus RNA Sequencing and Spatial Transcriptomics": Cluster3 GO top100.pdf

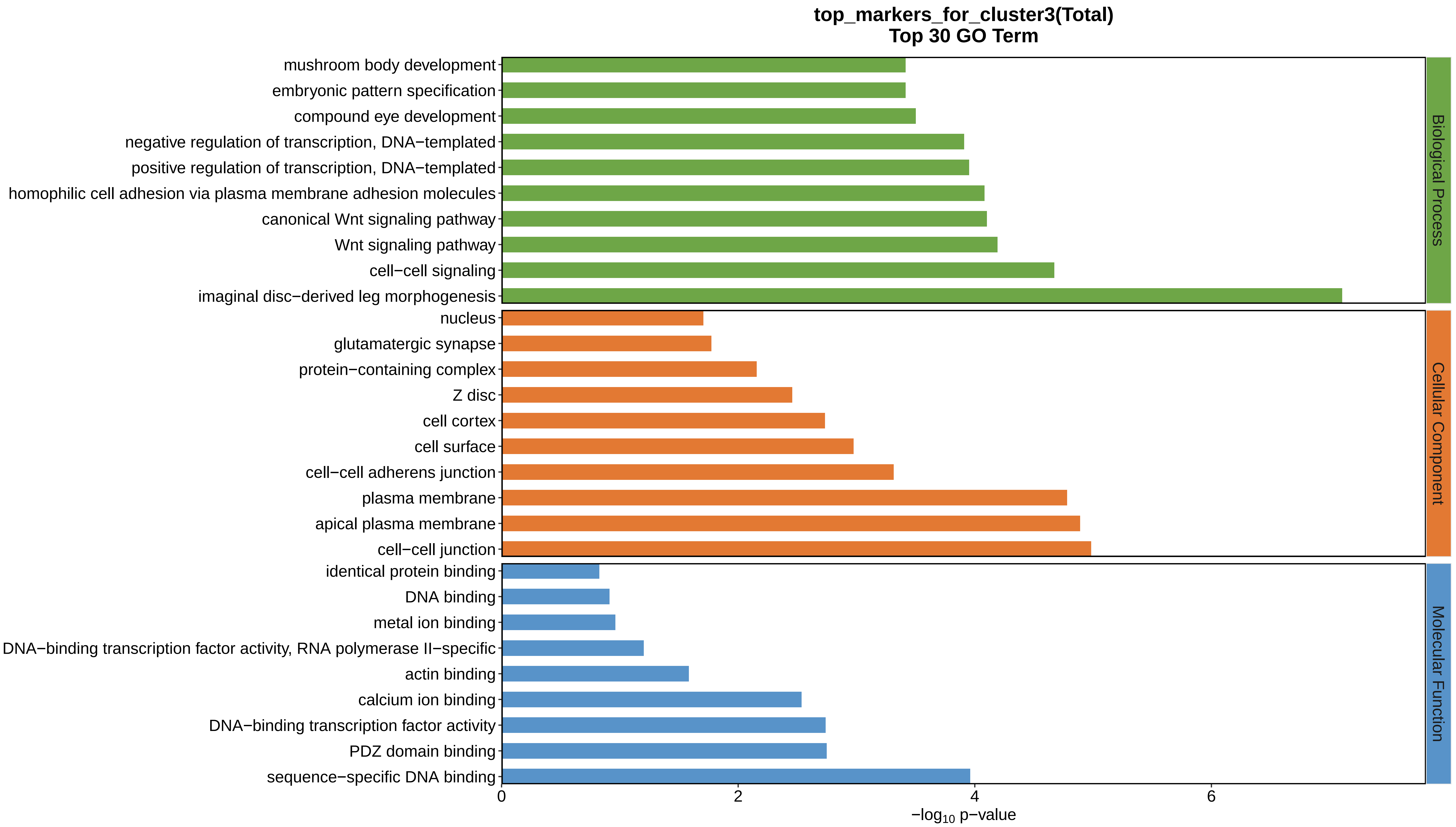
top_markers_for_cluster3(Total)
Top 30 GO Term
mushroom body development
embryonic pattern specification
compound eye development
negative regulation of transcription, DNA−templated
positive regulation of transcription, DNA−templated
homophilic cell adhesion via plasma membrane adhesion molecules
canonical Wnt signaling pathway
Wnt signaling pathway
cell−cell signaling
imaginal disc−derived leg morphogenesis
nucleus
glutamatergic synapse
protein−containing complex
Z disc
cell cortex
cell surface
cell−cell adherens junction
plasma membrane
apical plasma membrane
cell−cell junction
identical protein binding
DNA binding
metal ion binding
DNA−binding transcription factor activity, RNA polymerase II−specific
actin binding
calcium ion binding
DNA−binding transcription factor activity
PDZ domain binding
sequence−specific DNA binding
Biological Process
Cellular Component
Molecular Function
0
2
4
6
−log10 p−value
