## Supplemental Data 1-8 for "A Spatiotemporal Embryonic Atlas of Bombyx mori Integrating Single-Nucleus RNA Sequencing and Spatial Transcriptomics": Cluster4 GO top100.pdf

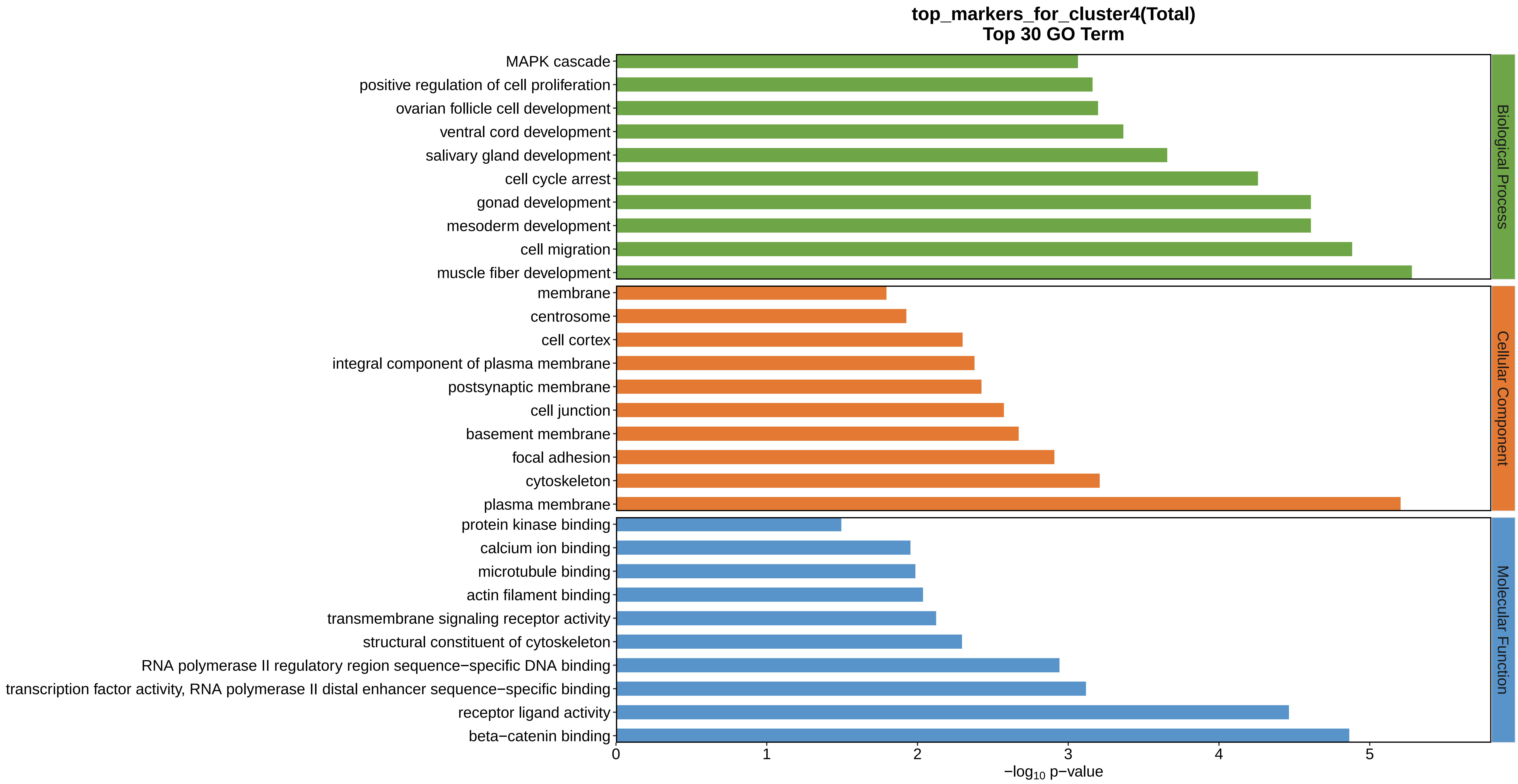
top_markers_for_cluster4(Total)
Top 30 GO Term
MAPK cascade
positive regulation of cell proliferation
ovarian follicle cell development
ventral cord development
salivary gland development
cell cycle arrest
gonad development
mesoderm development
cell migration
muscle fiber development
membrane
centrosome
cell cortex
integral component of plasma membrane
postsynaptic membrane
cell junction
basement membrane
focal adhesion
cytoskeleton
plasma membrane
protein kinase binding
calcium ion binding
microtubule binding
actin filament binding
transmembrane signaling receptor activity
structural constituent of cytoskeleton
RNA polymerase II regulatory region sequence−specific DNA binding
transcription factor activity, RNA polymerase II distal enhancer sequence−specific binding
receptor ligand activity
beta−catenin binding
Biological Process
Cellular Component
Molecular Function
0
1
2
3
4
5
$-\log_{10}$ p−value
