## Supplemental Data 1-8 for "A Spatiotemporal Embryonic Atlas of Bombyx mori Integrating Single-Nucleus RNA Sequencing and Spatial Transcriptomics": Cluster5 GO top100.pdf

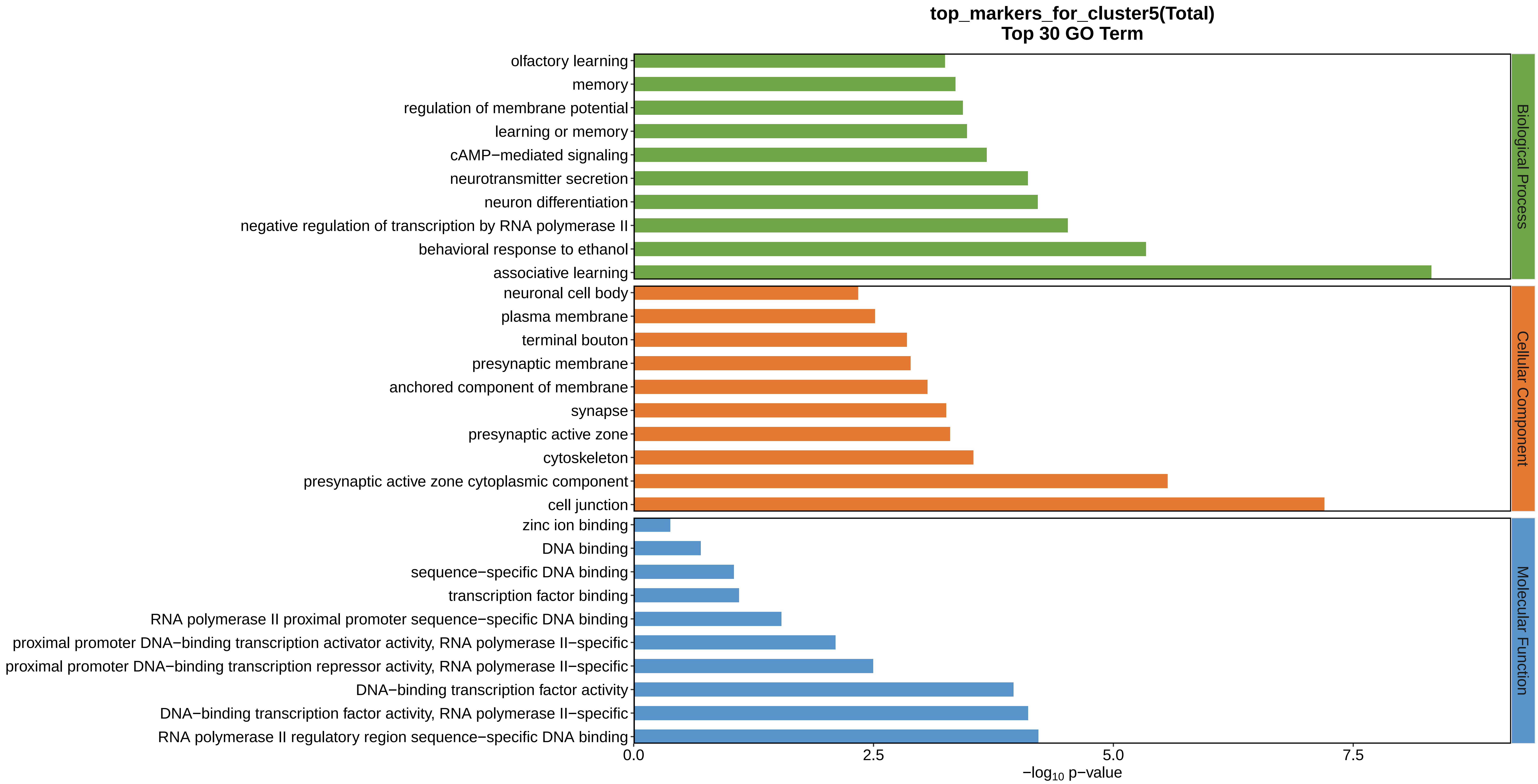
top_markers_for_cluster5(Total)
Top 30 GO Term
olfactory learning
memory
regulation of membrane potential
learning or memory
cAMP−mediated signaling
neurotransmitter secretion
neuron differentiation
negative regulation of transcription by RNA polymerase II
behavioral response to ethanol
associative learning
neuronal cell body
plasma membrane
terminal bouton
presynaptic membrane
anchored component of membrane
synapse
presynaptic active zone
cytoskeleton
presynaptic active zone cytoplasmic component
cell junction
zinc ion binding
DNA binding
sequence−specific DNA binding
transcription factor binding
RNA polymerase II proximal promoter sequence−specific DNA binding
proximal promoter DNA−binding transcription activator activity, RNA polymerase II−specific
proximal promoter DNA−binding transcription repressor activity, RNA polymerase II−specific
DNA−binding transcription factor activity
DNA−binding transcription factor activity, RNA polymerase II−specific
RNA polymerase II regulatory region sequence−specific DNA binding
Biological Process
Cellular Component
Molecular Function
0.0
2.5
5.0
7.5
−log10 p−value
