## Supplemental Data 1-8 for "A Spatiotemporal Embryonic Atlas of Bombyx mori Integrating Single-Nucleus RNA Sequencing and Spatial Transcriptomics": Cluster6 GO top100.pdf

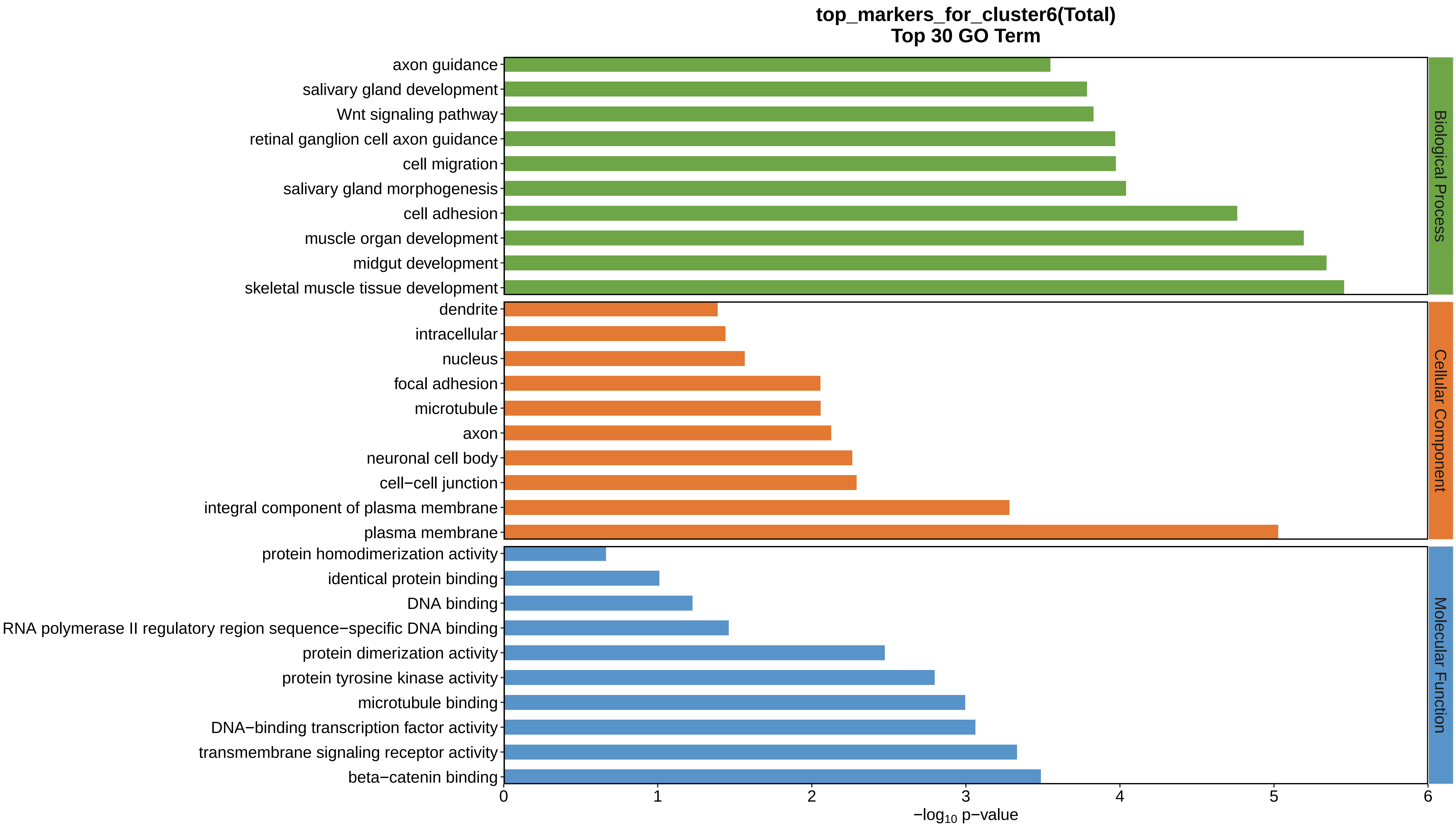

top_markers_for_cluster6(Total)
Top 30 GO Term
axon guidance
salivary gland development
Wnt signaling pathway
retinal ganglion cell axon guidance
cell migration
salivary gland morphogenesis
cell adhesion
muscle organ development
midgut development
skeletal muscle tissue development
dendrite
intracellular
nucleus
focal adhesion
microtubule
axon
neuronal cell body
cell−cell junction
integral component of plasma membrane
plasma membrane
protein homodimerization activity
identical protein binding
DNA binding
RNA polymerase II regulatory region sequence−specific DNA binding
protein dimerization activity
protein tyrosine kinase activity
microtubule binding
DNA−binding transcription factor activity
transmembrane signaling receptor activity
beta−catenin binding
0
1
2
3
4
5
6
$-\log_{10}$ p−value
Biological Process
Cellular Component
Molecular Function
