## Supplemental Data 1-8 for "A Spatiotemporal Embryonic Atlas of Bombyx mori Integrating Single-Nucleus RNA Sequencing and Spatial Transcriptomics": Cluster7 GO top100.pdf

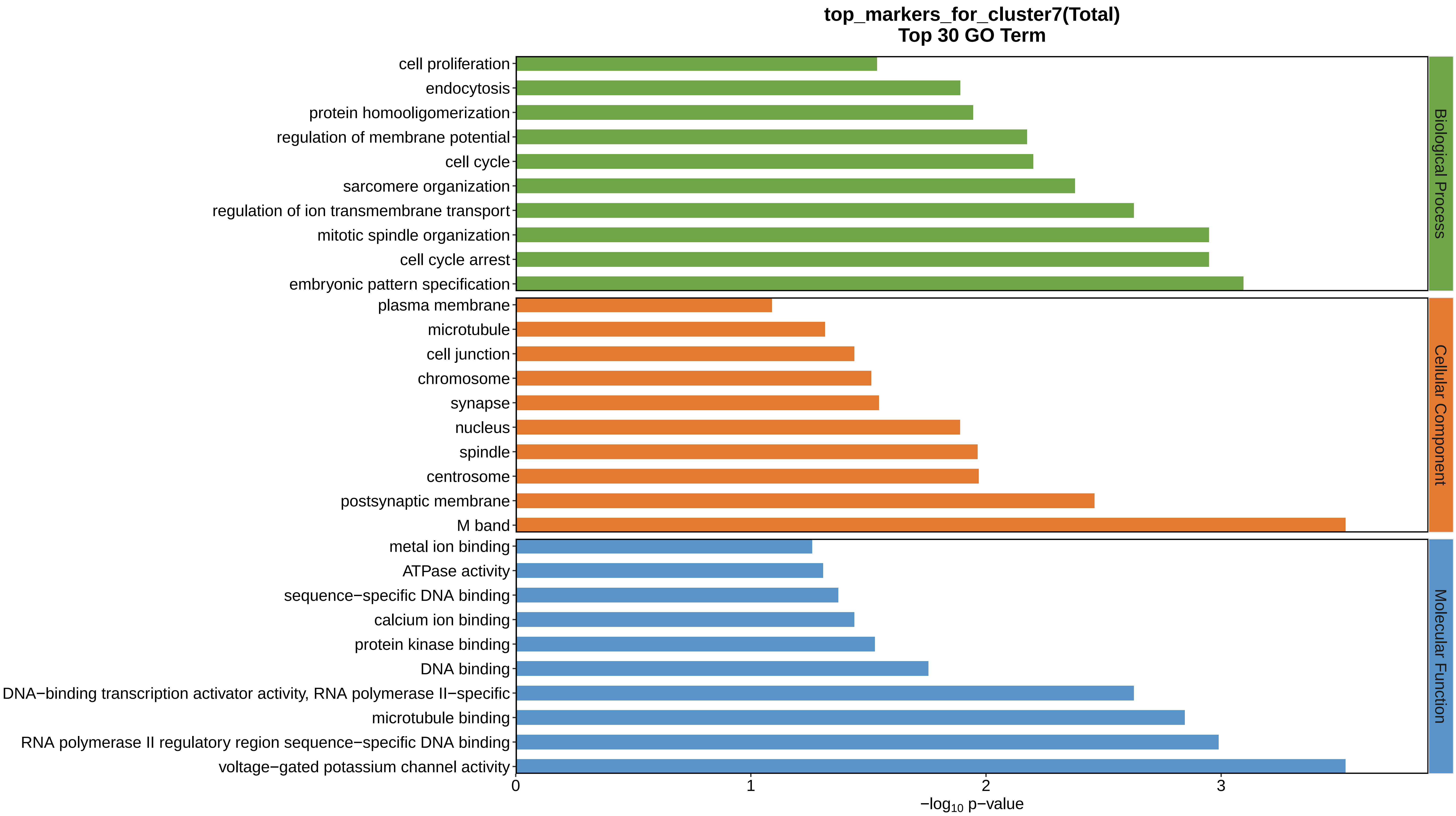
top_markers_for_cluster7(Total)
Top 30 GO Term
cell proliferation
endocytosis
protein homooligomerization
regulation of membrane potential
cell cycle
sarcomere organization
regulation of ion transmembrane transport
mitotic spindle organization
cell cycle arrest
embryonic pattern specification
plasma membrane
microtubule
cell junction
chromosome
synapse
nucleus
spindle
centrosome
postsynaptic membrane
M band
metal ion binding
ATPase activity
sequence−specific DNA binding
calcium ion binding
protein kinase binding
DNA binding
DNA−binding transcription activator activity, RNA polymerase II−specific
microtubule binding
RNA polymerase II regulatory region sequence−specific DNA binding
voltage−gated potassium channel activity
0
1
2
3
$-\log_{10}$ p−value
Biological Process
Cellular Component
Molecular Function
