## Supplemental Data 1-8 for "A Spatiotemporal Embryonic Atlas of Bombyx mori Integrating Single-Nucleus RNA Sequencing and Spatial Transcriptomics": Cluster8 GO top100.pdf

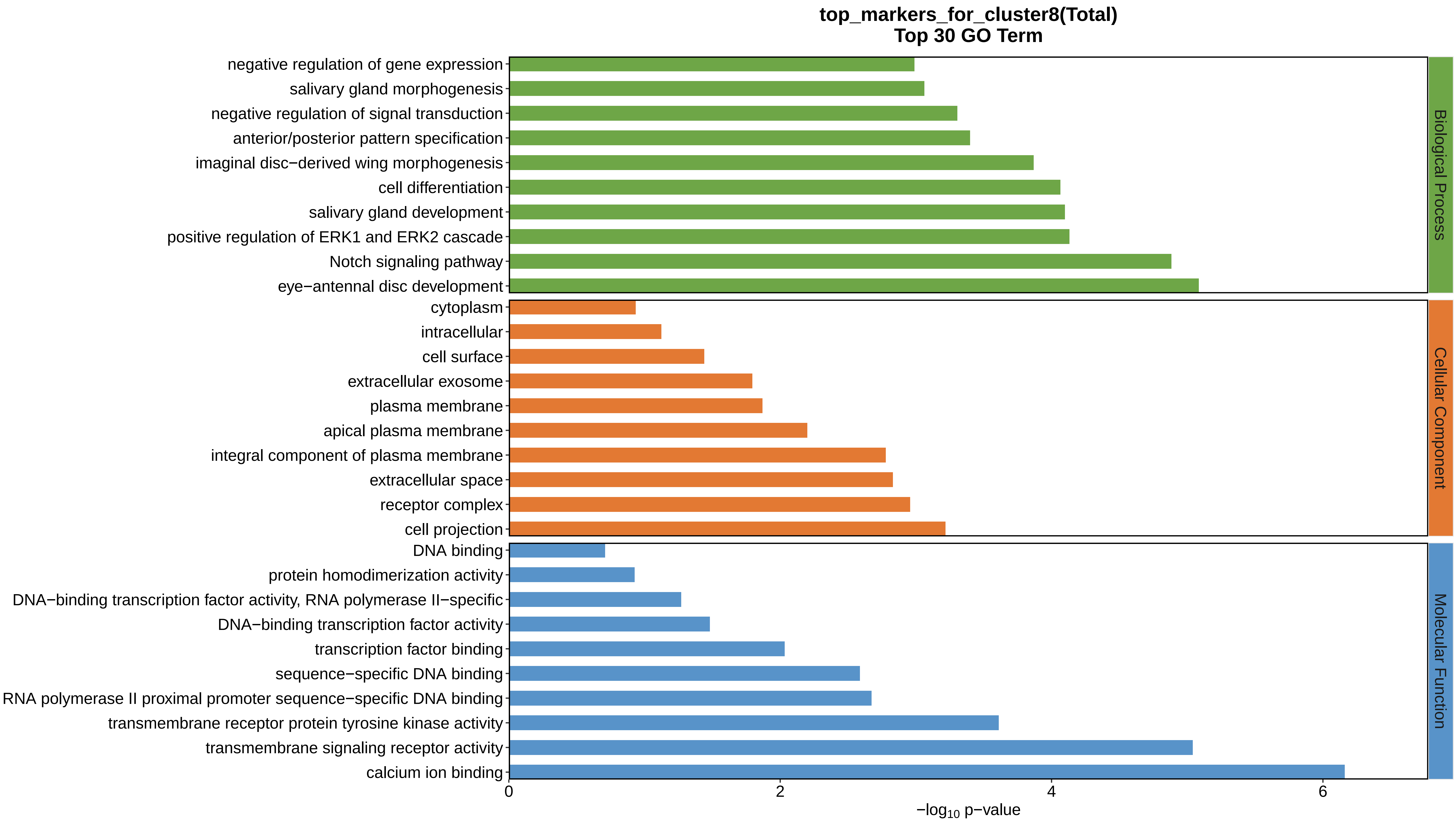

top_markers_for_cluster8(Total)
Top 30 GO Term
negative regulation of gene expression
salivary gland morphogenesis
negative regulation of signal transduction
anterior/posterior pattern specification
imaginal disc−derived wing morphogenesis
cell differentiation
salivary gland development
positive regulation of ERK1 and ERK2 cascade
Notch signaling pathway
eye−antennal disc development
cytoplasm
intracellular
cell surface
extracellular exosome
plasma membrane
apical plasma membrane
integral component of plasma membrane
extracellular space
receptor complex
cell projection
DNA binding
protein homodimerization activity
DNA−binding transcription factor activity, RNA polymerase II−specific
DNA−binding transcription factor activity
transcription factor binding
sequence−specific DNA binding
RNA polymerase II proximal promoter sequence−specific DNA binding
transmembrane receptor protein tyrosine kinase activity
transmembrane signaling receptor activity
calcium ion binding
Biological Process
Cellular Component
Molecular Function
0
2
4
6
$-\log_{10}$ p−value
