## Supplemental Data 1-8 for "A Spatiotemporal Embryonic Atlas of Bombyx mori Integrating Single-Nucleus RNA Sequencing and Spatial Transcriptomics": Cluster9 GO top100.pdf

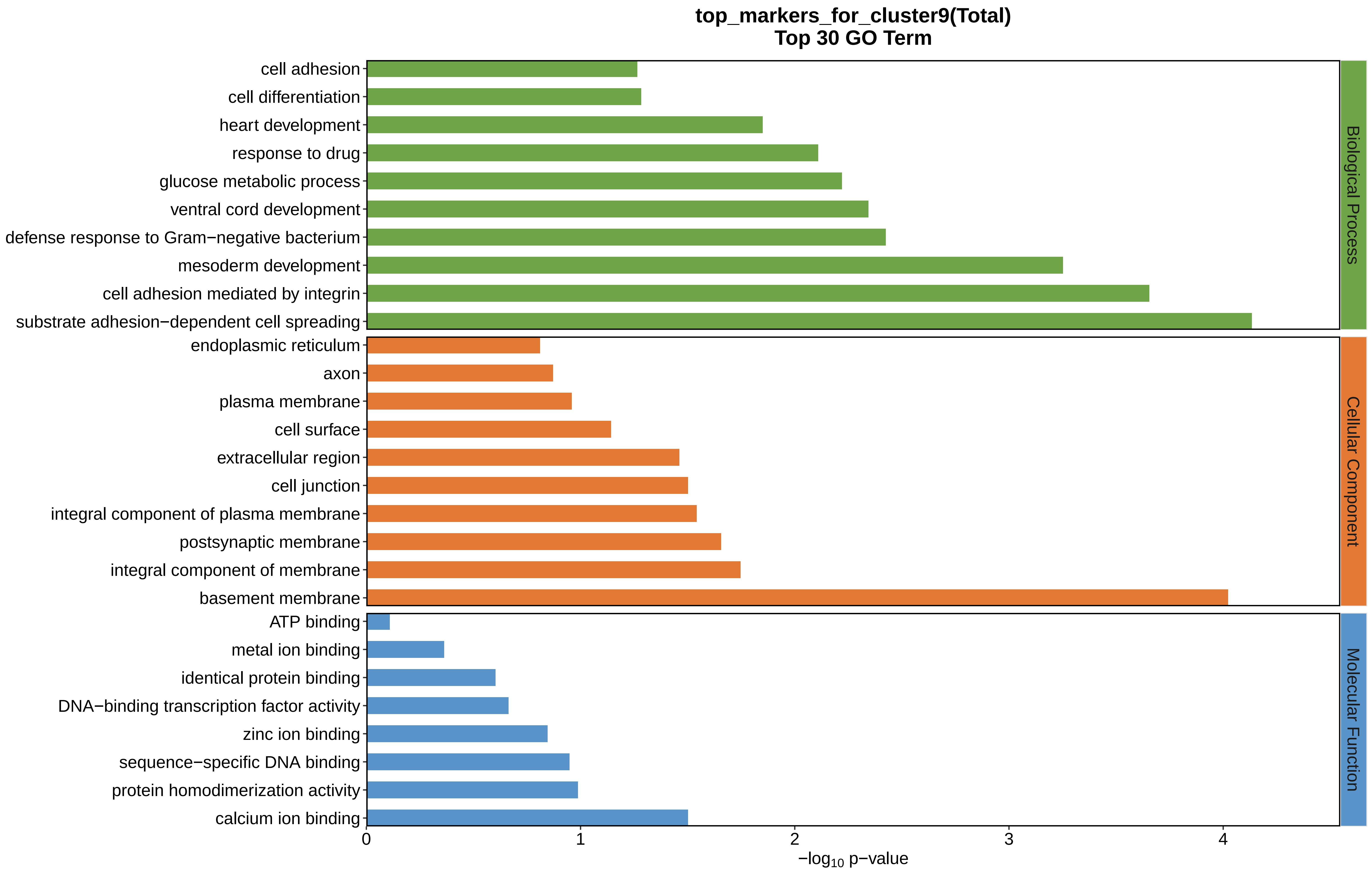
top_markers_for_cluster9(Total)
Top 30 GO Term
cell adhesion
cell differentiation
heart development
response to drug
glucose metabolic process
ventral cord development
defense response to Gram−negative bacterium
mesoderm development
cell adhesion mediated by integrin
substrate adhesion−dependent cell spreading
endoplasmic reticulum
axon
plasma membrane
cell surface
extracellular region
cell junction
integral component of plasma membrane
postsynaptic membrane
integral component of membrane
basement membrane
ATP binding
metal ion binding
identical protein binding
DNA−binding transcription factor activity
zinc ion binding
sequence−specific DNA binding
protein homodimerization activity
calcium ion binding
Biological Process
Cellular Component
Molecular Function
0
1
2
3
4
−log10 p−value
