## Supplemental Data 1-8 for "A Spatiotemporal Embryonic Atlas of Bombyx mori Integrating Single-Nucleus RNA Sequencing and Spatial Transcriptomics": Cluster10 GO top100.pdf

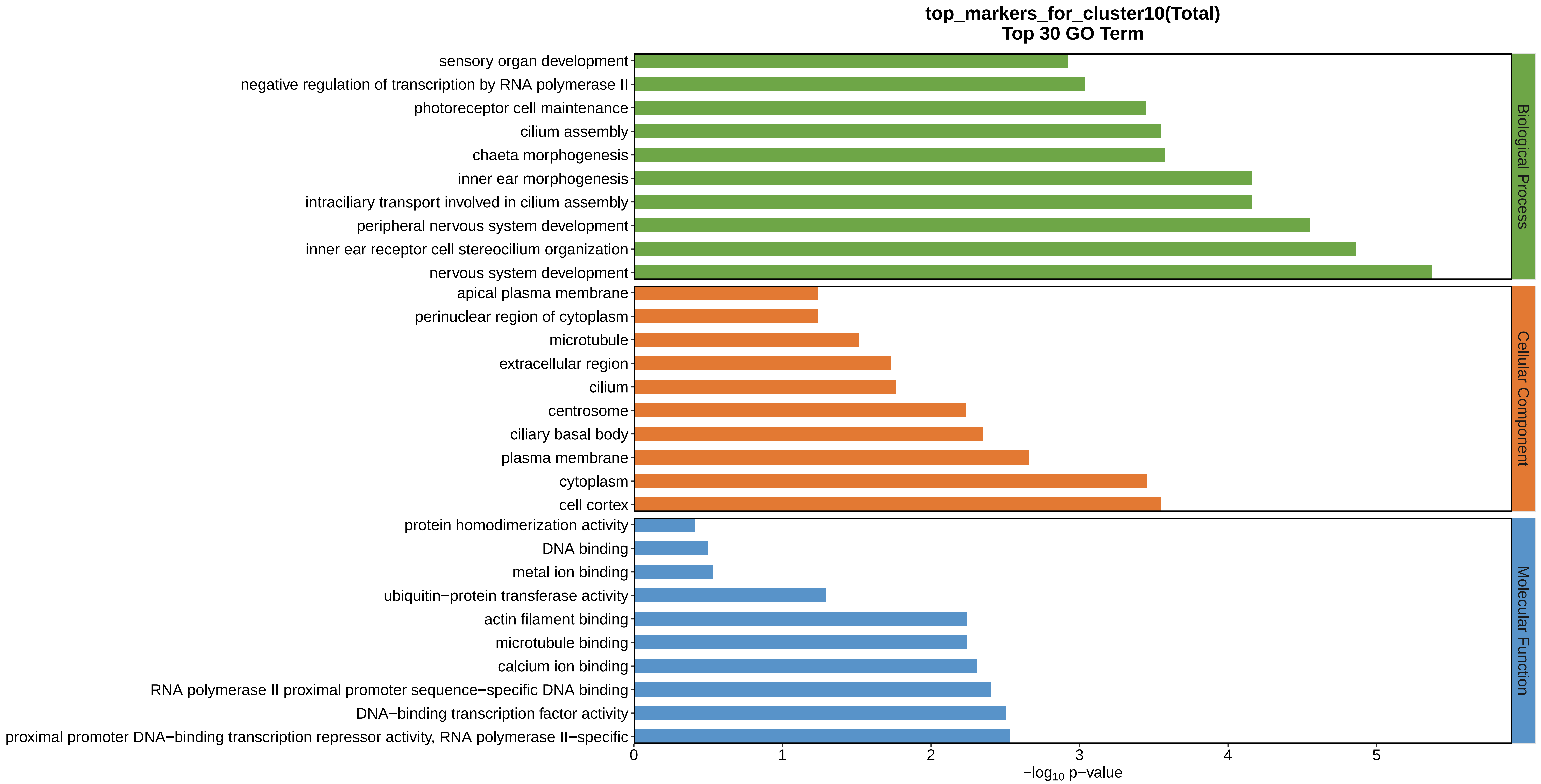
top_markers_for_cluster10(Total)
Top 30 GO Term
sensory organ development
negative regulation of transcription by RNA polymerase II
photoreceptor cell maintenance
cilium assembly
chaeta morphogenesis
inner ear morphogenesis
intraciliary transport involved in cilium assembly
peripheral nervous system development
inner ear receptor cell stereocilium organization
nervous system development
apical plasma membrane
perinuclear region of cytoplasm
microtubule
extracellular region
cilium
centrosome
ciliary basal body
plasma membrane
cytoplasm
cell cortex
protein homodimerization activity
DNA binding
metal ion binding
ubiquitin−protein transferase activity
actin filament binding
microtubule binding
calcium ion binding
RNA polymerase II proximal promoter sequence−specific DNA binding
DNA−binding transcription factor activity
proximal promoter DNA−binding transcription repressor activity, RNA polymerase II−specific
Biological Process
Cellular Component
Molecular Function
0
1
2
3
4
5
$-\log_{10}$ p−value
