## Supplemental Data 1-8 for "A Spatiotemporal Embryonic Atlas of Bombyx mori Integrating Single-Nucleus RNA Sequencing and Spatial Transcriptomics": Cluster11 GO top100.pdf

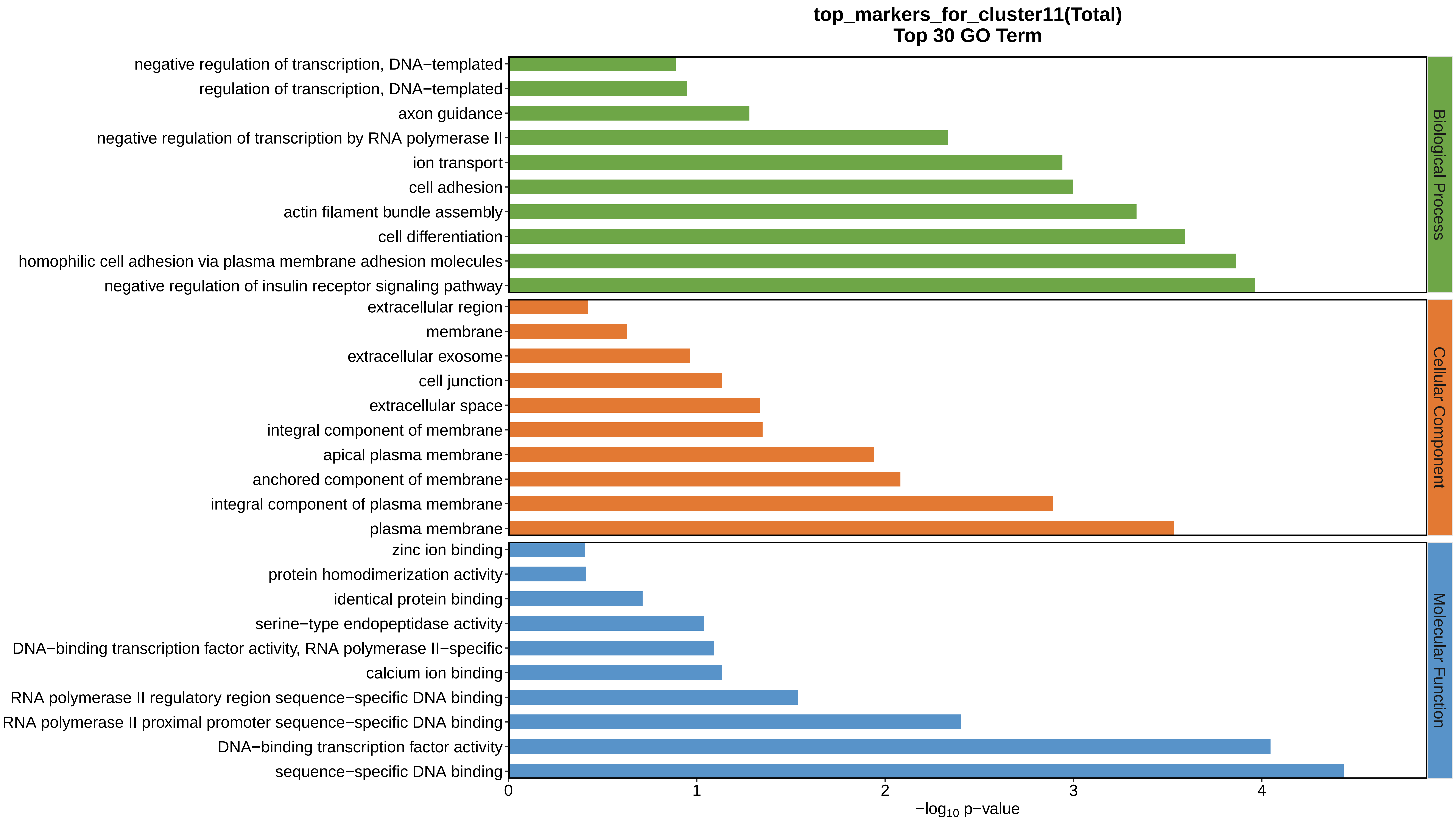
top_markers_for_cluster11(Total)
Top 30 GO Term
negative regulation of transcription, DNA−templated
regulation of transcription, DNA−templated
axon guidance
negative regulation of transcription by RNA polymerase II
ion transport
cell adhesion
actin filament bundle assembly
cell differentiation
homophilic cell adhesion via plasma membrane adhesion molecules
negative regulation of insulin receptor signaling pathway
extracellular region
membrane
extracellular exosome
cell junction
extracellular space
integral component of membrane
apical plasma membrane
anchored component of membrane
integral component of plasma membrane
plasma membrane
zinc ion binding
protein homodimerization activity
identical protein binding
serine−type endopeptidase activity
DNA−binding transcription factor activity, RNA polymerase II−specific
calcium ion binding
RNA polymerase II regulatory region sequence−specific DNA binding
RNA polymerase II proximal promoter sequence−specific DNA binding
DNA−binding transcription factor activity
sequence−specific DNA binding
Biological Process
Cellular Component
Molecular Function
0
1
2
3
4
−log10 p−value
