## Supplemental Data 1-8 for "A Spatiotemporal Embryonic Atlas of Bombyx mori Integrating Single-Nucleus RNA Sequencing and Spatial Transcriptomics": Cluster12 GO top100.pdf

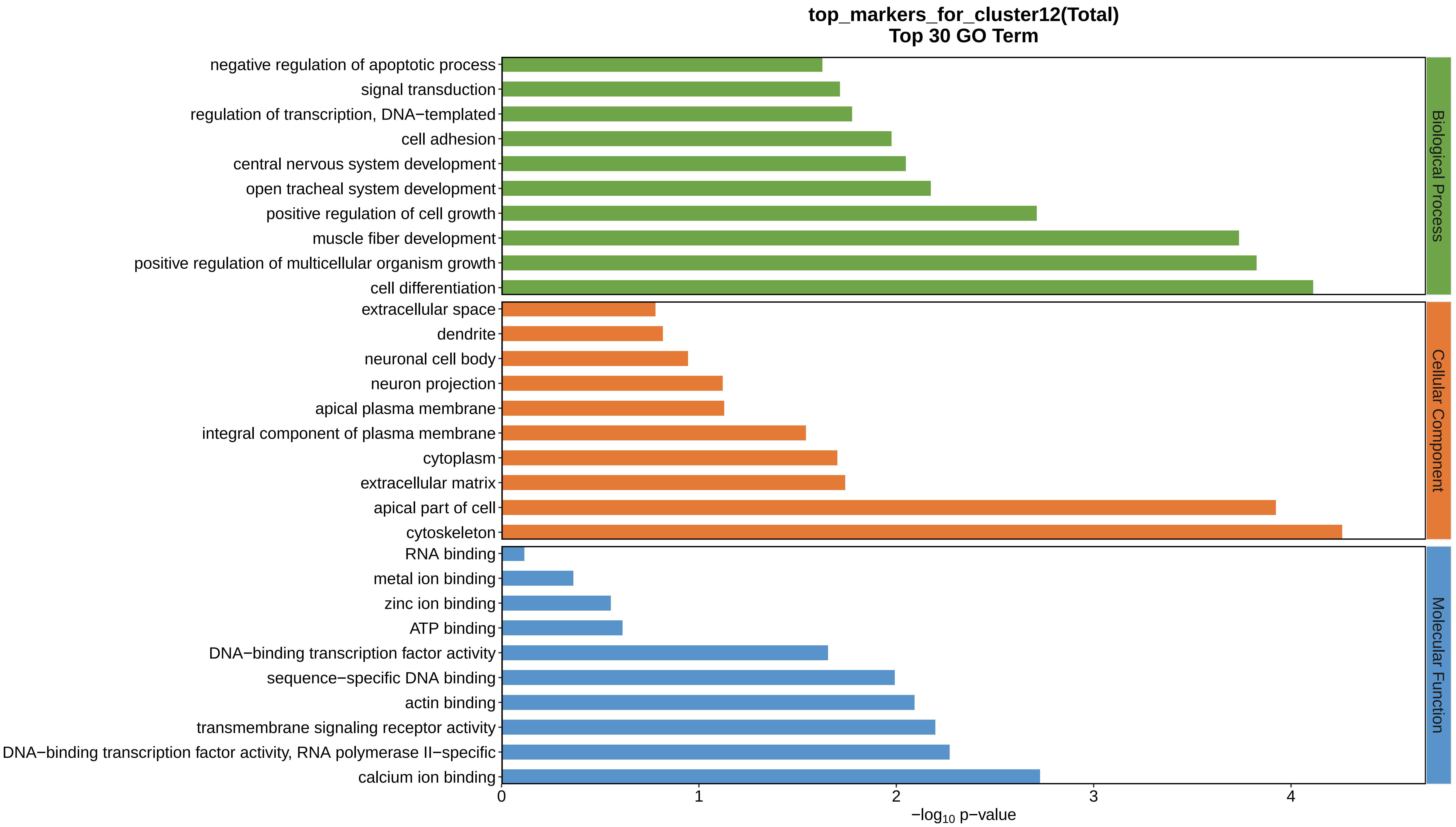
top_markers_for_cluster12(Total)
Top 30 GO Term
negative regulation of apoptotic process
signal transduction
regulation of transcription, DNA−templated
cell adhesion
central nervous system development
open tracheal system development
positive regulation of cell growth
muscle fiber development
positive regulation of multicellular organism growth
cell differentiation
extracellular space
dendrite
neuronal cell body
neuron projection
apical plasma membrane
integral component of plasma membrane
cytoplasm
extracellular matrix
apical part of cell
cytoskeleton
RNA binding
metal ion binding
zinc ion binding
ATP binding
DNA−binding transcription factor activity
sequence−specific DNA binding
actin binding
transmembrane signaling receptor activity
DNA−binding transcription factor activity, RNA polymerase II−specific
calcium ion binding
Biological Process
Cellular Component
Molecular Function
0
1
2
3
4
−log10 p−value
