## Supplemental Data 1-8 for "A Spatiotemporal Embryonic Atlas of Bombyx mori Integrating Single-Nucleus RNA Sequencing and Spatial Transcriptomics": Cluster13 GO top100.pdf

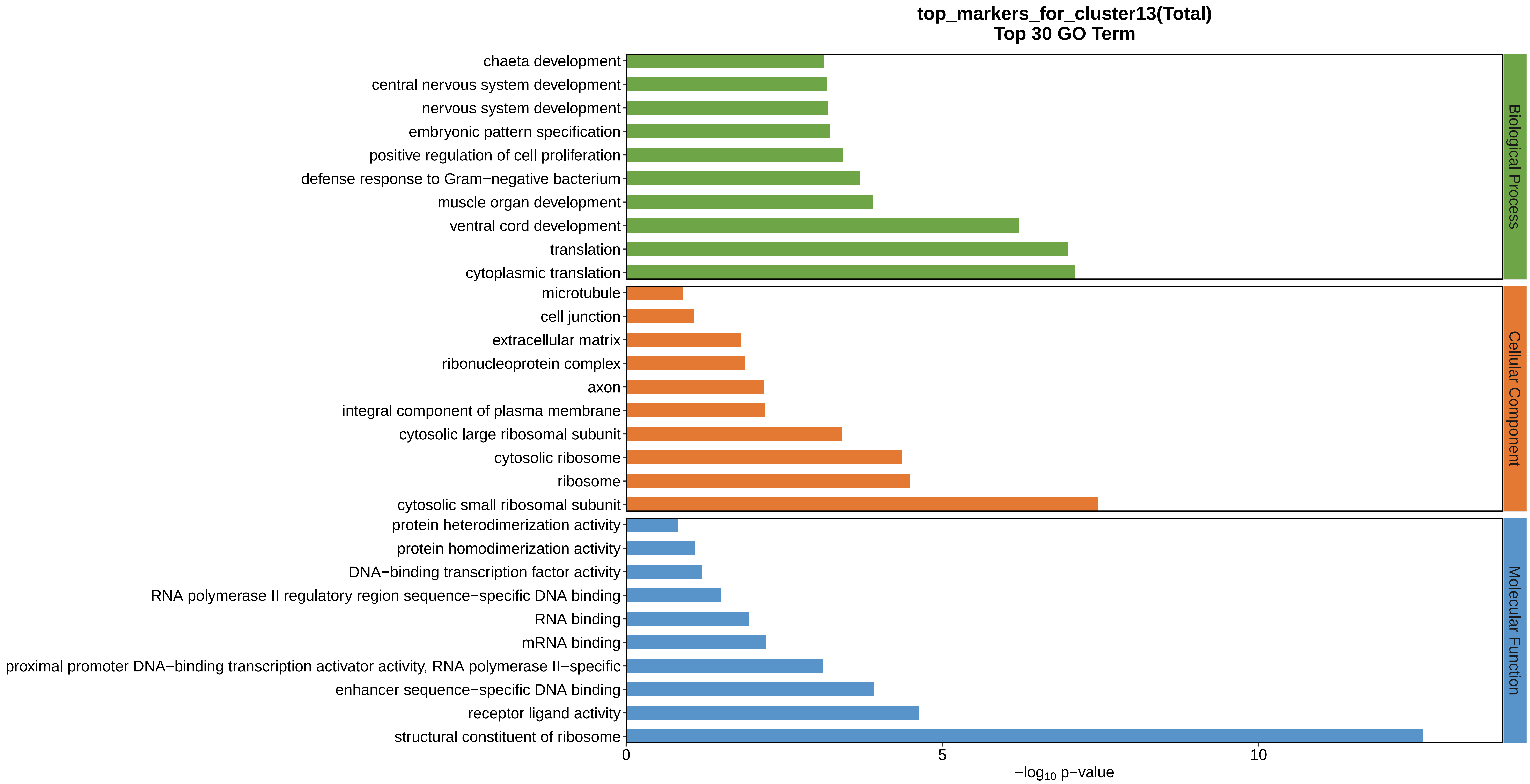

top_markers_for_cluster13(Total)
Top 30 GO Term
chaeta development
central nervous system development
nervous system development
embryonic pattern specification
positive regulation of cell proliferation
defense response to Gram−negative bacterium
muscle organ development
ventral cord development
translation
cytoplasmic translation
microtubule
cell junction
extracellular matrix
ribonucleoprotein complex
axon
integral component of plasma membrane
cytosolic large ribosomal subunit
cytosolic ribosome
ribosome
cytosolic small ribosomal subunit
protein heterodimerization activity
protein homodimerization activity
DNA−binding transcription factor activity
RNA polymerase II regulatory region sequence−specific DNA binding
RNA binding
mRNA binding
proximal promoter DNA−binding transcription activator activity, RNA polymerase II−specific
enhancer sequence−specific DNA binding
receptor ligand activity
structural constituent of ribosome
Biological Process
Cellular Component
Molecular Function
0
5
10
$-\log_{10}$ p−value
