## Supplemental Data 1-8 for "A Spatiotemporal Embryonic Atlas of Bombyx mori Integrating Single-Nucleus RNA Sequencing and Spatial Transcriptomics": Cluster14 GO top100.pdf

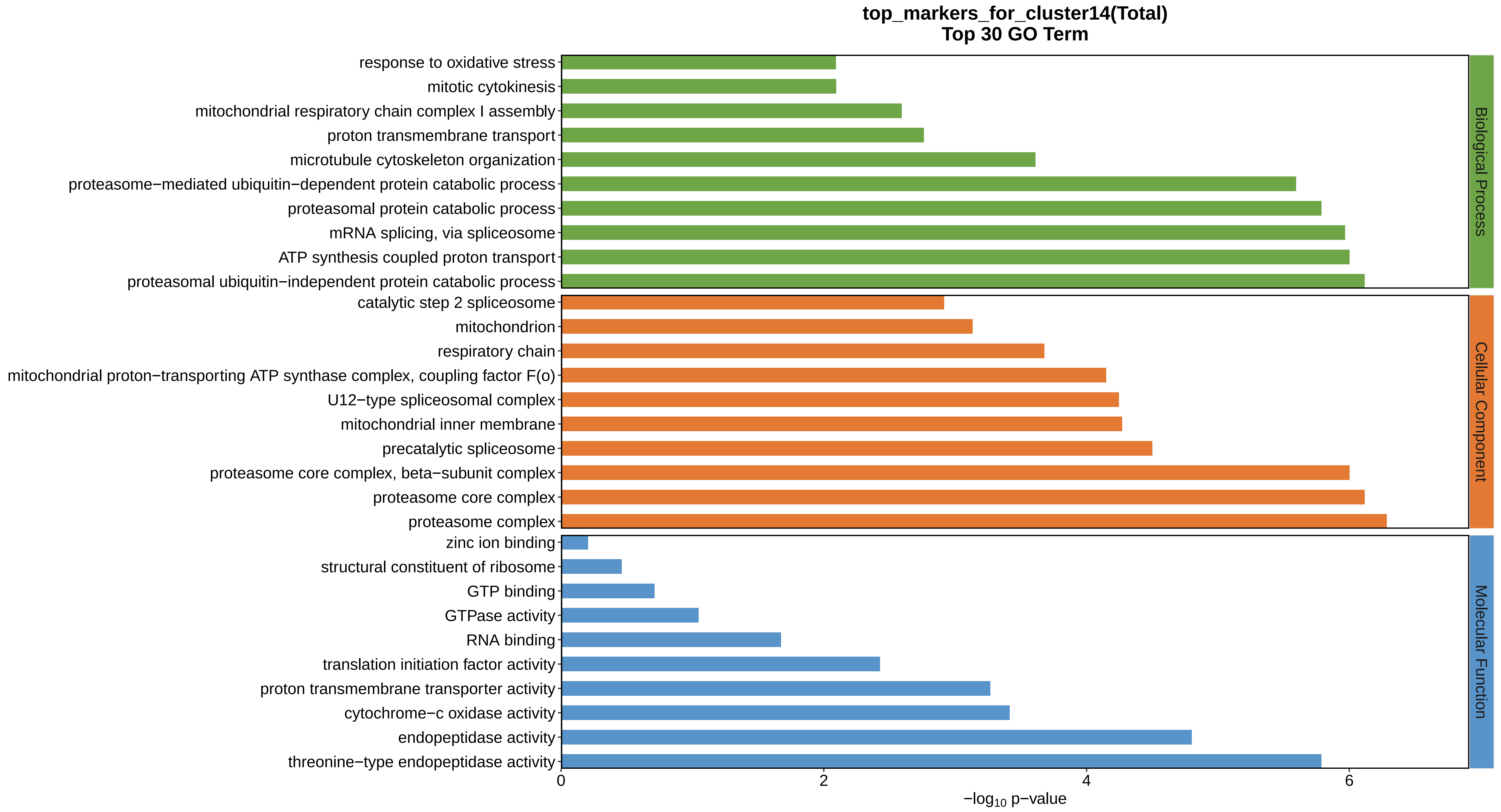
top_markers_for_cluster14(Total)
Top 30 GO Term
response to oxidative stress
mitotic cytokinesis
mitochondrial respiratory chain complex I assembly
proton transmembrane transport
microtubule cytoskeleton organization
proteasome−mediated ubiquitin−dependent protein catabolic process
proteasomal protein catabolic process
mRNA splicing, via spliceosome
ATP synthesis coupled proton transport
proteasomal ubiquitin−independent protein catabolic process
catalytic step 2 spliceosome
mitochondrion
respiratory chain
mitochondrial proton−transporting ATP synthase complex, coupling factor F(o)
U12−type spliceosomal complex
mitochondrial inner membrane
precatalytic spliceosome
proteasome core complex, beta−subunit complex
proteasome core complex
proteasome complex
zinc ion binding
structural constituent of ribosome
GTP binding
GTPase activity
RNA binding
translation initiation factor activity
proton transmembrane transporter activity
cytochrome−c oxidase activity
endopeptidase activity
threonine−type endopeptidase activity
Biological Process
Cellular Component
Molecular Function
0
2
4
6
−log10 p−value
