## Supplemental Data 1-8 for "A Spatiotemporal Embryonic Atlas of Bombyx mori Integrating Single-Nucleus RNA Sequencing and Spatial Transcriptomics": Cluster15 GO top100.pdf

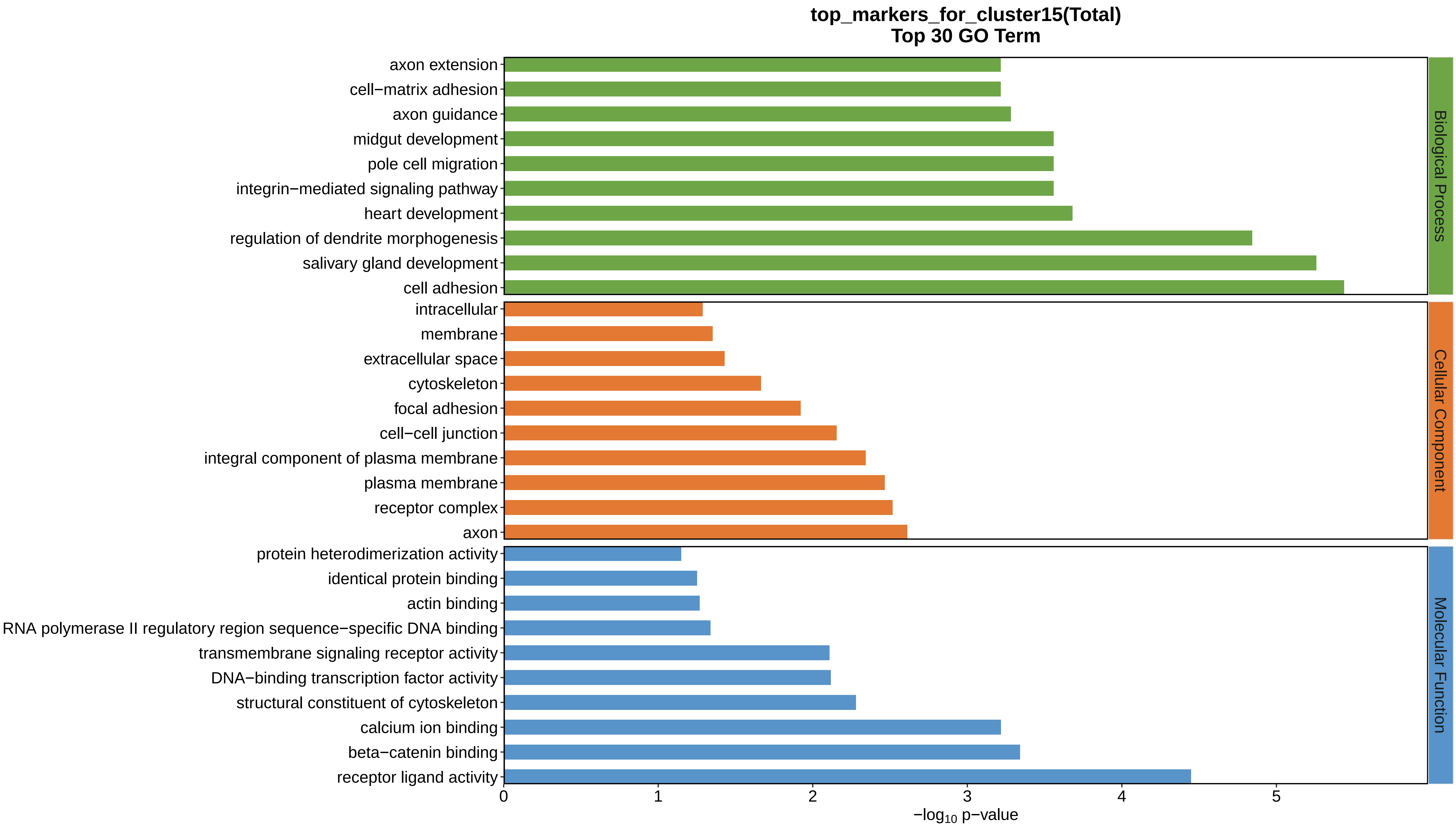
top_markers_for_cluster15(Total)
Top 30 GO Term
axon extension
cell−matrix adhesion
axon guidance
midgut development
pole cell migration
integrin−mediated signaling pathway
heart development
regulation of dendrite morphogenesis
salivary gland development
cell adhesion
intracellular
membrane
extracellular space
cytoskeleton
focal adhesion
cell−cell junction
integral component of plasma membrane
plasma membrane
receptor complex
axon
protein heterodimerization activity
identical protein binding
actin binding
RNA polymerase II regulatory region sequence−specific DNA binding
transmembrane signaling receptor activity
DNA−binding transcription factor activity
structural constituent of cytoskeleton
calcium ion binding
beta−catenin binding
receptor ligand activity
Biological Process
Cellular Component
Molecular Function
0
1
2
3
4
5
−log10 p−value
