## Supplemental Data 1-8 for "A Spatiotemporal Embryonic Atlas of Bombyx mori Integrating Single-Nucleus RNA Sequencing and Spatial Transcriptomics": Cluster16 GO top100.pdf

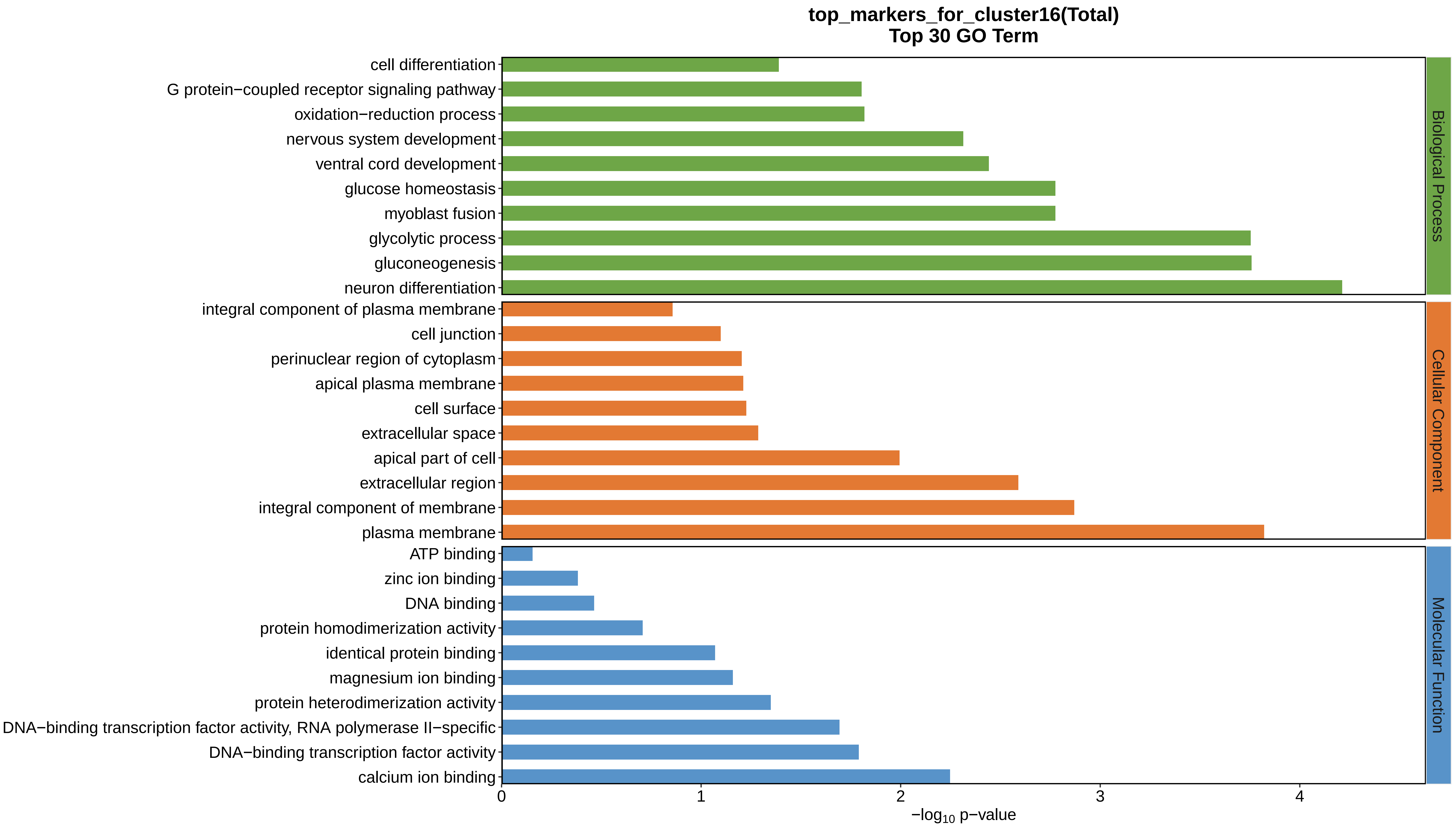
top_markers_for_cluster16(Total)
Top 30 GO Term
cell differentiation
G protein−coupled receptor signaling pathway
oxidation−reduction process
nervous system development
ventral cord development
glucose homeostasis
myoblast fusion
glycolytic process
gluconeogenesis
neuron differentiation
integral component of plasma membrane
cell junction
perinuclear region of cytoplasm
apical plasma membrane
cell surface
extracellular space
apical part of cell
extracellular region
integral component of membrane
plasma membrane
ATP binding
zinc ion binding
DNA binding
protein homodimerization activity
identical protein binding
magnesium ion binding
protein heterodimerization activity
DNA−binding transcription factor activity, RNA polymerase II−specific
DNA−binding transcription factor activity
calcium ion binding
0
1
2
3
4
−log10 p−value
Biological Process
Cellular Component
Molecular Function
