## Supplemental Data 1-8 for "A Spatiotemporal Embryonic Atlas of Bombyx mori Integrating Single-Nucleus RNA Sequencing and Spatial Transcriptomics": Cluster17 GO top100.pdf

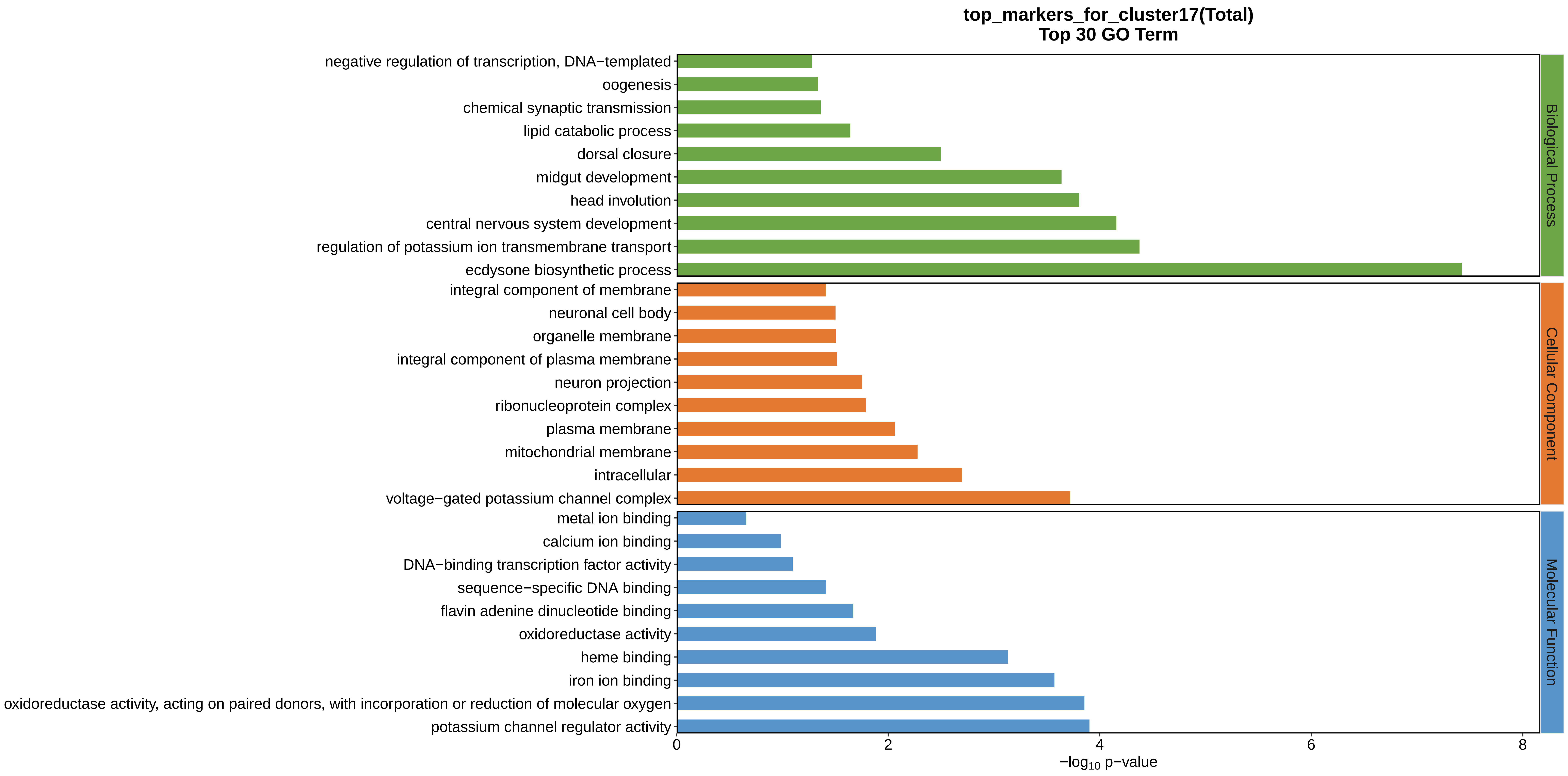

top_markers_for_cluster17(Total)
Top 30 GO Term
negative regulation of transcription, DNA−templated
oogenesis
chemical synaptic transmission
lipid catabolic process
dorsal closure
midgut development
head involution
central nervous system development
regulation of potassium ion transmembrane transport
ecdysone biosynthetic process
integral component of membrane
neuronal cell body
organelle membrane
integral component of plasma membrane
neuron projection
ribonucleoprotein complex
plasma membrane
mitochondrial membrane
intracellular
voltage−gated potassium channel complex
metal ion binding
calcium ion binding
DNA−binding transcription factor activity
sequence−specific DNA binding
flavin adenine dinucleotide binding
oxidoreductase activity
heme binding
iron ion binding
oxidoreductase activity, acting on paired donors, with incorporation or reduction of molecular oxygen
potassium channel regulator activity
Biological Process
Cellular Component
Molecular Function
0
2
4
6
8
−log10 p−value
