## Supplemental Data 1-8 for "A Spatiotemporal Embryonic Atlas of Bombyx mori Integrating Single-Nucleus RNA Sequencing and Spatial Transcriptomics": Cluster18 GO top100.pdf

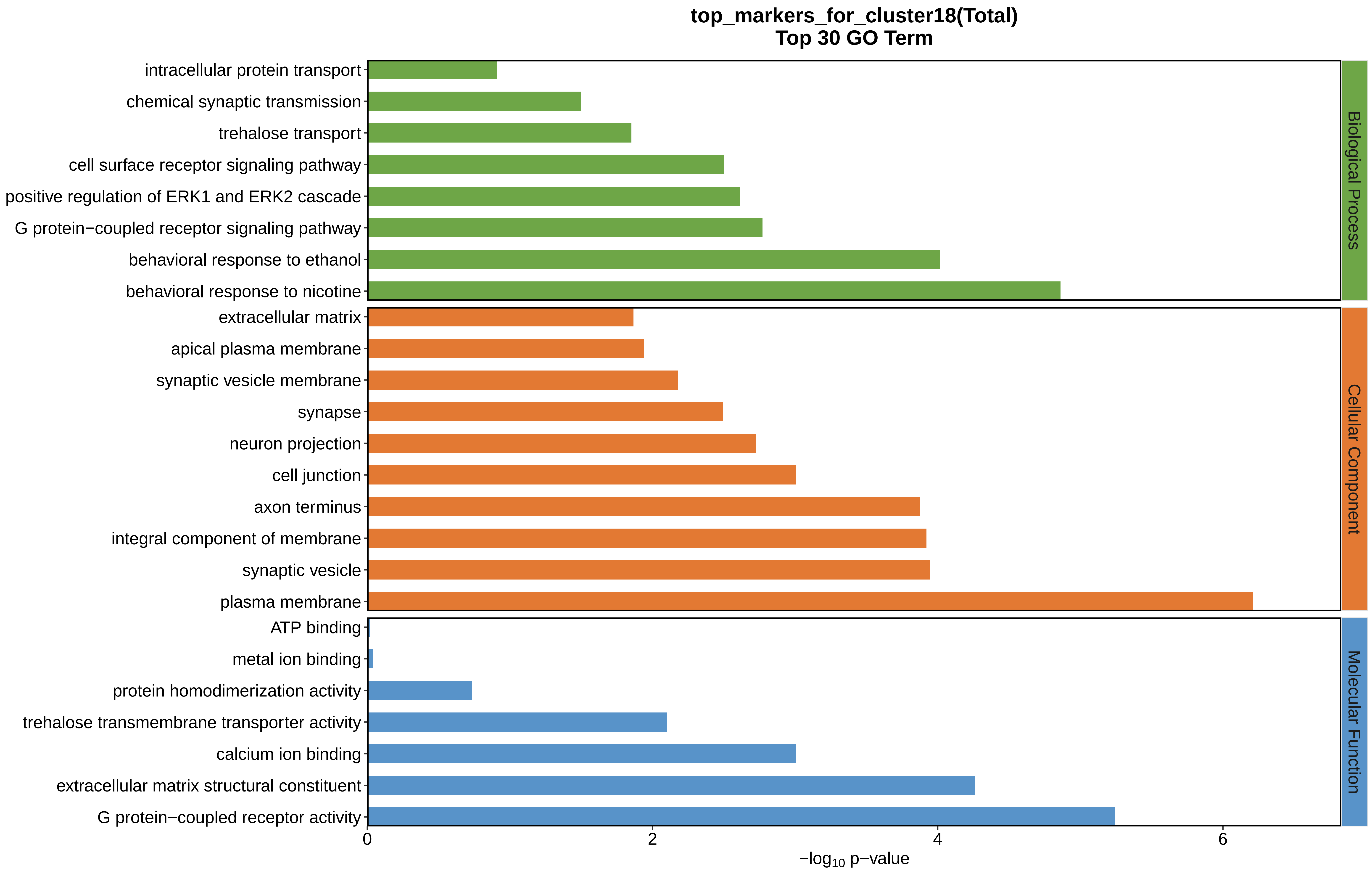
top_markers_for_cluster18(Total)
Top 30 GO Term
intracellular protein transport
chemical synaptic transmission
trehalose transport
cell surface receptor signaling pathway
positive regulation of ERK1 and ERK2 cascade
G protein−coupled receptor signaling pathway
behavioral response to ethanol
behavioral response to nicotine
extracellular matrix
apical plasma membrane
synaptic vesicle membrane
synapse
neuron projection
cell junction
axon terminus
integral component of membrane
synaptic vesicle
plasma membrane
ATP binding
metal ion binding
protein homodimerization activity
trehalose transmembrane transporter activity
calcium ion binding
extracellular matrix structural constituent
G protein−coupled receptor activity
Biological Process
Cellular Component
Molecular Function
0
2
4
6
−log10 p−value
