## Supplemental Data 1-8 for "A Spatiotemporal Embryonic Atlas of Bombyx mori Integrating Single-Nucleus RNA Sequencing and Spatial Transcriptomics": Cluster19 GO top100.pdf

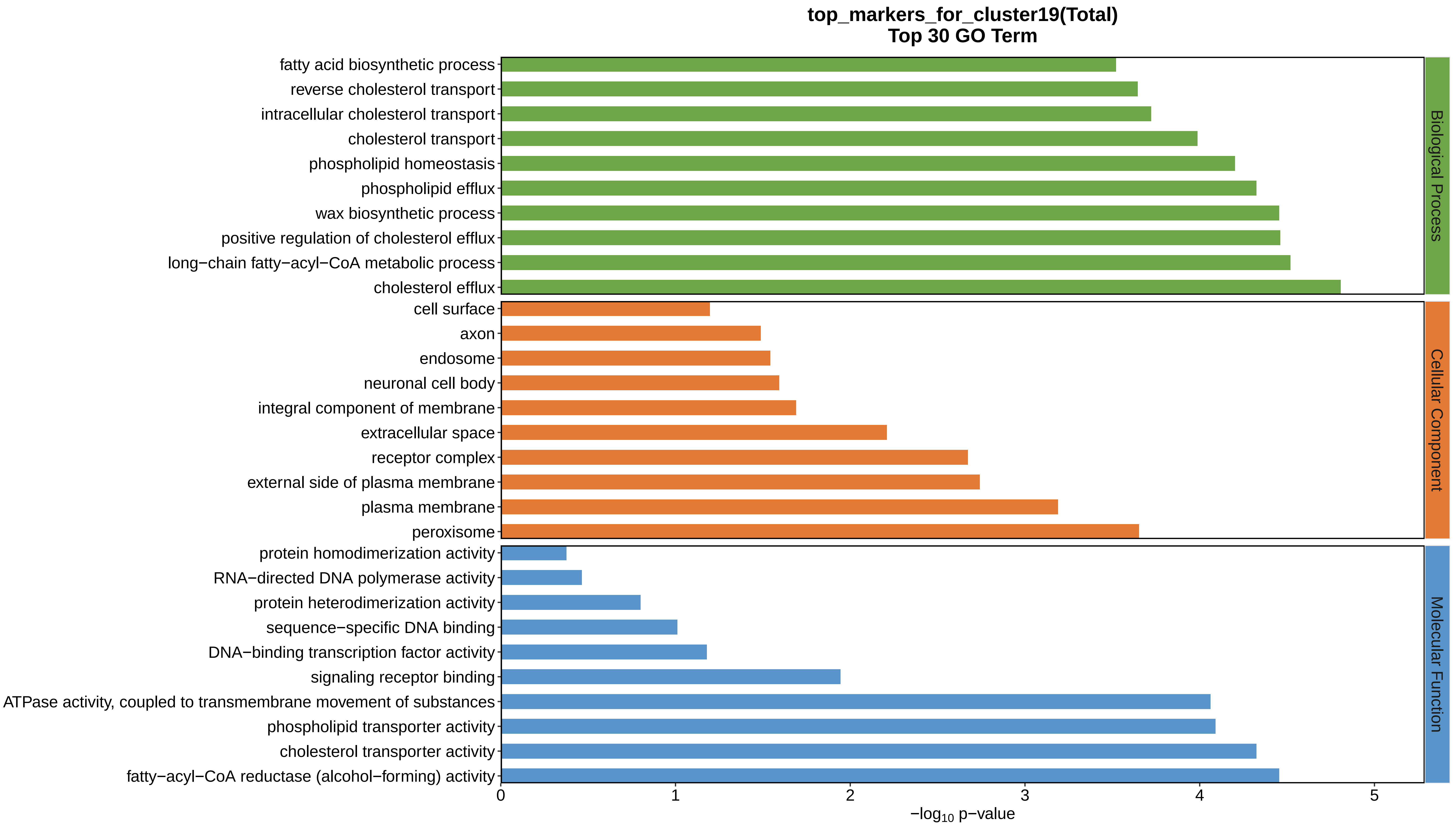
top_markers_for_cluster19(Total)
Top 30 GO Term
fatty acid biosynthetic process
reverse cholesterol transport
intracellular cholesterol transport
cholesterol transport
phospholipid homeostasis
phospholipid efflux
wax biosynthetic process
positive regulation of cholesterol efflux
long−chain fatty−acyl−CoA metabolic process
cholesterol efflux
cell surface
axon
endosome
neuronal cell body
integral component of membrane
extracellular space
receptor complex
external side of plasma membrane
plasma membrane
peroxisome
protein homodimerization activity
RNA−directed DNA polymerase activity
protein heterodimerization activity
sequence−specific DNA binding
DNA−binding transcription factor activity
signaling receptor binding
ATPase activity, coupled to transmembrane movement of substances
phospholipid transporter activity
cholesterol transporter activity
fatty−acyl−CoA reductase (alcohol−forming) activity
Biological Process
Cellular Component
Molecular Function
0
1
2
3
4
5
$-\log_{10}$ p−value
