## Supplemental Data 1-8 for "A Spatiotemporal Embryonic Atlas of Bombyx mori Integrating Single-Nucleus RNA Sequencing and Spatial Transcriptomics": Cluster20 GO top100.pdf

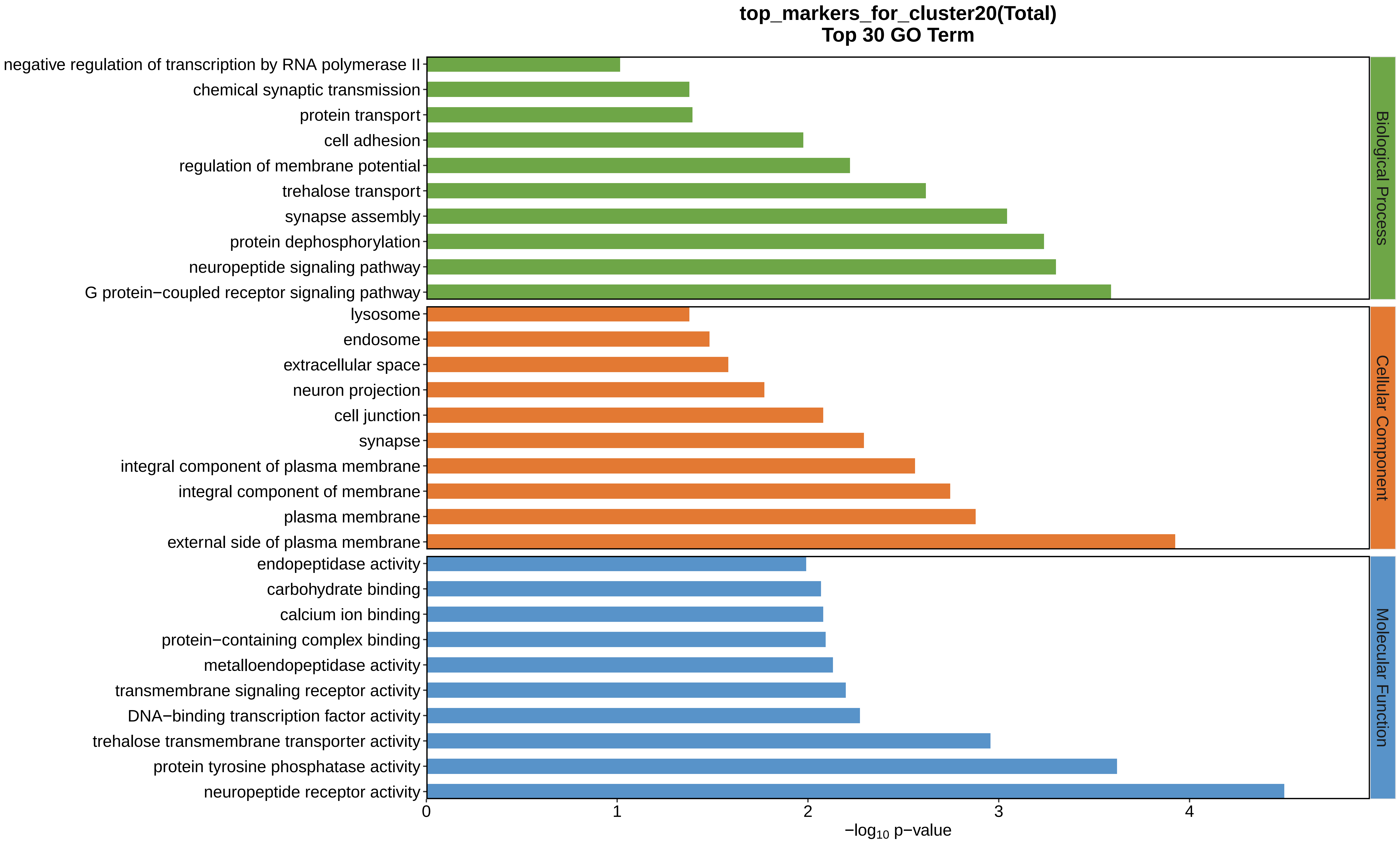

top_markers_for_cluster20(Total)
Top 30 GO Term
negative regulation of transcription by RNA polymerase II
chemical synaptic transmission
protein transport
cell adhesion
regulation of membrane potential
trehalose transport
synapse assembly
protein dephosphorylation
neuropeptide signaling pathway
G protein−coupled receptor signaling pathway
lysosome
endosome
extracellular space
neuron projection
cell junction
synapse
integral component of plasma membrane
integral component of membrane
plasma membrane
external side of plasma membrane
endopeptidase activity
carbohydrate binding
calcium ion binding
protein−containing complex binding
metalloendopeptidase activity
transmembrane signaling receptor activity
DNA−binding transcription factor activity
trehalose transmembrane transporter activity
protein tyrosine phosphatase activity
neuropeptide receptor activity
Biological Process
Cellular Component
Molecular Function
0
1
2
3
4
$-\log_{10}$ p−value
