## Supplemental Data 1-8 for "A Spatiotemporal Embryonic Atlas of Bombyx mori Integrating Single-Nucleus RNA Sequencing and Spatial Transcriptomics": Cluster21 GO top100.pdf

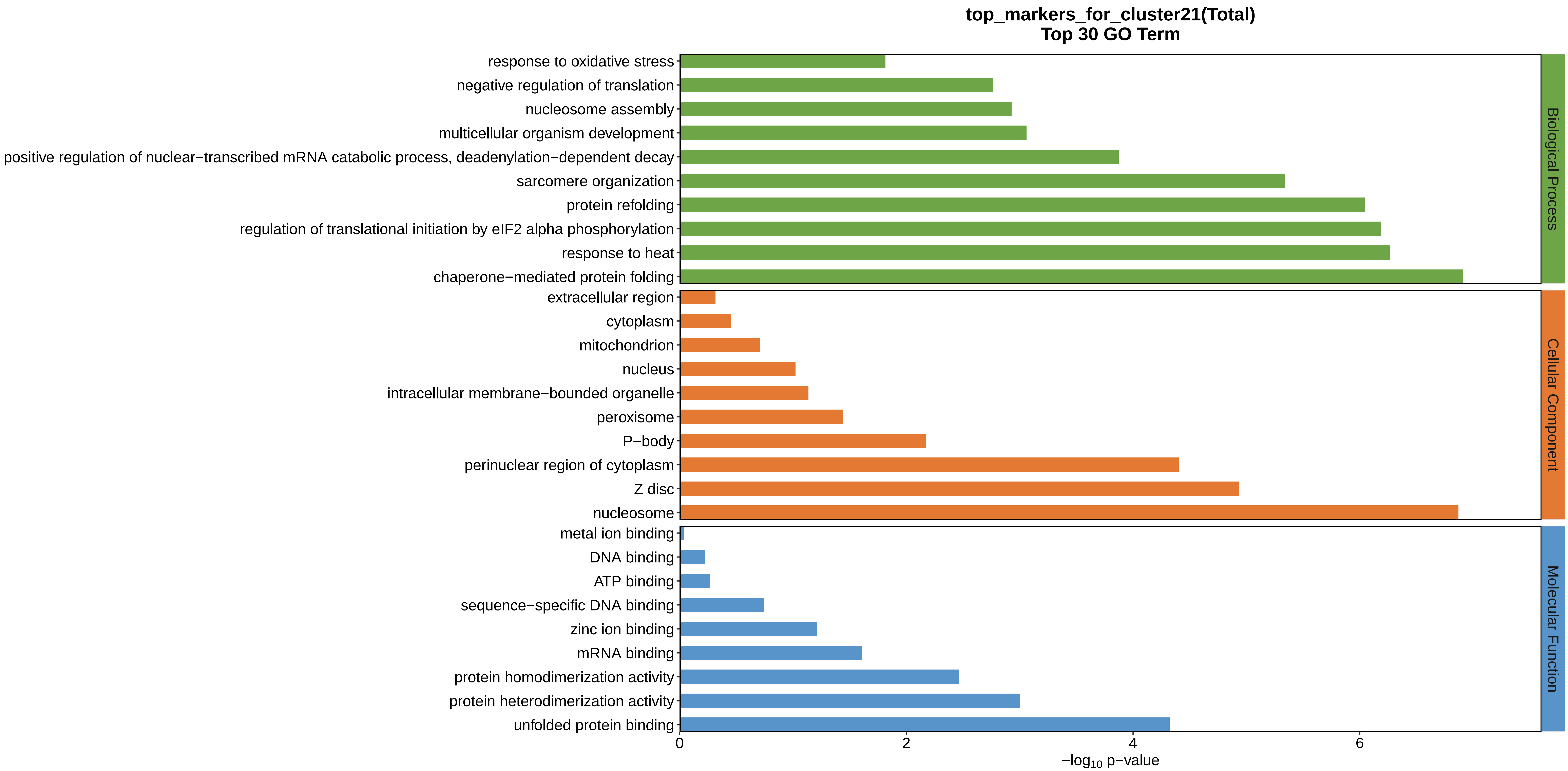
top_markers_for_cluster21(Total)
Top 30 GO Term
response to oxidative stress
negative regulation of translation
nucleosome assembly
multicellular organism development
positive regulation of nuclear−transcribed mRNA catabolic process, deadenylation−dependent decay
sarcomere organization
protein refolding
regulation of translational initiation by eIF2 alpha phosphorylation
response to heat
chaperone−mediated protein folding
extracellular region
cytoplasm
mitochondrion
nucleus
intracellular membrane−bounded organelle
peroxisome
P−body
perinuclear region of cytoplasm
Z disc
nucleosome
metal ion binding
DNA binding
ATP binding
sequence−specific DNA binding
zinc ion binding
mRNA binding
protein homodimerization activity
protein heterodimerization activity
unfolded protein binding
Biological Process
Cellular Component
Molecular Function
0
2
4
6
$-\log_{10}$ p−value
