## Supplemental Data 1-8 for "A Spatiotemporal Embryonic Atlas of Bombyx mori Integrating Single-Nucleus RNA Sequencing and Spatial Transcriptomics": Cluster22 GO top100.pdf

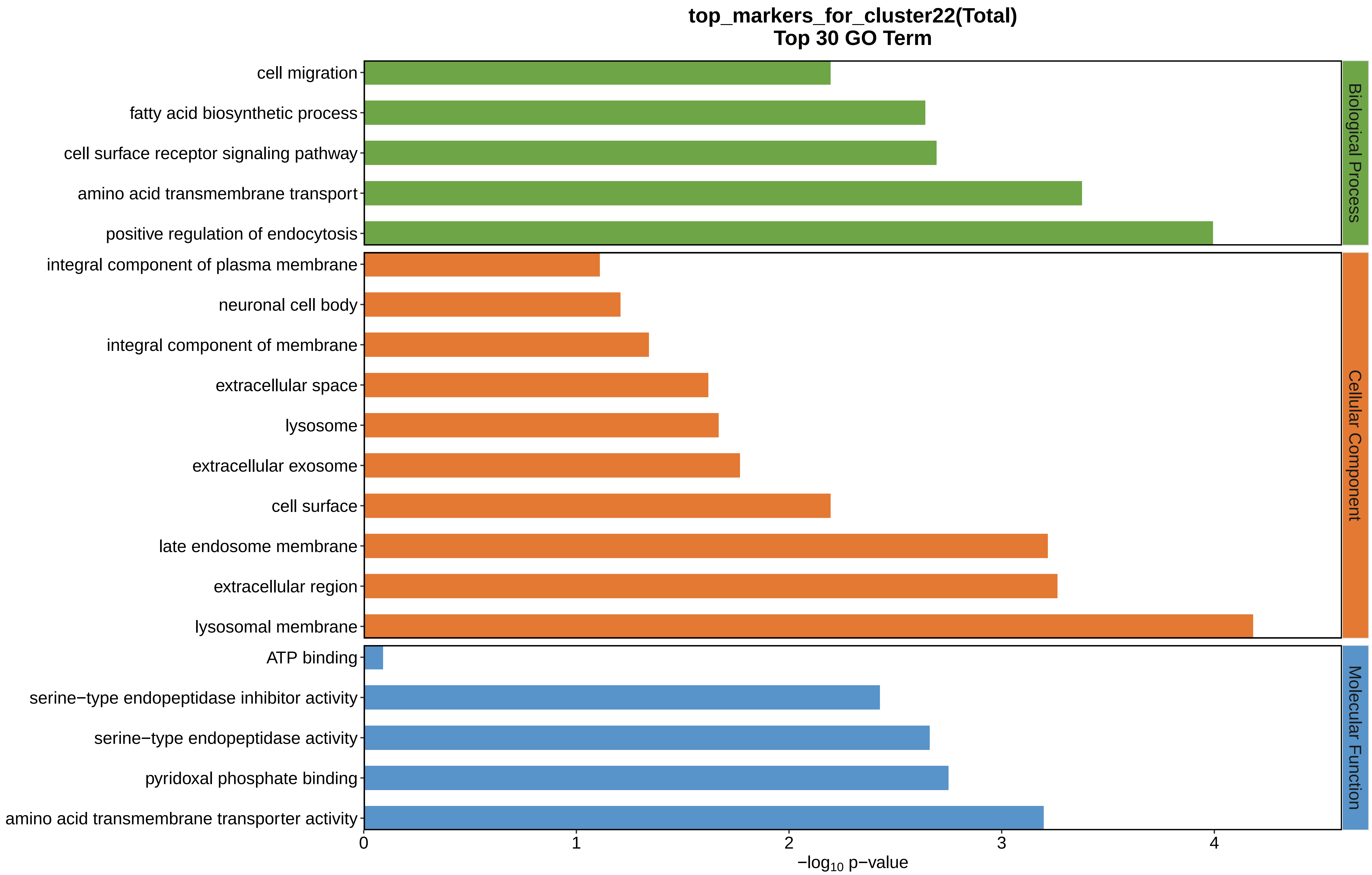

top_markers_for_cluster22(Total)
Top 30 GO Term
cell migration
fatty acid biosynthetic process
cell surface receptor signaling pathway
amino acid transmembrane transport
positive regulation of endocytosis
integral component of plasma membrane
neuronal cell body
integral component of membrane
extracellular space
lysosome
extracellular exosome
cell surface
late endosome membrane
extracellular region
lysosomal membrane
ATP binding
serine−type endopeptidase inhibitor activity
serine−type endopeptidase activity
pyridoxal phosphate binding
amino acid transmembrane transporter activity
Biological Process
Cellular Component
Molecular Function
0
1
2
3
4
−log10 p−value
