## Supplemental Data 1-8 for "A Spatiotemporal Embryonic Atlas of Bombyx mori Integrating Single-Nucleus RNA Sequencing and Spatial Transcriptomics": Cluster1 GO top100.pdf

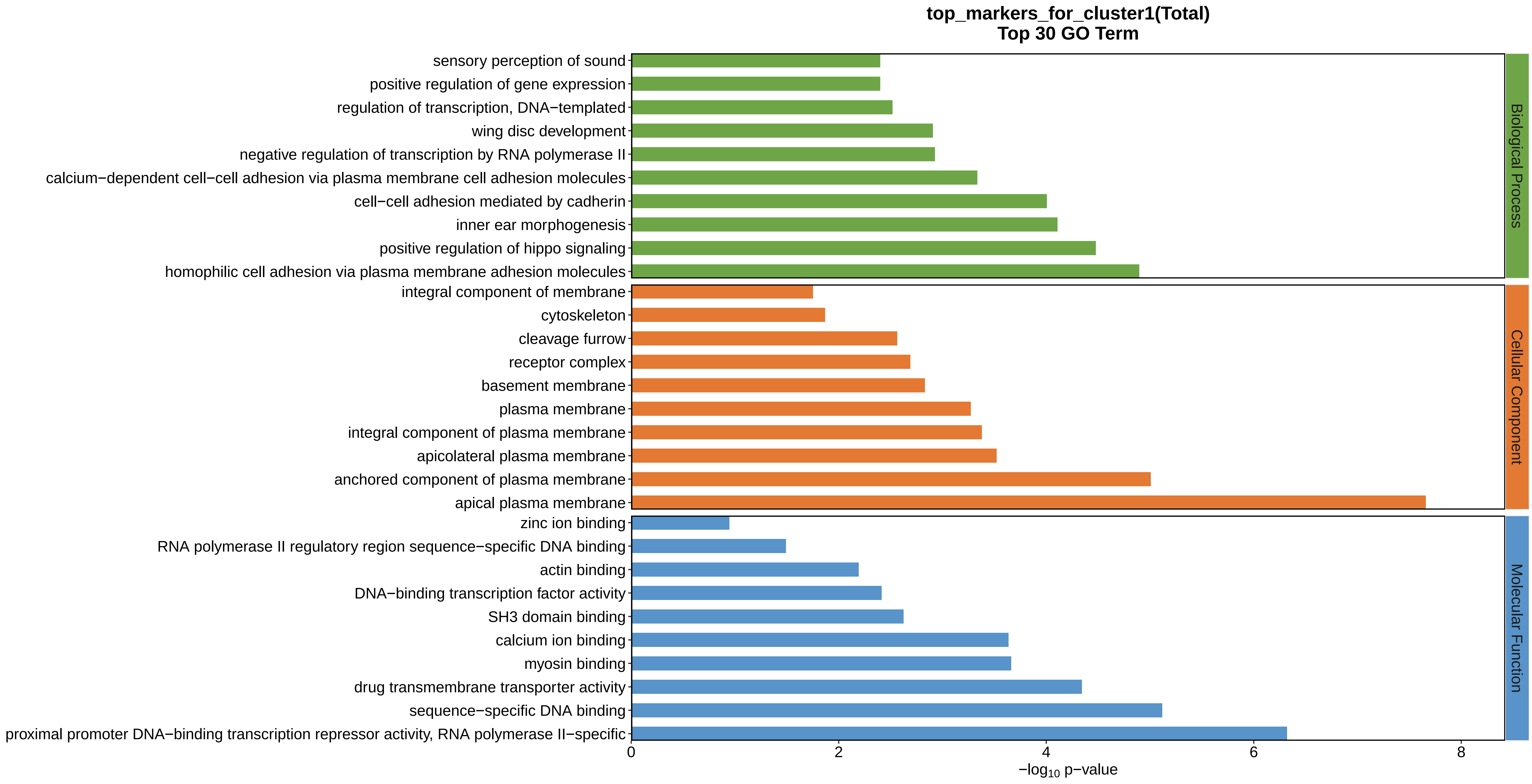

top_markers_for_cluster1(Total)
Top 30 GO Term
sensory perception of sound
positive regulation of gene expression
regulation of transcription, DNA−templated
wing disc development
negative regulation of transcription by RNA polymerase II
calcium−dependent cell−cell adhesion via plasma membrane cell adhesion molecules
cell−cell adhesion mediated by cadherin
inner ear morphogenesis
positive regulation of hippo signaling
homophilic cell adhesion via plasma membrane adhesion molecules
integral component of membrane
cytoskeleton
cleavage furrow
receptor complex
basement membrane
plasma membrane
integral component of plasma membrane
apicolateral plasma membrane
anchored component of plasma membrane
apical plasma membrane
zinc ion binding
RNA polymerase II regulatory region sequence−specific DNA binding
actin binding
DNA−binding transcription factor activity
SH3 domain binding
calcium ion binding
myosin binding
drug transmembrane transporter activity
sequence−specific DNA binding
proximal promoter DNA−binding transcription repressor activity, RNA polymerase II−specific
Biological Process
Cellular Component
Molecular Function
0
2
4
6
8
$-\log_{10}$ p−value
