## Supplemental Data 1-8 for "A Spatiotemporal Embryonic Atlas of Bombyx mori Integrating Single-Nucleus RNA Sequencing and Spatial Transcriptomics": Cluster2 GO top100.pdf

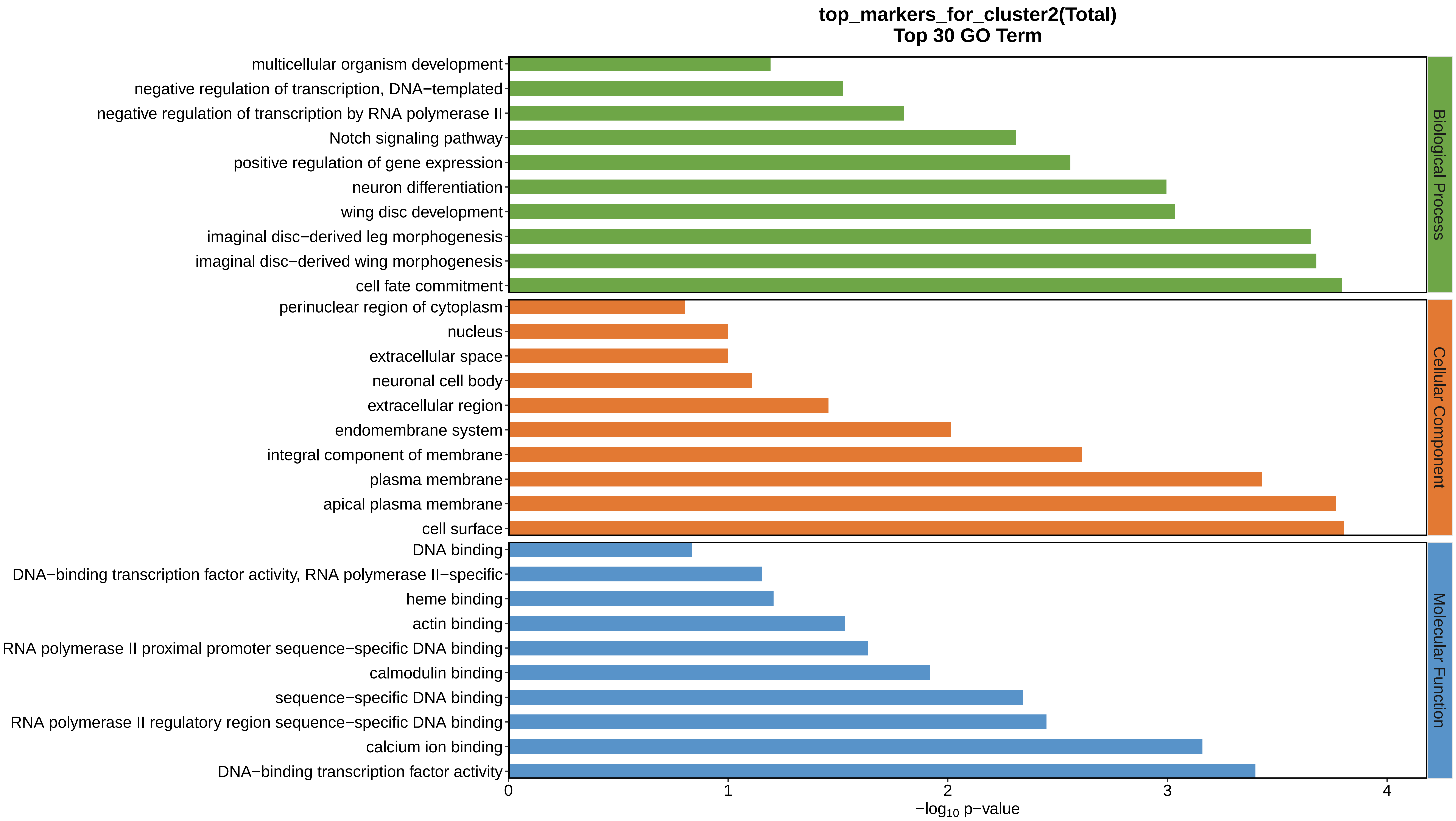
top_markers_for_cluster2(Total)
Top 30 GO Term
multicellular organism development
negative regulation of transcription, DNA−templated
negative regulation of transcription by RNA polymerase II
Notch signaling pathway
positive regulation of gene expression
neuron differentiation
wing disc development
imaginal disc−derived leg morphogenesis
imaginal disc−derived wing morphogenesis
cell fate commitment
perinuclear region of cytoplasm
nucleus
extracellular space
neuronal cell body
extracellular region
endomembrane system
integral component of membrane
plasma membrane
apical plasma membrane
cell surface
DNA binding
DNA−binding transcription factor activity, RNA polymerase II−specific
heme binding
actin binding
RNA polymerase II proximal promoter sequence−specific DNA binding
calmodulin binding
sequence−specific DNA binding
RNA polymerase II regulatory region sequence−specific DNA binding
calcium ion binding
DNA−binding transcription factor activity
Biological Process
Cellular Component
Molecular Function
0
1
2
3
4
$-\log_{10}$ p−value
