## Supplemental Data 1-8 for "A Spatiotemporal Embryonic Atlas of Bombyx mori Integrating Single-Nucleus RNA Sequencing and Spatial Transcriptomics": Cluster3 GO top100.pdf

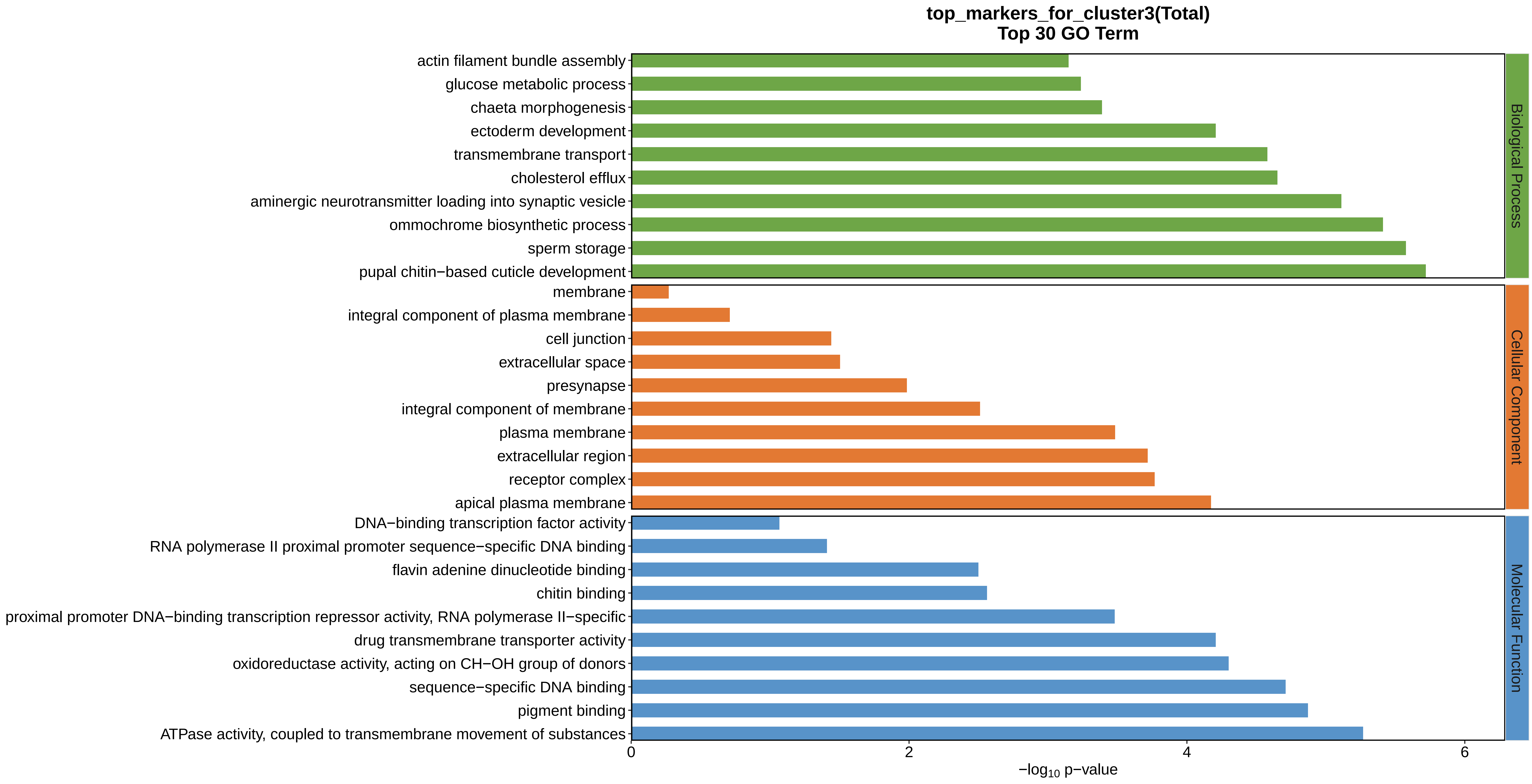
top_markers_for_cluster3(Total)
Top 30 GO Term
actin filament bundle assembly
glucose metabolic process
chaeta morphogenesis
ectoderm development
transmembrane transport
cholesterol efflux
aminergic neurotransmitter loading into synaptic vesicle
ommochrome biosynthetic process
sperm storage
pupal chitin−based cuticle development
membrane
integral component of plasma membrane
cell junction
extracellular space
presynapse
integral component of membrane
plasma membrane
extracellular region
receptor complex
apical plasma membrane
DNA−binding transcription factor activity
RNA polymerase II proximal promoter sequence−specific DNA binding
flavin adenine dinucleotide binding
chitin binding
proximal promoter DNA−binding transcription repressor activity, RNA polymerase II−specific
drug transmembrane transporter activity
oxidoreductase activity, acting on CH−OH group of donors
sequence−specific DNA binding
pigment binding
ATPase activity, coupled to transmembrane movement of substances
Biological Process
Cellular Component
Molecular Function
0
2
4
6
−log10 p−value
