## Supplemental Data 1-8 for "A Spatiotemporal Embryonic Atlas of Bombyx mori Integrating Single-Nucleus RNA Sequencing and Spatial Transcriptomics": Cluster4 GO top100.pdf

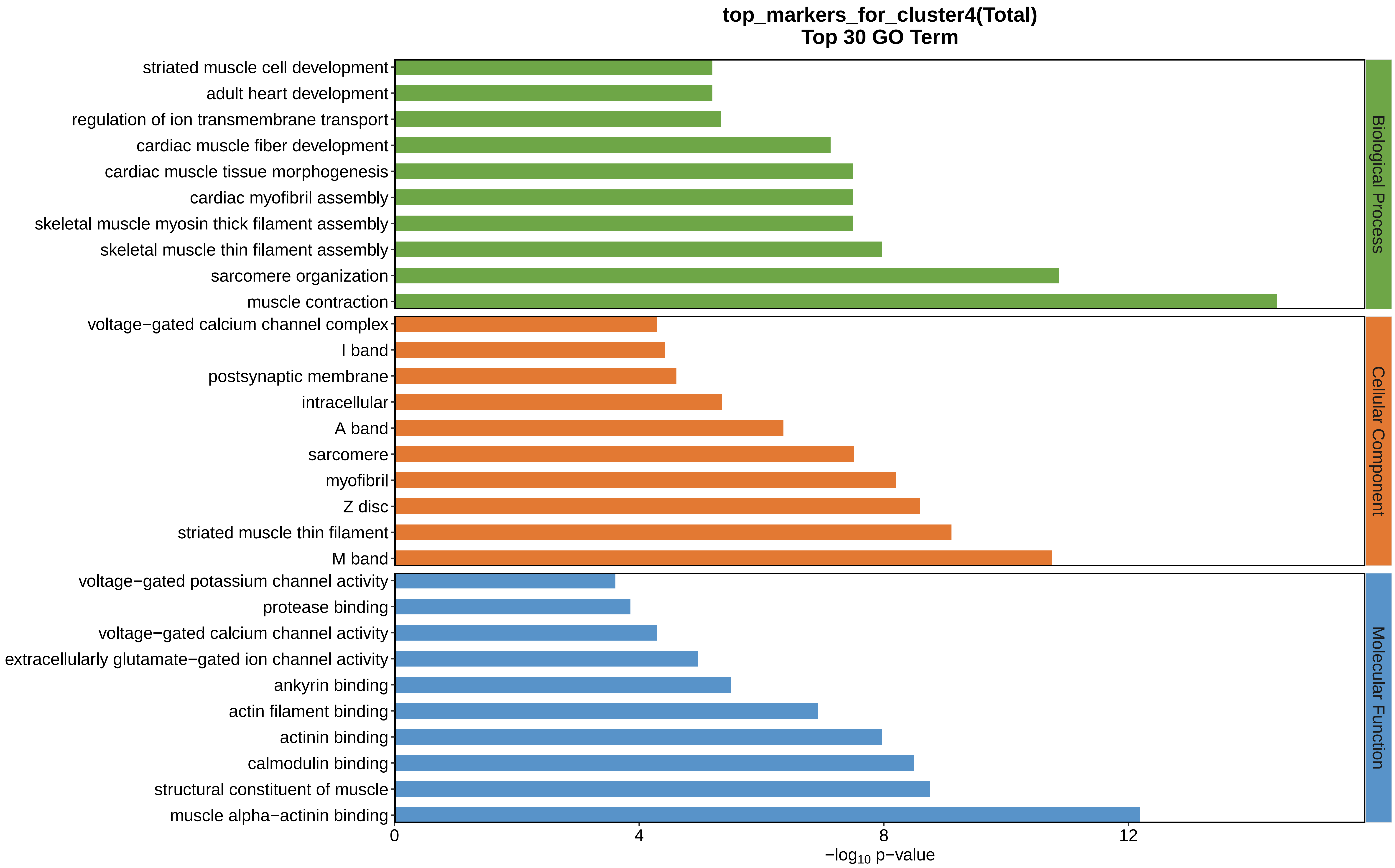
top_markers_for_cluster4(Total)
Top 30 GO Term
striated muscle cell development
adult heart development
regulation of ion transmembrane transport
cardiac muscle fiber development
cardiac muscle tissue morphogenesis
cardiac myofibril assembly
skeletal muscle myosin thick filament assembly
skeletal muscle thin filament assembly
sarcomere organization
muscle contraction
voltage−gated calcium channel complex
I band
postsynaptic membrane
intracellular
A band
sarcomere
myofibril
Z disc
striated muscle thin filament
M band
voltage−gated potassium channel activity
protease binding
voltage−gated calcium channel activity
extracellularly glutamate−gated ion channel activity
ankyrin binding
actin filament binding
actinin binding
calmodulin binding
structural constituent of muscle
muscle alpha−actinin binding
Biological Process
Cellular Component
Molecular Function
0
4
8
12
−log10 p−value
