## Supplemental Data 1-8 for "A Spatiotemporal Embryonic Atlas of Bombyx mori Integrating Single-Nucleus RNA Sequencing and Spatial Transcriptomics": Cluster5 GO top100.pdf

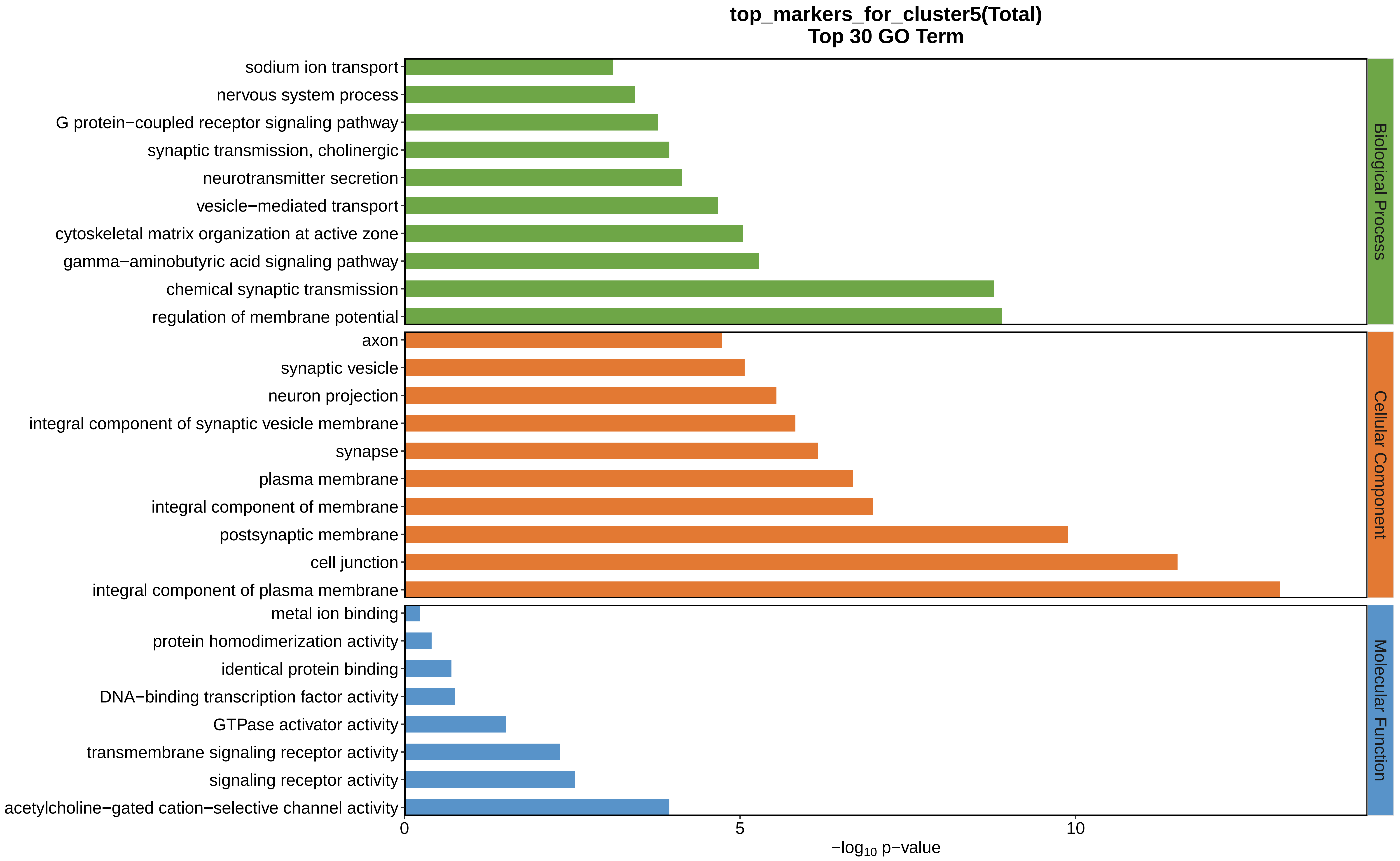

top_markers_for_cluster5(Total)
Top 30 GO Term
sodium ion transport
nervous system process
G protein−coupled receptor signaling pathway
synaptic transmission, cholinergic
neurotransmitter secretion
vesicle−mediated transport
cytoskeletal matrix organization at active zone
gamma−aminobutyric acid signaling pathway
chemical synaptic transmission
regulation of membrane potential
axon
synaptic vesicle
neuron projection
integral component of synaptic vesicle membrane
synapse
plasma membrane
integral component of membrane
postsynaptic membrane
cell junction
integral component of plasma membrane
metal ion binding
protein homodimerization activity
identical protein binding
DNA−binding transcription factor activity
GTPase activator activity
transmembrane signaling receptor activity
signaling receptor activity
acetylcholine−gated cation−selective channel activity
Biological Process
Cellular Component
Molecular Function
0
5
10
−log10 p−value
