## Supplemental Data 1-8 for "A Spatiotemporal Embryonic Atlas of Bombyx mori Integrating Single-Nucleus RNA Sequencing and Spatial Transcriptomics": Cluster6 GO top100.pdf

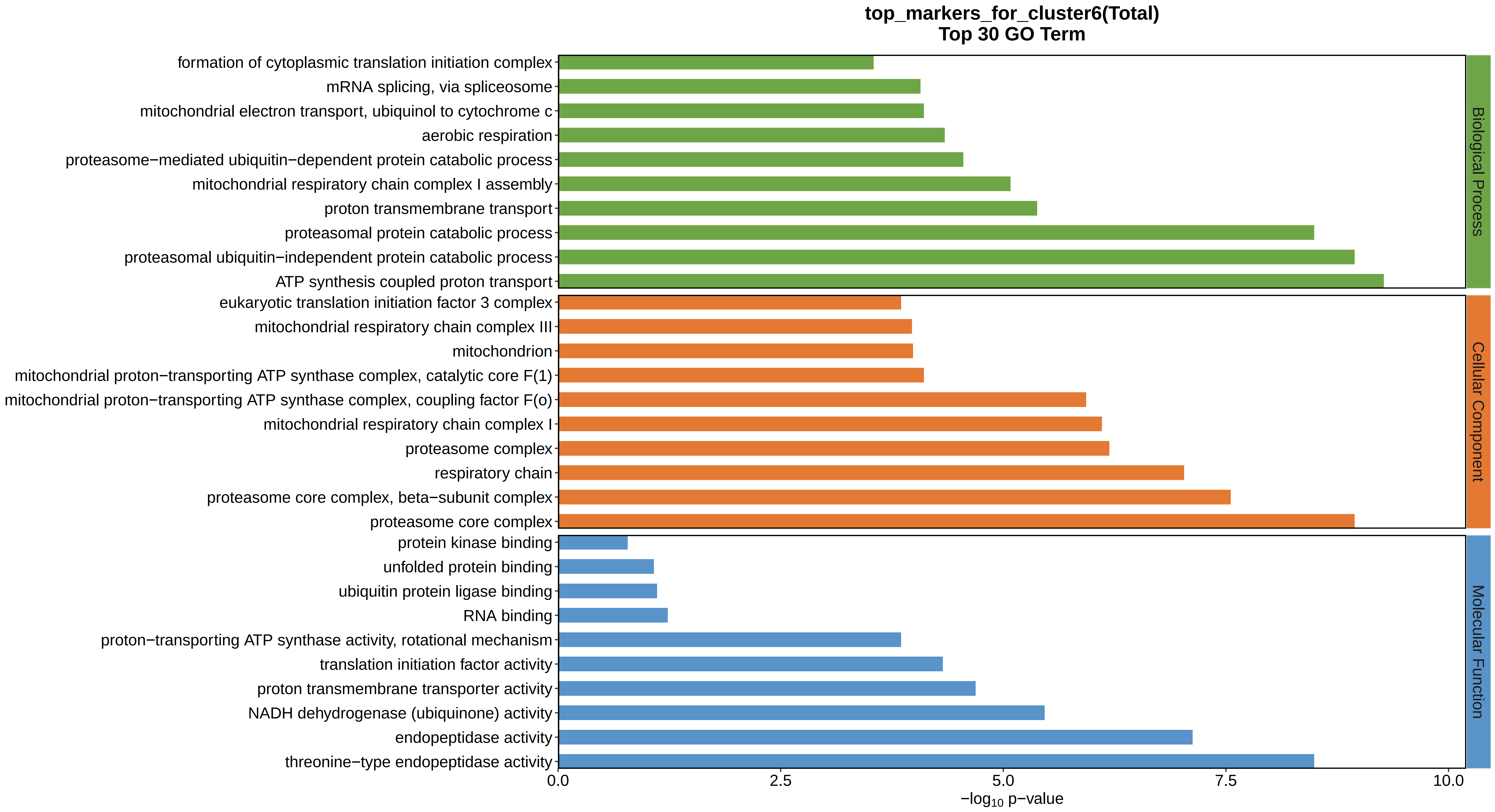
top_markers_for_cluster6(Total)
Top 30 GO Term
formation of cytoplasmic translation initiation complex
mRNA splicing, via spliceosome
mitochondrial electron transport, ubiquinol to cytochrome c
aerobic respiration
proteasome−mediated ubiquitin−dependent protein catabolic process
mitochondrial respiratory chain complex I assembly
proton transmembrane transport
proteasomal protein catabolic process
proteasomal ubiquitin−independent protein catabolic process
ATP synthesis coupled proton transport
eukaryotic translation initiation factor 3 complex
mitochondrial respiratory chain complex III
mitochondrion
mitochondrial proton−transporting ATP synthase complex, catalytic core F(1)
mitochondrial proton−transporting ATP synthase complex, coupling factor F(o)
mitochondrial respiratory chain complex I
proteasome complex
respiratory chain
proteasome core complex, beta−subunit complex
proteasome core complex
protein kinase binding
unfolded protein binding
ubiquitin protein ligase binding
RNA binding
proton−transporting ATP synthase activity, rotational mechanism
translation initiation factor activity
proton transmembrane transporter activity
NADH dehydrogenase (ubiquinone) activity
endopeptidase activity
threonine−type endopeptidase activity
Biological Process
Cellular Component
Molecular Function
0.0
2.5
5.0
7.5
10.0
−log10 p−value
