## Supplemental Data 1-8 for "A Spatiotemporal Embryonic Atlas of Bombyx mori Integrating Single-Nucleus RNA Sequencing and Spatial Transcriptomics": Cluster7 GO top100.pdf

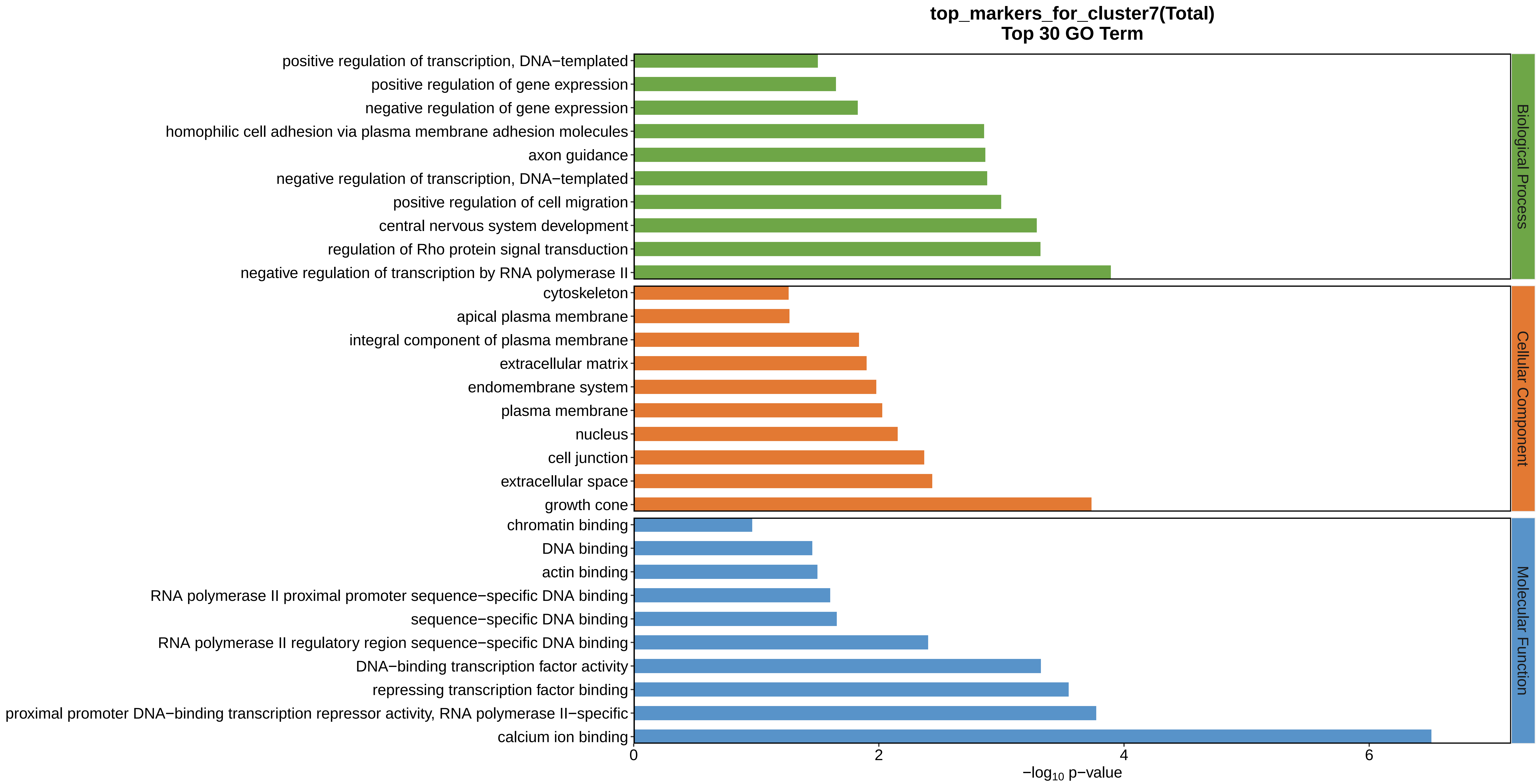
top_markers_for_cluster7(Total)
Top 30 GO Term
positive regulation of transcription, DNA−templated
positive regulation of gene expression
negative regulation of gene expression
homophilic cell adhesion via plasma membrane adhesion molecules
axon guidance
negative regulation of transcription, DNA−templated
positive regulation of cell migration
central nervous system development
regulation of Rho protein signal transduction
negative regulation of transcription by RNA polymerase II
cytoskeleton
apical plasma membrane
integral component of plasma membrane
extracellular matrix
endomembrane system
plasma membrane
nucleus
cell junction
extracellular space
growth cone
chromatin binding
DNA binding
actin binding
RNA polymerase II proximal promoter sequence−specific DNA binding
sequence−specific DNA binding
RNA polymerase II regulatory region sequence−specific DNA binding
DNA−binding transcription factor activity
repressing transcription factor binding
proximal promoter DNA−binding transcription repressor activity, RNA polymerase II−specific
calcium ion binding
Biological Process
Cellular Component
Molecular Function
0
2
4
6
−log10 p−value
