## Supplemental Data 1-8 for "A Spatiotemporal Embryonic Atlas of Bombyx mori Integrating Single-Nucleus RNA Sequencing and Spatial Transcriptomics": Cluster8 GO top100.pdf

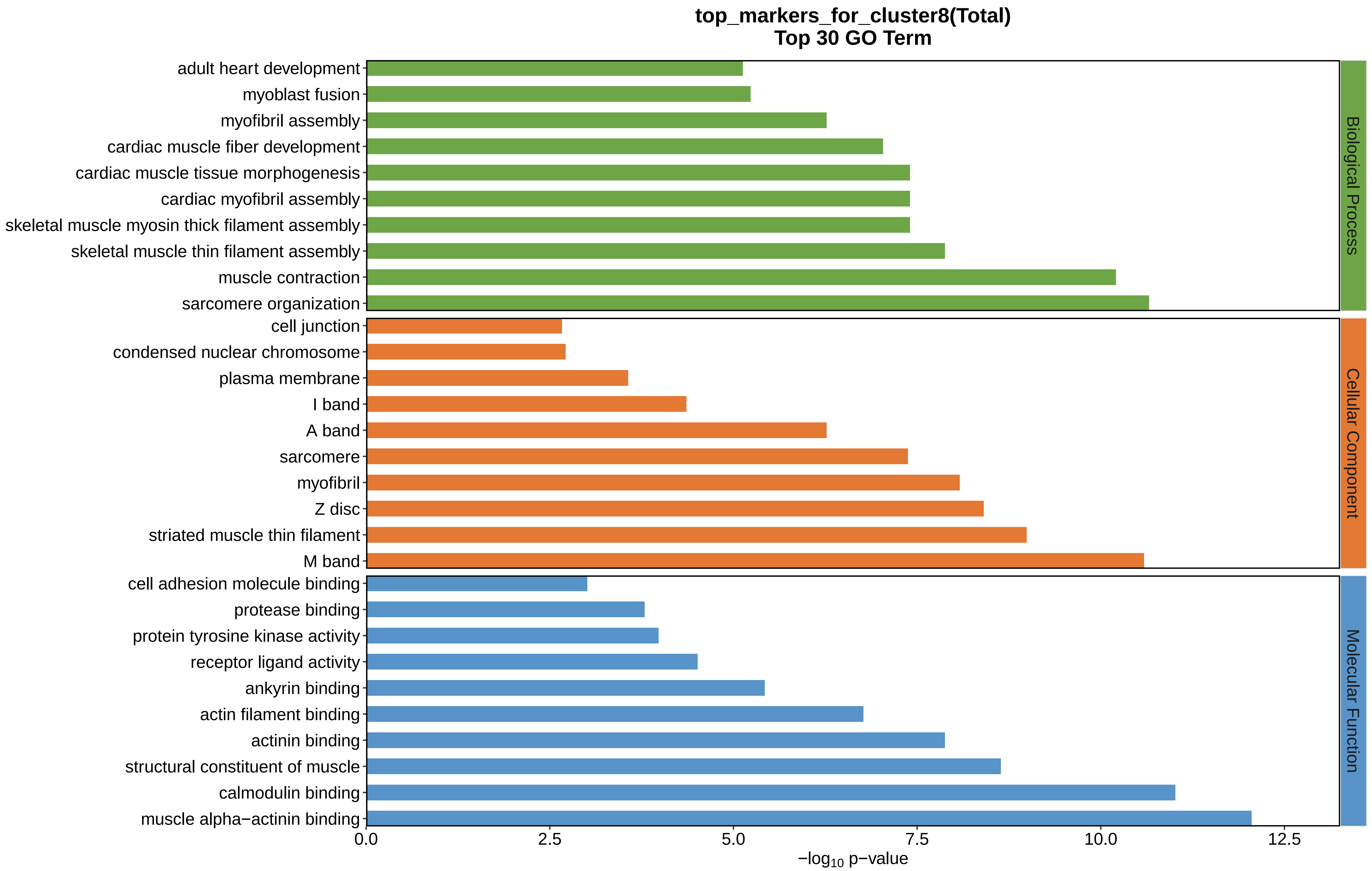

top_markers_for_cluster8(Total)
Top 30 GO Term
adult heart development
myoblast fusion
myofibril assembly
cardiac muscle fiber development
cardiac muscle tissue morphogenesis
cardiac myofibril assembly
skeletal muscle myosin thick filament assembly
skeletal muscle thin filament assembly
muscle contraction
sarcomere organization
cell junction
condensed nuclear chromosome
plasma membrane
I band
A band
sarcomere
myofibril
Z disc
striated muscle thin filament
M band
cell adhesion molecule binding
protease binding
protein tyrosine kinase activity
receptor ligand activity
ankyrin binding
actin filament binding
actinin binding
structural constituent of muscle
calmodulin binding
muscle alpha−actinin binding
Biological Process
Cellular Component
Molecular Function
0.0
2.5
5.0
7.5
10.0
12.5
$-\log_{10}$ p−value
