## Supplemental Data 1-8 for "A Spatiotemporal Embryonic Atlas of Bombyx mori Integrating Single-Nucleus RNA Sequencing and Spatial Transcriptomics": Cluster9 GO top100.pdf

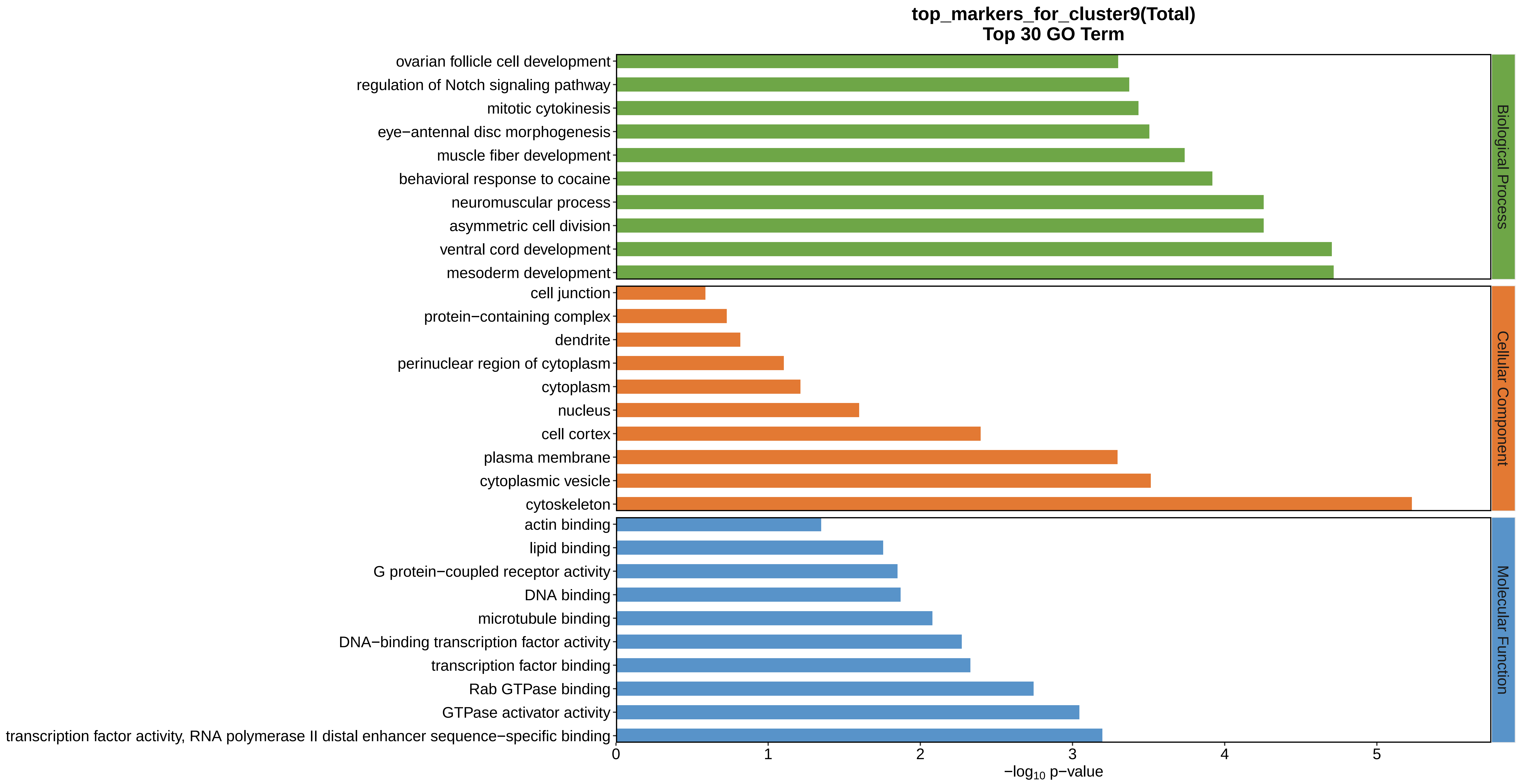
top_markers_for_cluster9(Total)
Top 30 GO Term
ovarian follicle cell development
regulation of Notch signaling pathway
mitotic cytokinesis
eye−antennal disc morphogenesis
muscle fiber development
behavioral response to cocaine
neuromuscular process
asymmetric cell division
ventral cord development
mesoderm development
cell junction
protein−containing complex
dendrite
perinuclear region of cytoplasm
cytoplasm
nucleus
cell cortex
plasma membrane
cytoplasmic vesicle
cytoskeleton
actin binding
lipid binding
G protein−coupled receptor activity
DNA binding
microtubule binding
DNA−binding transcription factor activity
transcription factor binding
Rab GTPase binding
GTPase activator activity
transcription factor activity, RNA polymerase II distal enhancer sequence−specific binding
Biological Process
Cellular Component
Molecular Function
0
1
2
3
4
5
−log10 p−value
