## Supplemental Data 1-8 for "A Spatiotemporal Embryonic Atlas of Bombyx mori Integrating Single-Nucleus RNA Sequencing and Spatial Transcriptomics": Cluster10 GO top100.pdf

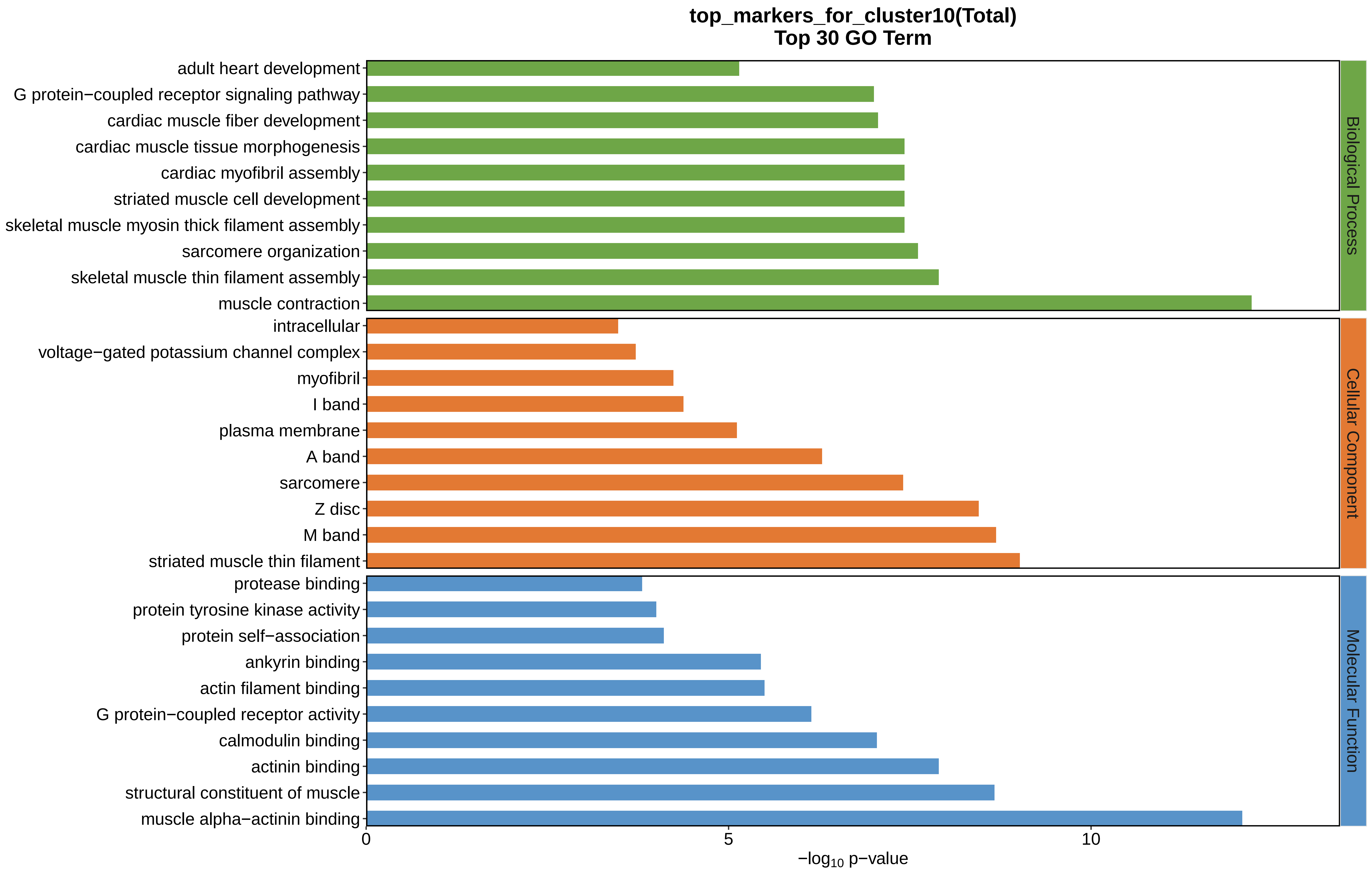
top_markers_for_cluster10(Total)
Top 30 GO Term
adult heart development
G protein−coupled receptor signaling pathway
cardiac muscle fiber development
cardiac muscle tissue morphogenesis
cardiac myofibril assembly
striated muscle cell development
skeletal muscle myosin thick filament assembly
sarcomere organization
skeletal muscle thin filament assembly
muscle contraction
intracellular
voltage−gated potassium channel complex
myofibril
I band
plasma membrane
A band
sarcomere
Z disc
M band
striated muscle thin filament
protease binding
protein tyrosine kinase activity
protein self−association
ankyrin binding
actin filament binding
G protein−coupled receptor activity
calmodulin binding
actinin binding
structural constituent of muscle
muscle alpha−actinin binding
Biological Process
Cellular Component
Molecular Function
0
5
10
$-\log_{10}$ p−value
