## Supplemental Data 1-8 for "A Spatiotemporal Embryonic Atlas of Bombyx mori Integrating Single-Nucleus RNA Sequencing and Spatial Transcriptomics": Cluster11 GO top100.pdf

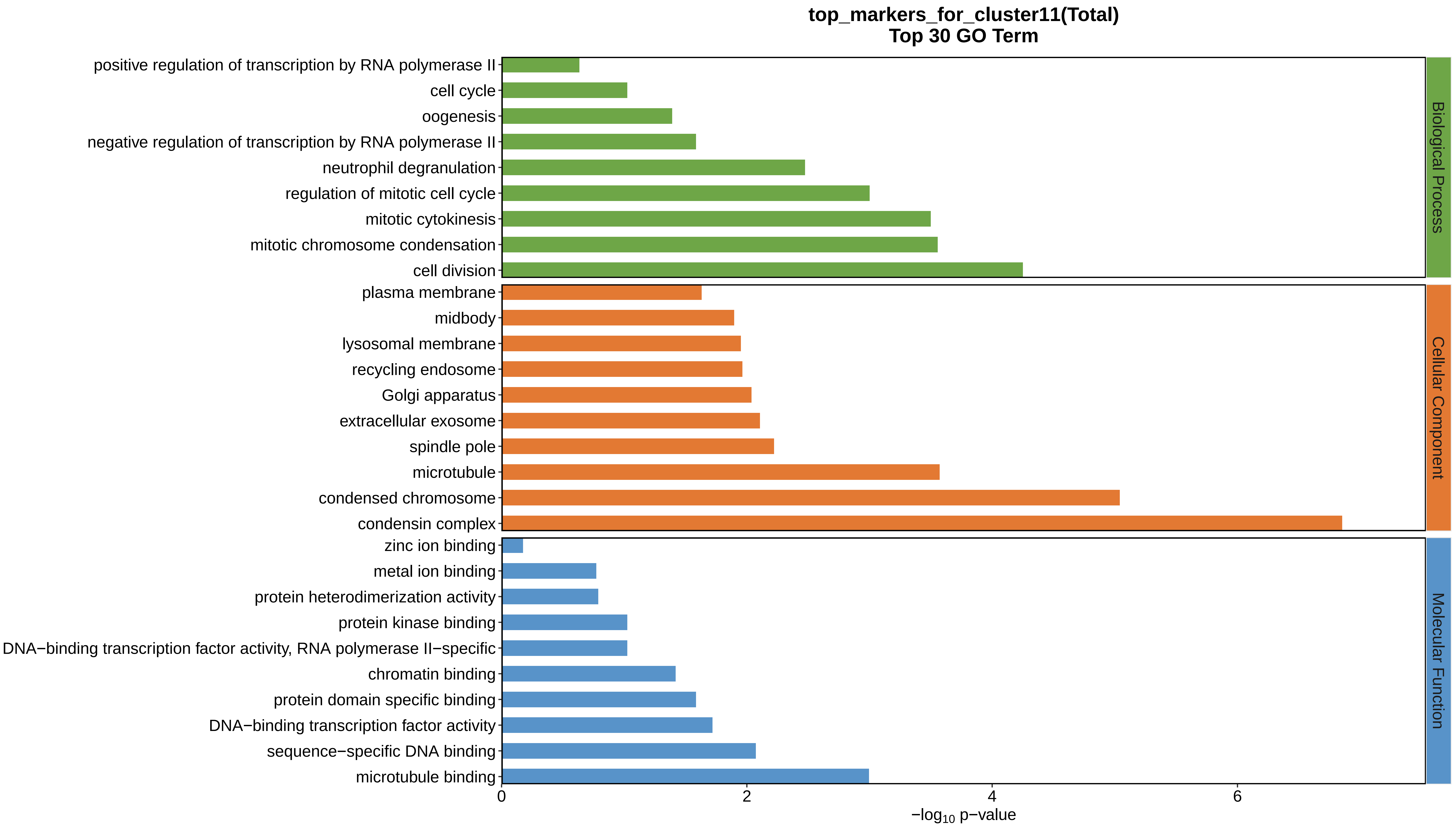
top_markers_for_cluster11(Total)
Top 30 GO Term
positive regulation of transcription by RNA polymerase II
cell cycle
oogenesis
negative regulation of transcription by RNA polymerase II
neutrophil degranulation
regulation of mitotic cell cycle
mitotic cytokinesis
mitotic chromosome condensation
cell division
plasma membrane
midbody
lysosomal membrane
recycling endosome
Golgi apparatus
extracellular exosome
spindle pole
microtubule
condensed chromosome
condensin complex
zinc ion binding
metal ion binding
protein heterodimerization activity
protein kinase binding
DNA−binding transcription factor activity, RNA polymerase II−specific
chromatin binding
protein domain specific binding
DNA−binding transcription factor activity
sequence−specific DNA binding
microtubule binding
Biological Process
Cellular Component
Molecular Function
0
2
4
6
−log10 p−value
