## Supplemental Data 1-8 for "A Spatiotemporal Embryonic Atlas of Bombyx mori Integrating Single-Nucleus RNA Sequencing and Spatial Transcriptomics": Cluster12 GO top100.pdf

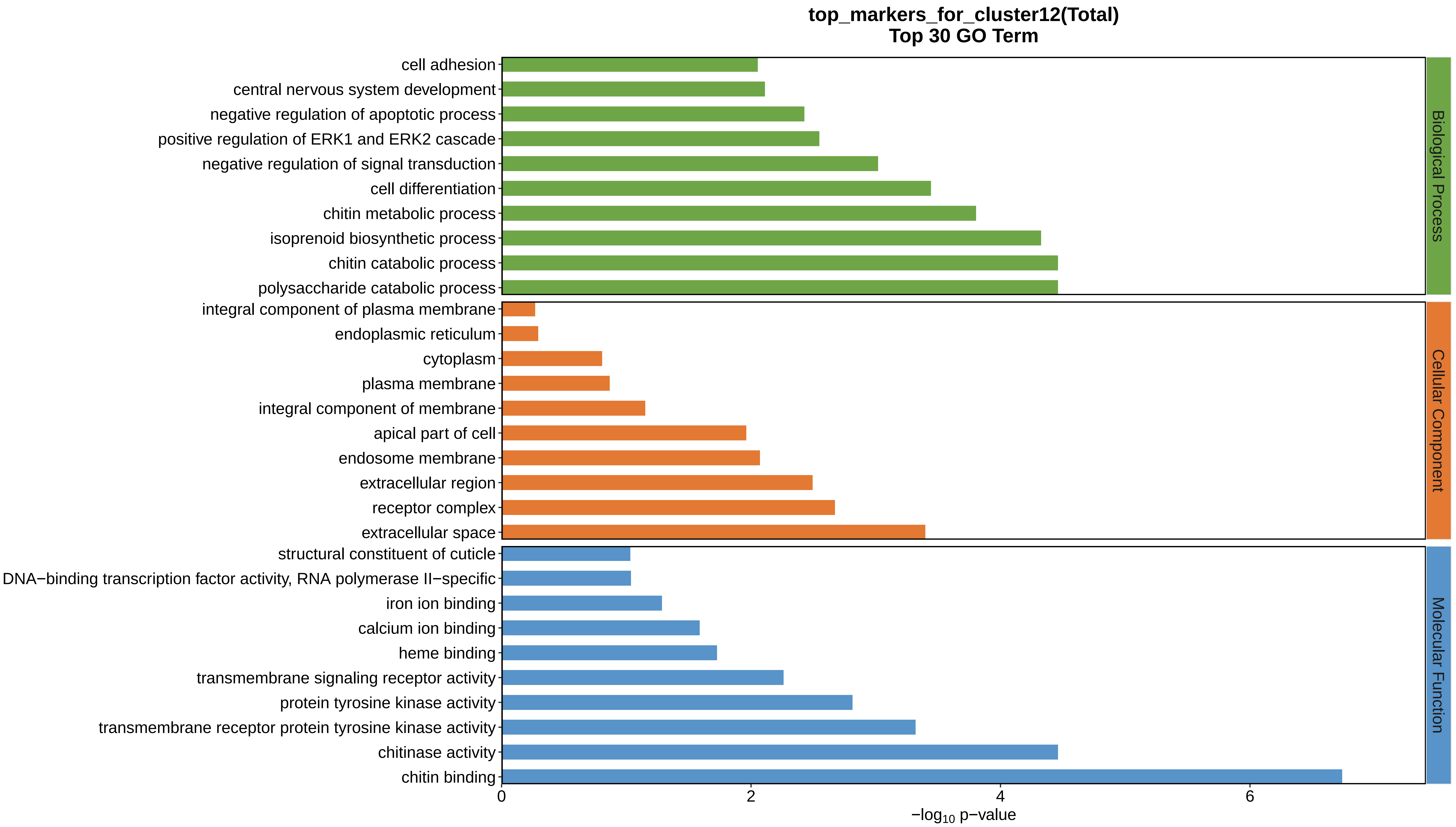
top_markers_for_cluster12(Total)
Top 30 GO Term
cell adhesion
central nervous system development
negative regulation of apoptotic process
positive regulation of ERK1 and ERK2 cascade
negative regulation of signal transduction
cell differentiation
chitin metabolic process
isoprenoid biosynthetic process
chitin catabolic process
polysaccharide catabolic process
integral component of plasma membrane
endoplasmic reticulum
cytoplasm
plasma membrane
integral component of membrane
apical part of cell
endosome membrane
extracellular region
receptor complex
extracellular space
structural constituent of cuticle
DNA−binding transcription factor activity, RNA polymerase II−specific
iron ion binding
calcium ion binding
heme binding
transmembrane signaling receptor activity
protein tyrosine kinase activity
transmembrane receptor protein tyrosine kinase activity
chitinase activity
chitin binding
0
2
4
6
−log10 p−value
Biological Process
Cellular Component
Molecular Function
