## Supplemental Data 1-8 for "A Spatiotemporal Embryonic Atlas of Bombyx mori Integrating Single-Nucleus RNA Sequencing and Spatial Transcriptomics": Cluster13 GO top100.pdf

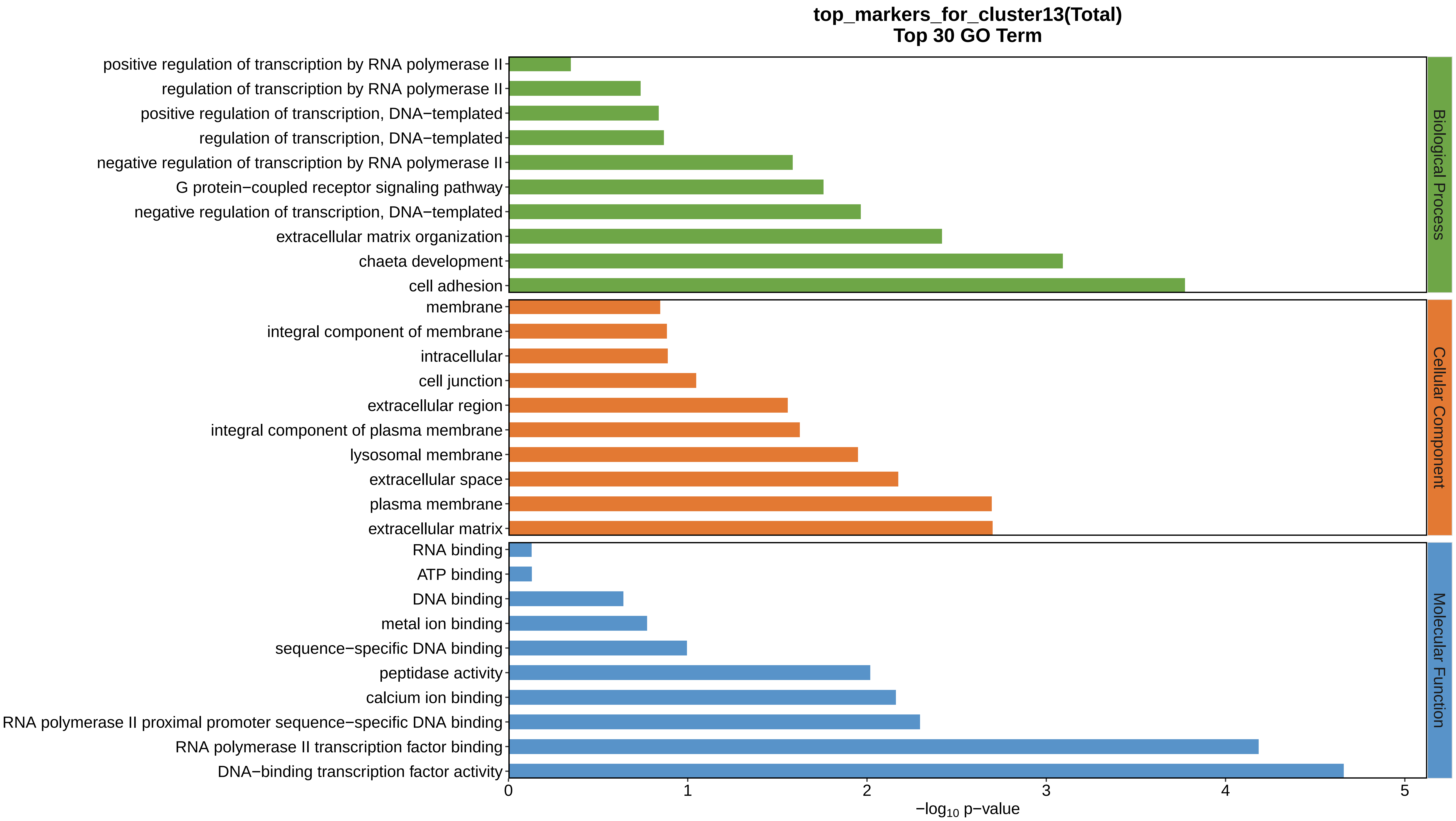

top_markers_for_cluster13(Total)
Top 30 GO Term
positive regulation of transcription by RNA polymerase II
regulation of transcription by RNA polymerase II
positive regulation of transcription, DNA−templated
regulation of transcription, DNA−templated
negative regulation of transcription by RNA polymerase II
G protein−coupled receptor signaling pathway
negative regulation of transcription, DNA−templated
extracellular matrix organization
chaeta development
cell adhesion
membrane
integral component of membrane
intracellular
cell junction
extracellular region
integral component of plasma membrane
lysosomal membrane
extracellular space
plasma membrane
extracellular matrix
RNA binding
ATP binding
DNA binding
metal ion binding
sequence−specific DNA binding
peptidase activity
calcium ion binding
RNA polymerase II proximal promoter sequence−specific DNA binding
RNA polymerase II transcription factor binding
DNA−binding transcription factor activity
Biological Process
Cellular Component
Molecular Function
0
1
2
3
4
5
−log10 p−value
