## Supplemental Data 1-8 for "A Spatiotemporal Embryonic Atlas of Bombyx mori Integrating Single-Nucleus RNA Sequencing and Spatial Transcriptomics": Cluster14 GO top100.pdf

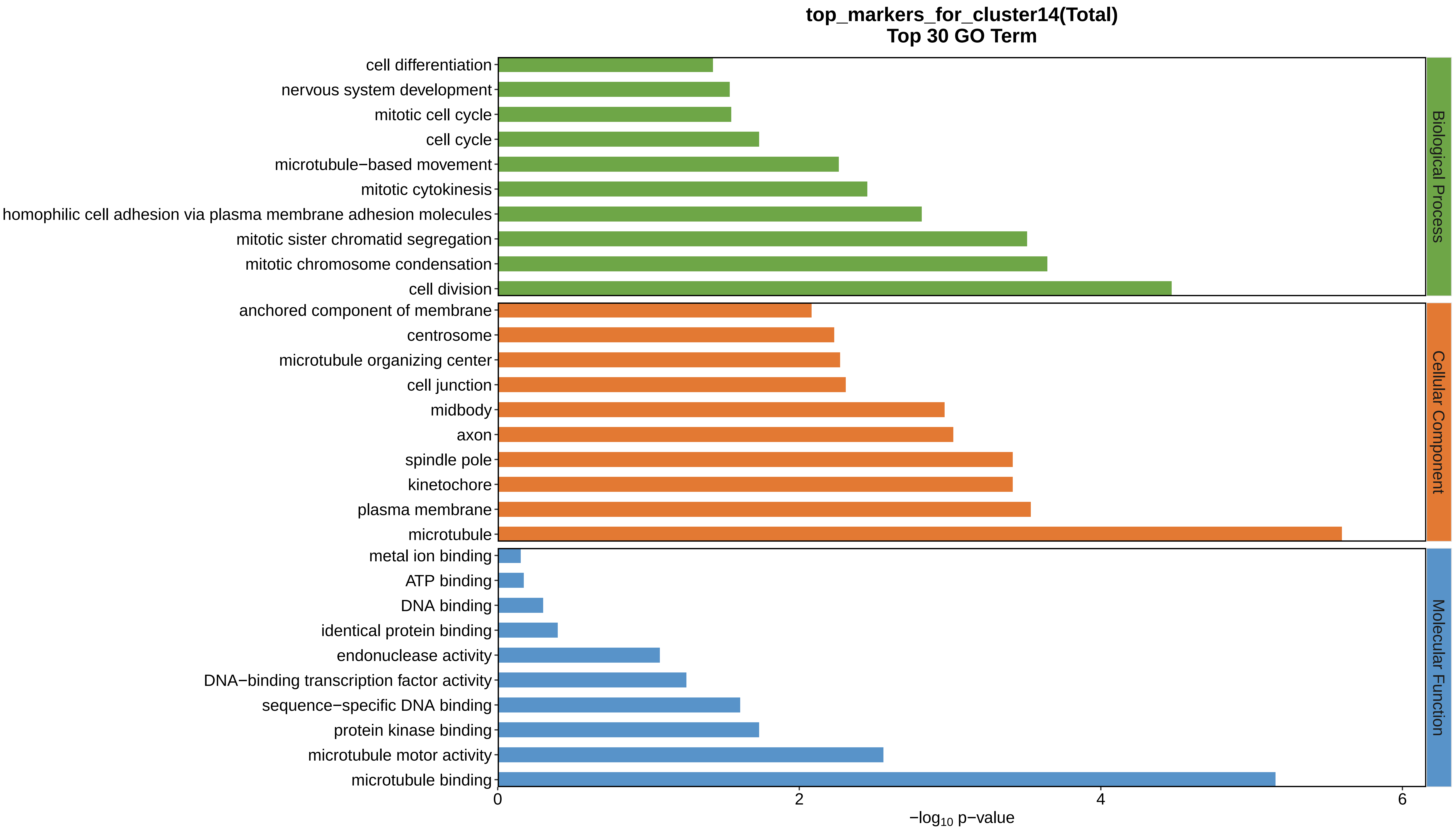
top_markers_for_cluster14(Total)
Top 30 GO Term
cell differentiation
nervous system development
mitotic cell cycle
cell cycle
microtubule−based movement
mitotic cytokinesis
homophilic cell adhesion via plasma membrane adhesion molecules
mitotic sister chromatid segregation
mitotic chromosome condensation
cell division
anchored component of membrane
centrosome
microtubule organizing center
cell junction
midbody
axon
spindle pole
kinetochore
plasma membrane
microtubule
metal ion binding
ATP binding
DNA binding
identical protein binding
endonuclease activity
DNA−binding transcription factor activity
sequence−specific DNA binding
protein kinase binding
microtubule motor activity
microtubule binding
Biological Process
Cellular Component
Molecular Function
0
2
4
6
−log10 p−value
