## Supplemental Data 1-8 for "A Spatiotemporal Embryonic Atlas of Bombyx mori Integrating Single-Nucleus RNA Sequencing and Spatial Transcriptomics": Cluster15 GO top100.pdf

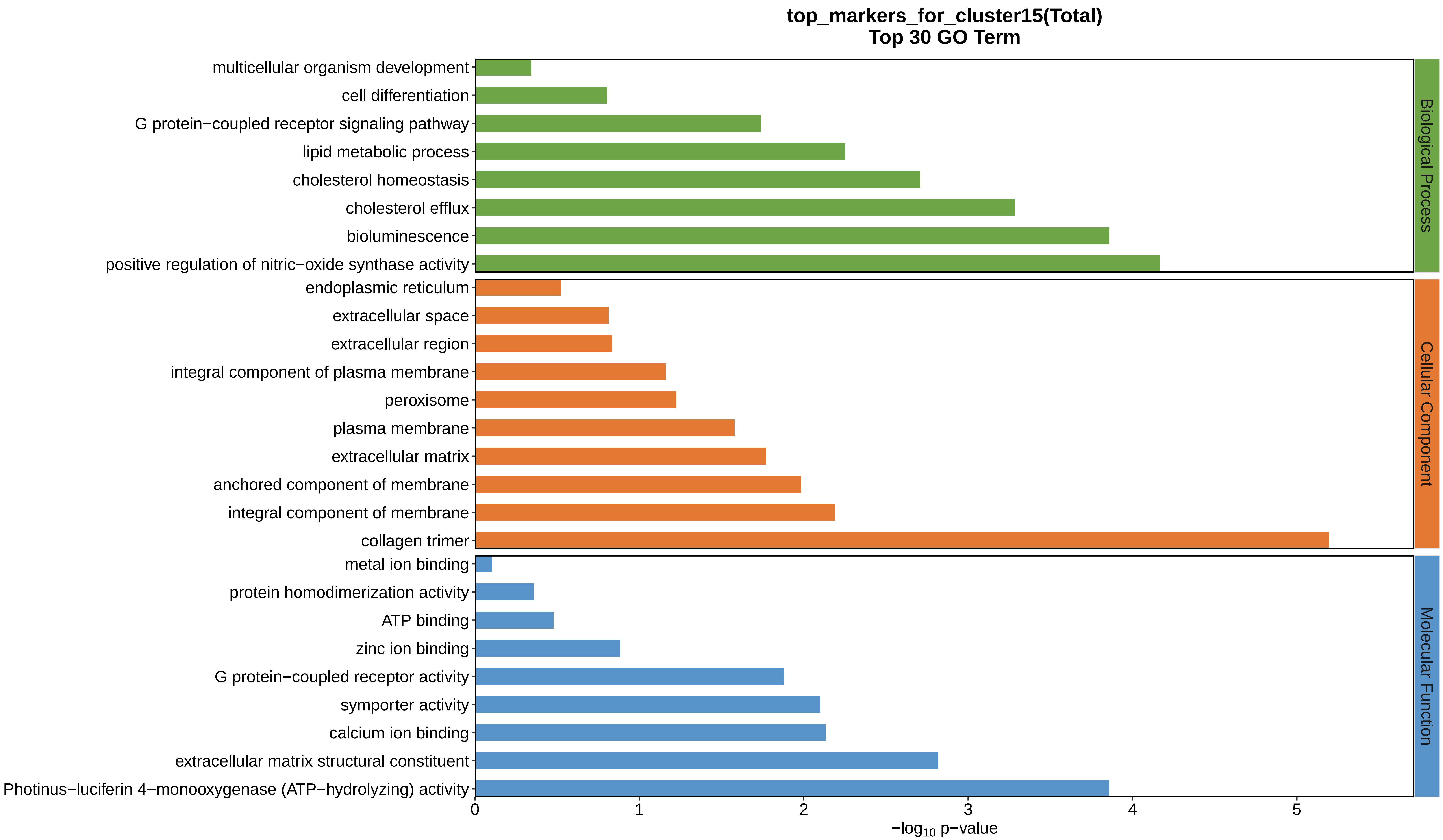
top_markers_for_cluster15(Total)
Top 30 GO Term
multicellular organism development
cell differentiation
G protein−coupled receptor signaling pathway
lipid metabolic process
cholesterol homeostasis
cholesterol efflux
bioluminescence
positive regulation of nitric−oxide synthase activity
endoplasmic reticulum
extracellular space
extracellular region
integral component of plasma membrane
peroxisome
plasma membrane
extracellular matrix
anchored component of membrane
integral component of membrane
collagen trimer
metal ion binding
protein homodimerization activity
ATP binding
zinc ion binding
G protein−coupled receptor activity
symporter activity
calcium ion binding
extracellular matrix structural constituent
Photinus−luciferin 4−monooxygenase (ATP−hydrolyzing) activity
0
1
2
3
4
5
$-\log_{10}$ p−value
Biological Process
Cellular Component
Molecular Function
