## Supplemental Data 1-8 for "A Spatiotemporal Embryonic Atlas of Bombyx mori Integrating Single-Nucleus RNA Sequencing and Spatial Transcriptomics": Cluster16 GO top100.pdf

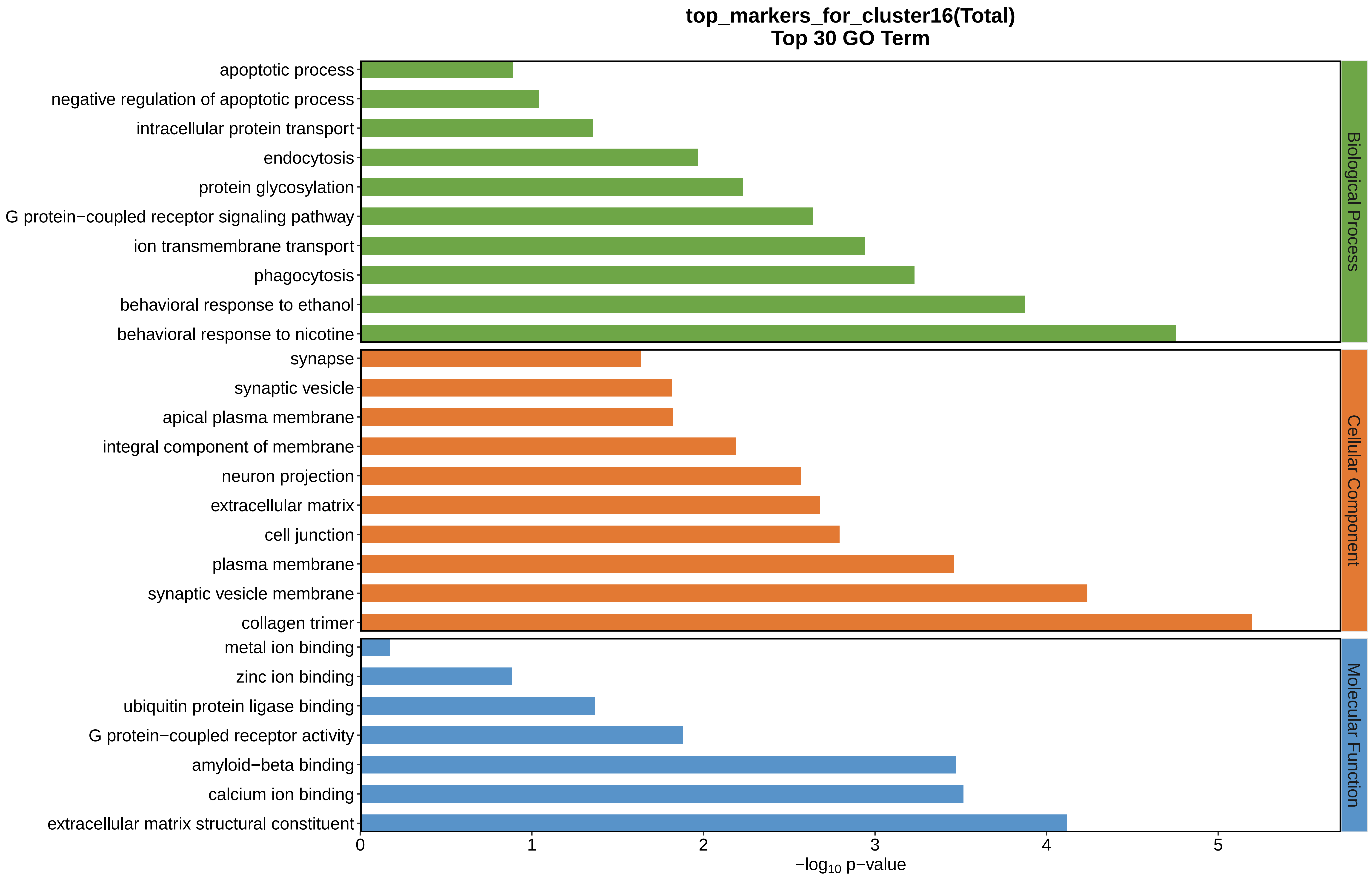
top_markers_for_cluster16(Total)
Top 30 GO Term
apoptotic process
negative regulation of apoptotic process
intracellular protein transport
endocytosis
protein glycosylation
G protein−coupled receptor signaling pathway
ion transmembrane transport
phagocytosis
behavioral response to ethanol
behavioral response to nicotine
synapse
synaptic vesicle
apical plasma membrane
integral component of membrane
neuron projection
extracellular matrix
cell junction
plasma membrane
synaptic vesicle membrane
collagen trimer
metal ion binding
zinc ion binding
ubiquitin protein ligase binding
G protein−coupled receptor activity
amyloid−beta binding
calcium ion binding
extracellular matrix structural constituent
Biological Process
Cellular Component
Molecular Function
0
1
2
3
4
5
−log10 p−value
