## Supplemental Data 1-8 for "A Spatiotemporal Embryonic Atlas of Bombyx mori Integrating Single-Nucleus RNA Sequencing and Spatial Transcriptomics": Cluster17 GO top100.pdf

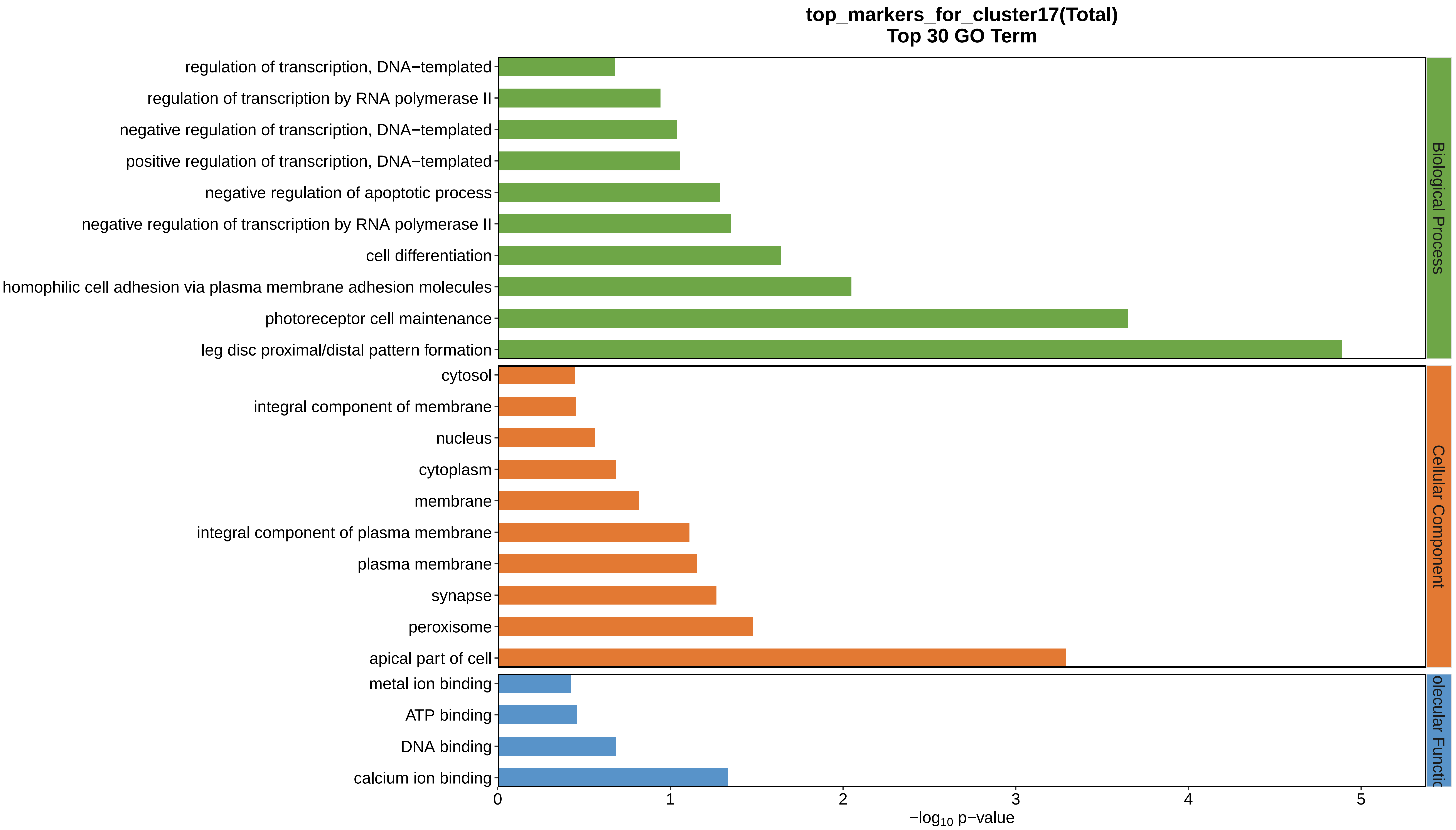
top_markers_for_cluster17(Total)
Top 30 GO Term
regulation of transcription, DNA−templated
regulation of transcription by RNA polymerase II
negative regulation of transcription, DNA−templated
positive regulation of transcription, DNA−templated
negative regulation of apoptotic process
negative regulation of transcription by RNA polymerase II
cell differentiation
homophilic cell adhesion via plasma membrane adhesion molecules
photoreceptor cell maintenance
leg disc proximal/distal pattern formation
cytosol
integral component of membrane
nucleus
cytoplasm
membrane
integral component of plasma membrane
plasma membrane
synapse
peroxisome
apical part of cell
metal ion binding
ATP binding
DNA binding
calcium ion binding
Biological Process
Cellular Component
olecular Functio
0
1
2
3
4
5
−log10 p−value
