## Supplemental Data 1-8 for "A Spatiotemporal Embryonic Atlas of Bombyx mori Integrating Single-Nucleus RNA Sequencing and Spatial Transcriptomics": Cluster18 GO top100.pdf

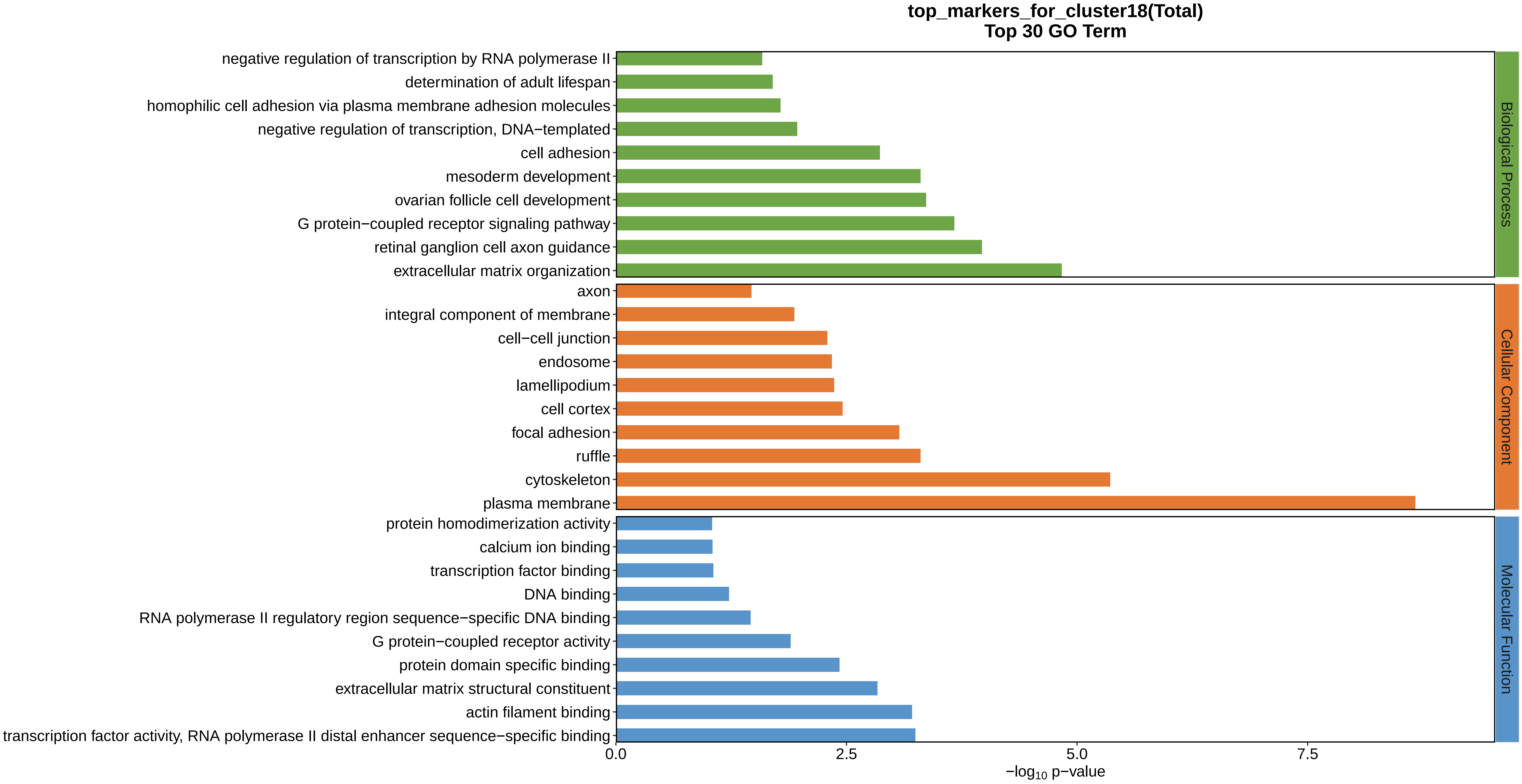
top_markers_for_cluster18(Total)
Top 30 GO Term
negative regulation of transcription by RNA polymerase II
determination of adult lifespan
homophilic cell adhesion via plasma membrane adhesion molecules
negative regulation of transcription, DNA−templated
cell adhesion
mesoderm development
ovarian follicle cell development
G protein−coupled receptor signaling pathway
retinal ganglion cell axon guidance
extracellular matrix organization
axon
integral component of membrane
cell−cell junction
endosome
lamellipodium
cell cortex
focal adhesion
ruffle
cytoskeleton
plasma membrane
protein homodimerization activity
calcium ion binding
transcription factor binding
DNA binding
RNA polymerase II regulatory region sequence−specific DNA binding
G protein−coupled receptor activity
protein domain specific binding
extracellular matrix structural constituent
actin filament binding
transcription factor activity, RNA polymerase II distal enhancer sequence−specific binding
Biological Process
Cellular Component
Molecular Function
0.0
2.5
5.0
7.5
$-\log_{10}$ p−value
