## Supplemental Data 1-8 for "A Spatiotemporal Embryonic Atlas of Bombyx mori Integrating Single-Nucleus RNA Sequencing and Spatial Transcriptomics": Cluster19 GO top100.pdf

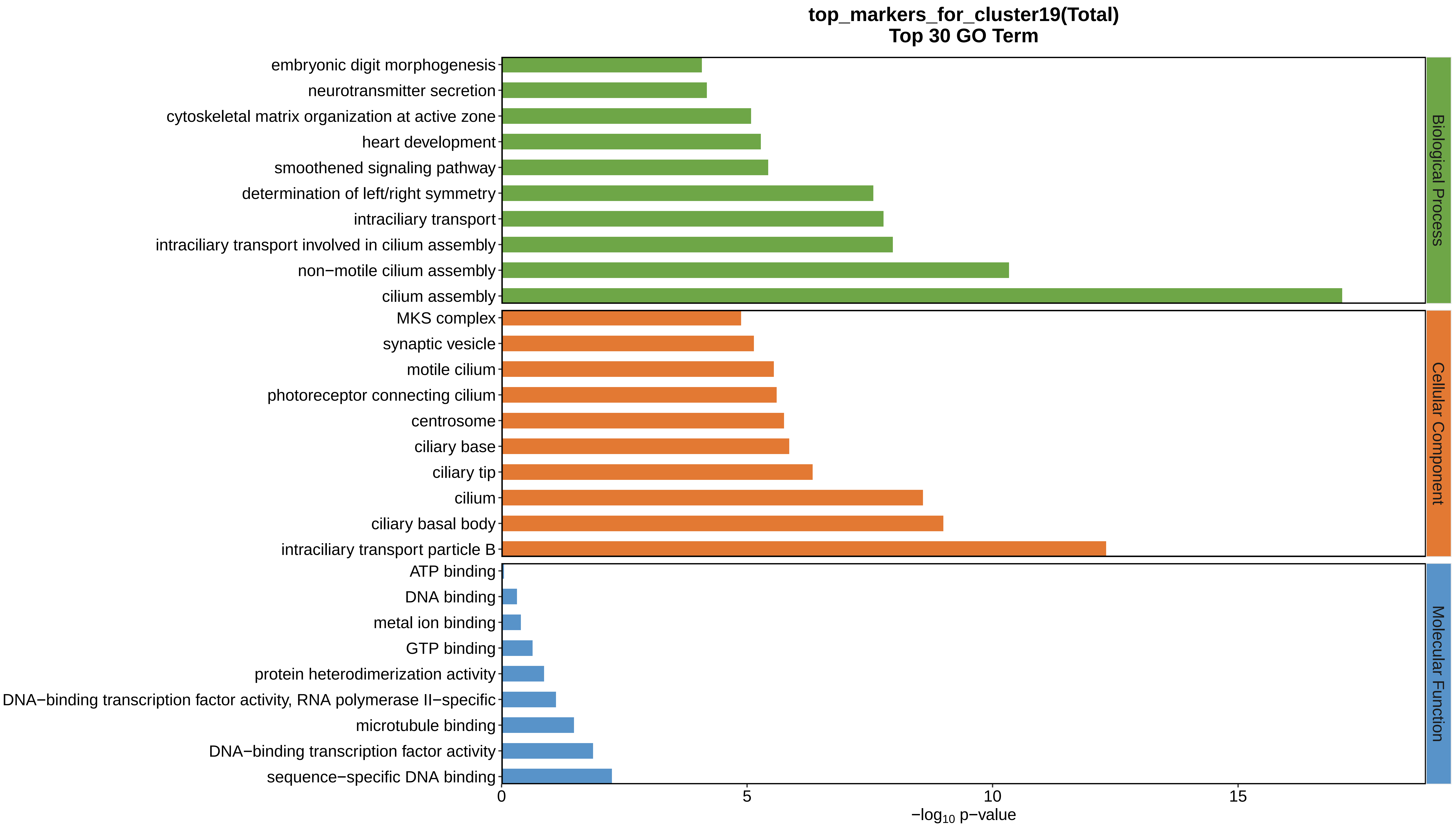
top_markers_for_cluster19(Total)
Top 30 GO Term
embryonic digit morphogenesis
neurotransmitter secretion
cytoskeletal matrix organization at active zone
heart development
smoothened signaling pathway
determination of left/right symmetry
intraciliary transport
intraciliary transport involved in cilium assembly
non−motile cilium assembly
cilium assembly
MKS complex
synaptic vesicle
motile cilium
photoreceptor connecting cilium
centrosome
ciliary base
ciliary tip
cilium
ciliary basal body
intraciliary transport particle B
ATP binding
DNA binding
metal ion binding
GTP binding
protein heterodimerization activity
DNA−binding transcription factor activity, RNA polymerase II−specific
microtubule binding
DNA−binding transcription factor activity
sequence−specific DNA binding
0
5
10
15
−log10 p−value
Biological Process
Cellular Component
Molecular Function
