## Supplemental Data 1-8 for "A Spatiotemporal Embryonic Atlas of Bombyx mori Integrating Single-Nucleus RNA Sequencing and Spatial Transcriptomics": Cluster20 GO top100.pdf

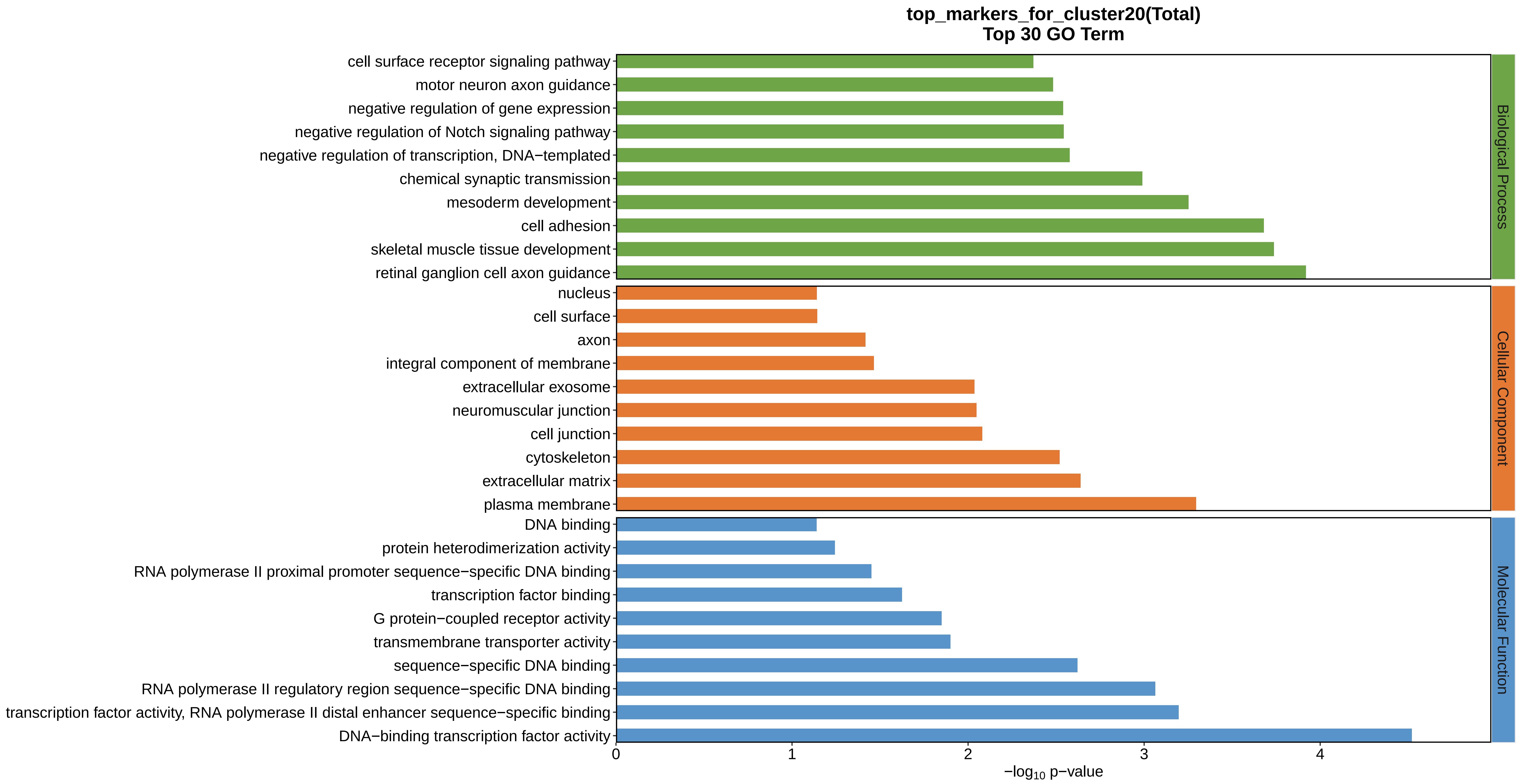
top_markers_for_cluster20(Total)
Top 30 GO Term
cell surface receptor signaling pathway
motor neuron axon guidance
negative regulation of gene expression
negative regulation of Notch signaling pathway
negative regulation of transcription, DNA−templated
chemical synaptic transmission
mesoderm development
cell adhesion
skeletal muscle tissue development
retinal ganglion cell axon guidance
nucleus
cell surface
axon
integral component of membrane
extracellular exosome
neuromuscular junction
cell junction
cytoskeleton
extracellular matrix
plasma membrane
DNA binding
protein heterodimerization activity
RNA polymerase II proximal promoter sequence−specific DNA binding
transcription factor binding
G protein−coupled receptor activity
transmembrane transporter activity
sequence−specific DNA binding
RNA polymerase II regulatory region sequence−specific DNA binding
transcription factor activity, RNA polymerase II distal enhancer sequence−specific binding
DNA−binding transcription factor activity
Biological Process
Cellular Component
Molecular Function
0
1
2
3
4
−log10 p−value
