## Supplemental Data 1-8 for "A Spatiotemporal Embryonic Atlas of Bombyx mori Integrating Single-Nucleus RNA Sequencing and Spatial Transcriptomics": Cluster21 GO top100.pdf

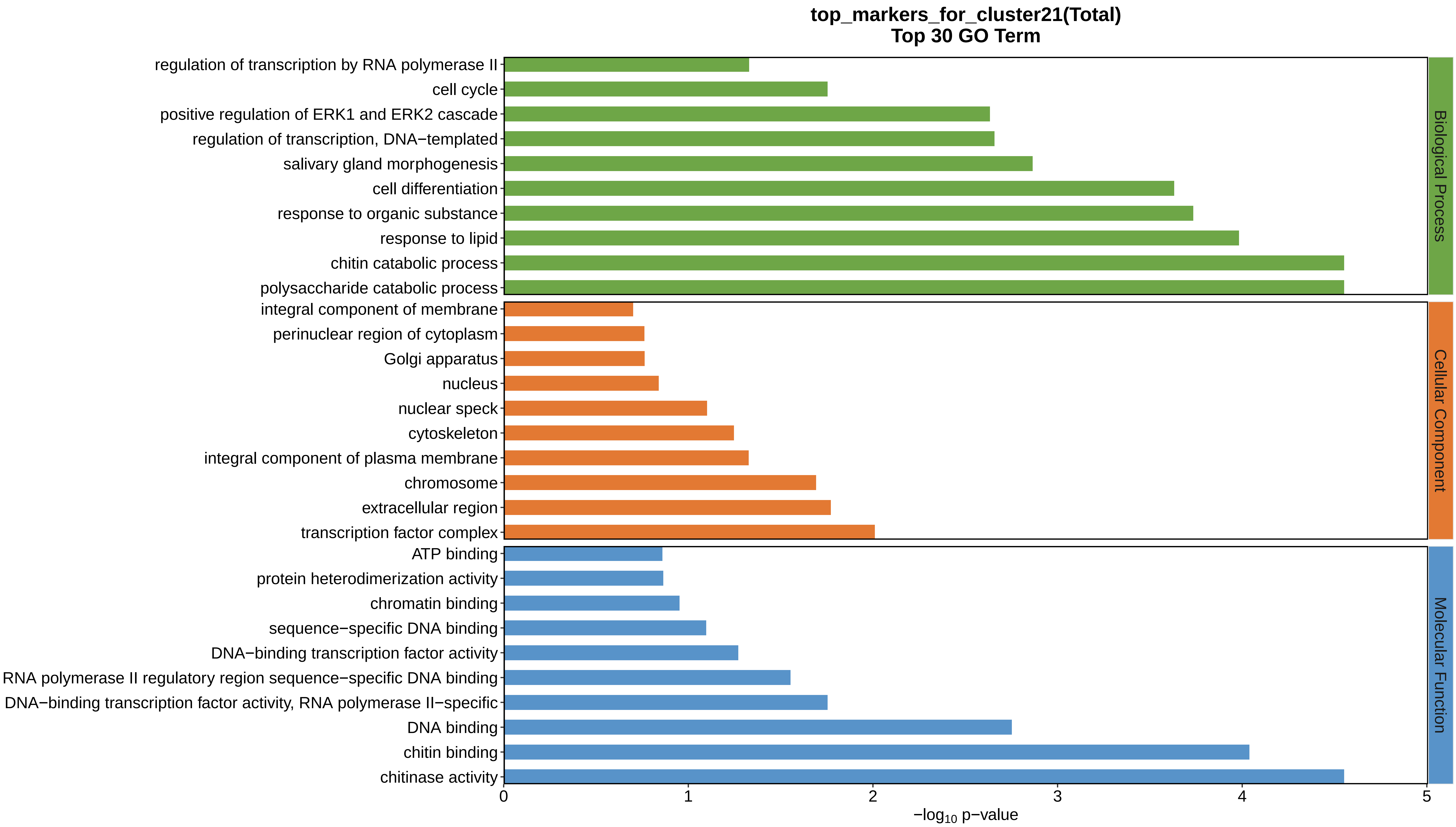
top_markers_for_cluster21(Total)
Top 30 GO Term
regulation of transcription by RNA polymerase II
cell cycle
positive regulation of ERK1 and ERK2 cascade
regulation of transcription, DNA−templated
salivary gland morphogenesis
cell differentiation
response to organic substance
response to lipid
chitin catabolic process
polysaccharide catabolic process
integral component of membrane
perinuclear region of cytoplasm
Golgi apparatus
nucleus
nuclear speck
cytoskeleton
integral component of plasma membrane
chromosome
extracellular region
transcription factor complex
ATP binding
protein heterodimerization activity
chromatin binding
sequence−specific DNA binding
DNA−binding transcription factor activity
RNA polymerase II regulatory region sequence−specific DNA binding
DNA−binding transcription factor activity, RNA polymerase II−specific
DNA binding
chitin binding
chitinase activity
Biological Process
Cellular Component
Molecular Function
0
1
2
3
4
5
−log10 p−value
