## Supplemental Data 1-8 for "A Spatiotemporal Embryonic Atlas of Bombyx mori Integrating Single-Nucleus RNA Sequencing and Spatial Transcriptomics": Cluster22 GO top100.pdf

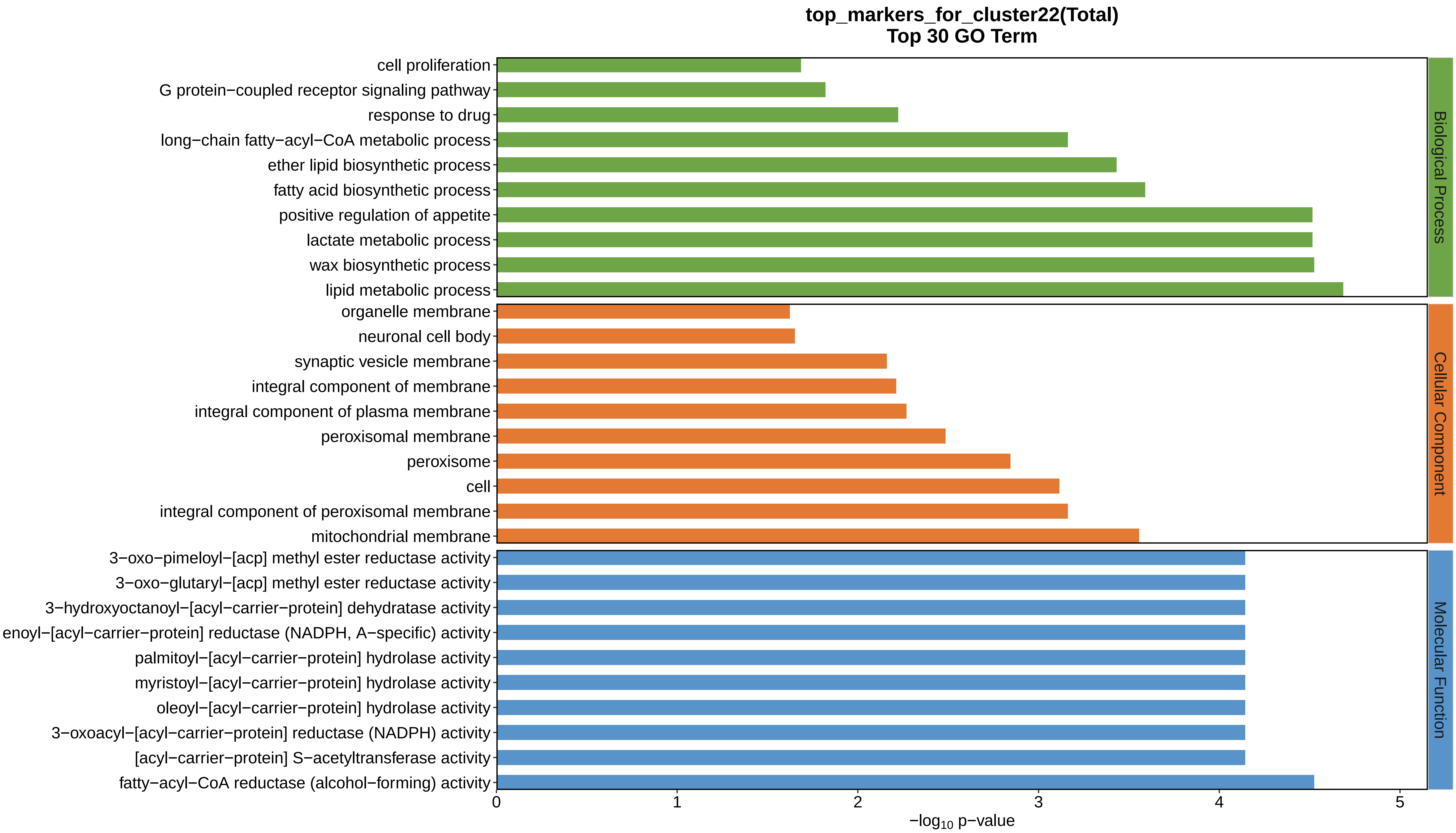
top_markers_for_cluster22(Total)
Top 30 GO Term
cell proliferation
G protein−coupled receptor signaling pathway
response to drug
long−chain fatty−acyl−CoA metabolic process
ether lipid biosynthetic process
fatty acid biosynthetic process
positive regulation of appetite
lactate metabolic process
wax biosynthetic process
lipid metabolic process
organelle membrane
neuronal cell body
synaptic vesicle membrane
integral component of membrane
integral component of plasma membrane
peroxisomal membrane
peroxisome
cell
integral component of peroxisomal membrane
mitochondrial membrane
3−oxo−pimeloyl−[acp] methyl ester reductase activity
3−oxo−glutaryl−[acp] methyl ester reductase activity
3−hydroxyoctanoyl−[acyl−carrier−protein] dehydratase activity
enoyl−[acyl−carrier−protein] reductase (NADPH, A−specific) activity
palmitoyl−[acyl−carrier−protein] hydrolase activity
myristoyl−[acyl−carrier−protein] hydrolase activity
oleoyl−[acyl−carrier−protein] hydrolase activity
3−oxoacyl−[acyl−carrier−protein] reductase (NADPH) activity
[acyl−carrier−protein] S−acetyltransferase activity
fatty−acyl−CoA reductase (alcohol−forming) activity
Biological Process
Cellular Component
Molecular Function
0
1
2
3
4
5
$-\log_{10}$ p−value
