## Supplemental Data 1-8 for "A Spatiotemporal Embryonic Atlas of Bombyx mori Integrating Single-Nucleus RNA Sequencing and Spatial Transcriptomics": Cluster1 GO top100.pdf

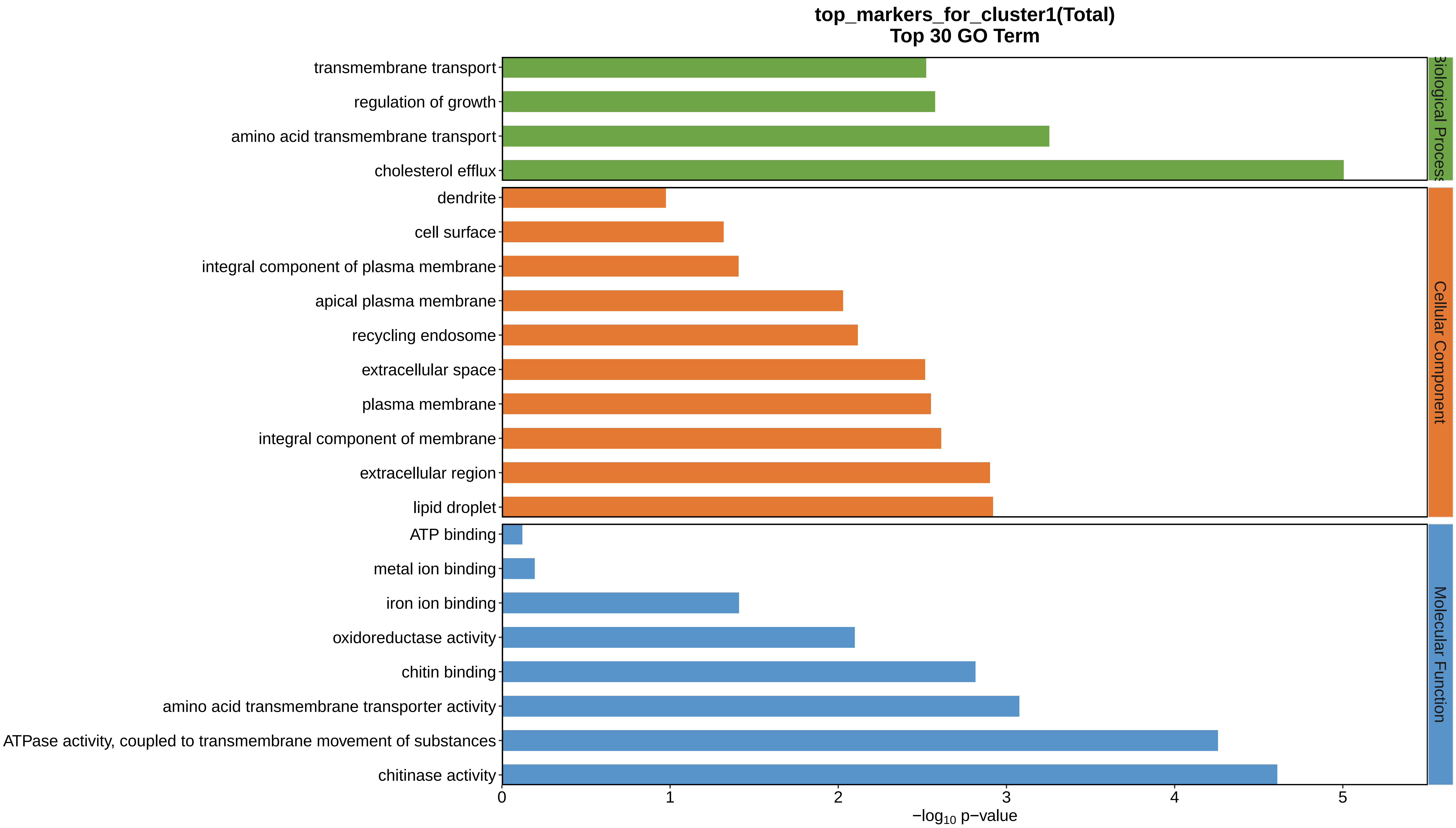
top_markers_for_cluster1(Total)
Top 30 GO Term
transmembrane transport
regulation of growth
amino acid transmembrane transport
cholesterol efflux
dendrite
cell surface
integral component of plasma membrane
apical plasma membrane
recycling endosome
extracellular space
plasma membrane
integral component of membrane
extracellular region
lipid droplet
ATP binding
metal ion binding
iron ion binding
oxidoreductase activity
chitin binding
amino acid transmembrane transporter activity
ATPase activity, coupled to transmembrane movement of substances
chitinase activity
Biological Process
Cellular Component
Molecular Function
0
1
2
3
4
5
−log10 p−value
