## Supplemental Data 1-8 for "A Spatiotemporal Embryonic Atlas of Bombyx mori Integrating Single-Nucleus RNA Sequencing and Spatial Transcriptomics": Cluster2 GO top100.pdf

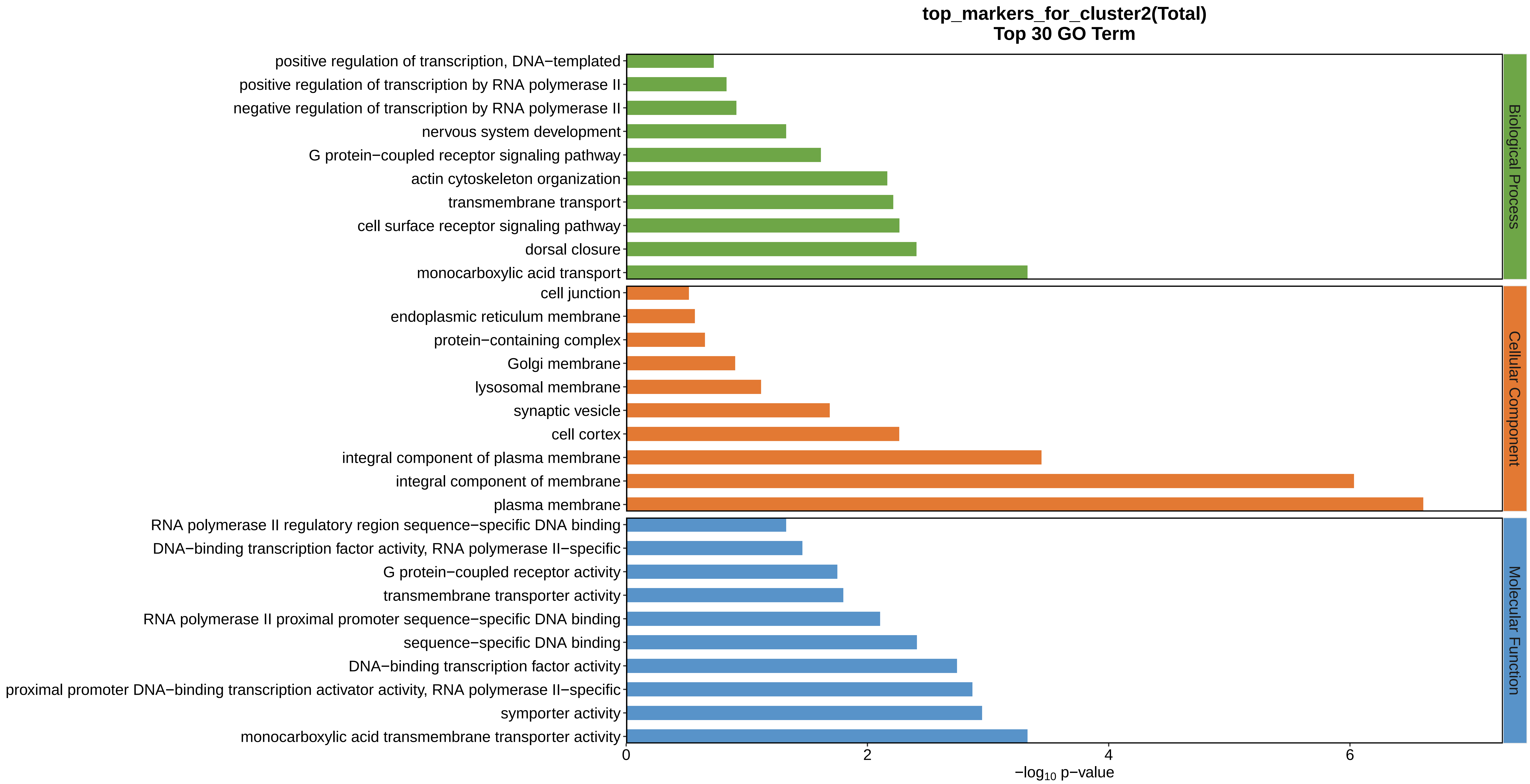

top_markers_for_cluster2(Total)
Top 30 GO Term
positive regulation of transcription, DNA−templated
positive regulation of transcription by RNA polymerase II
negative regulation of transcription by RNA polymerase II
nervous system development
G protein−coupled receptor signaling pathway
actin cytoskeleton organization
transmembrane transport
cell surface receptor signaling pathway
dorsal closure
monocarboxylic acid transport
cell junction
endoplasmic reticulum membrane
protein−containing complex
Golgi membrane
lysosomal membrane
synaptic vesicle
cell cortex
integral component of plasma membrane
integral component of membrane
plasma membrane
RNA polymerase II regulatory region sequence−specific DNA binding
DNA−binding transcription factor activity, RNA polymerase II−specific
G protein−coupled receptor activity
transmembrane transporter activity
RNA polymerase II proximal promoter sequence−specific DNA binding
sequence−specific DNA binding
DNA−binding transcription factor activity
proximal promoter DNA−binding transcription activator activity, RNA polymerase II−specific
symporter activity
monocarboxylic acid transmembrane transporter activity
Biological Process
Cellular Component
Molecular Function
0
2
4
6
$-\log_{10}$ p−value
