## Supplemental Data 1-8 for "A Spatiotemporal Embryonic Atlas of Bombyx mori Integrating Single-Nucleus RNA Sequencing and Spatial Transcriptomics": Cluster3 GO top100.pdf

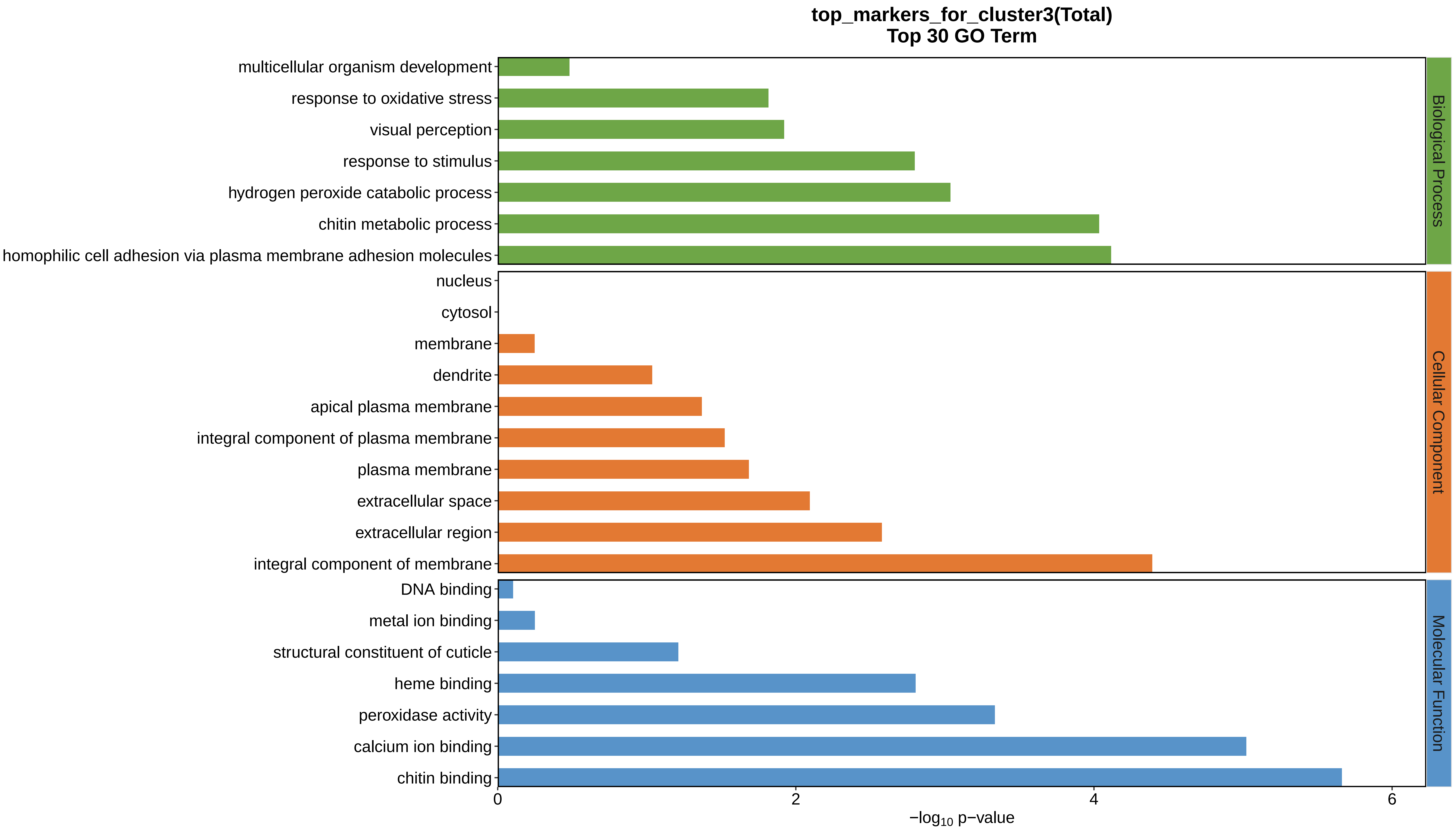
top_markers_for_cluster3(Total)
Top 30 GO Term
multicellular organism development
response to oxidative stress
visual perception
response to stimulus
hydrogen peroxide catabolic process
chitin metabolic process
homophilic cell adhesion via plasma membrane adhesion molecules
nucleus
cytosol
membrane
dendrite
apical plasma membrane
integral component of plasma membrane
plasma membrane
extracellular space
extracellular region
integral component of membrane
DNA binding
metal ion binding
structural constituent of cuticle
heme binding
peroxidase activity
calcium ion binding
chitin binding
Biological Process
Cellular Component
Molecular Function
0
2
4
6
−log10 p−value
