## Supplemental Data 1-8 for "A Spatiotemporal Embryonic Atlas of Bombyx mori Integrating Single-Nucleus RNA Sequencing and Spatial Transcriptomics": Cluster4 GO top100.pdf

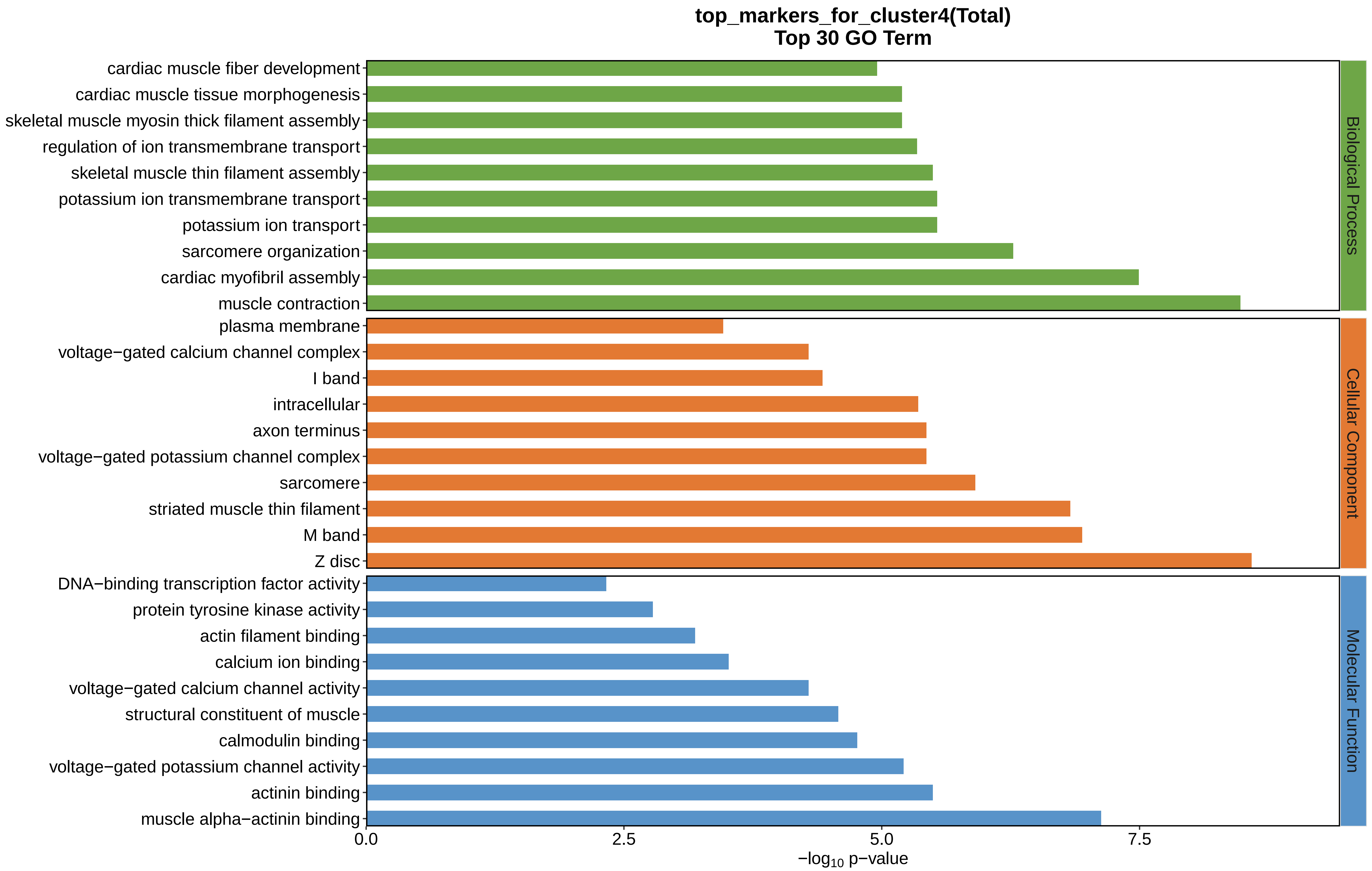

top_markers_for_cluster4(Total)
Top 30 GO Term
cardiac muscle fiber development
cardiac muscle tissue morphogenesis
skeletal muscle myosin thick filament assembly
regulation of ion transmembrane transport
skeletal muscle thin filament assembly
potassium ion transmembrane transport
potassium ion transport
sarcomere organization
cardiac myofibril assembly
muscle contraction
plasma membrane
voltage−gated calcium channel complex
I band
intracellular
axon terminus
voltage−gated potassium channel complex
sarcomere
striated muscle thin filament
M band
Z disc
DNA−binding transcription factor activity
protein tyrosine kinase activity
actin filament binding
calcium ion binding
voltage−gated calcium channel activity
structural constituent of muscle
calmodulin binding
voltage−gated potassium channel activity
actinin binding
muscle alpha−actinin binding
Biological Process
Cellular Component
Molecular Function
0.0
2.5
5.0
7.5
$-\log_{10}$ p−value
