## Supplemental Data 1-8 for "A Spatiotemporal Embryonic Atlas of Bombyx mori Integrating Single-Nucleus RNA Sequencing and Spatial Transcriptomics": Cluster5 GO top100.pdf

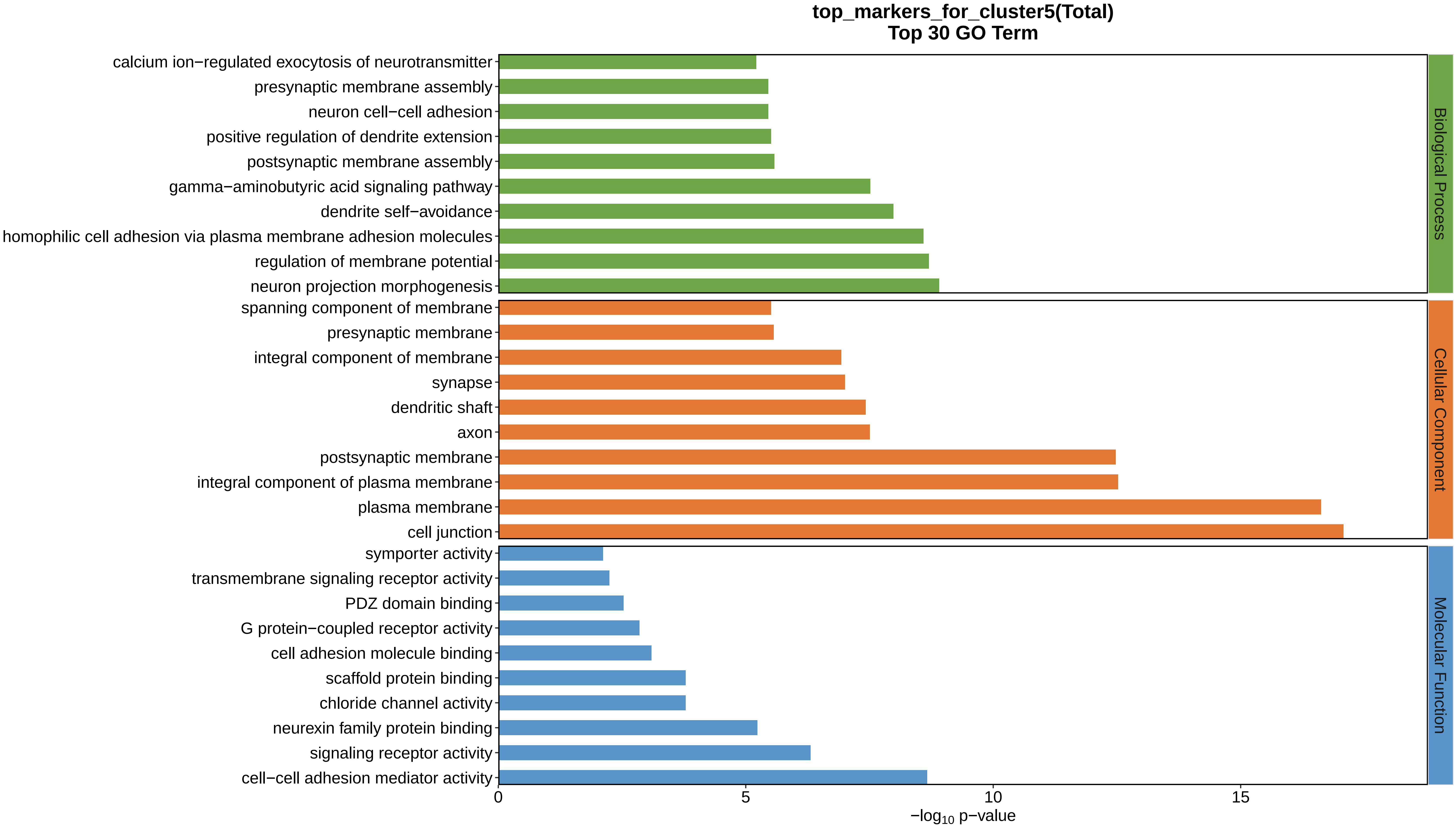
top_markers_for_cluster5(Total)
Top 30 GO Term
calcium ion−regulated exocytosis of neurotransmitter
presynaptic membrane assembly
neuron cell−cell adhesion
positive regulation of dendrite extension
postsynaptic membrane assembly
gamma−aminobutyric acid signaling pathway
dendrite self−avoidance
homophilic cell adhesion via plasma membrane adhesion molecules
regulation of membrane potential
neuron projection morphogenesis
spanning component of membrane
presynaptic membrane
integral component of membrane
synapse
dendritic shaft
axon
postsynaptic membrane
integral component of plasma membrane
plasma membrane
cell junction
symporter activity
transmembrane signaling receptor activity
PDZ domain binding
G protein−coupled receptor activity
cell adhesion molecule binding
scaffold protein binding
chloride channel activity
neurexin family protein binding
signaling receptor activity
cell−cell adhesion mediator activity
Biological Process
Cellular Component
Molecular Function
0
5
10
15
−log10 p−value
