## Supplemental Data 1-8 for "A Spatiotemporal Embryonic Atlas of Bombyx mori Integrating Single-Nucleus RNA Sequencing and Spatial Transcriptomics": Cluster6 GO top100.pdf

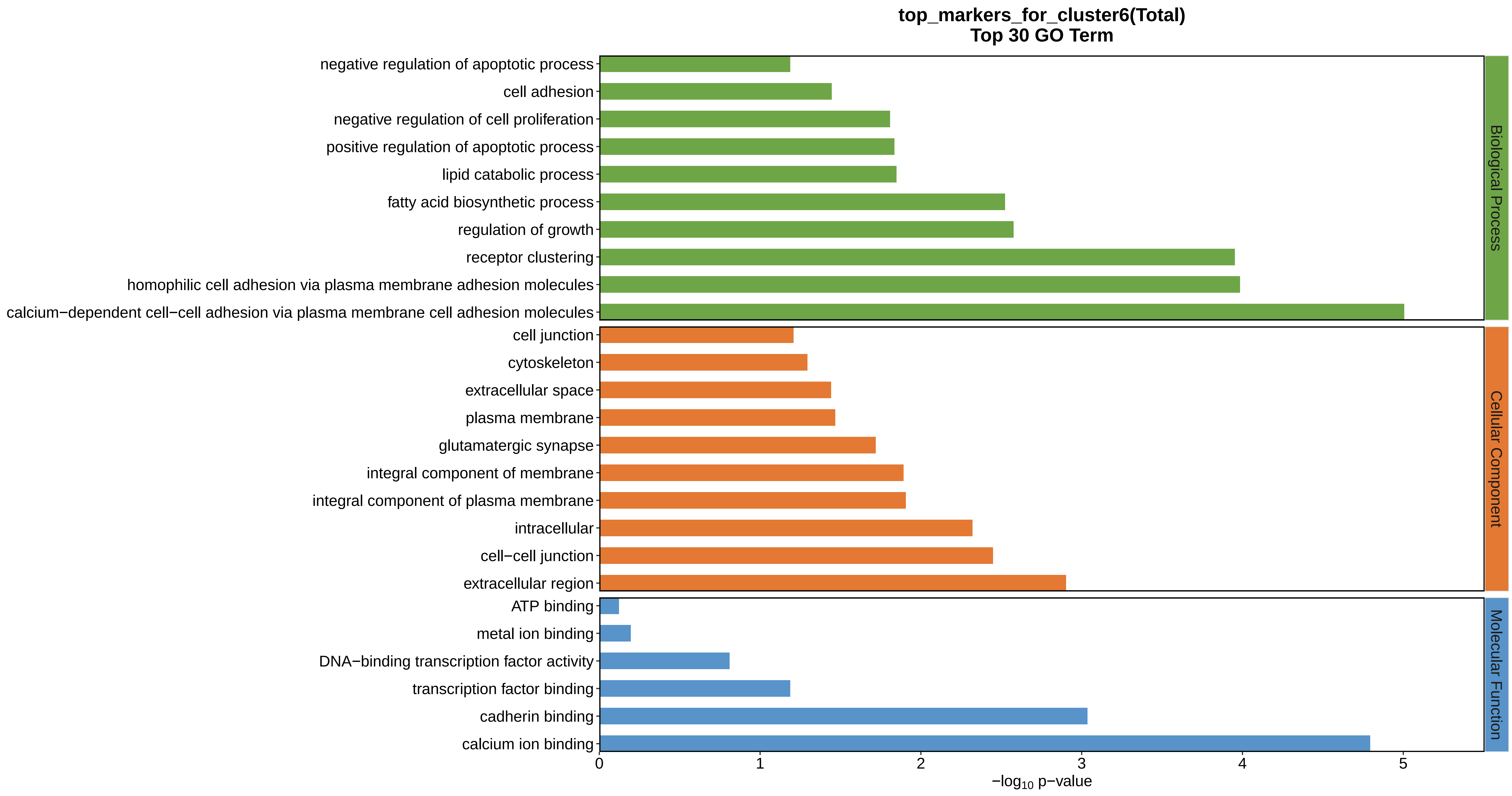
top_markers_for_cluster6(Total)
Top 30 GO Term
negative regulation of apoptotic process
cell adhesion
negative regulation of cell proliferation
positive regulation of apoptotic process
lipid catabolic process
fatty acid biosynthetic process
regulation of growth
receptor clustering
homophilic cell adhesion via plasma membrane adhesion molecules
calcium−dependent cell−cell adhesion via plasma membrane cell adhesion molecules
cell junction
cytoskeleton
extracellular space
plasma membrane
glutamatergic synapse
integral component of membrane
integral component of plasma membrane
intracellular
cell−cell junction
extracellular region
ATP binding
metal ion binding
DNA−binding transcription factor activity
transcription factor binding
cadherin binding
calcium ion binding
Biological Process
Cellular Component
Molecular Function
0
1
2
3
4
5
−log10 p−value
