## Supplemental Data 1-8 for "A Spatiotemporal Embryonic Atlas of Bombyx mori Integrating Single-Nucleus RNA Sequencing and Spatial Transcriptomics": Cluster7 GO top100.pdf

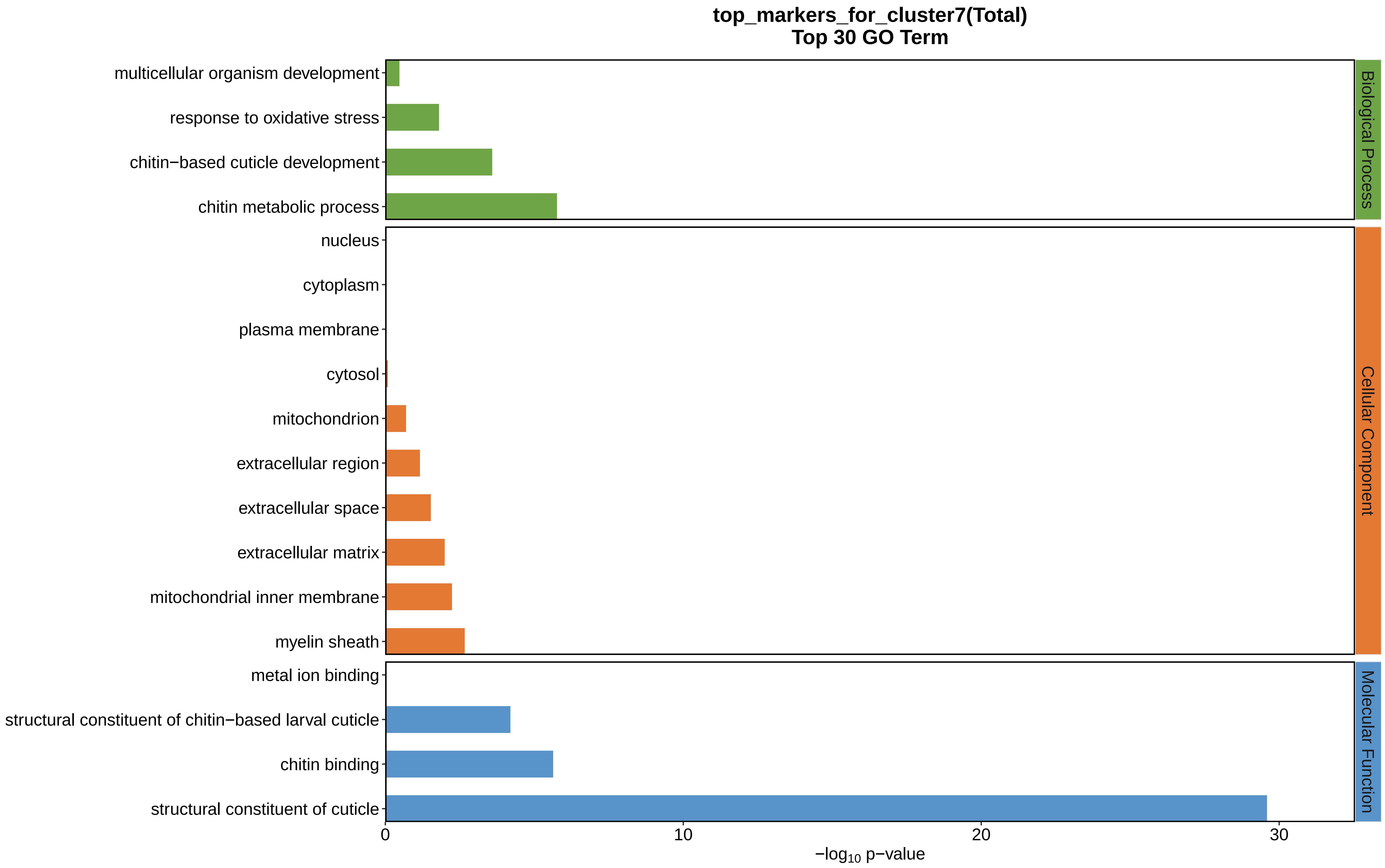
top_markers_for_cluster7(Total)
Top 30 GO Term
multicellular organism development
response to oxidative stress
chitin−based cuticle development
chitin metabolic process
nucleus
cytoplasm
plasma membrane
cytosol
mitochondrion
extracellular region
extracellular space
extracellular matrix
mitochondrial inner membrane
myelin sheath
metal ion binding
structural constituent of chitin−based larval cuticle
chitin binding
structural constituent of cuticle
Biological Process
Cellular Component
Molecular Function
0
10
20
30
−log10 p−value
