## Supplemental Data 1-8 for "A Spatiotemporal Embryonic Atlas of Bombyx mori Integrating Single-Nucleus RNA Sequencing and Spatial Transcriptomics": Cluster8 GO top100.pdf

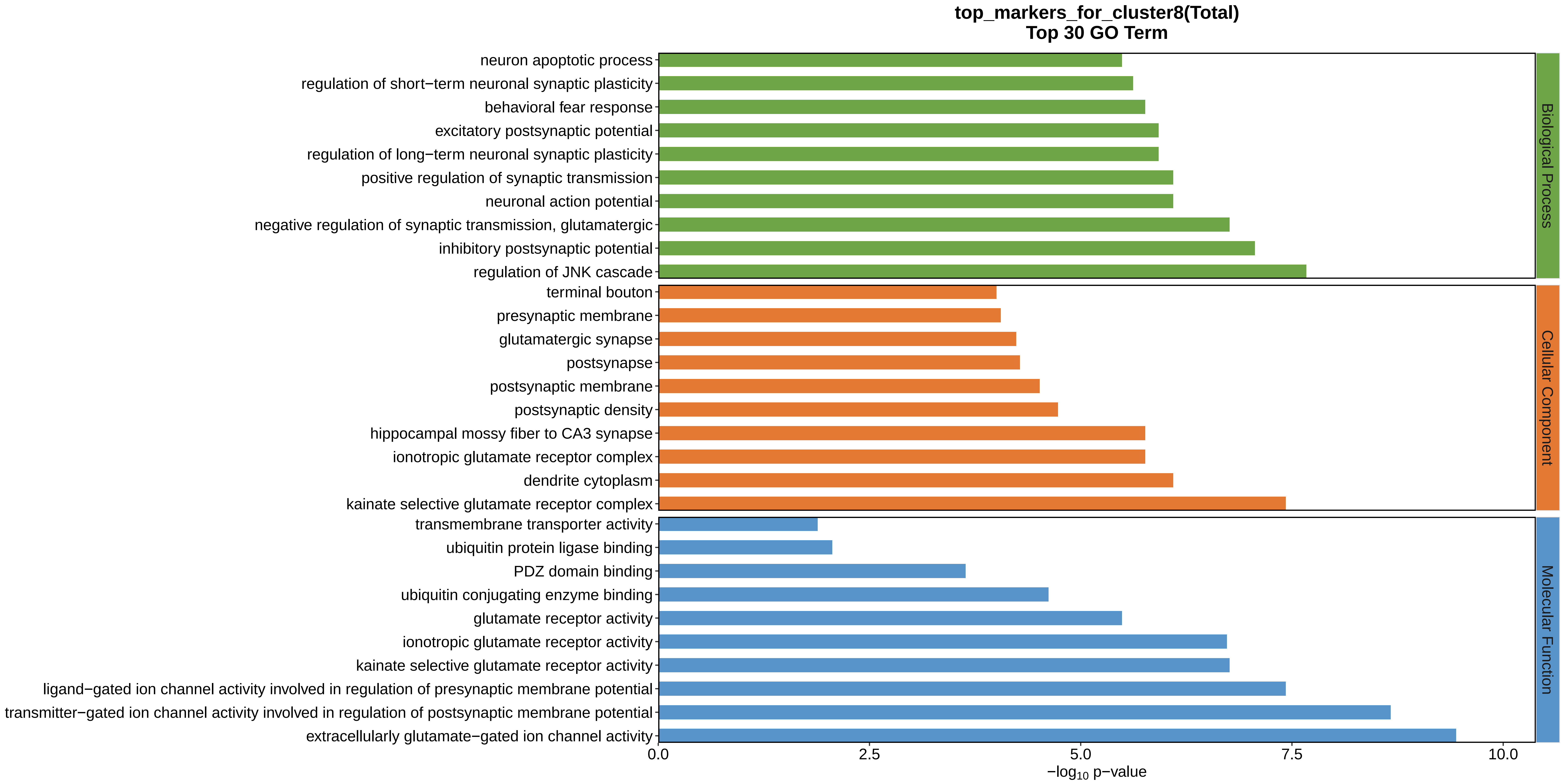
top_markers_for_cluster8(Total)
Top 30 GO Term
neuron apoptotic process
regulation of short−term neuronal synaptic plasticity
behavioral fear response
excitatory postsynaptic potential
regulation of long−term neuronal synaptic plasticity
positive regulation of synaptic transmission
neuronal action potential
negative regulation of synaptic transmission, glutamatergic
inhibitory postsynaptic potential
regulation of JNK cascade
terminal bouton
presynaptic membrane
glutamatergic synapse
postsynapse
postsynaptic membrane
postsynaptic density
hippocampal mossy fiber to CA3 synapse
ionotropic glutamate receptor complex
dendrite cytoplasm
kainate selective glutamate receptor complex
transmembrane transporter activity
ubiquitin protein ligase binding
PDZ domain binding
ubiquitin conjugating enzyme binding
glutamate receptor activity
ionotropic glutamate receptor activity
kainate selective glutamate receptor activity
ligand−gated ion channel activity involved in regulation of presynaptic membrane potential
transmitter−gated ion channel activity involved in regulation of postsynaptic membrane potential
extracellularly glutamate−gated ion channel activity
0.0
2.5
5.0
7.5
10.0
$-\log_{10}$ p−value
Biological Process
Cellular Component
Molecular Function
