## Supplemental Data 1-8 for "A Spatiotemporal Embryonic Atlas of Bombyx mori Integrating Single-Nucleus RNA Sequencing and Spatial Transcriptomics": Cluster9 GO top100.pdf

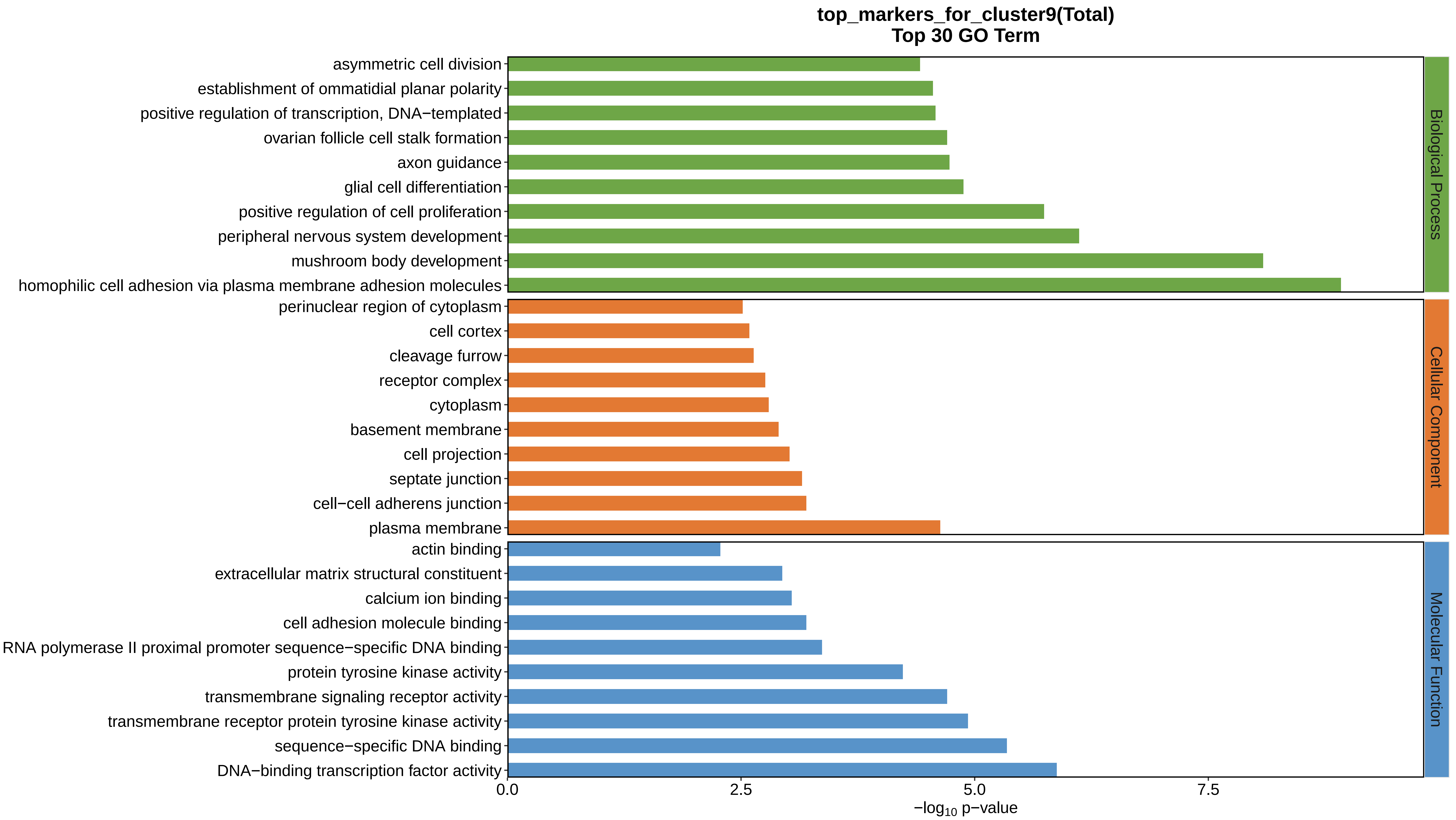

top_markers_for_cluster9(Total)
Top 30 GO Term
asymmetric cell division
establishment of ommatidial planar polarity
positive regulation of transcription, DNA−templated
ovarian follicle cell stalk formation
axon guidance
glial cell differentiation
positive regulation of cell proliferation
peripheral nervous system development
mushroom body development
homophilic cell adhesion via plasma membrane adhesion molecules
perinuclear region of cytoplasm
cell cortex
cleavage furrow
receptor complex
cytoplasm
basement membrane
cell projection
septate junction
cell−cell adherens junction
plasma membrane
actin binding
extracellular matrix structural constituent
calcium ion binding
cell adhesion molecule binding
RNA polymerase II proximal promoter sequence−specific DNA binding
protein tyrosine kinase activity
transmembrane signaling receptor activity
transmembrane receptor protein tyrosine kinase activity
sequence−specific DNA binding
DNA−binding transcription factor activity
Biological Process
Cellular Component
Molecular Function
0.0
2.5
5.0
7.5
−log10 p−value
