## Supplemental Data 1-8 for "A Spatiotemporal Embryonic Atlas of Bombyx mori Integrating Single-Nucleus RNA Sequencing and Spatial Transcriptomics": Cluster10 GO top100.pdf

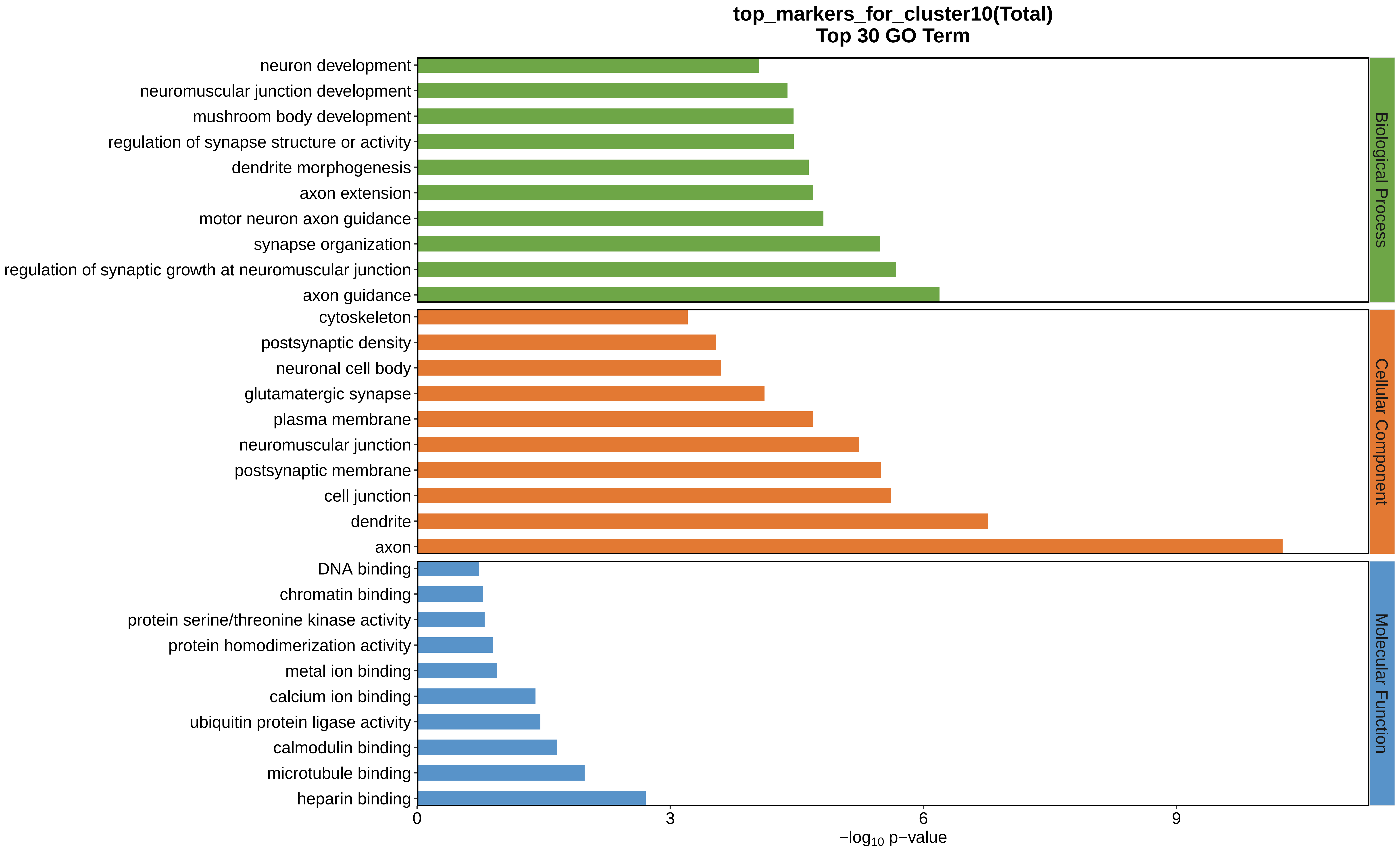
top_markers_for_cluster10(Total)
Top 30 GO Term
neuron development
neuromuscular junction development
mushroom body development
regulation of synapse structure or activity
dendrite morphogenesis
axon extension
motor neuron axon guidance
synapse organization
regulation of synaptic growth at neuromuscular junction
axon guidance
cytoskeleton
postsynaptic density
neuronal cell body
glutamatergic synapse
plasma membrane
neuromuscular junction
postsynaptic membrane
cell junction
dendrite
axon
DNA binding
chromatin binding
protein serine/threonine kinase activity
protein homodimerization activity
metal ion binding
calcium ion binding
ubiquitin protein ligase activity
calmodulin binding
microtubule binding
heparin binding
0
3
6
9
−log10 p−value
Biological Process
Cellular Component
Molecular Function
