## Supplemental Data 1-8 for "A Spatiotemporal Embryonic Atlas of Bombyx mori Integrating Single-Nucleus RNA Sequencing and Spatial Transcriptomics": Cluster11 GO top100.pdf

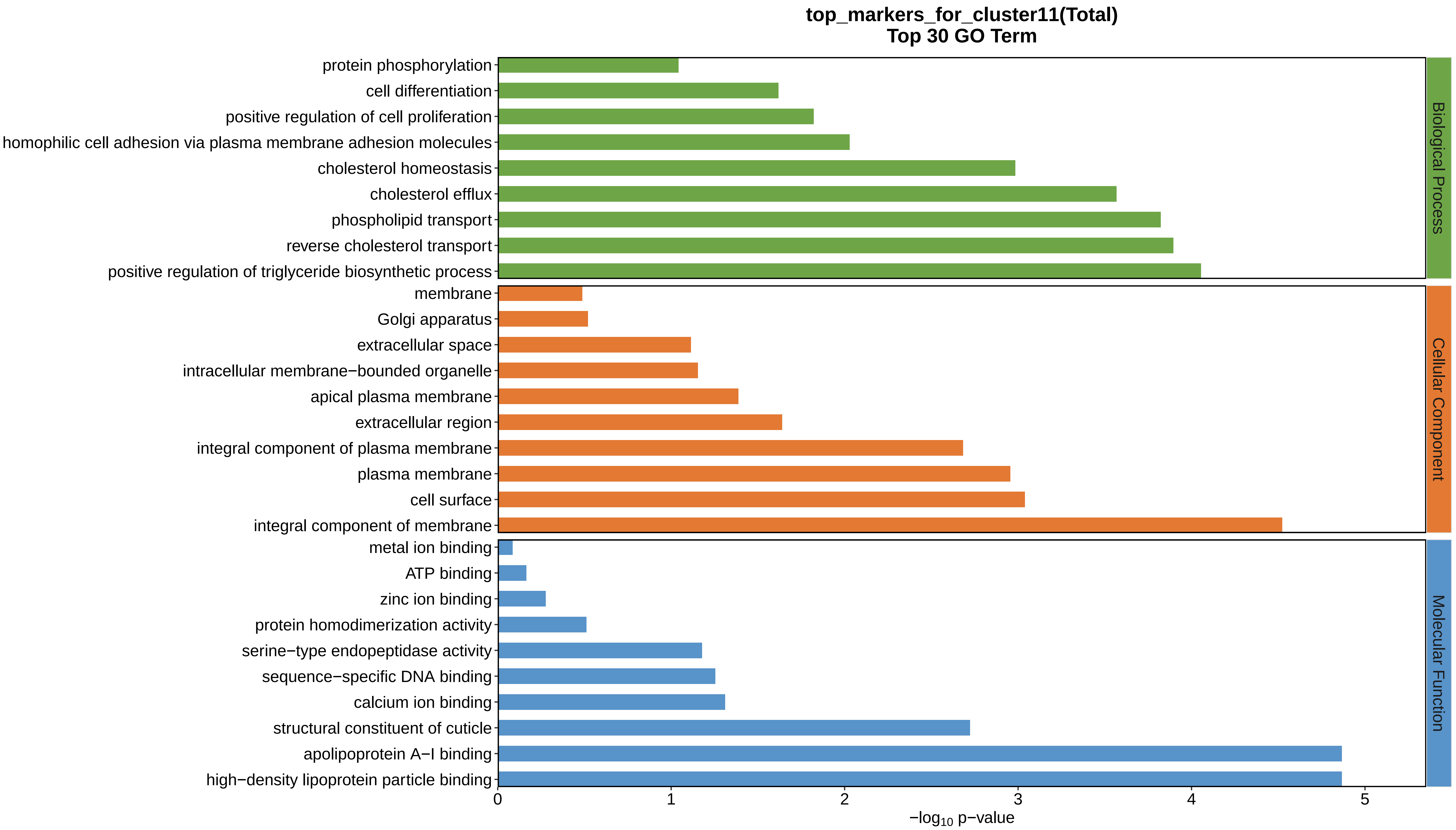
top_markers_for_cluster11(Total)
Top 30 GO Term
protein phosphorylation
cell differentiation
positive regulation of cell proliferation
homophilic cell adhesion via plasma membrane adhesion molecules
cholesterol homeostasis
cholesterol efflux
phospholipid transport
reverse cholesterol transport
positive regulation of triglyceride biosynthetic process
membrane
Golgi apparatus
extracellular space
intracellular membrane−bounded organelle
apical plasma membrane
extracellular region
integral component of plasma membrane
plasma membrane
cell surface
integral component of membrane
metal ion binding
ATP binding
zinc ion binding
protein homodimerization activity
serine−type endopeptidase activity
sequence−specific DNA binding
calcium ion binding
structural constituent of cuticle
apolipoprotein A−I binding
high−density lipoprotein particle binding
Biological Process
Cellular Component
Molecular Function
0
1
2
3
4
5
$-\log_{10}$ p−value
