## Supplemental Data 1-8 for "A Spatiotemporal Embryonic Atlas of Bombyx mori Integrating Single-Nucleus RNA Sequencing and Spatial Transcriptomics": Cluster12 GO top100.pdf

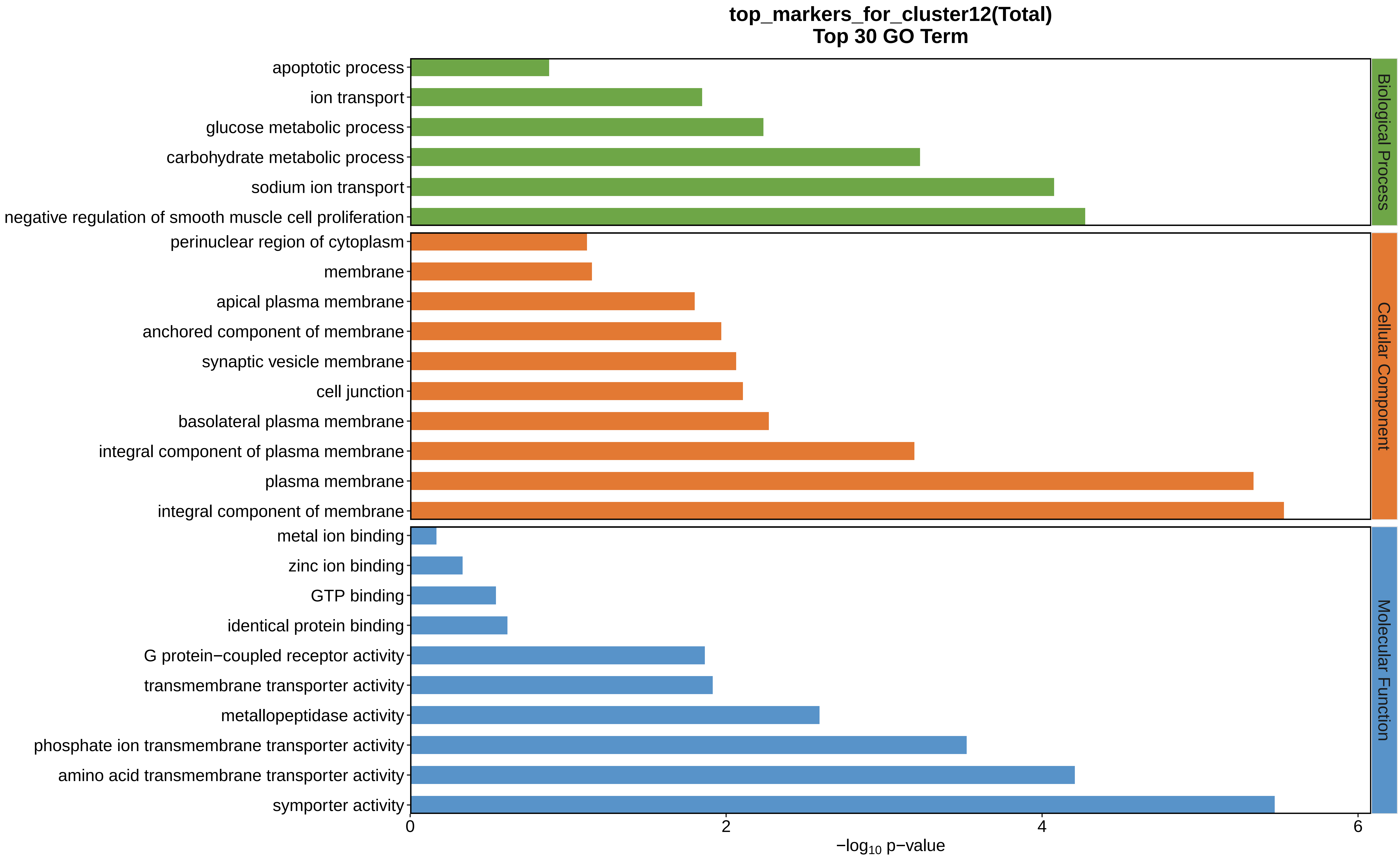
top_markers_for_cluster12(Total)
Top 30 GO Term
apoptotic process
ion transport
glucose metabolic process
carbohydrate metabolic process
sodium ion transport
negative regulation of smooth muscle cell proliferation
perinuclear region of cytoplasm
membrane
apical plasma membrane
anchored component of membrane
synaptic vesicle membrane
cell junction
basolateral plasma membrane
integral component of plasma membrane
plasma membrane
integral component of membrane
metal ion binding
zinc ion binding
GTP binding
identical protein binding
G protein−coupled receptor activity
transmembrane transporter activity
metallopeptidase activity
phosphate ion transmembrane transporter activity
amino acid transmembrane transporter activity
symporter activity
Biological Process
Cellular Component
Molecular Function
0
2
4
6
$-\log_{10}$ p−value
