## Supplemental Data 1-8 for "A Spatiotemporal Embryonic Atlas of Bombyx mori Integrating Single-Nucleus RNA Sequencing and Spatial Transcriptomics": Cluster13 GO top100.pdf

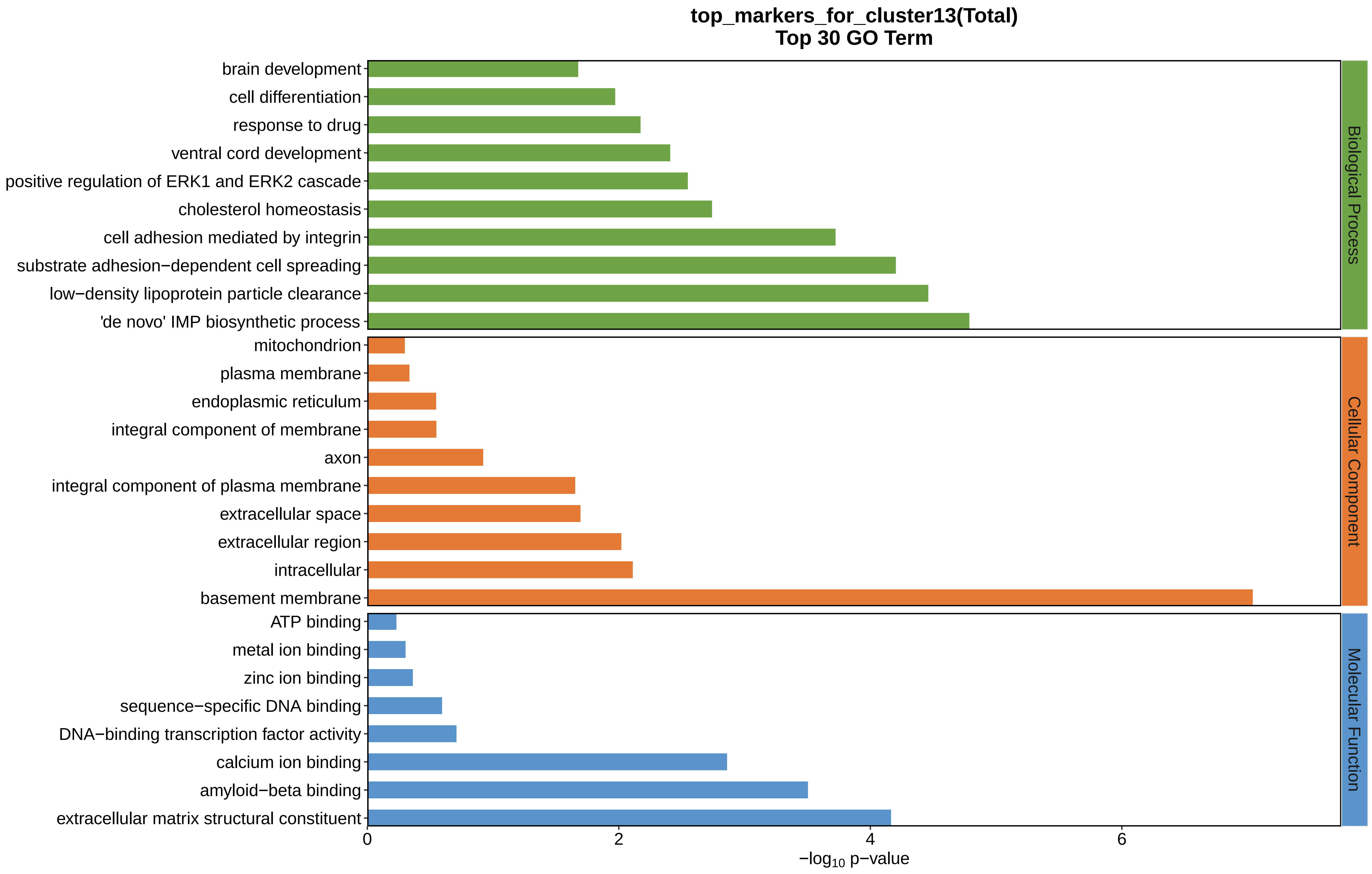
top_markers_for_cluster13(Total)
Top 30 GO Term
brain development
cell differentiation
response to drug
ventral cord development
positive regulation of ERK1 and ERK2 cascade
cholesterol homeostasis
cell adhesion mediated by integrin
substrate adhesion−dependent cell spreading
low−density lipoprotein particle clearance
'de novo' IMP biosynthetic process
mitochondrion
plasma membrane
endoplasmic reticulum
integral component of membrane
axon
integral component of plasma membrane
extracellular space
extracellular region
intracellular
basement membrane
ATP binding
metal ion binding
zinc ion binding
sequence−specific DNA binding
DNA−binding transcription factor activity
calcium ion binding
amyloid−beta binding
extracellular matrix structural constituent
Biological Process
Cellular Component
Molecular Function
0
2
4
6
−log10 p−value
