## Supplemental Data 1-8 for "A Spatiotemporal Embryonic Atlas of Bombyx mori Integrating Single-Nucleus RNA Sequencing and Spatial Transcriptomics": Cluster14 GO top100.pdf

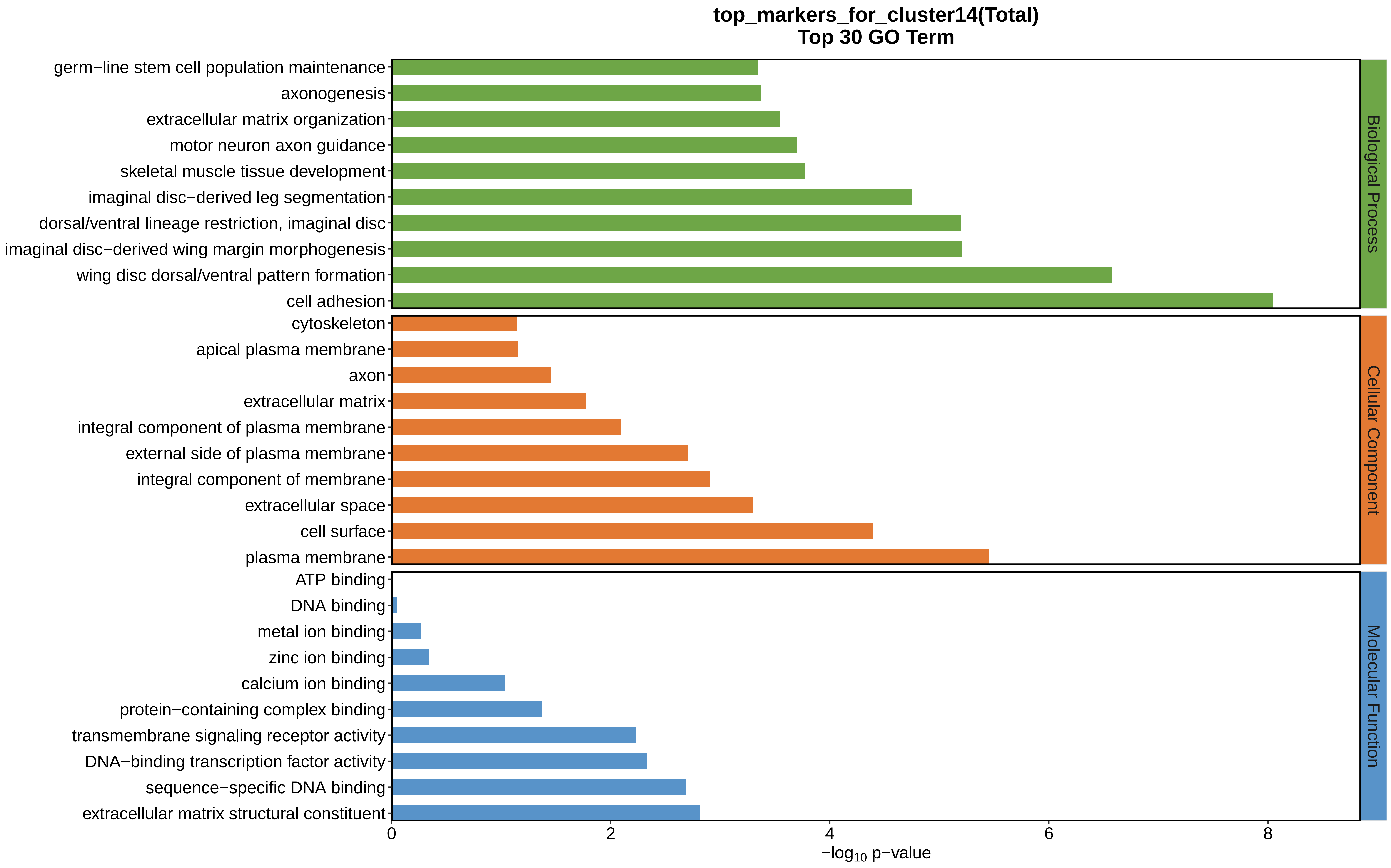

top_markers_for_cluster14(Total)
Top 30 GO Term
germ−line stem cell population maintenance
axonogenesis
extracellular matrix organization
motor neuron axon guidance
skeletal muscle tissue development
imaginal disc−derived leg segmentation
dorsal/ventral lineage restriction, imaginal disc
imaginal disc−derived wing margin morphogenesis
wing disc dorsal/ventral pattern formation
cell adhesion
cytoskeleton
apical plasma membrane
axon
extracellular matrix
integral component of plasma membrane
external side of plasma membrane
integral component of membrane
extracellular space
cell surface
plasma membrane
ATP binding
DNA binding
metal ion binding
zinc ion binding
calcium ion binding
protein−containing complex binding
transmembrane signaling receptor activity
DNA−binding transcription factor activity
sequence−specific DNA binding
extracellular matrix structural constituent
Biological Process
Cellular Component
Molecular Function
0
2
4
6
8
−log10 p−value
