## Supplemental Data 1-8 for "A Spatiotemporal Embryonic Atlas of Bombyx mori Integrating Single-Nucleus RNA Sequencing and Spatial Transcriptomics": Cluster15 GO top100.pdf

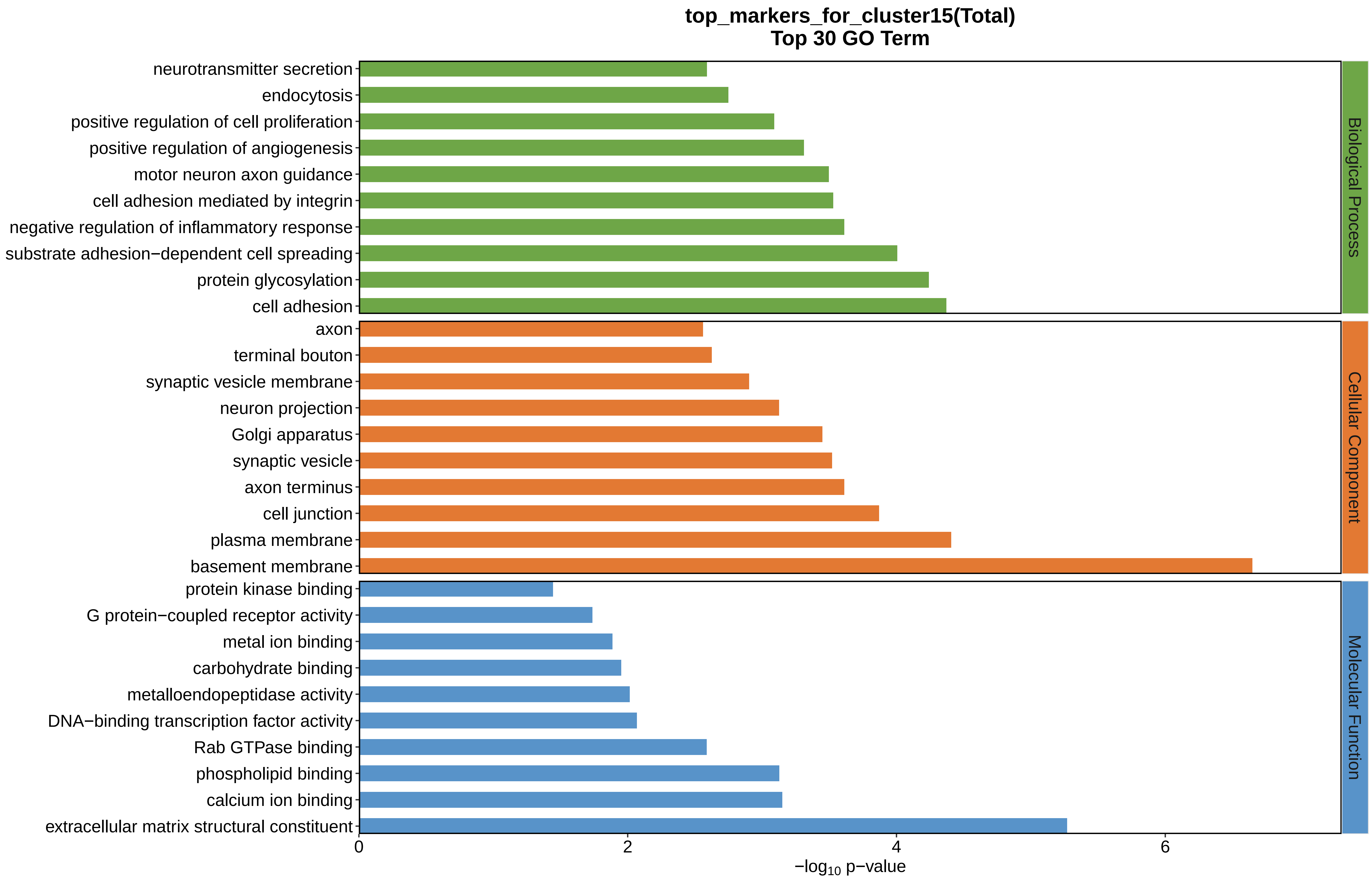
top_markers_for_cluster15(Total)
Top 30 GO Term
neurotransmitter secretion
endocytosis
positive regulation of cell proliferation
positive regulation of angiogenesis
motor neuron axon guidance
cell adhesion mediated by integrin
negative regulation of inflammatory response
substrate adhesion−dependent cell spreading
protein glycosylation
cell adhesion
axon
terminal bouton
synaptic vesicle membrane
neuron projection
Golgi apparatus
synaptic vesicle
axon terminus
cell junction
plasma membrane
basement membrane
protein kinase binding
G protein−coupled receptor activity
metal ion binding
carbohydrate binding
metalloendopeptidase activity
DNA−binding transcription factor activity
Rab GTPase binding
phospholipid binding
calcium ion binding
extracellular matrix structural constituent
Biological Process
Cellular Component
Molecular Function
0
2
4
6
$-\log_{10}$ p−value
