## Supplemental Data 1-8 for "A Spatiotemporal Embryonic Atlas of Bombyx mori Integrating Single-Nucleus RNA Sequencing and Spatial Transcriptomics": Cluster16 GO top100.pdf

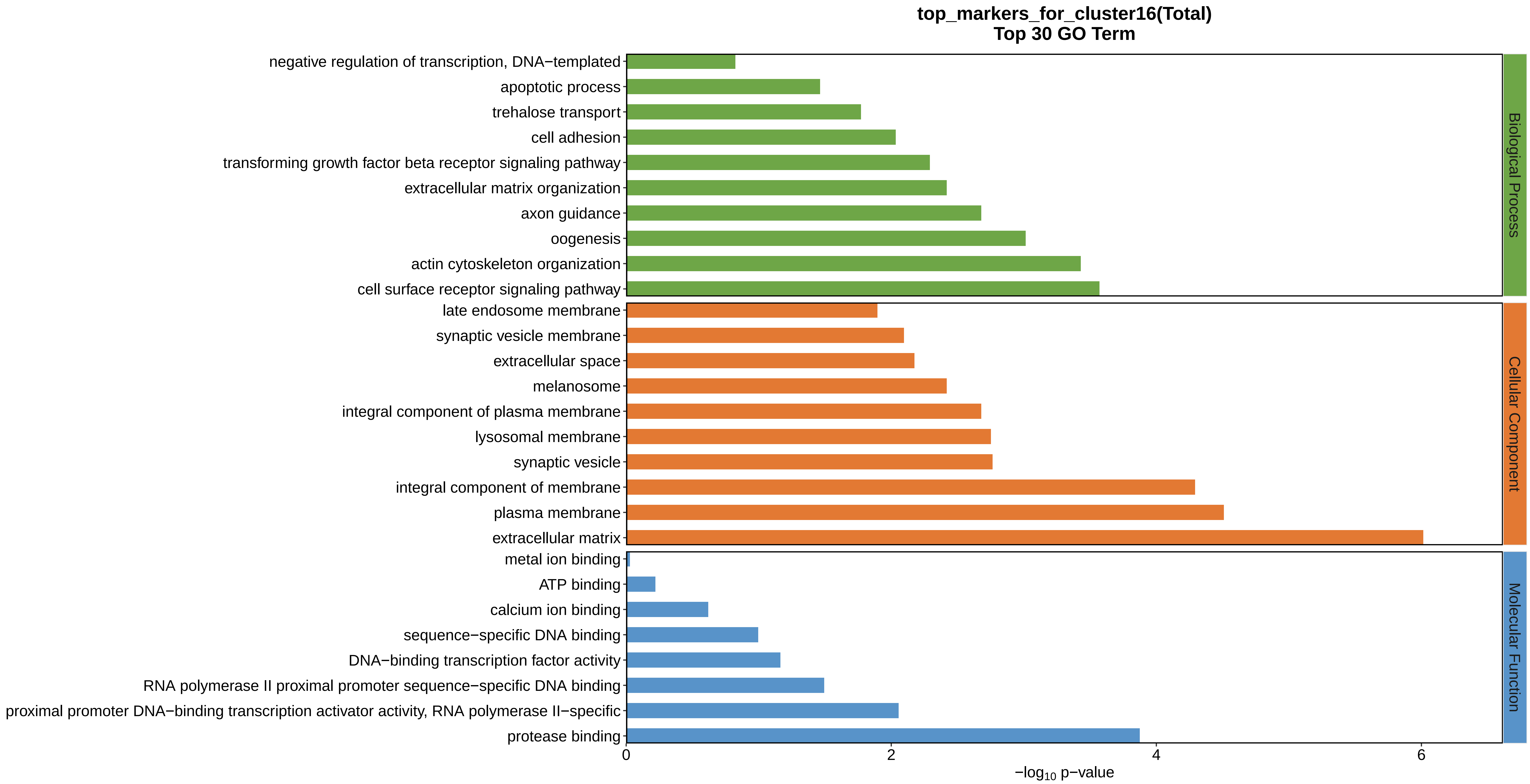
top_markers_for_cluster16(Total)
Top 30 GO Term
negative regulation of transcription, DNA−templated
apoptotic process
trehalose transport
cell adhesion
transforming growth factor beta receptor signaling pathway
extracellular matrix organization
axon guidance
oogenesis
actin cytoskeleton organization
cell surface receptor signaling pathway
late endosome membrane
synaptic vesicle membrane
extracellular space
melanosome
integral component of plasma membrane
lysosomal membrane
synaptic vesicle
integral component of membrane
plasma membrane
extracellular matrix
metal ion binding
ATP binding
calcium ion binding
sequence−specific DNA binding
DNA−binding transcription factor activity
RNA polymerase II proximal promoter sequence−specific DNA binding
proximal promoter DNA−binding transcription activator activity, RNA polymerase II−specific
protease binding
0
2
4
6
$-\log_{10}$ p−value
Biological Process
Cellular Component
Molecular Function
