## Supplemental Data 1-8 for "A Spatiotemporal Embryonic Atlas of Bombyx mori Integrating Single-Nucleus RNA Sequencing and Spatial Transcriptomics": Cluster17 GO top100.pdf

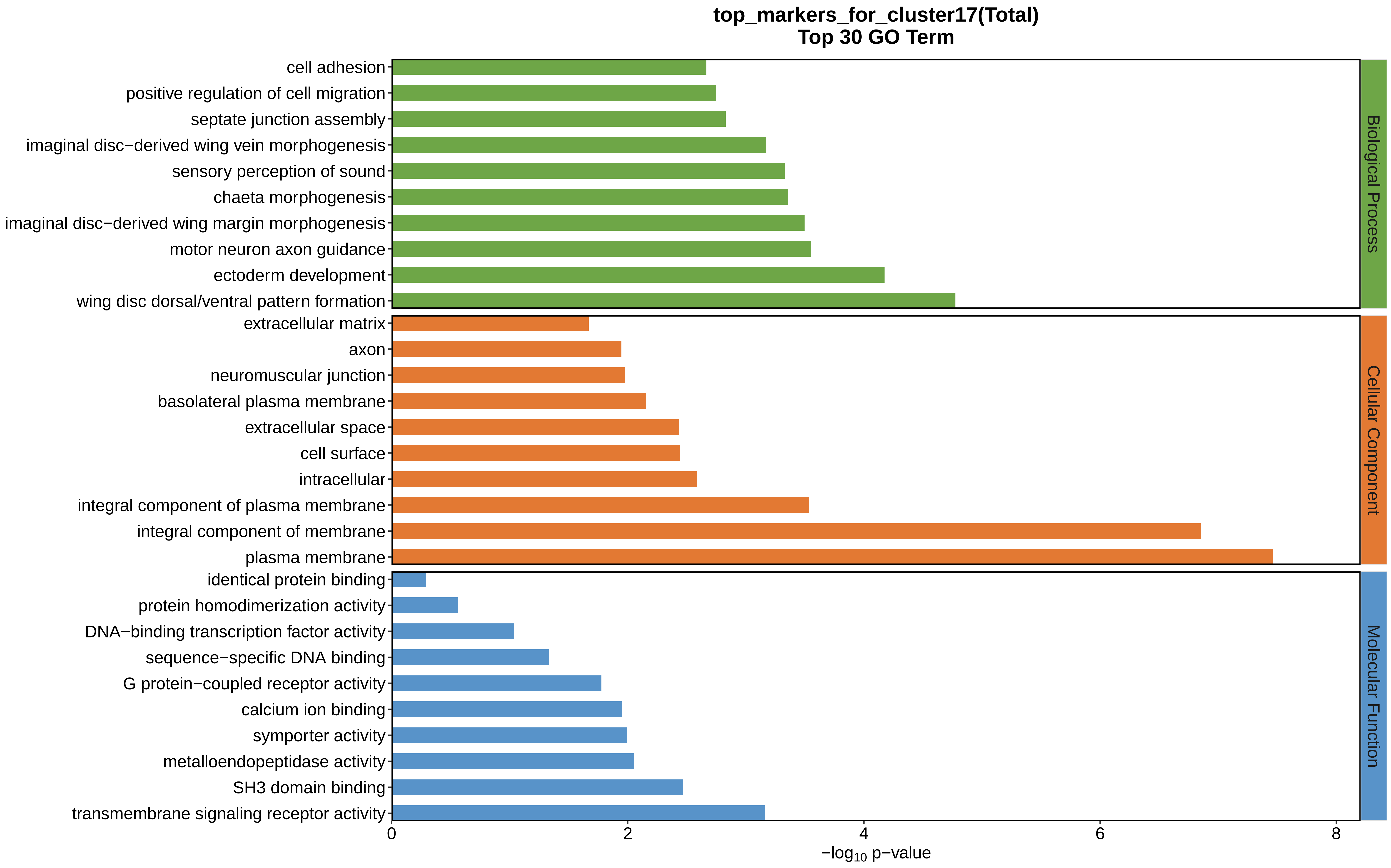

top_markers_for_cluster17(Total)
Top 30 GO Term
cell adhesion
positive regulation of cell migration
septate junction assembly
imaginal disc−derived wing vein morphogenesis
sensory perception of sound
chaeta morphogenesis
imaginal disc−derived wing margin morphogenesis
motor neuron axon guidance
ectoderm development
wing disc dorsal/ventral pattern formation
extracellular matrix
axon
neuromuscular junction
basolateral plasma membrane
extracellular space
cell surface
intracellular
integral component of plasma membrane
integral component of membrane
plasma membrane
identical protein binding
protein homodimerization activity
DNA−binding transcription factor activity
sequence−specific DNA binding
G protein−coupled receptor activity
calcium ion binding
symporter activity
metalloendopeptidase activity
SH3 domain binding
transmembrane signaling receptor activity
0
2
4
6
8
$-\log_{10}$ p−value
Biological Process
Cellular Component
Molecular Function
