## Supplemental Data 1-8 for "A Spatiotemporal Embryonic Atlas of Bombyx mori Integrating Single-Nucleus RNA Sequencing and Spatial Transcriptomics": Cluster18 GO top100.pdf

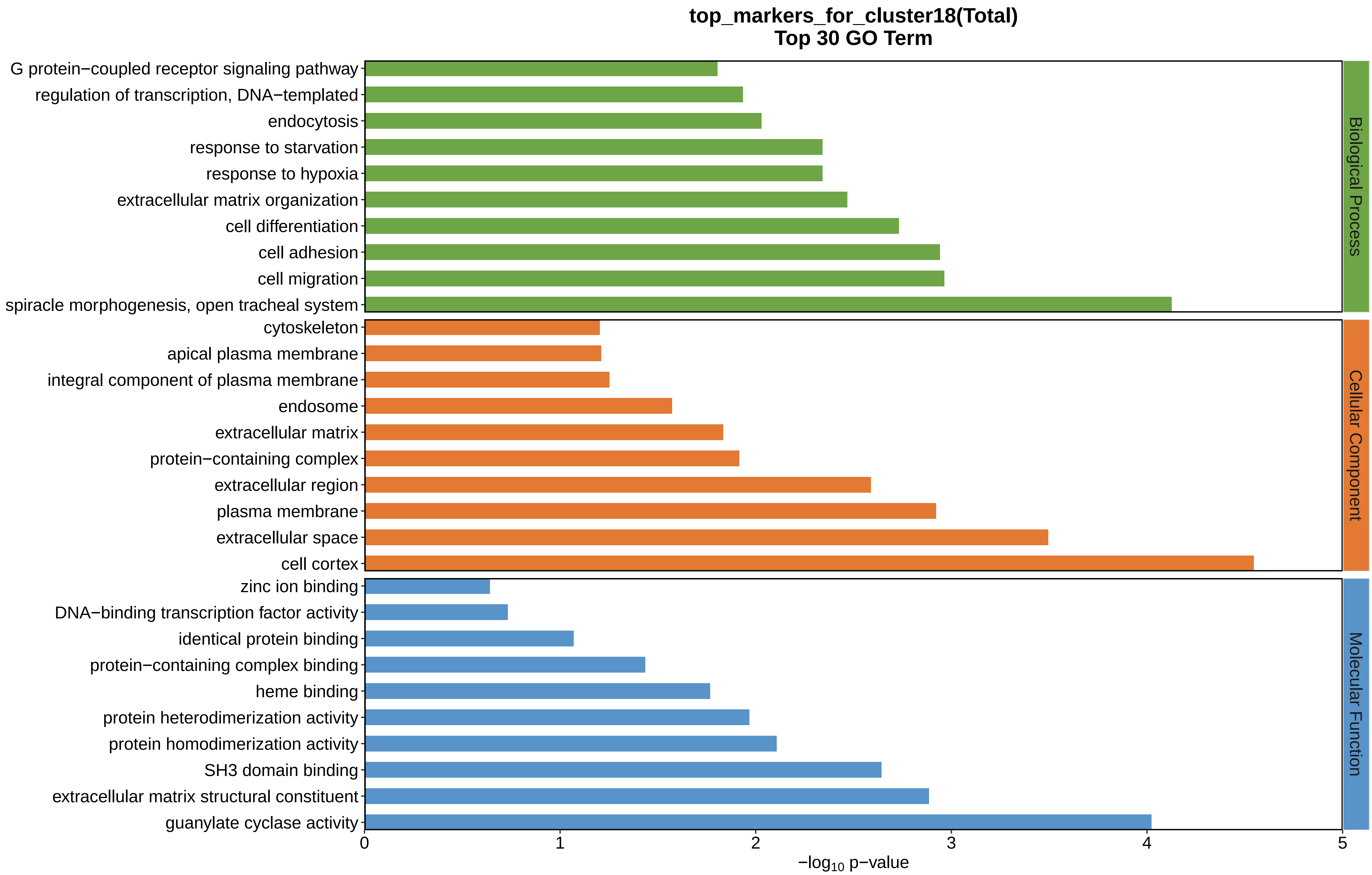
top_markers_for_cluster18(Total)
Top 30 GO Term
G protein−coupled receptor signaling pathway
regulation of transcription, DNA−templated
endocytosis
response to starvation
response to hypoxia
extracellular matrix organization
cell differentiation
cell adhesion
cell migration
spiracle morphogenesis, open tracheal system
cytoskeleton
apical plasma membrane
integral component of plasma membrane
endosome
extracellular matrix
protein−containing complex
extracellular region
plasma membrane
extracellular space
cell cortex
zinc ion binding
DNA−binding transcription factor activity
identical protein binding
protein−containing complex binding
heme binding
protein heterodimerization activity
protein homodimerization activity
SH3 domain binding
extracellular matrix structural constituent
guanylate cyclase activity
Biological Process
Cellular Component
Molecular Function
0
1
2
3
4
5
−log10 p−value
