## Supplemental Data 1-8 for "A Spatiotemporal Embryonic Atlas of Bombyx mori Integrating Single-Nucleus RNA Sequencing and Spatial Transcriptomics": Cluster19 GO top100.pdf

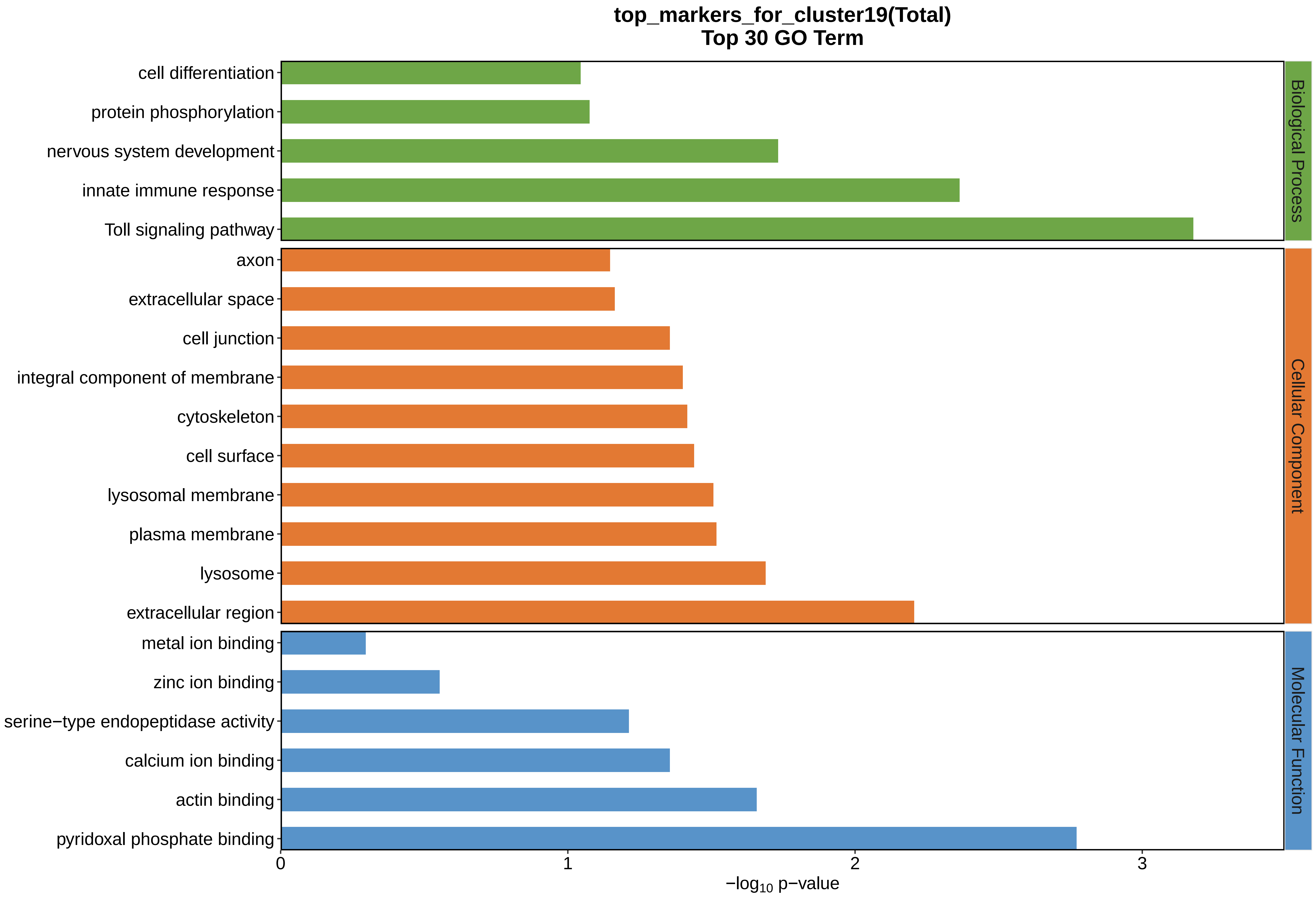
top_markers_for_cluster19(Total)
Top 30 GO Term
cell differentiation
protein phosphorylation
nervous system development
innate immune response
Toll signaling pathway
axon
extracellular space
cell junction
integral component of membrane
cytoskeleton
cell surface
lysosomal membrane
plasma membrane
lysosome
extracellular region
metal ion binding
zinc ion binding
serine−type endopeptidase activity
calcium ion binding
actin binding
pyridoxal phosphate binding
Biological Process
Cellular Component
Molecular Function
0
1
2
3
−log10 p−value
