## Supplemental Data 1-8 for "A Spatiotemporal Embryonic Atlas of Bombyx mori Integrating Single-Nucleus RNA Sequencing and Spatial Transcriptomics": Cluster20 GO top100.pdf

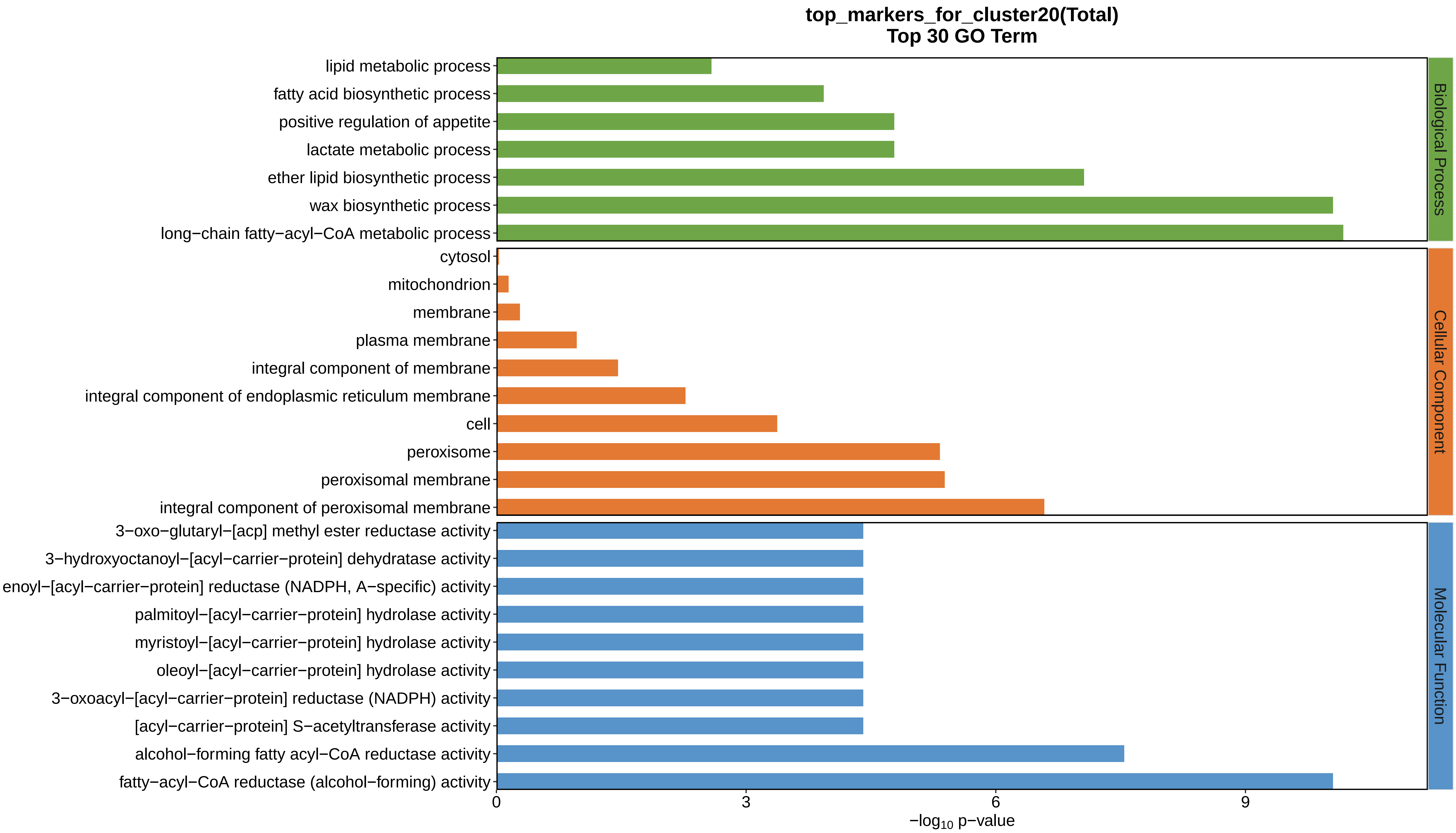

top_markers_for_cluster20(Total)
Top 30 GO Term
lipid metabolic process
fatty acid biosynthetic process
positive regulation of appetite
lactate metabolic process
ether lipid biosynthetic process
wax biosynthetic process
long−chain fatty−acyl−CoA metabolic process
cytosol
mitochondrion
membrane
plasma membrane
integral component of membrane
integral component of endoplasmic reticulum membrane
cell
peroxisome
peroxisomal membrane
integral component of peroxisomal membrane
3−oxo−glutaryl−[acp] methyl ester reductase activity
3−hydroxyoctanoyl−[acyl−carrier−protein] dehydratase activity
enoyl−[acyl−carrier−protein] reductase (NADPH, A−specific) activity
palmitoyl−[acyl−carrier−protein] hydrolase activity
myristoyl−[acyl−carrier−protein] hydrolase activity
oleoyl−[acyl−carrier−protein] hydrolase activity
3−oxoacyl−[acyl−carrier−protein] reductase (NADPH) activity
[acyl−carrier−protein] S−acetyltransferase activity
alcohol−forming fatty acyl−CoA reductase activity
fatty−acyl−CoA reductase (alcohol−forming) activity
Biological Process
Cellular Component
Molecular Function
0
3
6
9
$-\log_{10}$ p−value
