## Supplemental Data 1-8 for "A Spatiotemporal Embryonic Atlas of Bombyx mori Integrating Single-Nucleus RNA Sequencing and Spatial Transcriptomics": Cluster21 GO top100.pdf

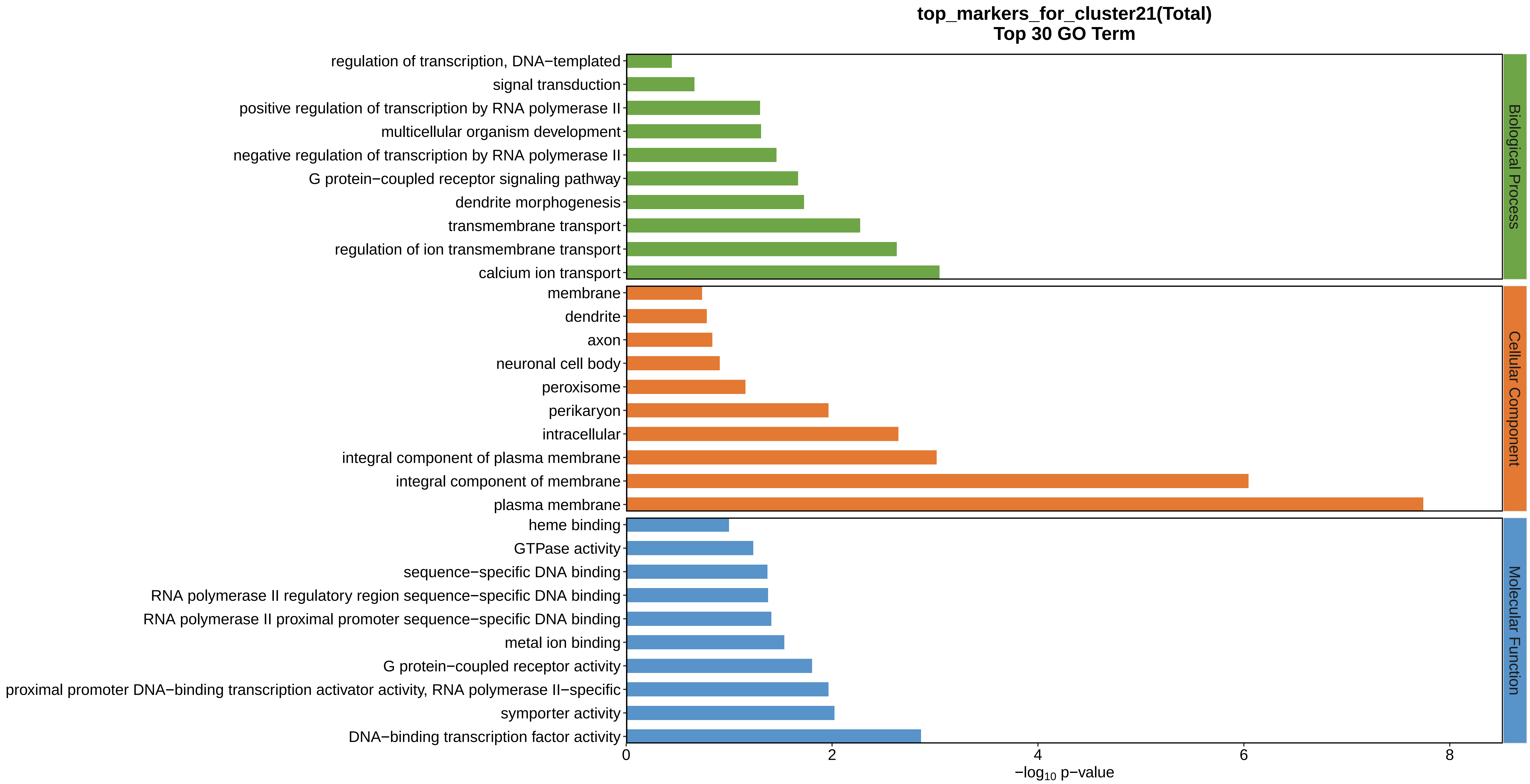
top_markers_for_cluster21(Total)
Top 30 GO Term
regulation of transcription, DNA−templated
signal transduction
positive regulation of transcription by RNA polymerase II
multicellular organism development
negative regulation of transcription by RNA polymerase II
G protein−coupled receptor signaling pathway
dendrite morphogenesis
transmembrane transport
regulation of ion transmembrane transport
calcium ion transport
membrane
dendrite
axon
neuronal cell body
peroxisome
perikaryon
intracellular
integral component of plasma membrane
integral component of membrane
plasma membrane
heme binding
GTPase activity
sequence−specific DNA binding
RNA polymerase II regulatory region sequence−specific DNA binding
RNA polymerase II proximal promoter sequence−specific DNA binding
metal ion binding
G protein−coupled receptor activity
proximal promoter DNA−binding transcription activator activity, RNA polymerase II−specific
symporter activity
DNA−binding transcription factor activity
Biological Process
Cellular Component
Molecular Function
0
2
4
6
8
−log10 p−value
