## Supplemental Data 1-8 for "A Spatiotemporal Embryonic Atlas of Bombyx mori Integrating Single-Nucleus RNA Sequencing and Spatial Transcriptomics": Cluster22 GO top100.pdf

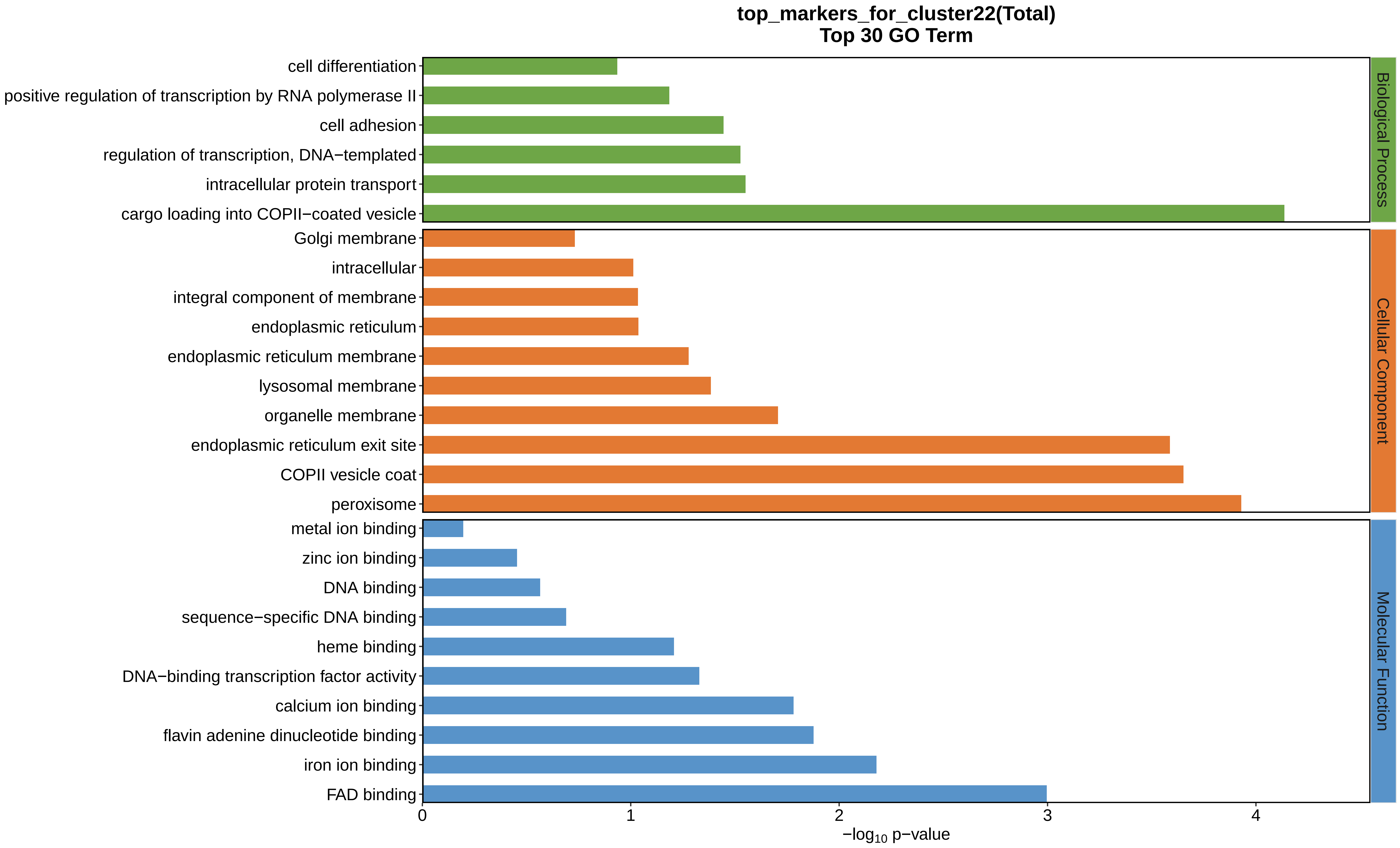
top_markers_for_cluster22(Total)
Top 30 GO Term
cell differentiation
positive regulation of transcription by RNA polymerase II
cell adhesion
regulation of transcription, DNA−templated
intracellular protein transport
cargo loading into COPII−coated vesicle
Golgi membrane
intracellular
integral component of membrane
endoplasmic reticulum
endoplasmic reticulum membrane
lysosomal membrane
organelle membrane
endoplasmic reticulum exit site
COPII vesicle coat
peroxisome
metal ion binding
zinc ion binding
DNA binding
sequence−specific DNA binding
heme binding
DNA−binding transcription factor activity
calcium ion binding
flavin adenine dinucleotide binding
iron ion binding
FAD binding
Biological Process
Cellular Component
Molecular Function
0
1
2
3
4
−log10 p−value
