## Supplemental Data 1-8 for "A Spatiotemporal Embryonic Atlas of Bombyx mori Integrating Single-Nucleus RNA Sequencing and Spatial Transcriptomics": Cluster23 GO top100.pdf

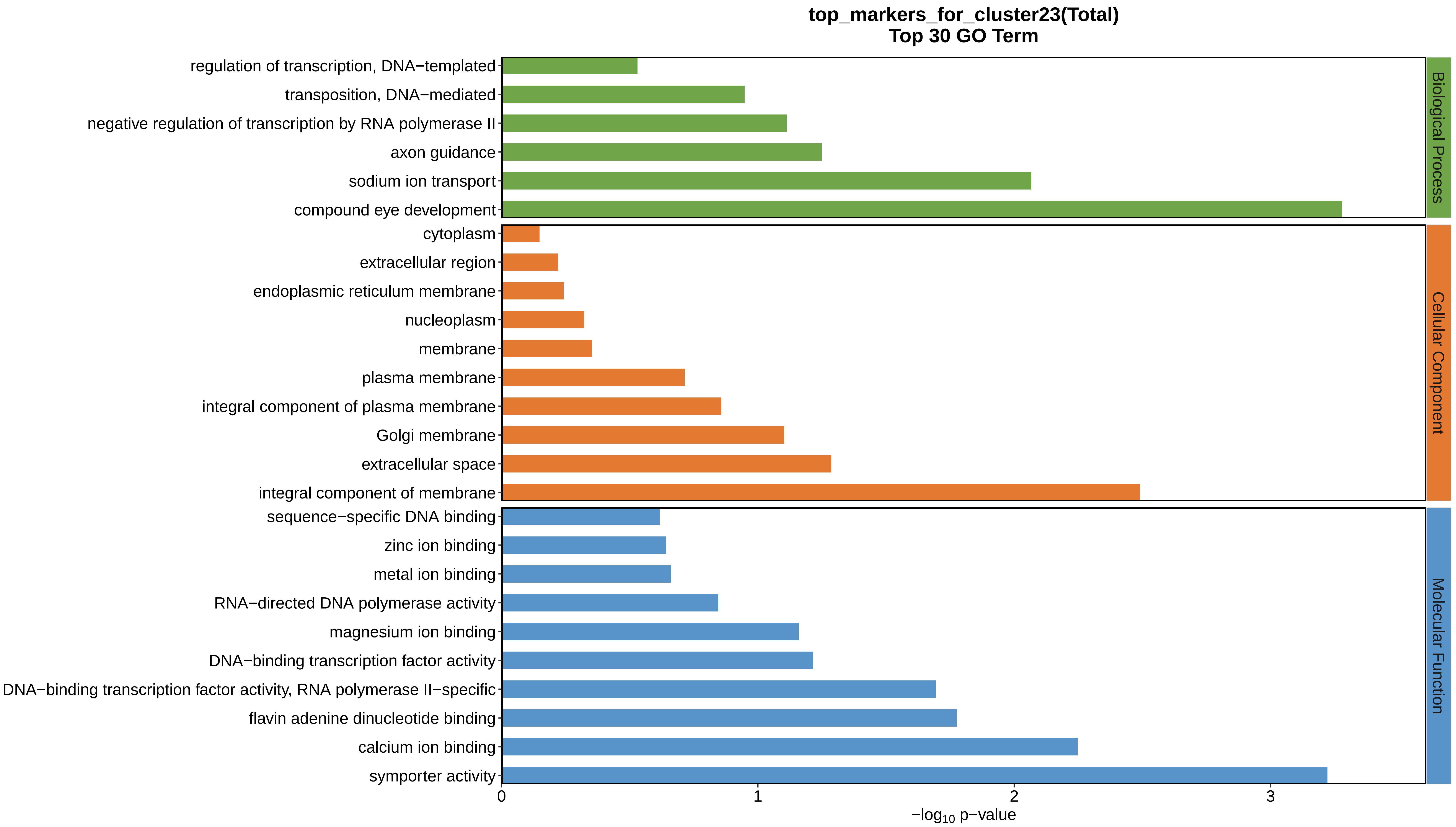
top_markers_for_cluster23(Total)
Top 30 GO Term
regulation of transcription, DNA−templated
transposition, DNA−mediated
negative regulation of transcription by RNA polymerase II
axon guidance
sodium ion transport
compound eye development
cytoplasm
extracellular region
endoplasmic reticulum membrane
nucleoplasm
membrane
plasma membrane
integral component of plasma membrane
Golgi membrane
extracellular space
integral component of membrane
sequence−specific DNA binding
zinc ion binding
metal ion binding
RNA−directed DNA polymerase activity
magnesium ion binding
DNA−binding transcription factor activity
DNA−binding transcription factor activity, RNA polymerase II−specific
flavin adenine dinucleotide binding
calcium ion binding
symporter activity
Biological Process
Cellular Component
Molecular Function
0
1
2
3
−log10 p−value
