## Supplemental Data 1-8 for "A Spatiotemporal Embryonic Atlas of Bombyx mori Integrating Single-Nucleus RNA Sequencing and Spatial Transcriptomics": Cluster24 GO top100.pdf

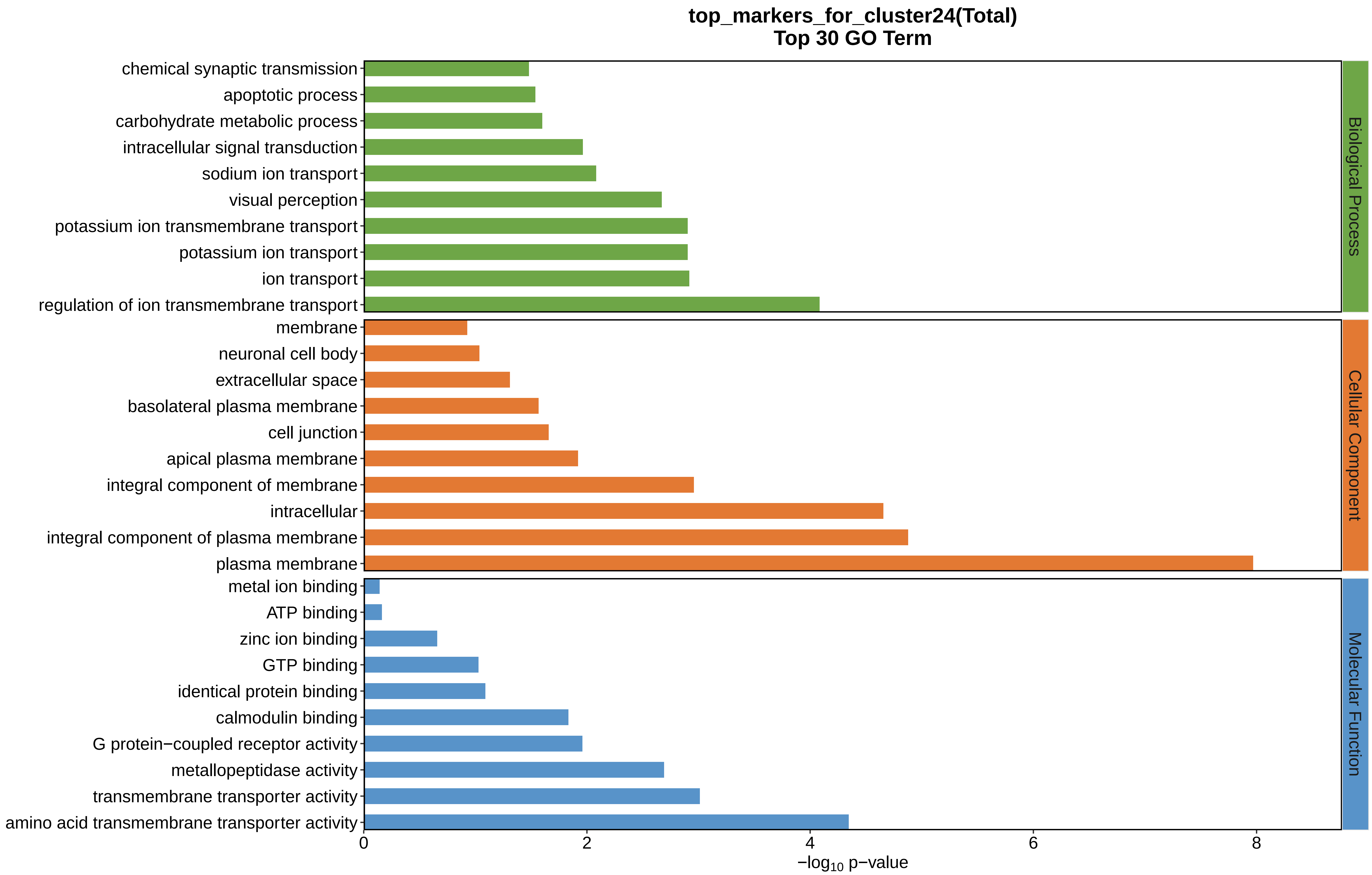
top_markers_for_cluster24(Total)
Top 30 GO Term
chemical synaptic transmission
apoptotic process
carbohydrate metabolic process
intracellular signal transduction
sodium ion transport
visual perception
potassium ion transmembrane transport
potassium ion transport
ion transport
regulation of ion transmembrane transport
membrane
neuronal cell body
extracellular space
basolateral plasma membrane
cell junction
apical plasma membrane
integral component of membrane
intracellular
integral component of plasma membrane
plasma membrane
metal ion binding
ATP binding
zinc ion binding
GTP binding
identical protein binding
calmodulin binding
G protein−coupled receptor activity
metallopeptidase activity
transmembrane transporter activity
amino acid transmembrane transporter activity
Biological Process
Cellular Component
Molecular Function
0
2
4
6
8
$-\log_{10}$ p−value
