## Supplemental Data 1-8 for "A Spatiotemporal Embryonic Atlas of Bombyx mori Integrating Single-Nucleus RNA Sequencing and Spatial Transcriptomics": Cluster1 GO top100.pdf

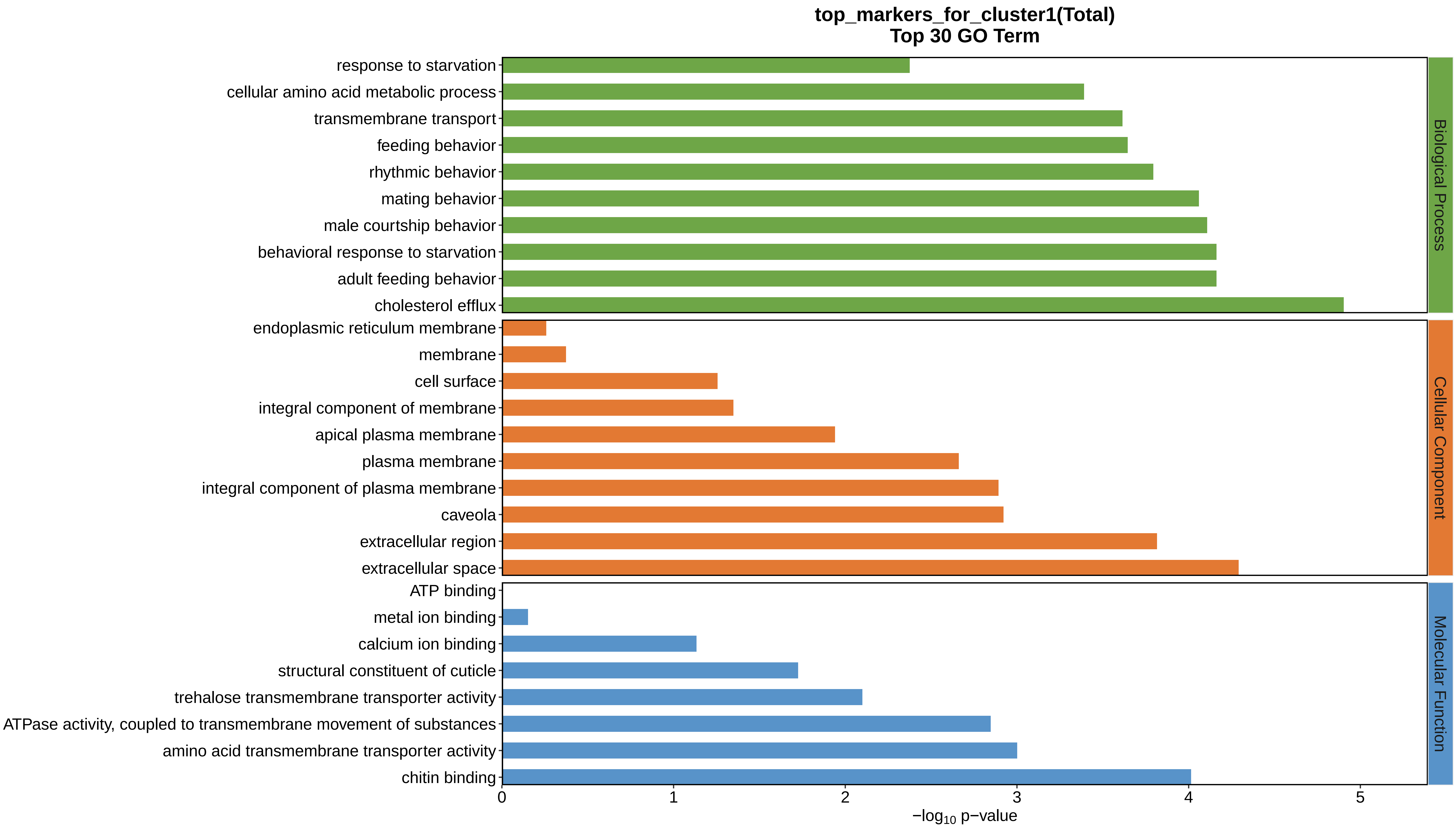
top_markers_for_cluster1(Total)
Top 30 GO Term
response to starvation
cellular amino acid metabolic process
transmembrane transport
feeding behavior
rhythmic behavior
mating behavior
male courtship behavior
behavioral response to starvation
adult feeding behavior
cholesterol efflux
endoplasmic reticulum membrane
membrane
cell surface
integral component of membrane
apical plasma membrane
plasma membrane
integral component of plasma membrane
caveola
extracellular region
extracellular space
ATP binding
metal ion binding
calcium ion binding
structural constituent of cuticle
trehalose transmembrane transporter activity
ATPase activity, coupled to transmembrane movement of substances
amino acid transmembrane transporter activity
chitin binding
Biological Process
Cellular Component
Molecular Function
0
1
2
3
4
5
−log10 p−value
