## Supplemental Data 1-8 for "A Spatiotemporal Embryonic Atlas of Bombyx mori Integrating Single-Nucleus RNA Sequencing and Spatial Transcriptomics": Cluster2 GO top100.pdf

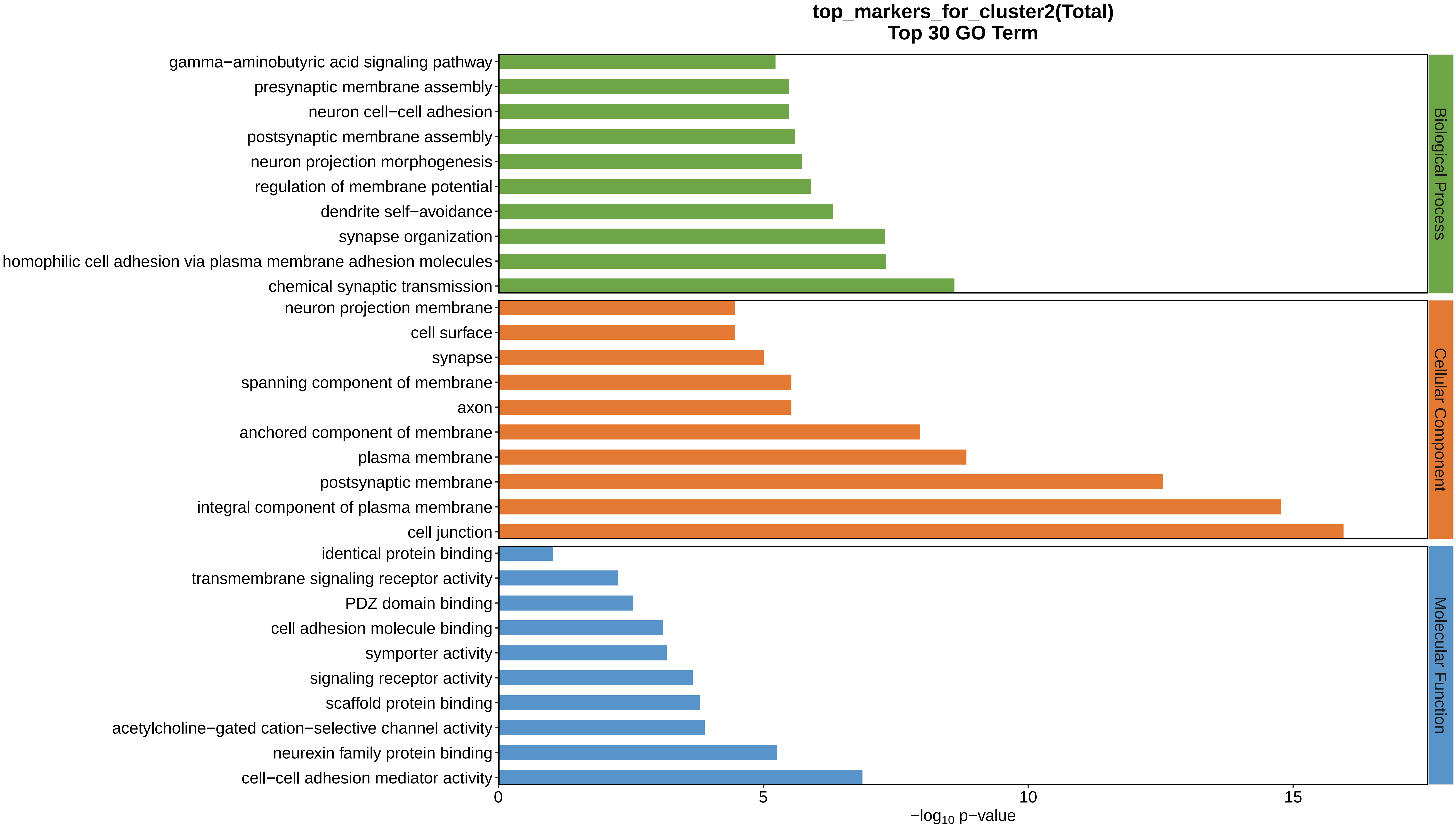
top_markers_for_cluster2(Total)
Top 30 GO Term
gamma−aminobutyric acid signaling pathway
presynaptic membrane assembly
neuron cell−cell adhesion
postsynaptic membrane assembly
neuron projection morphogenesis
regulation of membrane potential
dendrite self−avoidance
synapse organization
homophilic cell adhesion via plasma membrane adhesion molecules
chemical synaptic transmission
neuron projection membrane
cell surface
synapse
spanning component of membrane
axon
anchored component of membrane
plasma membrane
postsynaptic membrane
integral component of plasma membrane
cell junction
identical protein binding
transmembrane signaling receptor activity
PDZ domain binding
cell adhesion molecule binding
symporter activity
signaling receptor activity
scaffold protein binding
acetylcholine−gated cation−selective channel activity
neurexin family protein binding
cell−cell adhesion mediator activity
Biological Process
Cellular Component
Molecular Function
0
5
10
15
$-\log_{10}$ p−value
