## Supplemental Data 1-8 for "A Spatiotemporal Embryonic Atlas of Bombyx mori Integrating Single-Nucleus RNA Sequencing and Spatial Transcriptomics": Cluster10 GO top100.pdf

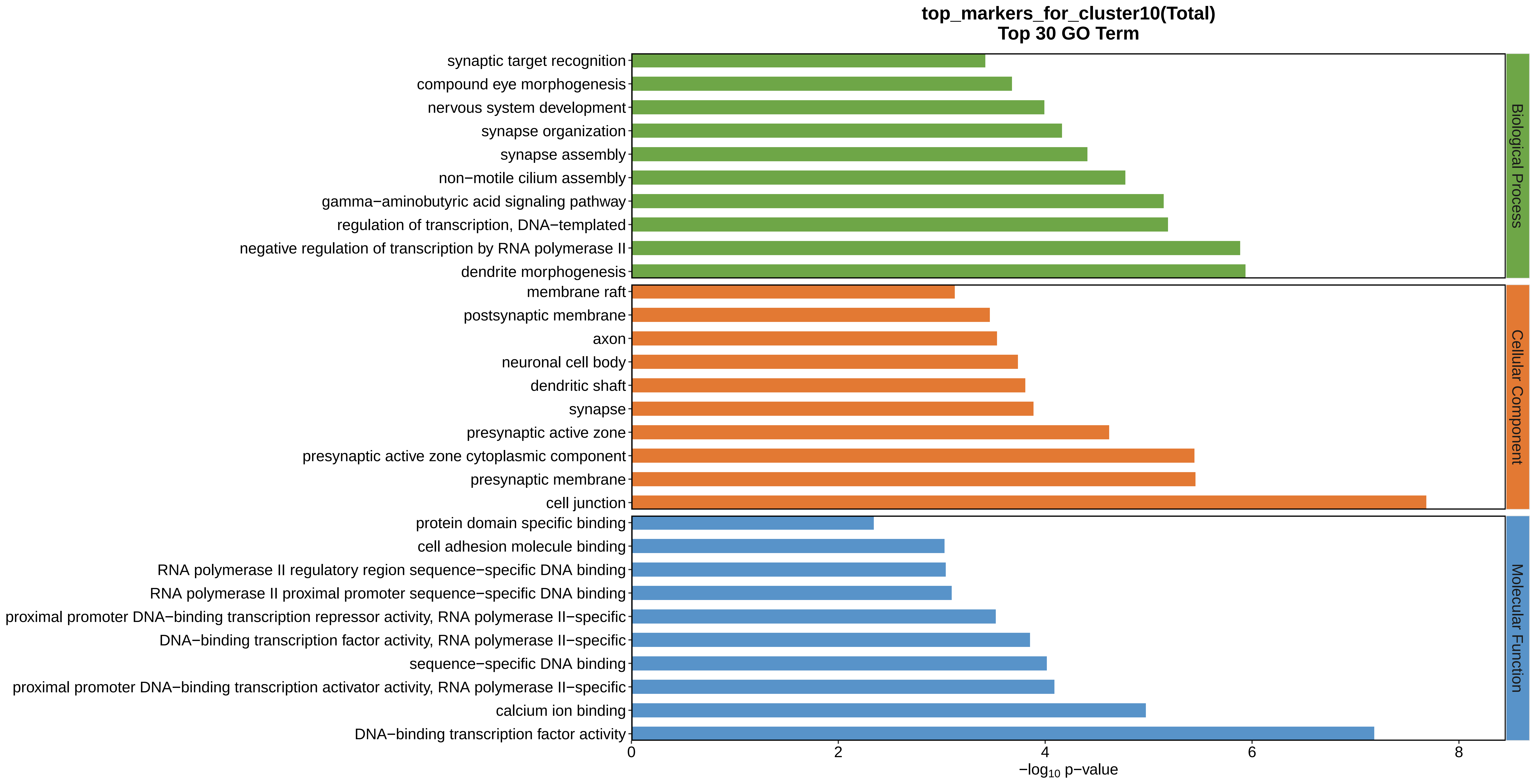

top_markers_for_cluster10(Total)
Top 30 GO Term
synaptic target recognition
compound eye morphogenesis
nervous system development
synapse organization
synapse assembly
non−motile cilium assembly
gamma−aminobutyric acid signaling pathway
regulation of transcription, DNA−templated
negative regulation of transcription by RNA polymerase II
dendrite morphogenesis
membrane raft
postsynaptic membrane
axon
neuronal cell body
dendritic shaft
synapse
presynaptic active zone
presynaptic active zone cytoplasmic component
presynaptic membrane
cell junction
protein domain specific binding
cell adhesion molecule binding
RNA polymerase II regulatory region sequence−specific DNA binding
RNA polymerase II proximal promoter sequence−specific DNA binding
proximal promoter DNA−binding transcription repressor activity, RNA polymerase II−specific
DNA−binding transcription factor activity, RNA polymerase II−specific
sequence−specific DNA binding
proximal promoter DNA−binding transcription activator activity, RNA polymerase II−specific
calcium ion binding
DNA−binding transcription factor activity
Biological Process
Cellular Component
Molecular Function
0
2
4
6
8
−log10 p−value
