## Supplemental Data 1-8 for "A Spatiotemporal Embryonic Atlas of Bombyx mori Integrating Single-Nucleus RNA Sequencing and Spatial Transcriptomics": Cluster11 GO top100.pdf

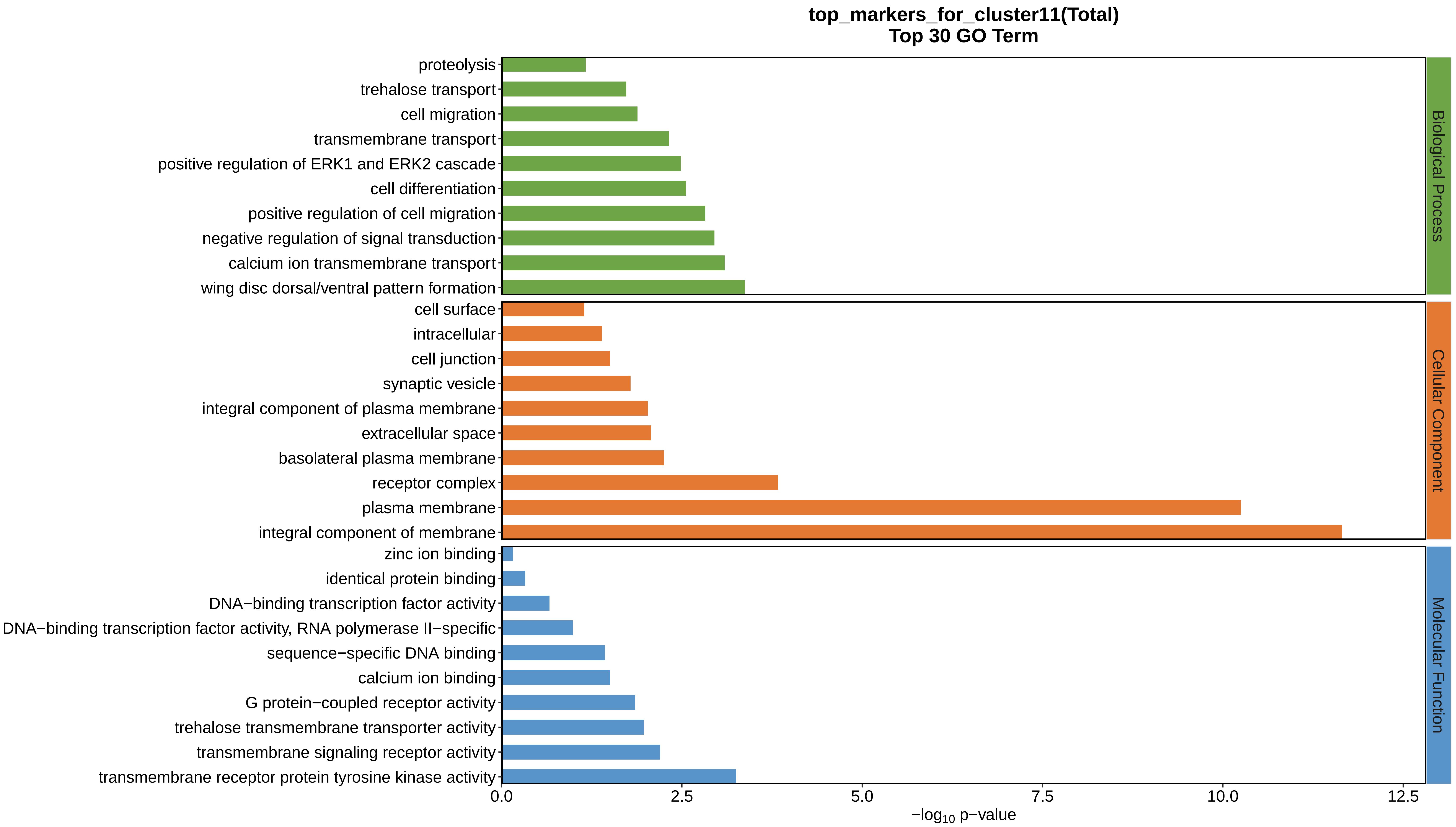
top_markers_for_cluster11(Total)
Top 30 GO Term
proteolysis
trehalose transport
cell migration
transmembrane transport
positive regulation of ERK1 and ERK2 cascade
cell differentiation
positive regulation of cell migration
negative regulation of signal transduction
calcium ion transmembrane transport
wing disc dorsal/ventral pattern formation
cell surface
intracellular
cell junction
synaptic vesicle
integral component of plasma membrane
extracellular space
basolateral plasma membrane
receptor complex
plasma membrane
integral component of membrane
zinc ion binding
identical protein binding
DNA−binding transcription factor activity
DNA−binding transcription factor activity, RNA polymerase II−specific
sequence−specific DNA binding
calcium ion binding
G protein−coupled receptor activity
trehalose transmembrane transporter activity
transmembrane signaling receptor activity
transmembrane receptor protein tyrosine kinase activity
Biological Process
Cellular Component
Molecular Function
0.0
2.5
5.0
7.5
10.0
12.5
−log10 p−value
