## Supplemental Data 1-8 for "A Spatiotemporal Embryonic Atlas of Bombyx mori Integrating Single-Nucleus RNA Sequencing and Spatial Transcriptomics": Cluster12 GO top100.pdf

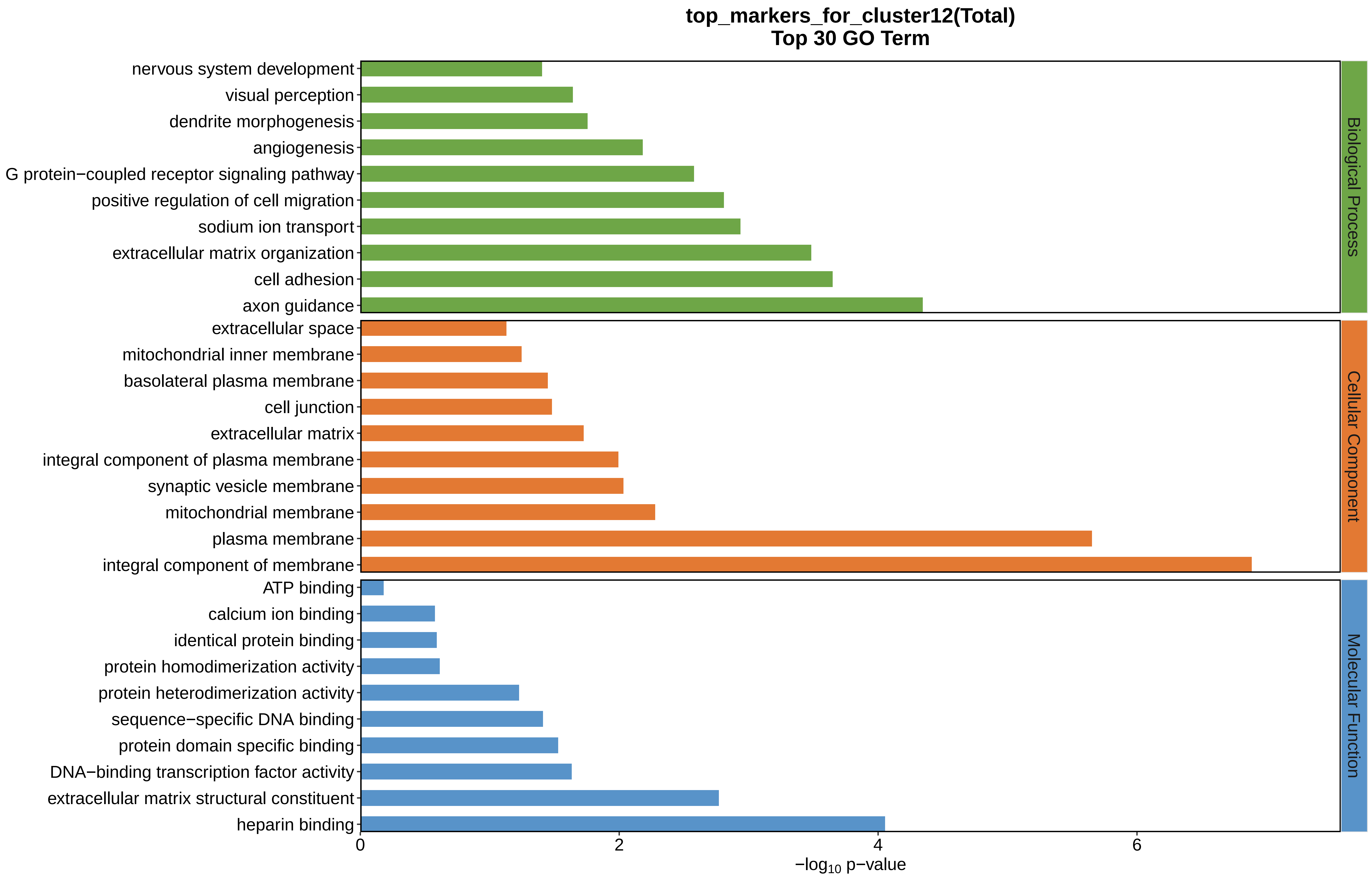
top_markers_for_cluster12(Total)
Top 30 GO Term
nervous system development
visual perception
dendrite morphogenesis
angiogenesis
G protein−coupled receptor signaling pathway
positive regulation of cell migration
sodium ion transport
extracellular matrix organization
cell adhesion
axon guidance
extracellular space
mitochondrial inner membrane
basolateral plasma membrane
cell junction
extracellular matrix
integral component of plasma membrane
synaptic vesicle membrane
mitochondrial membrane
plasma membrane
integral component of membrane
ATP binding
calcium ion binding
identical protein binding
protein homodimerization activity
protein heterodimerization activity
sequence−specific DNA binding
protein domain specific binding
DNA−binding transcription factor activity
extracellular matrix structural constituent
heparin binding
Biological Process
Cellular Component
Molecular Function
0
2
4
6
$-\log_{10}$ p−value
