## Supplemental Data 1-8 for "A Spatiotemporal Embryonic Atlas of Bombyx mori Integrating Single-Nucleus RNA Sequencing and Spatial Transcriptomics": Cluster13 GO top100.pdf

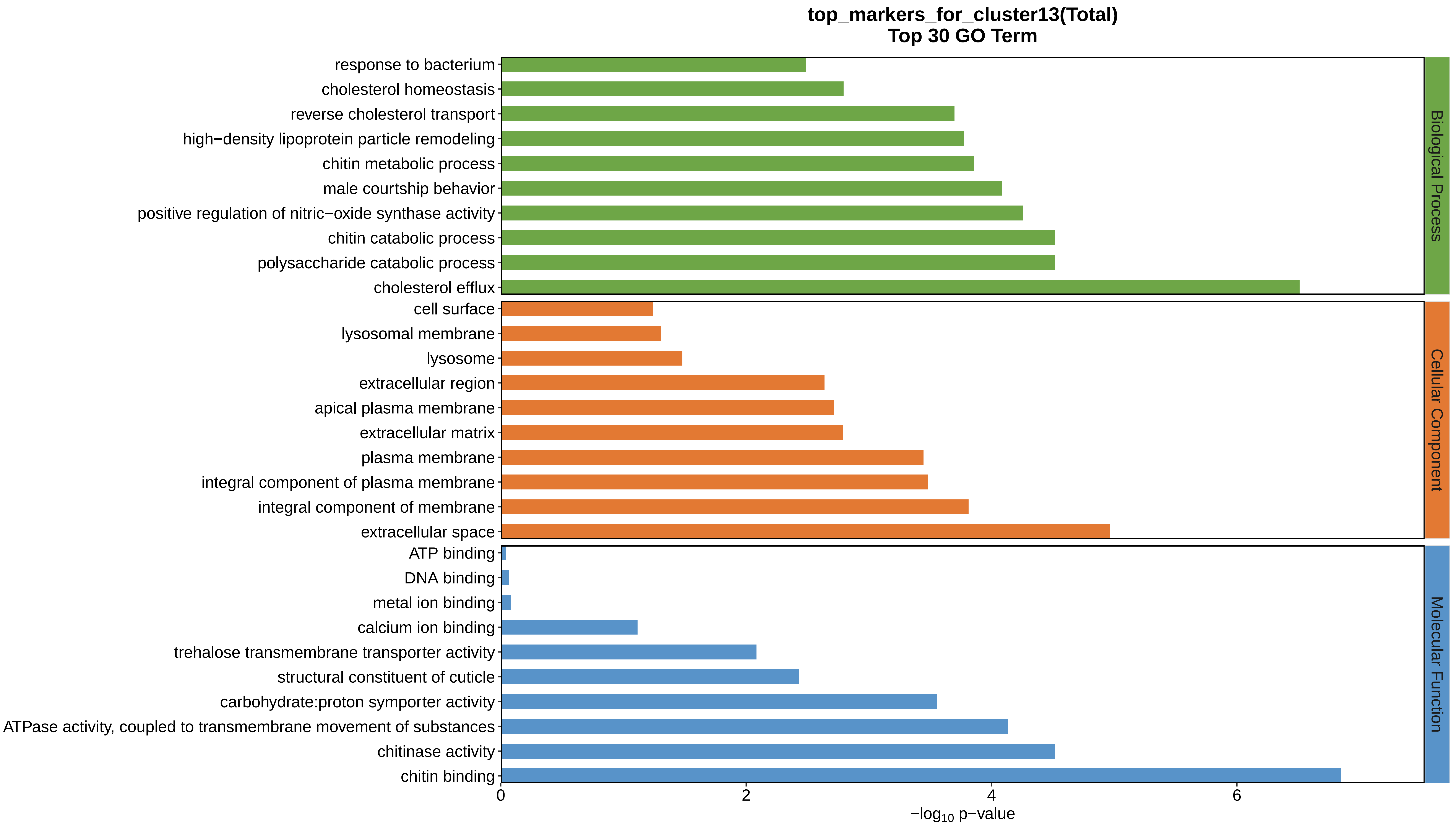
top_markers_for_cluster13(Total)
Top 30 GO Term
response to bacterium
cholesterol homeostasis
reverse cholesterol transport
high−density lipoprotein particle remodeling
chitin metabolic process
male courtship behavior
positive regulation of nitric−oxide synthase activity
chitin catabolic process
polysaccharide catabolic process
cholesterol efflux
cell surface
lysosomal membrane
lysosome
extracellular region
apical plasma membrane
extracellular matrix
plasma membrane
integral component of plasma membrane
integral component of membrane
extracellular space
ATP binding
DNA binding
metal ion binding
calcium ion binding
trehalose transmembrane transporter activity
structural constituent of cuticle
carbohydrate:proton symporter activity
ATPase activity, coupled to transmembrane movement of substances
chitinase activity
chitin binding
Biological Process
Cellular Component
Molecular Function
0
2
4
6
$-\log_{10}$ p−value
