## Supplemental Data 1-8 for "A Spatiotemporal Embryonic Atlas of Bombyx mori Integrating Single-Nucleus RNA Sequencing and Spatial Transcriptomics": Cluster14 GO top100.pdf

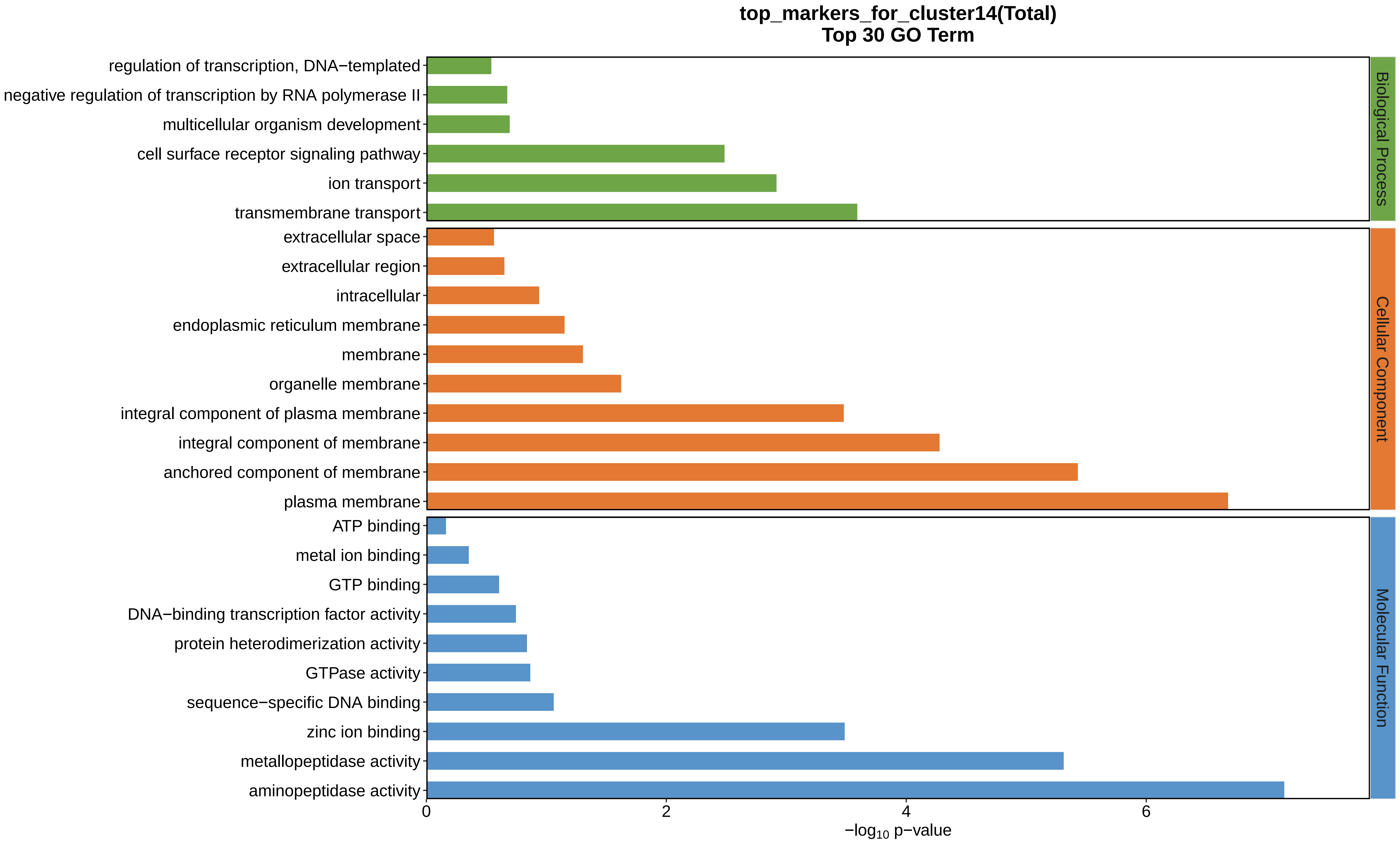
top_markers_for_cluster14(Total)
Top 30 GO Term
regulation of transcription, DNA−templated
negative regulation of transcription by RNA polymerase II
multicellular organism development
cell surface receptor signaling pathway
ion transport
transmembrane transport
extracellular space
extracellular region
intracellular
endoplasmic reticulum membrane
membrane
organelle membrane
integral component of plasma membrane
integral component of membrane
anchored component of membrane
plasma membrane
ATP binding
metal ion binding
GTP binding
DNA−binding transcription factor activity
protein heterodimerization activity
GTPase activity
sequence−specific DNA binding
zinc ion binding
metallopeptidase activity
aminopeptidase activity
Biological Process
Cellular Component
Molecular Function
0
2
4
6
−log10 p−value
