## Supplemental Data 1-8 for "A Spatiotemporal Embryonic Atlas of Bombyx mori Integrating Single-Nucleus RNA Sequencing and Spatial Transcriptomics": Cluster15 GO top100.pdf

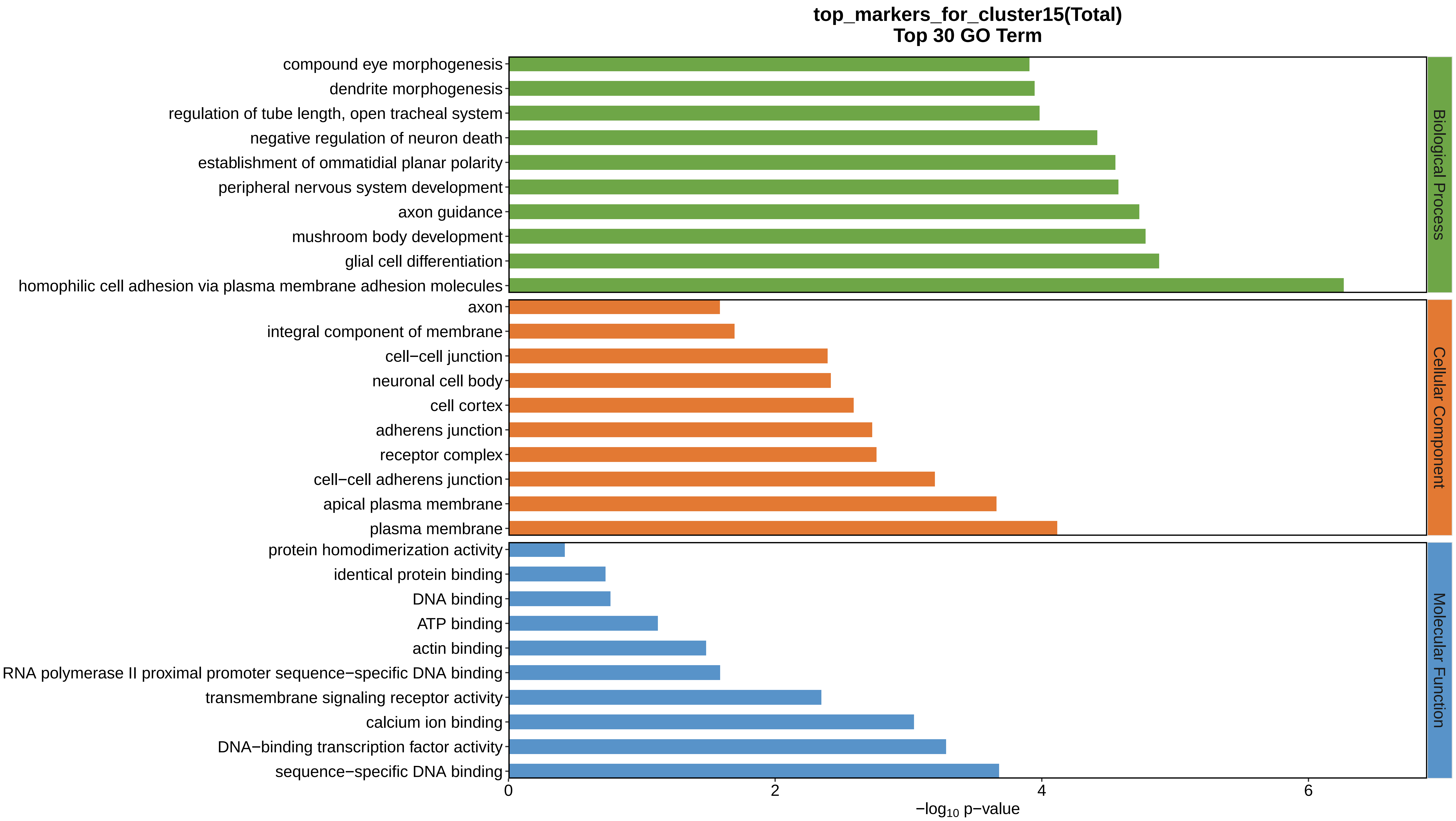
top_markers_for_cluster15(Total)
Top 30 GO Term
compound eye morphogenesis
dendrite morphogenesis
regulation of tube length, open tracheal system
negative regulation of neuron death
establishment of ommatidial planar polarity
peripheral nervous system development
axon guidance
mushroom body development
glial cell differentiation
homophilic cell adhesion via plasma membrane adhesion molecules
axon
integral component of membrane
cell−cell junction
neuronal cell body
cell cortex
adherens junction
receptor complex
cell−cell adherens junction
apical plasma membrane
plasma membrane
protein homodimerization activity
identical protein binding
DNA binding
ATP binding
actin binding
RNA polymerase II proximal promoter sequence−specific DNA binding
transmembrane signaling receptor activity
calcium ion binding
DNA−binding transcription factor activity
sequence−specific DNA binding
0
2
4
6
−log10 p−value
Biological Process
Cellular Component
Molecular Function
