## Supplemental Data 1-8 for "A Spatiotemporal Embryonic Atlas of Bombyx mori Integrating Single-Nucleus RNA Sequencing and Spatial Transcriptomics": Cluster16 GO top100.pdf

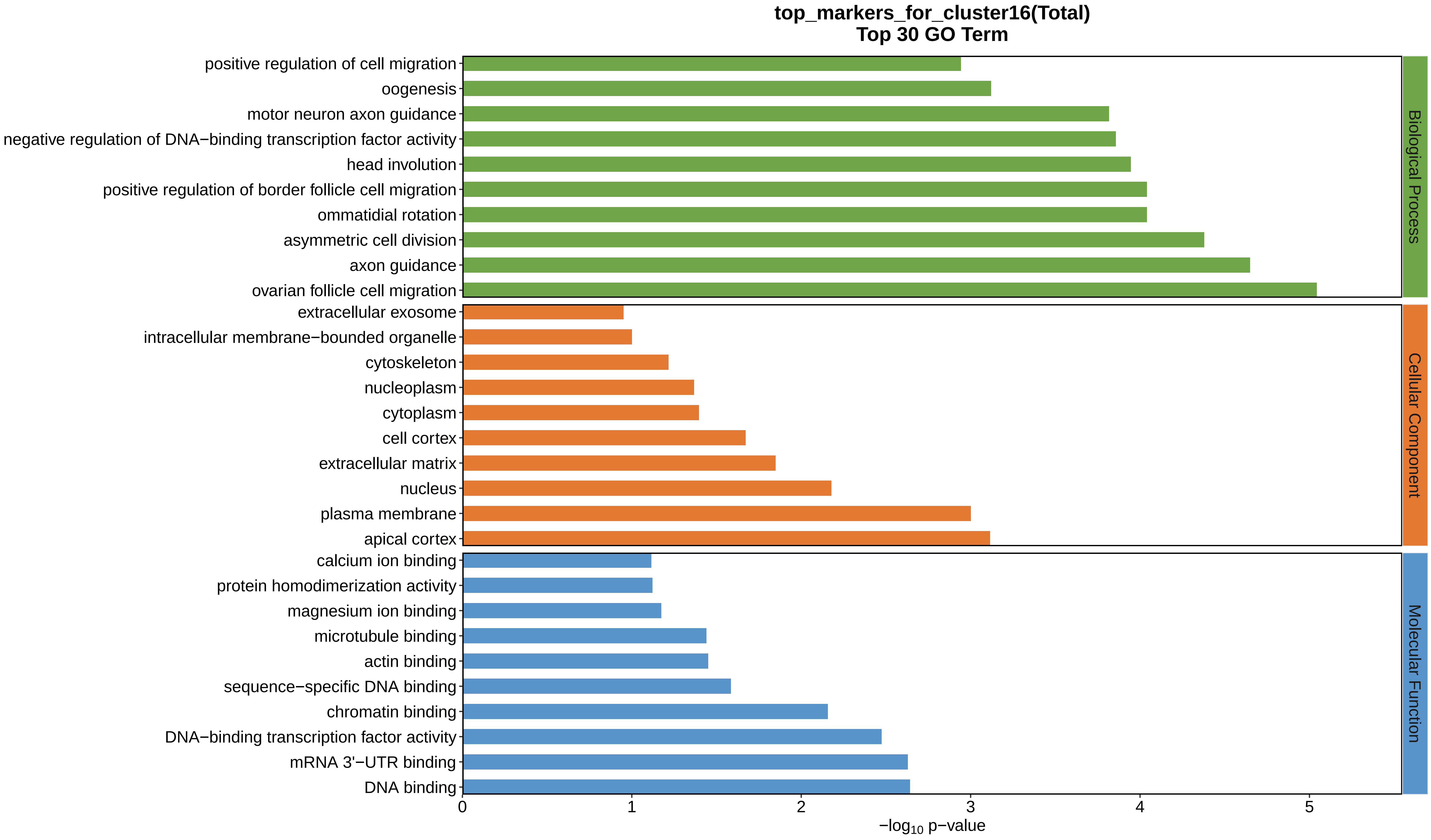
top_markers_for_cluster16(Total)
Top 30 GO Term
positive regulation of cell migration
oogenesis
motor neuron axon guidance
negative regulation of DNA−binding transcription factor activity
head involution
positive regulation of border follicle cell migration
ommatidial rotation
asymmetric cell division
axon guidance
ovarian follicle cell migration
extracellular exosome
intracellular membrane−bounded organelle
cytoskeleton
nucleoplasm
cytoplasm
cell cortex
extracellular matrix
nucleus
plasma membrane
apical cortex
calcium ion binding
protein homodimerization activity
magnesium ion binding
microtubule binding
actin binding
sequence−specific DNA binding
chromatin binding
DNA−binding transcription factor activity
mRNA 3'−UTR binding
DNA binding
Biological Process
Cellular Component
Molecular Function
0
1
2
3
4
5
$-\log_{10}$ p−value
