## Supplemental Data 1-8 for "A Spatiotemporal Embryonic Atlas of Bombyx mori Integrating Single-Nucleus RNA Sequencing and Spatial Transcriptomics": Cluster17 GO top100.pdf

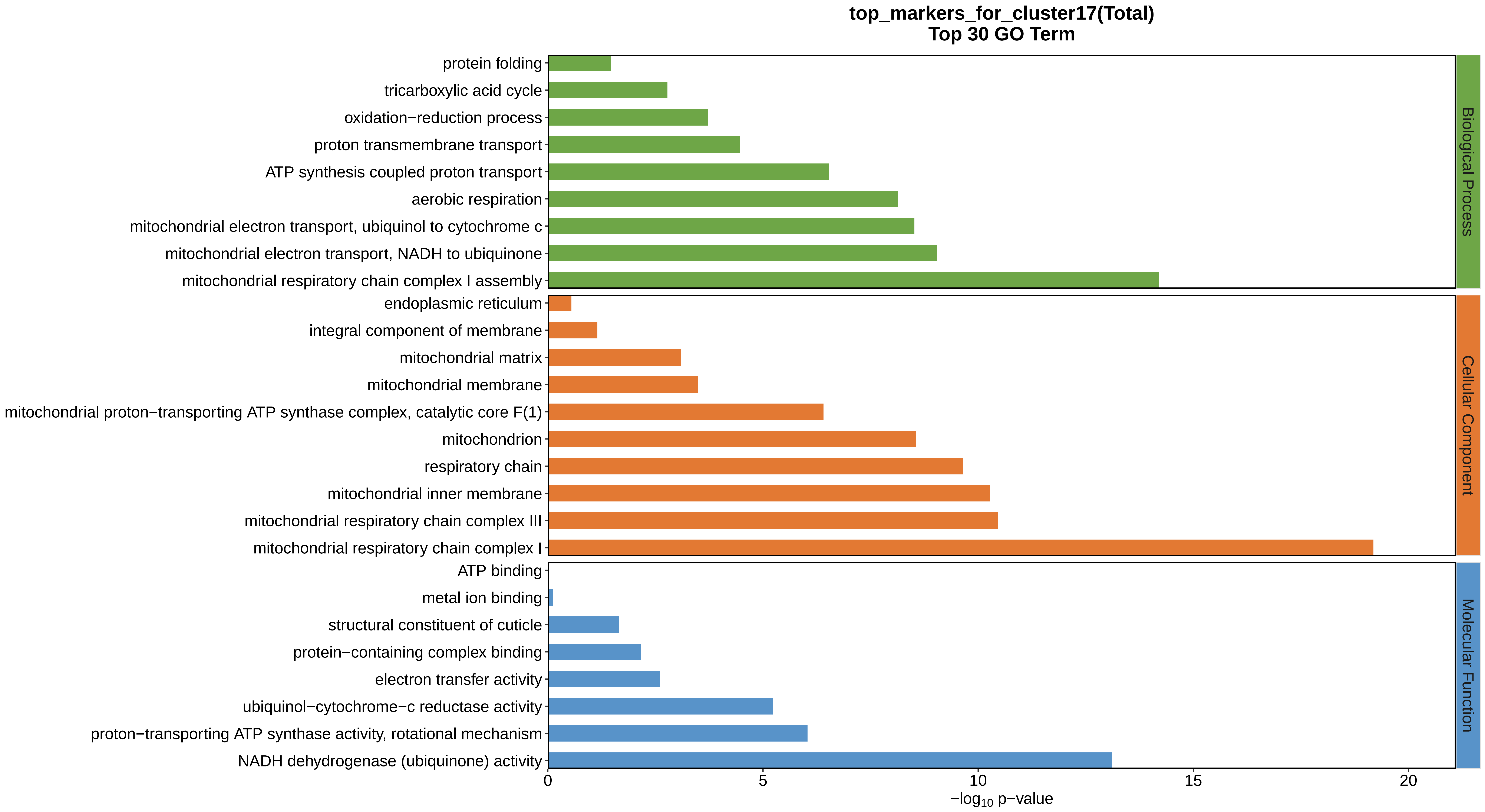
top_markers_for_cluster17(Total)
Top 30 GO Term
protein folding
tricarboxylic acid cycle
oxidation−reduction process
proton transmembrane transport
ATP synthesis coupled proton transport
aerobic respiration
mitochondrial electron transport, ubiquinol to cytochrome c
mitochondrial electron transport, NADH to ubiquinone
mitochondrial respiratory chain complex I assembly
endoplasmic reticulum
integral component of membrane
mitochondrial matrix
mitochondrial membrane
mitochondrial proton−transporting ATP synthase complex, catalytic core F(1)
mitochondrion
respiratory chain
mitochondrial inner membrane
mitochondrial respiratory chain complex III
mitochondrial respiratory chain complex I
ATP binding
metal ion binding
structural constituent of cuticle
protein−containing complex binding
electron transfer activity
ubiquinol−cytochrome−c reductase activity
proton−transporting ATP synthase activity, rotational mechanism
NADH dehydrogenase (ubiquinone) activity
Biological Process
Cellular Component
Molecular Function
0
5
10
15
20
−log10 p−value
