## Supplemental Data 1-8 for "A Spatiotemporal Embryonic Atlas of Bombyx mori Integrating Single-Nucleus RNA Sequencing and Spatial Transcriptomics": Cluster18 GO top100.pdf

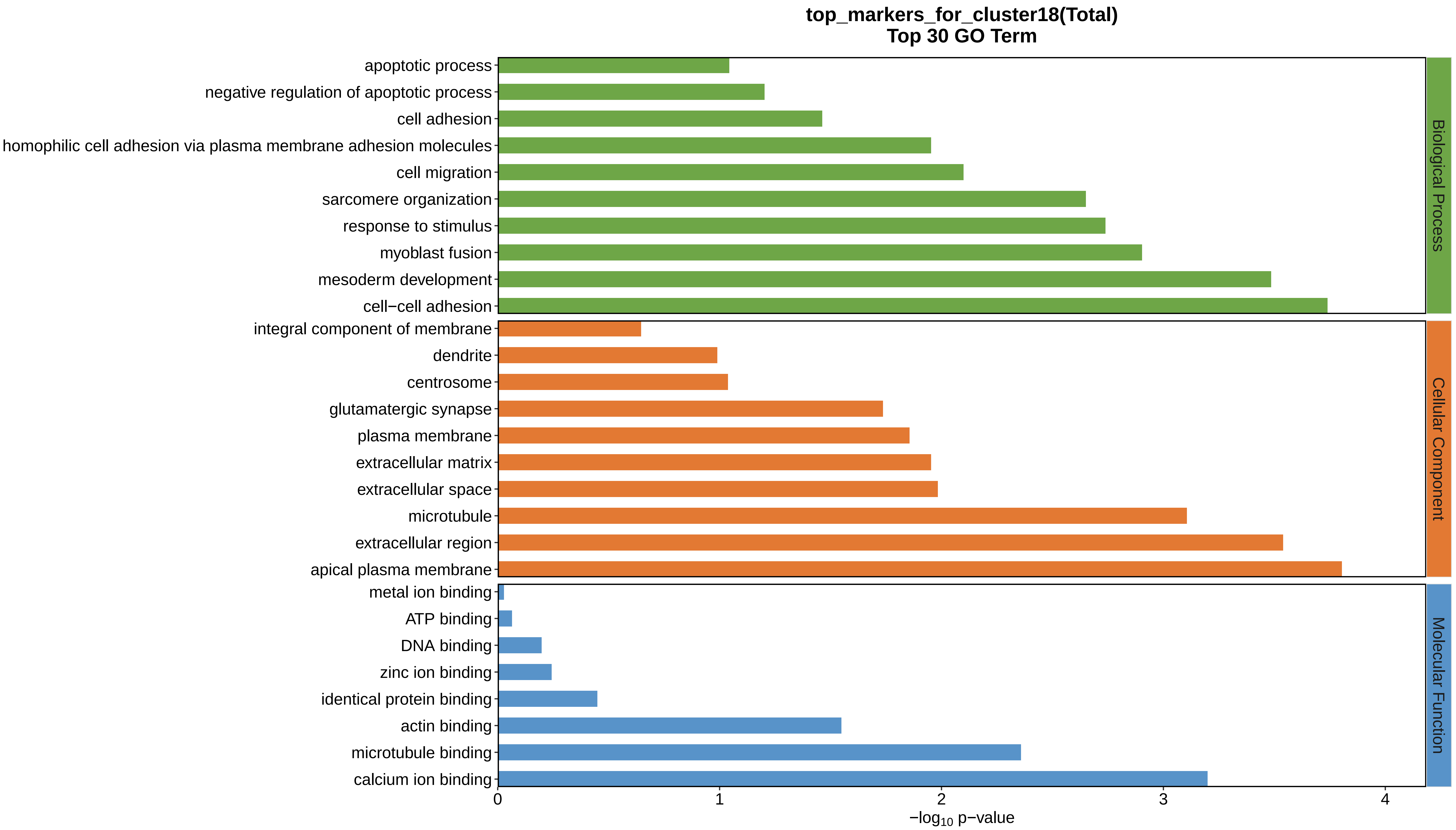
top_markers_for_cluster18(Total)
Top 30 GO Term
apoptotic process
negative regulation of apoptotic process
cell adhesion
homophilic cell adhesion via plasma membrane adhesion molecules
cell migration
sarcomere organization
response to stimulus
myoblast fusion
mesoderm development
cell−cell adhesion
integral component of membrane
dendrite
centrosome
glutamatergic synapse
plasma membrane
extracellular matrix
extracellular space
microtubule
extracellular region
apical plasma membrane
metal ion binding
ATP binding
DNA binding
zinc ion binding
identical protein binding
actin binding
microtubule binding
calcium ion binding
Biological Process
Cellular Component
Molecular Function
0
1
2
3
4
−log10 p−value
