## Supplemental Data 1-8 for "A Spatiotemporal Embryonic Atlas of Bombyx mori Integrating Single-Nucleus RNA Sequencing and Spatial Transcriptomics": Cluster19 GO top100.pdf

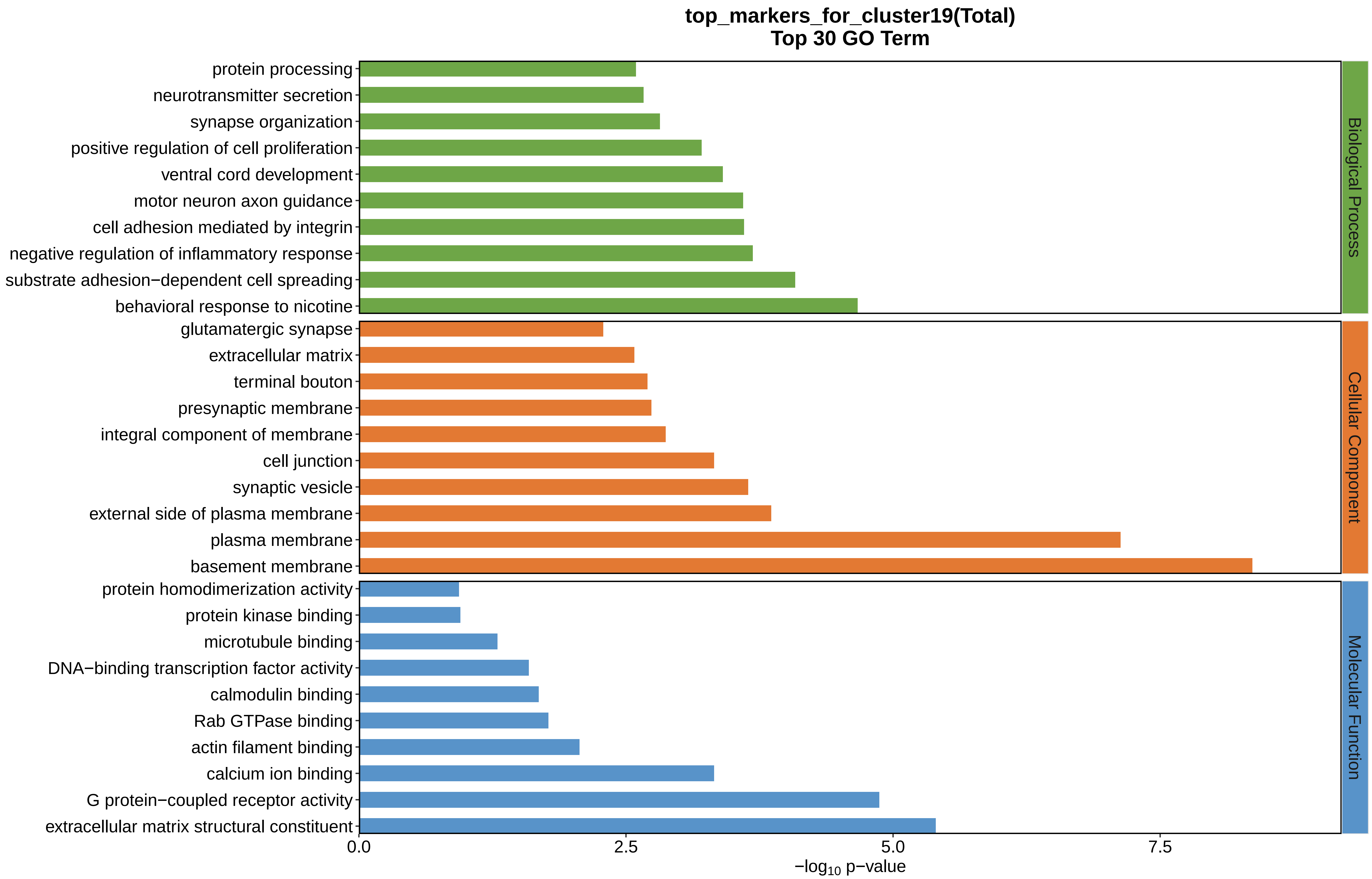

top_markers_for_cluster19(Total)
Top 30 GO Term
protein processing
neurotransmitter secretion
synapse organization
positive regulation of cell proliferation
ventral cord development
motor neuron axon guidance
cell adhesion mediated by integrin
negative regulation of inflammatory response
substrate adhesion−dependent cell spreading
behavioral response to nicotine
glutamatergic synapse
extracellular matrix
terminal bouton
presynaptic membrane
integral component of membrane
cell junction
synaptic vesicle
external side of plasma membrane
plasma membrane
basement membrane
protein homodimerization activity
protein kinase binding
microtubule binding
DNA−binding transcription factor activity
calmodulin binding
Rab GTPase binding
actin filament binding
calcium ion binding
G protein−coupled receptor activity
extracellular matrix structural constituent
Biological Process
Cellular Component
Molecular Function
0.0
2.5
5.0
7.5
−log10 p−value
