## Supplemental Data 1-8 for "A Spatiotemporal Embryonic Atlas of Bombyx mori Integrating Single-Nucleus RNA Sequencing and Spatial Transcriptomics": Cluster20 GO top100.pdf

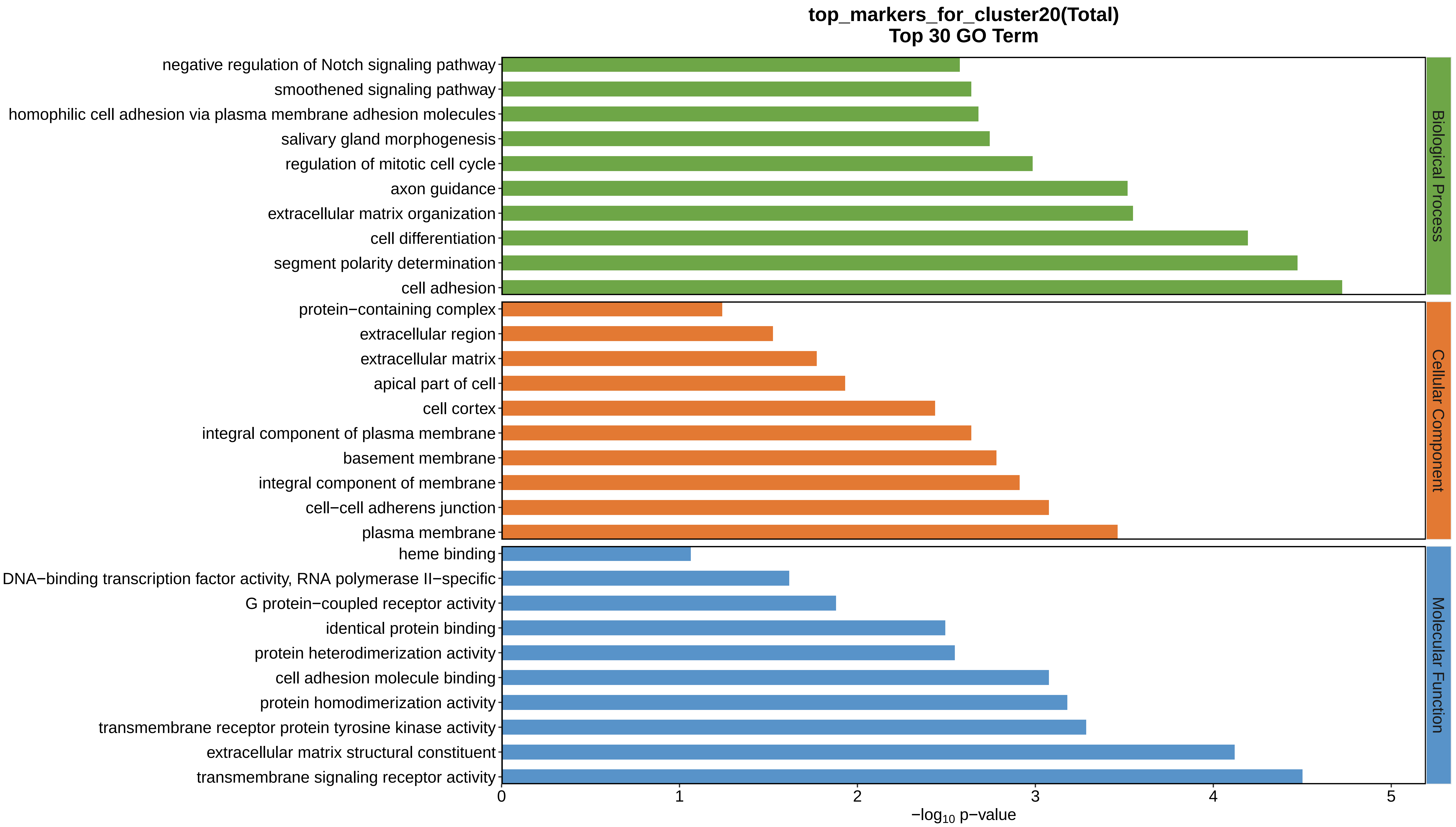
top_markers_for_cluster20(Total)
Top 30 GO Term
negative regulation of Notch signaling pathway
smoothened signaling pathway
homophilic cell adhesion via plasma membrane adhesion molecules
salivary gland morphogenesis
regulation of mitotic cell cycle
axon guidance
extracellular matrix organization
cell differentiation
segment polarity determination
cell adhesion
protein−containing complex
extracellular region
extracellular matrix
apical part of cell
cell cortex
integral component of plasma membrane
basement membrane
integral component of membrane
cell−cell adherens junction
plasma membrane
heme binding
DNA−binding transcription factor activity, RNA polymerase II−specific
G protein−coupled receptor activity
identical protein binding
protein heterodimerization activity
cell adhesion molecule binding
protein homodimerization activity
transmembrane receptor protein tyrosine kinase activity
extracellular matrix structural constituent
transmembrane signaling receptor activity
Biological Process
Cellular Component
Molecular Function
0
1
2
3
4
5
$-\log_{10}$ p−value
