## Supplemental Data 1-8 for "A Spatiotemporal Embryonic Atlas of Bombyx mori Integrating Single-Nucleus RNA Sequencing and Spatial Transcriptomics": Cluster21 GO top100.pdf

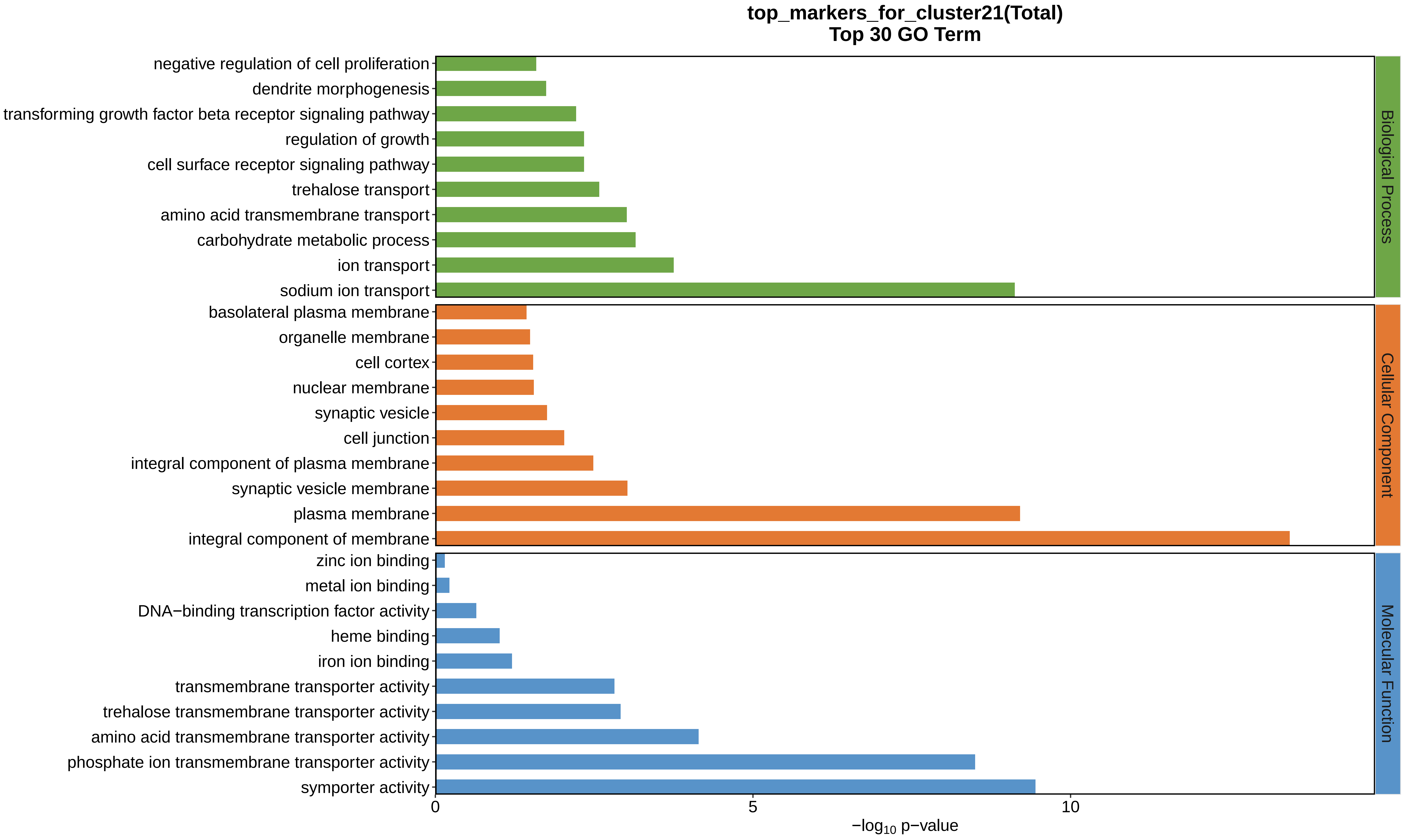
top_markers_for_cluster21(Total)
Top 30 GO Term
negative regulation of cell proliferation
dendrite morphogenesis
transforming growth factor beta receptor signaling pathway
regulation of growth
cell surface receptor signaling pathway
trehalose transport
amino acid transmembrane transport
carbohydrate metabolic process
ion transport
sodium ion transport
basolateral plasma membrane
organelle membrane
cell cortex
nuclear membrane
synaptic vesicle
cell junction
integral component of plasma membrane
synaptic vesicle membrane
plasma membrane
integral component of membrane
zinc ion binding
metal ion binding
DNA−binding transcription factor activity
heme binding
iron ion binding
transmembrane transporter activity
trehalose transmembrane transporter activity
amino acid transmembrane transporter activity
phosphate ion transmembrane transporter activity
symporter activity
Biological Process
Cellular Component
Molecular Function
0
5
10
−log10 p−value
