## Supplemental Data 1-8 for "A Spatiotemporal Embryonic Atlas of Bombyx mori Integrating Single-Nucleus RNA Sequencing and Spatial Transcriptomics": Cluster22 GO top100.pdf

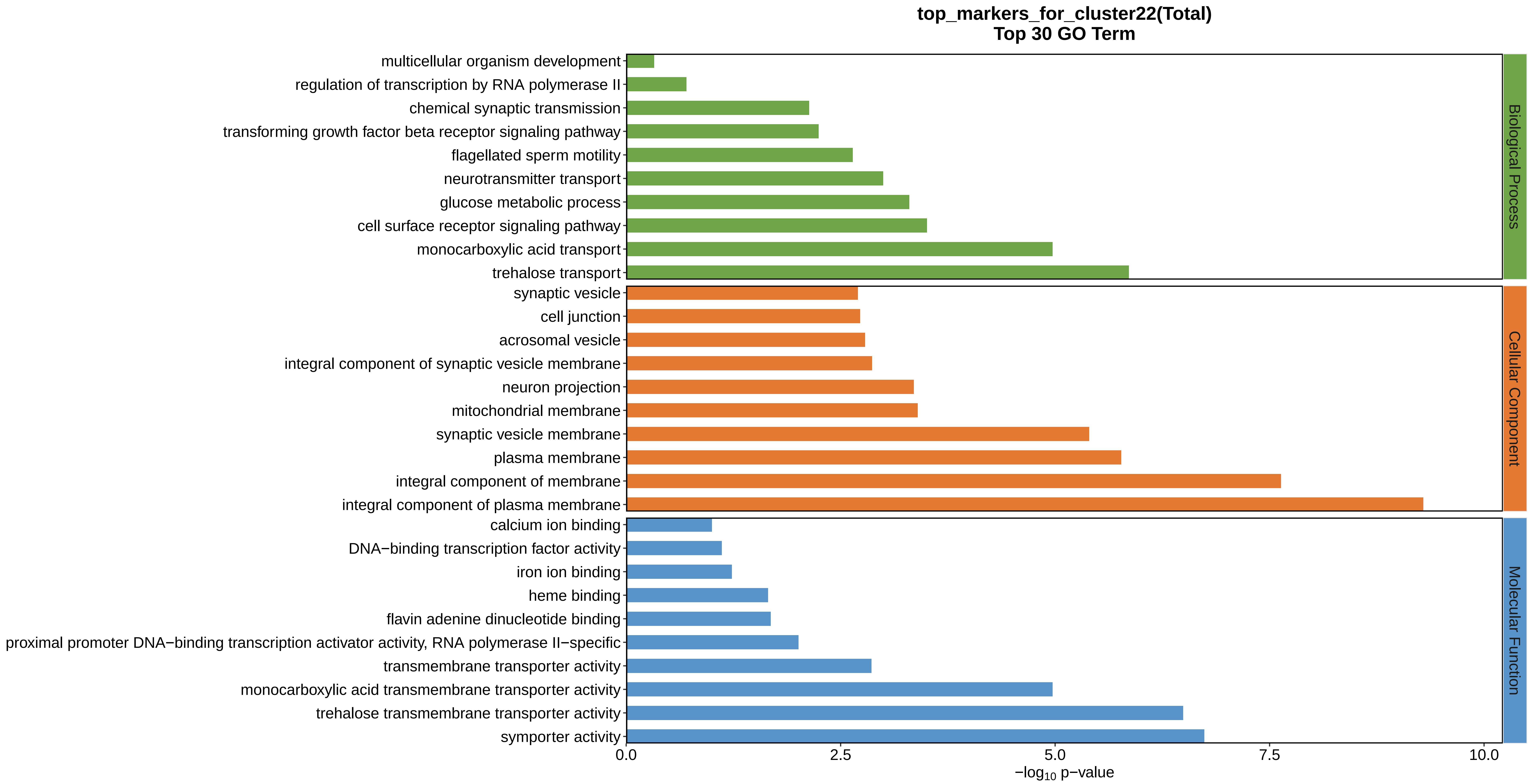
top_markers_for_cluster22(Total)
Top 30 GO Term
multicellular organism development
regulation of transcription by RNA polymerase II
chemical synaptic transmission
transforming growth factor beta receptor signaling pathway
flagellated sperm motility
neurotransmitter transport
glucose metabolic process
cell surface receptor signaling pathway
monocarboxylic acid transport
trehalose transport
synaptic vesicle
cell junction
acrosomal vesicle
integral component of synaptic vesicle membrane
neuron projection
mitochondrial membrane
synaptic vesicle membrane
plasma membrane
integral component of membrane
integral component of plasma membrane
calcium ion binding
DNA−binding transcription factor activity
iron ion binding
heme binding
flavin adenine dinucleotide binding
proximal promoter DNA−binding transcription activator activity, RNA polymerase II−specific
transmembrane transporter activity
monocarboxylic acid transmembrane transporter activity
trehalose transmembrane transporter activity
symporter activity
Biological Process
Cellular Component
Molecular Function
0.0
2.5
5.0
7.5
10.0
$-\log_{10}$ p−value
