## Supplemental Data 1-8 for "A Spatiotemporal Embryonic Atlas of Bombyx mori Integrating Single-Nucleus RNA Sequencing and Spatial Transcriptomics": Cluster23 GO top100.pdf

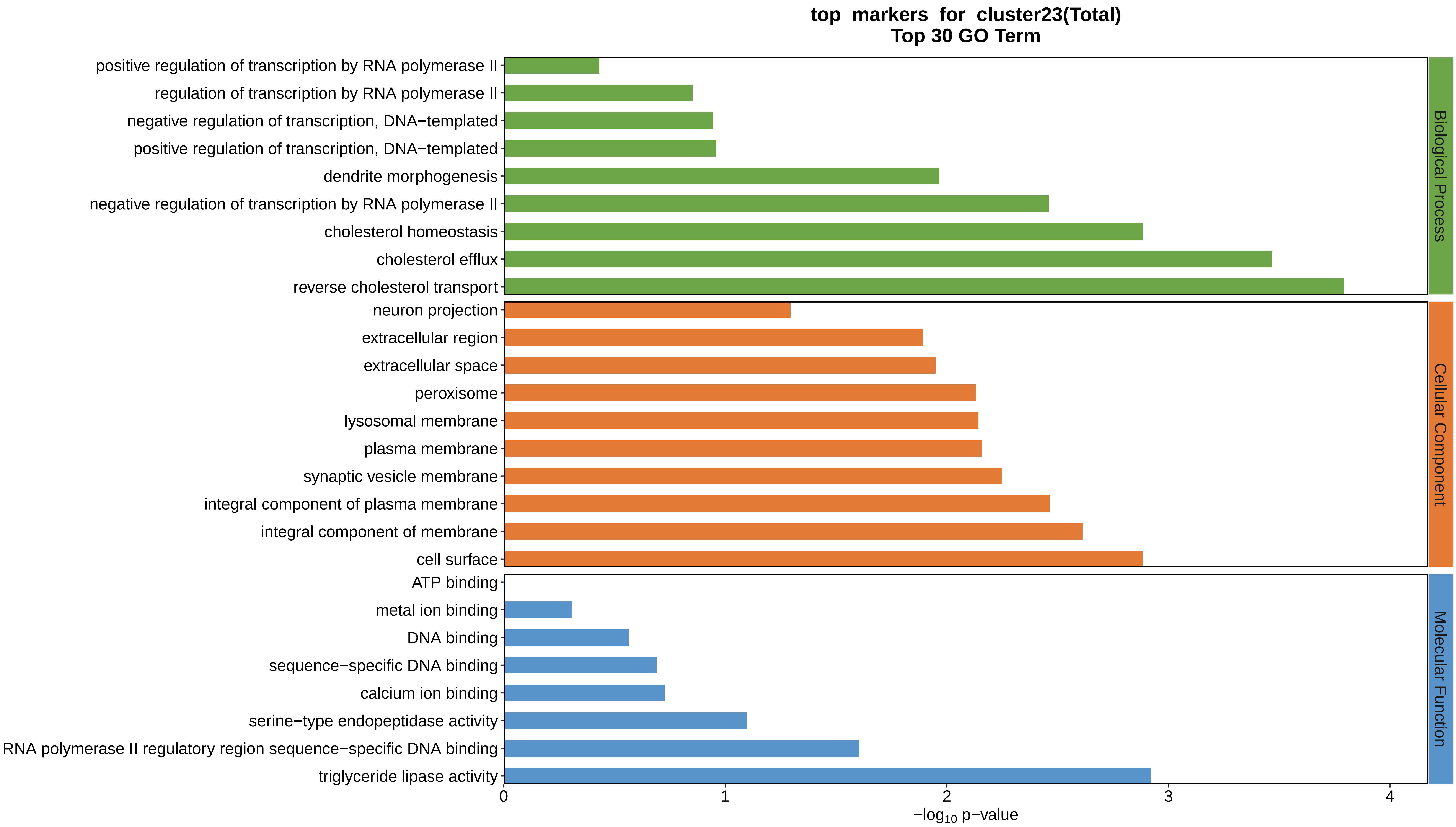

top_markers_for_cluster23(Total)
Top 30 GO Term
positive regulation of transcription by RNA polymerase II
regulation of transcription by RNA polymerase II
negative regulation of transcription, DNA−templated
positive regulation of transcription, DNA−templated
dendrite morphogenesis
negative regulation of transcription by RNA polymerase II
cholesterol homeostasis
cholesterol efflux
reverse cholesterol transport
neuron projection
extracellular region
extracellular space
peroxisome
lysosomal membrane
plasma membrane
synaptic vesicle membrane
integral component of plasma membrane
integral component of membrane
cell surface
ATP binding
metal ion binding
DNA binding
sequence−specific DNA binding
calcium ion binding
serine−type endopeptidase activity
RNA polymerase II regulatory region sequence−specific DNA binding
triglyceride lipase activity
Biological Process
Cellular Component
Molecular Function
0
1
2
3
4
−log10 p−value
