## Supplementary figures and images for "A Spatiotemporal Embryonic Atlas of Bombyx mori Integrating Single-Nucleus RNA Sequencing and Spatial Transcriptomics"

### Cluster1 GO top100.png

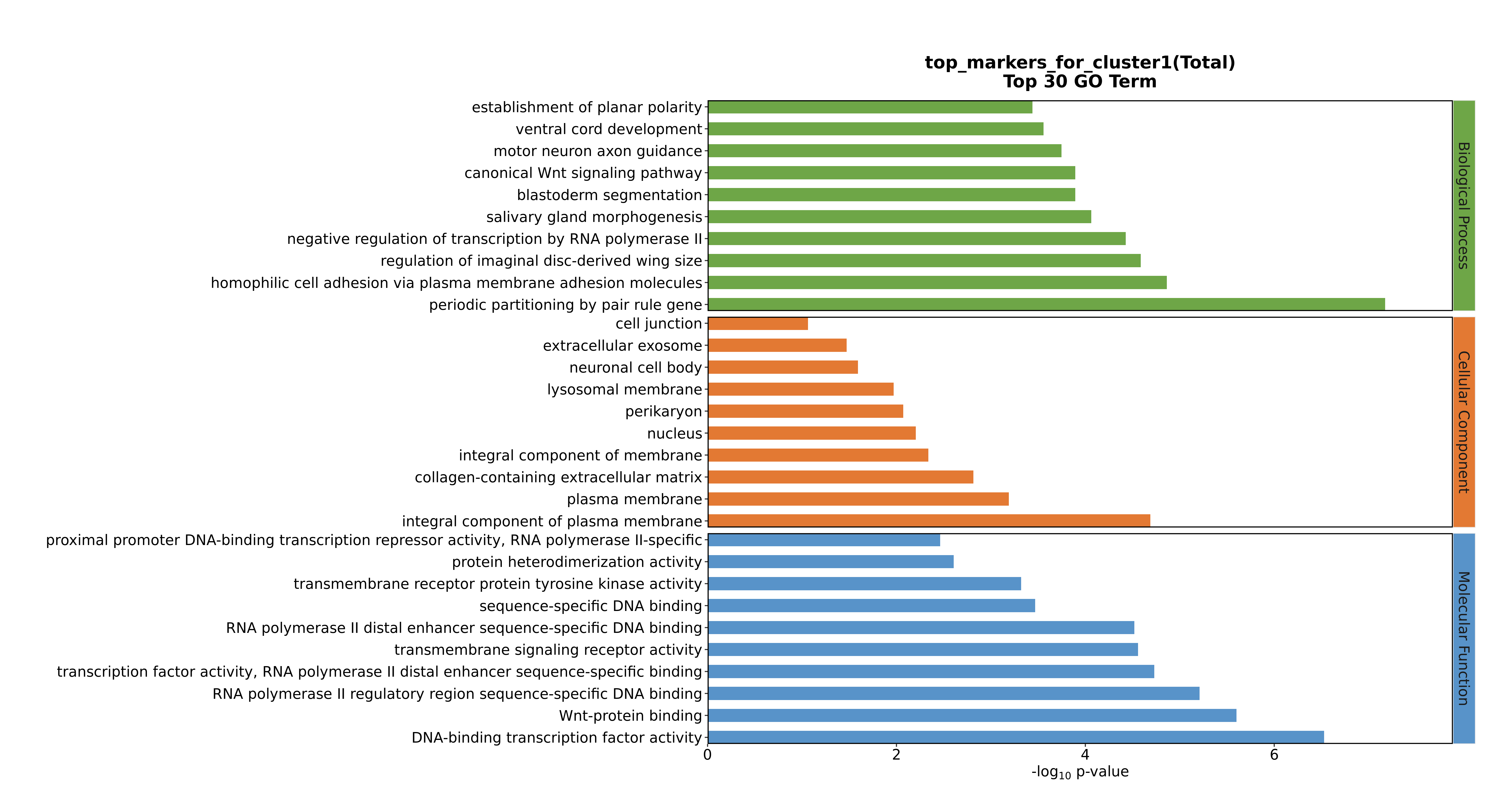

### Cluster1 GO top100.png

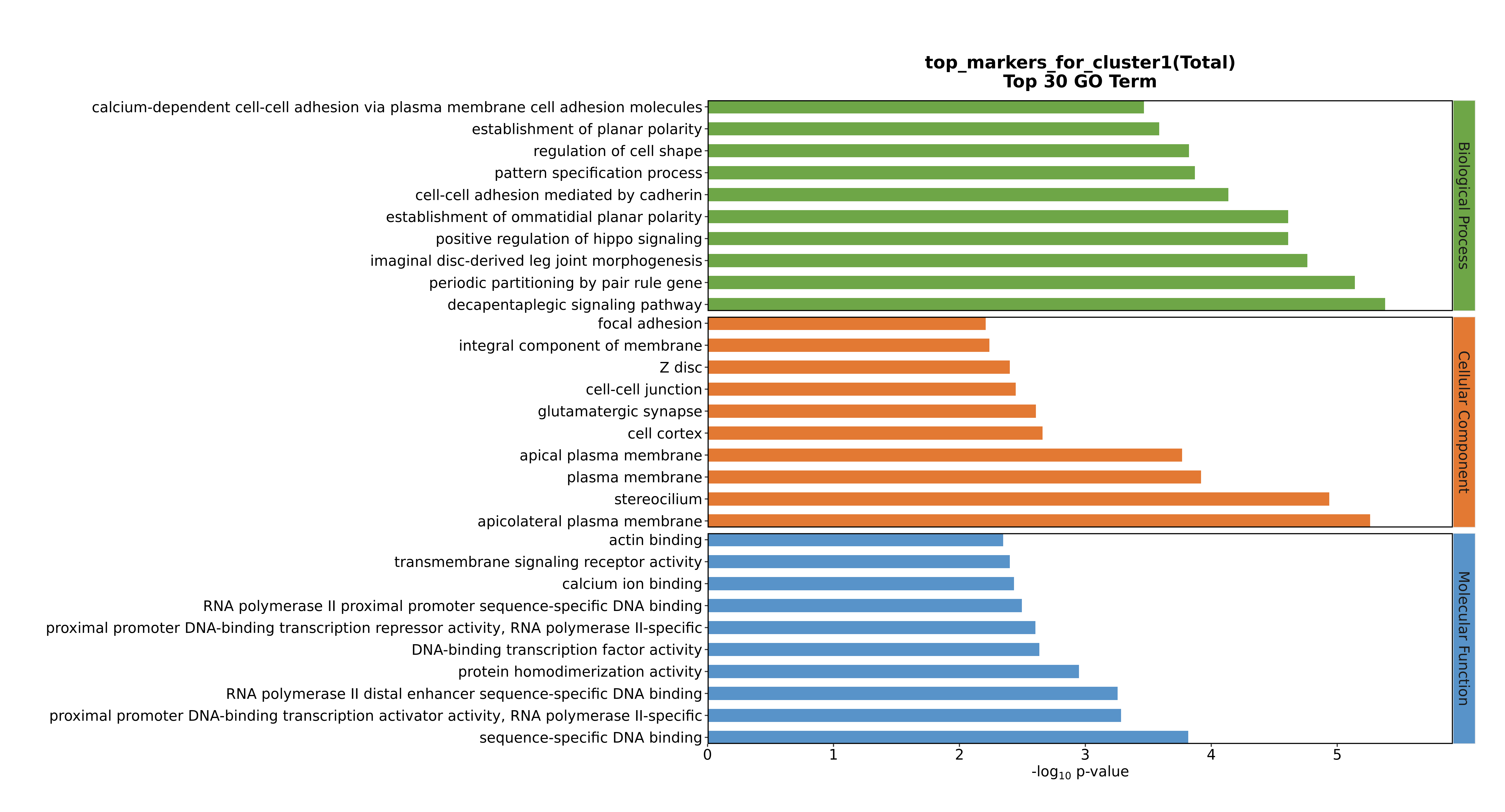

### Cluster1 GO top100.png

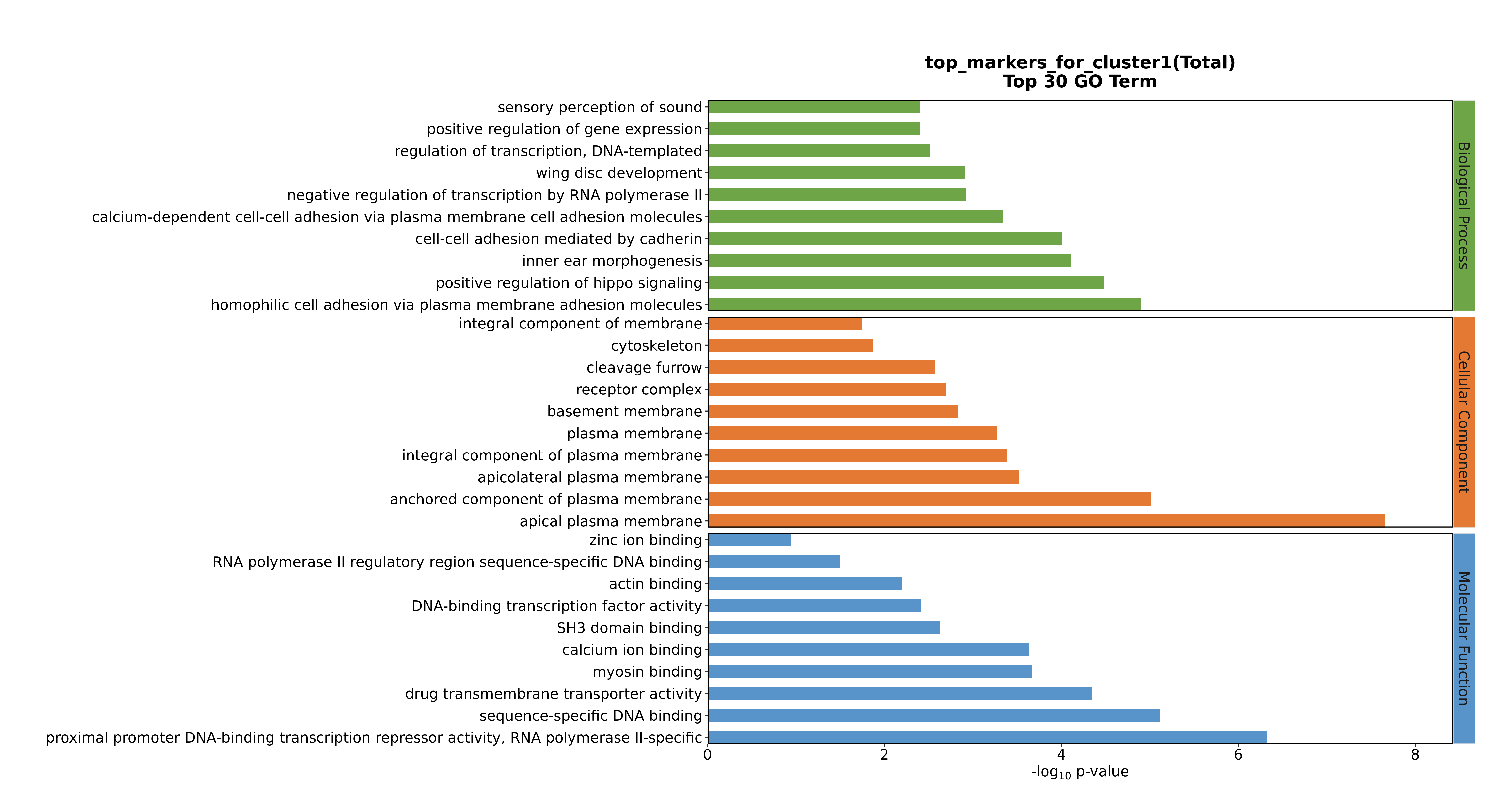

### Cluster1 GO top100.png

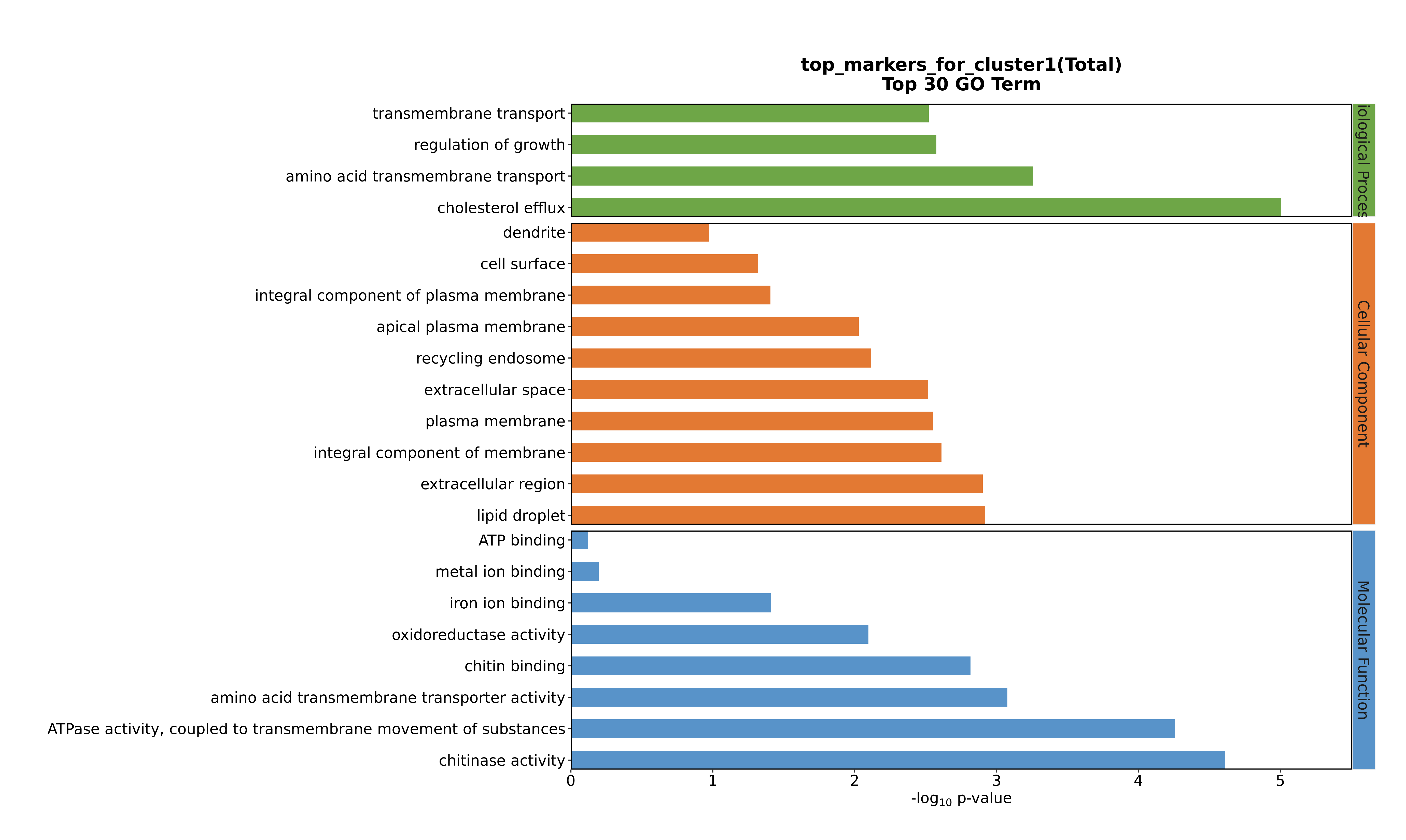

### Cluster1 GO top100.png

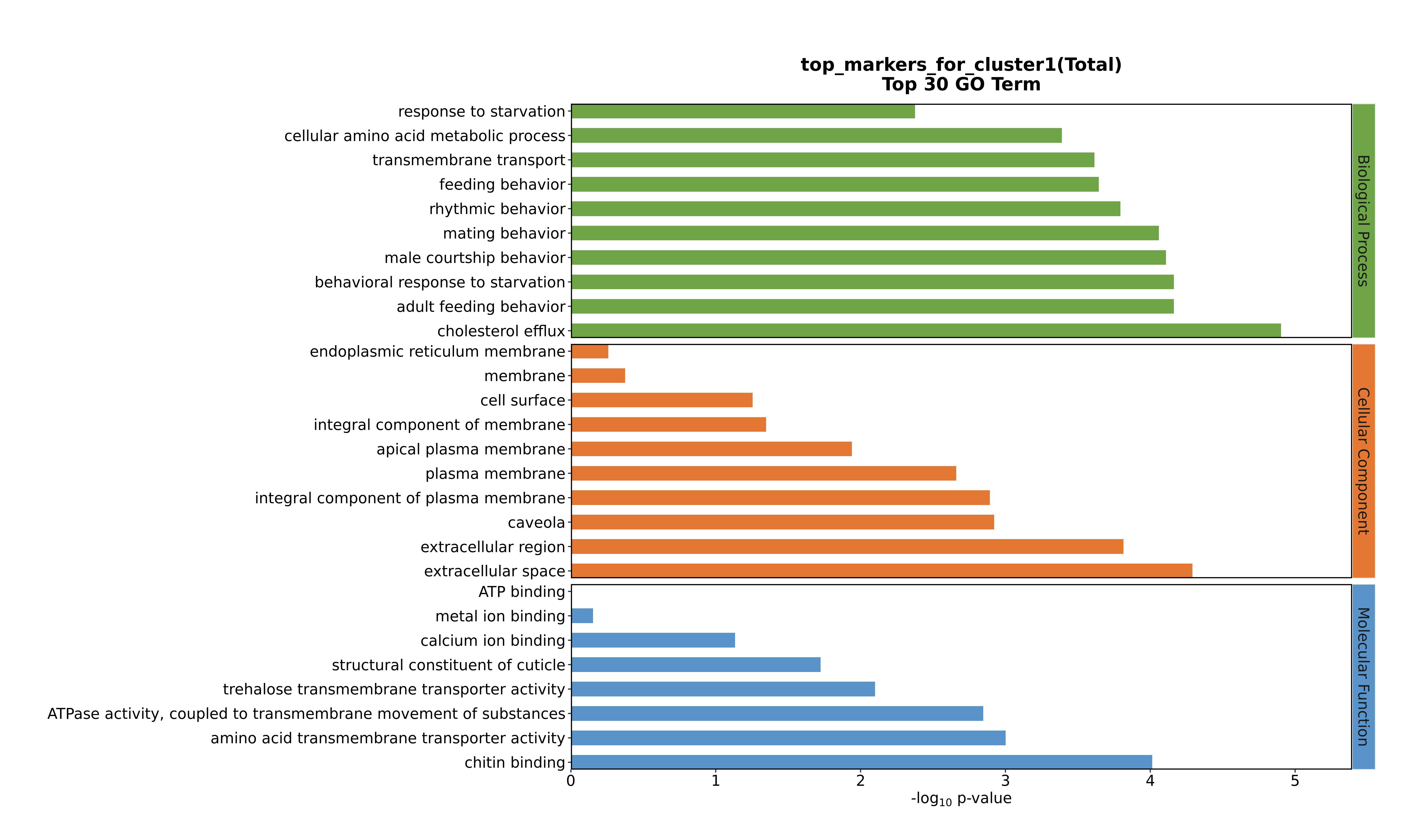

### Cluster2 GO top100.png

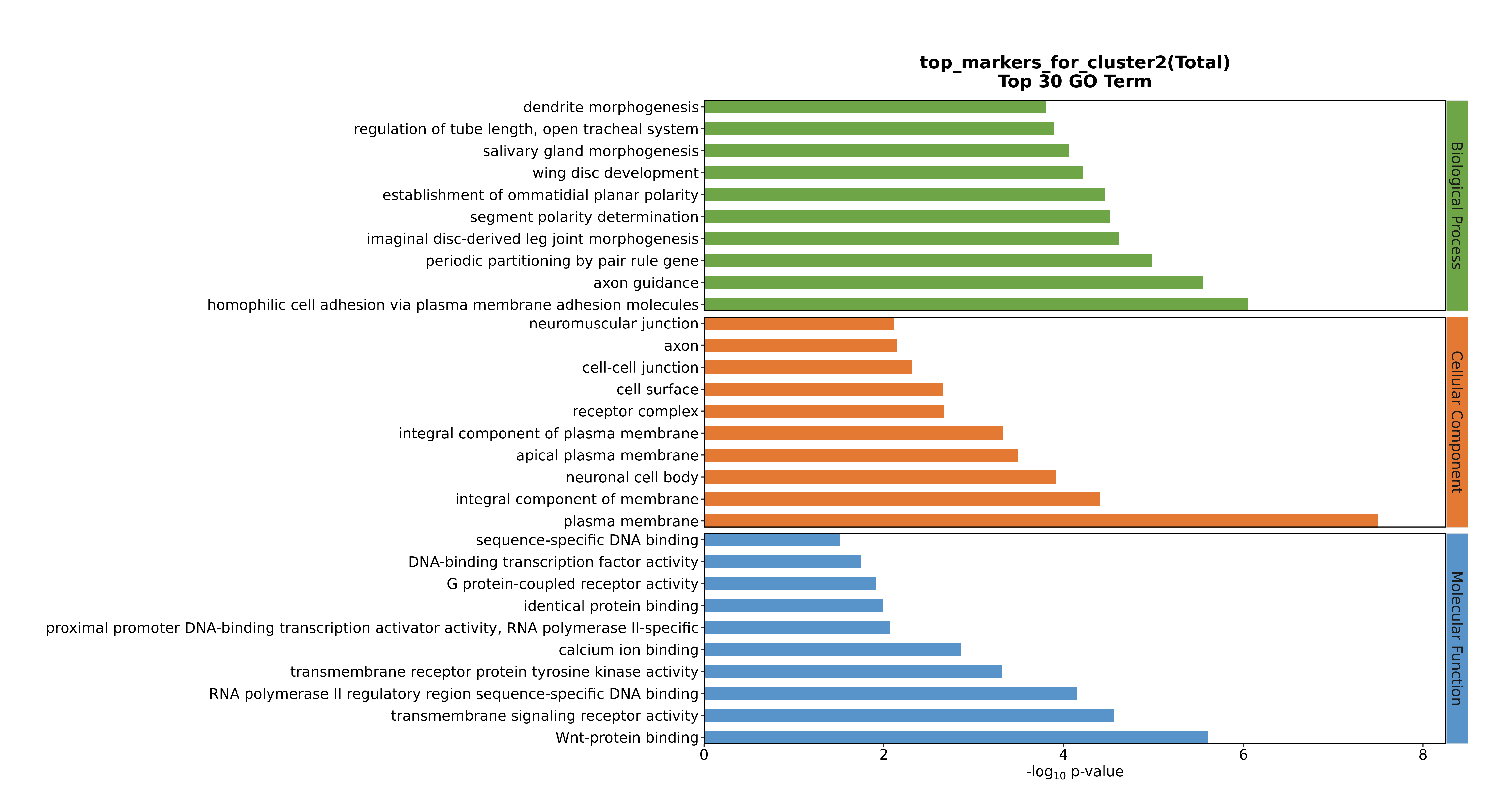

### Cluster2 GO top100.png

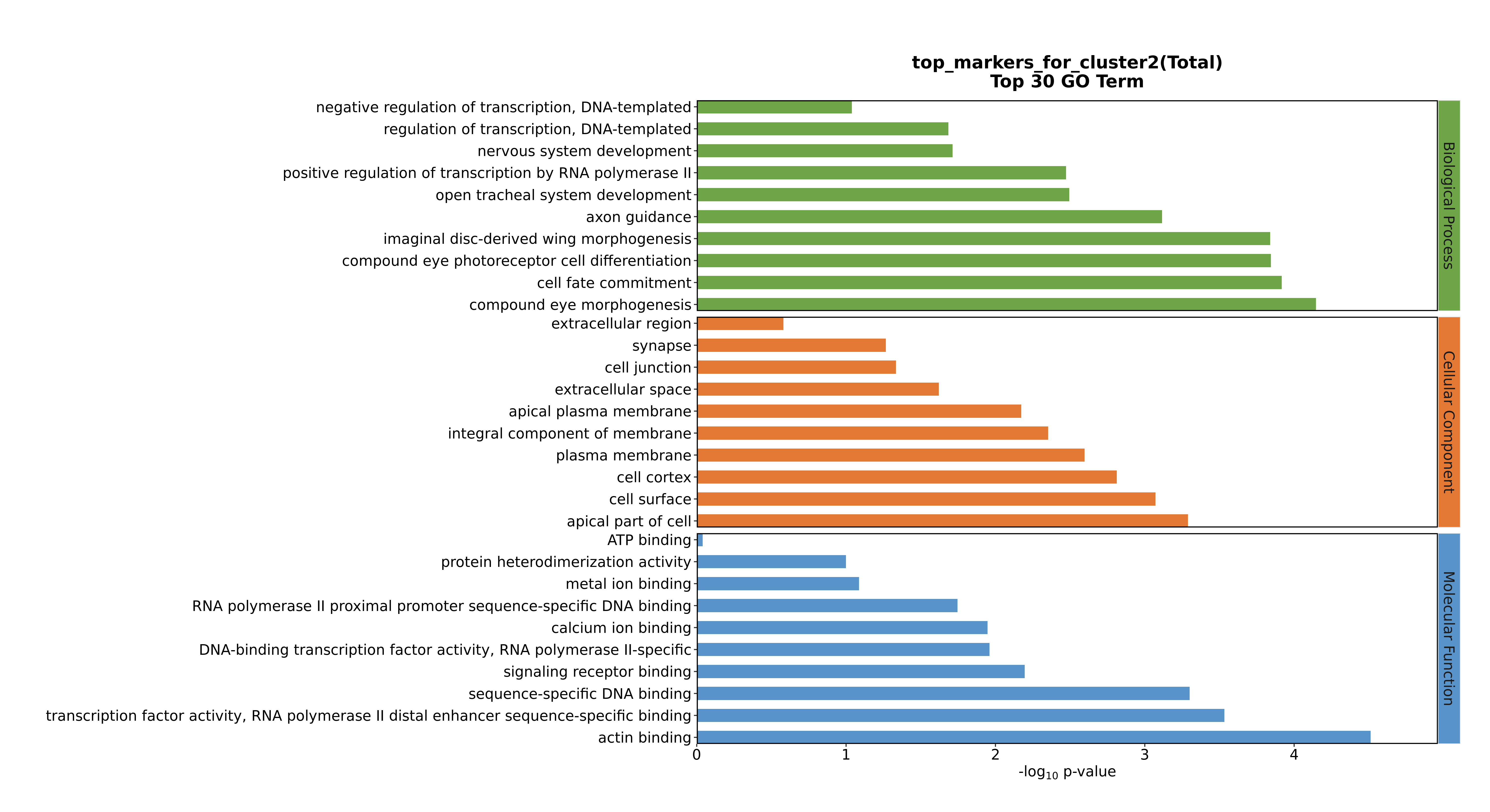

### Cluster2 GO top100.png

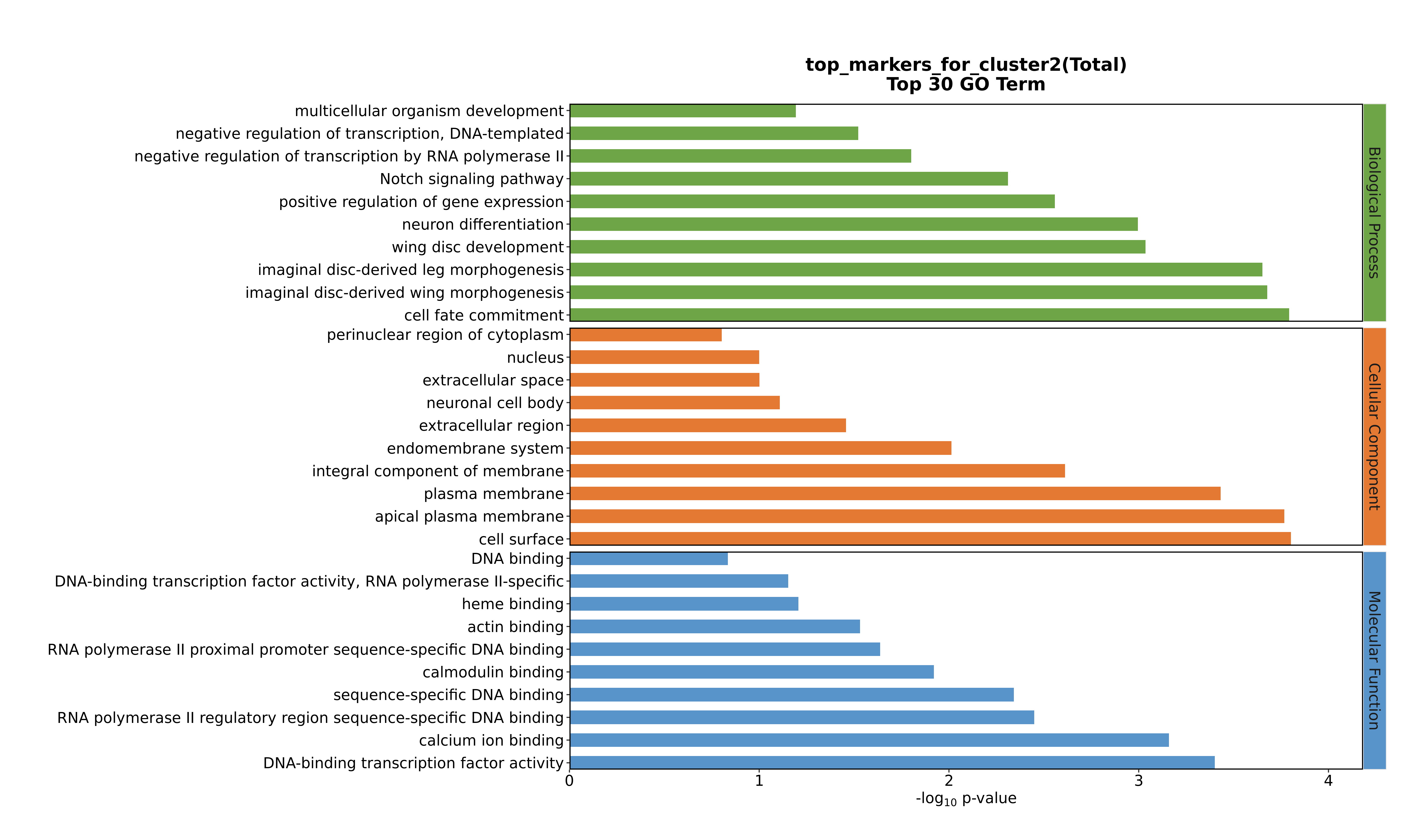

### Cluster2 GO top100.png

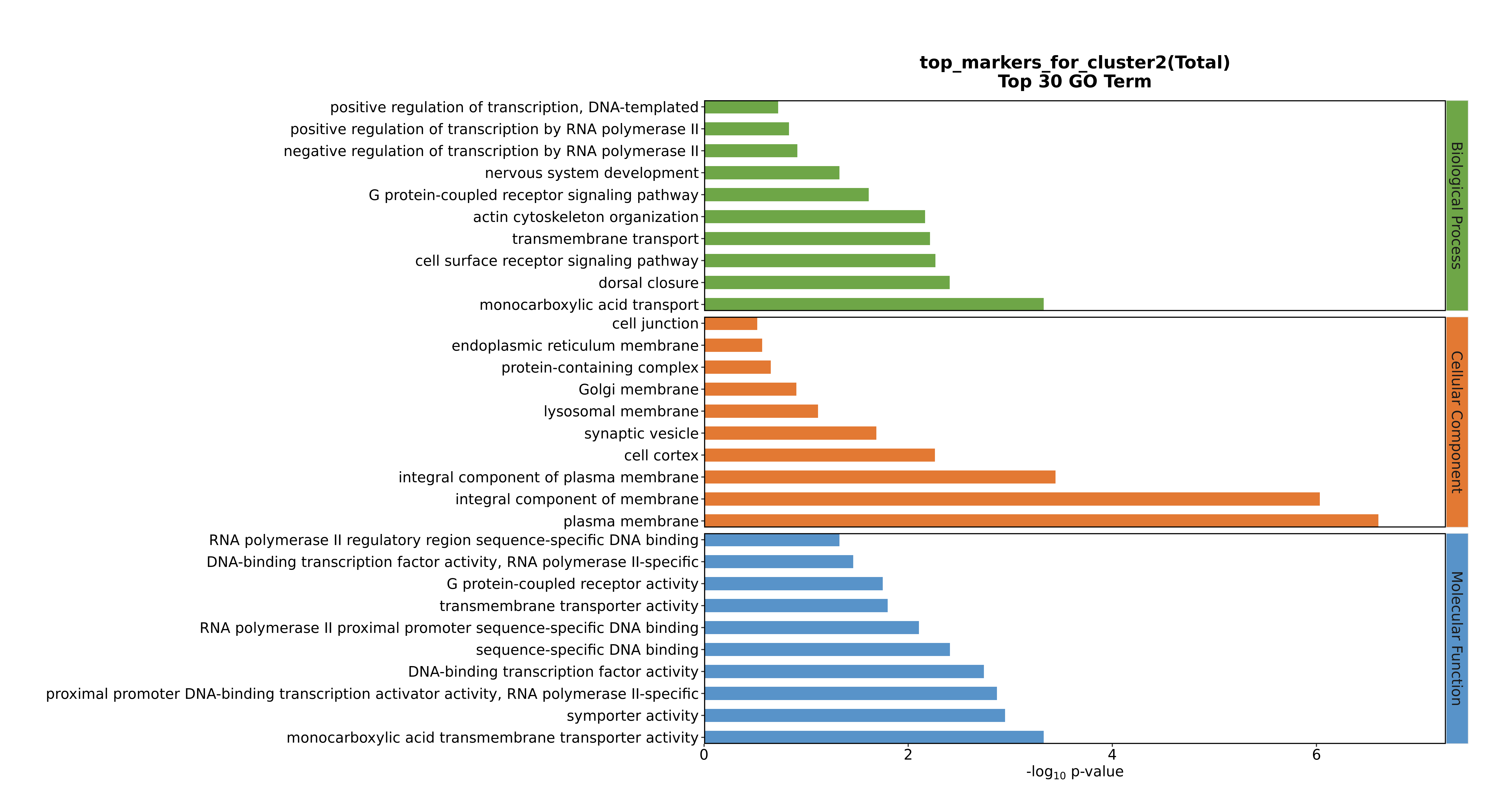

### Cluster2 GO top100.png

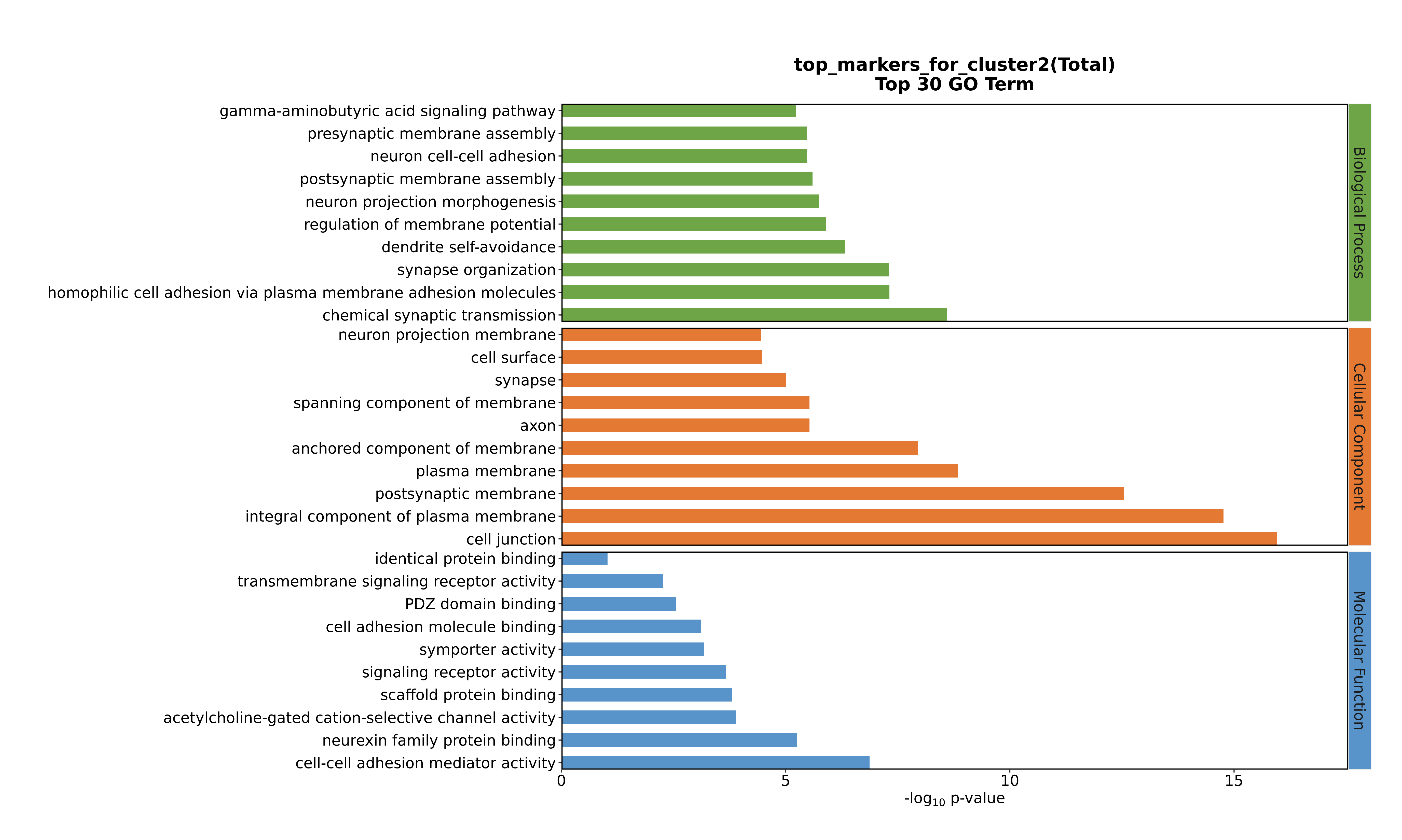

### Cluster3 GO top100.png

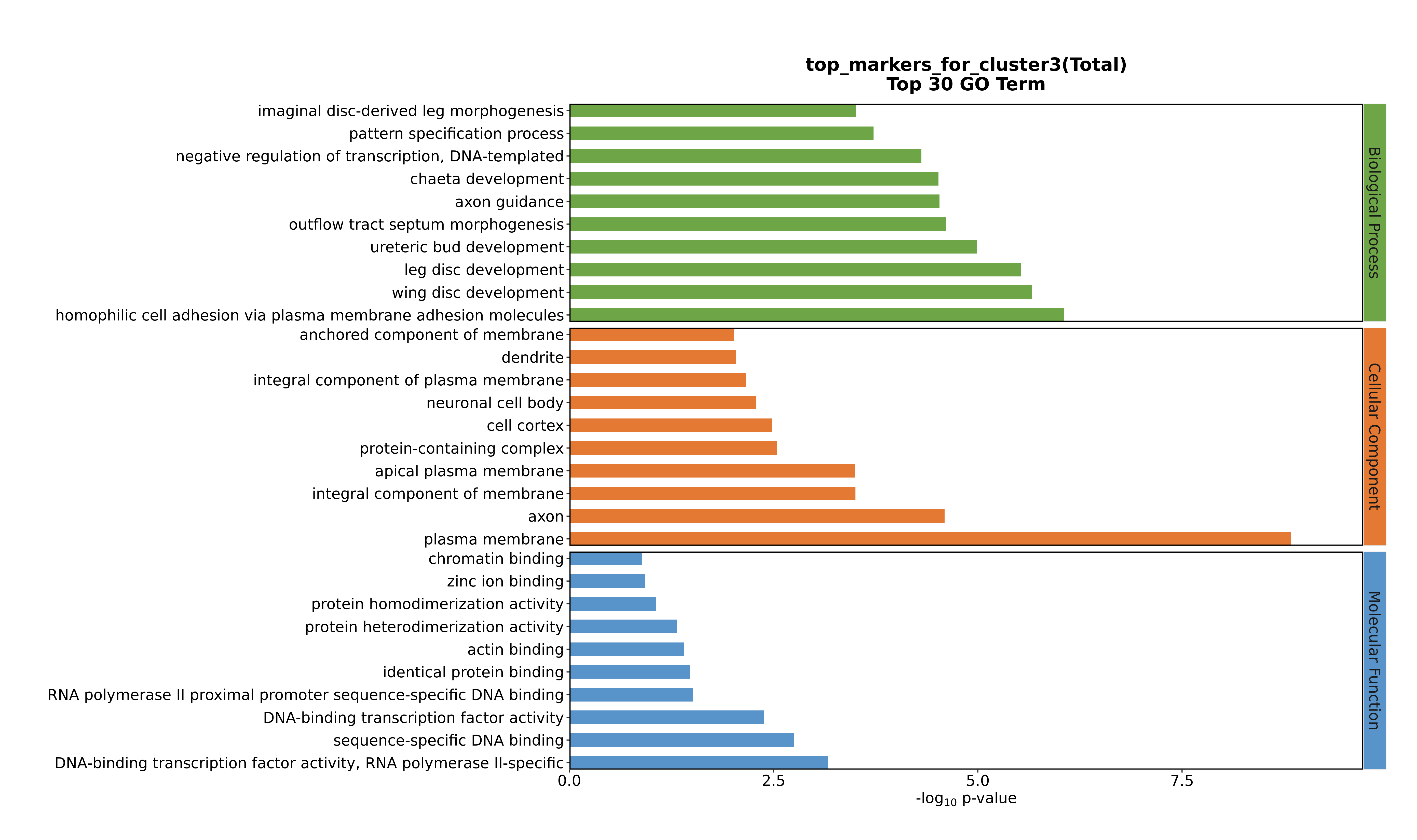

### Cluster3 GO top100.png

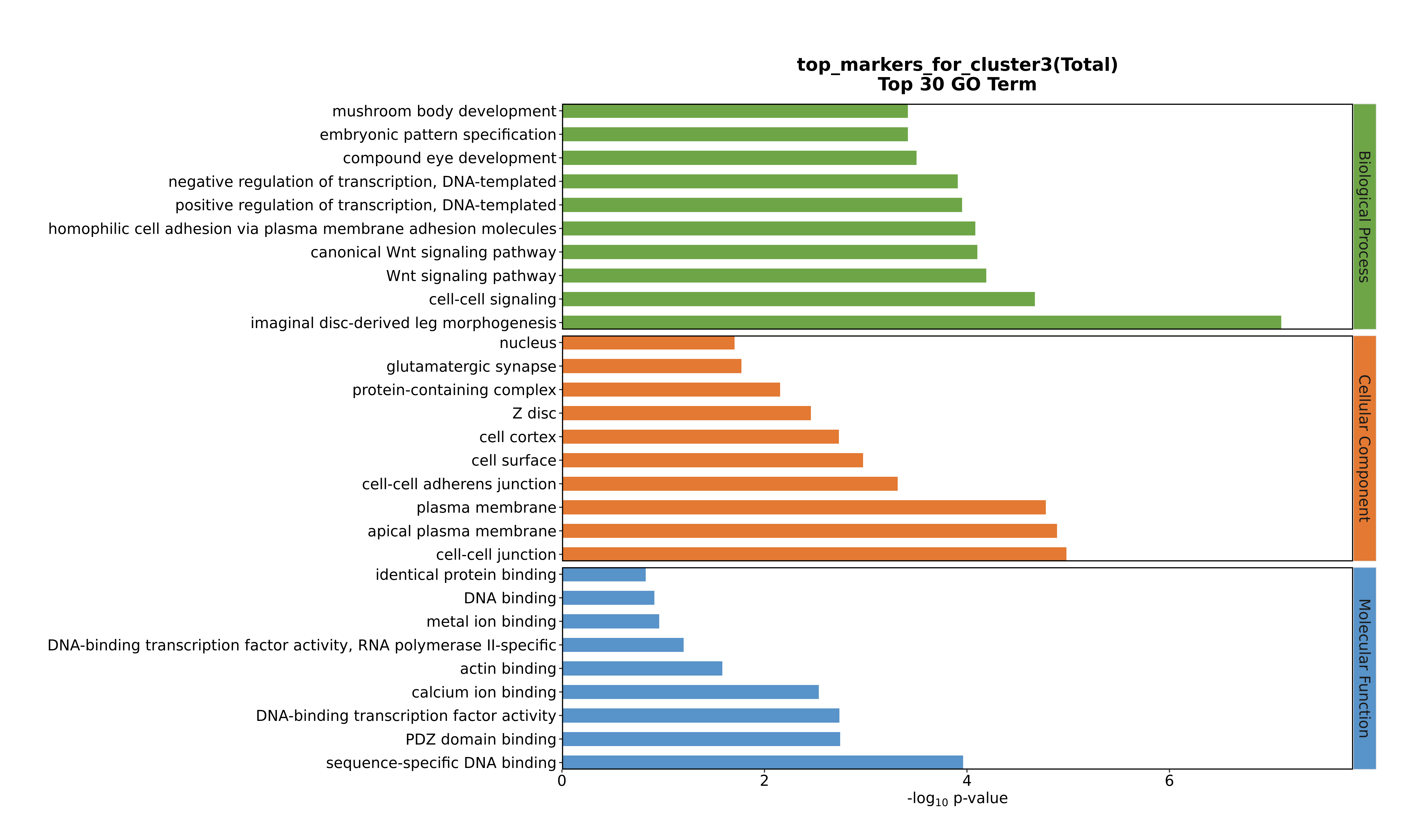

### Cluster3 GO top100.png

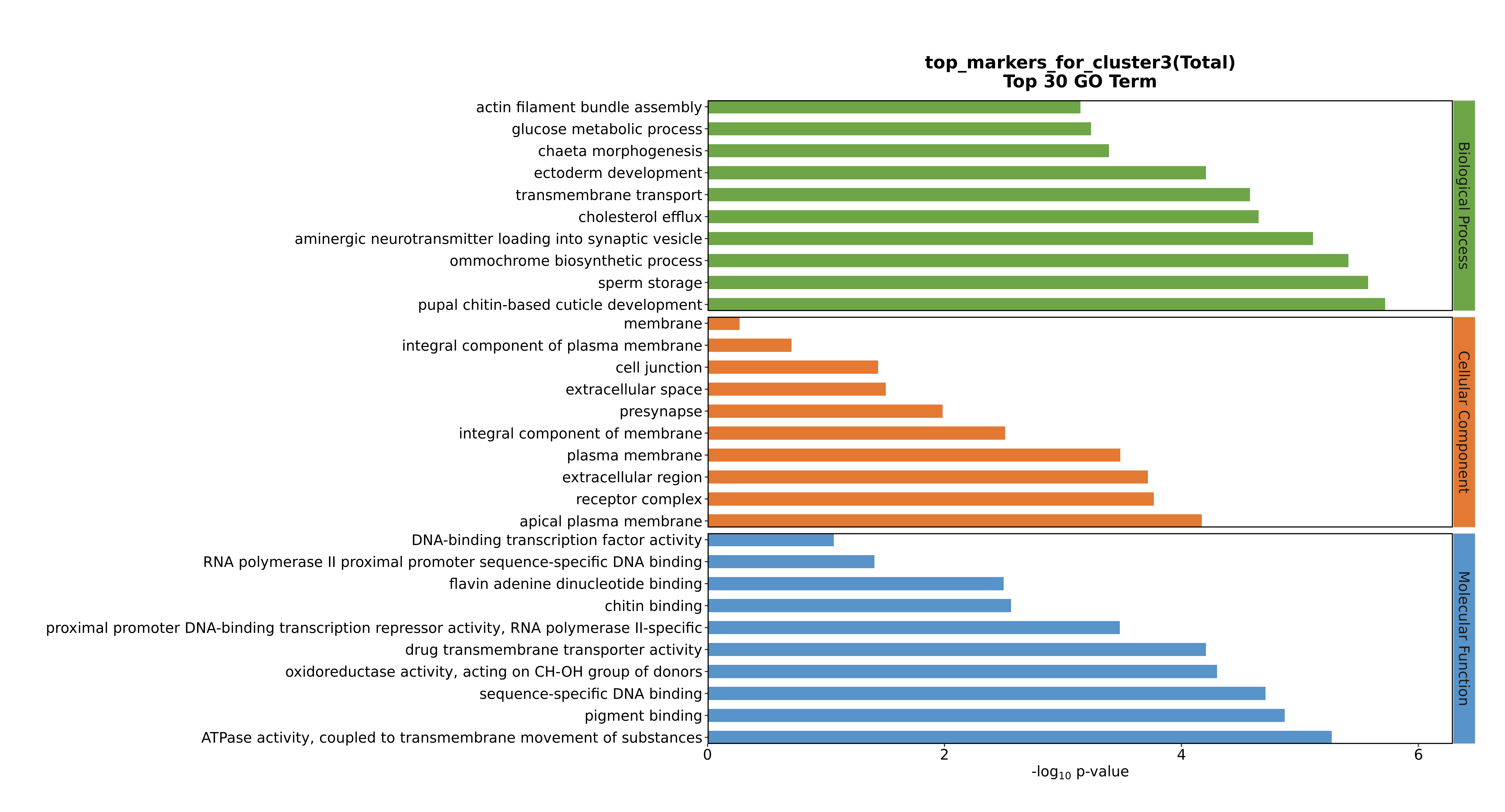

### Cluster3 GO top100.png

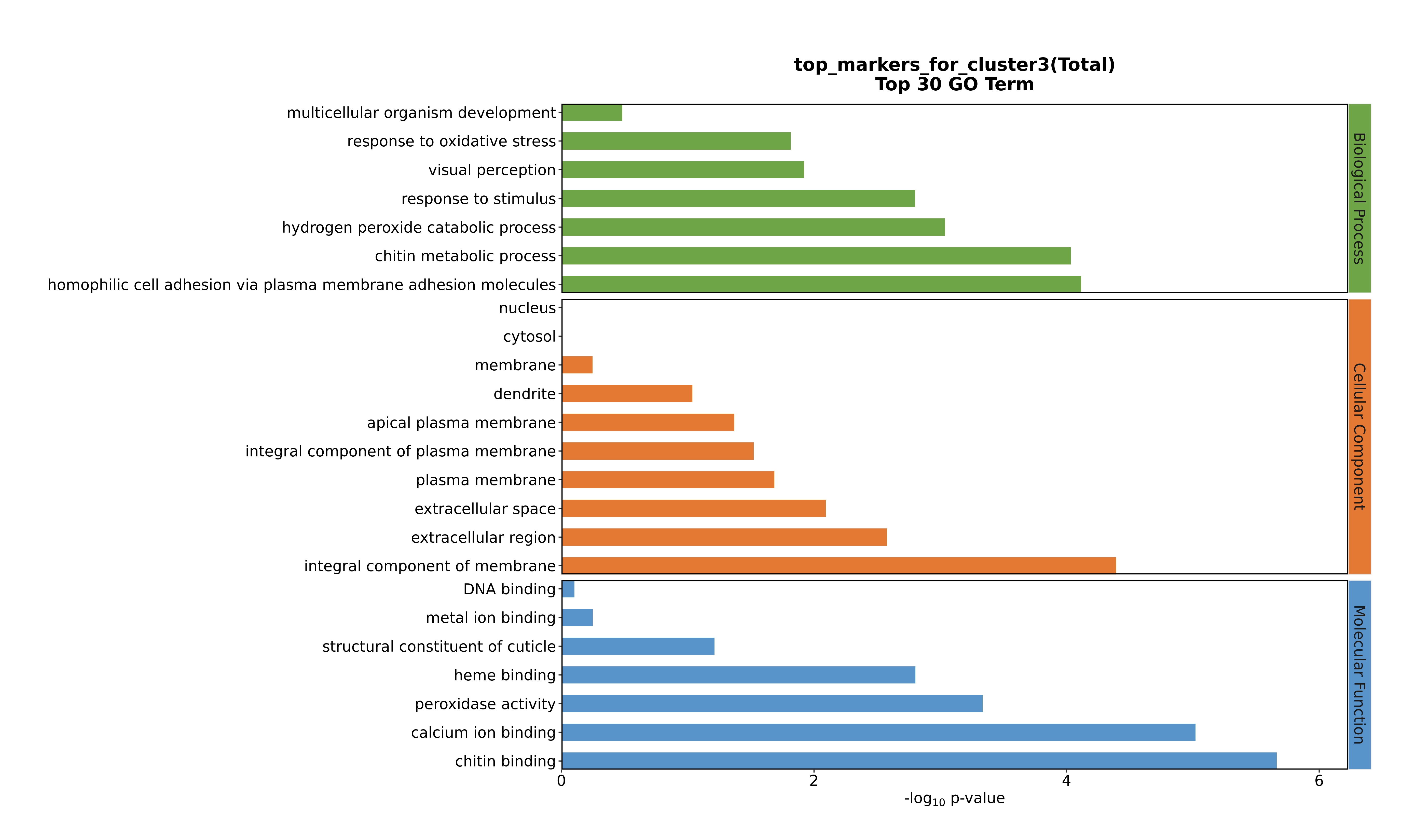

### Cluster4 GO top100.png

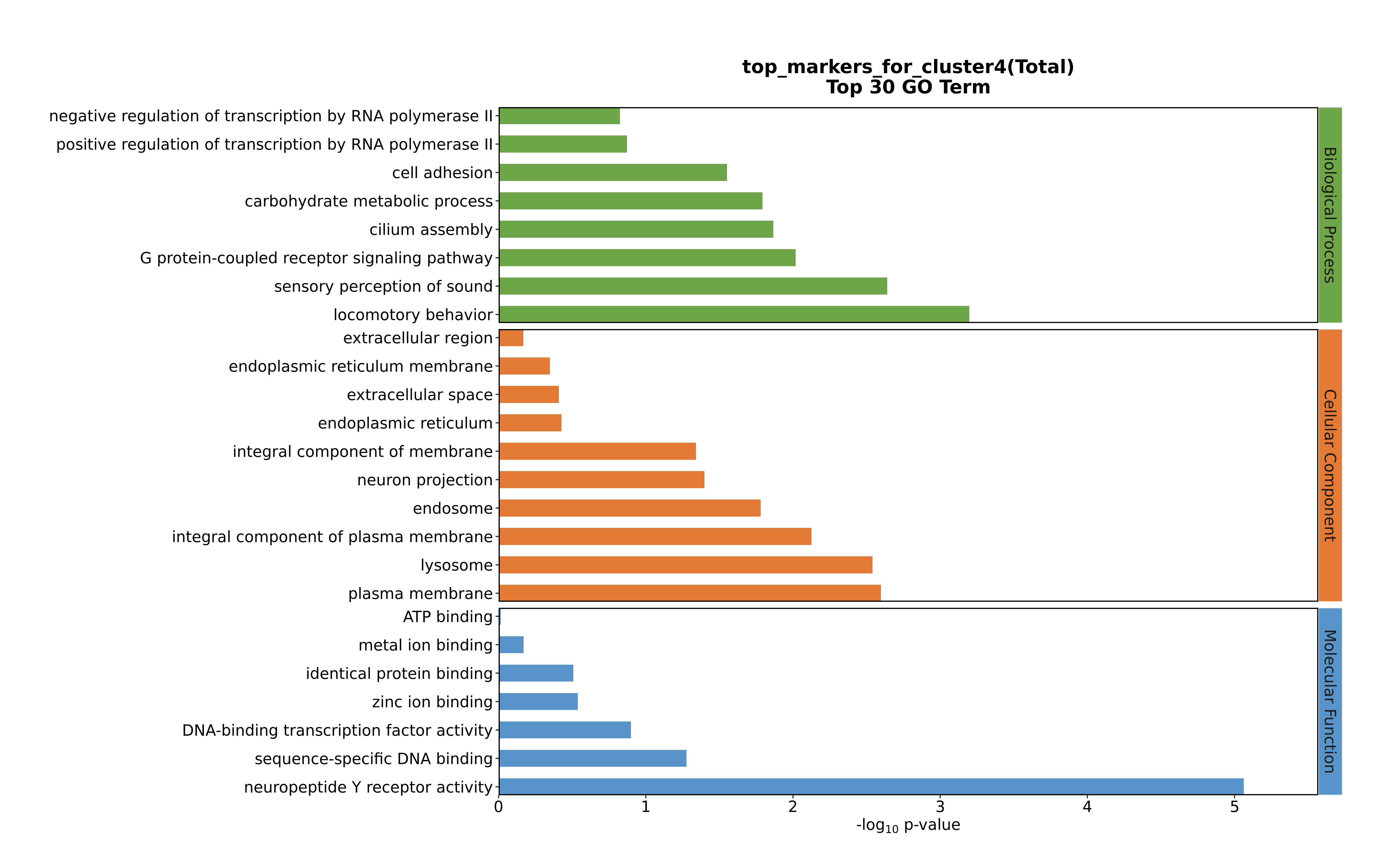

### Cluster4 GO top100.png

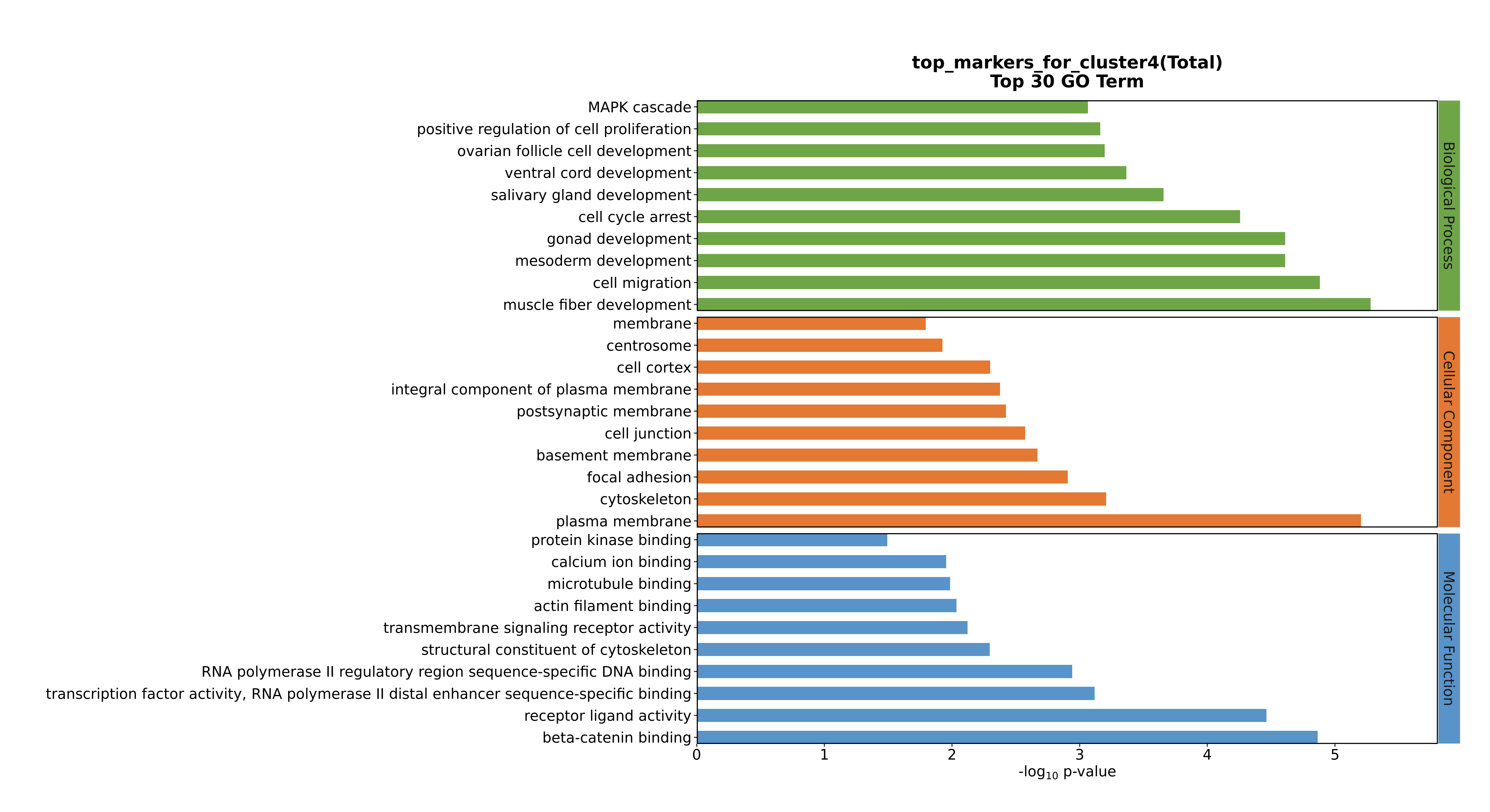

### Cluster4 GO top100.png

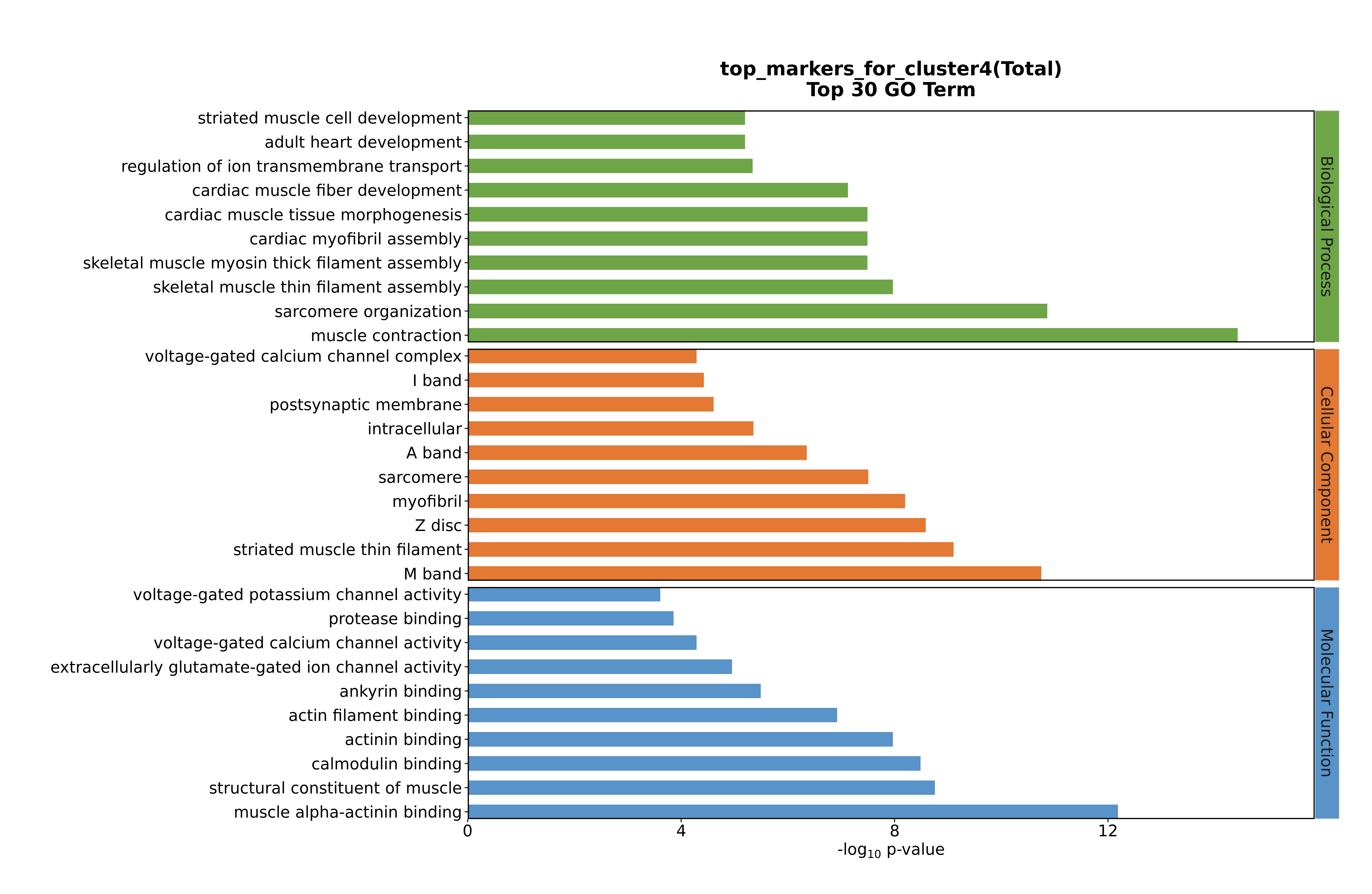

### Cluster4 GO top100.png

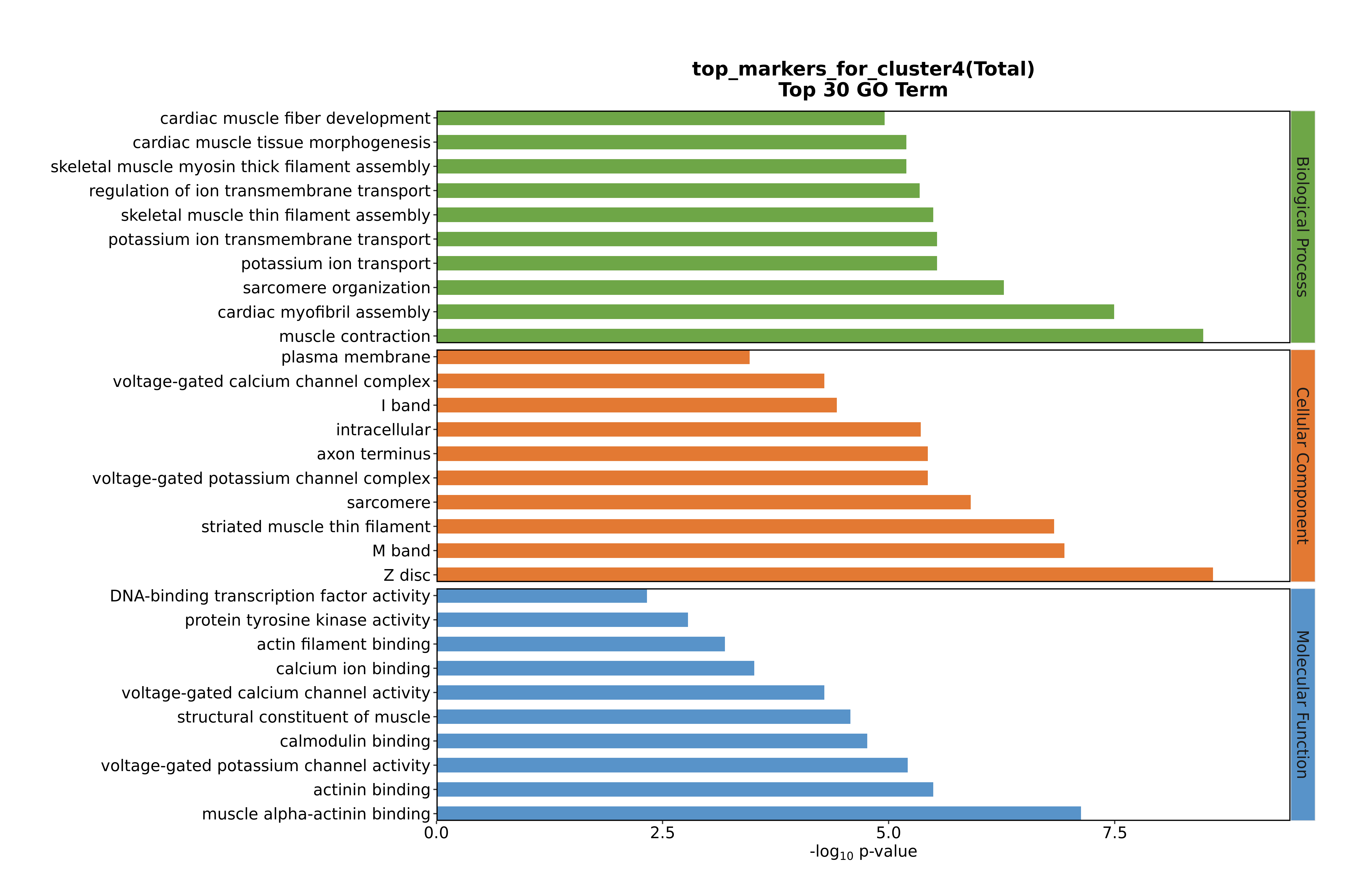

### Cluster5 GO top100.png

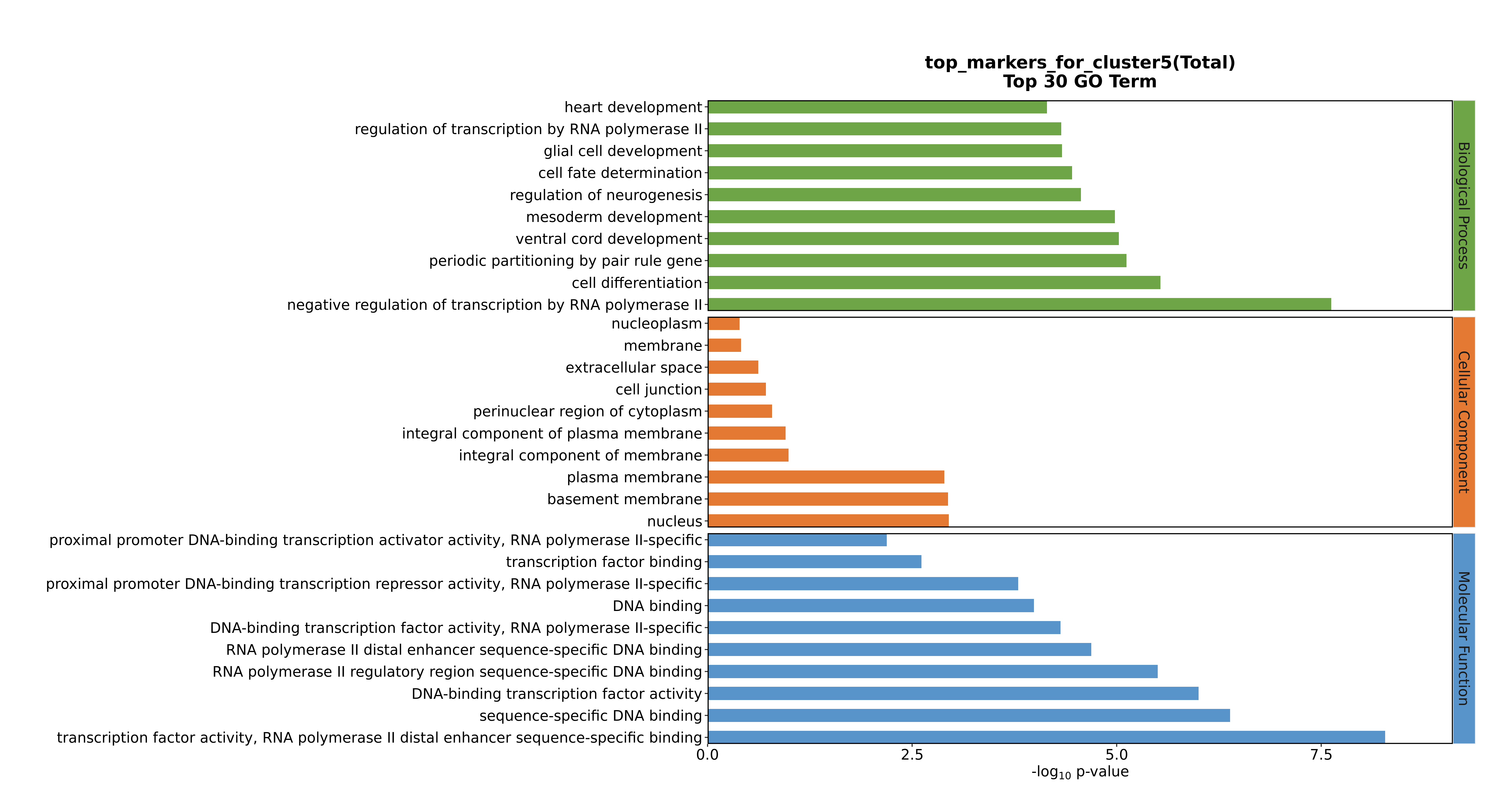

### Cluster5 GO top100.png

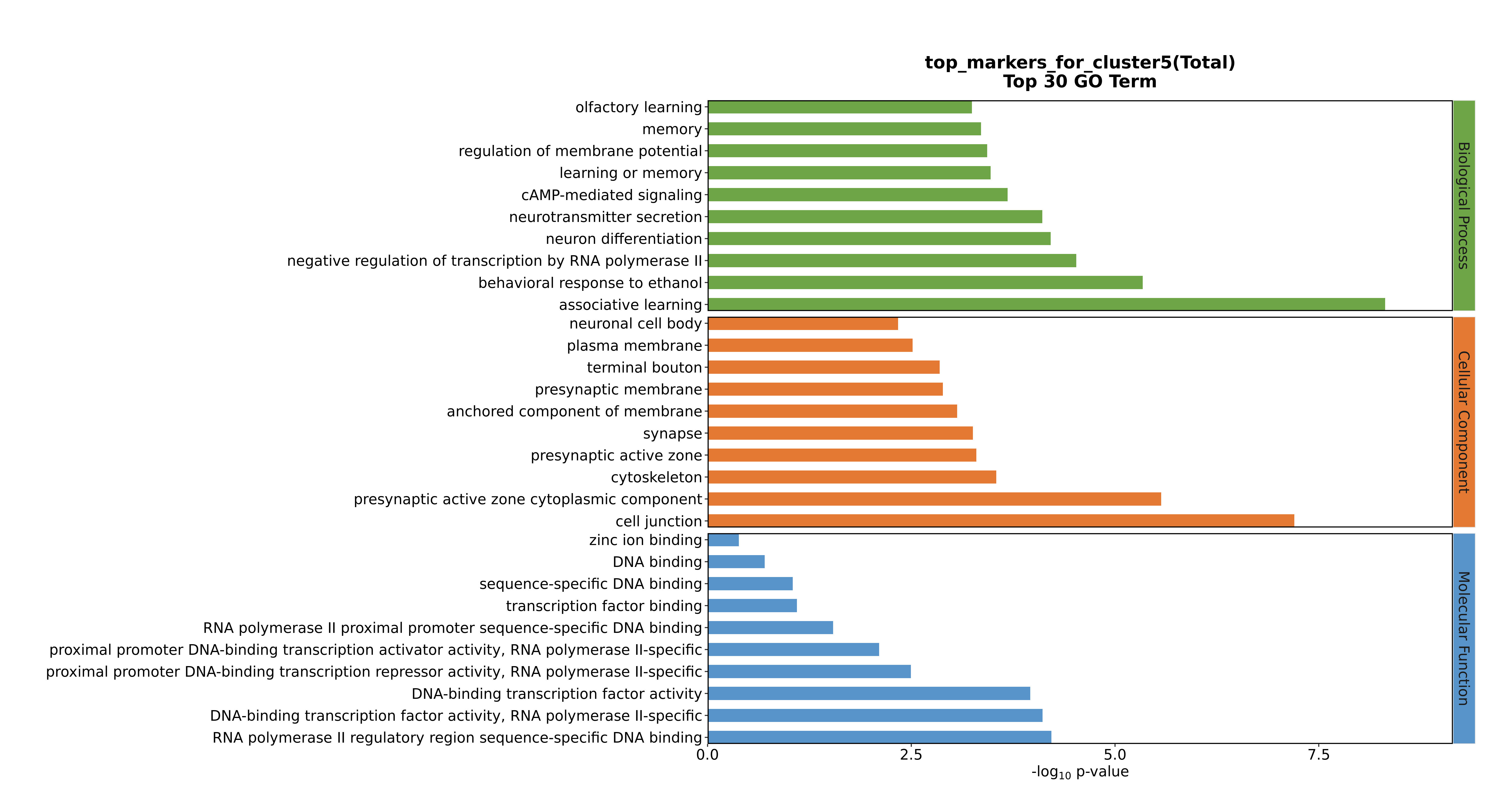

### Cluster5 GO top100.png

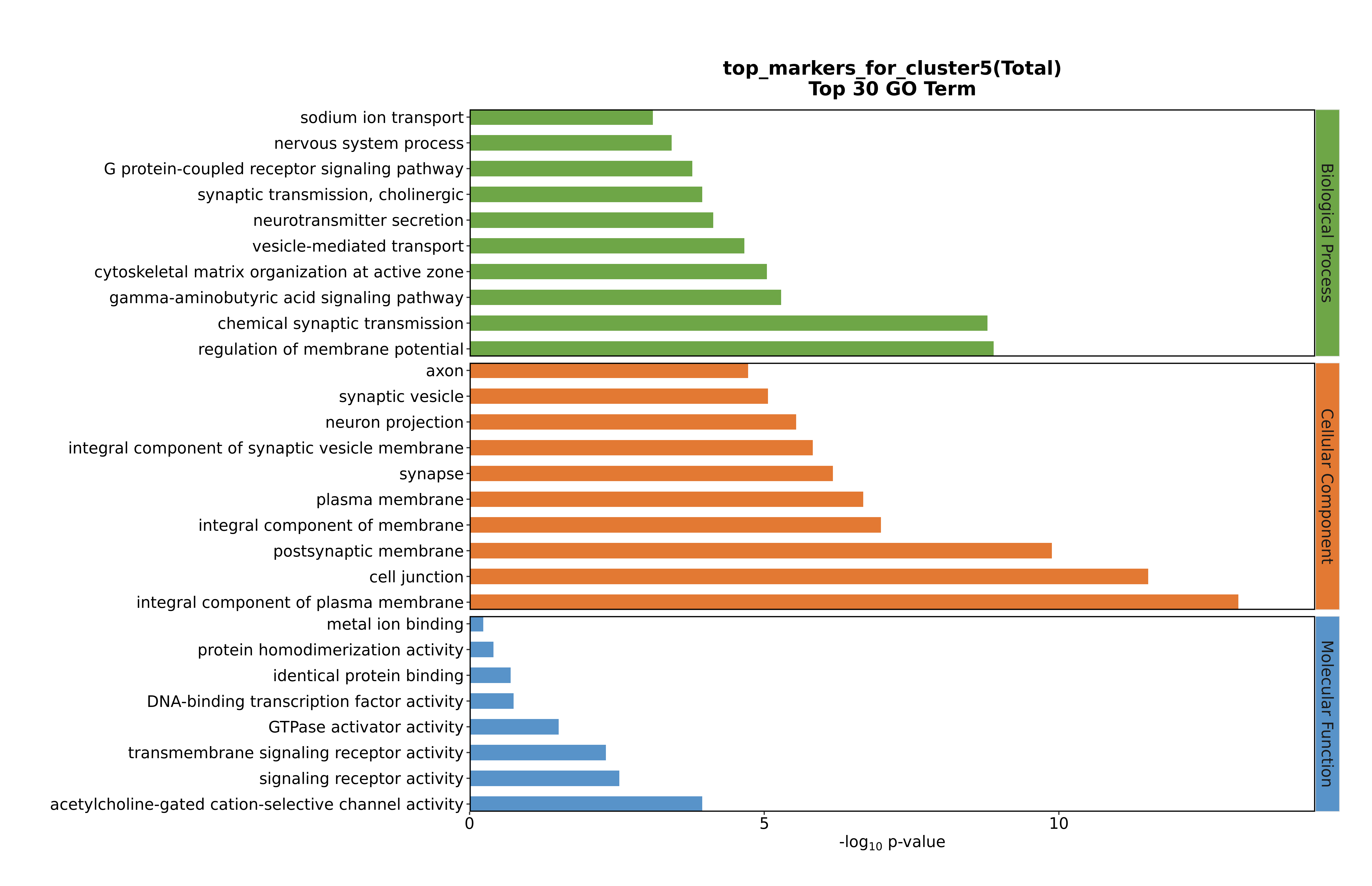

### Cluster5 GO top100.png

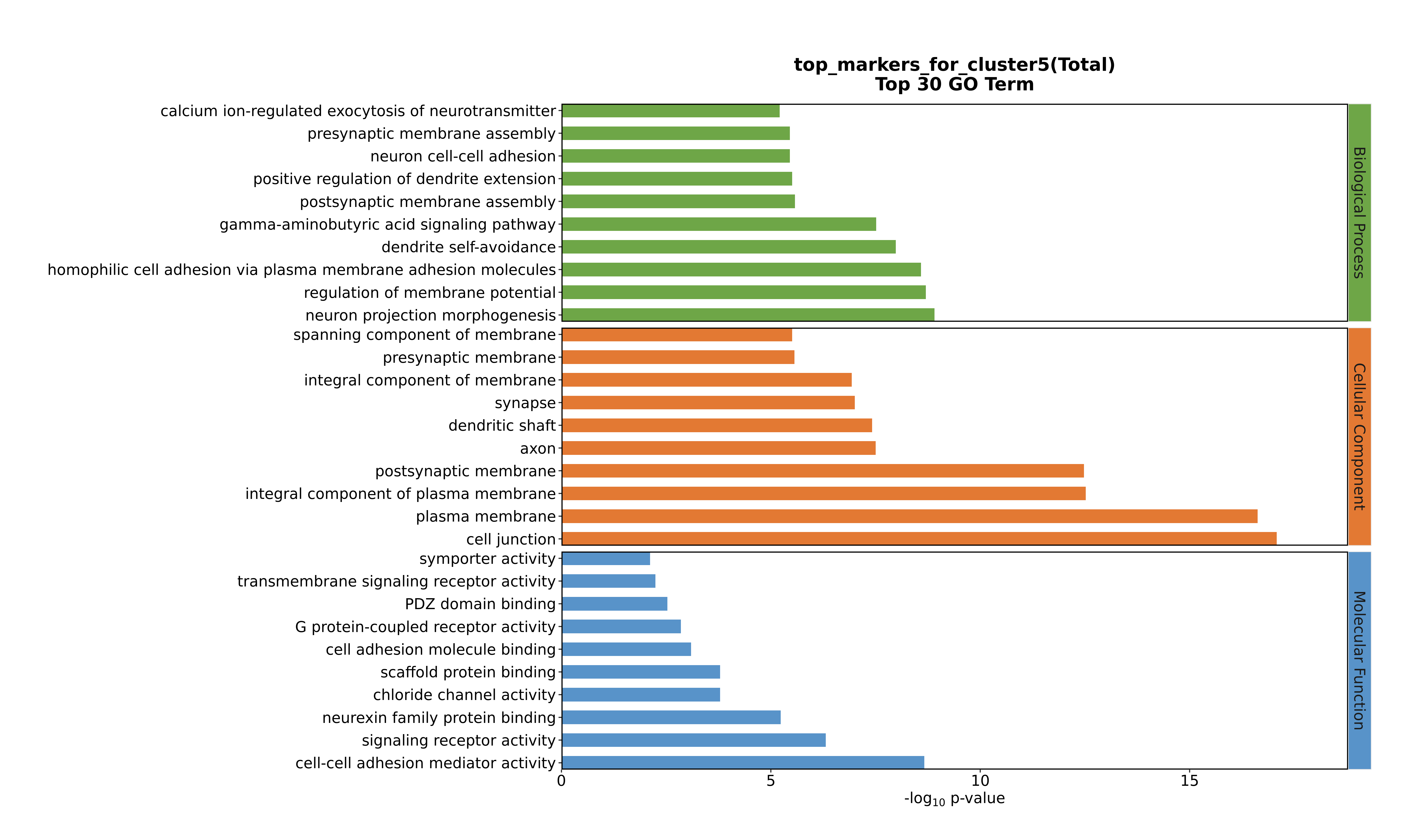

### Cluster6 GO top100.png

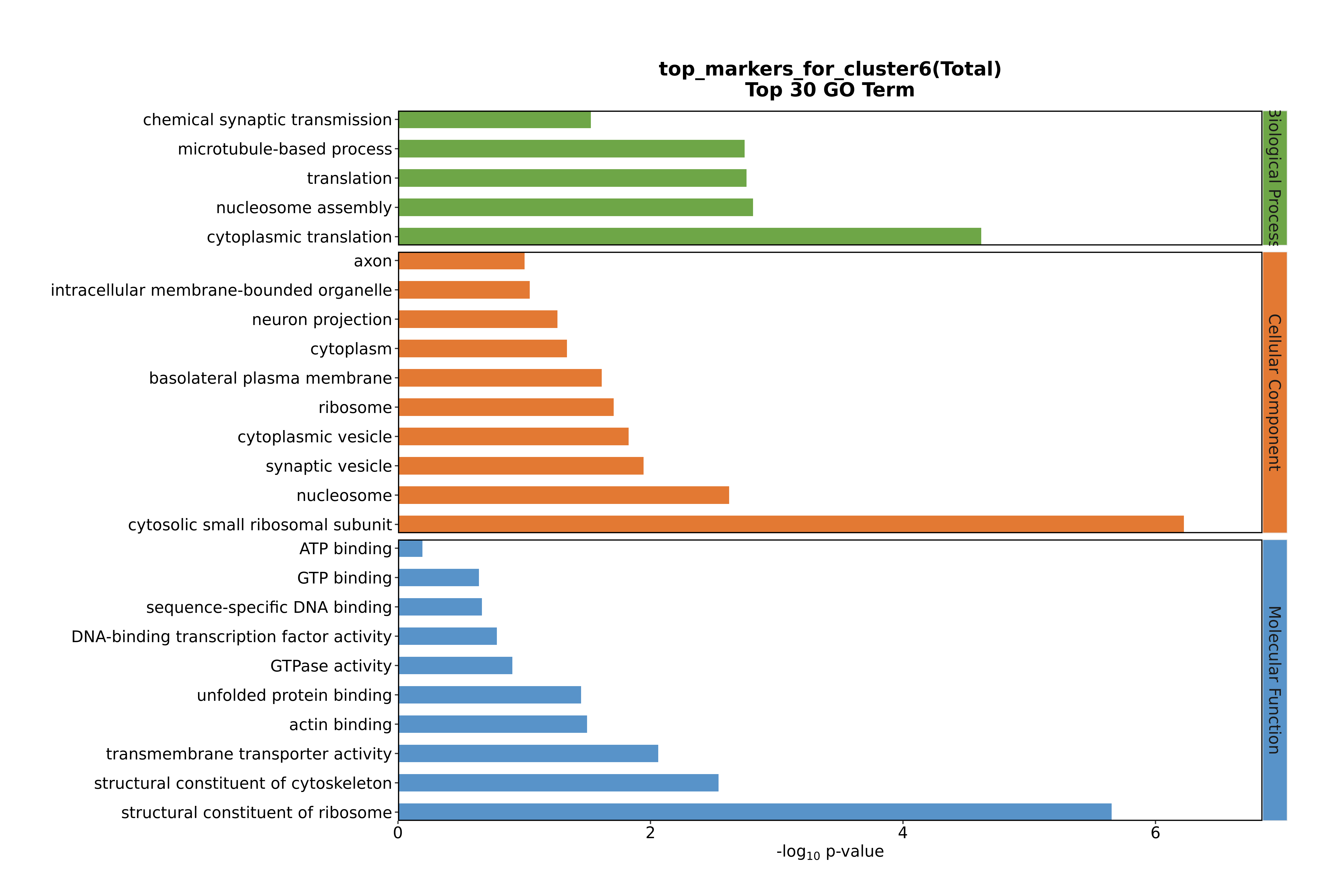

### Cluster6 GO top100.png

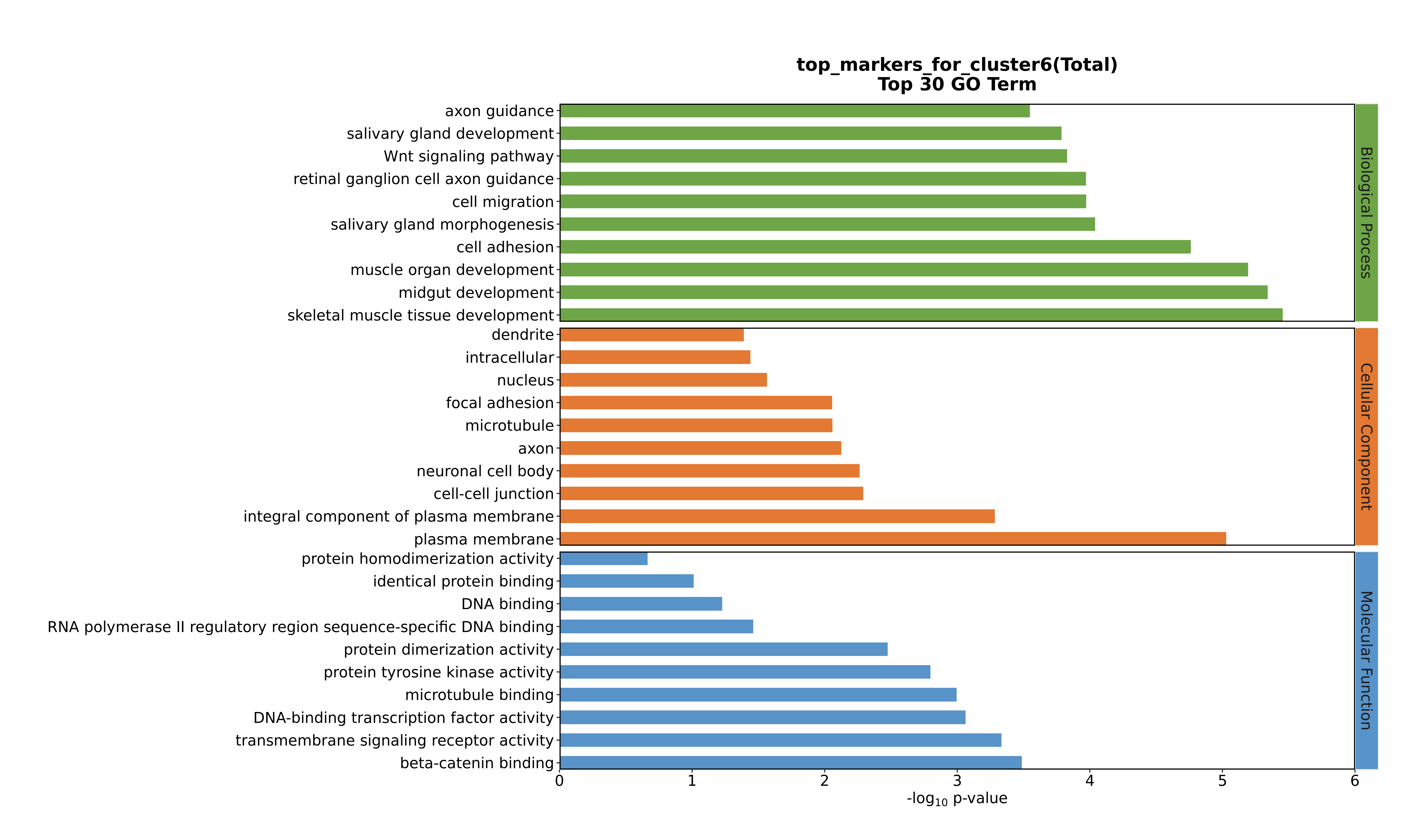

### Cluster6 GO top100.png

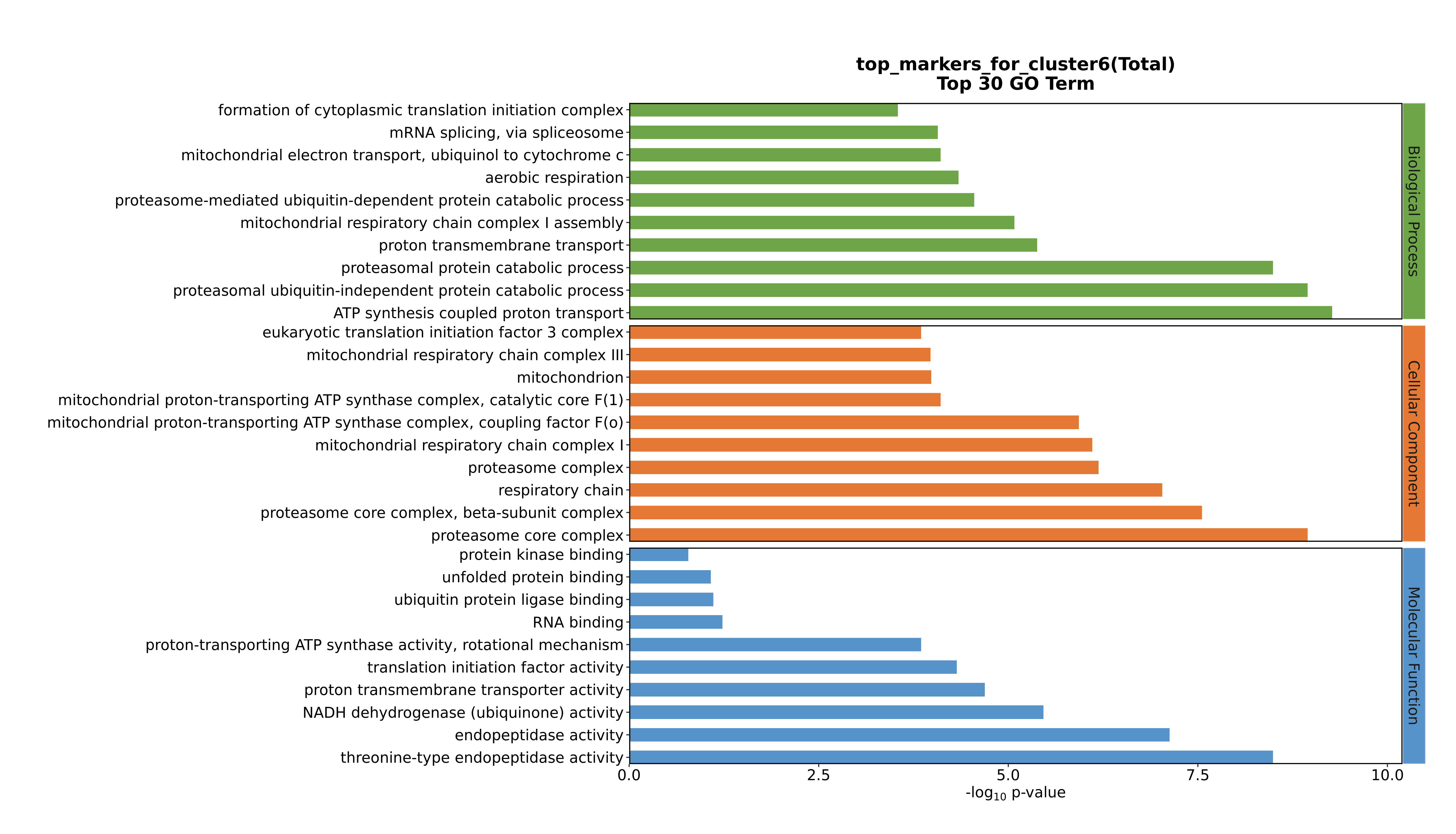

### Cluster6 GO top100.png

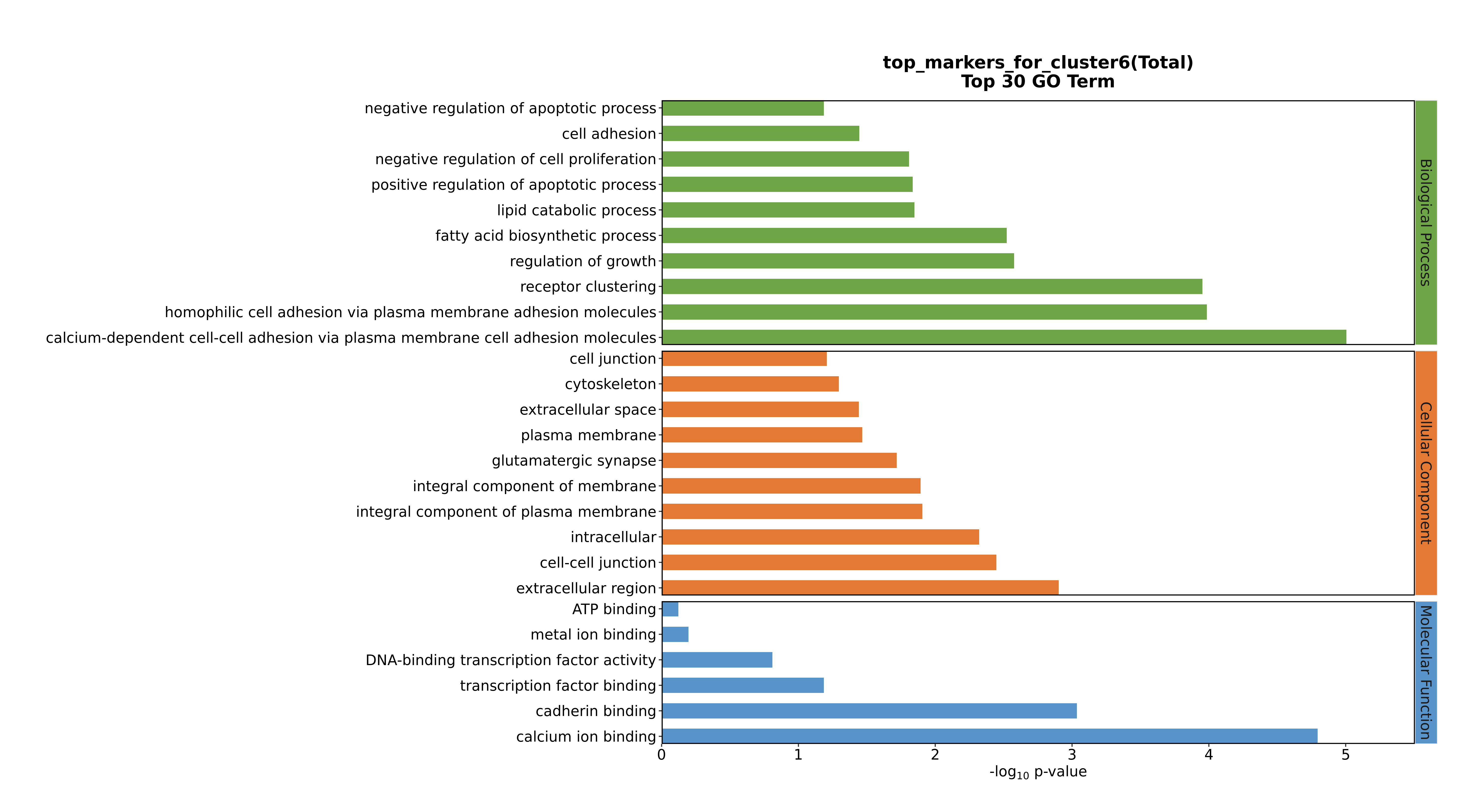

### Cluster7 GO top100.png

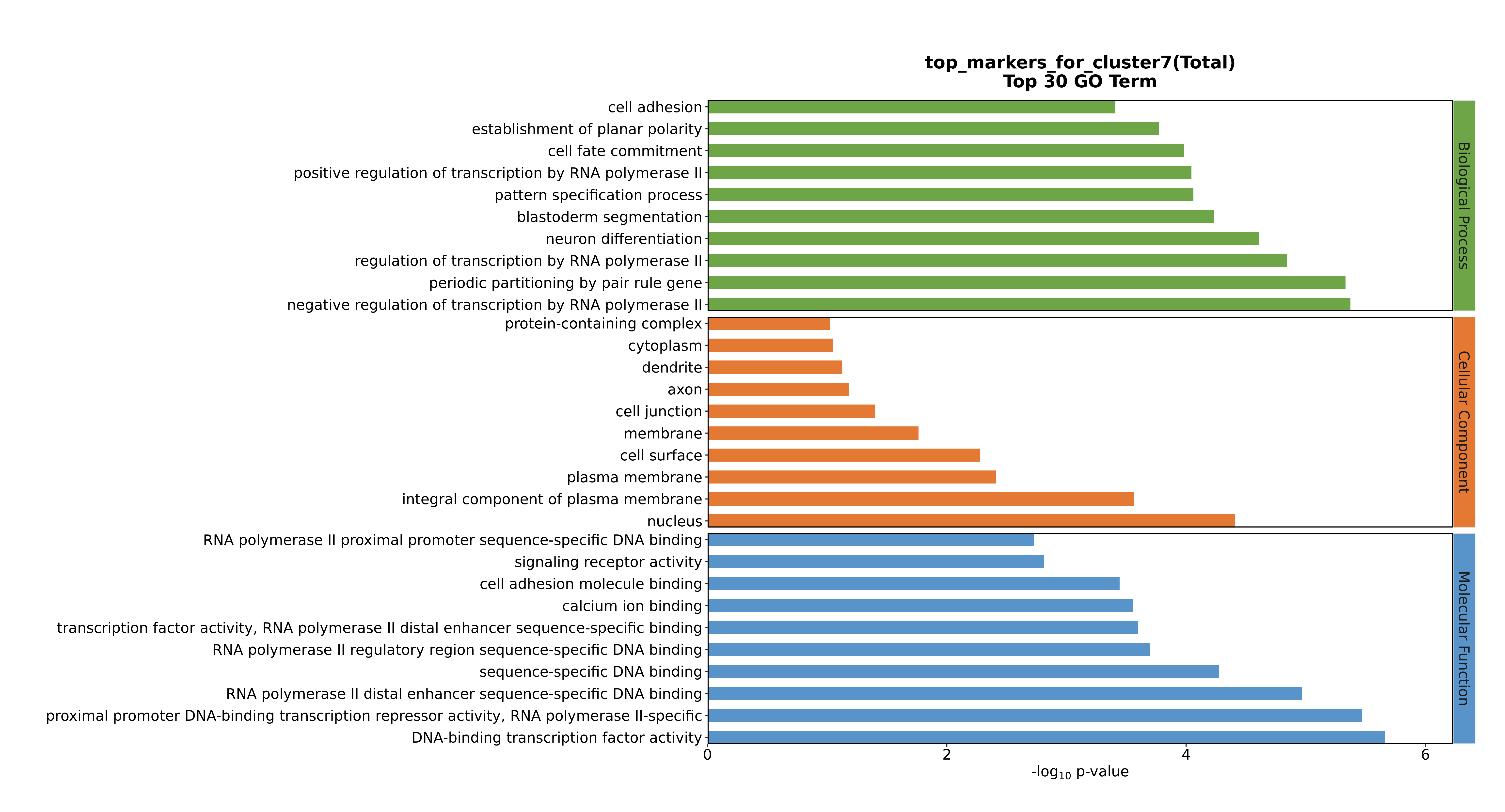

### Cluster7 GO top100.png

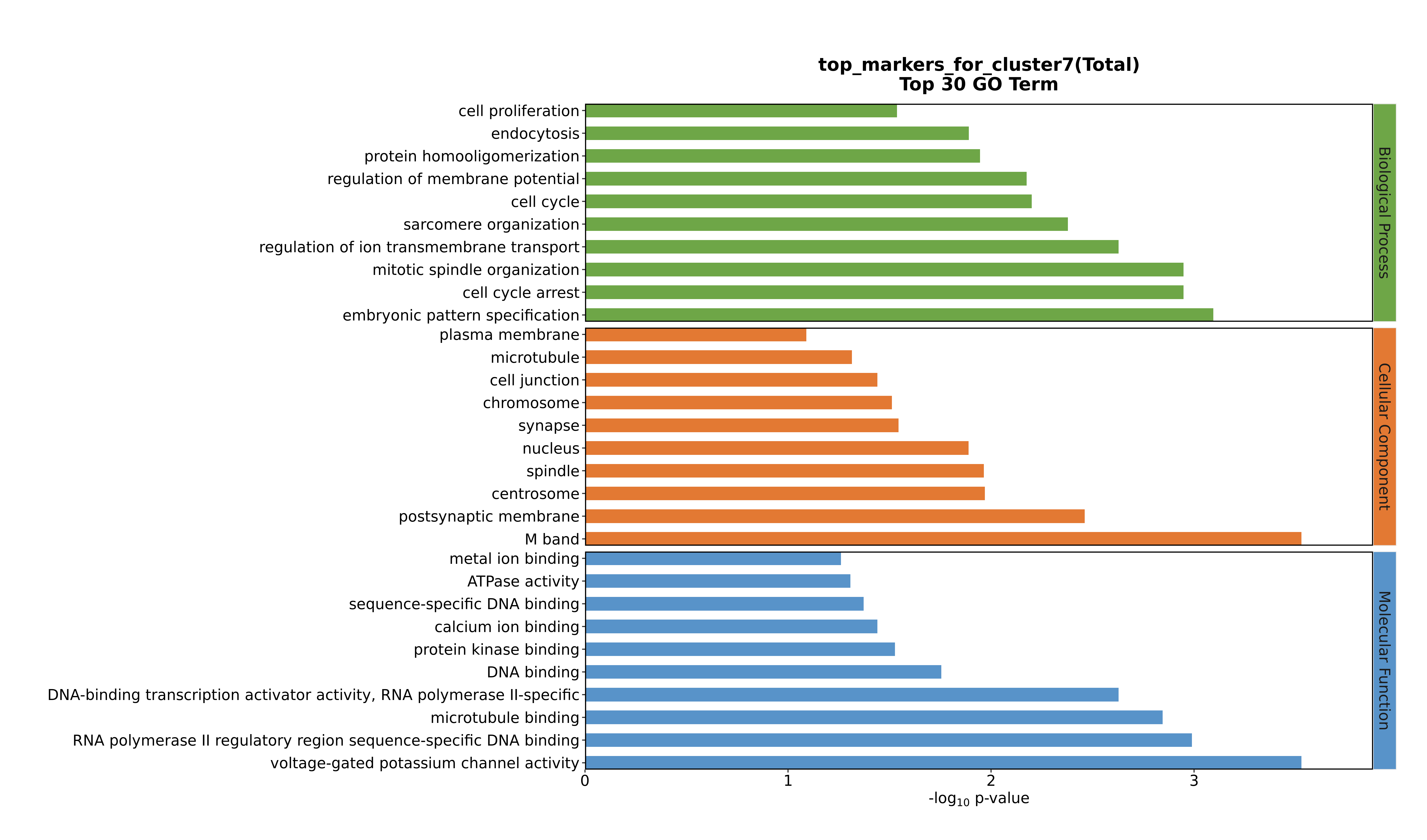

### Cluster7 GO top100.png

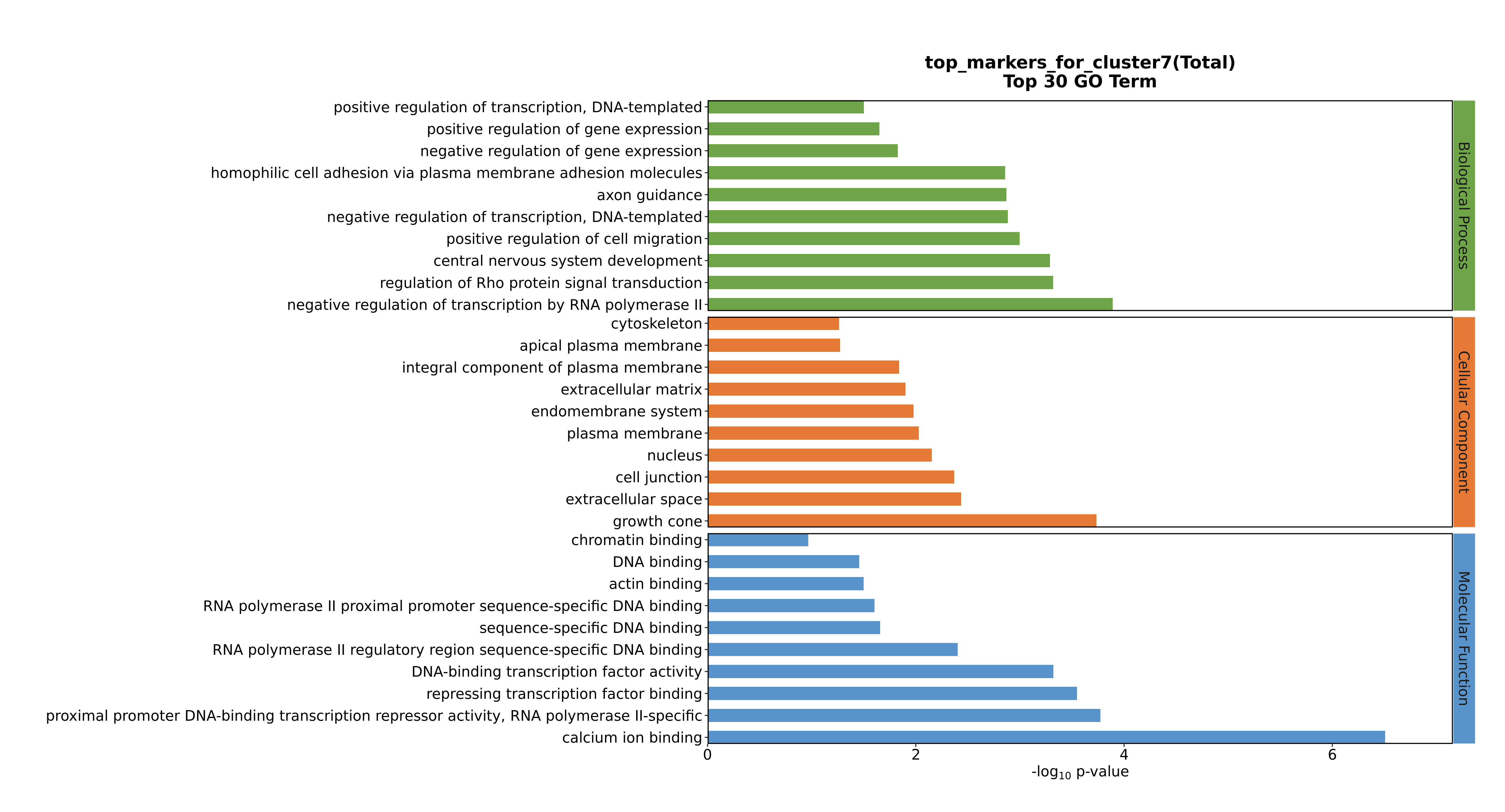

### Cluster7 GO top100.png

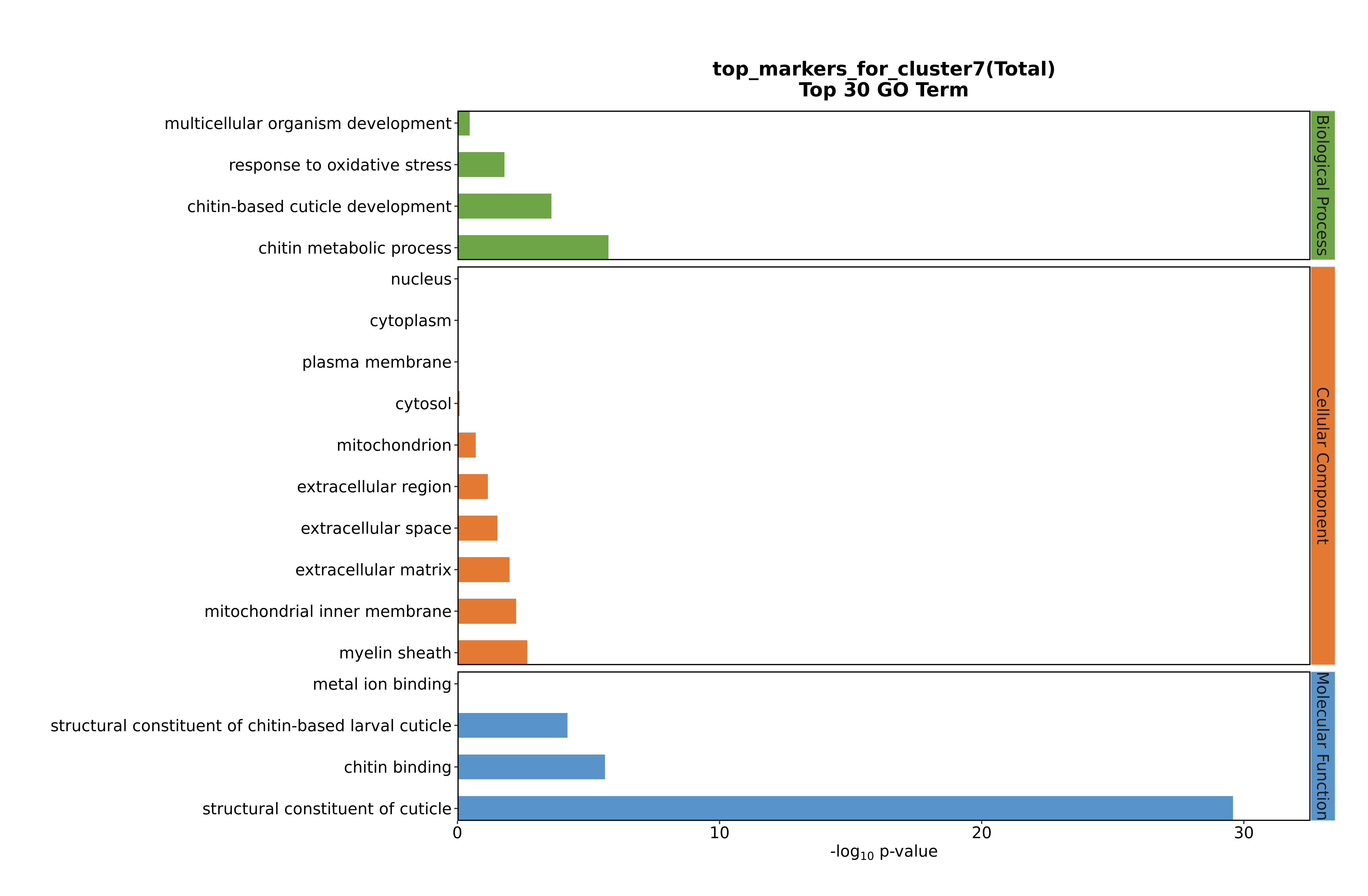

### Cluster8 GO top100.png

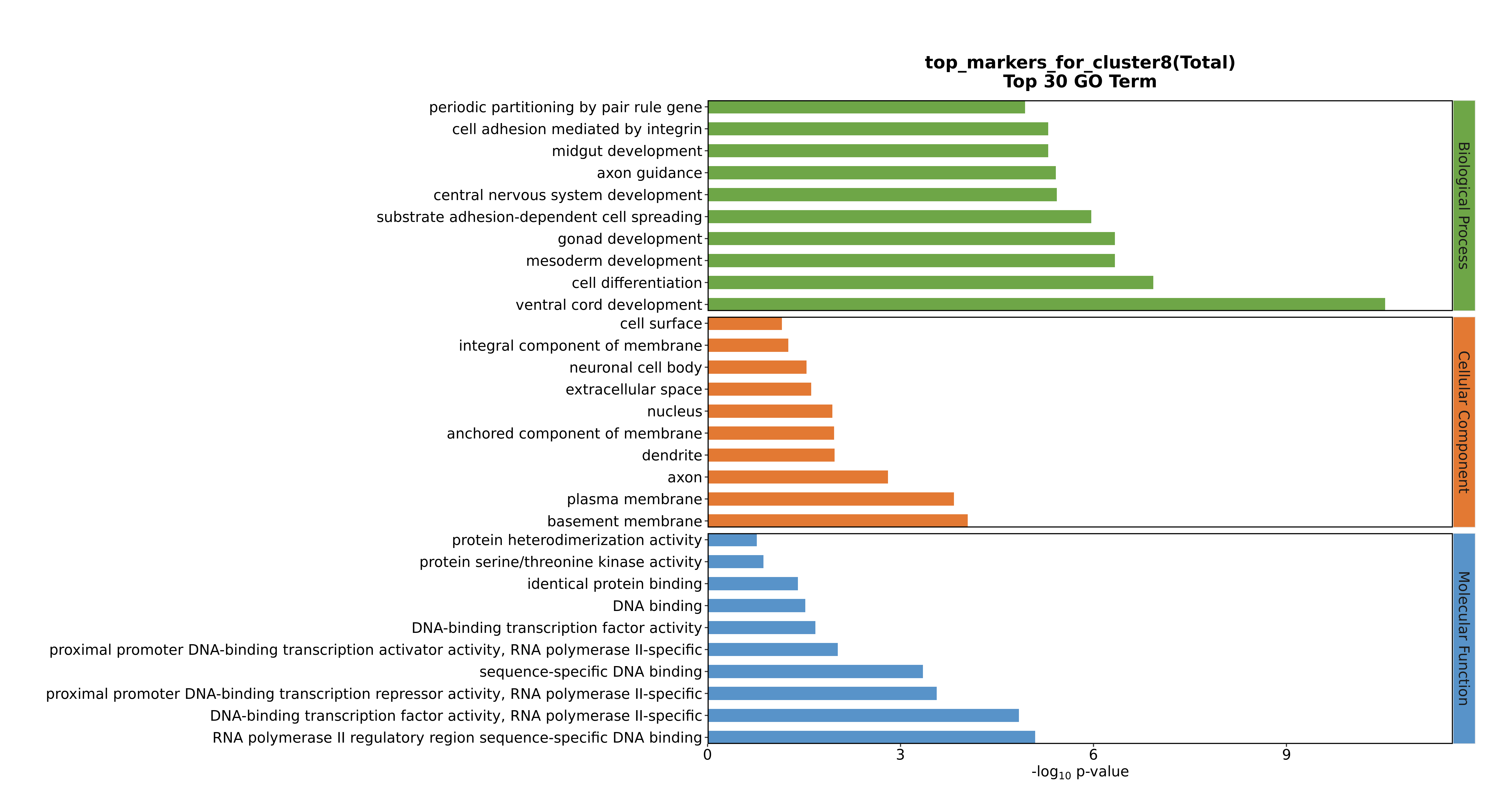

### Cluster8 GO top100.png

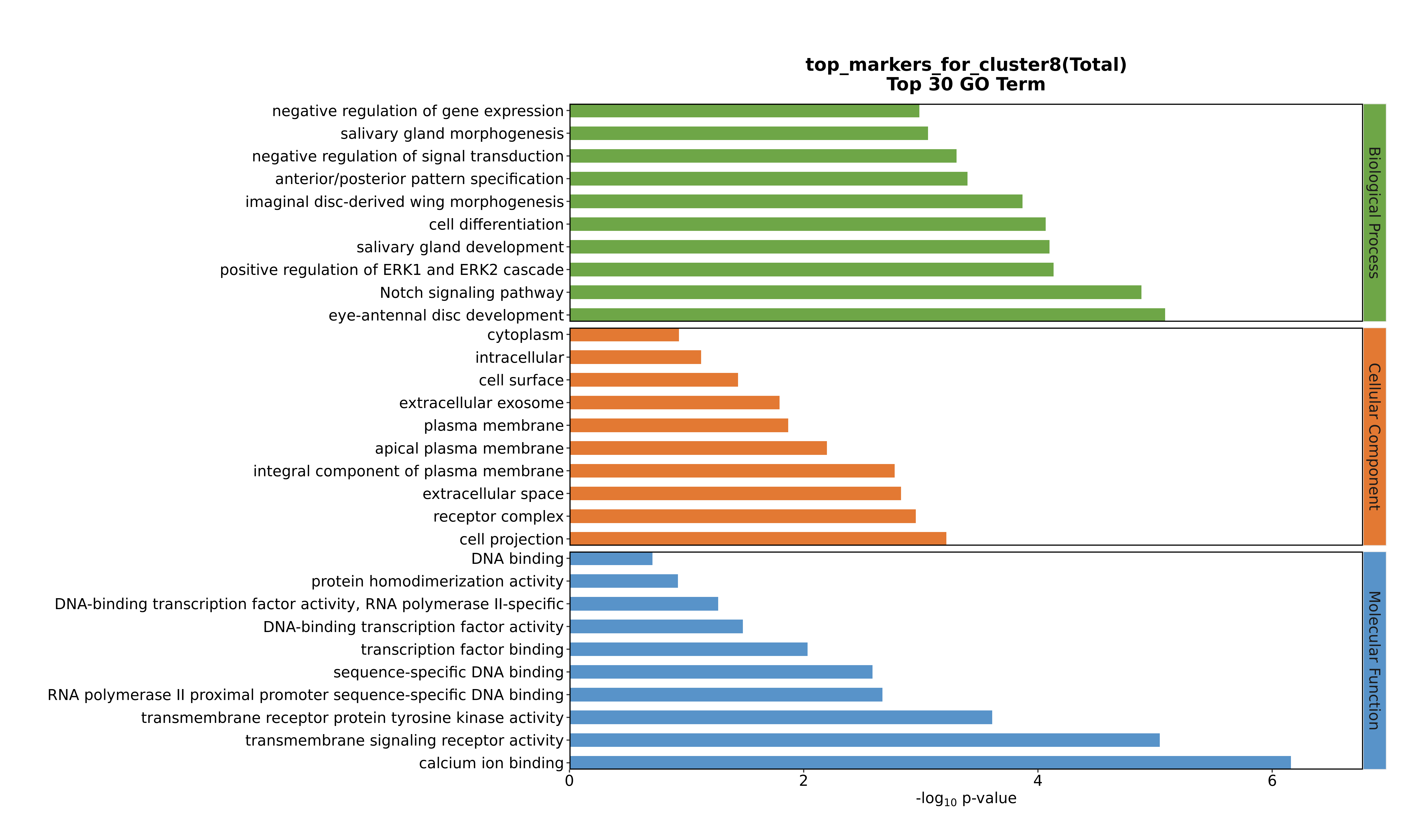

### Cluster8 GO top100.png

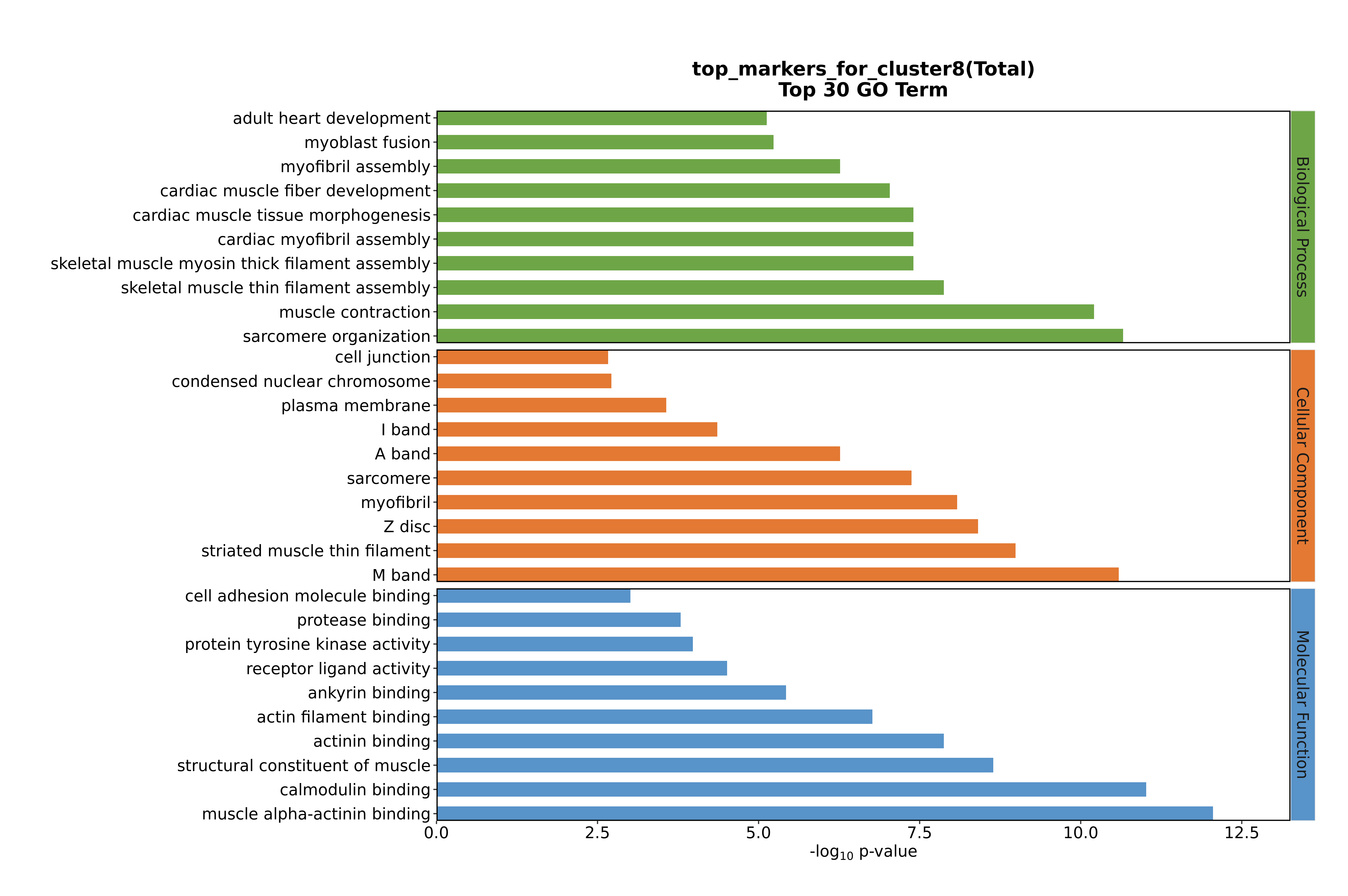

### Cluster8 GO top100.png

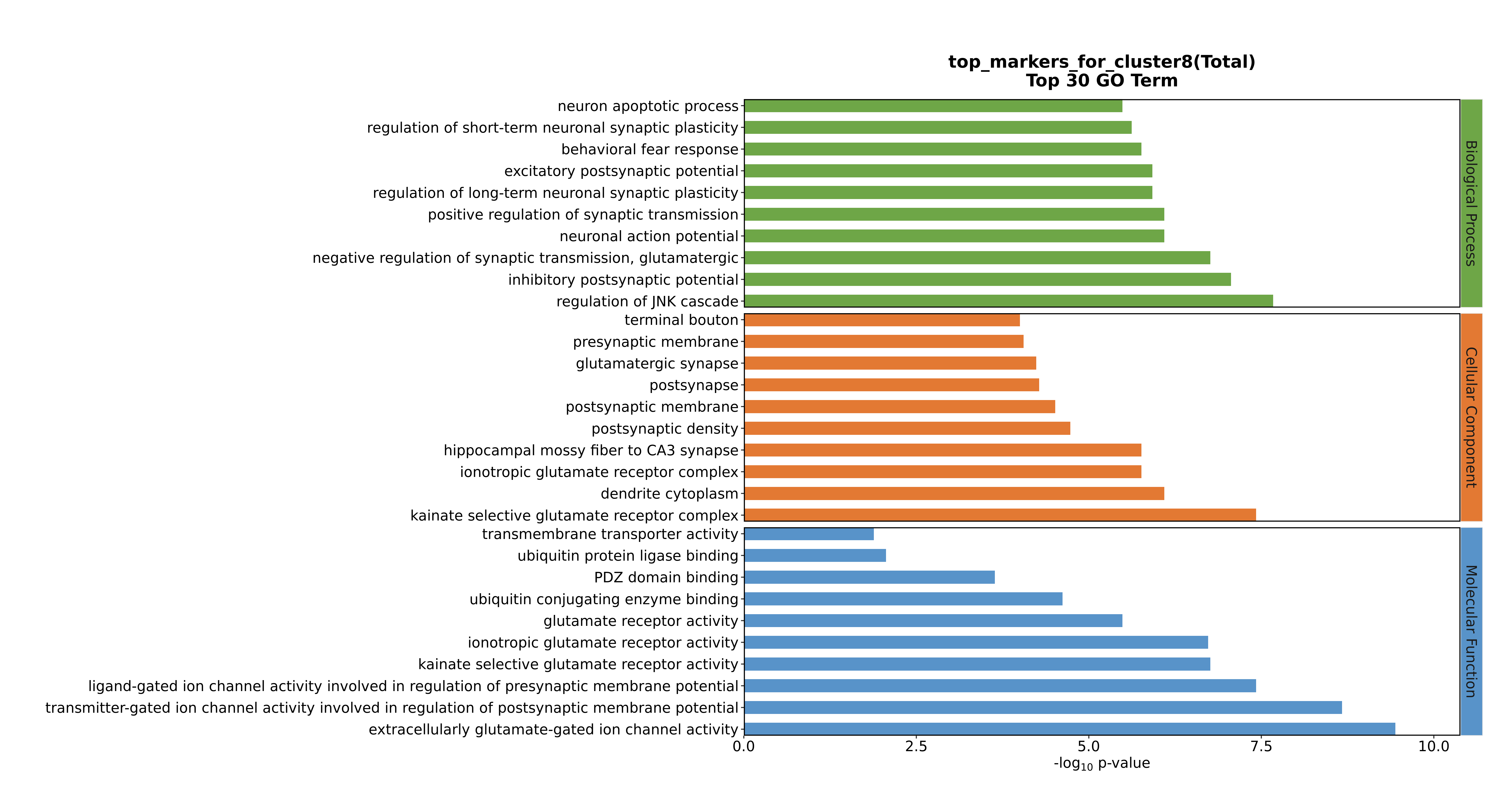

### Cluster9 GO top100.png

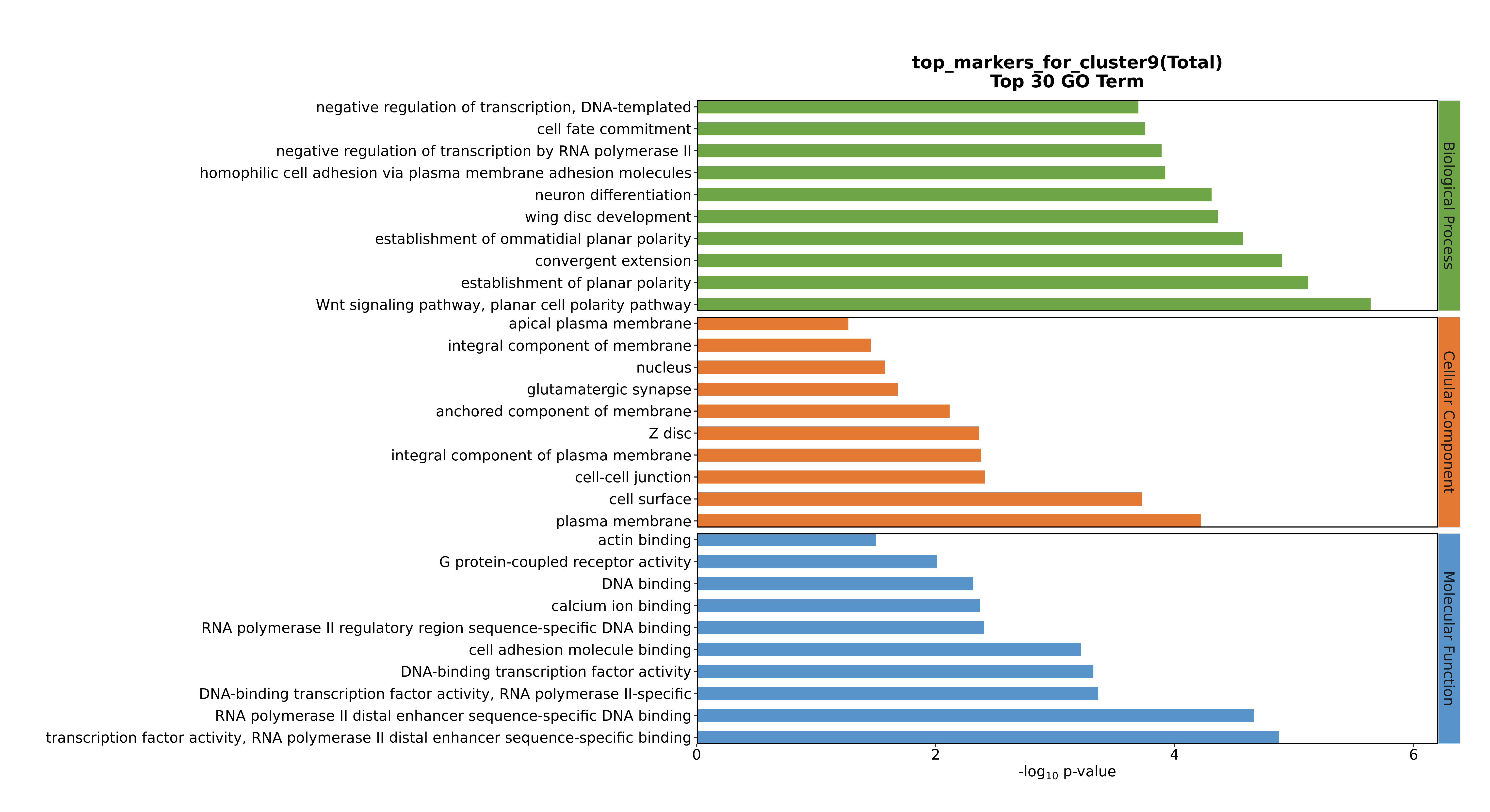

### Cluster9 GO top100.png

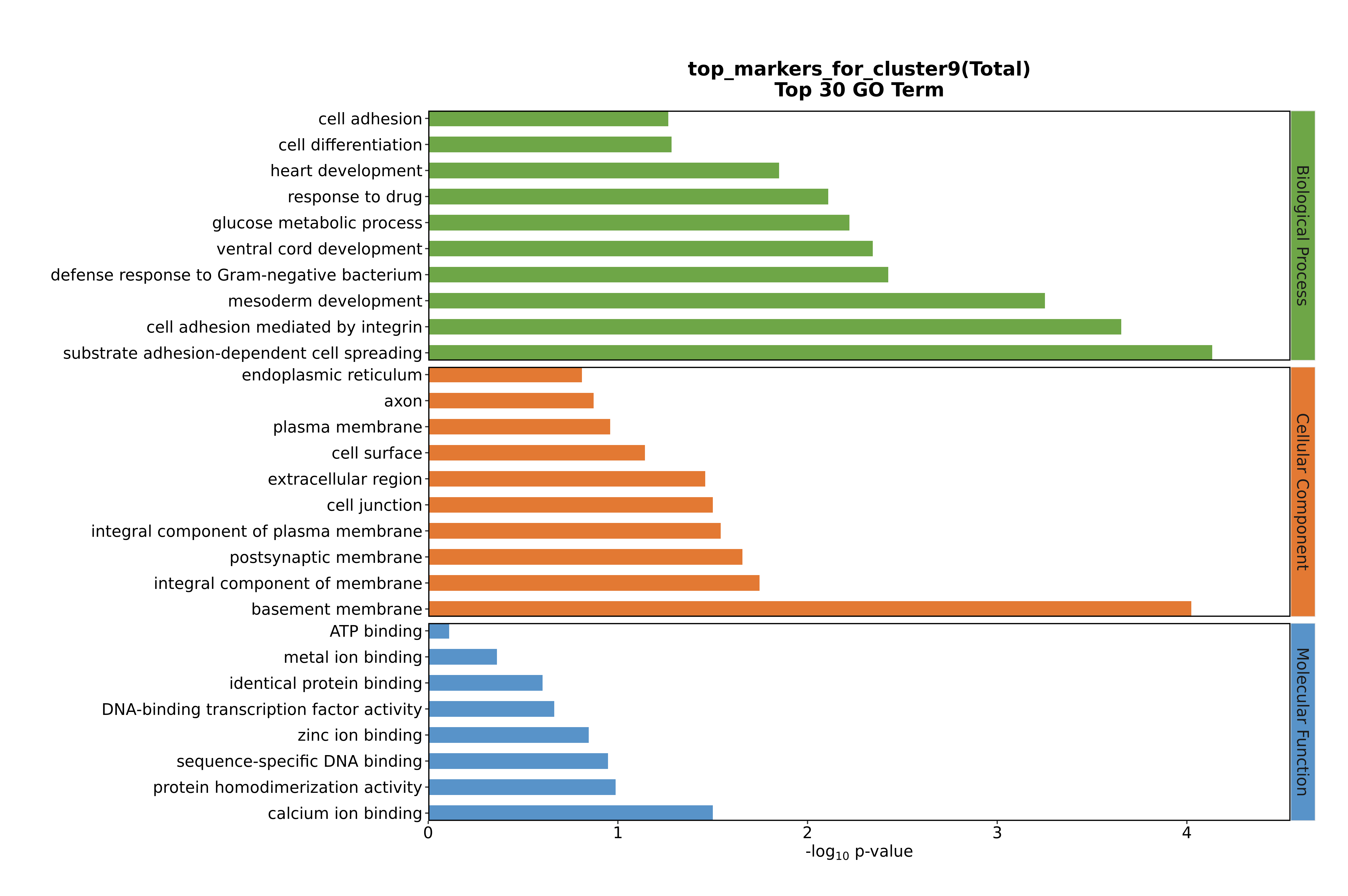

### Cluster9 GO top100.png

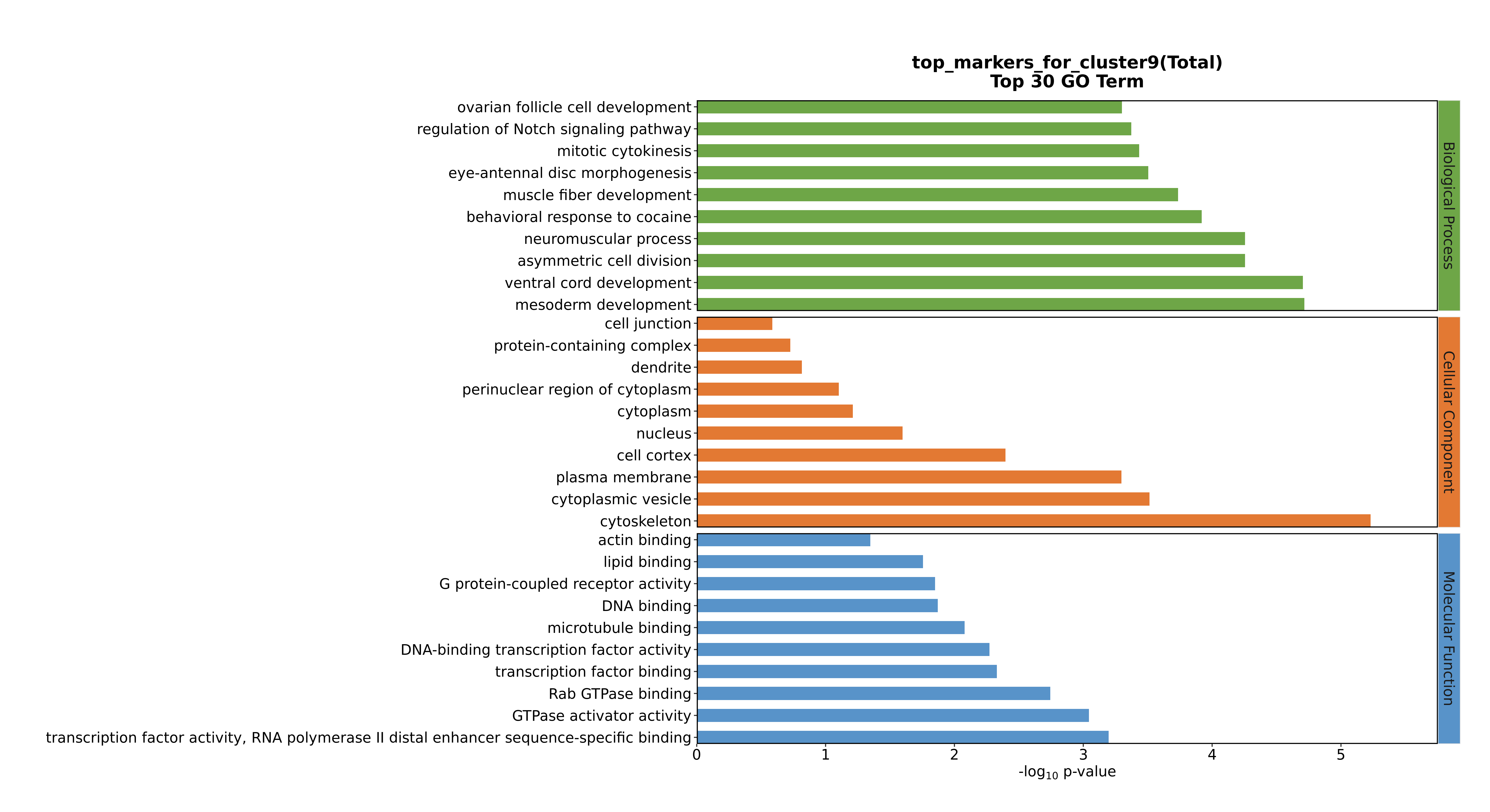

### Cluster9 GO top100.png

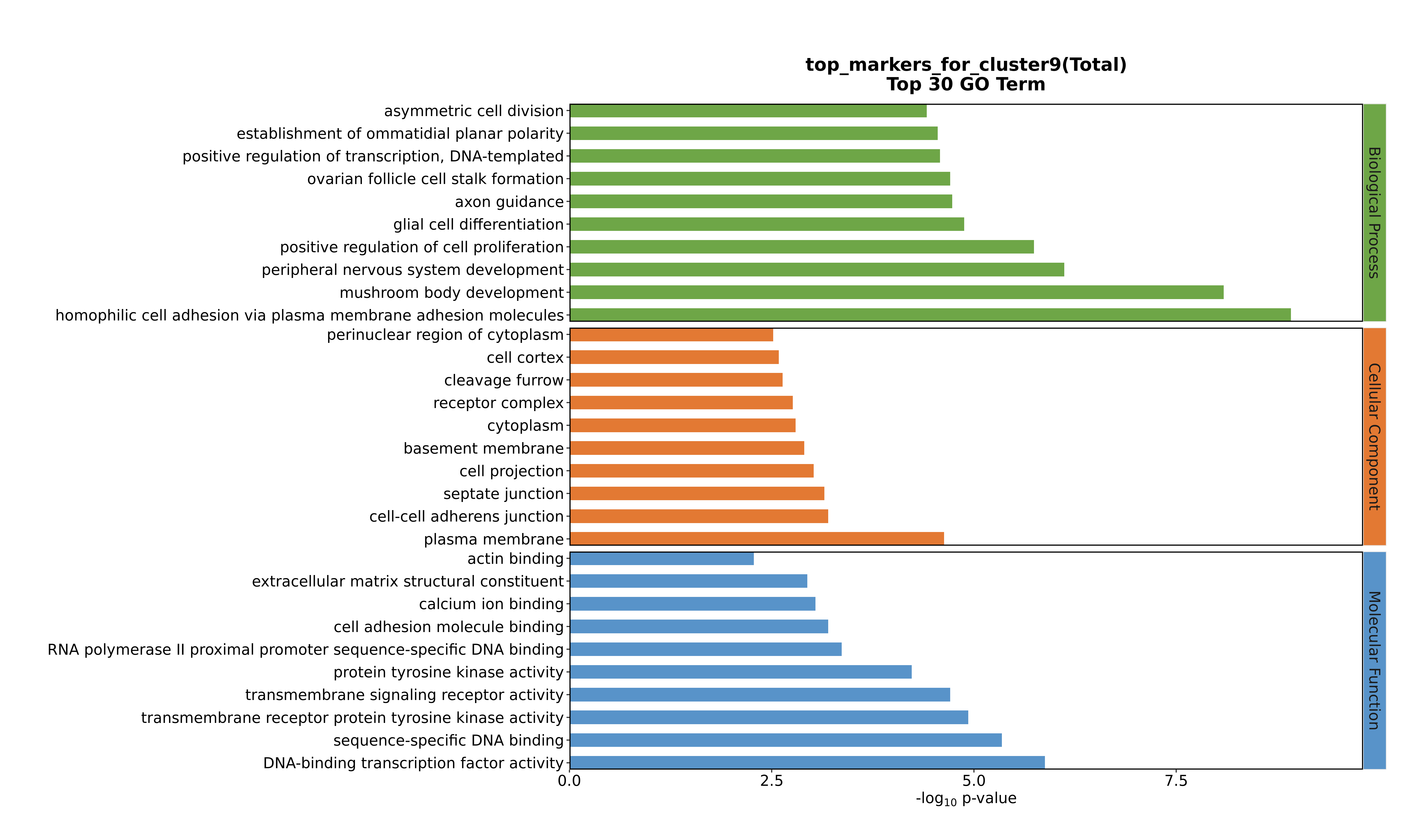

### Cluster10 GO top100.png

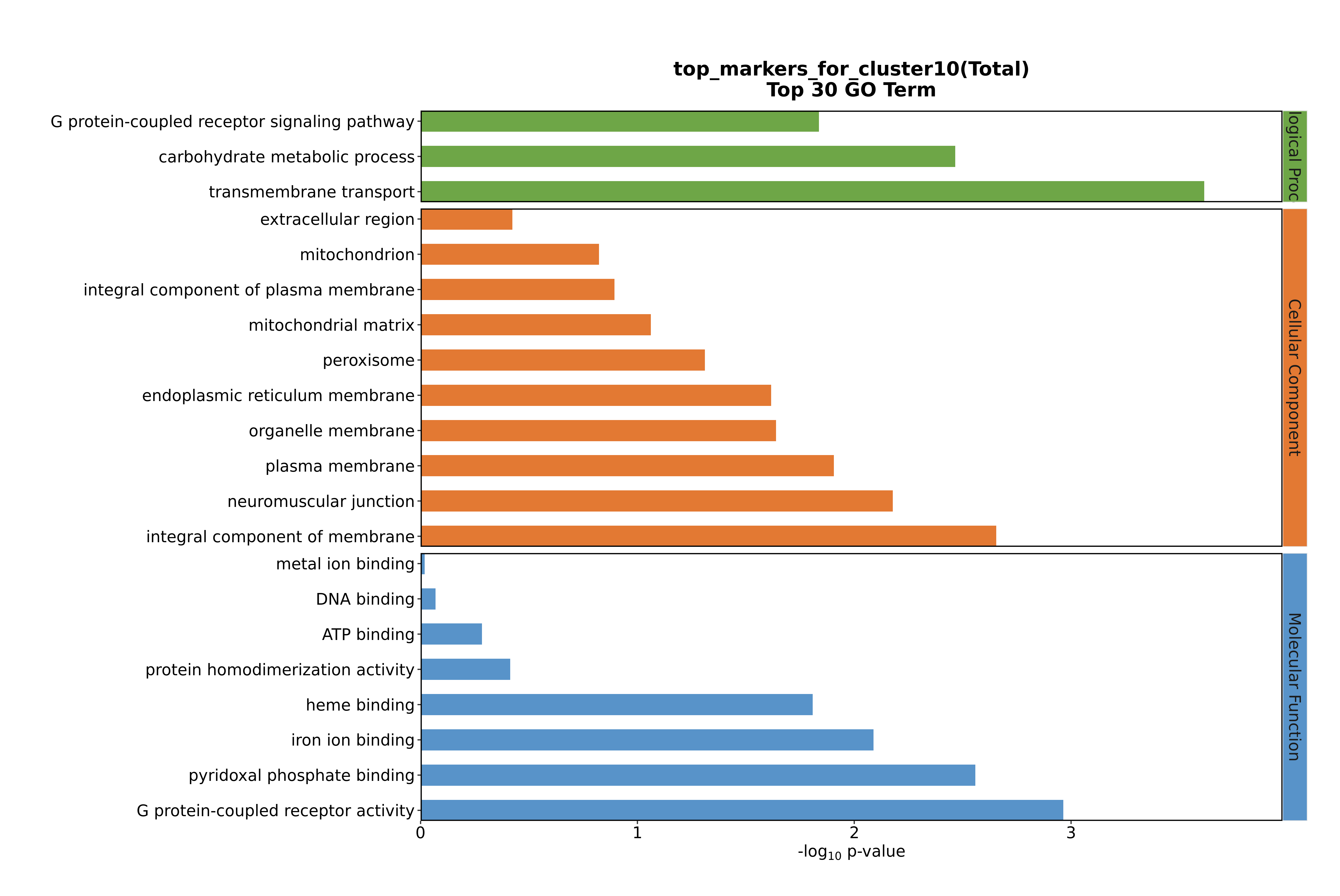

### Cluster10 GO top100.png

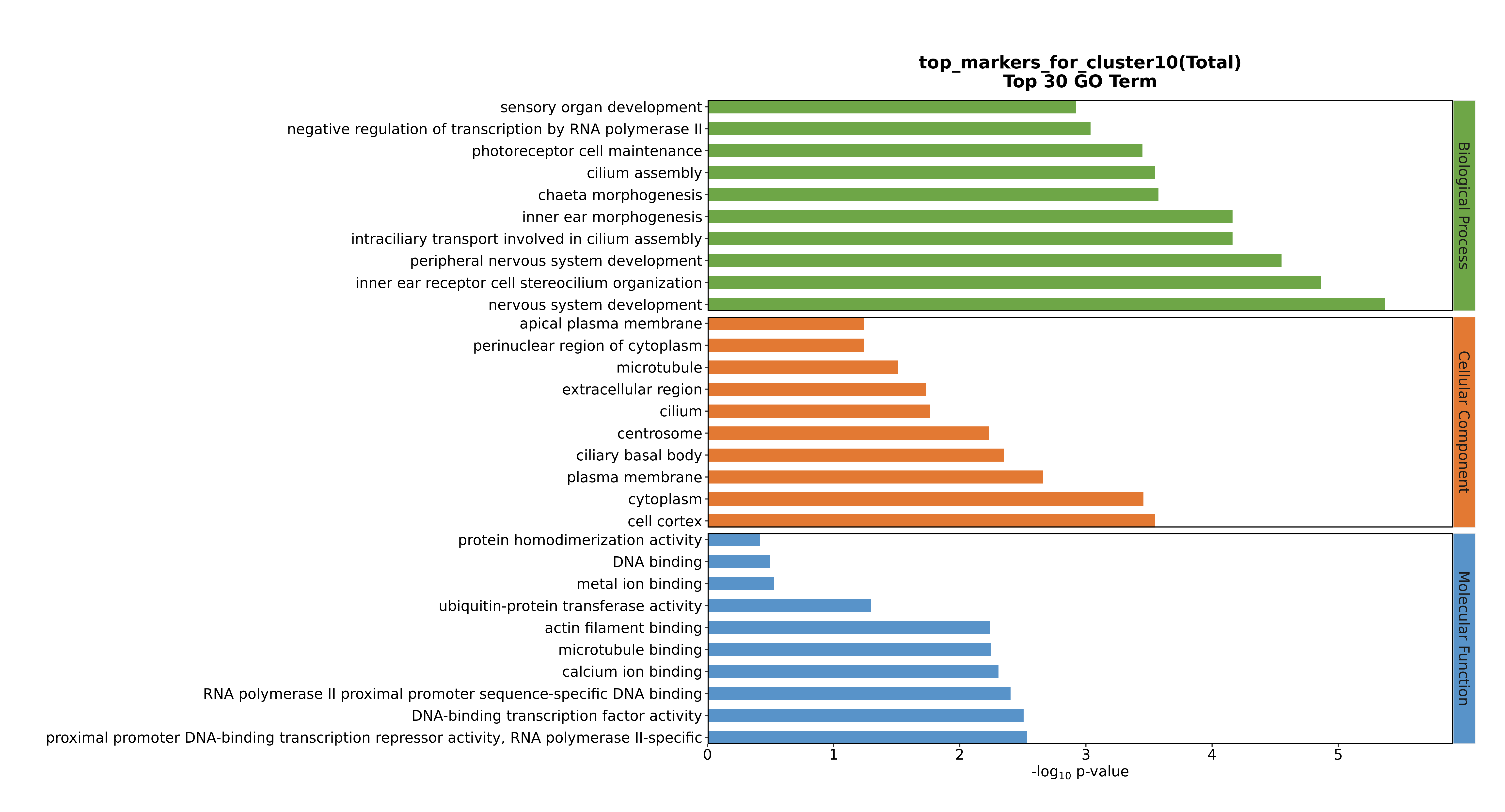

### Cluster10 GO top100.png

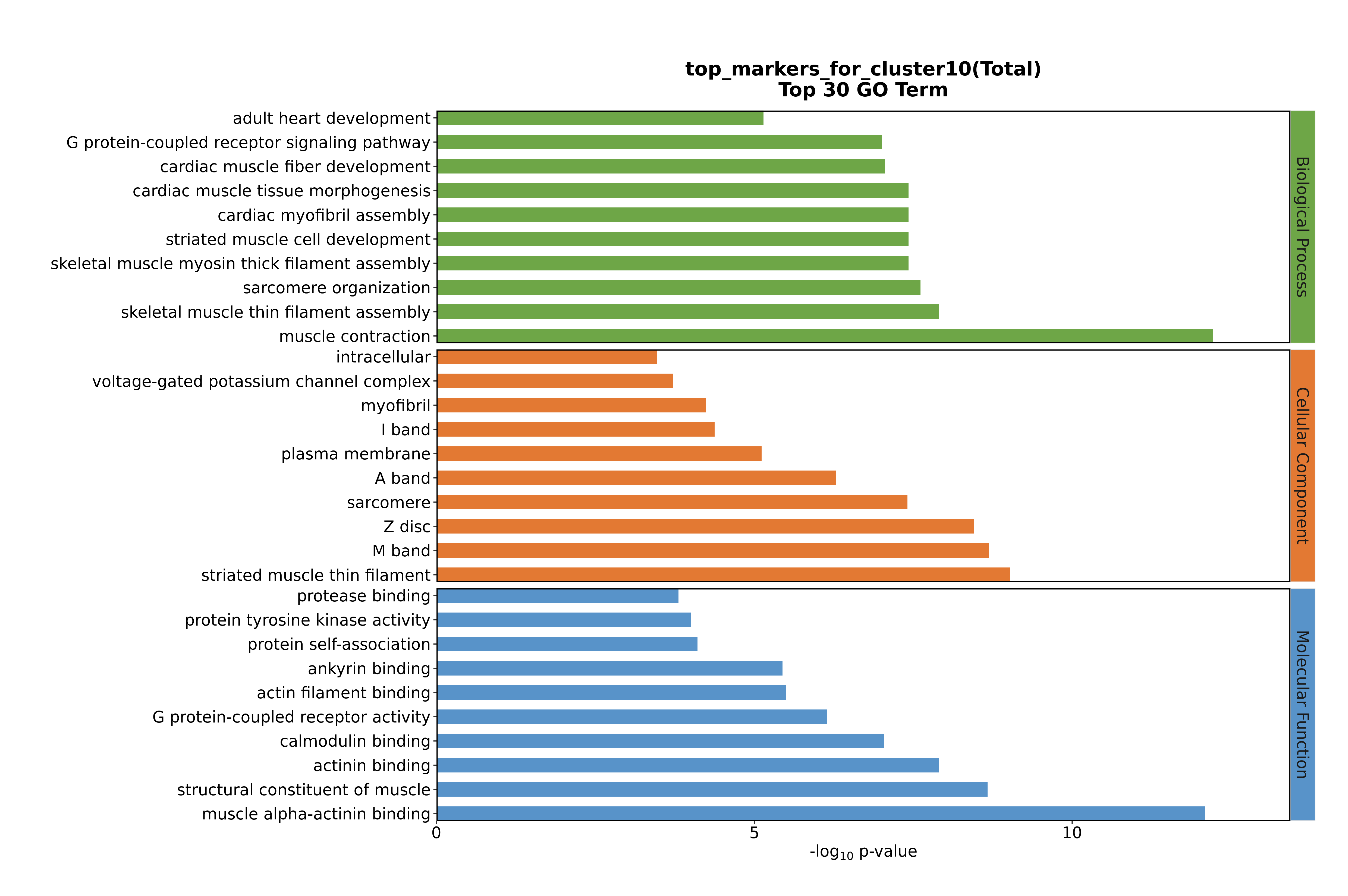

### Cluster10 GO top100.png

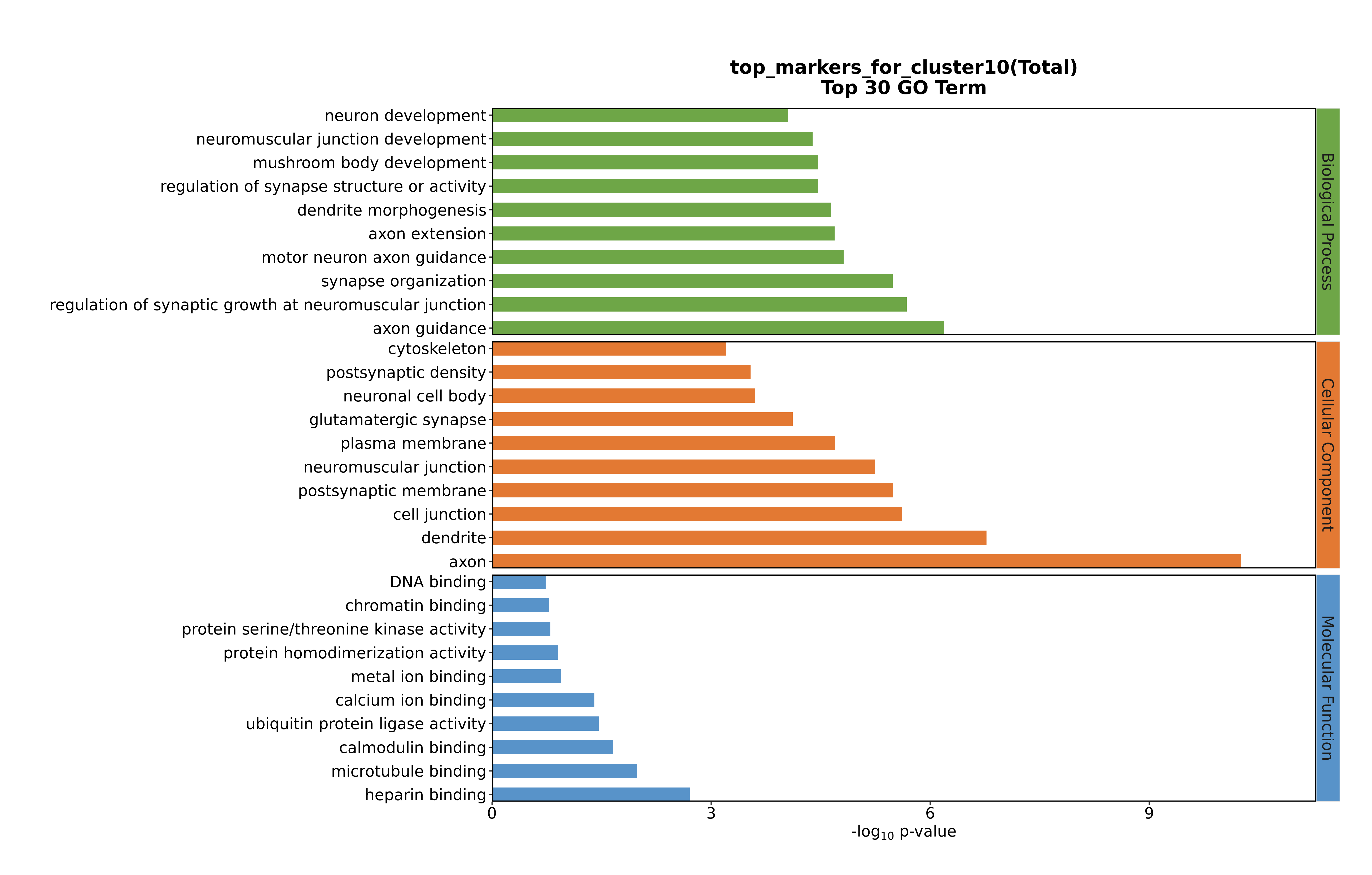

### Cluster10 GO top100.png

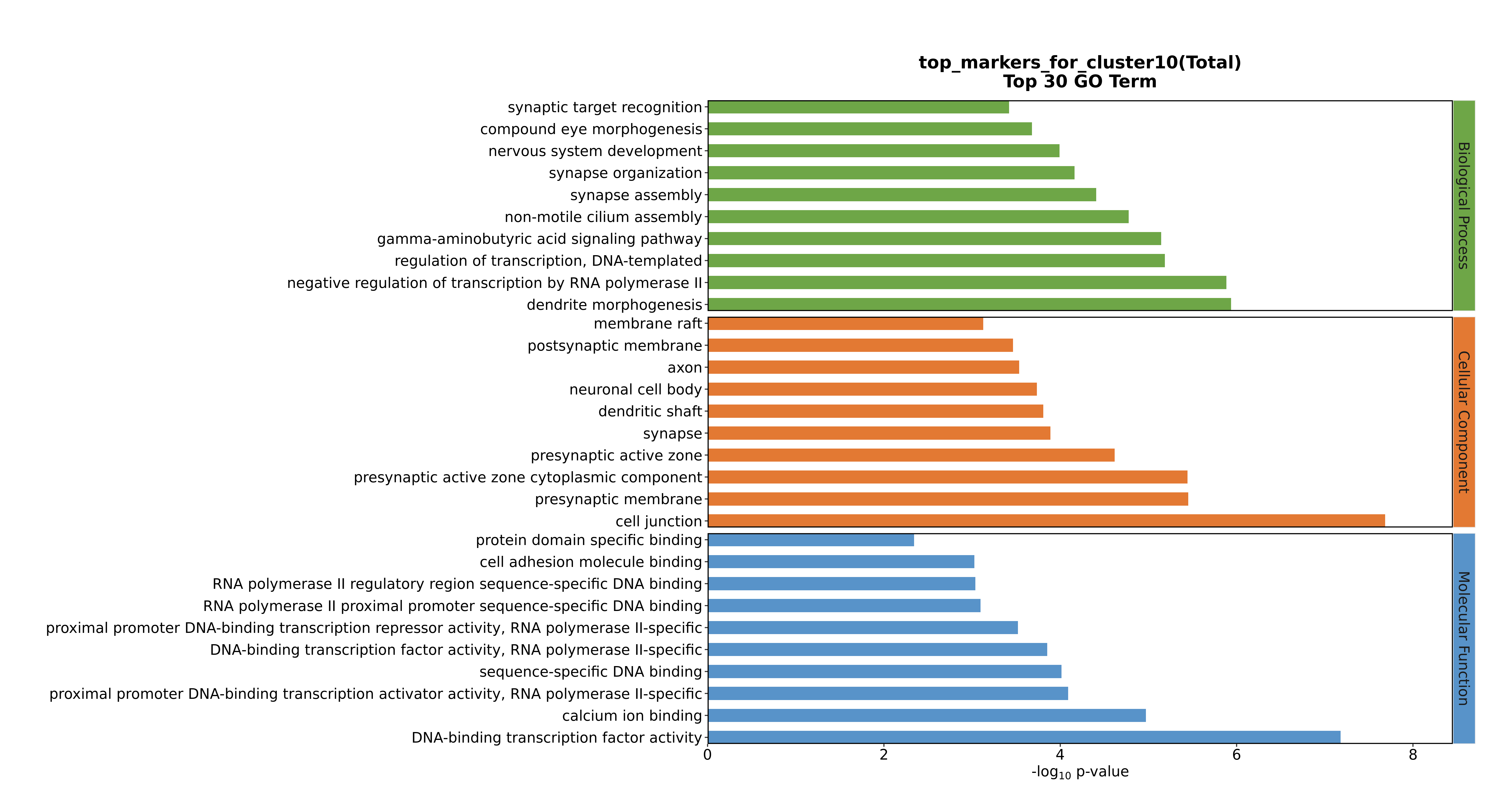

### Cluster11 GO top100.png

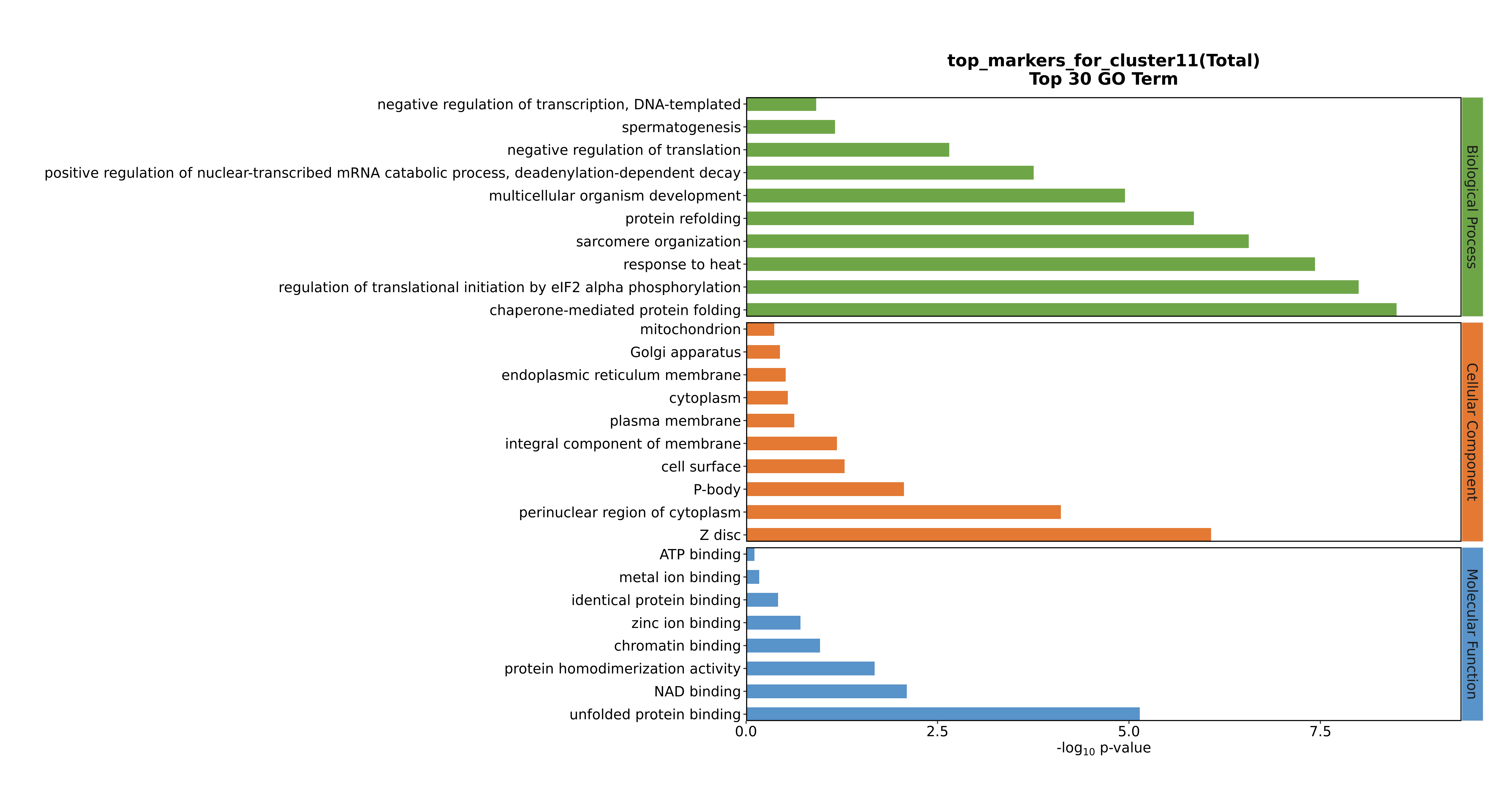

### Cluster11 GO top100.png

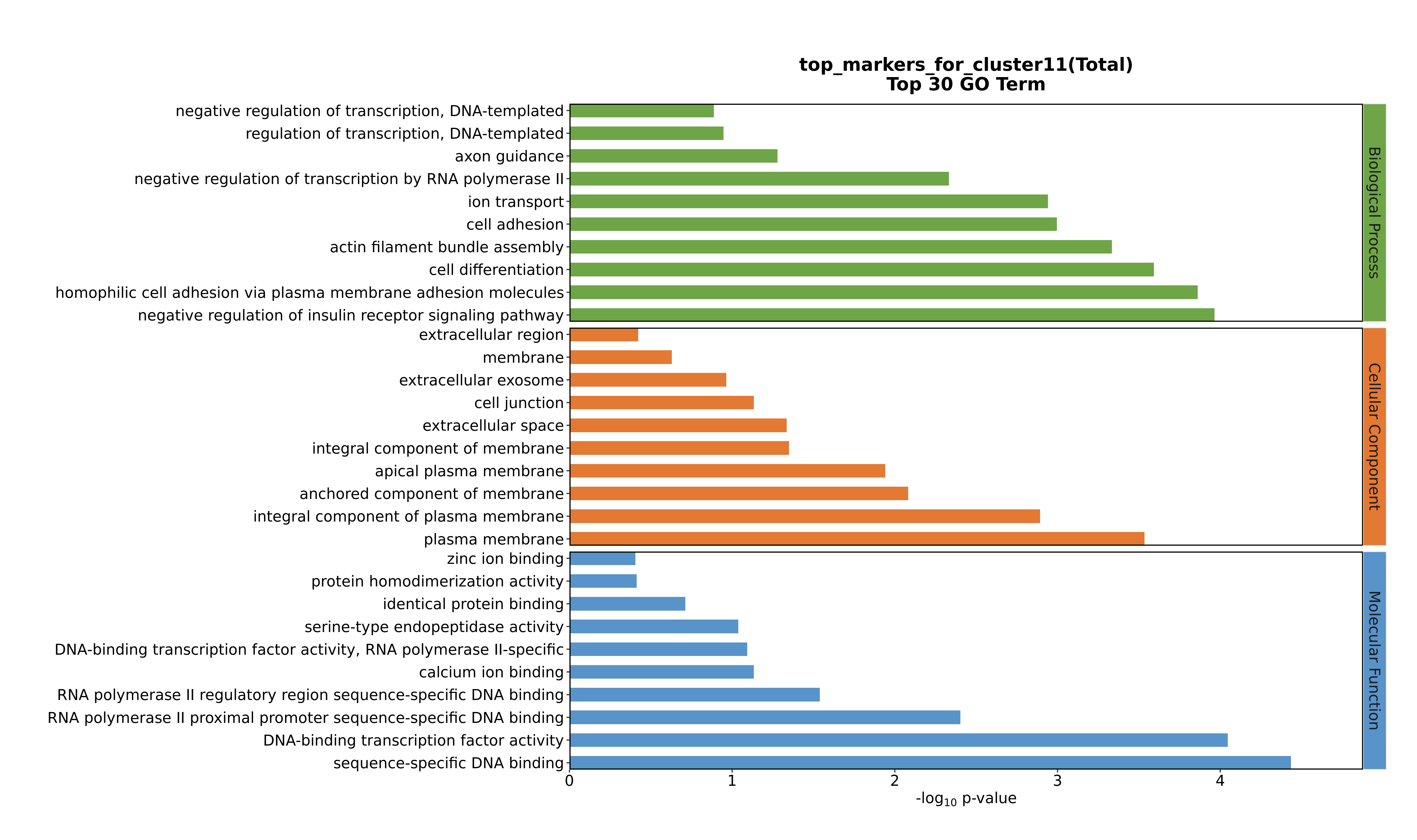

### Cluster11 GO top100.png

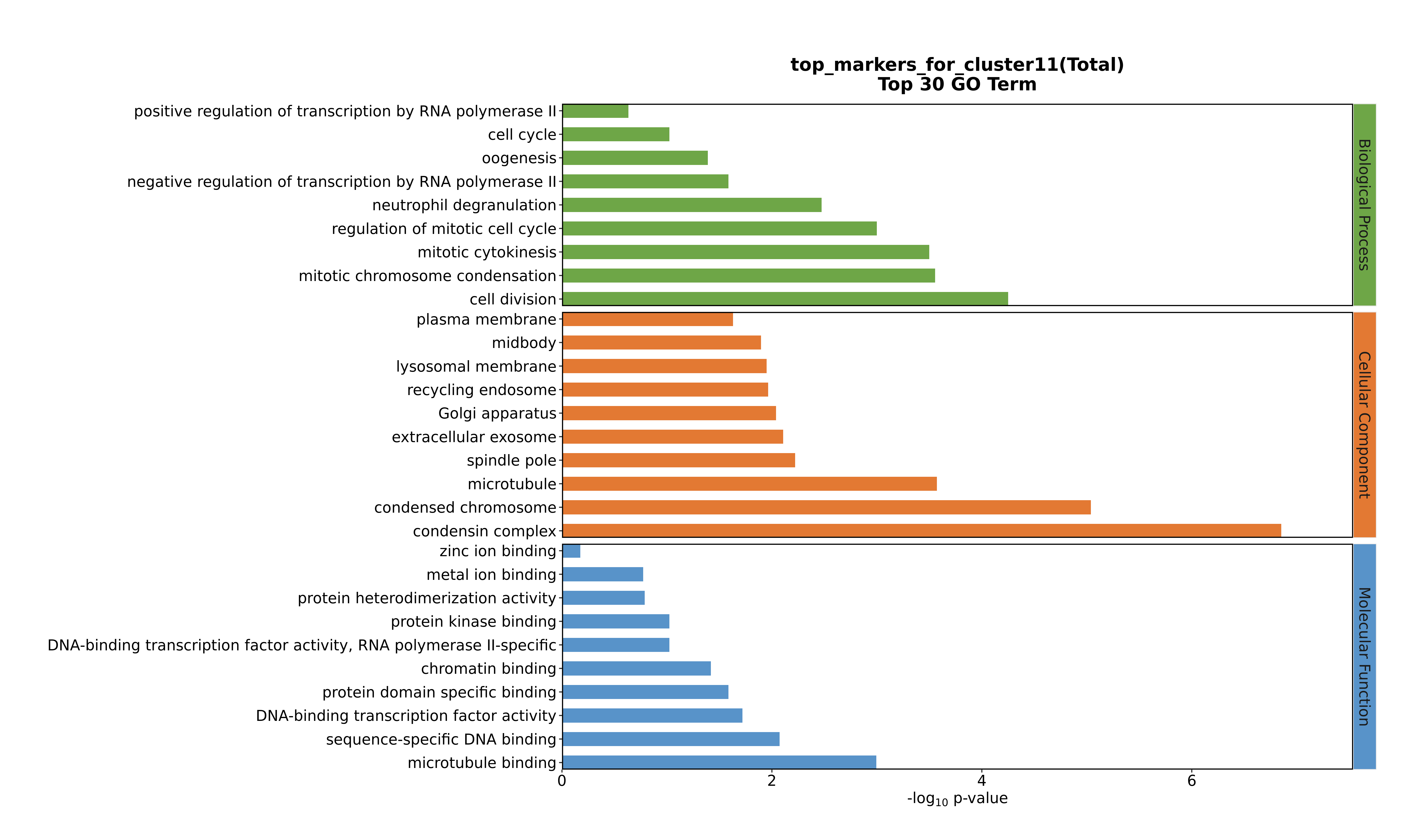

### Cluster11 GO top100.png

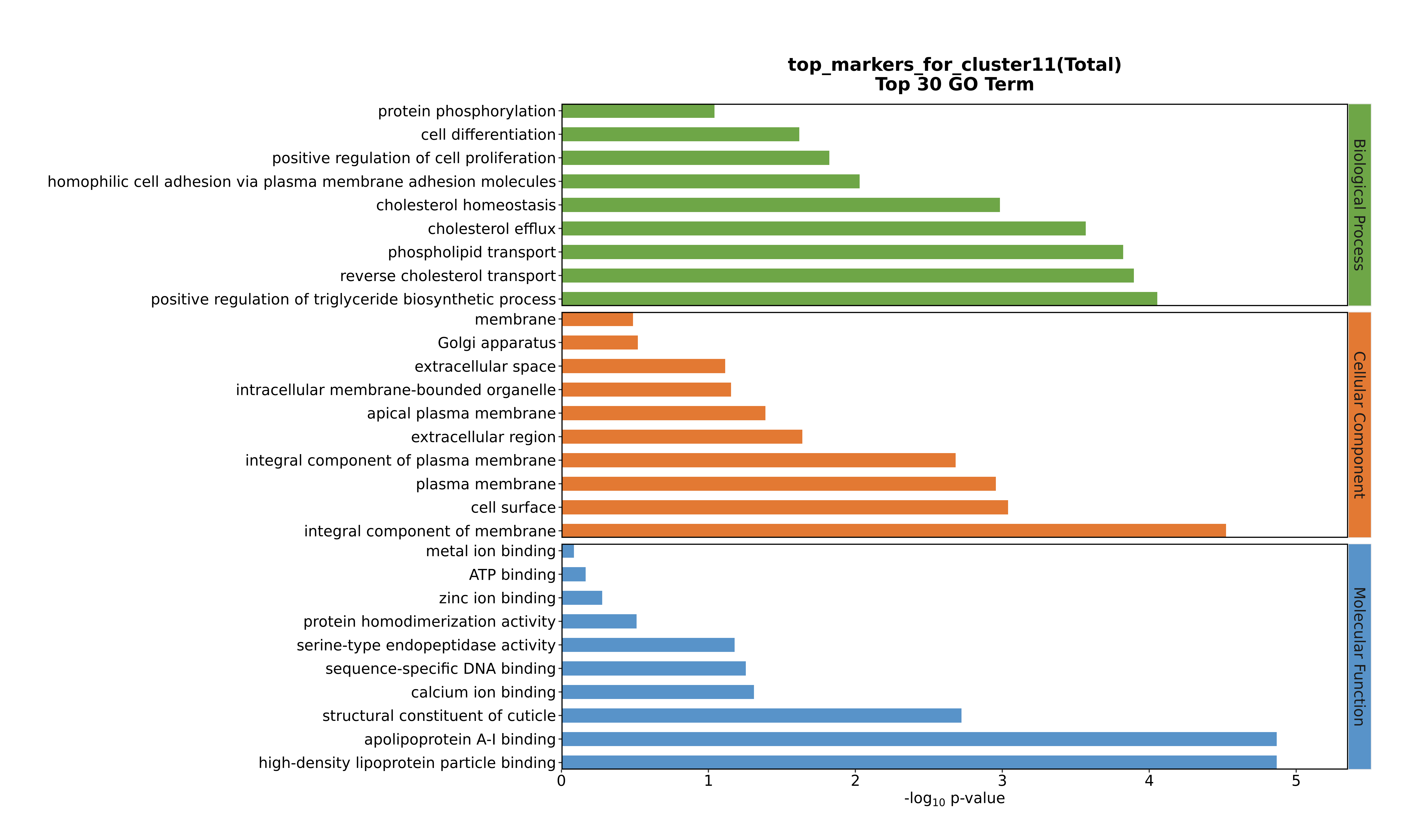

### Cluster11 GO top100.png

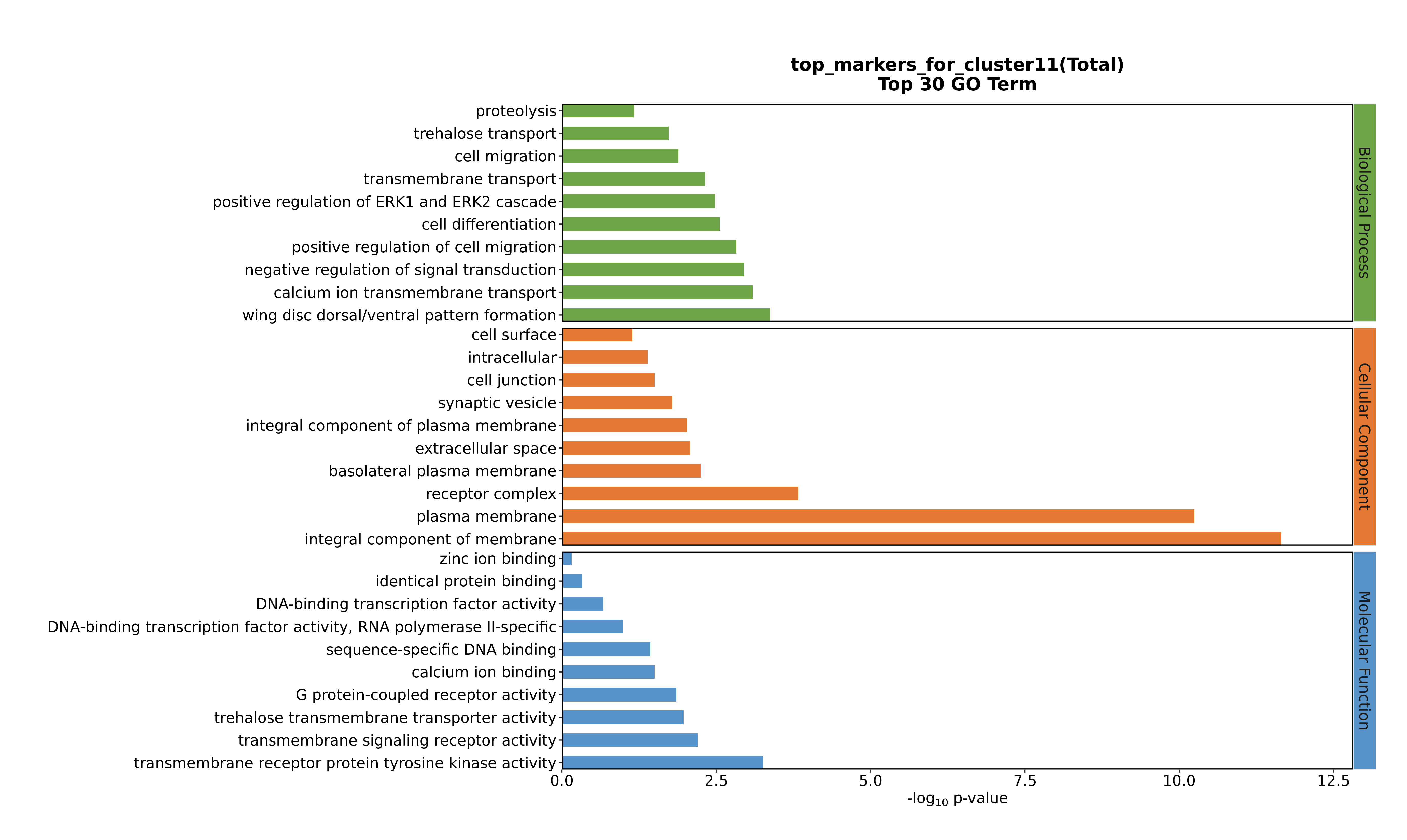

### Cluster12 GO top100.png

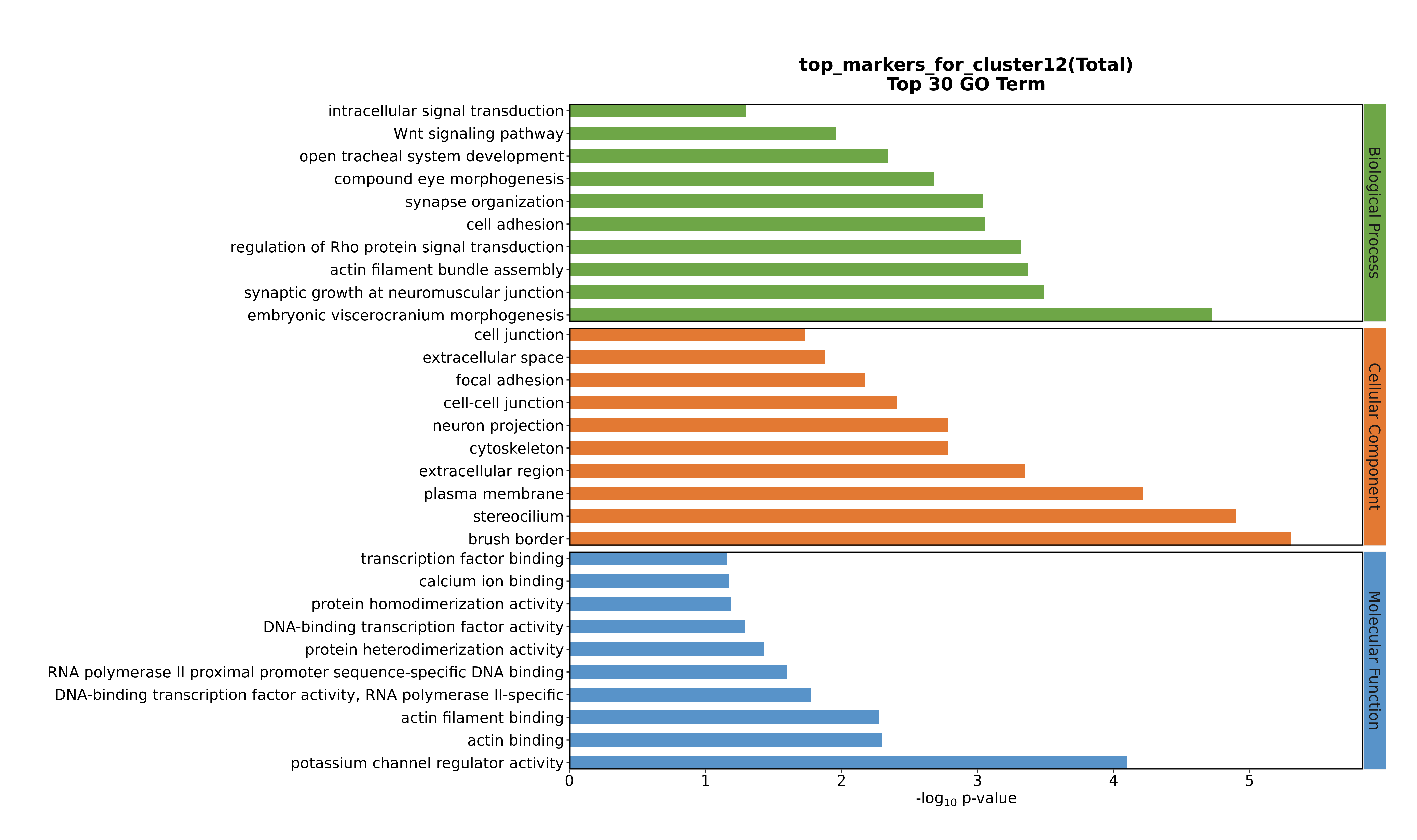

### Cluster12 GO top100.png

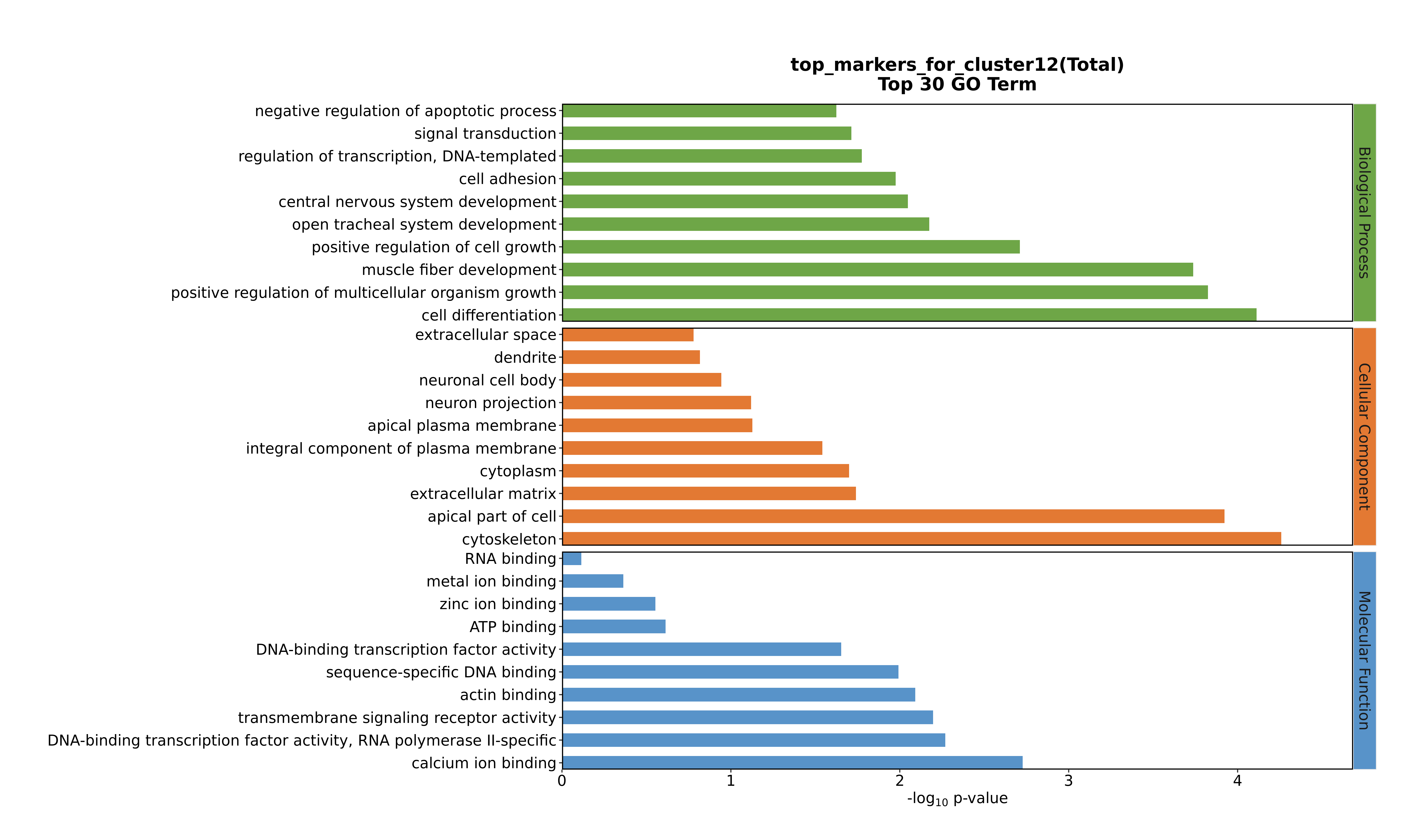

### Cluster12 GO top100.png

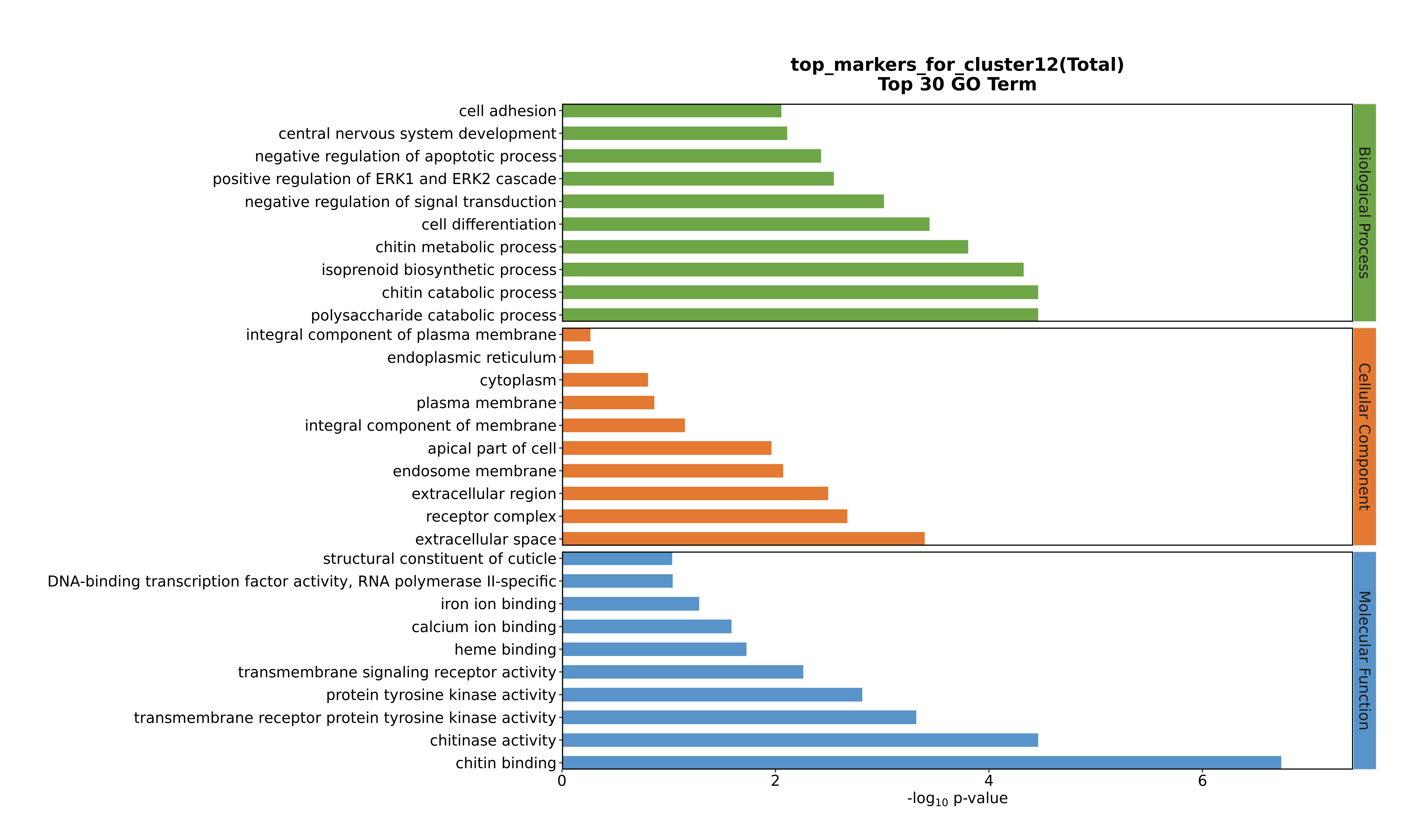

### Cluster12 GO top100.png

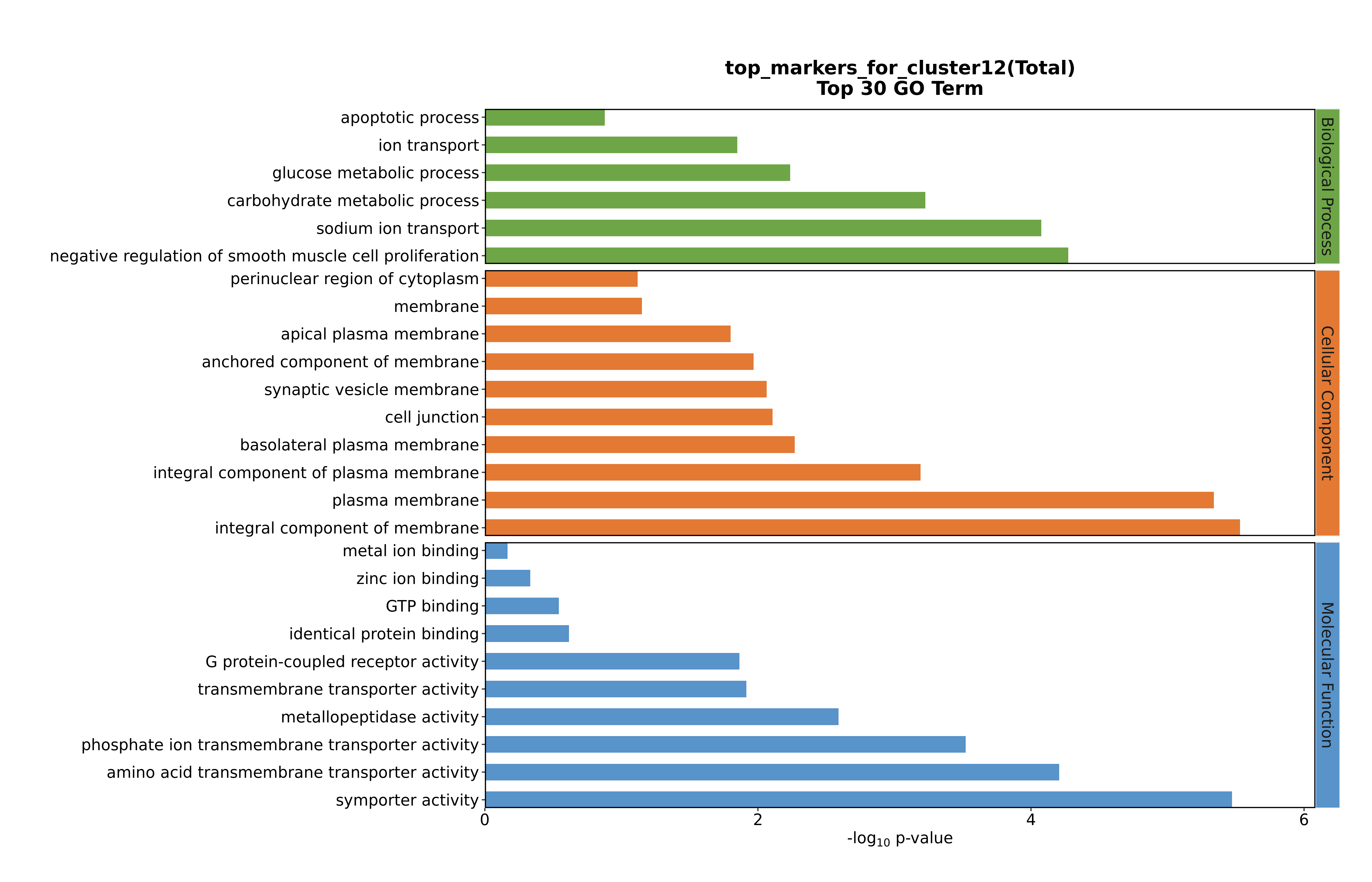

### Cluster12 GO top100.png

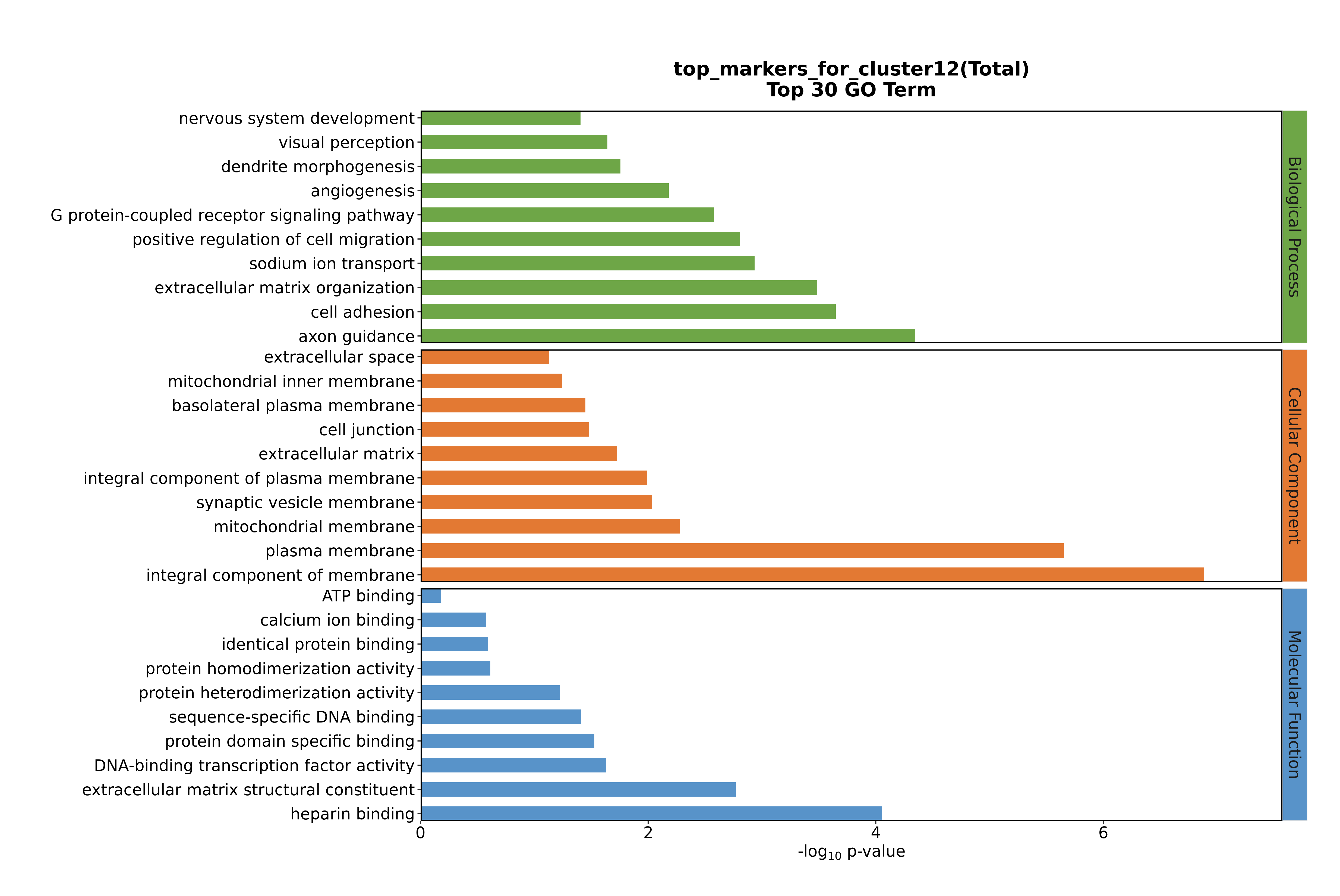

### Cluster13 GO top100.png

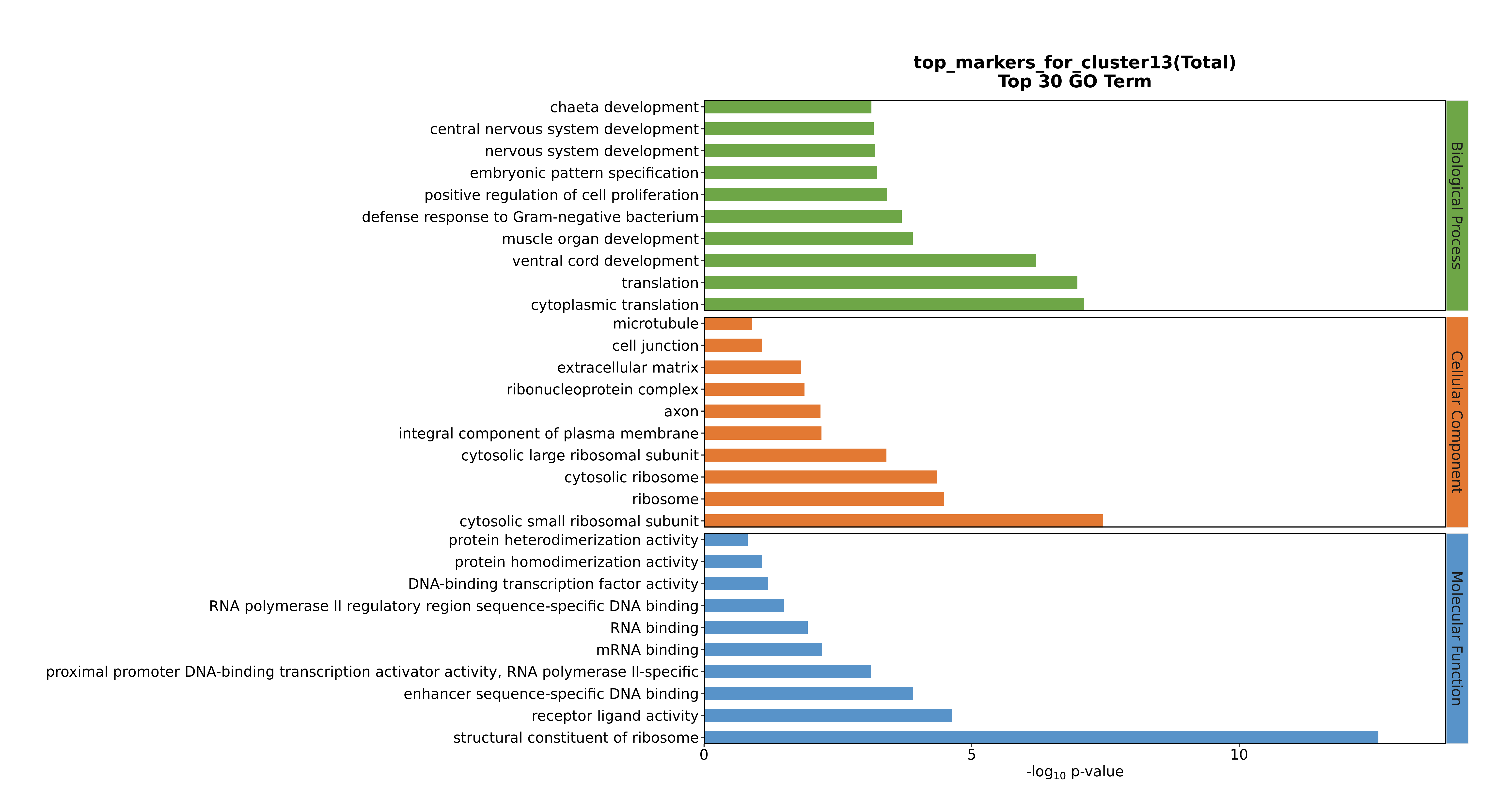

### Cluster13 GO top100.png

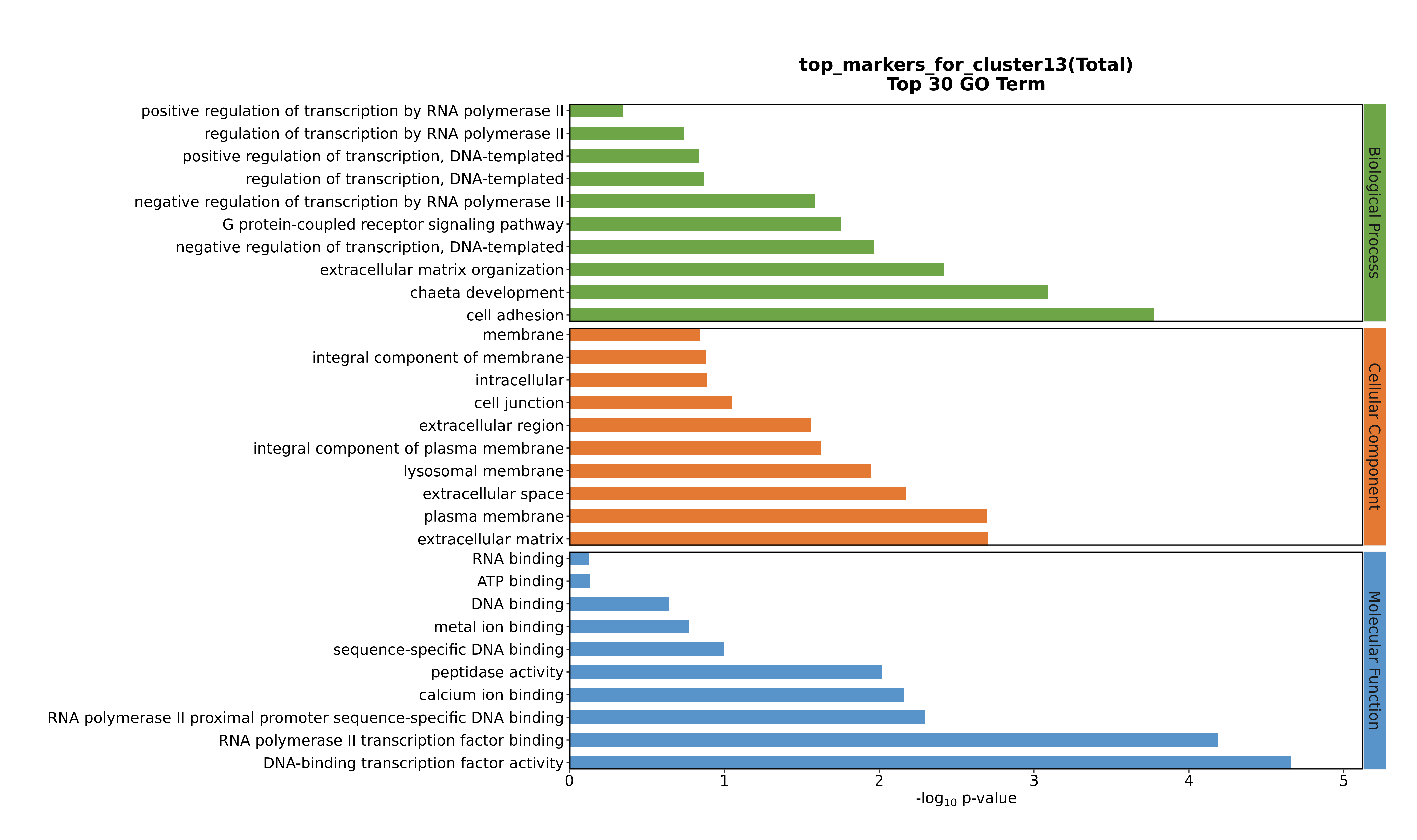

### Cluster13 GO top100.png

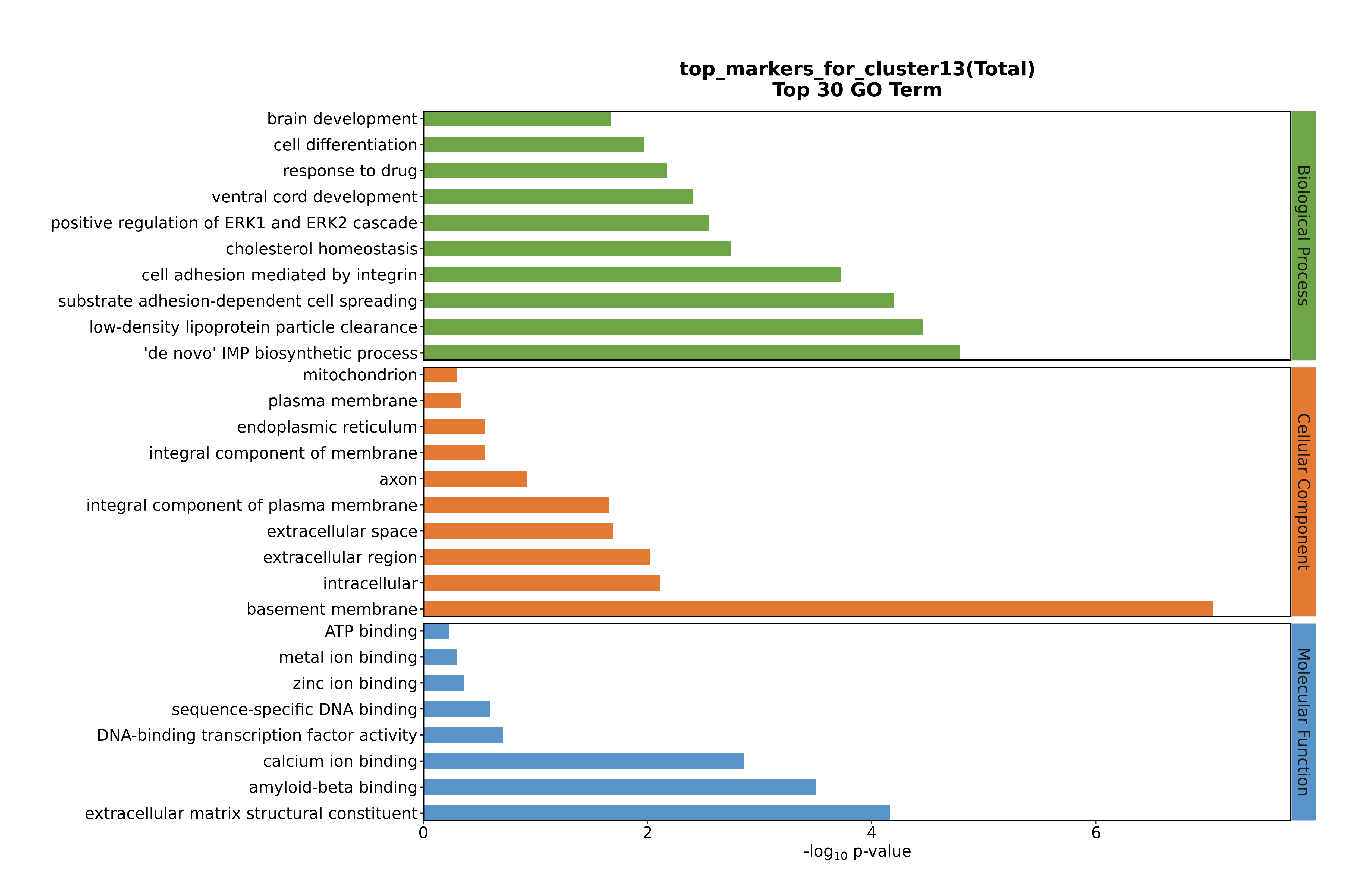

### Cluster13 GO top100.png

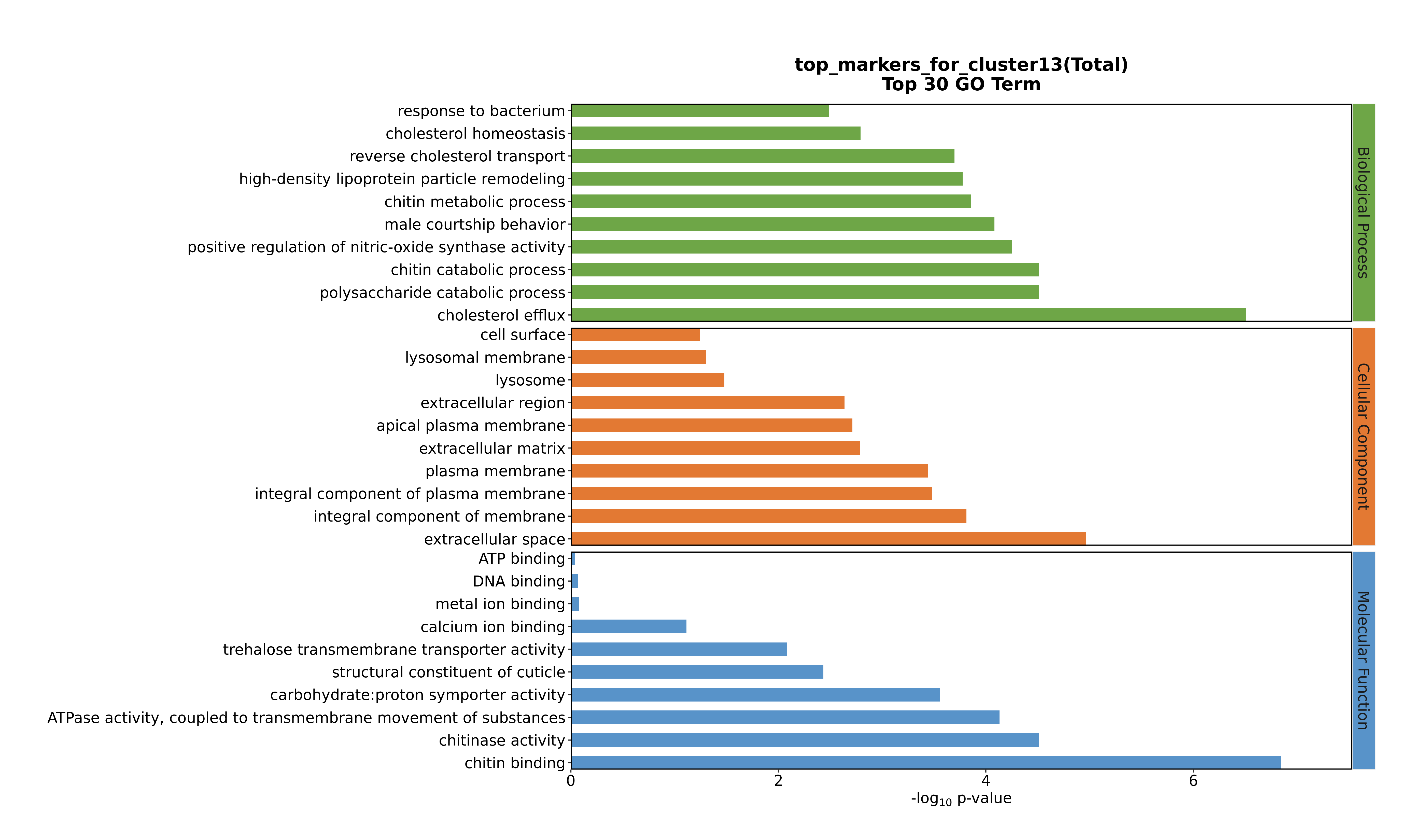

### Cluster14 GO top100.png

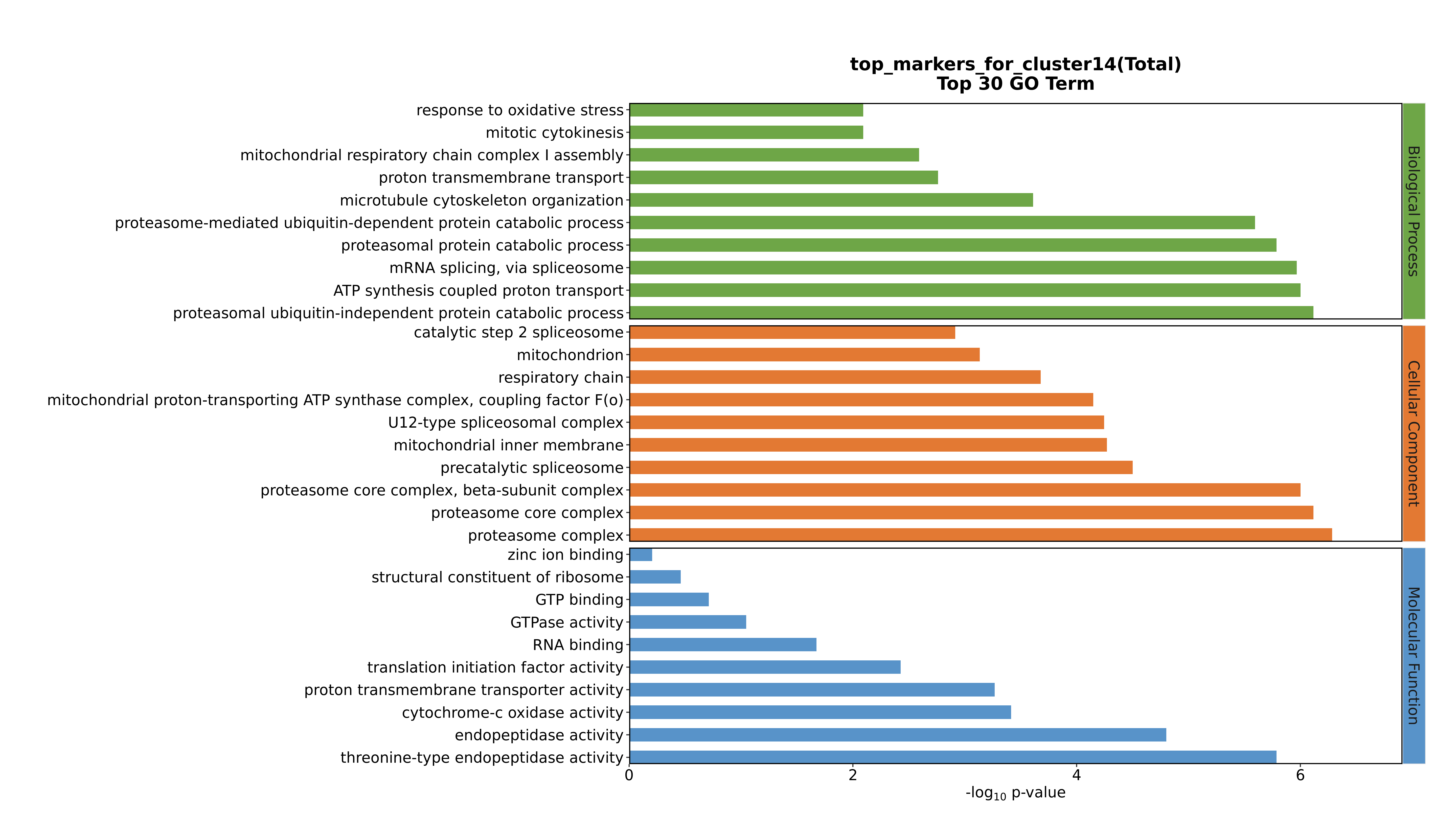

### Cluster14 GO top100.png

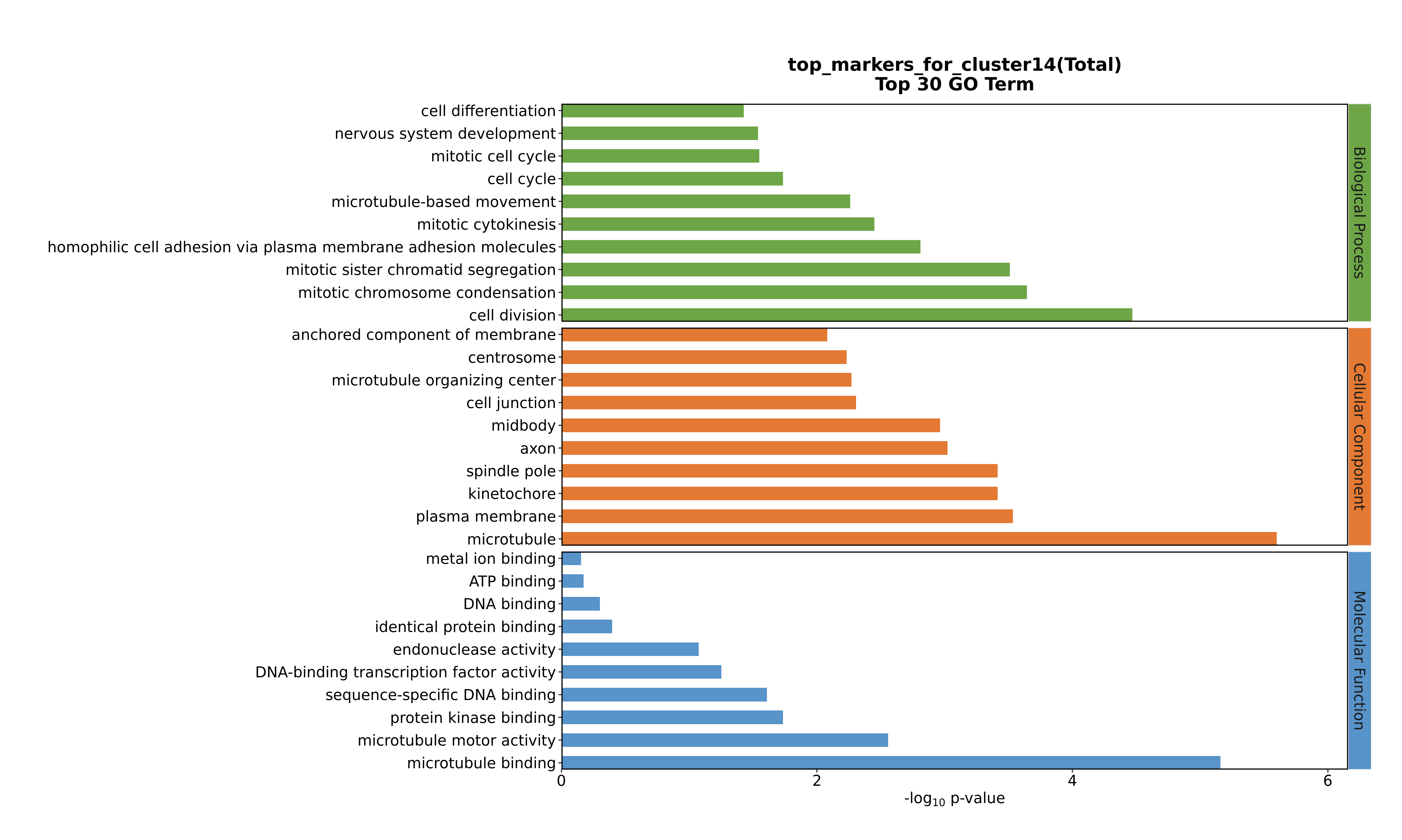

### Cluster14 GO top100.png

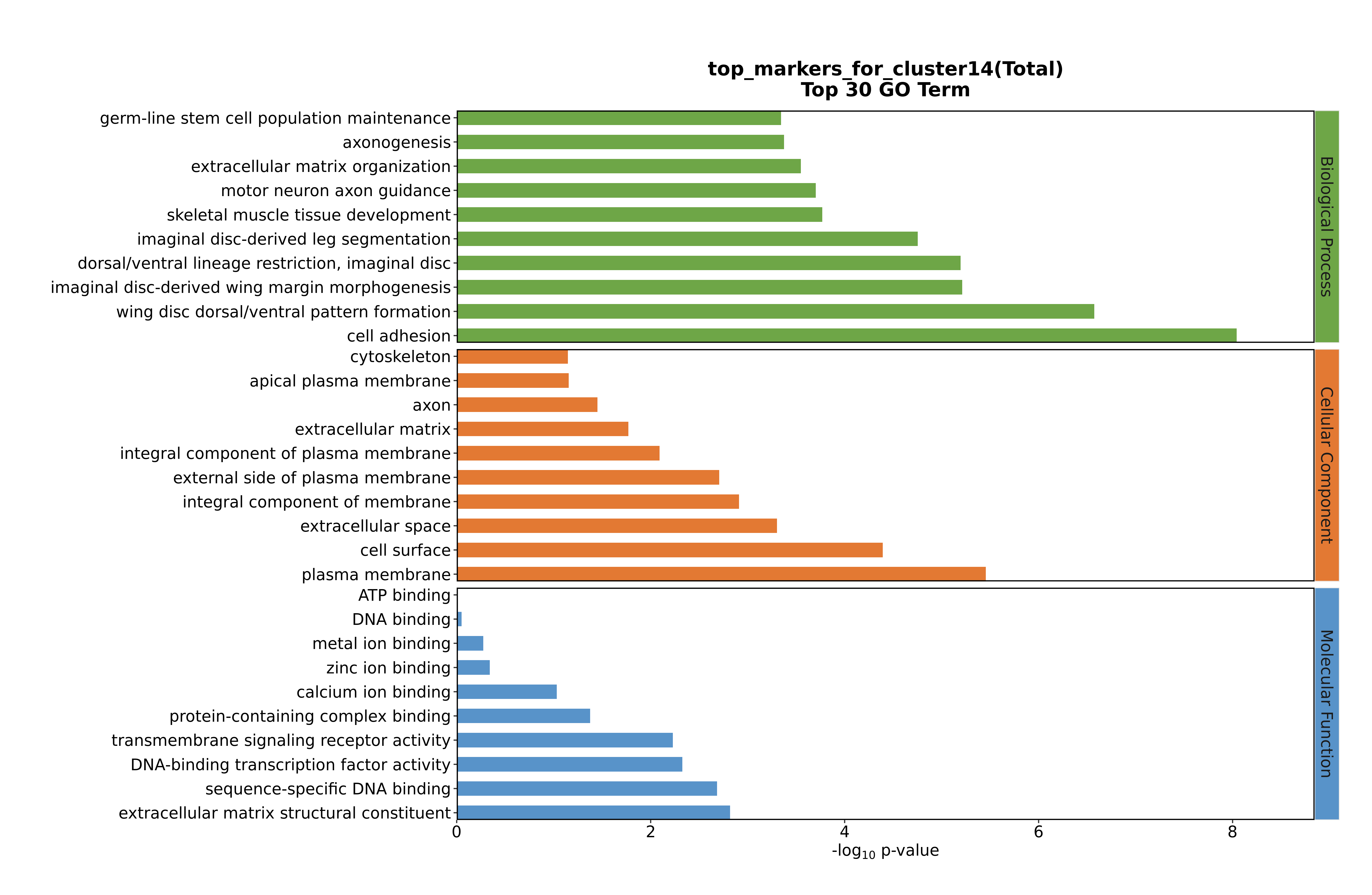

### Cluster14 GO top100.png

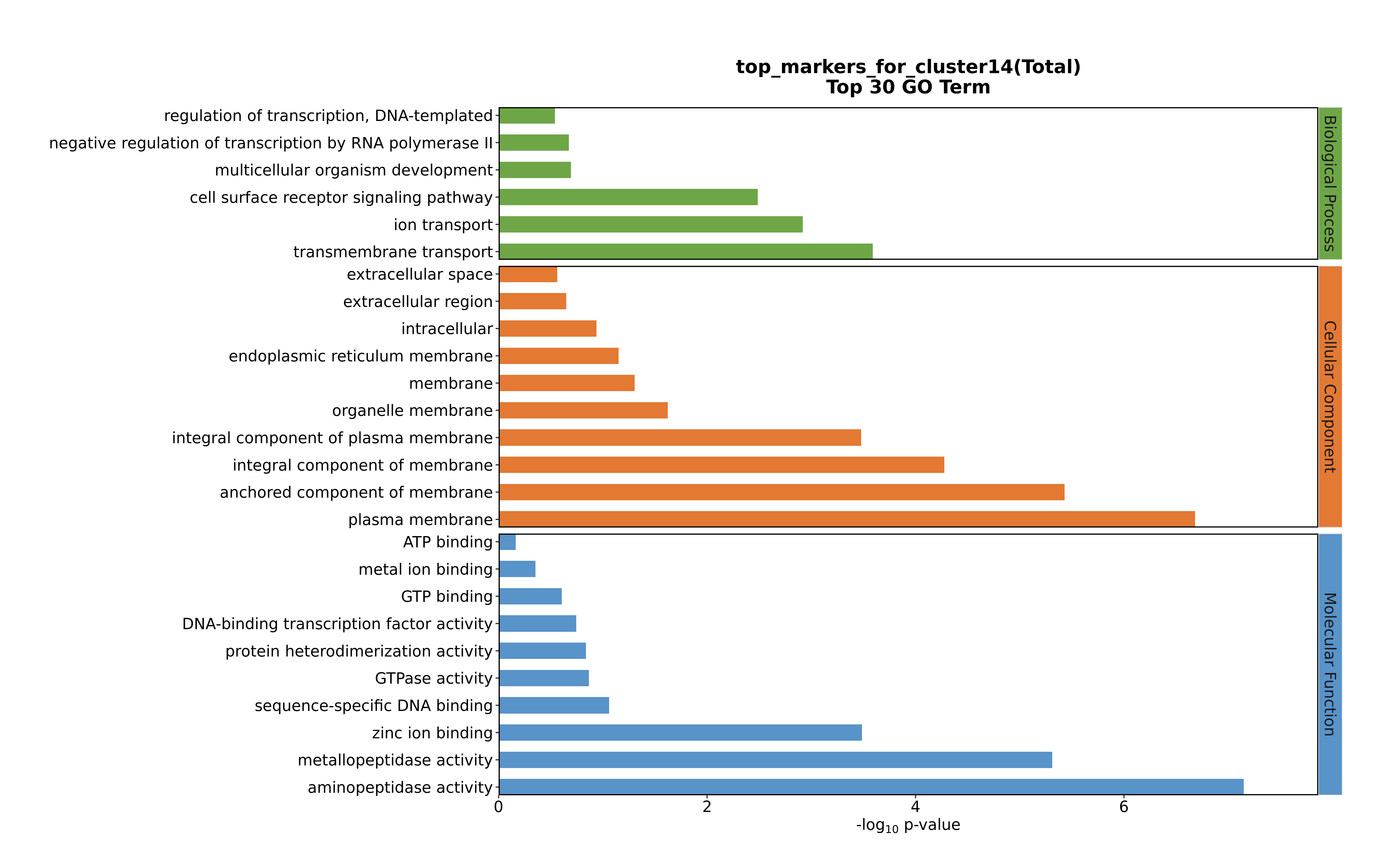

### Cluster15 GO top100.png

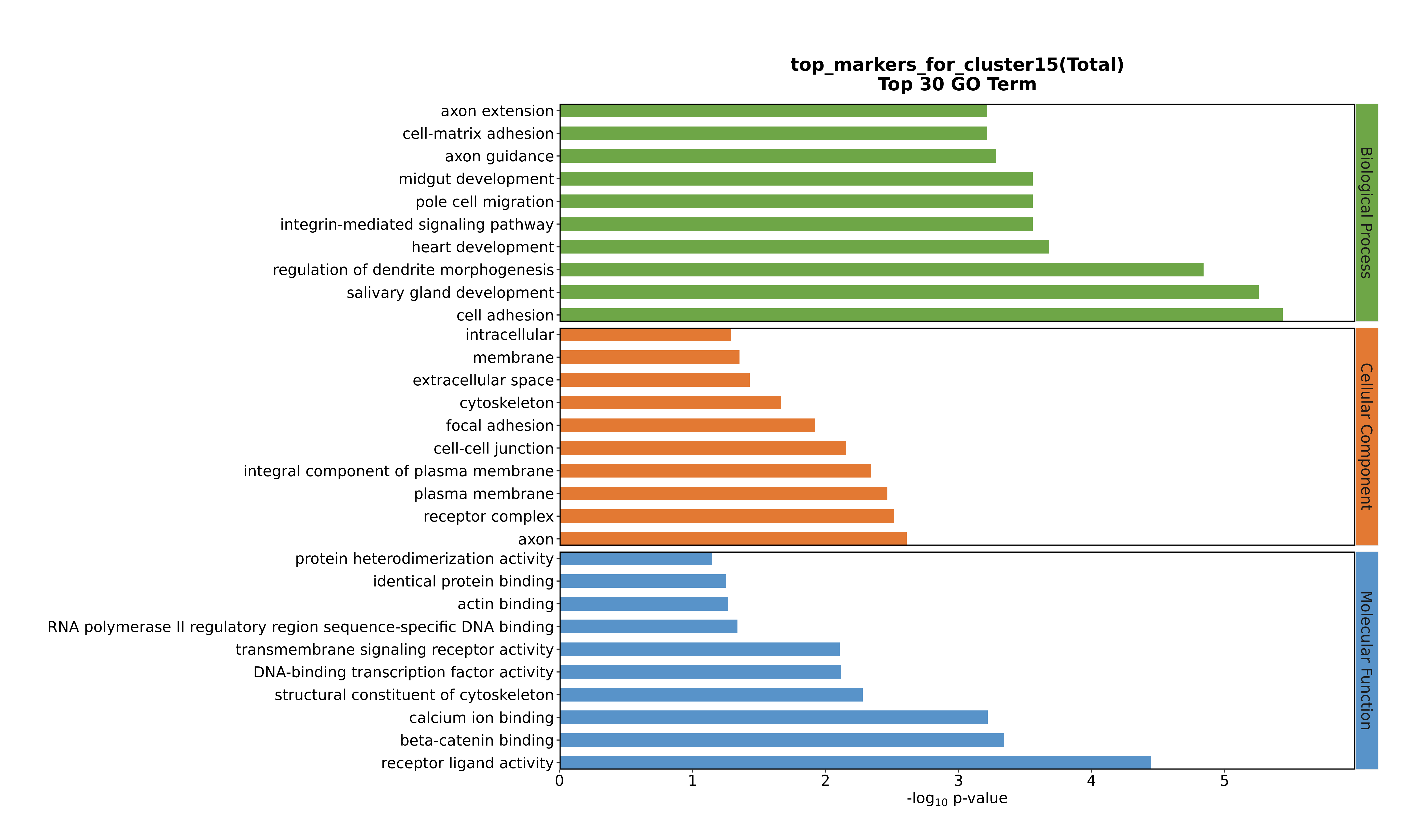

### Cluster15 GO top100.png

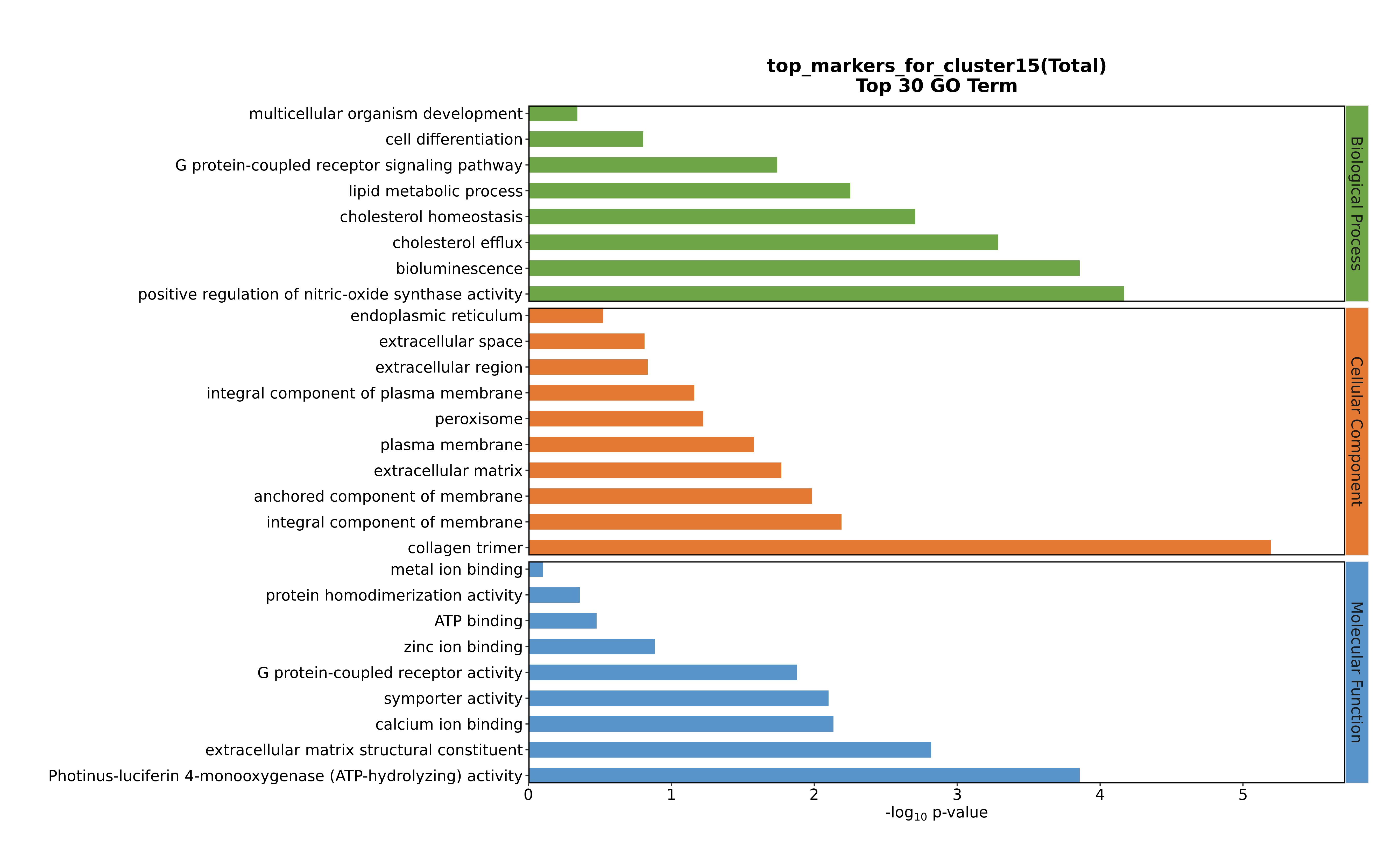

### Cluster15 GO top100.png

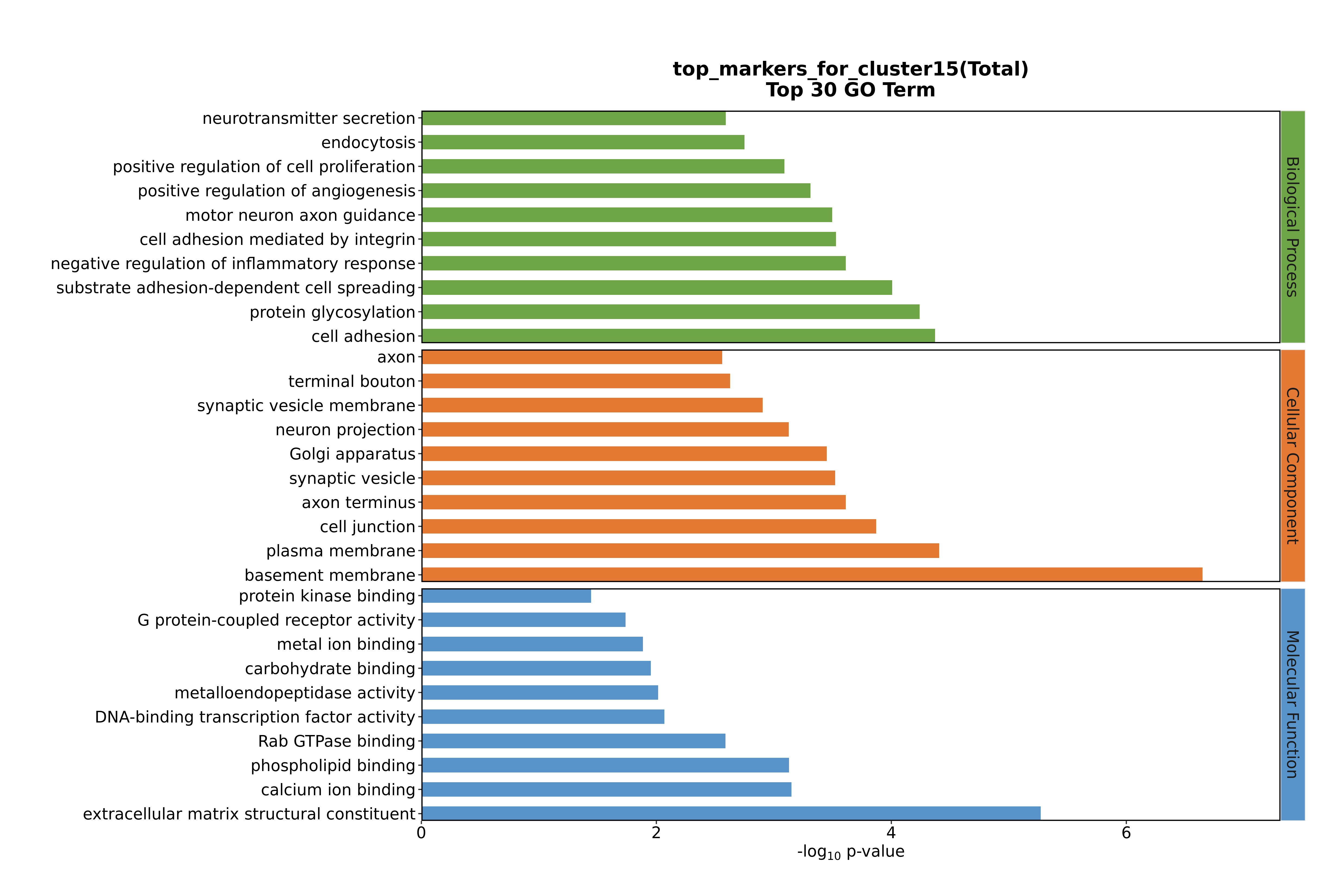

### Cluster15 GO top100.png

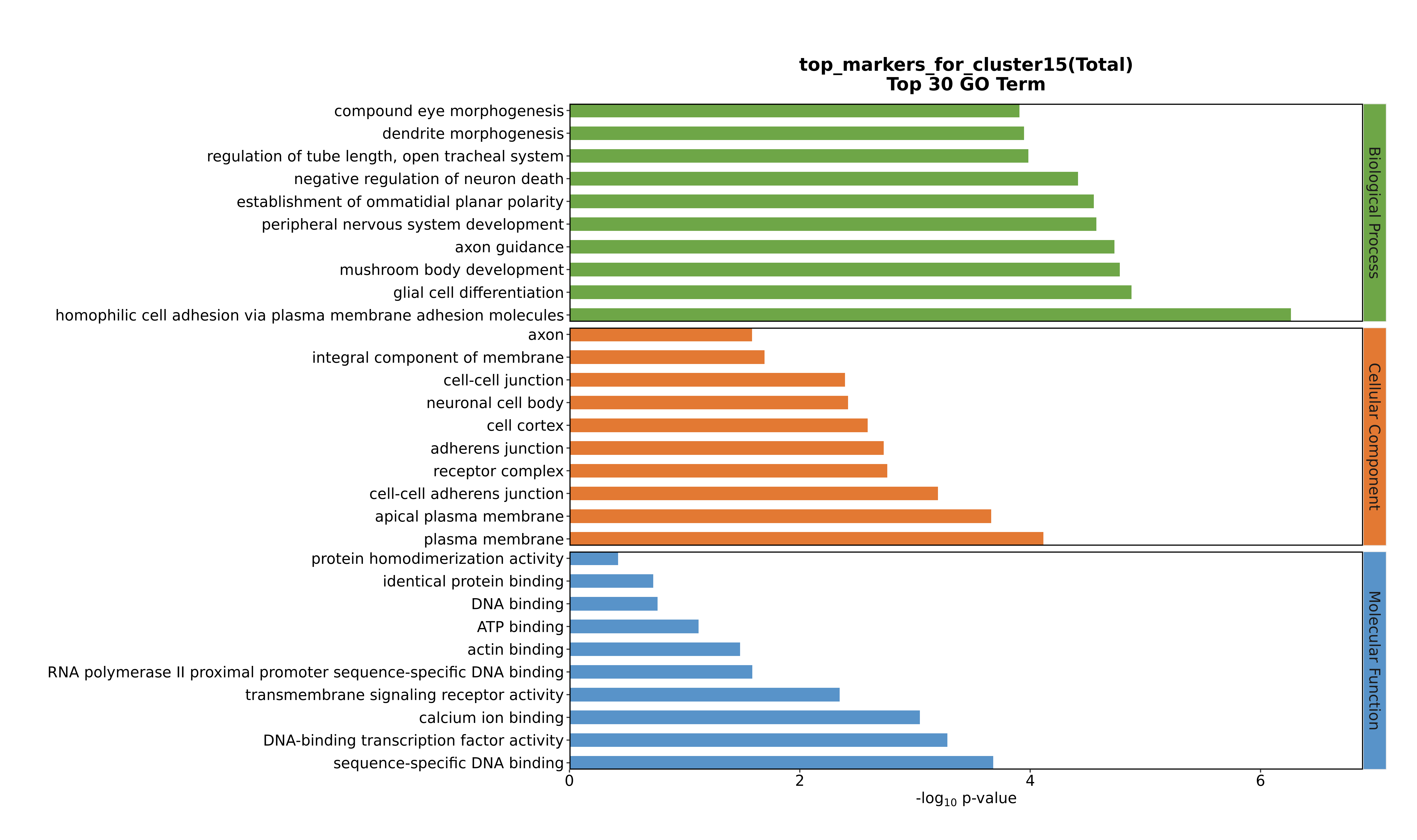

### Cluster16 GO top100.png

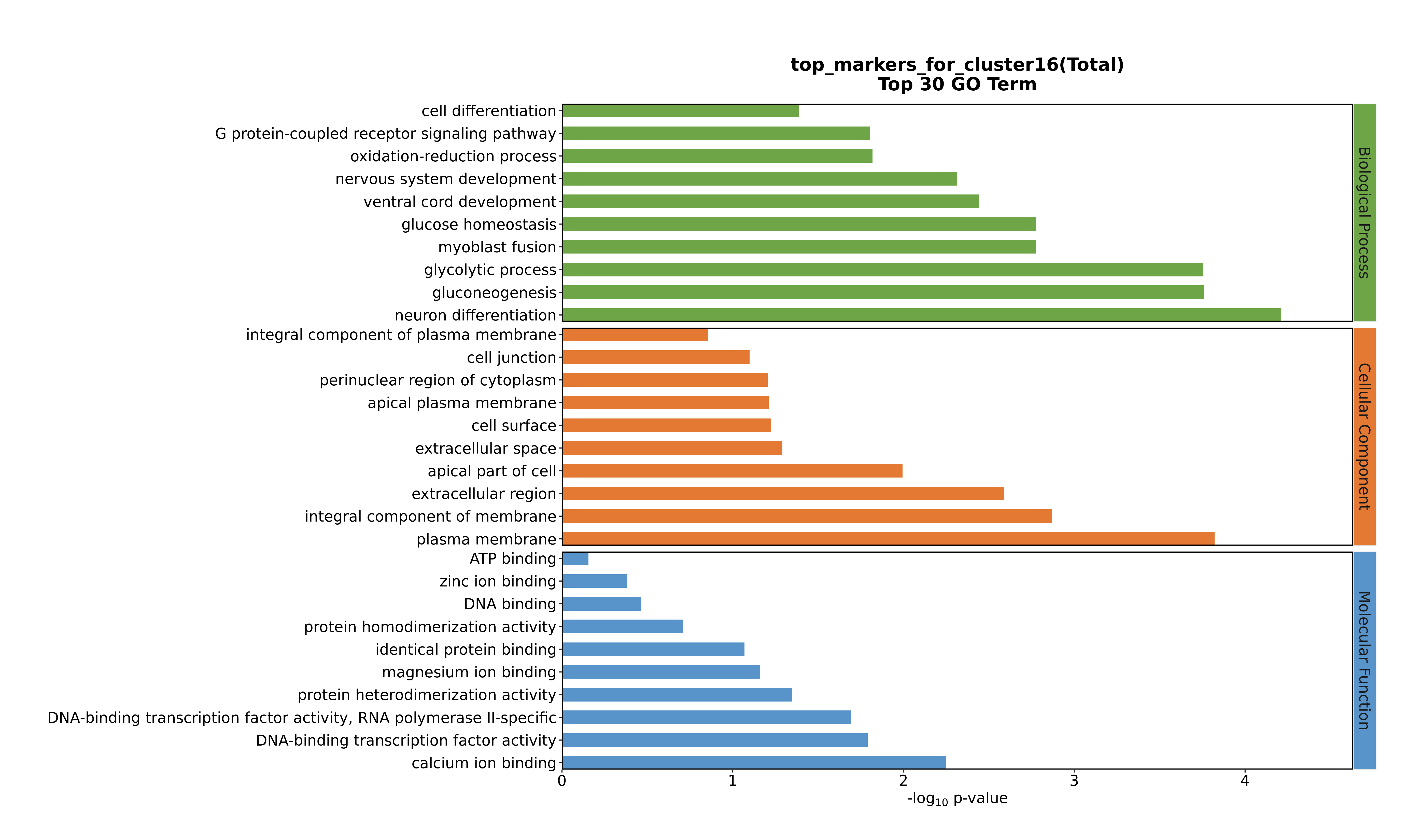

### Cluster16 GO top100.png

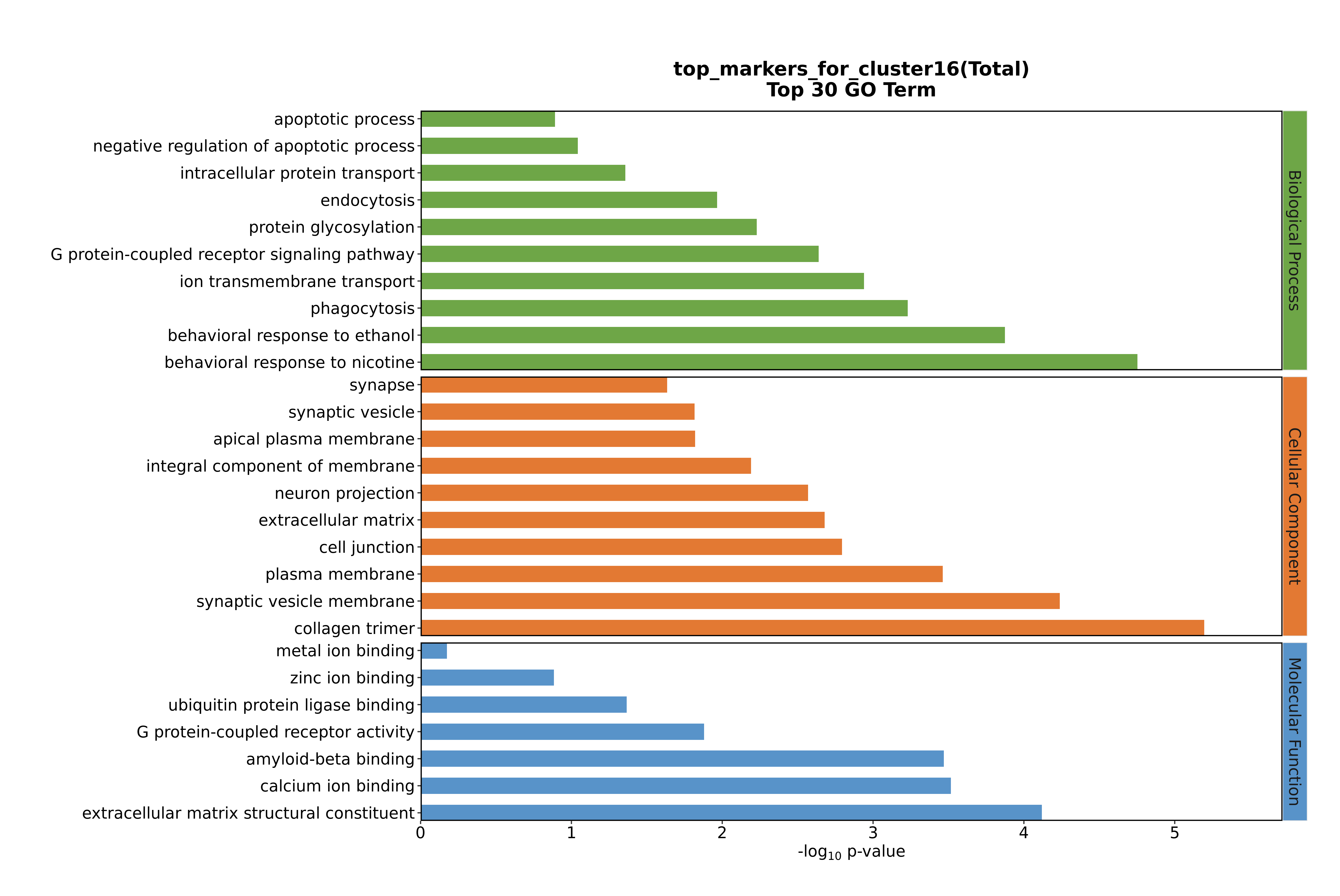

### Cluster16 GO top100.png

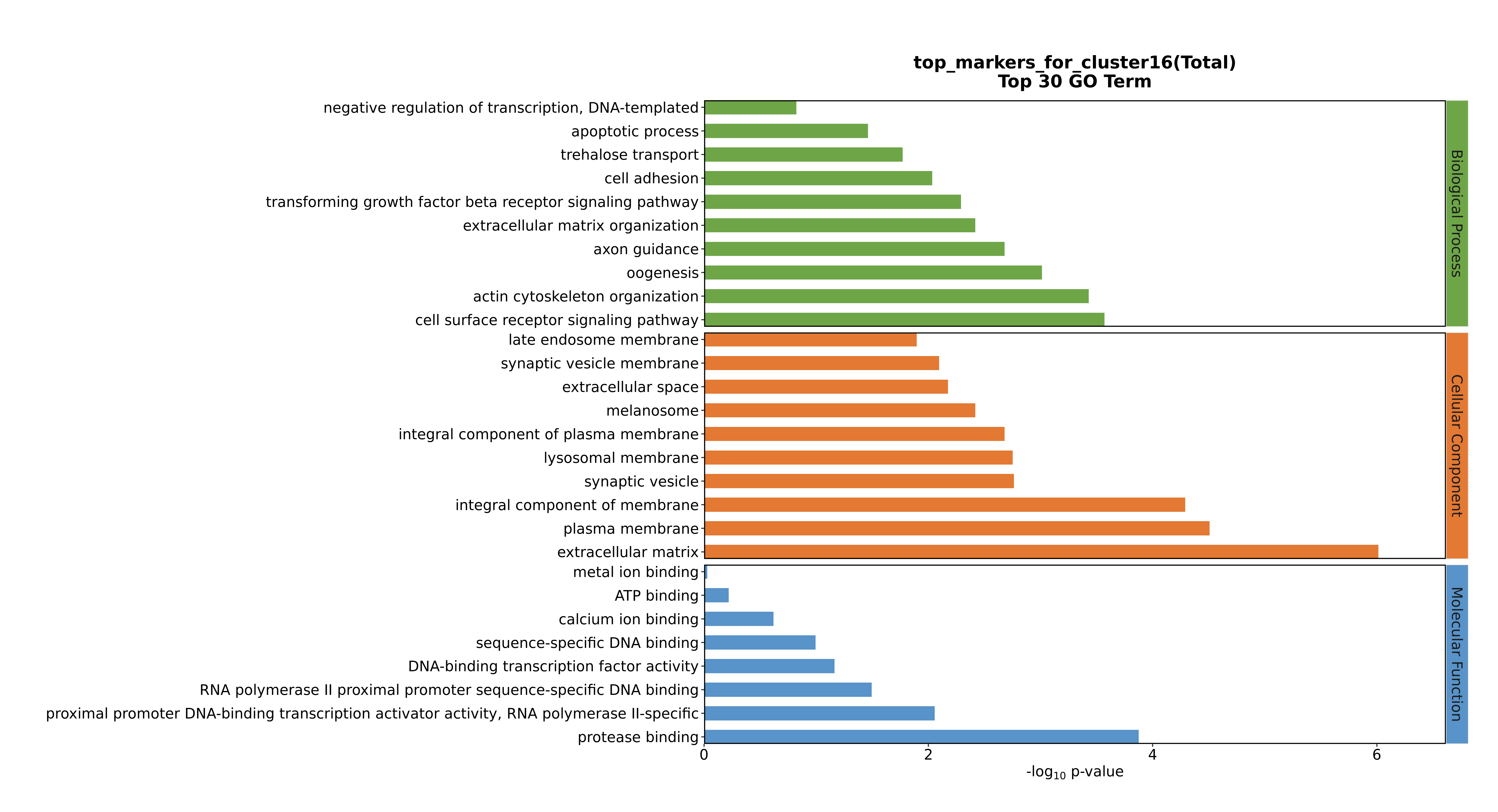

### Cluster16 GO top100.png

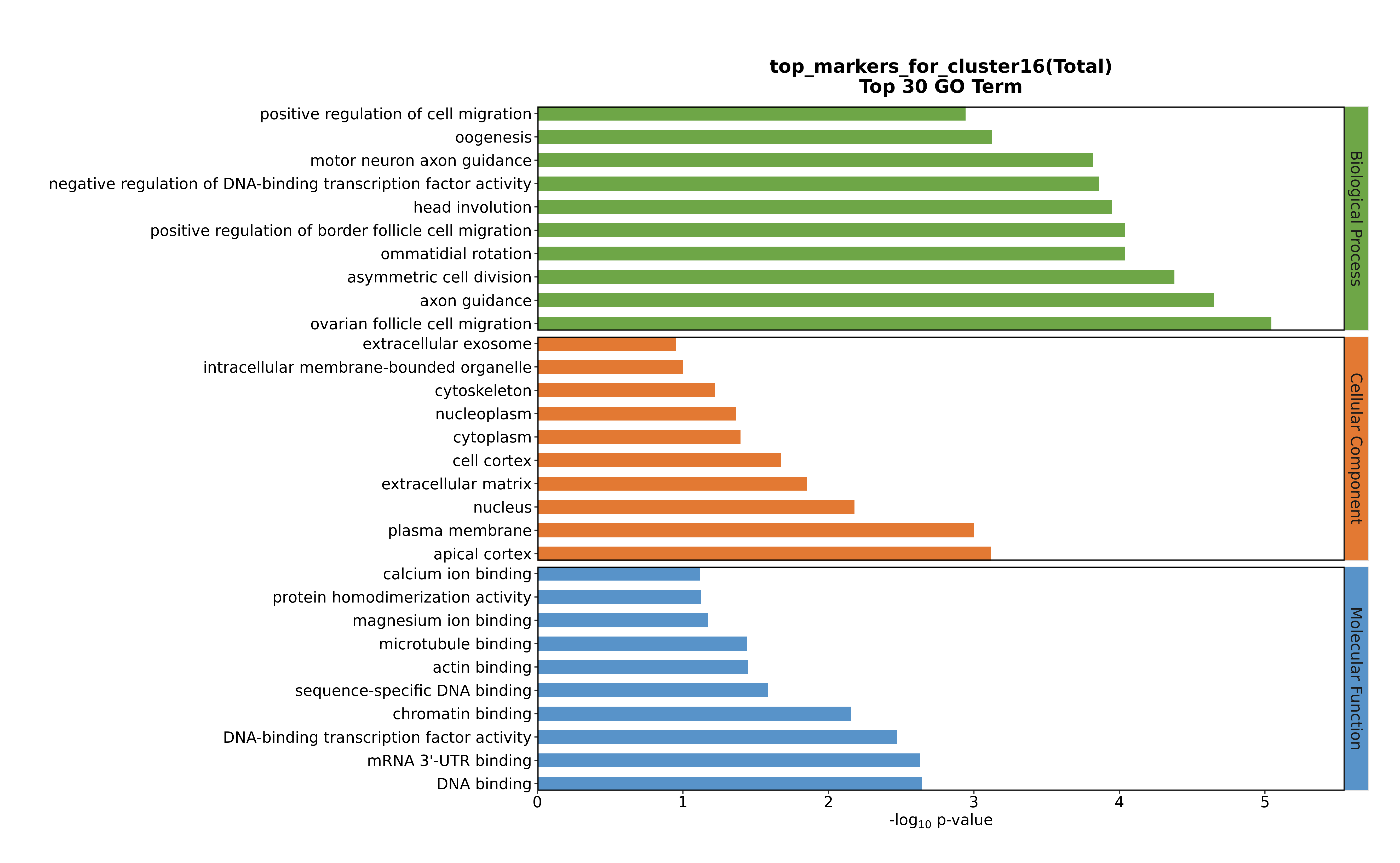

### Cluster17 GO top100.png

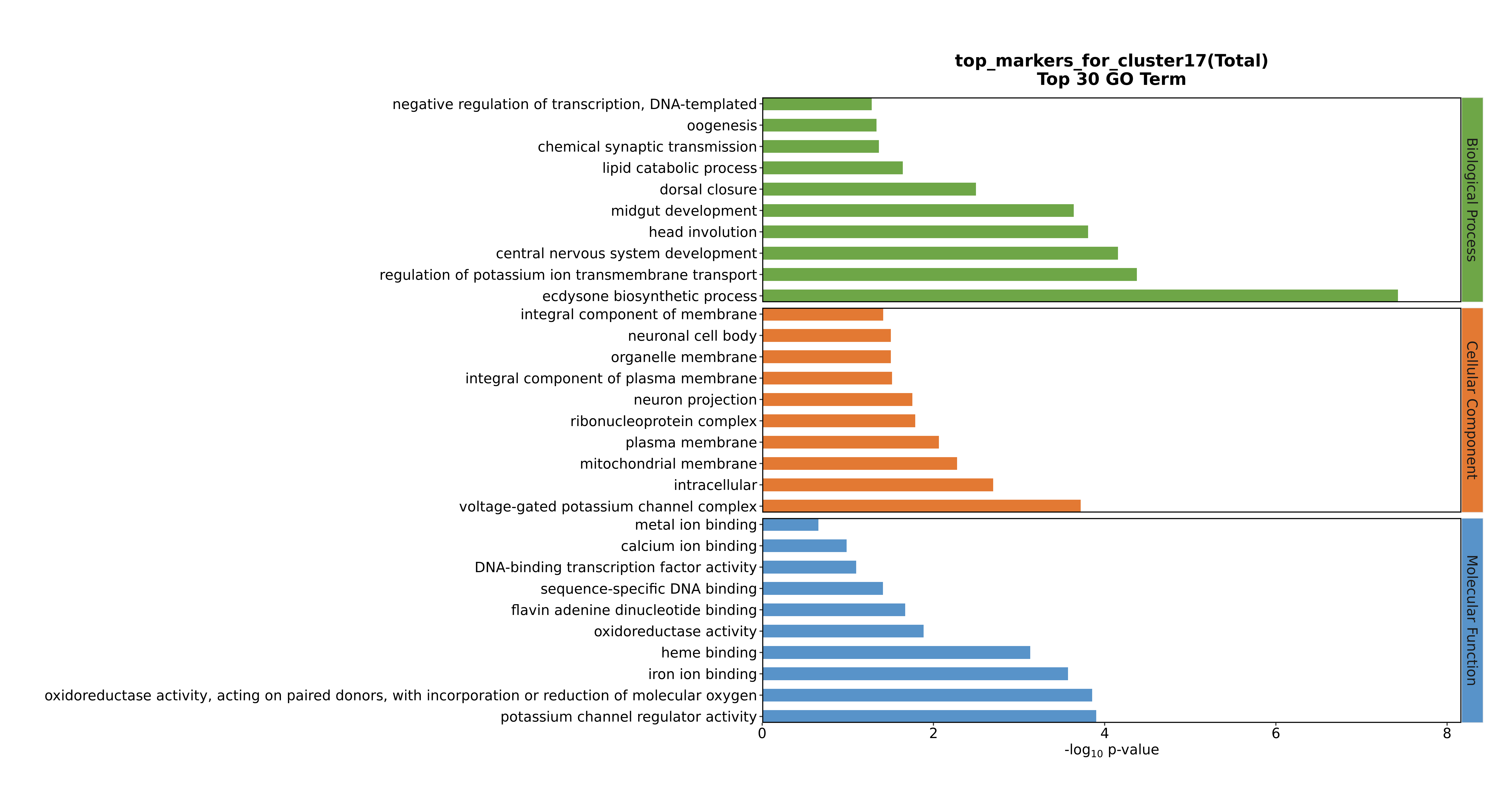

### Cluster17 GO top100.png

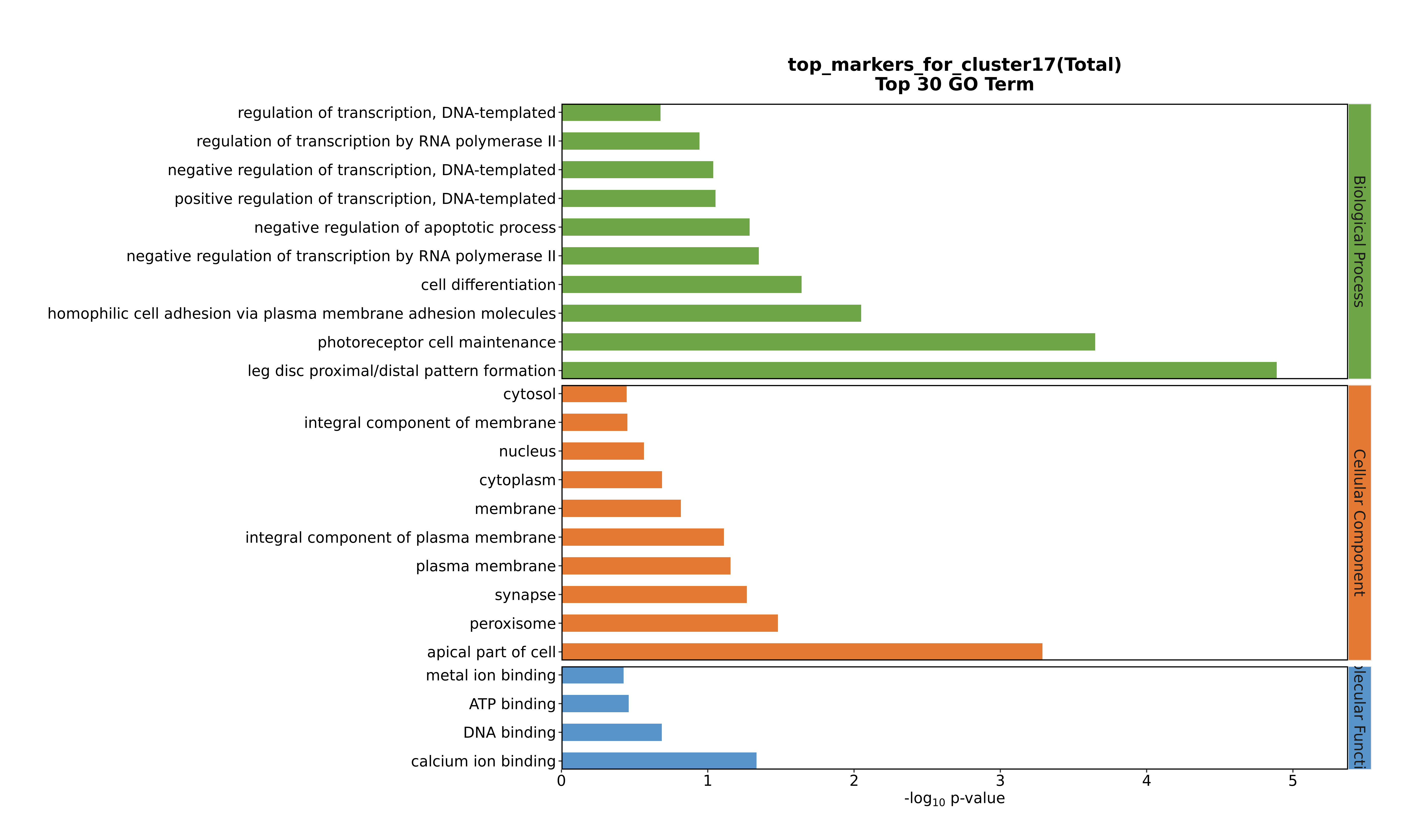

### Cluster17 GO top100.png

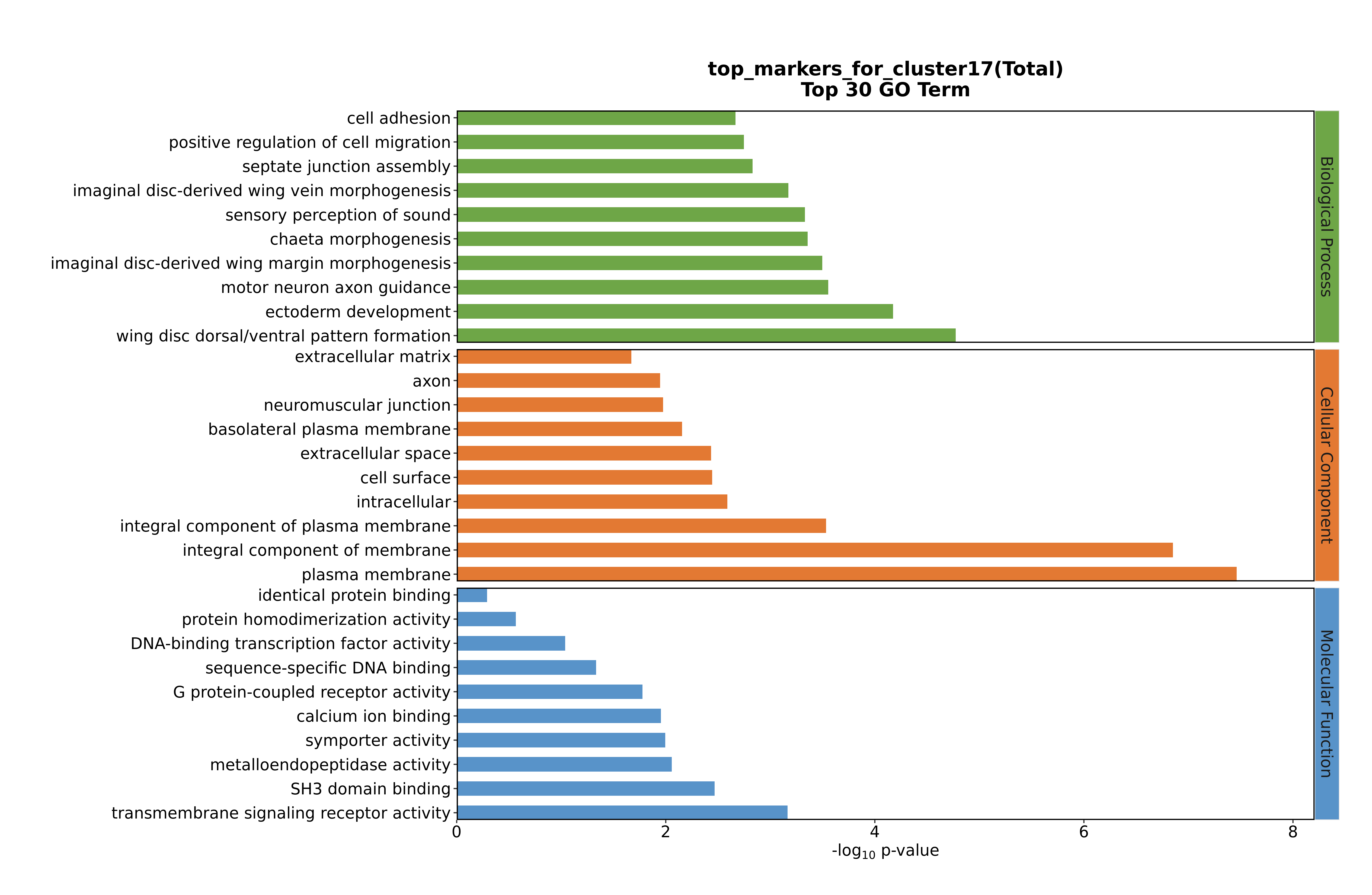

### Cluster17 GO top100.png

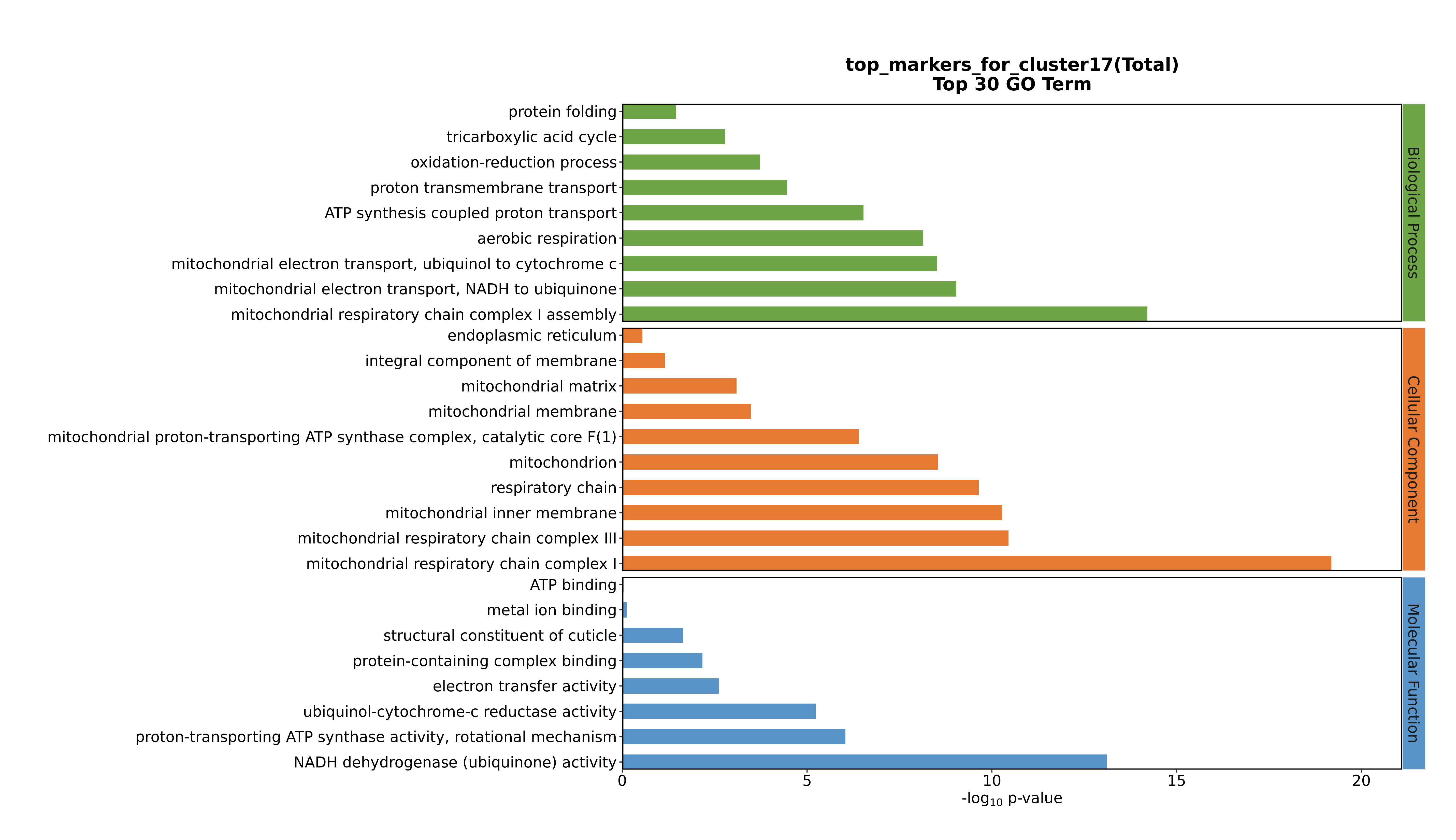

### Cluster18 GO top100.png

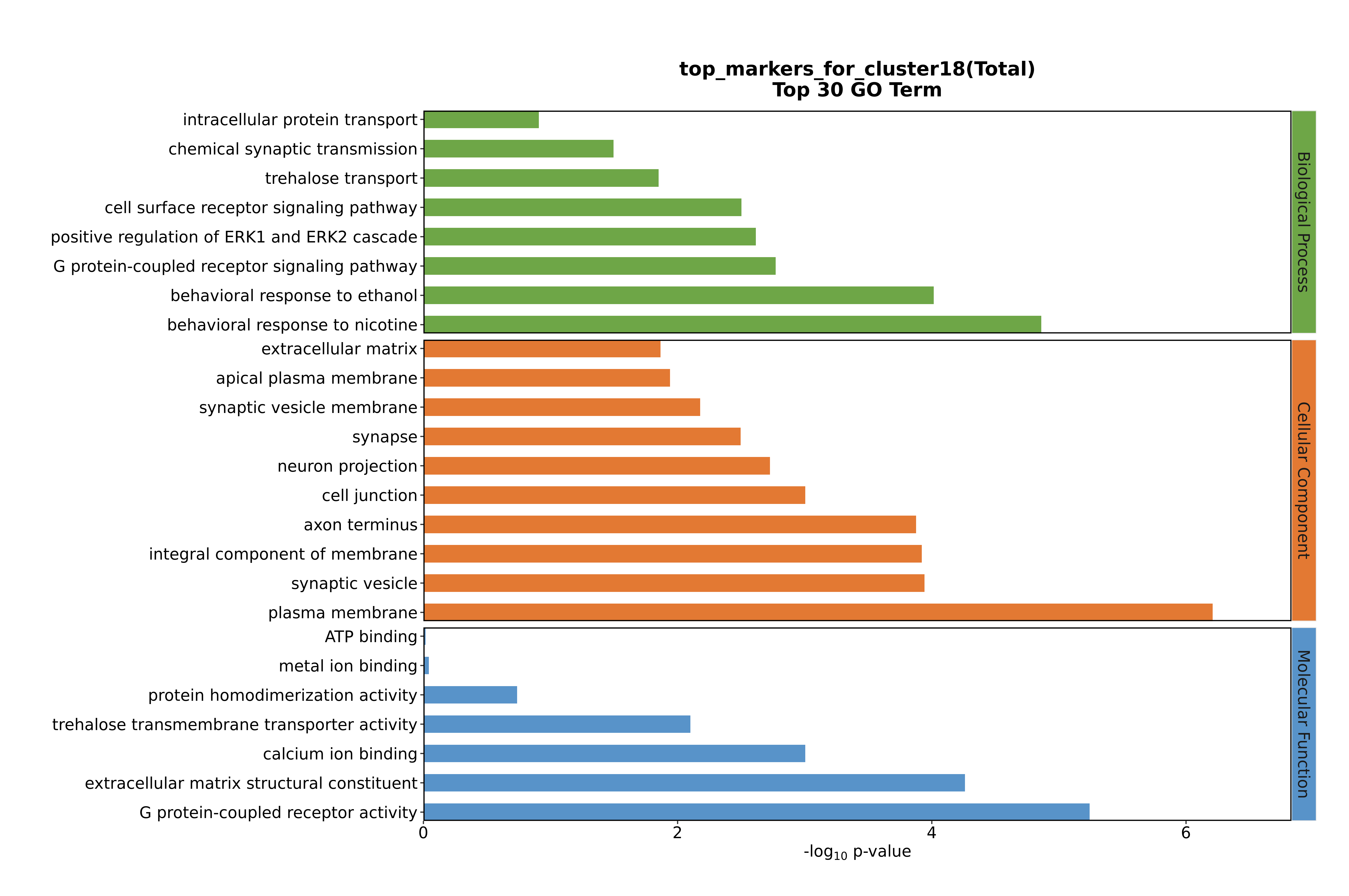

### Cluster18 GO top100.png

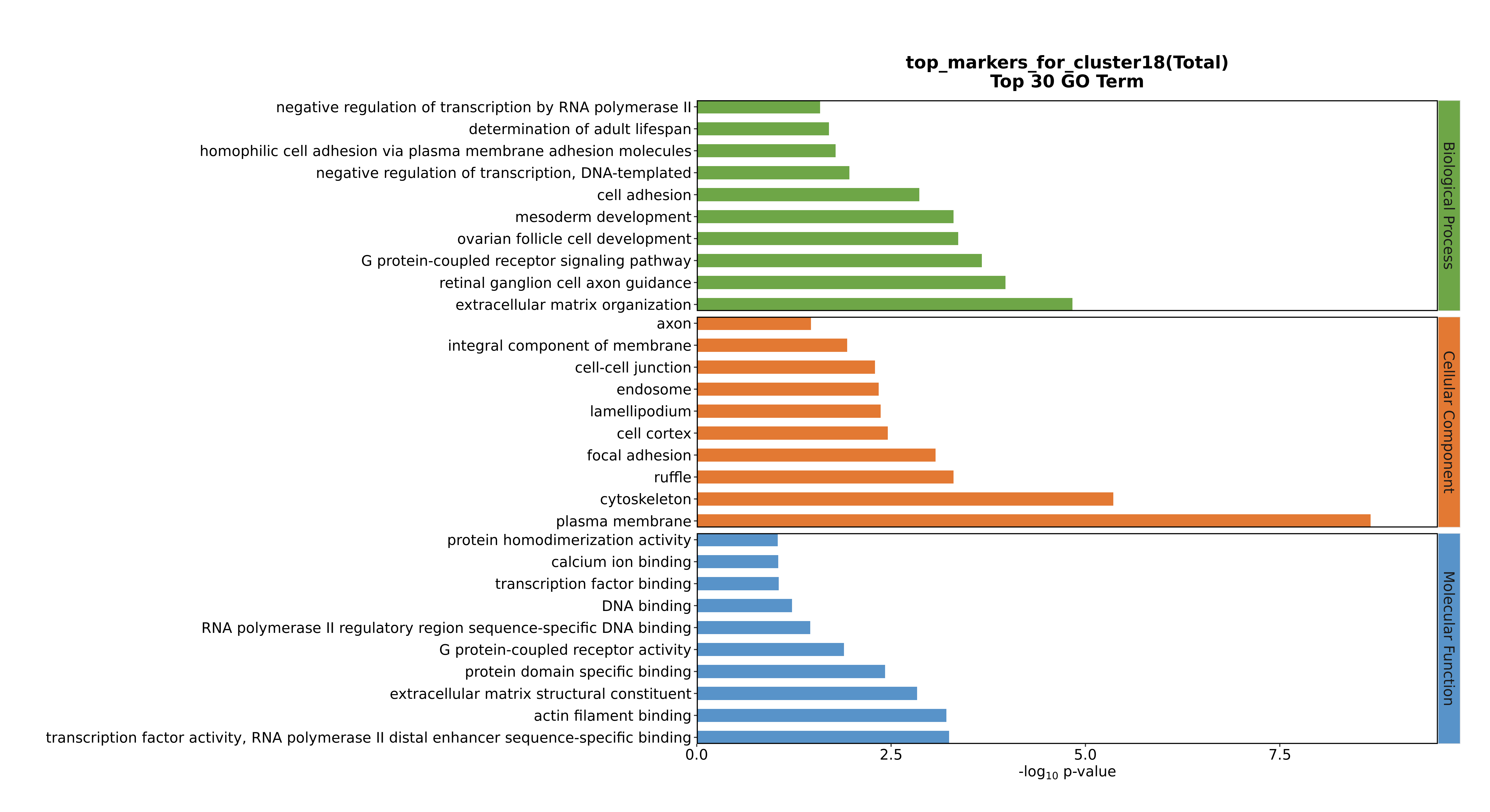

### Cluster18 GO top100.png

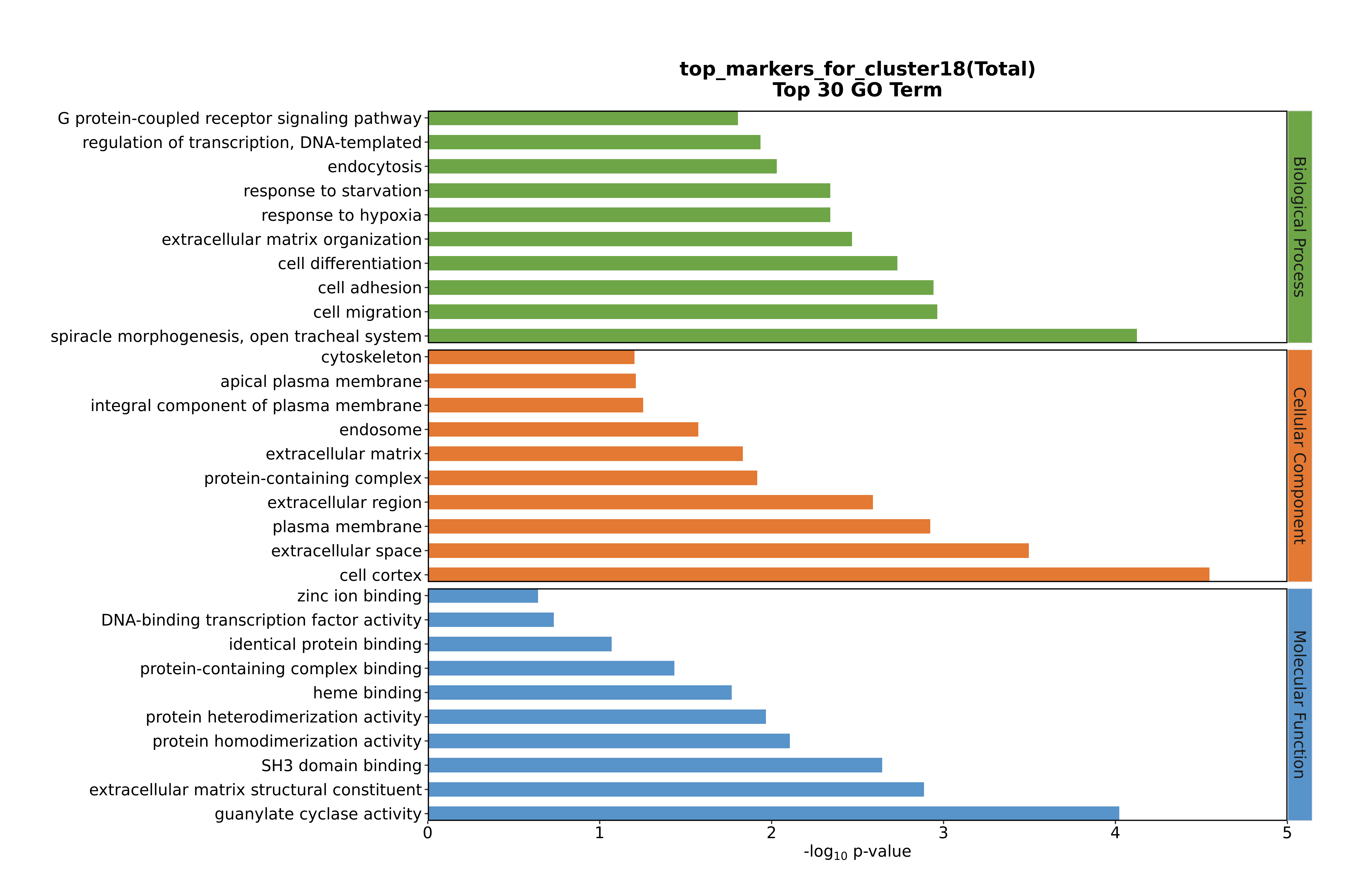

### Cluster18 GO top100.png

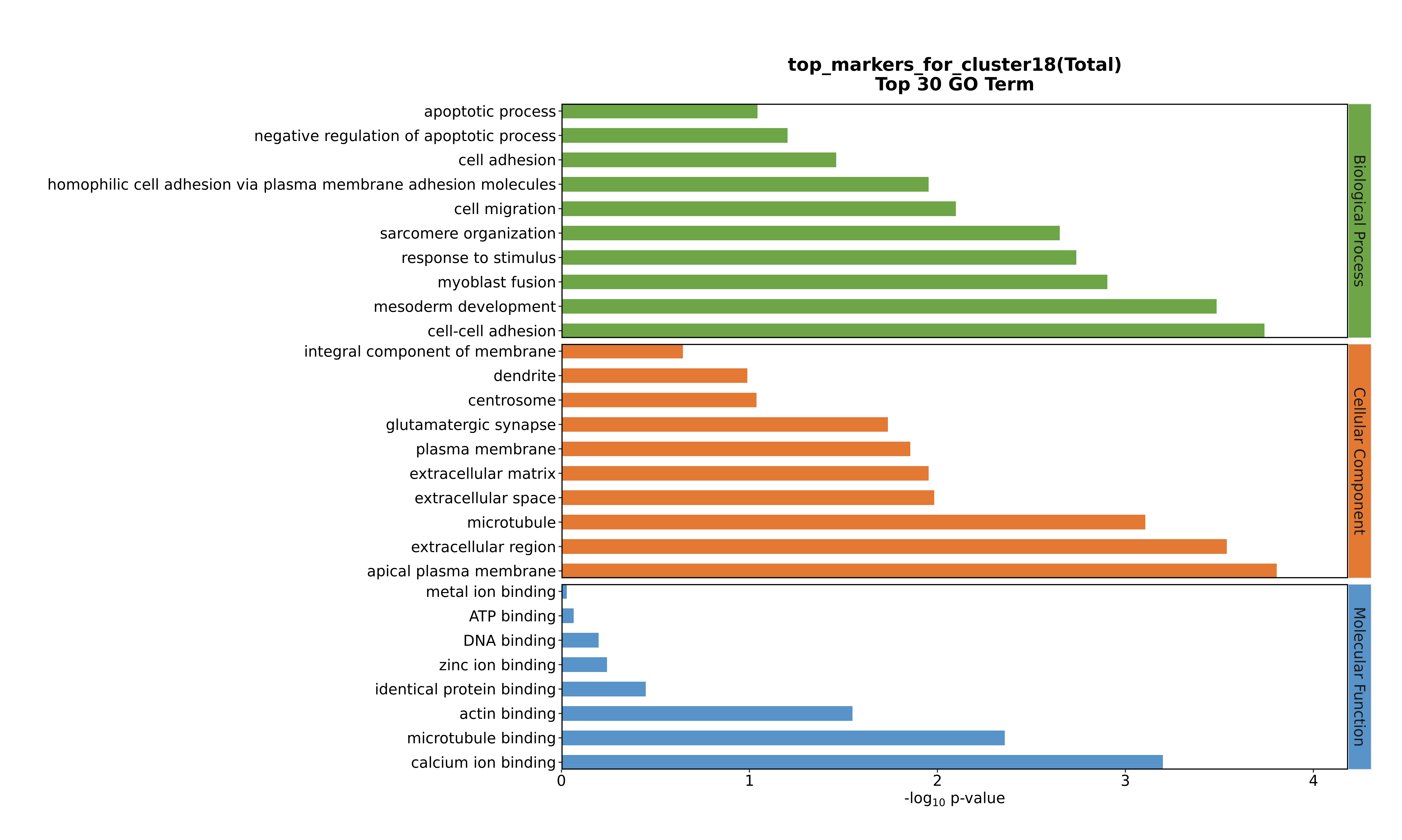

### Cluster19 GO top100.png

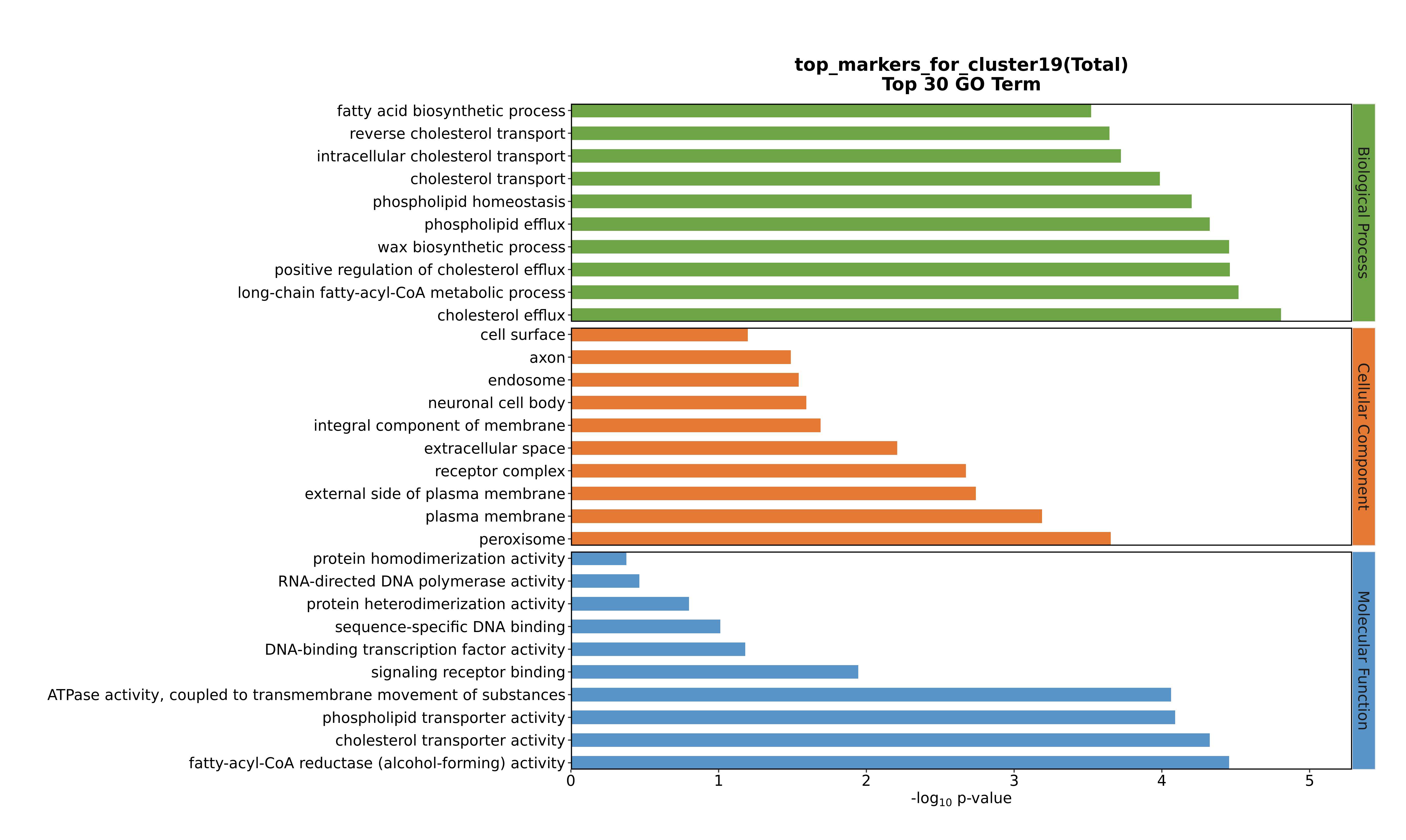

### Cluster19 GO top100.png

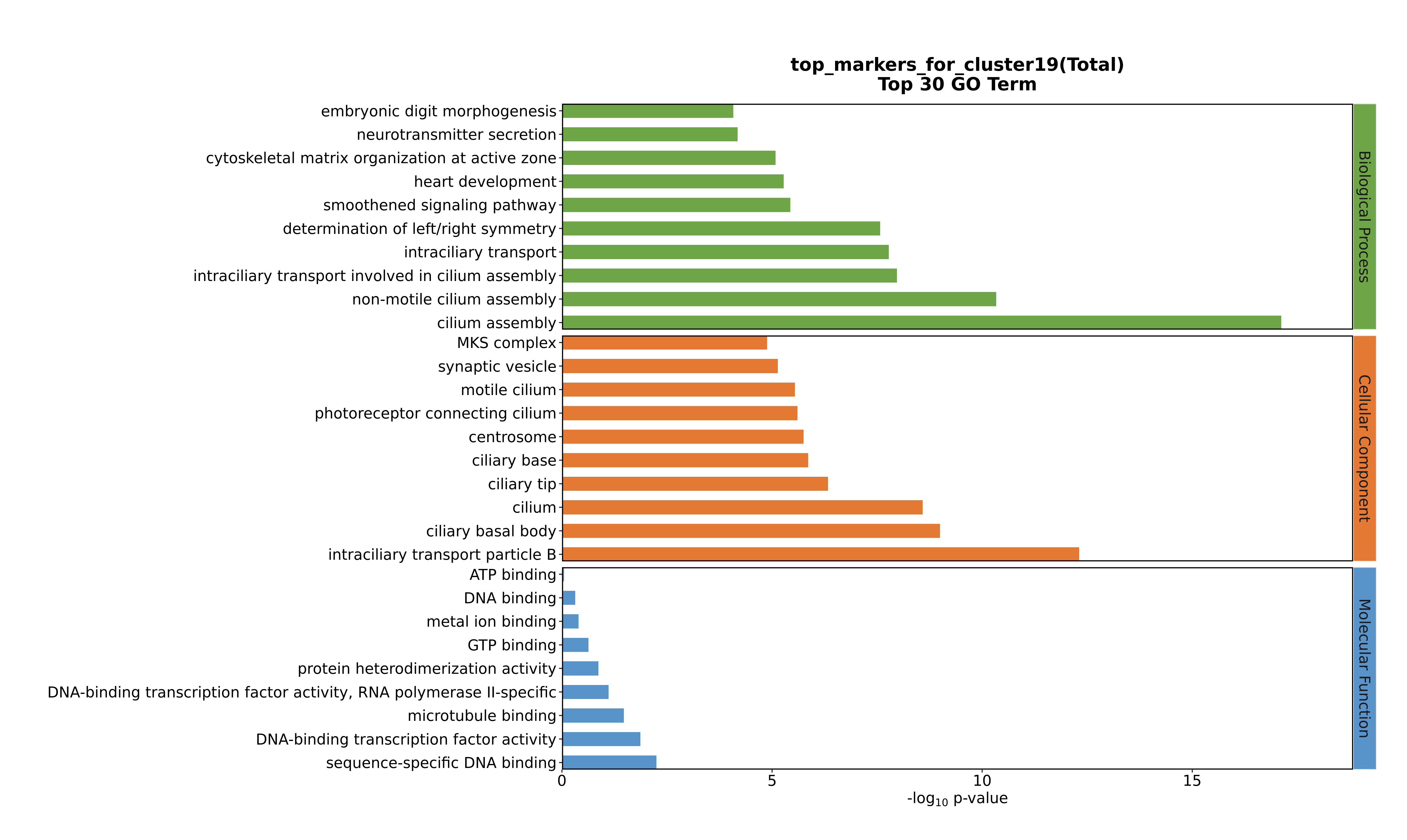

### Cluster19 GO top100.png

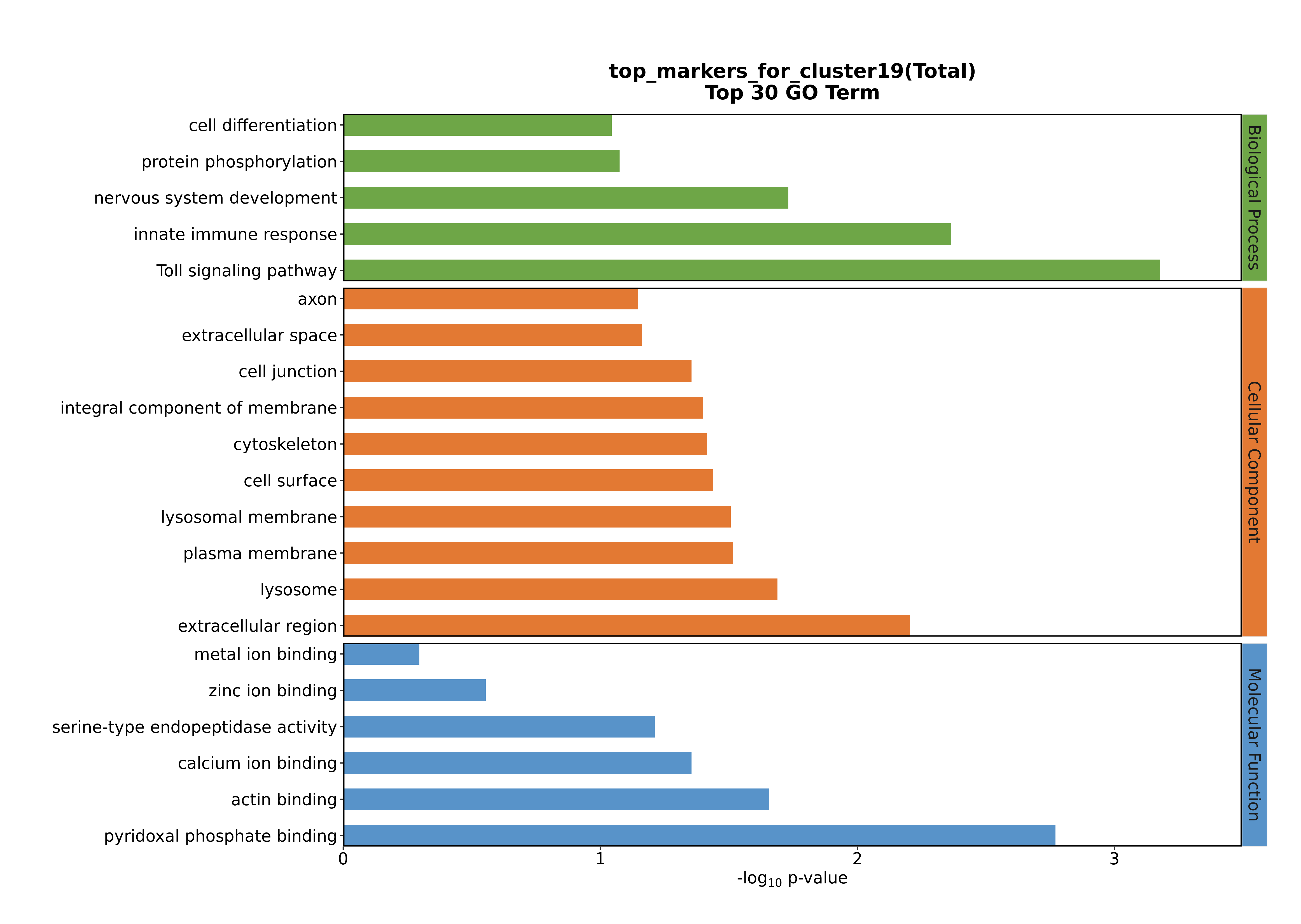

### Cluster19 GO top100.png

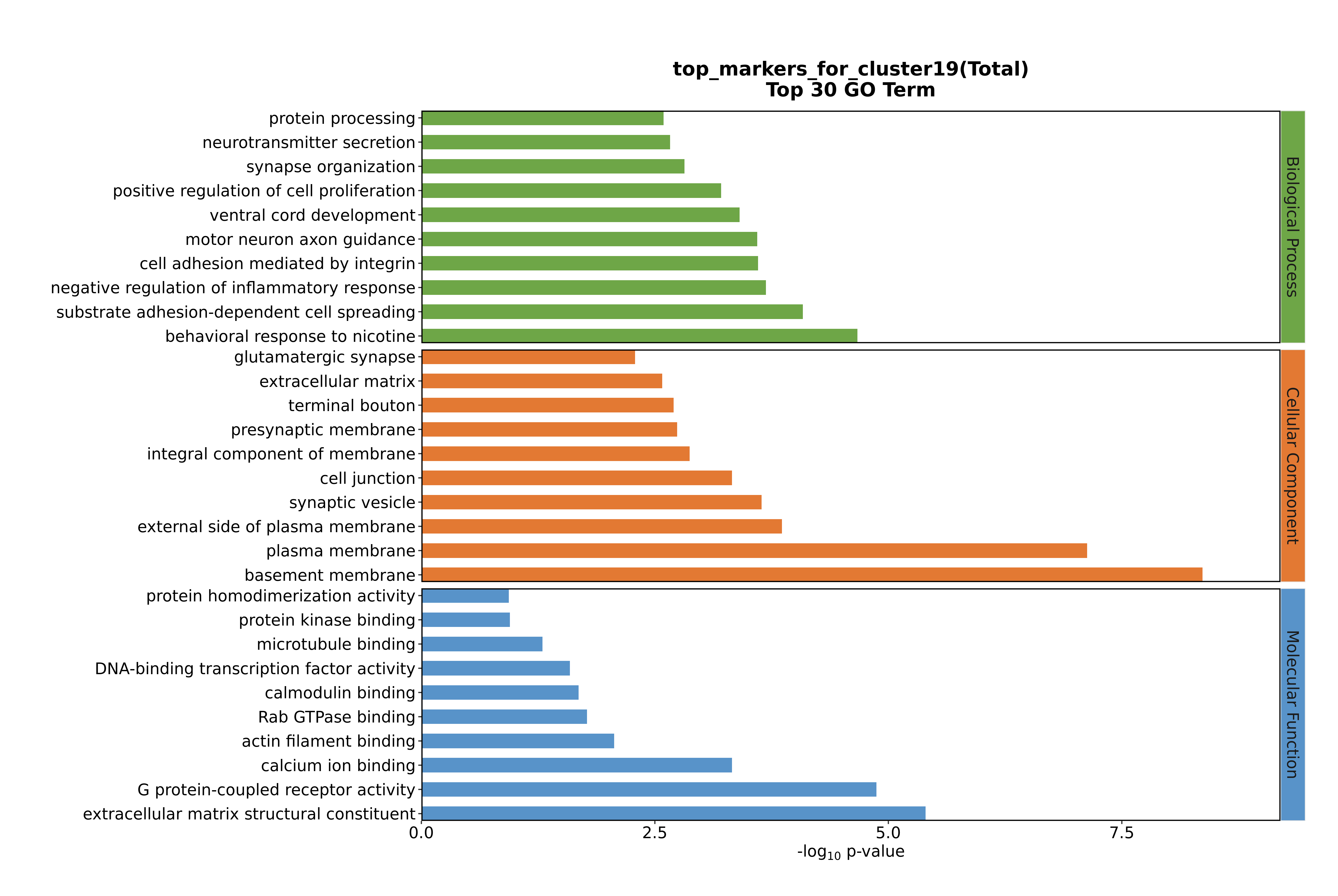

### Cluster20 GO top100.png

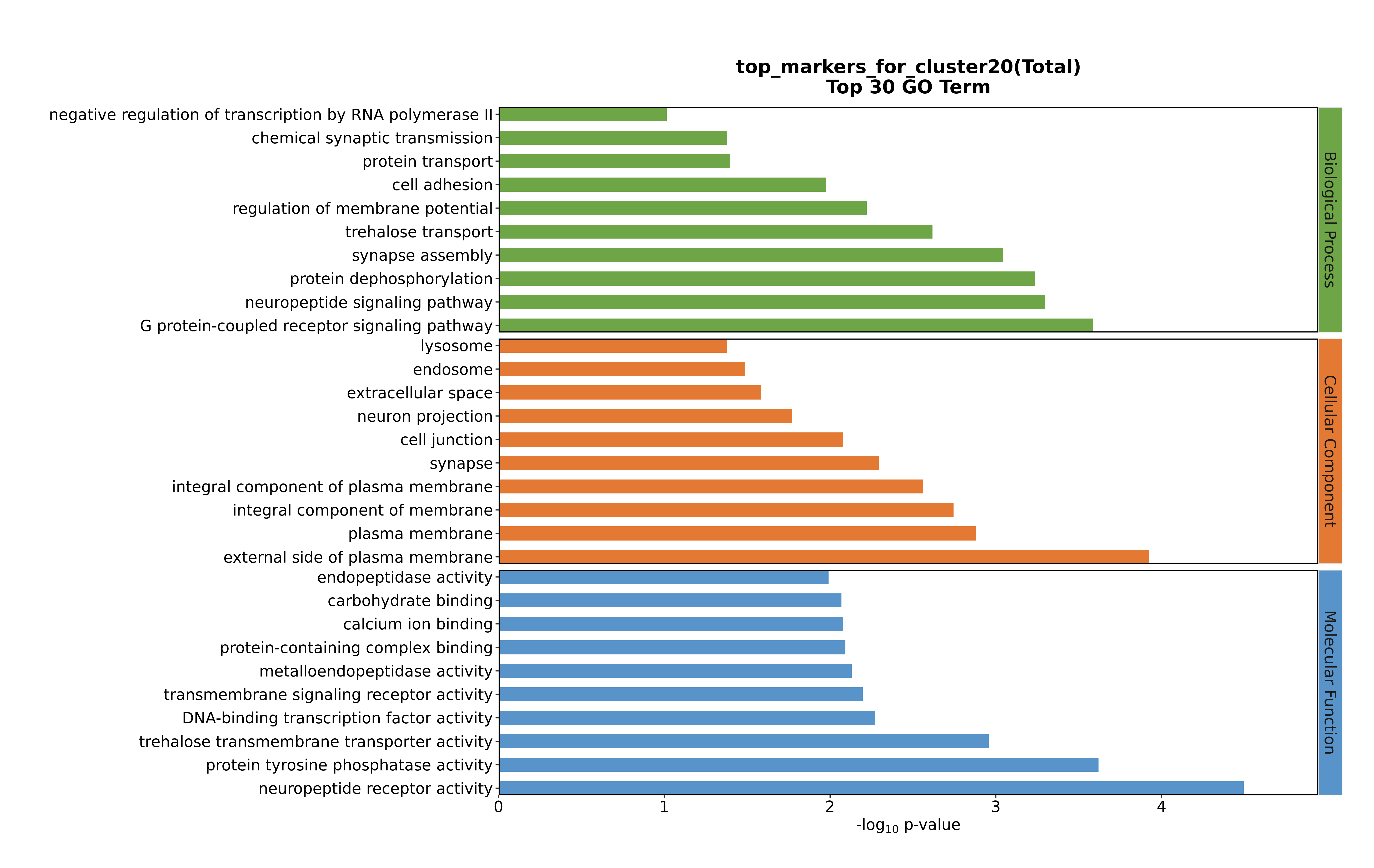

### Cluster20 GO top100.png

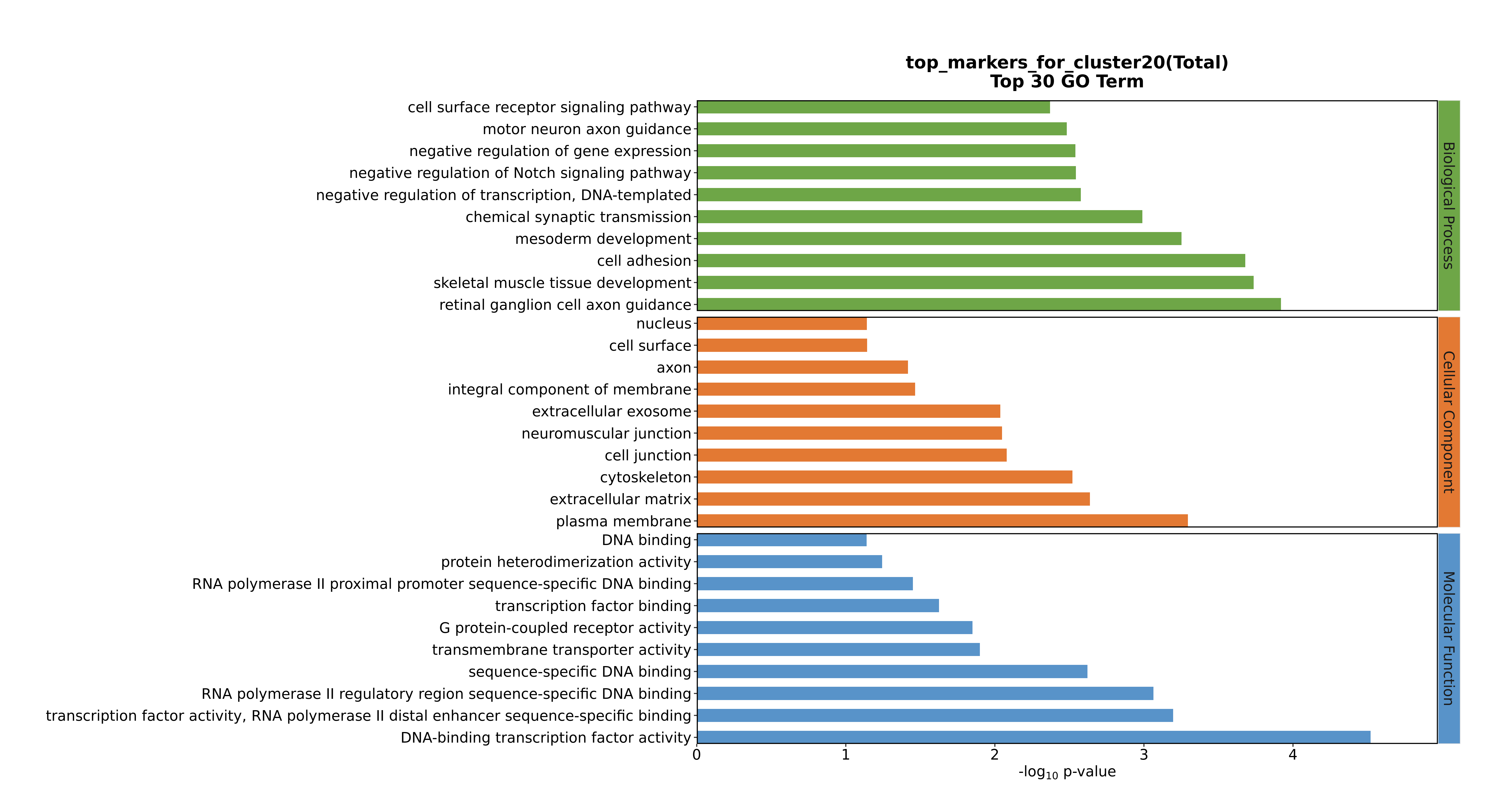

### Cluster20 GO top100.png

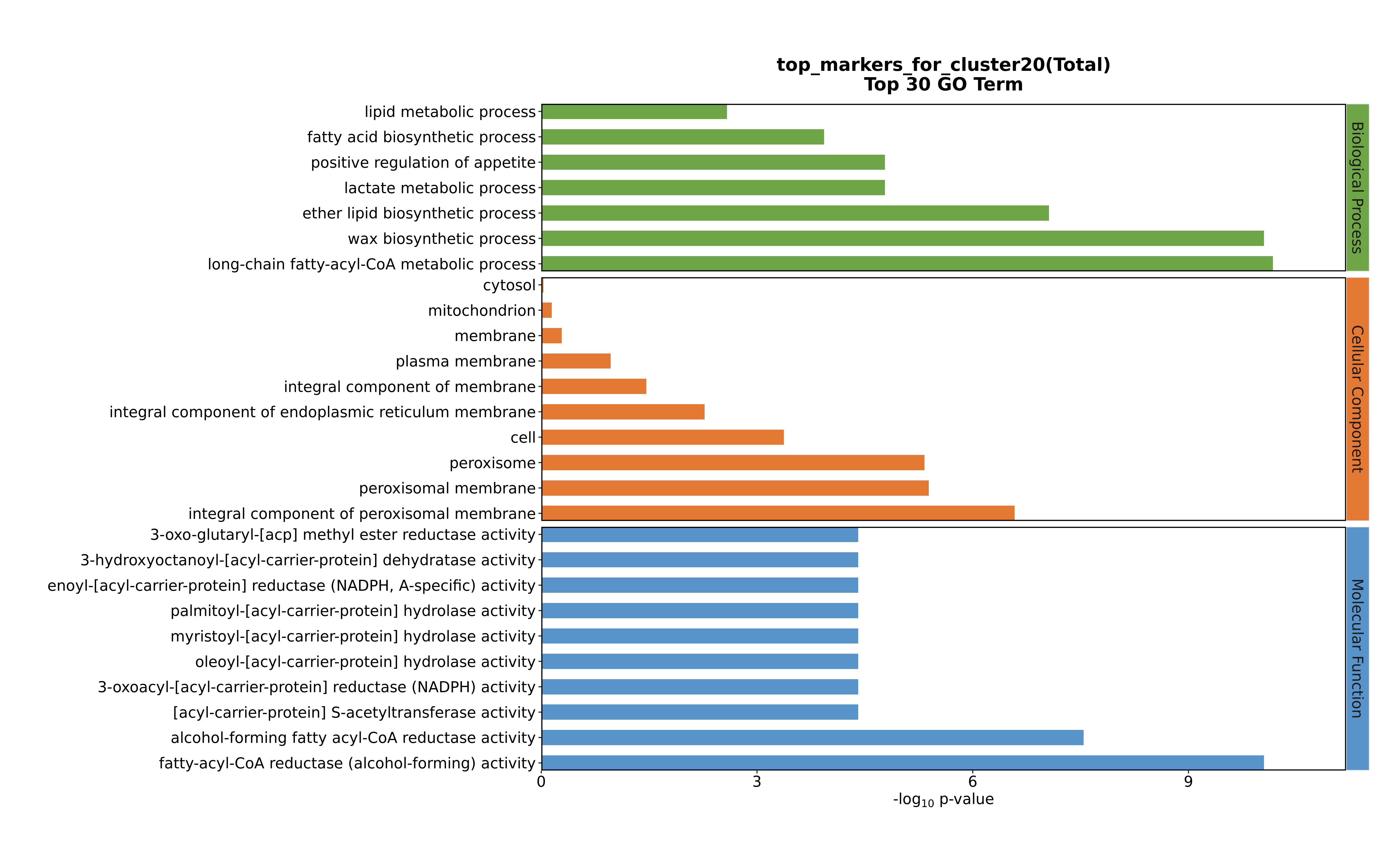

### Cluster20 GO top100.png

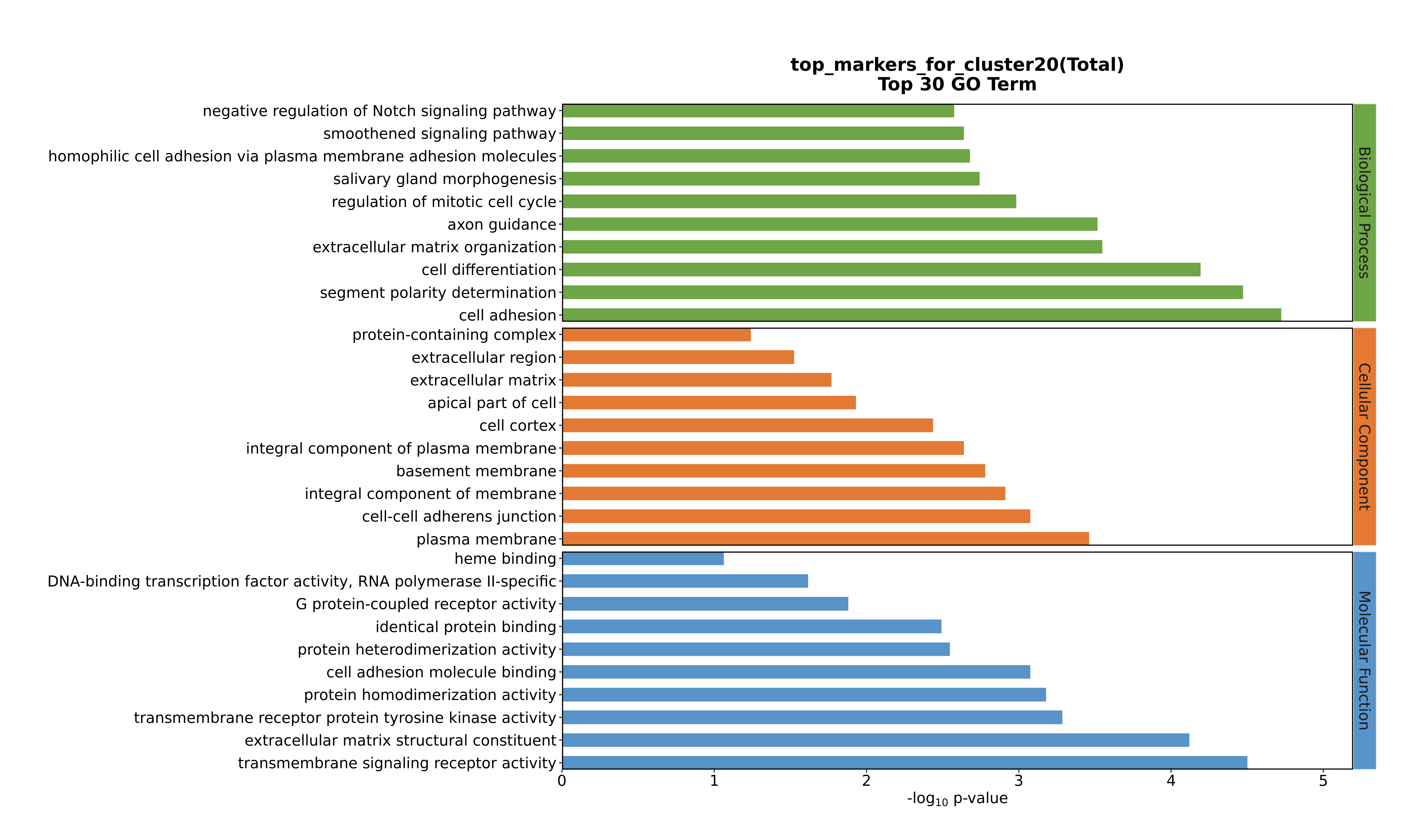

### Cluster21 GO top100.png

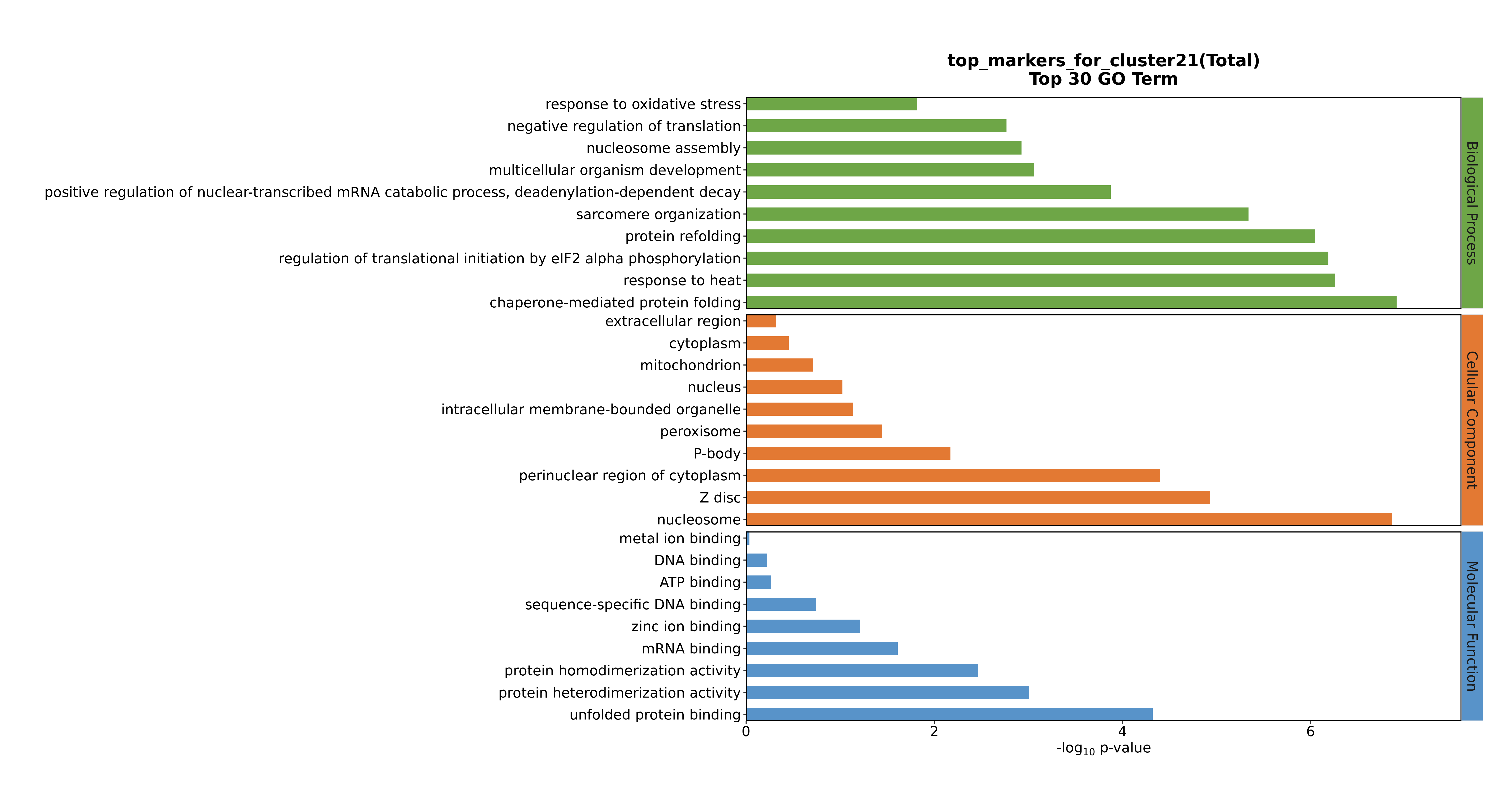

### Cluster21 GO top100.png

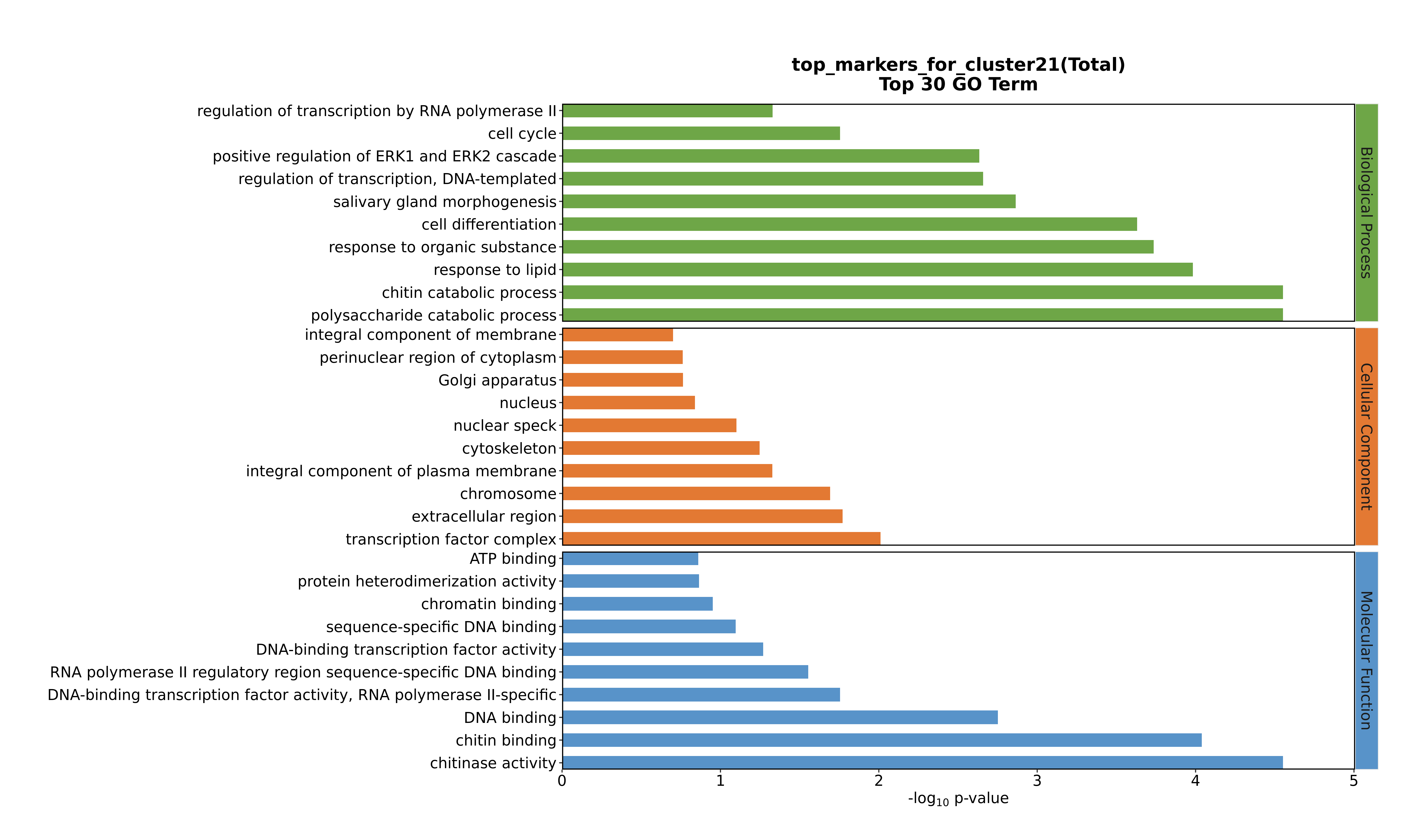

### Cluster21 GO top100.png

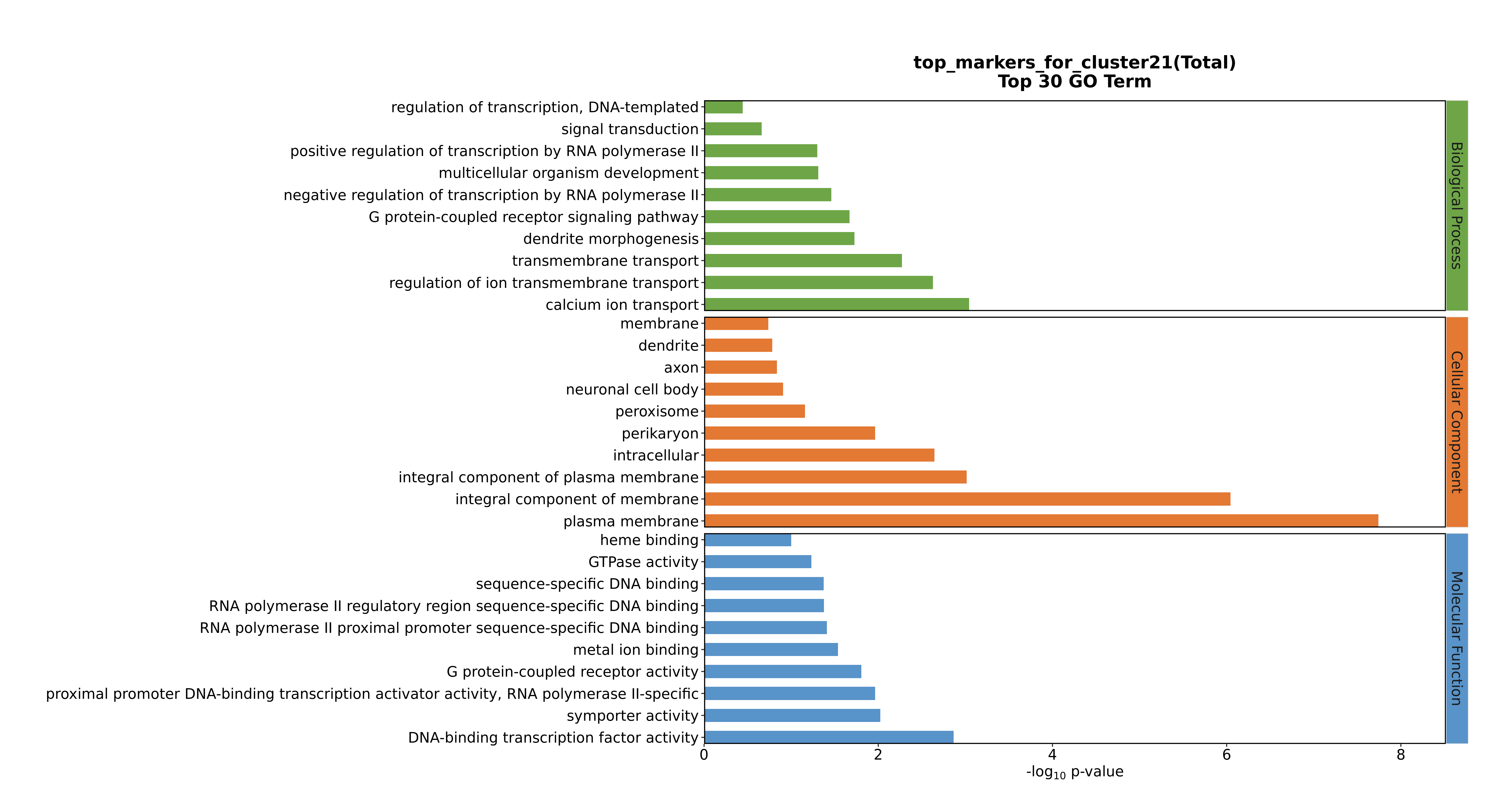

### Cluster21 GO top100.png

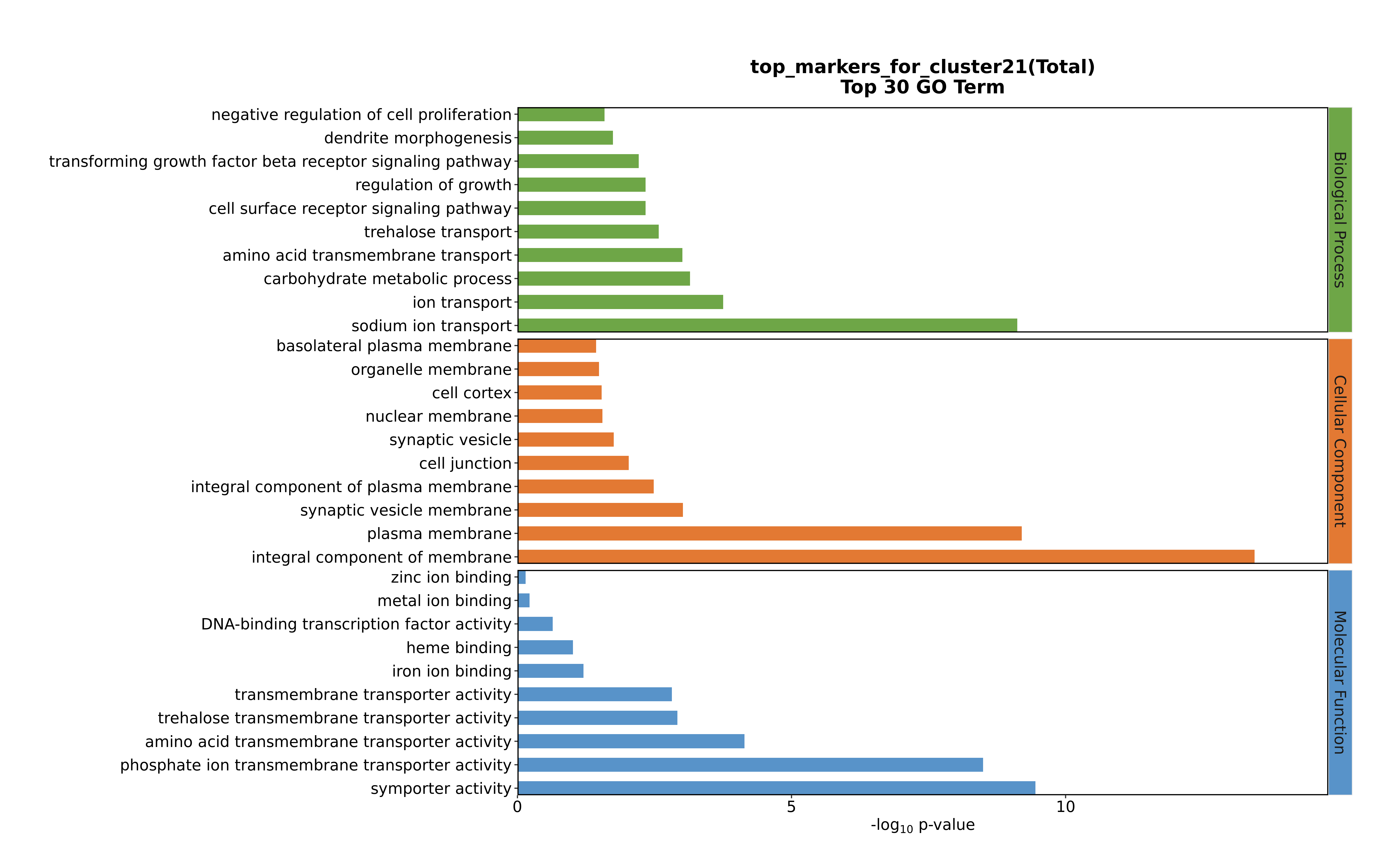

### Cluster22 GO top100.png

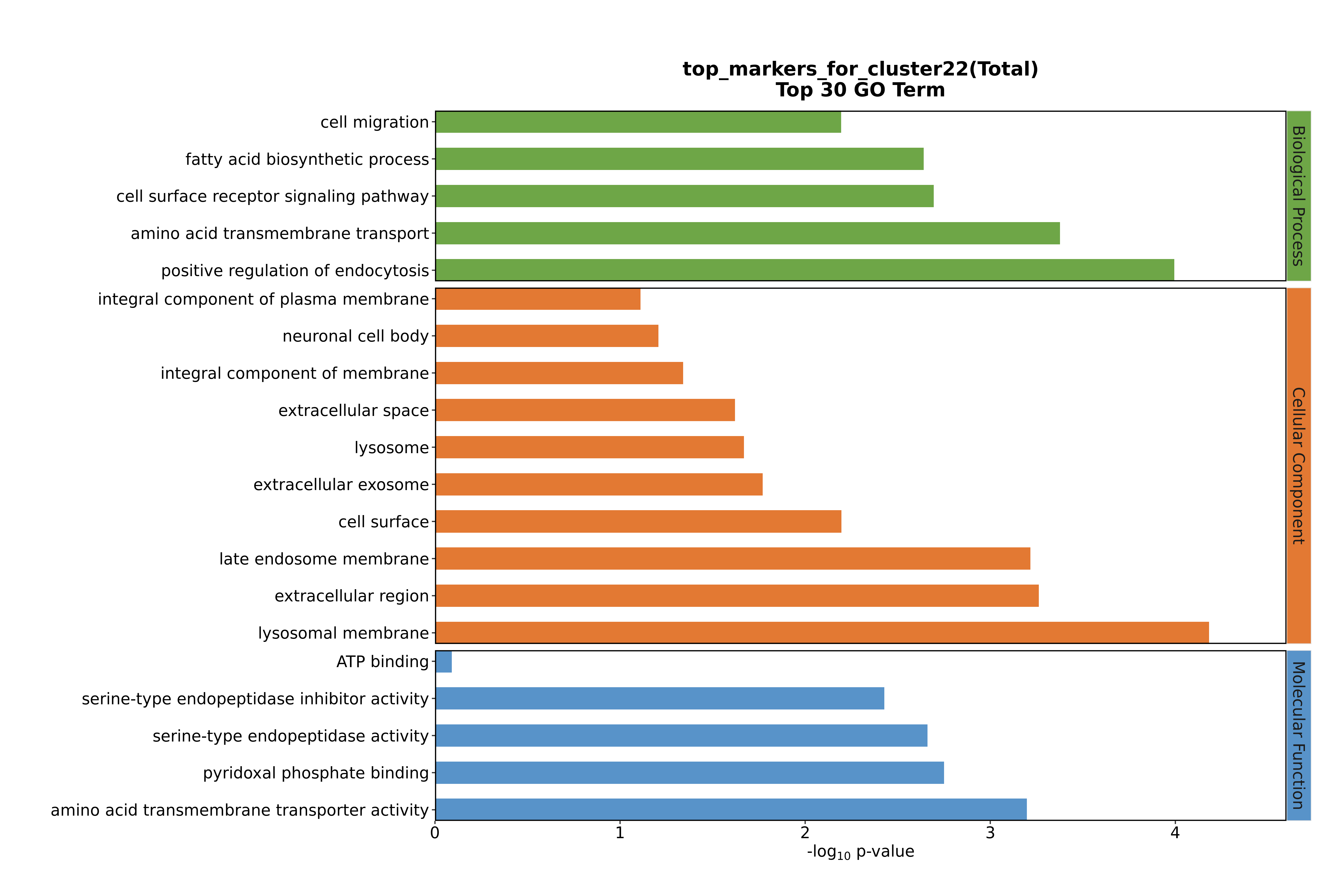

### Cluster22 GO top100.png

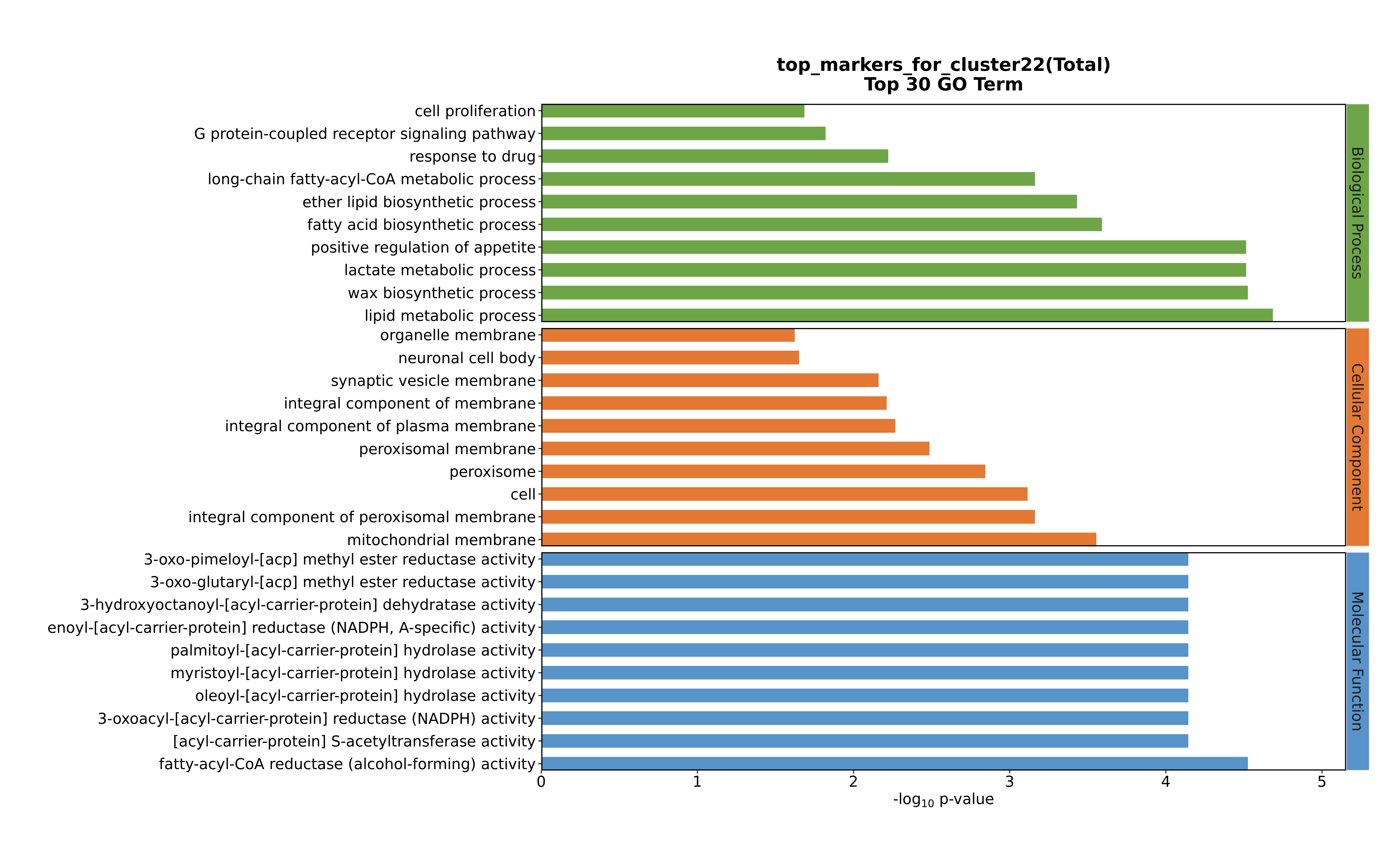

### Cluster22 GO top100.png

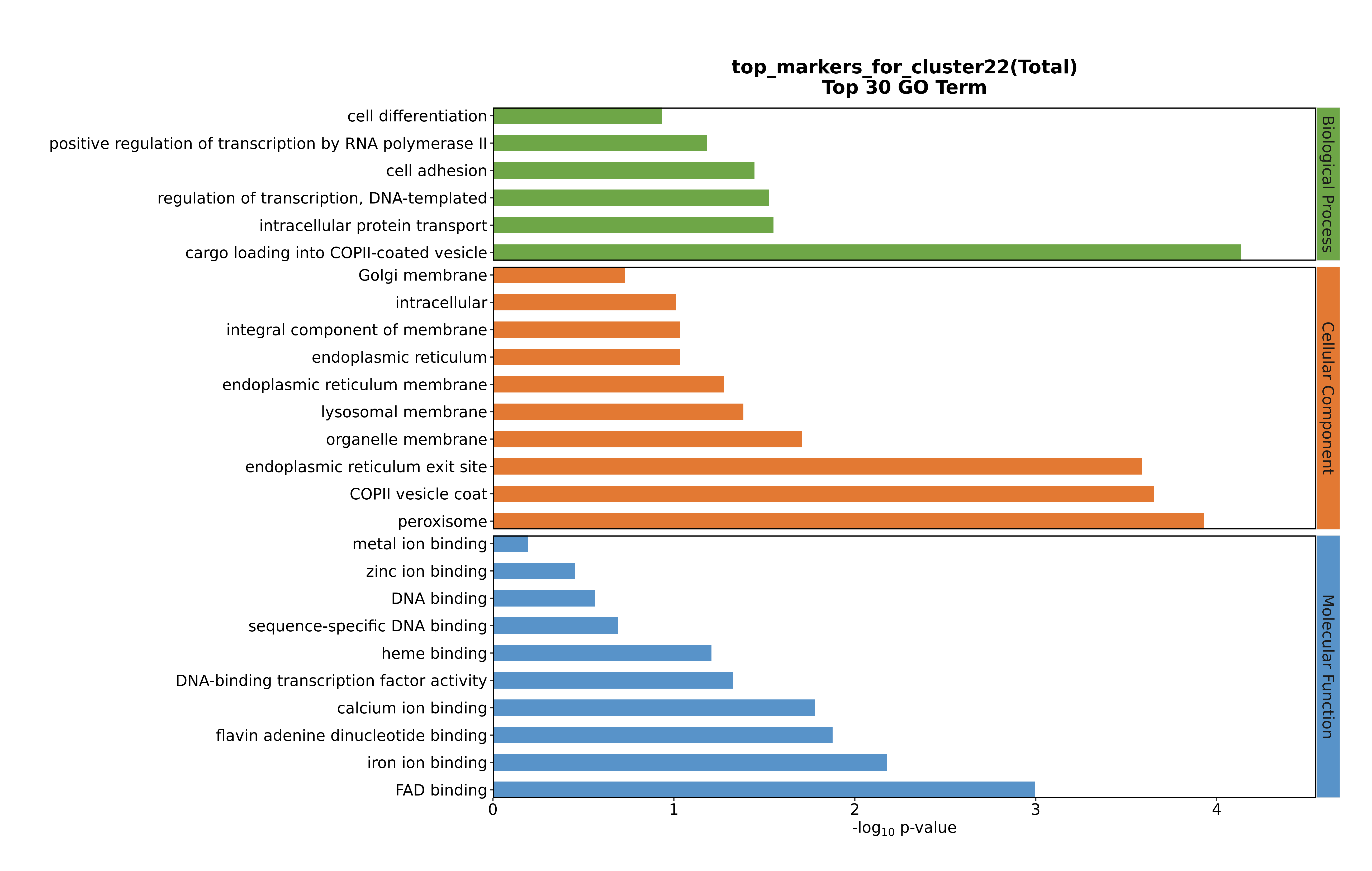

### Cluster22 GO top100.png

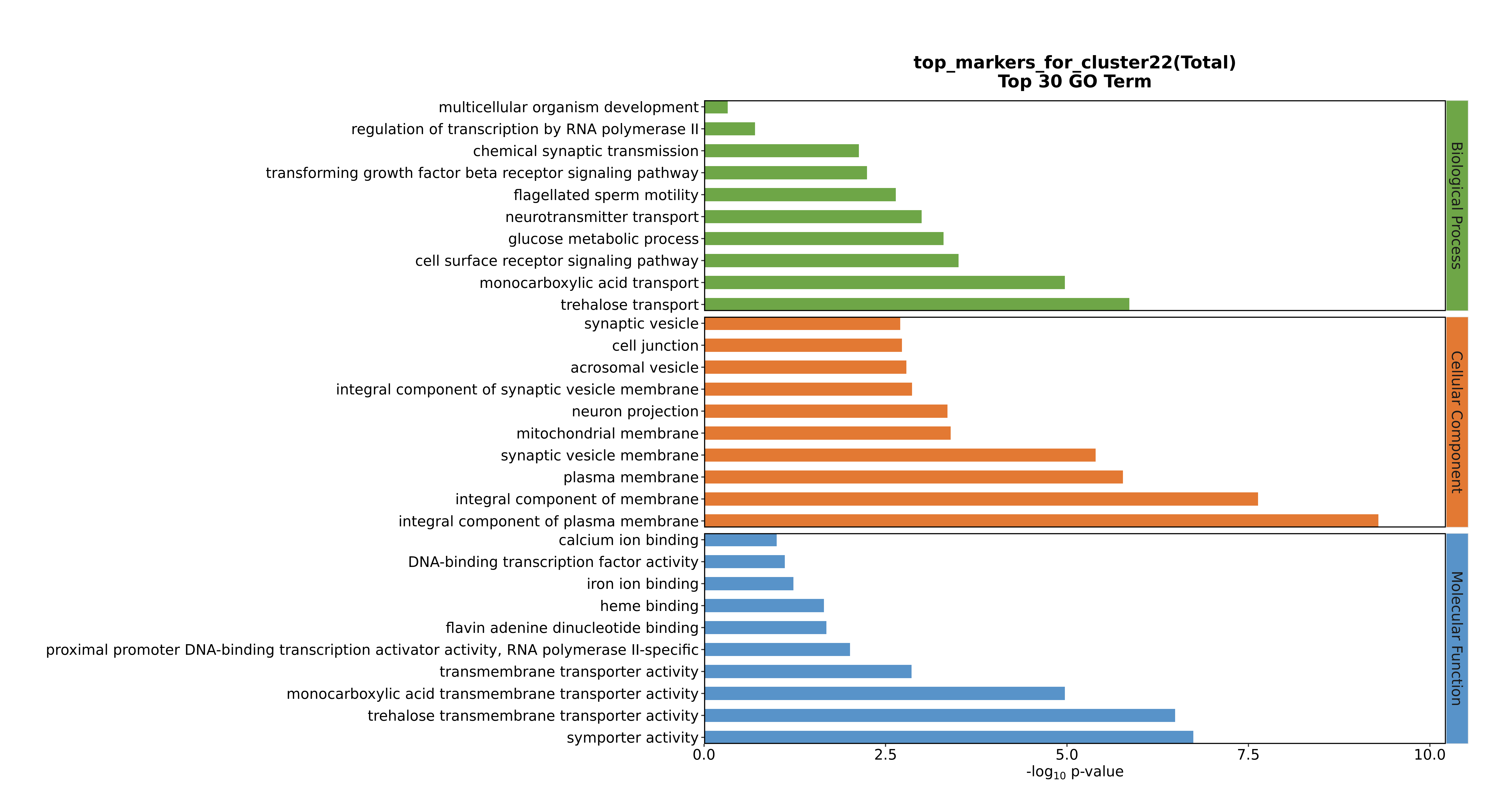

### Cluster23 GO top100.png

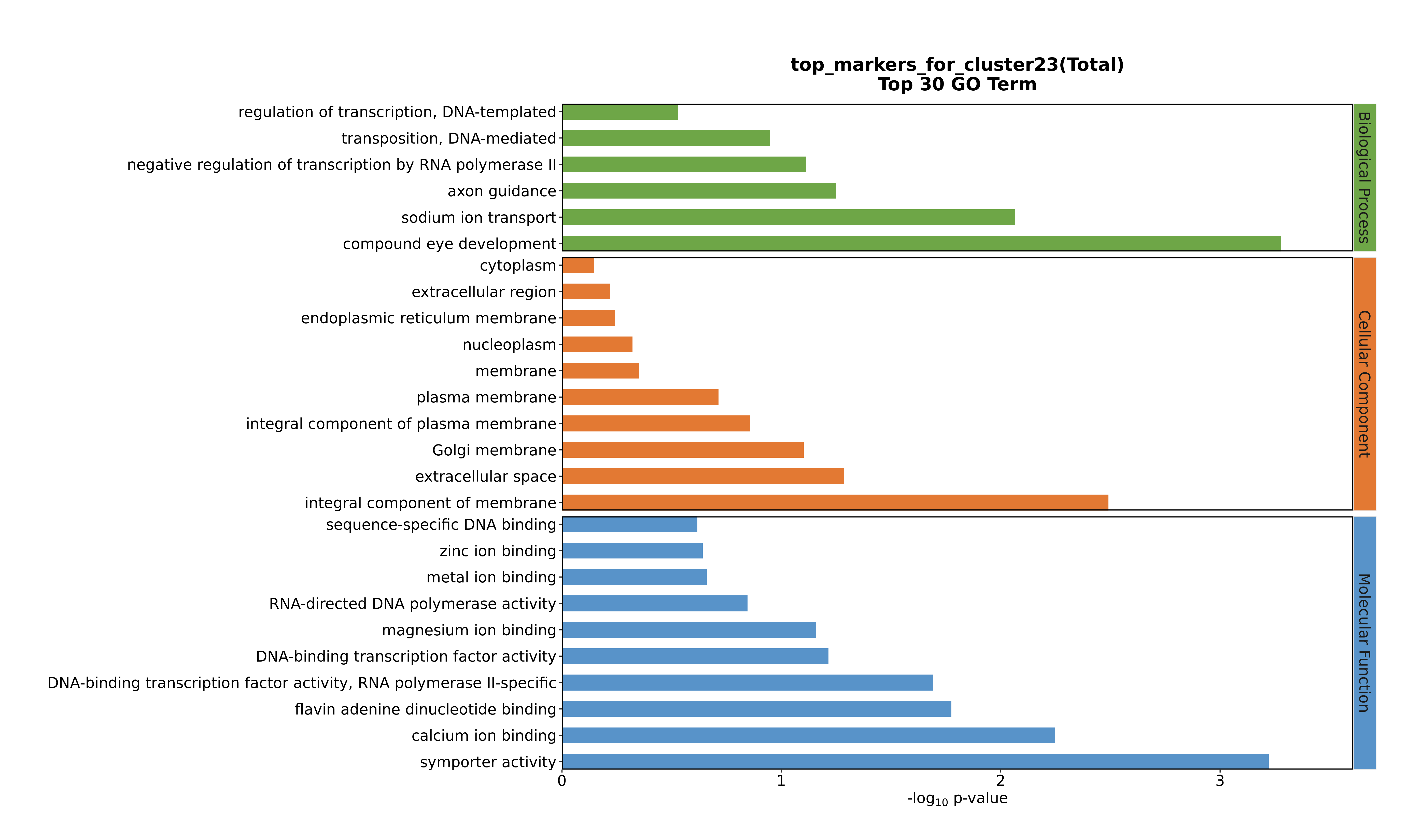

### Cluster24 GO top100.png

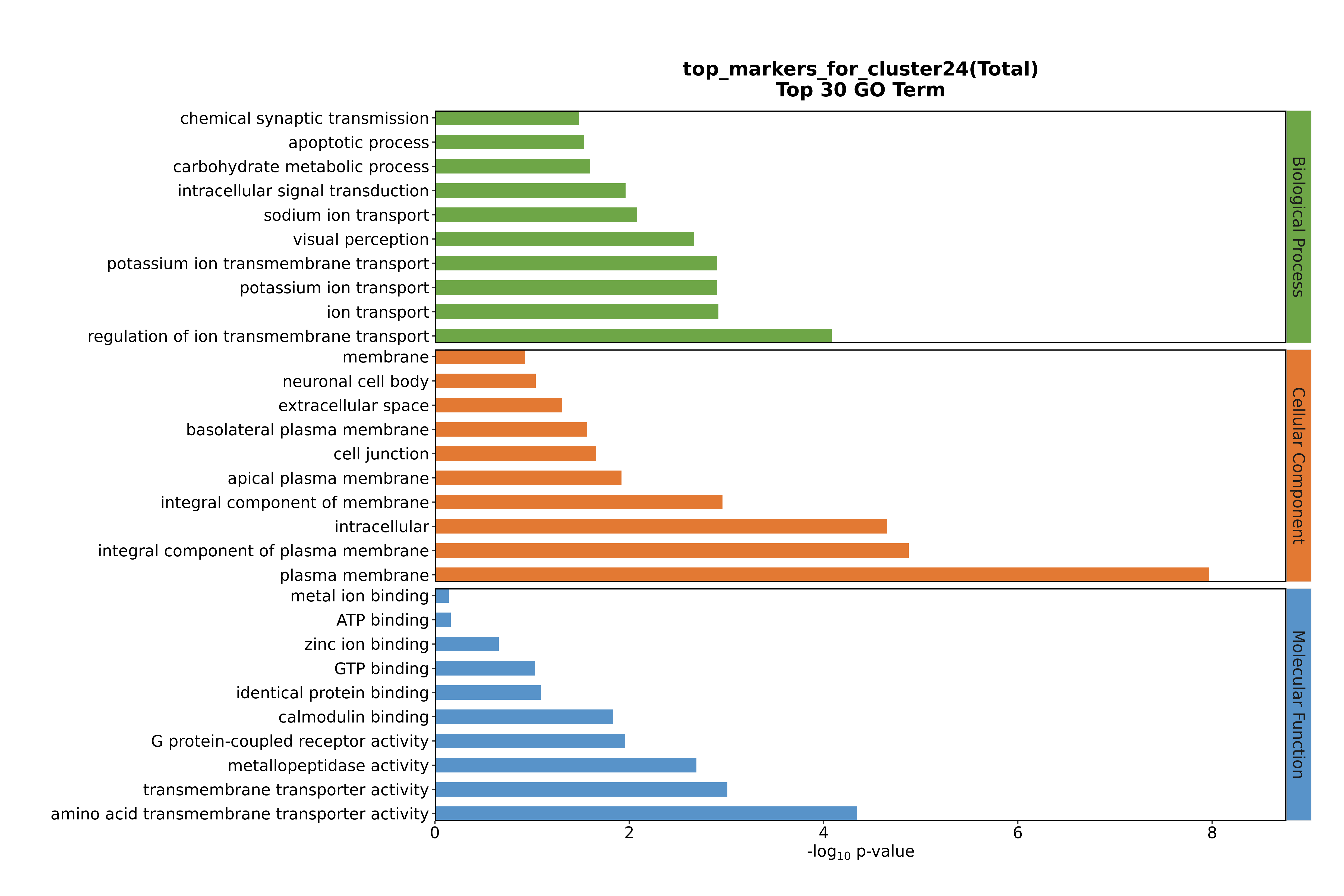
